# Enhanced longitudinal differential expression detection in proteomics with robust reproducibility optimization regression

**DOI:** 10.1101/2021.04.19.440388

**Authors:** Tommi Välikangas, Tomi Suomi, Courtney E. Chandler, Alison J Scott, Bao Q. Tran, Robert K. Ernst, David R. Goodlett, Laura L. Elo

## Abstract

Quantitative proteomics has matured into an established tool and longitudinal proteomic experiments have begun to emerge. However, no effective, simple-to-use differential expression method for longitudinal proteomics data has been released. Typically, such data is noisy, contains missing values, has only few time points and biological replicates. To address this need, we provide a comprehensive evaluation of several existing differential expression methods for high-throughput longitudinal omics data and introduce a new method, Robust longitudinal Differential Expression (RolDE). The methods were evaluated using nearly 2000 semi-simulated spike-in proteomic datasets and a large experimental dataset. The RolDE method performed overall best; it was most tolerant to missing values, displayed good reproducibility and was the top method in ranking the results in a biologically meaningful way. Furthermore, contrary to many approaches, the open source RolDE does not require prior knowledge concerning the types of differences searched, but can easily be applied even by non-experienced users.

## 1. Introduction

In the course of the past few decades, mass spectrometry (MS)-based proteomics has developed significantly and emerged as a powerful tool for clinical biomarker discovery [1]. Currently, MS-powered quantitative proteomics can be considered as an established method routinely used for proteome exploration in biomedical research [2,3].

Longitudinal study designs are generally regarded as having more statistical power to detect differences between the examined study groups than cross-sectional designs [4,5]. While requiring more measurements per individual, less individuals are required to achieve the same statistical power as in cross-sectional studies [4,5]. In addition to having more statistical power, longitudinal study designs deliver information concerning the changes in the studied individuals over time. In the context of high-throughput transcriptomics, longitudinal experiments for detecting time-resolved gene expression changes have been performed already for more than two decades [6–8]. For proteomic profiling, most of the experiments thus far have utilized cross-sectional study designs, but longitudinal proteomic experiments with two [3] or multiple [9,10] time points have begun to emerge.

A recent study comparing longitudinal methods for RNA-sequencing (RNA-seq) gene expression experiments discovered that most of the specific longitudinal methods performed worse than the timepoint-wise analysis of the data using traditional pairwise differential expression tools when the number of time points was small (< 8) [11]. With only few time points, most of the tested longitudinal methods produced a high number of false positives [11]. When the number of time points was increased, the performance of many of the longitudinal methods also improved [11]. However, as currently relatively few time points are typical for a biomedical study, the usability of such methods which cannot perform well on short time series data is limited and new approaches are required.

In the context of proteomics data, another limitation of the longitudinal methods for RNA-seq data [11] is that many of them are specifically designed for discrete negative binomially distributed count data and, as such, are not directly applicable. Proteomics data is typically close to normally distributed after logarithm transformation and/or normalization [12], making methods originally proposed for the analysis of longitudinal gene expression microarray data better suited. Among those methods, BETR (Bayesian Estimation of Temporal Regulation) [13] and Timecourse [14] rely on a Bayesian framework, and Microarray Significant Profiles (MaSigPro) builds on a two-step regression strategy [15]. Additionally, the popular R package for the analysis of microarray and sequencing data, Limma [16], also contains tools for analyzing longitudinal differential expression. However, since these methods do not take into account the characteristics of MS proteomics data, they might not be optimal for proteomics experiments. In particular, as has been extensively observed in previous studies [17,18], missing values are prevalent in proteomics data and their handling is not trivial [18]. This is particularly the case with the data dependent acquisition (DDA) label-free proteomics approach, which is popular due to its cost efficiency, speed and ability to handle complex samples [19]. Furthermore, MS data is prone to noise in quantification [20,21], rendering especially the lowly abundant proteins subject to false positive detections [22].

In addition to the methods specially designed for longitudinal omics data, different statistical modelling approaches have been utilized in their analysis. Several types of linear and non-linear regression based approaches, with or without random effects, have been applied in various contexts [10,23–25]. For example, Liu et al. [10] used mixed effects regression modelling with quadratic random terms to detect trends and differential expression patterns in the longitudinal expression of proteins between children developing type 1 Diabetes (T1D) and healthy controls. However, no comprehensive comparison on the performance and application of the different approaches exist and, therefore, specific standard practices for analyzing differential expression in longitudinal omics data with regression modelling have not been established. Furthermore, the statistical frameworks developed for the analysis of longitudinal clinical variables might not be best suited for the analysis of longitudinal omics data, in which the number of individuals and the number of measured time points are typically small, but the number of simultaneous variables (e.g. genes or proteins) is very large.

To address the need for a robust method applicable to noisy longitudinal proteomics data, we introduce here a new method, Robust longitudinal Differential Expression (RolDE), which combines the detection power of reproducibility optimization and longitudinal regression modelling. The method builds on our reproducibility optimization procedure (ROTS) [26,27], which has been shown to be especially effective in controlling the number of false positive detections in cross-sectional omics studies by us and others [12,28–31], including proteomics [12,29,30]. To demonstrate the benefits of RolDE in the analysis of longitudinal proteomics data, we performed an extensive evaluation of its performance against multiple existing longitudinal approaches and a baseline cross-sectional method using nearly 2000 semi-simulated proteomics spike-in datasets as well as real experimental biological data. The examined datasets included a large variety of linear and non-linear longitudinal trend differences between the examined conditions. Our results show that RolDE robustly detects true differential expression between the conditions, is tolerant against missing values, has great reproducibility and is suitable for both shorter and longer time series data. Furthermore, RolDE does not require prior knowledge about the type of differences searched from the data but can still effectively detect both linear and non-linear differences simultaneously. RolDE is freely available as an R package in GitHub (https://github.com/elolab/RolDE).

## 2. Results

The proposed new method, RolDE, is a composite method, consisting of three independent modules, *RegROTS, DiffROTS and PolyReg*, with different approaches in detecting longitudinal differential expression. The combination of these diverse modules allows RolDE to robustly detect varying types of differences in longitudinal trends and expression levels in diverse experimental settings (**Figure 1A**). We have extensively evaluated the performance of RolDE against several existing specialized longitudinal differential expression methods and modelling approaches: Reproducibility Optimized Test Statistic (BaselineROTS), Bayesian Estimation of Temporal Regulation (BETR), Linear Models for Microarray Data (Limma, LimmaSplines), Timecourse (Timecourse), Microarray Significant Profiles (MaSigPro), Linear mixed effects regression modelling (Lme), and Polynomial mixed effects modelling (Pme). The performance evaluation of the methods was conducted using nearly 2000 semi-simulated spike-in proteomics datasets (**Figure 1B).** Reproducibility and biological relevance of the methods findings was further assessed with the large experimental Fn dataset (**Figure 1C**). Additionally, the ability of RolDE to perform well and produce meaningful findings even in data with non-aligned time points was demonstrated using the previously published longitudinal T1D proteomics dataset of [10] (**Figure 1D**).

**Figure 1.**
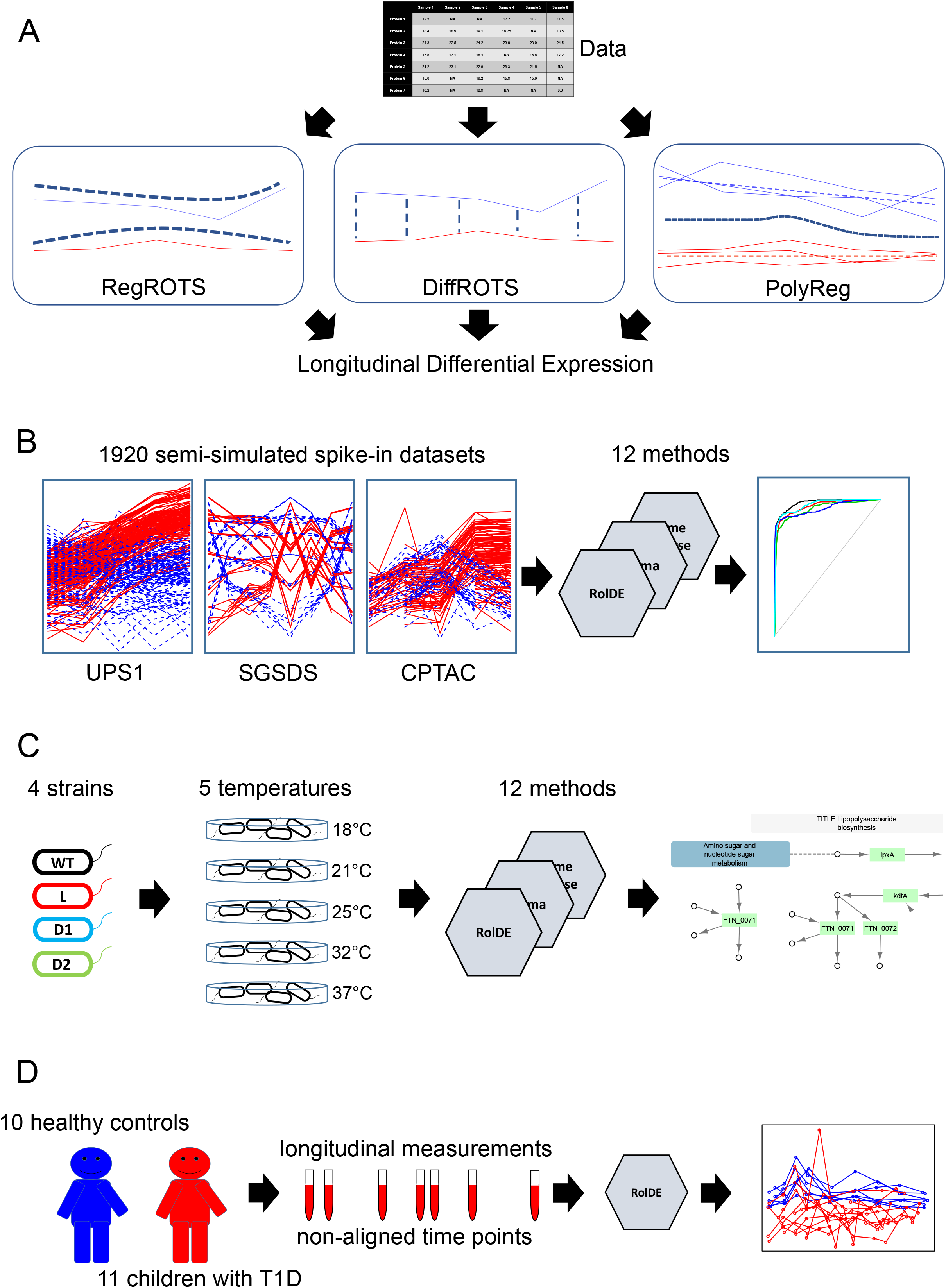
An illustration of the new proposed method and the study design. **(A)** The proposed new method, Robust longitudinal Differential Expression (RolDE), is a composite method, consisting of three independent modules with different approaches to detecting longitudinal differential expression. The **RegROTS** module combines individual regression modelling with the power of the established differential expression method Reproduciblity Optimized Test Statistic (ROTS) [26,27]. In the **DiffROTS** module, the expression between all the individuals in the different conditions is directly compared at all timepoints. The **PolyReg** module uses polynomial regression modelling to evaluate longitudinal differential expression. The combination of these modules allows RolDE to robustly detect differences in longitudinal trends and expression levels in diverse data types and experimental settings. **(B)** 1920 semi-simulated spike-in proteomics datasets with two conditions, five (UPS1 [29] and CPTAC [55]) or eight (SGSDS [54]) time points in each condition and varying trend differences between the conditions were generated for an extensive evaluation of the performances of 11 established approaches together with the proposed method in correctly detecting known longitudinal differential expression. **(C)** The reproducibility of the methods and their ability to provide biologically relevant findings was evaluated in an experimental *Francisella tularensis* subspecies *novicida (Fn)* dataset. The dataset consisted of a wild type and three null mutant strains of N-acyltransferases, lpxD1 (D1), lpxD2 (D2) and lpxL (L), involved in the production of a key factor, lipid A. Each strain was grown in five temperatures: 18°C, 21°C, 25°C, 32°C, 37°C to activate the temperature sensitive enzymes. Technical replicate datasets were used to evaluate the reproducibility of the method’s findings between the strains, while the growth at different temperatures offered a surrogate for the time points. Biological relevance of the method’s findings was evaluated using the associated KEGG [69] pathway “Lipopolysacchararide biosynthesis – *Francisella tularensis* subsp. *novicida* U112” and the knockout pathway “Lipopolysaccharide biosynthesis knockout pathway” (ko00540). **(D)** The ability of the new proposed method RolDE, to detect longitudinal differential expression even when the time points in the data are not aligned, was further demonstrated using a previously published longitudinal type 1 diabetes (T1D) proteomics data of [10].

### 2.1 Performance in the semi-simulated spike-in proteomics data with single trend categories

First, we investigated the performance of the methods in the filtered semi-simulated spike-in datasets, where a single trend category per condition was generated for each dataset (Stable, Linear, LogLike, Poly2, Sigmoid, or PolyHigher; **Supplementary File 1**) and only proteins with no missing values were included in the analysis.

In the UPS1-based filtered datasets, the overall highest pAUCs were obtained with the new proposed method RolDE with an IQR mean pAUC of 0.976 (**Figure 2A**), performing significantly better than the second best method Timecourse with an IQR mean pAUC of 0.973 (p = 0.038, one-tailed paired Mann-Whitney *U* test). The baseline test method ROTS ignoring longitudinal trends also performed well with an IQR mean pAUC of 0.924, while the difference to the best performing method RolDE was highly significant (p < 10^−15^). Among the regression-based models, the lower order regression models (denoted by the extension L) performed overall worse than models of higher polynomial degree (denoted with the extension H).

**Figure 2.**
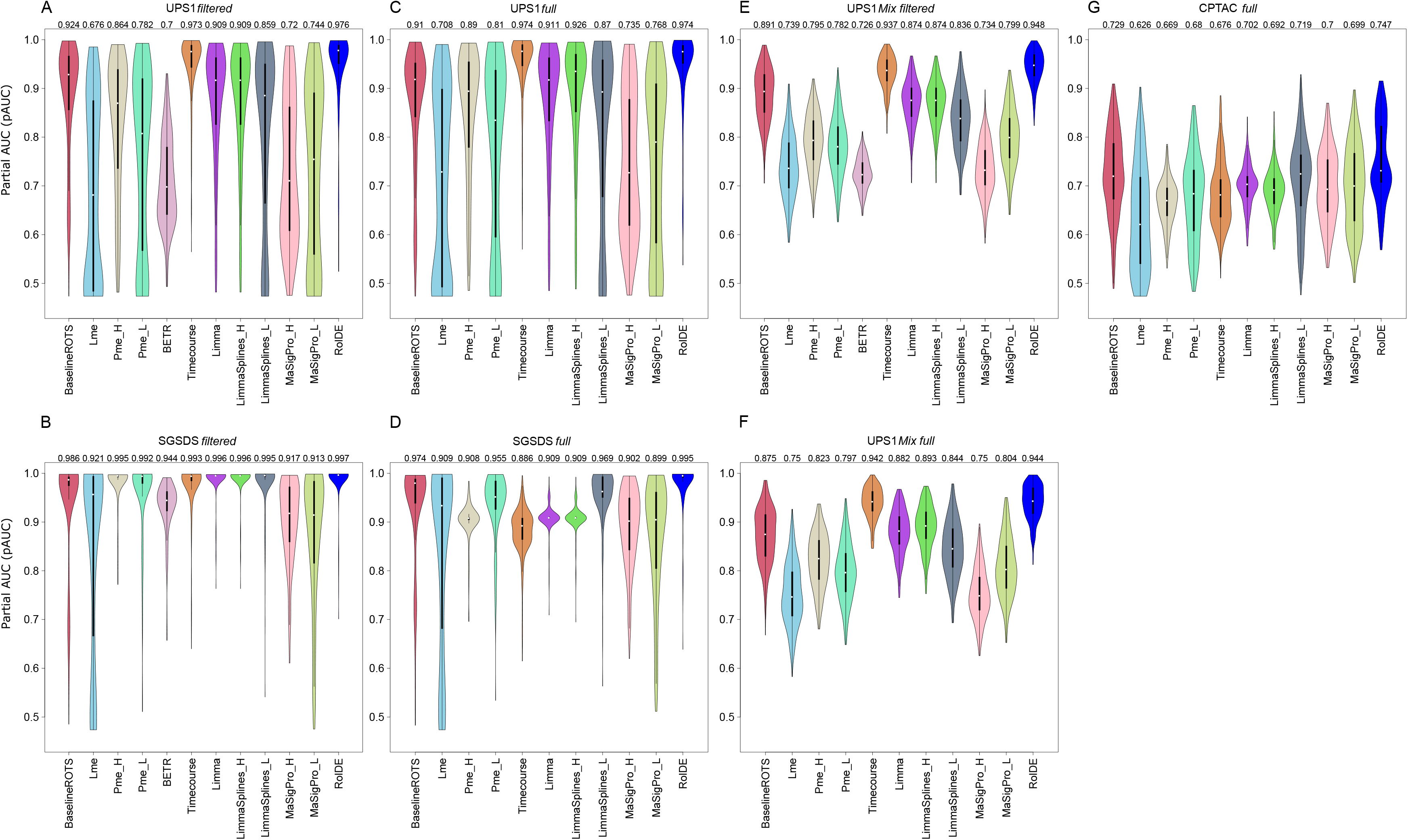
The performance of the examined methods in the semi-simulated spike-in datasets. **(A)** UPS1 filtered, **(B)** SGSDS filtered, **(C)** UPS1 full, **(D)** SGSDS full, **(E)** UPS1 Mix filtered, **(F)** UPS1 Mix full, (**G**) CPTAC full. The methods were examined in their ability to detect true (known) longitudinal differential expression using receiver operating characteristic (ROC) analysis across the UPS1-based [29] (300 filtered, 300 full, 300 mix filtered and 300 mix full), SGSDS-based [54] (210 filtered and 210 full) and the CPTAC-based [55] (300 full) datasets with varying longitudinal trend differences between the conditions in the spike-in proteins. The partial areas under the ROC curves (pAUC) between the specificity of 1 and 0.9 were used. The violin plots display the population of pAUCs for each method in each dataset. Black boxes represent the interquartile ranges, white circles represent median values and wider areas of the violins contain larger densities of values than narrow areas. The interquartile range (IQR) means of the pAUCs for each method are presented for each dataset. The IQR means represent typical performances the methods in the different dataset types.

In the SGSDS-based filtered datasets, all the tested methods performed relatively well (**Figure 2B**). Again, the performance of RolDE with IQR mean of pAUC 0.997 was significantly better than the next best methods Limma and LimmaSplines_H (p < 10^−12^).

A closer look at the performance of the methods in the different trend categories suggested that the new proposed method RolDE, together with Timecourse and BaselineROTS, performed consistently well in every category in both the UPS1-based and SGSDS-based datasets (**Supplementary Figure 1A-B**). For the general regression-based approaches, the performance was in concordance with the degree of the regression, as expected. The linear approach Lme performed well when the categories were linear or close to linear; the higher order polynomial regression Pme_L performed better when the examined categories were linear or close to second order polynomial. The highest order polynomial regression Pme_H performed well on the broadest spectrum of categories, but the performance was best when the examined categories were of higher order polynomial.

Second, we investigated the performance of the methods in the full semi-simulated spike-in datasets with a single trend category per condition but also proteins with missing values included in the analysis. Similarly as with the filtered datasets, RolDE performed overall best in the full datasets (**Figure 2C-D**). In the UPS1-based datasets, with missing values only in the true negative proteins, Timecourse and RolDE performed comparably best, both methods having an IQR mean pAUC of 0.974 and no significant differences in the overall performance (p = 0.185). In the SGSDS-based full datasets, with missing values also in the true positive spike-in proteins, RolDE with an IQR mean pAUC of 0.995 clearly outperformed the other methods and performed significantly better than the second best method BaselineROTS (p < 10^−15^). BETR does not tolerate missing values and was therefore excluded from the analysis of the full datasets.

Importantly, investigation of the performance of the methods in the different trend categories confirmed that the performance of RolDE in the full datasets remained on par to the filtered datasets without missing values, whereas most of the other methods experienced a decrease in their performance over all categories in the SGSDS-based full datasets when compared to the corresponding filtered datasets (**Supplementary Figure 1B,D**).

### 2.2 Performance in the semi-simulated spike-in proteomics data with mixed trend categories

Next, to examine the performance of the methods in more depth, semi-simulated UPS1-based datasets with mixed trend differences in the spike-proteins were generated to reflect typical real longitudinal proteomics data where proteins with multiple different types of longitudinal trends coexist (UPS1 Mix, **Supplementary File 1**).

In the filtered UPS1 Mix datasets without missing values, RolDE with an IQR mean pAUC of 0.948 performed significantly better than the second best method Timecourse with and an IQR mean pAUC of 0.937 (p = 10^−15^, **Figure 2E**). Similarly to datasets with only single trend category per condition, the higher order regression models outperformed the lower order models.

In the full UPS1 Mix datasets including missing values (**Figure 2F**), RolDE with an IQR mean pAUC of 0.944 performed best, slightly outperforming the second best method Timecourse with an IQR mean pAUC of 0.942; however, the overall differences between the two methods were not significant (p = 0.066). In the UPS1 Mix datasets, RolDE and Timecourse clearly outperformed the other methods, with the rest of the methods having IQR mean pAUCs below 0.9 in both the filtered and full datasets.

While RolDE displayed the strongest balanced performance over all the trend categories in the UPS1 Mix datasets, also Timecourse, Limma, LimmaSplines_H and BaselineROTS performed well across the categories (**Supplementary Figure 1E-F**). Similarly as with the single-category datasets, the complexity of the trends largely defined in which categories the regression based approaches performed well; more complex models performed better on a broader spectrum of categories, while the simpler models struggled when the polynomial complexity of the trends increased. Overall, proteins with Linear_Sigmoid and Linear_LogLike trend differences were most challenging across all the methods.

### 2.3 Missing values in the datasets

The original UPS1 and SGSDS datasets contained moderate to low numbers of missing values (14.1% and 3.4%, respectively) and only few missing values in the spike-in proteins (0% and 4.2%, respectively). Therefore, to further push the methods, semi-simulated datasets with a larger proportion of missing values were generate using the CPTAC dataset, which had a larger overall proportion of missing values (19.5%) and a considerably larger proportion of missing values in the spike-in proteins (29.4%) than the UPS1 and SGSDS datasets.

In the presence of such a large proportion of missing values (median of 28.5% for the spike-in proteins in the semi-simulated datasets, **Table 1**) all methods performed clearly worse than in the UPS1 and the SGSDS datasets (**Figure 2G**). RolDE performed overall best with an IQR mean pAUC of 0.747, followed by BaselineROTS with a significantly lower IQR mean pAUCs of 0.729 (p < 10^−15^).

**Table 1.**
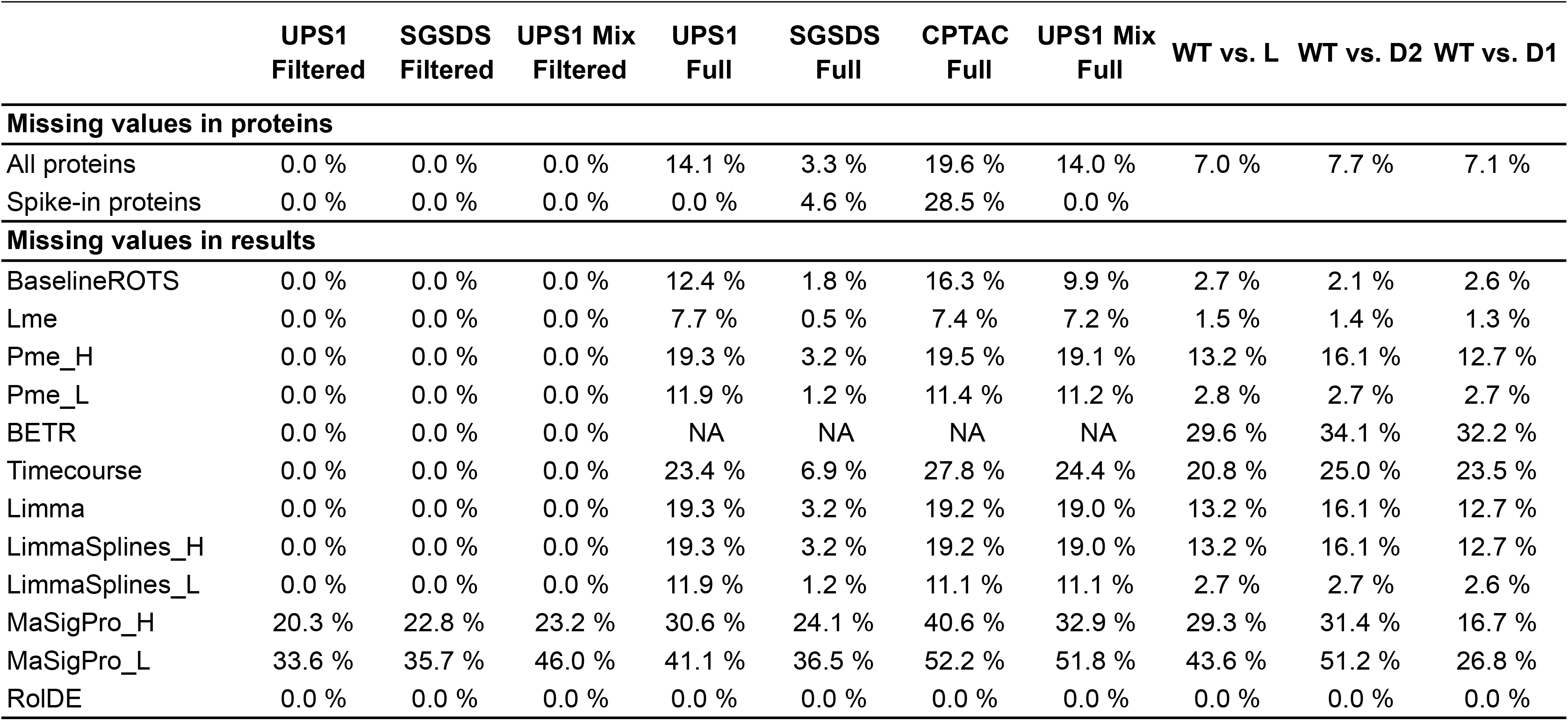
Median proportions of missing values and proportions of missing valid result scores in the semi-simulated spike-in datasets and the experimental *Francisella tularensis* subspecies *novicida* dataset. For the semi-simulated spike-in datasets, interquartile (IQR) mean proportions of missing result scores are shown for each method as representatives of the typical proportion of missing scores for the methods. NA refers to no scores delivered at all by the method.

Furthermore, many of the evaluated methods struggle to provide valid scores for proteins in the presence of missing values in the data. To evaluate how consistently the different methods were able to provide scores for the proteins in the datasets, IQR mean proportions of valid scores for each method and dataset type were calculated. In addition to RolDE, which by default generates a ranking for all the proteins in a dataset, the linear regression method Lme was most often able to provide a valid ranking for the proteins in all the datasets (**Table 1**). Overall, the lower order models, Pme_L and LimmaSplines_L, were able to provide a ranking more often than their higher order counterparts (Pme_H and LimmaSplines_H).

### 2.4 Reproducibility between technical replicate *Fn* datasets

To examine the overall reproducibility of the results produced by the tested methods, technical replicate datasets of the *Fn* data were analyzed separately to detect longitudinal differential expression between all possible pairwise combinations of strains (WT vs. D1, WT vs. D2, WT vs. L, D1 vs D2, D1 vs. L, D2 vs. L).

The overall proportion of missing values in the *Fn* data was 10.8%, being highest in the 37°C samples with more than 25% of all the values missing in all strains but L (**Supplementary Figure 2)**. The proportion of valid rankings provided by the different methods were consistent with the full semi-simulated spike-in benchmark datasets (**Table 1**): RolDE, Lme, BaselineROTS, Pme_L and LimmaSplines_L provided a ranking for more than 97% of all the proteins in each pairwise comparison; Limma, LimmaSplines_H and Pme_H provided a ranking for 84%-87% of the proteins; Timecourse and MaSigPro provided a ranking for less than 80% of the proteins.

The Spearman correlations between the technical replicate lists were on average highest with RolDE and different variants of limma (LimmaSplines_L, LimmaSplines_H and Limma) with IQR mean correlations of 0.793-0.813 (**Figure 3A**); the differences between these methods were not significant (p-values > 0.09). In addition to the overall correlations of the result lists, we examined the median overlaps of the top findings between the replicate datasets, when the size of the overlap was varied from 1 to the total number of proteins in the dataset. In line with the correlation results, RolDE and the different variants of Limma were found to have a high of overlap between their result lists, especially among the top findings (**Figure 3B**).

**Figure 3.**
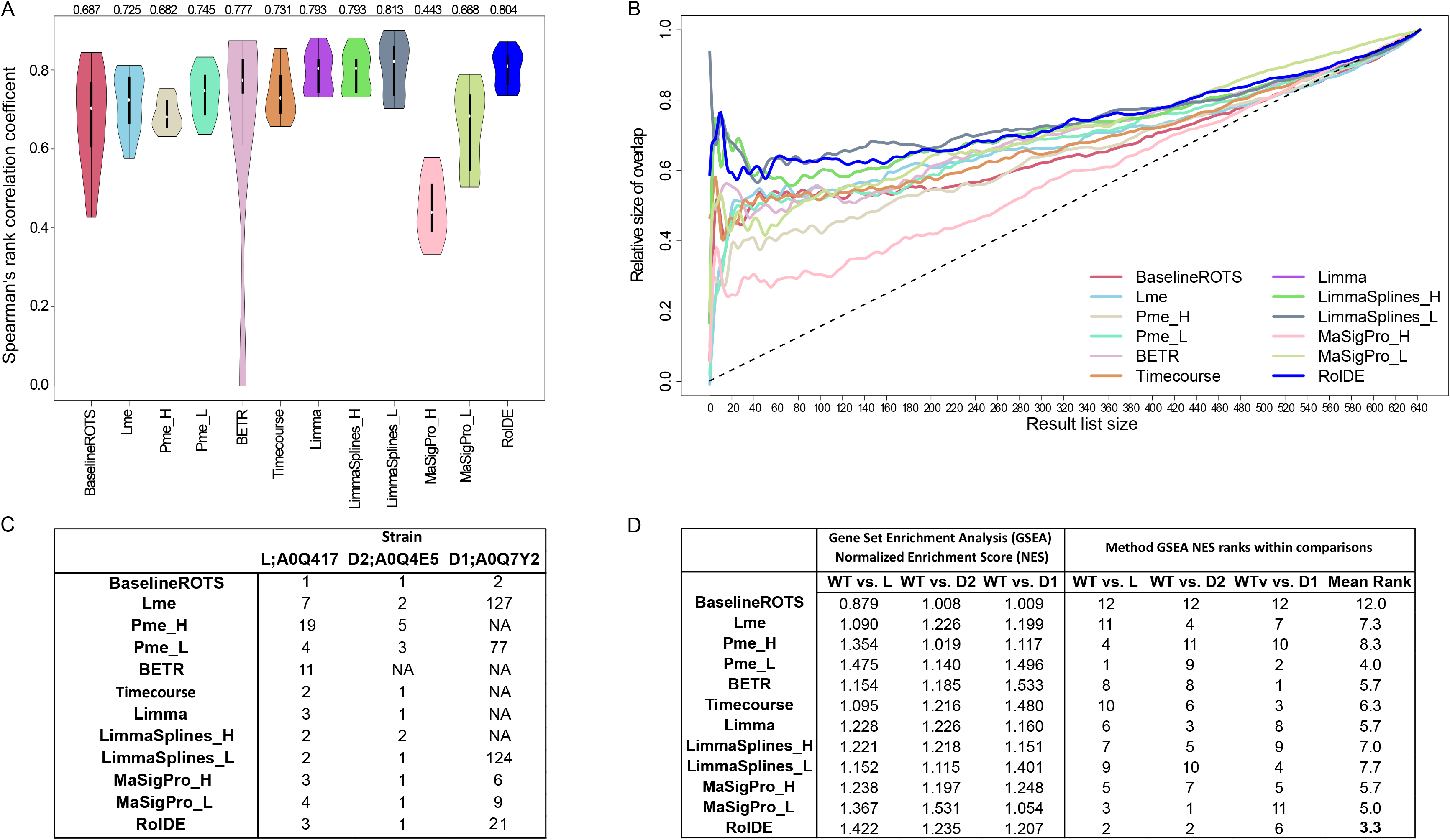
Reproducibility and biological relevance of the examined longitudinal differential expression methods in the experimental *Francisella tularensis* subspecies *novicida (Fn)* proteomics data. **(A)** Reproducibility of the methods was evaluated using the Spearman’s rank correlation coefficient between technical replicate dataset result lists. The violin plots display the population of correlations for each method. Black boxes represent the interquartile ranges, white circles represent median values and wider areas of the violins contain larger densities of values than narrow areas. The interquartile range (IQR) means of the correlations for each method are shown. The IQR means represent typical reproducibility for each method. **(B)** Median proportional overlaps of the top *k* findings between technical replicate dataset result lists. The included number of proteins, *k*, was varied from 1 to the length of the entire dataset. For (A) and (B), all proteins with missing values were filtered out from the dataset prior to the reproducibility testing. Reproducibility of the results were examined over all the possible pairwise comparisons of strains. **(C)** Rankings of the proteins related to the modified acyltransferases in the result lists of the different methods when longitudinal differential expression was examined by each method pairwise between the null mutant strains LpxD1 (D1), LpxD2 (D2), LpxL (L) and the wild type (WT) over the 5 temperatures. The UniProt accessions of each protein related to the modified null mutant is shown in the table headings. NA refers to not detected at all by the method. **(D)** The normalized enrichment scores (NES) from the gene set enrichment analysis (GSEA) of the Lipopolysaccharide synthesis pathway and the associated knockout pathway proteins among the findings of the different methods in comparisons of the acyltransferase null mutant strains and the wild type. 16 proteins from the “KEGG Lipopolysaccharide synthesis pathway (ftn00540)” [69] complemented with 2 proteins from the associated “Lipopolysaccharide biosynthesis knockout pathway (ko00540)” was used as a custom pathway for GSEA [70]. The NES of the methods were ranked within each comparison and a mean rank was calculated over all the comparisons for each method.

### 2.5 Biological relevance of the findings in the *Fn* data

Typically, the top findings in any experiment are the most interesting ones and consistency of a method in delivering the same findings at the top of the list is a highly desirable quality. However, reproducibility alone does not guarantee biological relevance. Therefore, we also examined how the different methods detected the proteins related to the modified acyltransferases in the *Fn* data. For this purpose, longitudinal differential expression over the temperatures was examined pairwise between the wild type (WT) and the null mutants of lpxD1 (D1), lpxD2 (D2) and lpxL (L) in the full data with technical replicates averaged for each biological replicate. The result lists from each method were ordered based on the strength of differential expression given by the method.

When investigating the acyltransferases themselves, most of the methods ranked the proteins related to the acyltransferases lpxL and lpxD2 to the top of their result lists, while more variation was observed in how the methods ranked the protein related to lpxD1, with BETR, PmeFull, Timecourse, Limma and LimmaSplines_H failing to provide a score for the protein (**Figure 3C**).

In addition to the modified proteins themselves, we also investigated how the different methods ranked the related pathways. All the modified acyltransferases were related to the synthesis of lipid A and the endotoxin lipopolysaccharide (LPS). Accordingly, we examined how the altogether 18 unique proteins in the KEGG pathway “Lipopolysacchararide biosynthesis – *Francisella tularensis* subsp. *novicida* U112” (ftn00540) and the associated knockout pathway “Lipopolysaccharide biosynthesis knockout pathway” (ko00540) were detected in our *Fn* dataset. The highest normalized enrichment scores (NES) over all the comparisons were detected in RolDE (**Figure 3D**). In each comparison of the wild type and mutant strains, RolDE consistently provided top findings with high biological relevance when compared to the other methods. Of the other methods, also Pme_L and MaSigPro_L provided highly biologically relevant top results in most comparisons while performing more poorly in some comparisons. Overall, these results suggest the ability of RolDE to consistently provide biologically meaningful results, in agreement to its excellent performance in the semi-simulated spike-in datasets and detecting the modified proteins themselves consistently in different comparisons.

### 2.6 Application of RolDE to longitudinal type 1 diabetes proteomics data with unaligned time points

While the semi-simulated spike-in datasets and the experimental *Fn* data contained perfectly aligned time points (or their surrogates), this is not always the case in various real experimental settings. Therefore, we finally demonstrate the applicability of RolDE in a previously published longitudinal type 1 diabetes (T1D) proteomics data with non-aligned time points [10]. The dataset contains blood plasma protein expression measurements of 11 children developing T1D and 10 matched controls [10]. Nine longitudinal samples were collected for each child with non-aligned time points. The aim was to detect early T1D related biomarkers from the blood even before seroconversion and T1D diagnosis, which would allow earlier disease prediction and intervention.

RolDE detected a total of 23 proteins with longitudinal differential expression between the T1D cases and controls at false discovery rate (FDR) of 0.05 (**Supplementary Table 1**). These included proteins with clear trend differences as well as proteins with clear expression level differences between the cases and controls. Two keratins, KRT31 and KRT86, were detected as highly differentially expressed by RolDE (**Figure 4A, Supplementary Table 1**). Decreased levels of keratins [32] as well as increased keratin metabolism [33] have been previously associated with diabetes and hyperglycemia with keratinocyte proliferation, differentiation and function [34]. Downregulation of another RolDE detected protein (**Figure 4B**), cell growth regulator with EF-hand domain 1 (CGREF1), has been associated with hyperglycemia in humans [35]. Significant decrease in serotransferrin (TF) in T1D cases was also detected by RolDE (**Figure 4C**). TF has been associated with diabetes and glucose metabolism levels in multiple studies [36–40]; Metz et al. reported TF to be downregulated by two-fold in the plasma of T1D patients compared to controls [40]. Serum Amyloid A1 (SAA1, **Figure 4D**), a top differential expression finding of RolDE, has been previously linked to diabetes and blood glucose levels in multiple studies [41–43]. Furthermore, sodium channel and clathrin linker 1 (SCLT1) was detected as highly significant by RolDE (**Figure 4E**). It has been linked to ciliopathies in human [44] and many ciliopathies have been linked to obesity and insulin secretion in the pancreas by the regulating effect of pancreatic cili [45].

**Figure 4.**
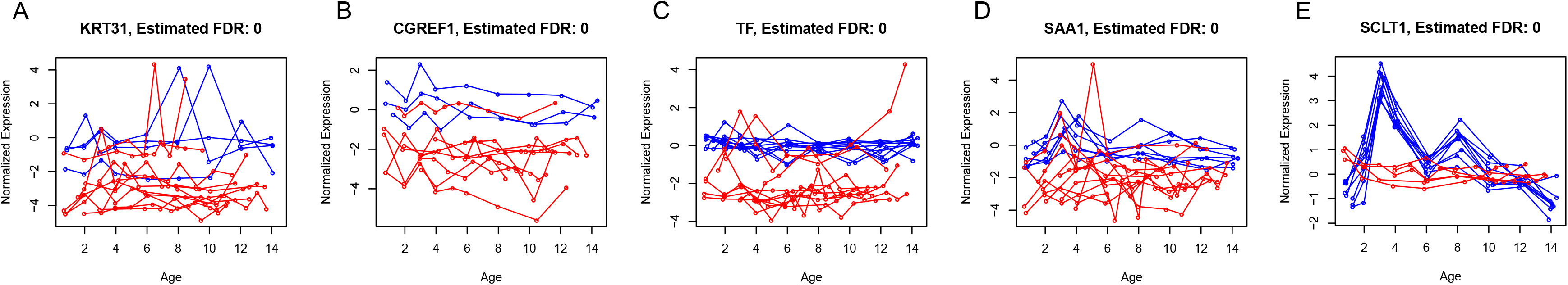
Longitudinal expression of representative examples of the top differential expression detections by the Reproducibility optimized longitudinal Differential Expression method RolDE in the longitudinal type 1 diabetes blood plasma proteomics data of Liu et al [10]. Blue color represents individuals in the control group while red color corresponds to children in the type 1 diabetes group.

Finally, we compared the RolDE results to the results from the original study by Liu et al. [10]. Using mixed effects modelling with linear and quadratic random effects in addition to the fixed effects, the original study detected two proteins with differential expression between T1D cases and controls at FDR of 0.05 [10] [Supplementary Table 3]. One of these, SCLT1, was also detected by RolDE, whereas the other one, CBR1 (**Supplementary Figure 3**), had FDR of 0.12 with RolDE.

Taken together, these top findings clearly demonstrate the ability of RolDE to detect proteins with various kinds of differential expression patterns also in data with non-aligned time points.

## 3. Discussion and conclusions

We have comprehensively evaluated the performance of eleven previously established approaches together with our new method RolDE for detecting differential expression between two conditions in longitudinal omics data. Altogether 1920 semi-simulated proteomics benchmark datasets were used to evaluate the ability of the different approaches to detect the truly differentially expressed spike-in proteins with various longitudinal expression patterns and differences, both with and without missing values present in the data. Furthermore, the reproducibility and biological relevance of the results of the different methods was evaluated using a longitudinal experimental dataset based on mutants of *Francisella tularensis* subspecies *novicida* at various temperatures known to change the expression of proteins in the Raetz pathway responsible for Lipid A Biosynthesis [46]. Additionally, we demonstrated the ability of RolDE to detect differential expression even in data where the time points in different conditions are not aligned using a previously published T1D dataset [10].

Overall, our new proposed robust longitudinal differential expression method RolDE performed best in the spike-in datasets, including datasets with missing values in the true positive spike-in proteins (**Figure 2D,G**). Missing values are a typical occurrence in proteomic data, especially in the popular label-free data dependent acquisition (DDA) approach. Furthermore, as the proteins of interest are typically unknown beforehand and may also contain missing values (such as in the SGSDS datasets or in the *Fn* dataset), the ability of a differential expression method to deal with versatile missing values might be crucial in detecting relevant proteins to the experimental question. RolDE also displayed the most balanced performance of detecting trend differences of all possible types with good consistency (**Supplementary Figure 1**). It is typical that a researcher does not know beforehand what types of differences in longitudinal expression are to be expected during the course of a study. The ability of a method to detect various types of trend differences is thus essential for a comprehensive analysis of the data. In addition, RolDE had good overall and top reproducibility (**Figure 3A-B**), was among the top methods in ranking the results in a biologically meaningful way in the experimental *Fn* dataset (**Figure 3C-D**) and delivered interesting findings even when the time points in the data were not aligned (**Figure 4**).

Timecourse, a longitudinal differential expression method developed for microarrays and utilizing a Bayesian framework, performed also generally well on the tested datasets. However, missing values in the spike-in proteins clearly hindered the performance of the method and Timecourse was able to provide a ranking only for ~72%-79% of proteins in the UPS1 and CPTAC datasets and the experimental *Fn* data when missing values were present (**Table 1**). Furthermore, Timecourse was not able to deliver a ranking for all the modified acyltransferases known to change between the conditions in the *Fn* dataset. A method that is able to deal with different kinds of missingness in the data might be impractical in the context of proteomics data, where missing values can occur due to various reasons [18,47,48]. Timecourse does not provide a significance estimate, which might also limit its usability.

Limma provides two different approaches to detect longitudinal differential expression. In general, the group mean parametrization approach (Limma) performed similar to Limma with cubic splines (LimmaSplines) when the data did not contain missing values. Some differences in performance could be observed when missing values were present in the data. In particular, even though performing worse in datasets with no missing values and when missing values were present only in the background proteins (UPS1-based datasets), the Limma regression splines approach with less degrees of freedom (LimmaSplines_L) was best among the Limma-based approaches when missing values were present also in the spike-in proteins in the SGSDS and CPTAC datasets (**Figure 2**).

There was considerable variation in the performance of the various regression modelling approaches tested. Overall, the higher order polynomial regression models outperformed the lower degree models. Only in the SGSDS full datasets, the overall performance of Pme_L was better than the performance of Pme_H. This reversal in performance was most likely related to missing values in the spike-in proteins in the SGSDS full datasets and the inability to reliably define the more complex models related to Pme_H for all the spike-in proteins. Furthermore, the performance of the full regression modelling approach was better over a broader spectrum of trend difference categories than the performance of the lower degree models (**Supplementary Figure 1**). The degree of the regression was directly associated to how complex trend differences could be detected between the conditions by the models, as expected. All polynomial regression model approaches utilized orthogonal polynomials. This likely allowed the detection of both lower and higher order differential trend patterns simultaneously by reducing collinearity between the coefficients of different polynomial degrees as compared to using raw polynomials [49,50].

The overall reproducibility of the results in the biological *Fn* data as well as the reproducibility of the top results was best with RolDE and the Limma-based approaches. The relevance of the findings in terms of the modified proteins and the directly associated KEGG “Lipopolysaccharide Biosynthesis” and “Lipopolysaccharide Biosynthesis knockout pathway” proteins was highest with RolDE, Pme_L and MaSigPro_L. LPS is an the essential part of the Gram negative bacteria outer membrane and forms an amphipathic interface between the bacteria and the environment [51,52]. LPS is composed of three structural parts: the core polysaccharide, O-antigen, and the membrane anchor lipid A [51]. Modifications of lipid A can alter the structure and pathogenicity of the bacterium and are influenced by environmental conditions such as temperature [53]. In *Fn*, temperature has been shown to alter the composition of lipid A [53]. The chosen KEGG pathway focusing on LPS biosynthesis therefore served as valuable readout for exploring and validating the performance of the different methods and also highlight the consistency of the generated proteomics data within our understanding of *Fn* biology field as a whole.

Most of the examined approaches in this study require a suitable polynomial degree or degrees of freedom to be defined. However, our analysis shows that, through the use of orthogonal polynomials, while not completely removing issues related to collinearity, differential expression related to different polynomial degrees can be effectively detected simultaneously within a single model (**Supplementary Figure 1**). Moreover, the performance of RolDE was not overly sensitive to the degree used. By default, RolDE defines the degrees by itself, which should be suitable for most use cases. However, the user is given full control of RolDE through modifying the default parameters.

We provide RolDE as a freely available R package in GitHub (https://github.com/elolab/RolDE). The default use of RolDE requires the user to provide only a design matrix linking the samples in the data to the time points and conditions, and information whether the time points are aligned or not. Importantly, the user does not need to specify what kind of differences are searched from the data, what degree should be used, or any other prior information. For more experienced users, however, we also provide an opportunity to utilize RolDE in a more custom-tailored way in various experimental settings.

RolDE is a relatively fast method. One semi-simulated UPS1 dataset with 1581 proteins and 30 samples can be analyzed in ~2 minutes with a standard desktop computer using parallel processing and 3 threads (Intel i7-6700, 32GB Ram). The slightly larger SGSDS dataset with 3487 proteins and 48 samples takes ~3 minutes. The analysis of the non-aligned longitudinal dataset of Liu et al. [10] with 2235 proteins and 189 samples lasted ~5 minutes using 7 threads on the same computer.

To summarize, RolDE is an easy-to-use, relatively fast longitudinal differential expression method with excellent performance applicable for both time point aligned as well as non-aligned longitudinal data in various types of experimental settings.

## 4. Methods

### 4.1 Semi-simulated spike-in proteomics datasets

Three spike-in proteomics datasets were used as a basis for generating the semi-simulated longitudinal spike-in proteomics datasets.

#### The Universal Proteomics Standard Set (UPS1) data

The UPS1 spike-in data includes 48 Universal Proteomics Standard Set (UPS1) proteins spiked into a yeast proteome digest with five varying concentrations: 2, 4, 10, 25 and 50 fmol/μl [29]. Three technical replicates of each concentration were analyzed using a LTQ Orbitrap Velos MS. After preprocessing, 47 spike-in proteins and 1581 proteins remained in the UPS1 data.

#### The Shotgun Standard Set Dataset (SGSDS)

In the SGSDS spike-in data, 12 nonhuman proteins were spiked into a stable human background (human embryonic kidney, HEK-293) in eight different sample groups with known concentrations [54]. Three technical replicates of each sample group were analyzed using a Q Exactive Orbitrap MS both in the DDA and data-independent acquisition (DIA) modes. In this study, the DDA shotgun proteomics data was used. After preprocessing, all 12 spike-in proteins and 3487 proteins remained in the SGSDS data.

#### The Clinical Proteomic Tumor Analysis Consortium (CPTAC) dataset

The CPTAC data from study six contains 48 UPS1 proteins spiked into stable yeast proteome digest in five different concentrations: 0.25, 0.74, 2.2, 6.7 and 20 fmol/μl [55]. Three technical replicates of each sample group were analyzed using a LTQ Orbitrap mass spectrometer (test site 86). After preprocessing, 41 spike-in proteins and 1247 proteins remained in the CPTAC data.

The default peak-picking settings were applied to process the raw MS files of all the datasets in MaxQuant [56] version 1.5.3.30. Peptide identifications were performed using the Andromeda search engine [57]. Match type was ‘match from and to’. Match time window size was by default 0.7 min, and alignment time window size was 20 min. ‘Require MS/MS for comparisons’ was on, and decoy mode was ‘revert’. False discovery rate (FDR) of 0.01 was set as a threshold for peptide and protein identifications. MaxQuant was allowed to automatically align the runs. A FASTA database of the yeast *Saccharomyces cerevisiae* protein sequences merged with the Sigma-Aldrich 48 UPS1 protein sequences was used to search protein identifications for the UPS1 and CPTAC data. For the SGSDS data, a FASTA database of the human HEK-293 cell proteins merged with the sequences of the nonhuman spike-in proteins was used.

MaxLFQ was on. ‘Advanced ratio estimation’, ‘stabilize large LFQ ratios’ and ‘advanced site intensities’ were on. Non-normalized protein intensities were extracted from MaxQuant and imported into the R statistical programming software environment. The datasets were normalized using the variance stabilization normalization (vsn) [58] shown to perform well with proteomics spike-in data [12].

### 4.2 Generation of the semi-simulated datasets with varying longitudinal trends

The normalized spike-in datasets were used to create semi-simulated datasets with varying pre-defined longitudinal trends in the spike-in proteins (**Figure 1B**). Unlike the spike-in proteins, the expression of the background proteins was expected to remain stable in all sample groups. It should be noted, however, that due to experimental noise and fluctuations, there is always some variation also in the abundance of the background proteins, which reflects the nature of MS data in a real biological experimental setting.

For the creation of the semi-simulated datasets, the means and standard deviations of the proteins in the sample groups of the original normalized spike-in data were used. More precisely, the simulated expression of protein *i* in sample group *j* for each replicate was drawn from a normal distribution 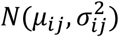, where the protein and sample group specific mean μ_*ij*_ and variance 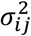 were calculated from the original spike-in dataset. The longitudinal trends were created by reorganizing the semi-simulated sample groups into desired combinations. For example, to create a simple linear trend using the UPS1 data, we sequentially combined the simulated 2 fmol to 50 fmol sample groups. This approach allowed the generation of a plethora of semi-simulated proteomic datasets with different longitudinal trends for the spike-in proteins as well as a constant but noisy background and, therefore, a realistic benchmarking of the longitudinal methods.

Six basic trend categories were introduced for the UPS1 and CPTAC datasets: Stable, Linear, LogLike, Sigmoid, Poly2, and PolyHigher (**Supplementary File 1**). For the SGSDS datasets, five basic trend categories were used: Stable, LogLike, Sigmoid, Poly2, and PolyHigher. For the SGSDS datasets, a longitudinal linear increase or decrease (Linear) was unattainable due to the uneven concentration differences of the spike-in proteins between the samples in the original data [54]. In total, this approach resulted in 21 trend difference combinations between two simulated conditions for the UPS1 and CPTAC datasets and 15 combinations for the SGSDS dataset. When each dataset contained only one type of trend for a condition, a total of 300 semi-simulated datasets with two conditions and different longitudinal trends and/or expression levels were generated for the UPS1 and CPTAC datasets and 210 for the SGSDS dataset (**Supplementary File 1**).

Additionally, we generated 300 semi-simulated datasets with varying trends within a condition and varying trend differences between the conditions to reflect typical real experimental data where proteins with multiple different types of longitudinal trends coexist. These datasets were generated by randomly selecting 10 different trend differences for the spike-in proteins using the UPS1 dataset, referred to as UPS1 Mix in the results section (**Supplementary File 1**).

For benchmarking the methods, two versions of each semi-simulated UPS1 and SGSDS dataset were generated, referred to as *full* and *filtered* in the results section. In the full datasets, none of the proteins were filtered from the data (i.e. proteins with missing values were included), whereas the filtered datasets contained only proteins with no missing values.

Finally, as missing values are a common occurrence in proteomics data [17,18,48], the CPTAC dataset with relatively large numbers of missing values in the spike-in proteins was used to generate semi-simulated datasets with high proportions of missing values to explore the tolerance of the different methods against such missing values. Here, only *full* datasets were considered.

### 4.2 Longitudinal *Francisella* proteomics data

To evaluate the longitudinal differential expression methods in a real proteomics study setting, we generated and used an experimental membrane enriched longitudinal proteomics data of *Francisella tularensis* subspecies *novicida (Fn)* (**Figure 1C**). For evaluation of the methods, differential expression between the null mutant and the wild type strains was investigated, while their growth at different temperatures offered a surrogate for the different time points. Technical replicates of the samples were used to form technical replicate datasets analyzed with the different methods to evaluate the reproducibility of the methods. In addition, the *Fn* dataset with known protein modifications, was used to explore the evaluated methods ability to detect these proteins and other closely associated pathway proteins expected to be affected.

*Francisella tularensis* subspecies *tularensis (Ft)* is a highly pathogenic Gram negative bacterial agent responsible for the disease tularemia in humans. Research interest on *Ft* has peaked during the past decades due to the possible application of the bacterial agent in biological warfare and bioterrorism [59]. As reliable methods for the genetic manipulation of the highly virulent subspecies *Ft* have been lacking, the more responsive and avirulent subspecies *Fn* was used as a proxy in an effort to shed light into the mechanisms of pathogenicity of *Ft* [59]. We considered two temperature regulated N-acyltransferases, designated lpxD1 and lpxD2, which have been identified in *Fn*. LpxD is involved in the production of a key virulence factor, lipid A, in *Fn* [52]. The null mutant of lpxD1 has been shown to be more sensitive to environmental factors and attenuated in virulence when compared to the wild type [52,53]. In addition, we studied a late acyltransferase lpxL, which has been discovered in *Ft* [60] and shown to be essential for cell viability at temperatures above 33°C [60]. Thus, the dataset consisted of four strains [61]: the wild type (WT) and null mutants of lpxD1 (D1), lpxD2 (D2) [53] and lpxL (L). In order to activate the temperature sensitive enzymes responsible for the production of the lipopolysaccharide, the global protein expression of the *Fn* strains was examined in five temperatures: 18°C, 21°C, 25°C, 32°C, 37°C (surrogates for time points). Given that LPS is assembled in the membrane of Gram negative bacteria, we chose to isolate and generate data only for a membrane-enriched fraction.

Each mutant and wild type strain consisted of three biological replicates (surrogates for individuals) of membrane only fractions of proteins, and each biological replicate was measured in three technical replicates resulting in a total of 180 unique samples. The membrane samples were prepared for proteomic analysis by high performance liquid chromatograpghy (HPLC)-tandem mass spectrometry [62] according to previously published protocols [63] for analysis on an LTQ Orbitrap Elite.

The raw MS data files were processed with the MaxQuant [56] software (version 1.6.5.0) and similar settings to the spike-in datasets (section 2.1). A SwissProt/UniProt [64] FASTA database of all the reviewed and unreviewed protein sequences for *Francisella tularensi*s subspecies *novicida* strain U112 (April 2019) was used for the peptide and protein identifications. Trypsin digestion with a maximum of two missed cleavages, carbamidomethylation of cysteine as a fixed modification, methionine oxidation and N-terminal acetylation as variable modifications were as search parameters.

Non-normalized protein intensities were extracted from the MaxQuant output and normalized using vsn [58] shown to perform well with proteomics data [12]

For more details concerning the *Fn* sample preparation and dataset generation, see **Supplementary Methods**.

### 4.4 Longitudinal human type 1 diabetes proteomics data

Finally, we evaluated the performance of RolDE on human type 1 diabetes (T1D) proteomics data from a recent study by Liu et al. [10] (**Figure 1D**). The data contained longitudinal blood plasma protein expression measurements from 11 children (aged 0.8-14.4 years) developing T1D and 10 matched controls. Nine longitudinal samples were collected for each child, with the T1D case samples covering seroconversion and clinical T1D diagnosis. The samples were analyzed using a TMT-10plex-based LC-MS/MS approach with a Q Exactive HF mass spectrometer with all time points of one individual and a common pooled reference sample included in one TMT-10plex MS run [10]. Altogether, 189 samples were analyzed. The data was normalized and filtered similarly to the original study: the reporter ion intensities were standardized to the reference sample, log2-transformed and median normalized, and only proteins with at least two individuals from either group with at least seven non-missing values were included in the analysis [10].

### 4.5 Robust longitudinal Differential Expression (RolDE)

The proposed new method, RolDE, is a composite method, consisting of three independent modules with different approaches to detect longitudinal differential expression (see **Figure 1A** for a schematic illustration). While each module typically performs well already on its own in ranking the true findings high, there can also be some additional unwanted noise detections (false positives) associated with each module. Since such false positives are typically specific for a single module, combining the modules creates a balanced composite method, enriches the true signal at the top of the results and allows RolDE to robustly detect varying differences in longitudinal trends and expression levels in diverse data types and experimental settings. The three modules of RolDE, named *RegROTS*, *DiffROTS* and *PolyReg*, and their combination are described in more detail below.

#### 4.5.1 RegROTS

The RegROTS module combines the power of polynomial regression modelling and the reproducibility optimization of ROTS [26,27]. First, a polynomial regression model for each protein is separately fitted for each individual *u* over time *t*:

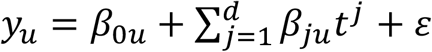

where *d* is the polynomial degree of the model, β_*ju*_ are the regression coefficients, and ε is the error term. The degree *d* is by default defined as: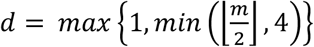 where *m* is the median number of time points over all the individuals. Orthogonal polynomials are used to reduce multicollinearity and allow for more independent exploration of coefficients of different polynomials of the same variable [49,50].

Next, all coefficients of the same degree *j* are compared across the individual models by considering all possible pairs of individuals *u* and *v* between the two conditions *C*_1_ and *C*_2_ under comparison: Δβ_*juv*_ = β_*ju*_ − β_*jv*_, where *u* ∈ *C*_1_ and *v* ∈ *C*_2_. To preserve the proper degrees of freedom for statistical testing, the replicate comparisons are divided into multiple different *runs* so that each individual is used only once in each run. For instance, with three individuals in both conditions, we consider three runs with the following combinations: run 1 *u_1_-v_1_, u_2_-v_2_, u_3_-v_3_*; run 2 *u_1_-v_2_, u_2_-v_3_, u_3_-v_1_*; and run 3 *u_1_-v_3_, u_2_-v_1_, u_3_-v_2_*. Within each run, multigroup ROTS is used to detect differences in the estimated coefficients between the conditions with the null hypothesis that there are no differences between the conditions, i.e. Δβ=_0_ Δβ_1_ = ⋯ = Δβ_*d*_= 0. Finally, the results from the different runs are combined using the geometric mean of the ranks of the significance values from each run *R_i_*, producing the final score *S*_*RegROTS*_ for a protein in the RegROTS module:

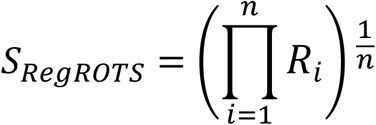

where *n* is the number of runs.

#### 4.5.2 DiffROTS

The DiffROTS module examines directly the expression differences between the conditions at each time point. Similar to the RegROTS module, all the replicates in condition *C*_1_ are compared to all the replicates in condition *C_2_*, but instead of comparing the coefficients of the fitted models, the expression measurements *y*_*tu*_ for each individual *u* at time point *t* are used directly: Δ*y*_*tuv*_ = *y*_*tu*_ − *y*_*tv*_. Within each run, multigroup ROTS is is used to detect differences in the expression levels between the conditions with the null hypothesis that there are no differences between the conditions, i.e. Δ*y*_1_= Δ*y*_2_ = ⋯ = Δ*y*_*T*_= 0, where *T* is the total number of time points considered. If the time points in the data are not aligned between the individuals, the expression levels between conditions are estimated after polynomial interpolation. Finally, the results from the different runs are combined using the geometric mean of the ranks of the significance values from each run *R_i_*, producing the final score *S*_*DiffROTS*_ for a protein in the DiffROTS module:

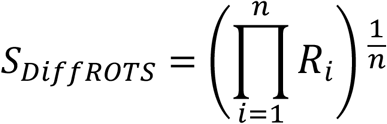

where *n* is the number of runs.

#### 4.5.3 PolyReg

The PolyReg module applies polynomial regression modelling to each protein to detect longitudinal differential expression over time *t* across the conditions *c*:

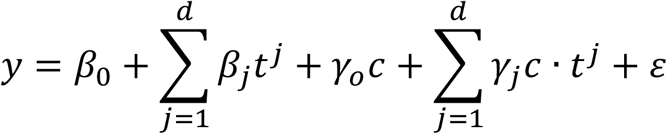

where *d* is the polynomial degree of the model, β_*j*_ and γ_*j*_ are the time and condition-related regression coefficients, respectively, and ε is the error term. Individual variation can be taken into account by adding a random effect for the individual baseline or slope. The degree *d* is by default defined as *d* = *max*{2, *min*(*m* − 1, 5)}, where *m* is the median number of time points over all the replicates. To detect any differential expression between the conditions, the minimum over the significance values of the condition-related regression coefficients *p*(γ_*j*_) is used as the final score *S*_*PolyReg*_ for a protein in the PolyReg module:

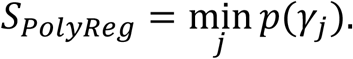

#### 4.5.4 The composite method RolDE

Finally, for a comprehensive inspection of differential expression between the conditions, the results from the different modules are combined using the geometric mean of the ranks of the module scores:

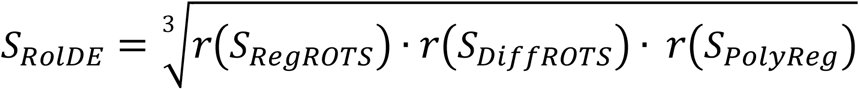

To estimate the significance of the RolDE score, a simulation procedure is used. Given the null hypothesis is true, the significance value distribution is (approximately) uniform. Based on this, a simulated internal rank product is calculated for the RegROTS and DiffROTS modules as follows. First, an equal number of simulated significance values as there are experimental significance values in the corresponding run are generated from the uniform distribution. Second, the simulated significance values within each run are ordered so that the ranks of the proteins according to the experimental significance values are retained to account for the dependencies between runs. Finally, each simulated significance value within each run is replaced with the rank of the closest experimental significance value and these are used to calculate the simulated internal rank products. Given the null hypothesis is valid, the experimental and simulated internal rank products are similar (**Supplementary Figure 4A**). For the PolyReg module, the representative simulated significance values are calculated in a similar fashion. After acquiring the simulated internal rank products or representative significance values for each module, final simulated rank products are calculated equivalently to the experimental rank products and utilized to estimate the overall significance values. This is done by generating 500000 (by default) simulated rank products and then calculating the fraction of simulated rank products smaller or equal to each experimental rank product. Given the null hypothesis is valid, the estimated significance value distribution for RolDE will be approximately uniform (**Supplementary Figure 4B**). The estimated significance values are adjusted for multiple hypothesis testing by using the Benjamini-Hochberg procedure [65]. Alternatively, Bonferroni correction, Q-value adjustment [66] or any of the other methods included in the R statistical programming language stats package can be used.

### 4.6 Existing methods for detecting longitudinal differential expression tested

#### Reproducibility Optimized Test Statistic (BaselineROTS)

ROTS is a well-established differential expression method aiming to maximize the reproducibility of the top detections using a modified t-statistic and group preserving bootstraps [26,27]. It has been observed to perform well in multiple types of omics data, including proteomics [12,28–30]. In this study, ROTS was used as a baseline cross-sectional method at each time point against which the longitudinal methods were compared to. Version 1.12.0 of the Bioconductor R package ROTS was used in the study.

#### Bayesian Estimation of Temporal Regulation (BETR)

BETR utilizes the Bayesian framework and was first introduced for the analysis of times series DNA microarray data [13]. It calculates the probability of differential expression for each feature (e.g. gene or protein), taking correlations between time points into account; the magnitude of expression at time points closer to each other are assumed to be more correlated than those further apart [13]. Version 1.32.0 of the Bioconductor R package betr was used in this study.

#### Linear Models for Microarray Data (Limma, LimmaSplines)

Limma is a popular toolset used especially in the analysis of gene expression microarray and RNA-seq data. Longitudinal differential expression can be examined in two different ways [16]. The first option is through group-mean parametrization, where the design matrix for the linear models is designed is such a way that a separate coefficient exists for each time point and condition combination and various contrasts between the coefficients can be defined to examine differential expression between the conditions. We utilized a similar approach as described by [67], where the contrasts were defined as differences in expression between the conditions at each time point and differential expression between the conditions was determined by examining whether all contrasts were zero simultaneously using the moderated F-statistic. The second option is to use regression splines as described in the limma R package user manual [16]. Version 3.40.2 of the Bioconductor limma R package was used in this study.

#### Timecourse (Timecourse)

Timecourse ranks features (e.g. genes or proteins) according to probabilities for differential expression using the Maxwell–Boltzmann or the Hotelling’s T^2^ statistics through a multivariate empirical Bayes approach, taking replicate variances and correlations in expression levels between time points into account [14]. The method borrows information across features to better estimate the variance-covariance matrices to reduce the number of false positives and false negatives [14]. Version 1.56.0 of the Bioconductor timecourse R package was used in this study.

#### Microarray Significant Profiles (MaSigPro)

MaSigPro is a method originally developed for the analysis of time series gene expression microarray data [15]. It follows a two-step regression approach to detect longitudinal differential expression between the conditions [15]. In the first step, a general regression model is defined for each feature (e.g. gene or protein). In the second step, only the significant models from the first step are then modelled using polynomial regression to find differences between the compared conditions [15].

Version 1.56.0 of the Bioconductor maSigPro R package was used in this study. For each protein, a minimum over the condition-related significance values was used as a representative significance value.

#### Linear mixed effects regression modelling (Lme)

Linear mixed effects regression is a popular approach in modeling longitudinal data, where individual variability can be incorporated into the model in the form of random effects [10,68]. We considered two different types of models. In the first variant, a random effect only for the individual baseline (the intercept) was allowed. In the second variant, a random effect was allowed also for the slope. Finally, a likelihood ratio test was conducted to examine if adding a random effect for the slope yielded a significantly better fit at the significance level of 0.05. If this was not the case or the second model could not be defined due to insufficient number of measurements (caused e.g. by missing values), the model with a random effect only for the intercept was used for the protein. For each protein, a minimum over the condition-related significance values was used as a representative significance value.

#### Polynomial mixed effects modelling (Pme)

In the polynomial mixed effects approach, we added polynomial fixed terms for the time-related effects in the models. Again, two different types of models were considered: models with a random effect only for the individual intercept and models with a random effect also for the slope. Similarly as with Lme, a likelihood ratio test was conducted to examine if adding a random effect for the slope yielded a significantly better fit at the significance level of 0.05. We did not consider random effects for the higher order polynomials as there was typically not enough data in the relatively short time series to define such models. For each protein, a minimum over the condition-related significance values was used as a representative significance value.

For each polynomial regression-based approach (LimmaSplines, MaSigPro, Pme), we explored two different levels of complexity: a more complex model (denoted by the extension H in the results) with the degree of the polynomial set to *T-1*, and a less complex one (denoted by the extension L) with the degree set to ⌊*T*/2⌋, where *T* is the number of time points.

### 4.7 Evaluation of the methods in the semi-simulated spike-in proteomics datasets

In the semi-simulated datasets, we evaluated the performance of the longitudinal methods in their ability to correctly detect true differential expression (the spike-in proteins) using receiver operating characteristic (ROC) analysis. In the ROC analysis, the sensitivity (i.e. true positive rate) was plotted against the specificity (i.e. true negative rate). The area under the ROC curve (AUC) was used to measure how well a given method was able to distinguish the true signal of interest when the detection threshold (e.g. significance value) was varied. As typically the interest of a differential expression analysis is focused on the top findings, we used a partial AUC (pAUC) between specificity values 1 and 0.9 to score the methods on the essential part of the ROC curve. The calculated pAUCs were summarized using the interquartile range (IQR) mean across the datasets of each type.

Different methods had varying abilities to calculate a score (a test statistic or a ranking) for the examined proteins due to missing values or other reasons. To ensure comparability, a full result list was expected from each method, including a score for all the proteins in the examined dataset. If a method could not produce a score for a protein, a random score larger than the maximum observed score for the method in the dataset was generated, placing the protein at the end of the result list. Thus, all proteins without valid scores for a given method, were placed randomly at the end of the result list.

### 4.8 Reproducibility and biological relevance in the longitudinal *Francisella* data

In the experimental longitudinal *Fn* dataset, we assessed the reproducibility of the results of the methods using the three technical replicate datasets. After filtering out all proteins with missing values, three completely separate technical datasets were formed, which were assumed to be similar to each other. Longitudinal differential expression between all possible pairwise combinations of strains (WT vs. D1, WT vs. D2, WT vs. L, D1 vs D2, D1 vs. L, D2 vs. L) were considered. To estimate the overall reproducibility of each method, the similarity of their outputs in the replicate datasets was assessed using the Spearman’s rank correlation coefficient. To evaluate the reproducibility of the top differential expression findings, the median proportional overlap between the top *k* findings was calculated over the replicate datasets, when the examined top list size *k* was varied from 1 to the number of proteins in the complete dataset. The proportional overlap at each value of *k* was calculated as the median overlap over the replicate datasets divided by *k.*

To examine the biological relevance of the findings by each method in the *Fn* dataset, we examined how the proteins in the KEGG pathway [69] “Lipopolysaccharide biosynthesis - *Francisella tularensis* subsp. novicida U112 (ftn00540)” complemented with the proteins in the associated “Lipopolysaccharide biosynthesis knockout pathway (ko00540)” (**Supplementary Table 2**) were ranked by the different methods. One of the key virulence factors in *Francisella* is lipid A, which is part of the endotoxin lipopolysaccharide (LPS), whereas LPS is an essential part of the Gram negative bacterial outer membrane and forms an amphipathic interface between the bacteria and the environment [51,52]. As the modified null mutants of acyltransferases in the *Fn* data (D1, D2, L) are associated with lipid A and LPS, the 18 proteins in the selected ftn00540 and ko00540 pathways are assumed to be affected by the modifications. To explore this, we used Gene Set Enrichment Analysis (GSEA) [70] to investigate enrichment of the pathway proteins in each wild type to mutant strain comparison (D1, D2, L) in the normalized data with technical replicates averaged for each biological replicate. The result lists for each method in each comparison were ranked according to the strength of the provided differential expression score and the ranks were used for preranked GSEA. Similarly to calculating the pAUCs, if a method failed to provide a score for a protein (due to missing values or other reasons), random ranks larger than the maximum observed rank for the method in the comparison was generated for such proteins, placing these proteins at the end of the result list. Normalized Enrichment Scores (NES) from the GSEA analysis within each comparison were used to reflect the methods ability in detecting the relevant pathway proteins and in producing biologically meaningful results.

## Author contributions

TV planned the computational experiments, developed the RolDE method, performed the computational analysis, interpreted the results and drafted the first version of the manuscript.

TS planned the computational experiments, participated in the computational analysis and method development, interpreted the results and participated in writing the manuscript.

C.E.C planned the experiments, generated the experimental *Fn* data and participated in analyzing the *Fn* data and writing the manuscript.

A.J.S constructed the mutants, planned the experiments for the *Fn* data and interpreted the Fn data.

B.Q.T was responsible for the experimental design and data acquisition of the *Fn* data.

R.K.E planned the experiments for the *Fn* data, interpreted the *Fn* data and participated in writing the manuscript.

D.R.G planned the experiments for the *Fn* data, interpreted the *Fn* data and participated in writing the manuscript.

L.L.E conceived, designed and supervised the study, participated in the method development, interpreted the results and participated in writing the manuscript.

## Data availability

The UPS1 spike-in dataset is archived in the PRoteomics IDentification Database (PRIDE) and is available with the identifier PXD002099.

The SGSDS profiling standard is available from PeptideAtlas: No. PASS00589 (username PASS00589, password WF6554orn).

The CPTAC spike-in dataset (study 6, at test site 86) is available from the CPTAC Portal.

The new generated Francisella novicida *(Fn*) dataset has been deposited to the ProteomeXchange Consortium via the PRIDE partner repository and will be made available upon publication with the dataset identifier PXD025439.

## Code availability

The new proposed method RolDE, is freely available as an R package in GitHub (https://github.com/elolab/RolDE).

## Acknowledgements

We thank Esko Pakarinen for the technical support provided during the course of the project.

