## Supplementary Material for "Enhanced longitudinal differential expression detection in proteomics with robust reproducibility optimization regression"

### Supplementary Methods

#### *Francisella tularensis* subspecies *novicida* longitudinal proteomics dataset

##### Bacterial strains and growth conditions

Wild-type *Franicsella tularensis* subspecies *novicida* (Fn) strain U112 was originally obtained from Professor Francis Nano (University of Victoria, Canada) [1] and maintained at -80C. The lpxD1, lpxD2 [2], and lpxL-null mutants were generated previously [3]. The lpxL strain used was a transposon insertion mutant of FTN_0071 (*lpxL*) identified as: tnfn1_pw060510p04q127 carrying a kanamycin cassette disrupting the coding sequence. The mutant was functionally characterized as having an lpxL knockout lipid A phenotype. All bacterial strains were grown in tryptic soy broth supplemented with 0.1% cysteine at the designated temperatures (18°C, 21°C, 25°C, 32°C, 37°C). Cultures were harvested at log phase, supernatant was aspirated, and cell pellets were flash frozen and stored at -80C until processing.

##### MS Analysis

To isolate a fraction of proteins enriched in membrane proteins, cell pellets were first fractionated along the lines of a prior report [4] with the following adjustments. Briefly, cell pellets were resuspended in 0.1M NaPO4, 0.05M MgSOD4, DNaseI digested, and sonicated. Unbroken cells were removed by centrifugation and the supernatant was again subjected to centrifugation at 39,000xg for 45 minutes. The resulting pellet, which contained the membrane envelope, was resuspended in 50mM ammonium bicarbonate and subjected to trypsin digestion as previously described. After digestion, samples were desalted using MACROspin C18 columns (The Nest Group, Southborough, MA). The flow-through was collected, concentrated in a speedvac to near dryness, and resuspended in 5% acetonitrile/0.1% formic acid for MS analysis. Samples were stored at -80C until MS analysis.

##### Data Analysis

The raw MS data files were processed with the MaxQuant [5] software (version 1.6.5.0). The peptide and protein identifications were performed using the Andromeda search engine [6] with a SwissProt/UniProt [7,8] FASTA database of all the reviewed and unreviewed protein sequences for *Francisella tularensi*s subspecies *novicida* strain U112 (April 2019). Trypsin digestion with a maximum of two missed cleavages, carbamidomethylation of cysteine as a fixed modification, and methionine oxidation and N-terminal acetylation as variable modifications were used as search parameters. A false discovery rate (FDR) of 0.01 at the peptide and protein level was applied. The 'match between the runs' option was enabled with the default time window of 0.7 min and the alignment time window size was 20 min. MaxQuant was allowed to automatically align the runs. Match type was 'match from and to'. 'Require MS/MS for comparisons' was on, and decoy mode was 'revert'. MaxQuant's label-free quantification (LFQ) algorithm [9] was used to calculate the relative protein intensity profiles across the samples. 'Advanced ratio estimation', 'stabilize large LFQ ratios' and 'advanced site intensities' were on.

Non-normalized protein intensities were extracted from the MaxQuant output and imported into the R statistical programming software envinronment version 3.6 [10]. Reverse protein hits, known contaminants and proteins with less than two peptides and at least one unique peptide were filtered out. The dataset was normalized using the variance stabilization normalization (vsn) [11] shown to perform well with proteomics data [12].

The dataset consisted of the null mutants of lpxD1 (D1), lpxD2 (D2) and lpxL (L) and the wild type (WT) measured in in five temperatures (18°C, 21°C, 25°C,, 32°C, 37°C). Each strain had three biological replicates measured in each temperature and each biological replicate was measured in three technical replicates. Each temperature consisted thus of 36 samples and the dataset had 180 samples in total.

For the evaluation of the reproducibility of the longitudinal differential expression methods, the dataset was divided into three technical replicate datasets. For the final longitudinal differential expression analysis and the associated pathway analysis, the technical replicates for each biological replicate in each temperature were averaged.

**Exploration of the dataset**

The proportion of missing values in the dataset was 10,8% and after averaging the technical replicates 6,9%. The majority of the missing values were located in the 37°C sample groups (**Supplementary Figure 2**).

**R packages**

R statistical programming software environment 3.6 [10] was used for the computational analysis performed in this study.

The R package pROC version 1.15.3 was used to conduct the ROC analysis [13]. The calculated pAUCs were visualized using the vioplot [14] package version 0.3.2. The gene set analysis was performed using the Bioconductor R package fgsea version 1.10.1 [15]. Version 3.1-142 of the R package nlme [16] was used for all the mixed effects modelling approaches in this study.

#### Supplementary Methods References

10. R Core Team. R: A Language and Environment for Statistical Computing. 2019;

11. Huber W, von Heydebreck A, Sültmann H, et al. Variance stabilization applied to microarray data calibration and to the quantification of differential expression. Bioinformatics 2002; 18 Suppl 1:S96–S104

12. Välikangas T, Suomi T, Elo LL. A systematic evaluation of normalization methods in quantitative label-free proteomics. Brief. Bioinform. 2018; 19:1–11

13. Robin X, Turck N, Hainard A, et al. pROC: an open-source package for R and S+ to analyze and compare ROC curves. BMC Bioinformatics 2011; 12:77

14. Adler D, Kelly ST. vioplot: violin plot. 2018;

15. Korotkevich G, Sukhov V, Budin N, et al. Fast gene set enrichment analysis. bioRxiv 2021; 60012

16. Pinheiro J, Bates D, DebRoy S, et al. {nlme}: Linear and Nonlinear Mixed Effects Models. 2019;

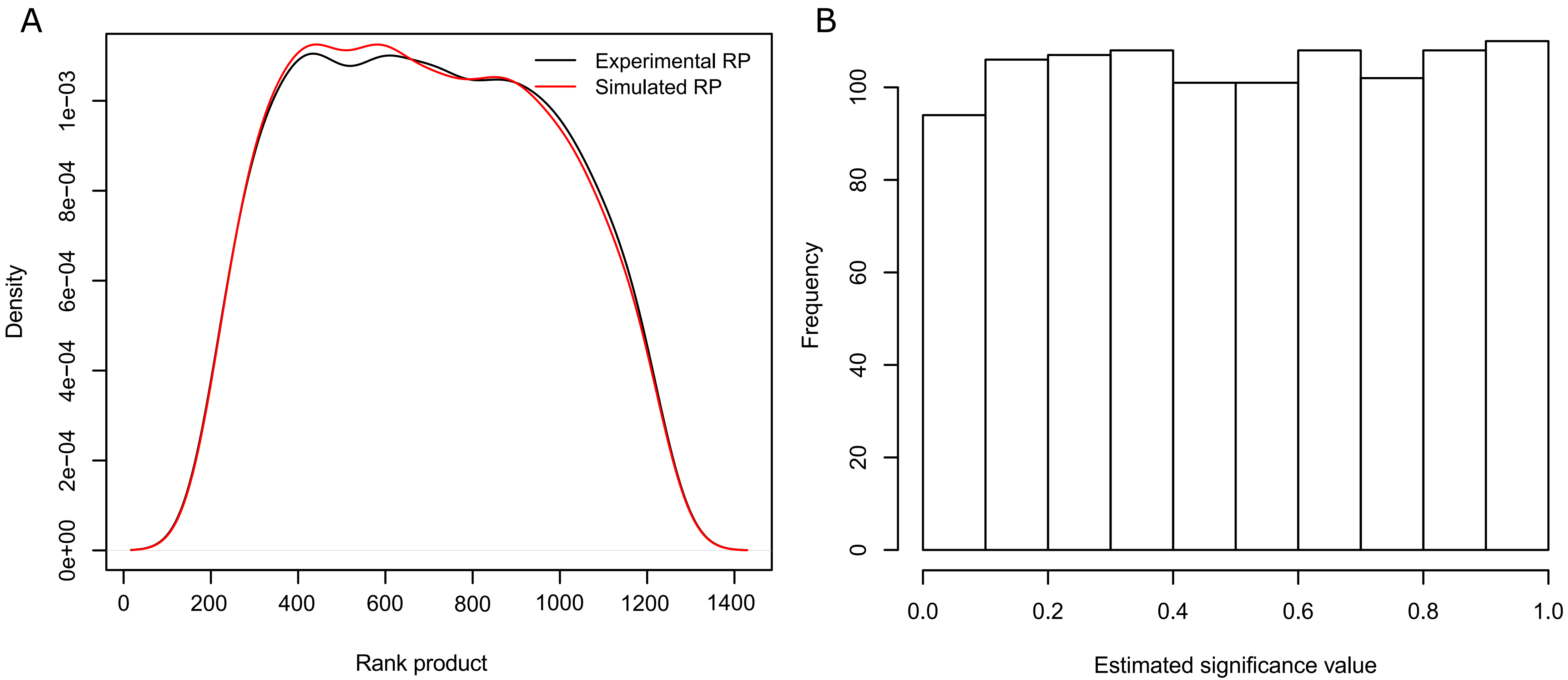


**Supplementary Figure 1.** Significance value estimation example in the Robust longitudinal Differential Expression method (RolDE). **(A)** Densities of experimental and simulated internal rank products for the RegROTS module in a simulated random longitudinal data. **(B)** Estimated significance values for RolDE in the random data of (A). Similar to the UPS1-based datasets, the random dataset contained two conditions, 5 timepoints and three replicates in each condition. The protein expression values for the random dataset were drawn from a normal distribution with a mean of 22 and a standard deviation of 1.5 with the rnorm() function in the stats-package in the R statistical programming language, resembling real $\log_2$ transformed experimental label free mass spectrometry (MS) intensity values. In figure (A) black lines annotate densities for the experimental rank products and red lines for the corresponding simulated rank products.

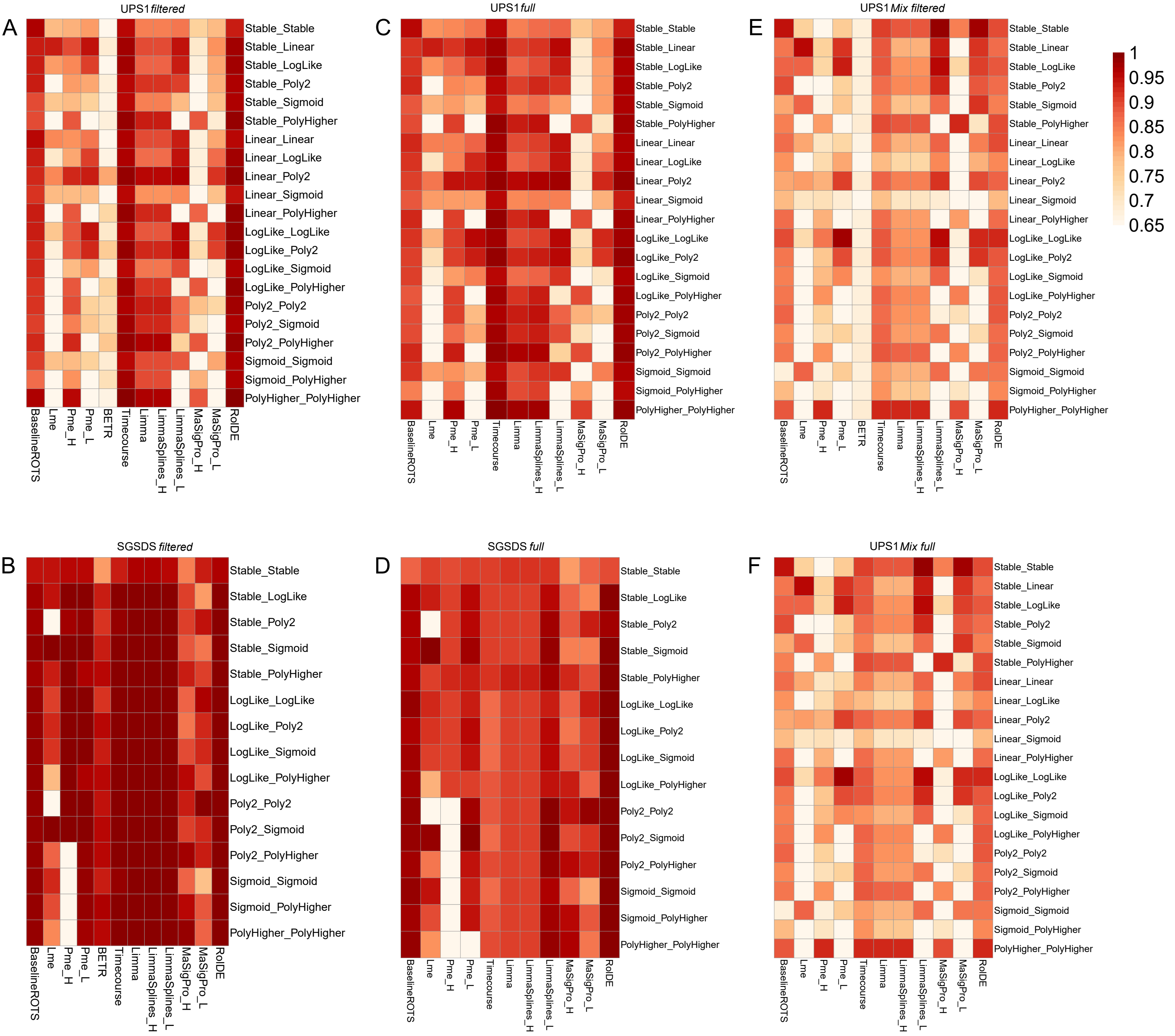


**Supplementary Figure 2.** The performance of the examined methods across the trend difference categories in the semi-simulated spike-in datasets. **(A)** UPS1 filtered, **(B)** SGSDS filtered, **(C)** UPS1 full, **(D)** SGSDS full, **(E)** UPS1 Mix filtered, **(F)** UPS1 Mix full. The methods were examined in their ability to detect true (known) longitudinal differential expression using receiver operating characteristic (ROC) analysis across the UPS1-based [29] (300 filtered, 300 full, 300 mix filtered and 300 mix full) and SGSDS-based [54] (210 filtered and 210 full) datasets with varying longitudinal trend differences in the spike-in proteins. The partial areas under the ROC curves (pAUC) between the specificity of 1 and 0.9 was used to measure the performance of the methods in the different categories. The interquartile range (IQR) means of the pAUCs for each method are presented.

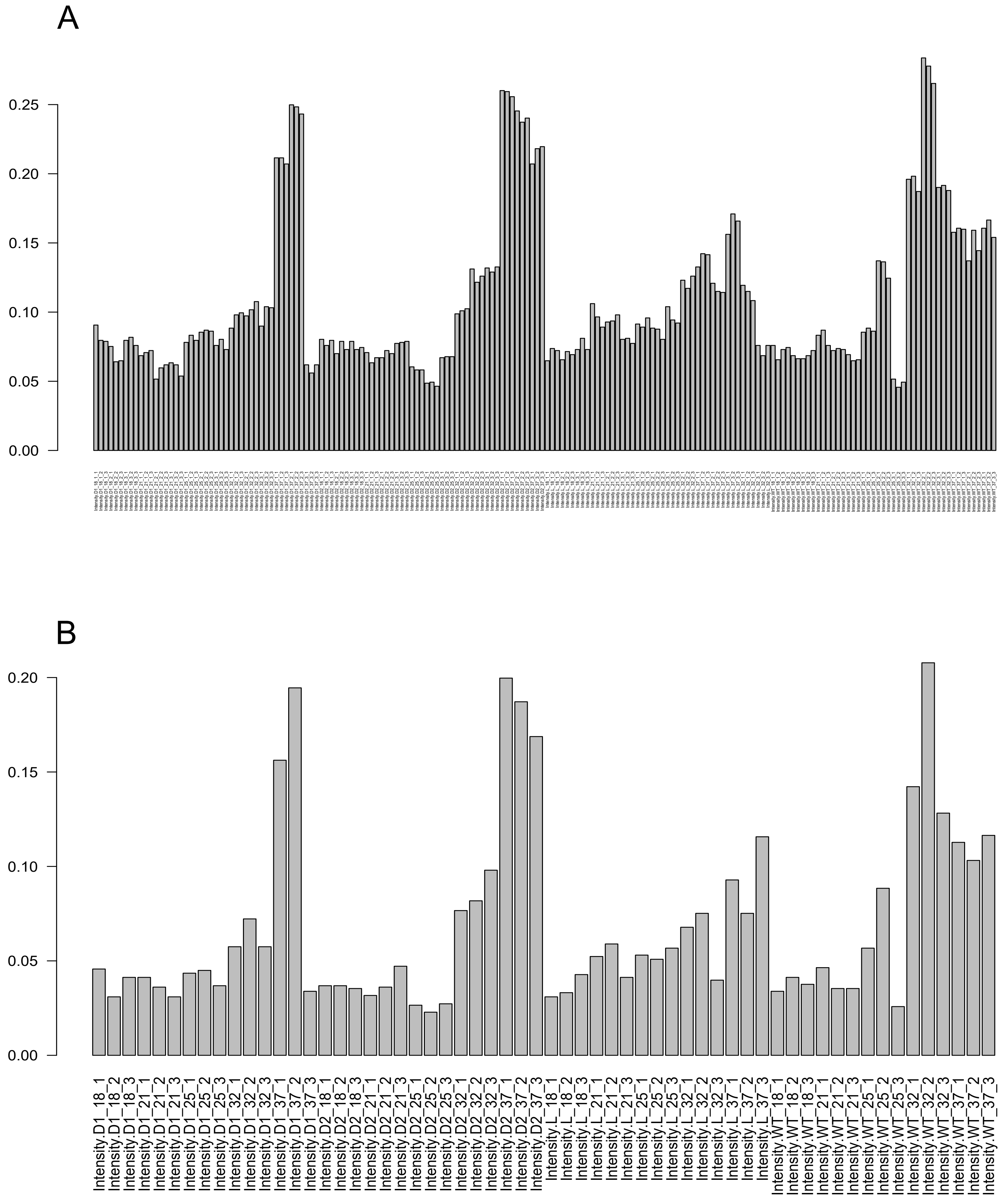


**Supplementary Figure 3.** Missing value proportions in the samples of the longitudinal *Francisella tularensis* subspecies *novicida* proteomics dataset. Missing value proportions in **(A)** all the samples, and **(B)** after averaging over the technical replicates for a biological replicate.

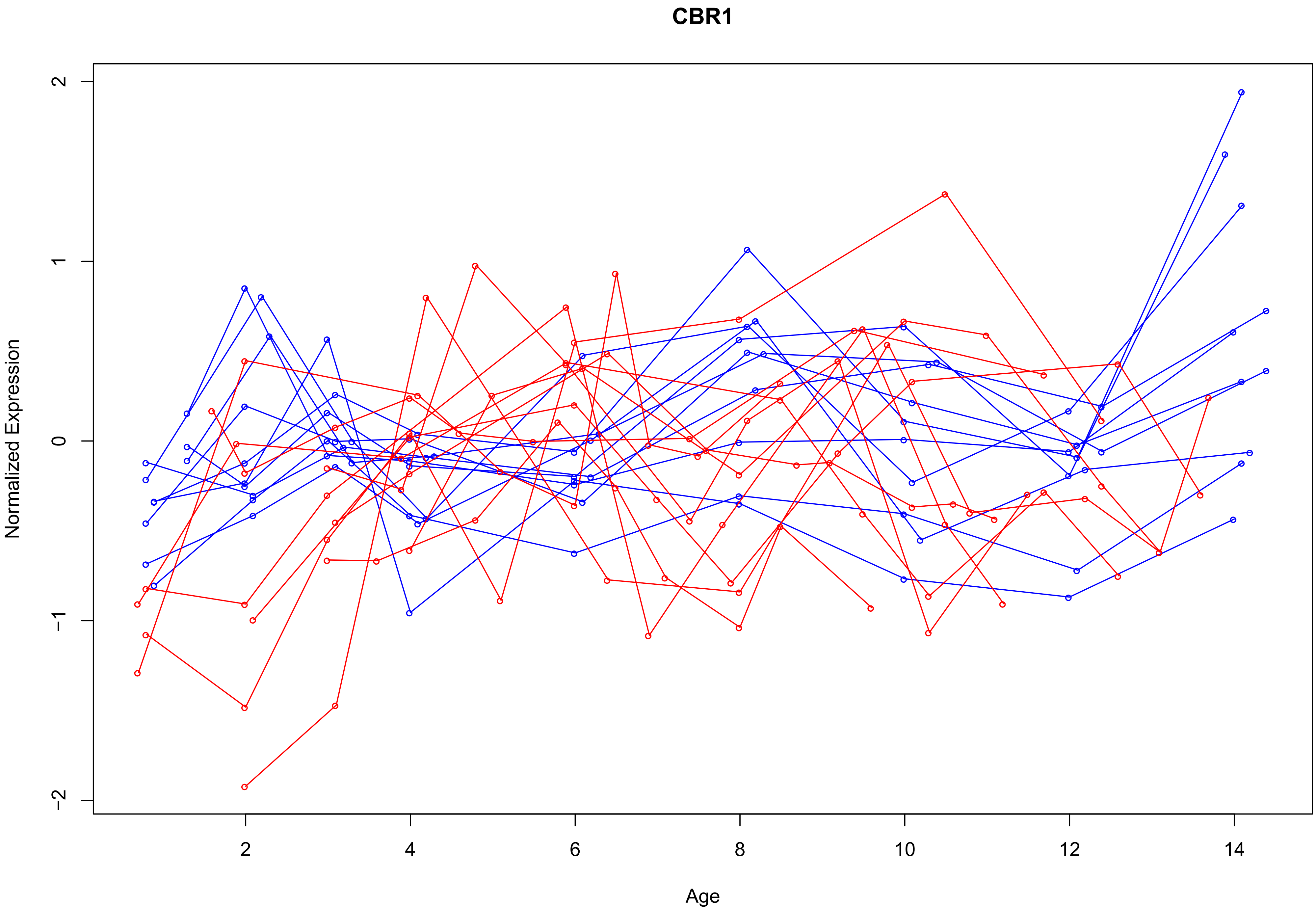


**Supplementary Figure 4.** A significant finding (FDR 0.04) by Liu et al. [10] [Supplementary Table 3] in their differential pattern analysis of the longitudinal blood plasma proteome of 11 children developing T1D and 10 matched controls. CBR1 RolDE FDR 0.12 for longitudinal differential expression.

**Supplementary Table 1.** The detected longitudinally differentially expressed proteins at FDR 0.05 by the new proposed differential expression method, Robust longitudinal Differential Expression (RolDE), in the longitudinal type 1 diabetes blood plasma proteomics data of Liu et al [10].

| Feature ID | RolDE Rank Product | Estimated false discovery rate |
|---|---|---|
| CGREF1 | 5.3 | $<10^{-16}$ |
| KRT31 | 7.7 | $<10^{-16}$ |
| TF | 7.7 | $<10^{-16}$ |
| SAA1 | 7.7 | $<10^{-16}$ |
| KRT86 | 16.6 | $<10^{-16}$ |
| HLA-A | 20.2 | $<10^{-16}$ |
| SCLT1 | 21.8 | $<10^{-16}$ |
| TSKU | 25.0 | $<10^{-16}$ |
| LY6G6F | 29.1 | $<10^{-16}$ |
| JAG2 | 31.4 | $<10^{-16}$ |
| HP;HPR | 33.4 | $<10^{-16}$ |
| HLA-B | 36.2 | $<10^{-16}$ |
| FTL | 40.7 | $<10^{-16}$ |
| TRY8;PRSS2 | 41.9 | $<10^{-16}$ |
| CCDC126 | 42.7 | $<10^{-16}$ |
| A0A0G2JH38 | 43.3 | $<10^{-16}$ |
| PNLIP | 45.7 | $<10^{-16}$ |
| RPS14 | 46.5 | $<10^{-16}$ |
| PEF1 | 47.7 | $<10^{-16}$ |
| RAB2A | 48.5 | $<10^{-16}$ |
| HBG2 | 48.7 | $<10^{-16}$ |
| CS | 50.6 | $<10^{-16}$ |
| ELTD1 | 52.5 | 0.043 |

**Supplementary Table 2.** All the unique proteins in the "KEGG Lipopolysaccharide synthesis pathway (ftn00540)" [69] complemented with unique proteins from the associated "Lipopolysaccharide biosynthesis knockout pathway (ko00540)". Proteins belonging to the "KEGG Lipopolysaccharide synthesis pathway (ftn00540)" are highlighted.

| Entry | Entry_name | Status | Gene_names | Gene_names_primary |
|---|---|---|---|---|
| A0Q7Y0 | A0Q7Y0_FRATN | unreviewed | lpxA FTN_1478 | lpxA |
| A0Q4B0 | A0Q4B0_FRATN | unreviewed | lpxC FTN_0165 | lpxC |
| A0Q7Y2 | A0Q7Y2_FRATN | unreviewed | lpxD FTN_1480 | lpxD |
| A0Q4E5 | A0Q4E5_FRATN | unreviewed | lpxD FTN_0200 | lpxD |
| A0Q5A8 | A0Q5A8_FRATN | unreviewed | lpxH FTN_0528 | lpxH |
| A0Q7X9 | LPXB_FRATN | reviewed | lpxB FTN_1477 | lpxB |
| A0Q8A0 | LPXK_FRATN | reviewed | lpxK FTN_1605 | lpxK |
| A0Q788 | A0Q788_FRATN | unreviewed | kpsF FTN_1222 | kpsF |
| A0Q5J1 | KDSA_FRATN | reviewed | kdsA FTN_0611 | kdsA |
| A0Q6C8 | A0Q6C8_FRATN | unreviewed | yrbI FTN_0905 | yrbI |
| A0Q5R0 | KDSB_FRATN | reviewed | kdsB FTN_0683 | kdsB |
| A0Q7X2 | A0Q7X2_FRATN | unreviewed | kdtA FTN_1469 | kdtA |
| A0Q418 | A0Q418_FRATN | unreviewed | FTN_0072 | #N/A |
| A0Q417 | A0Q417_FRATN | unreviewed | FTN_0071 | #N/A |
| A0Q450 | A0Q450_FRATN | unreviewed | FTN_0104 | #N/A |
| A0Q504 | A0Q504_FRATN | unreviewed | lpxE FTN_0416 | lpxE |
| A0Q576 | A0Q576_FRATN | unreviewed | kdoH1 FTN_0495 | kdoH1 |
| A0Q4N6 | LPXF_FRATN | reviewed | lpxF FTN_0295 AW25_1746 | lpxF |

**Supplementary File 1.** Including:

**Supplementary_File_SemiSimulated_Spike_In_Data_Trends)** All the generated trends and combinations of trends for the spike-in proteins in the semi-simulated datasets.

**Supplementary_Table_SemiSimulated_Spike_In_Data_SampleGroups)** All the generated trends, their categories and all the generated combinations of trends for spike-in proteins for the semi-simulated datasets based on the UPS1, SGSDS and the CPTAC datasets.

**Supplementary_Table_SemiSimulated_Mix_Datasets_Trends_SampleGroups)** All the generated trends for each spike-in protein for each semi-simulated UPS1 Mix dataset.
