## Supplementary File 1 for "Enhanced longitudinal differential expression detection in proteomics with robust reproducibility optimization regression": Supplementary_File_SemiSimulated_Spike_In_Data_Trends.pdf

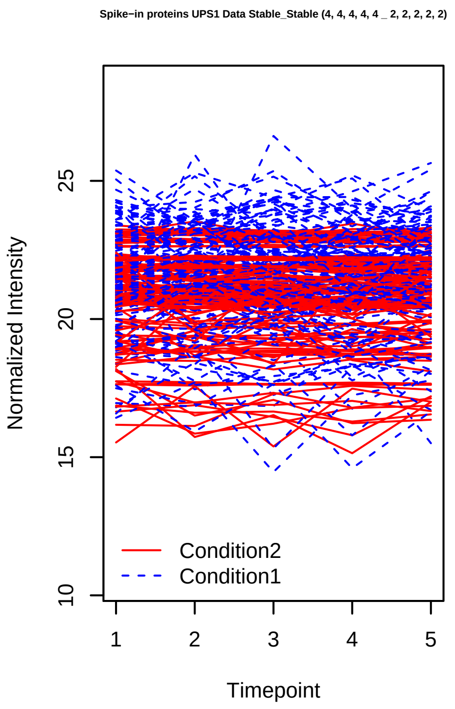
Spike−in proteins UPS1 Data Stable_Stable (4, 4, 4, 4, 4 _ 2, 2, 2, 2, 2)
Normalized Intensity
Timepoint
Condition2
Condition1

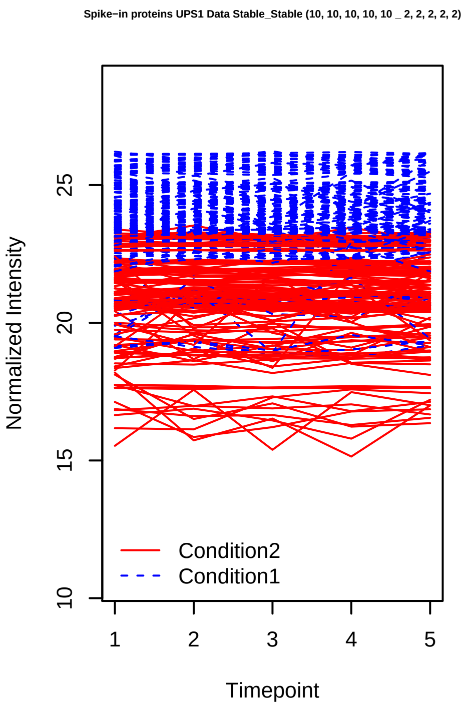
Spike−in proteins UPS1 Data Stable_Stable (10, 10, 10, 10, 10 _ 2, 2, 2, 2, 2)
Normalized Intensity
Timepoint
Condition2
Condition1

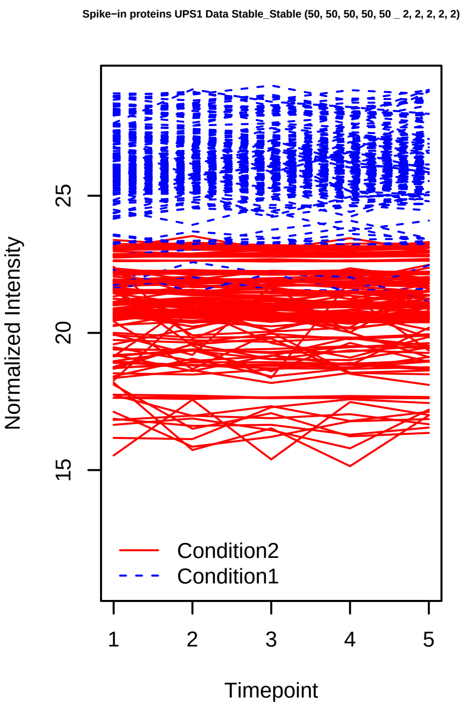
Spike−in proteins UPS1 Data Stable_Stable (50, 50, 50, 50, 50 _ 2, 2, 2, 2, 2)
Normalized Intensity
Timepoint
Condition2
Condition1

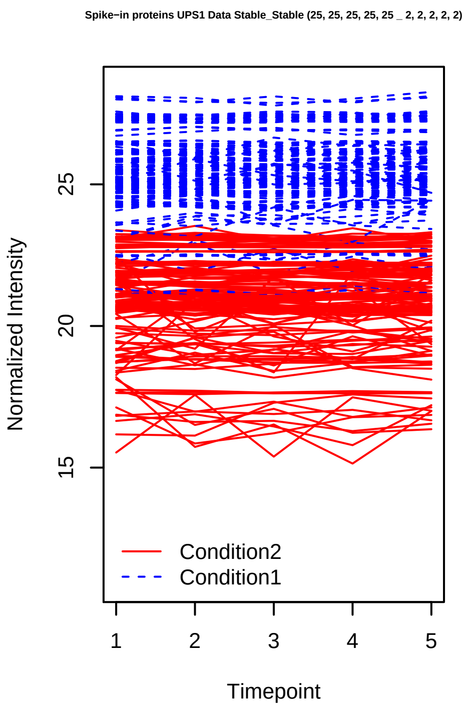
Spike−in proteins UPS1 Data Stable_Stable (25, 25, 25, 25, 25 _ 2, 2, 2, 2, 2)
Normalized Intensity
Timepoint
Condition2
Condition1

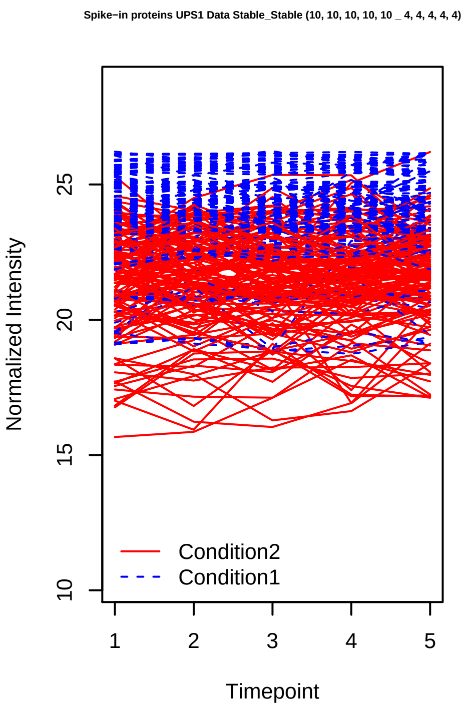
Spike−in proteins UPS1 Data Stable_Stable (10, 10, 10, 10, 10 _ 4, 4, 4, 4, 4)
Normalized Intensity
Timepoint
Condition2
Condition1

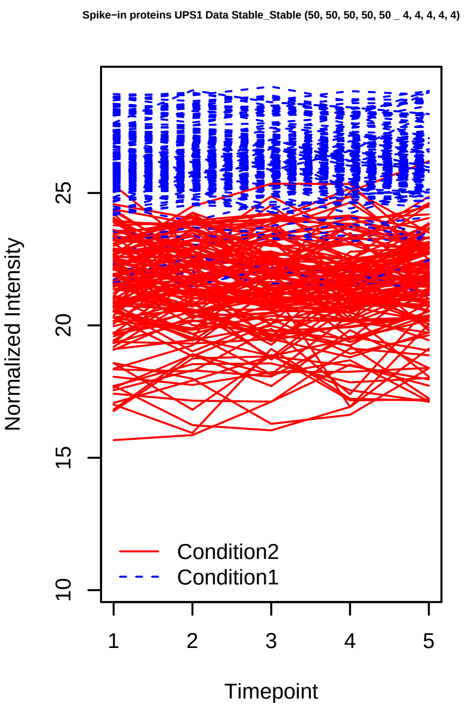
Spike−in proteins UPS1 Data Stable_Stable (50, 50, 50, 50, 50 _ 4, 4, 4, 4, 4)
Normalized Intensity
Timepoint
Condition2
Condition1

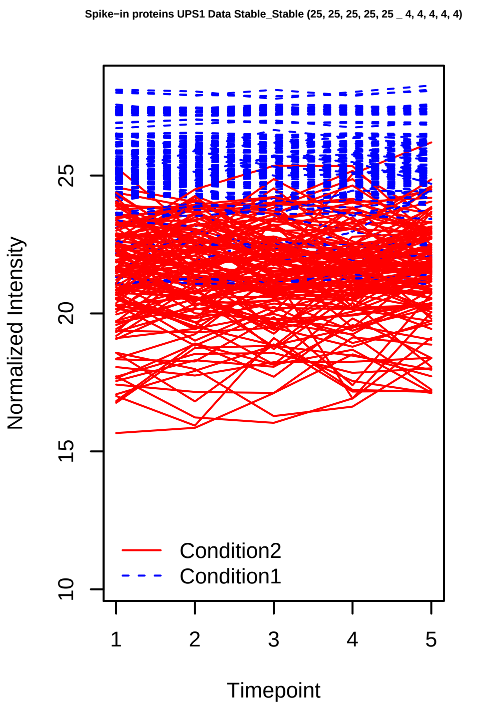
Spike−in proteins UPS1 Data Stable_Stable (25, 25, 25, 25, 25 _ 4, 4, 4, 4, 4)
Normalized Intensity
10
15
20
25
Timepoint
1
2
3
4
5
Condition2
Condition1

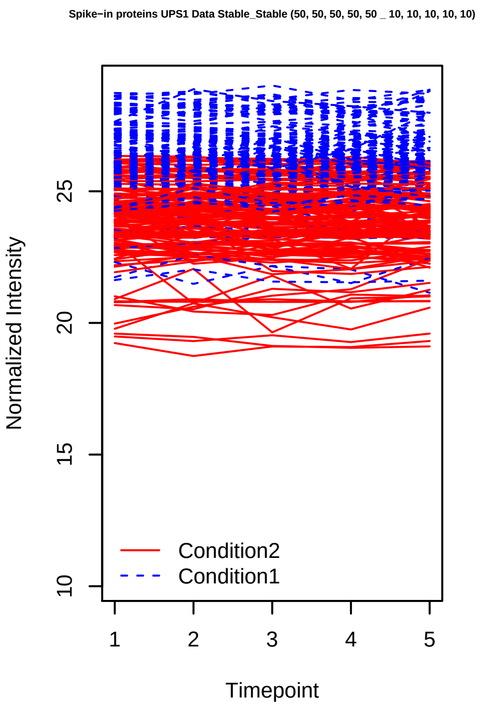
Spike−in proteins UPS1 Data Stable_Stable (50, 50, 50, 50, 50 _ 10, 10, 10, 10, 10)
Normalized Intensity
10
15
20
25
Timepoint
1
2
3
4
5
Condition2
Condition1

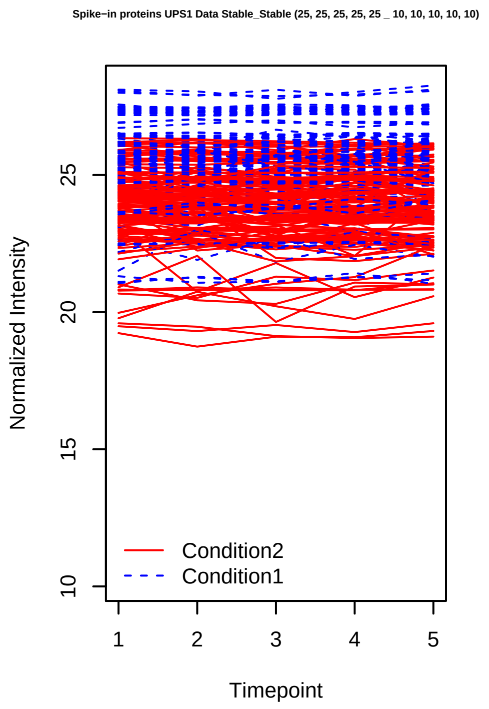
Spike−in proteins UPS1 Data Stable_Stable (25, 25, 25, 25, 25 _ 10, 10, 10, 10, 10)
Normalized Intensity
10
15
20
25
Timepoint
1
2
3
4
5
Condition2
Condition1

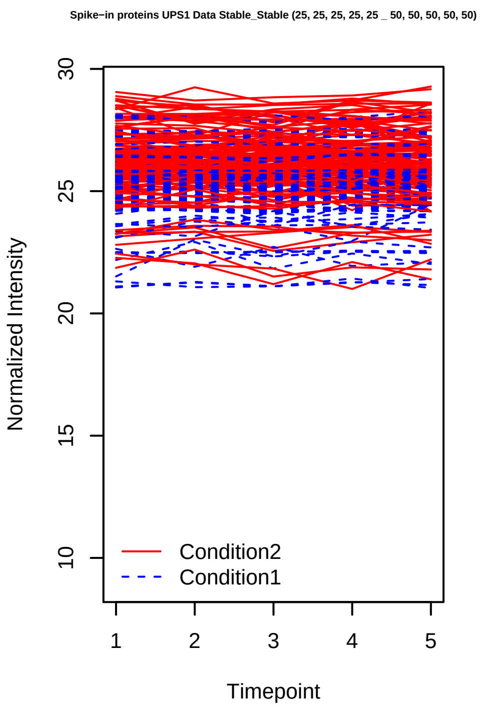
Spike−in proteins UPS1 Data Stable_Stable (25, 25, 25, 25, 25 _ 50, 50, 50, 50, 50)
Normalized Intensity
10
15
20
25
30
Timepoint
1
2
3
4
5
Condition2
Condition1

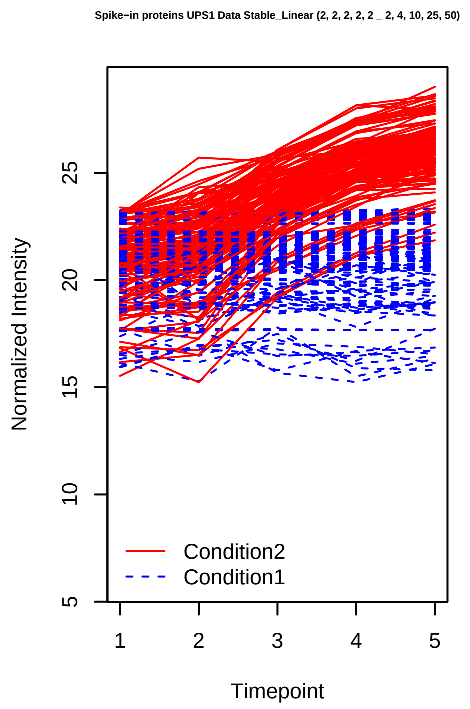
Spike−in proteins UPS1 Data Stable_Linear (2, 2, 2, 2, 2 _ 2, 4, 10, 25, 50)
Normalized Intensity
5
10
15
20
25
Timepoint
1
2
3
4
5
Condition2
Condition1

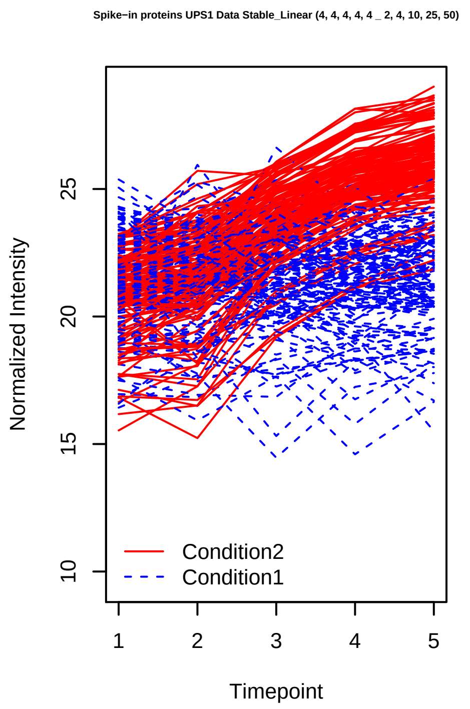
Spike−in proteins UPS1 Data Stable_Linear (4, 4, 4, 4, 4 _ 2, 4, 10, 25, 50)
Normalized Intensity
10
15
20
25
Timepoint
1
2
3
4
5
Condition2
Condition1

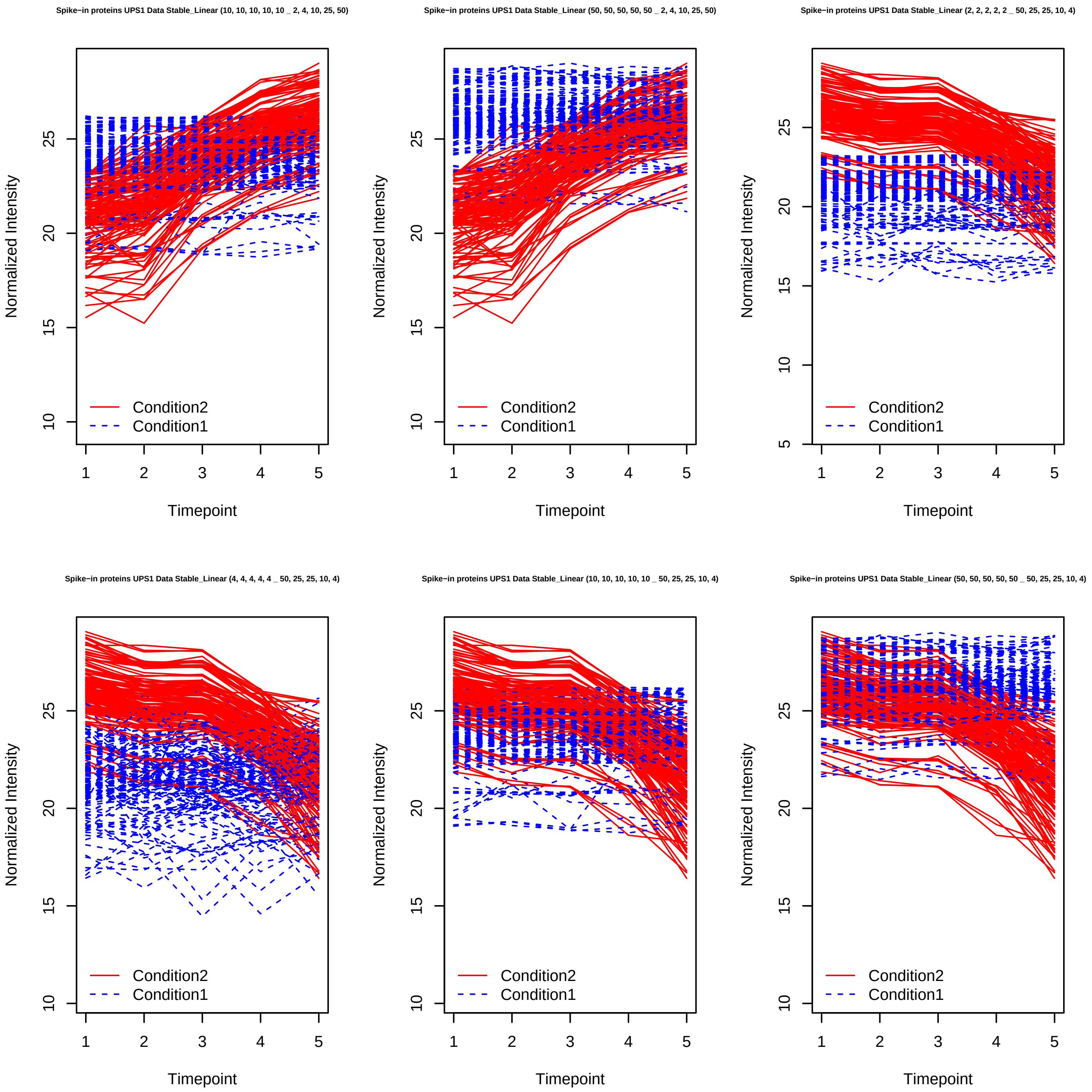

Spike−in proteins UPS1 Data Stable_Linear (10, 10, 10, 10, 10 _ 2, 4, 10, 25, 50)
Spike−in proteins UPS1 Data Stable_Linear (50, 50, 50, 50, 50 _ 2, 4, 10, 25, 50)
Spike−in proteins UPS1 Data Stable_Linear (2, 2, 2, 2, 2 _ 50, 25, 25, 10, 4)
Spike−in proteins UPS1 Data Stable_Linear (4, 4, 4, 4, 4 _ 50, 25, 25, 10, 4)
Spike−in proteins UPS1 Data Stable_Linear (10, 10, 10, 10, 10 _ 50, 25, 25, 10, 4)
Spike−in proteins UPS1 Data Stable_Linear (50, 50, 50, 50, 50 _ 50, 25, 25, 10, 4)
Normalized Intensity
Timepoint
Condition2
Condition1

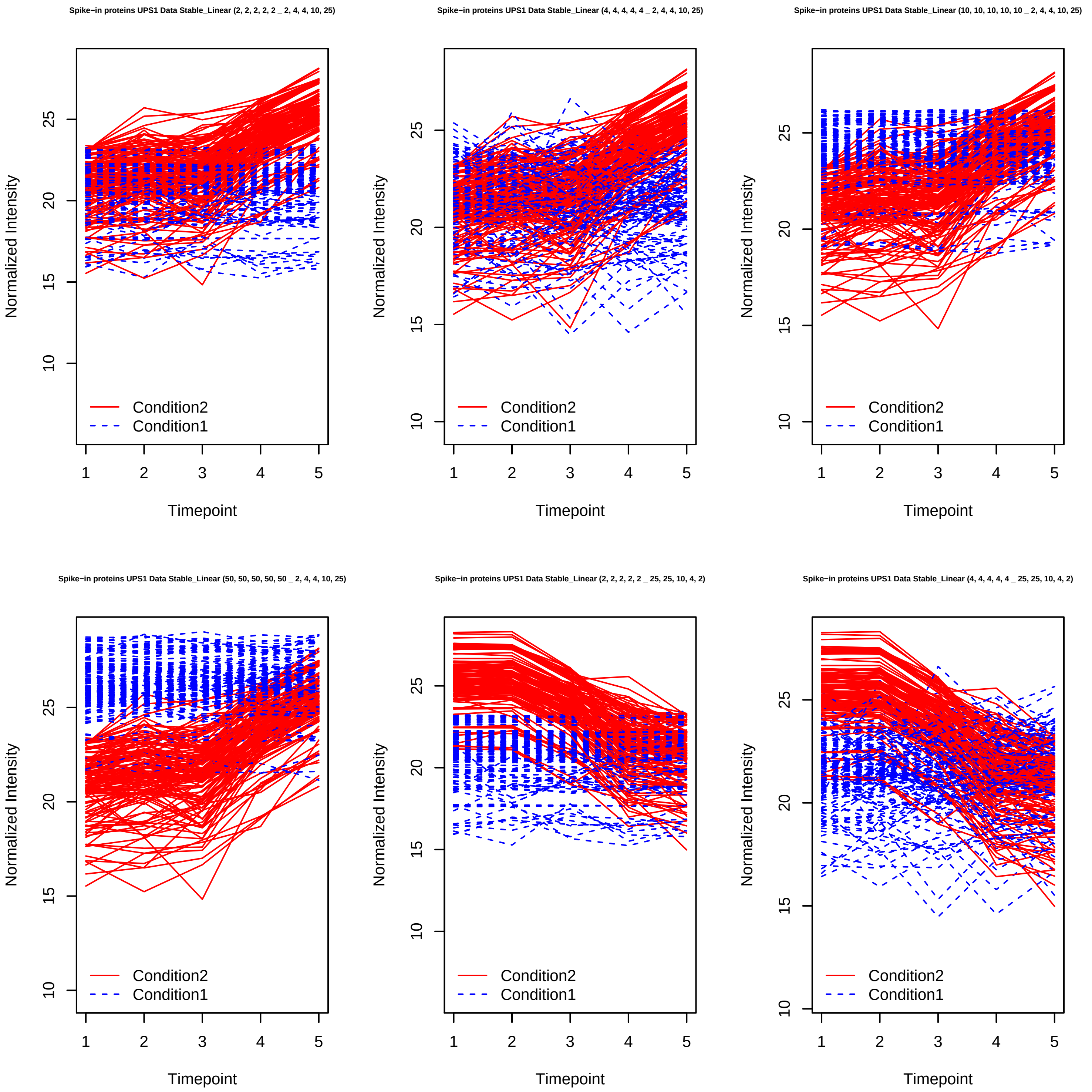
Spike−in proteins UPS1 Data Stable_Linear (2, 2, 2, 2, 2 _ 2, 4, 4, 10, 25)
Spike−in proteins UPS1 Data Stable_Linear (4, 4, 4, 4, 4 _ 2, 4, 4, 10, 25)
Spike−in proteins UPS1 Data Stable_Linear (10, 10, 10, 10, 10 _ 2, 4, 4, 10, 25)
Spike−in proteins UPS1 Data Stable_Linear (50, 50, 50, 50, 50 _ 2, 4, 4, 10, 25)
Spike−in proteins UPS1 Data Stable_Linear (2, 2, 2, 2, 2 _ 25, 25, 10, 4, 2)
Spike−in proteins UPS1 Data Stable_Linear (4, 4, 4, 4, 4 _ 25, 25, 10, 4, 2)
Normalized Intensity
Timepoint
Condition2
Condition1
10
15
20
25
1
2
3
4
5

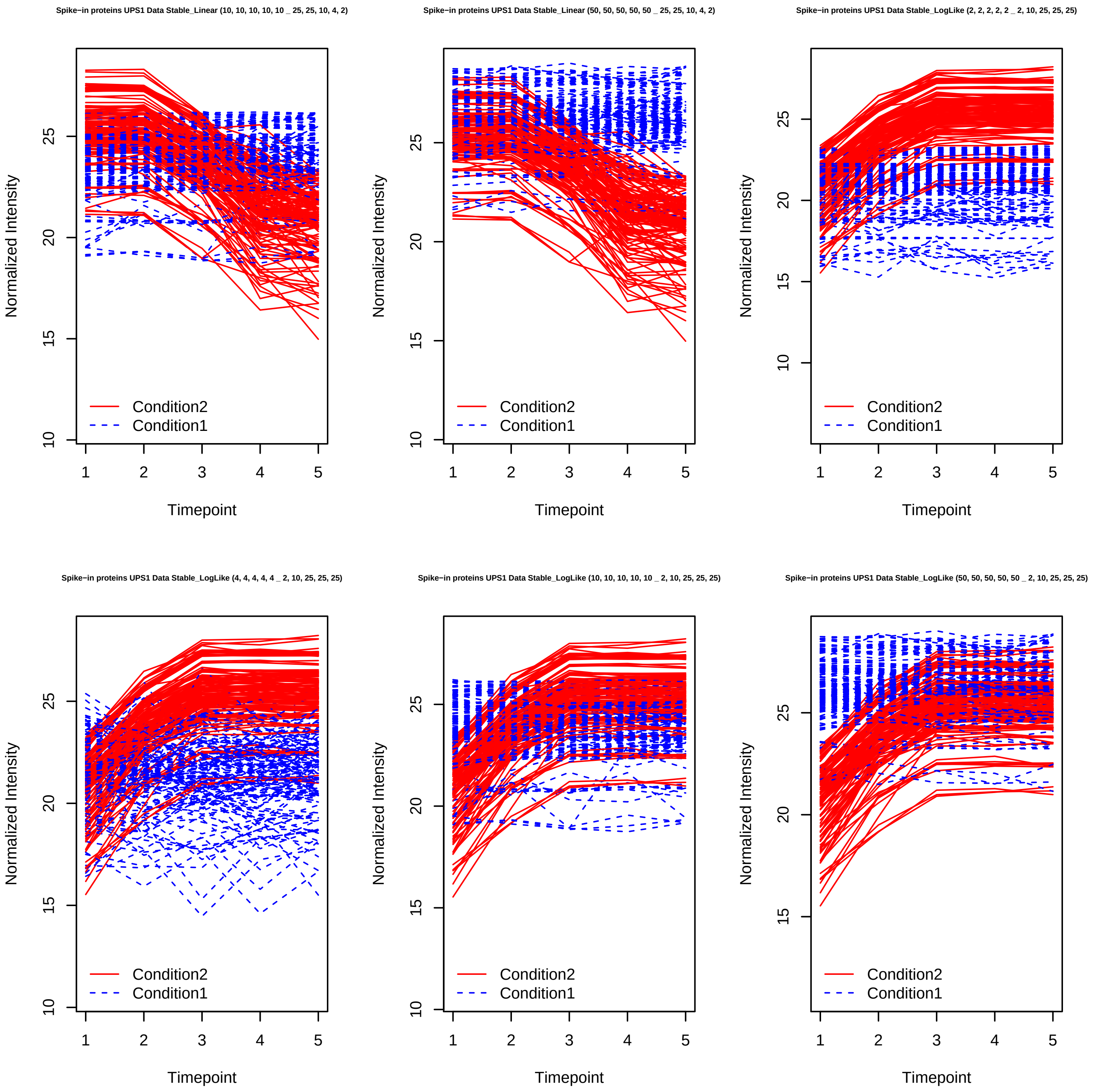
Spike−in proteins UPS1 Data Stable_Linear (10, 10, 10, 10, 10 _ 25, 25, 10, 4, 2)
Spike−in proteins UPS1 Data Stable_Linear (50, 50, 50, 50, 50 _ 25, 25, 10, 4, 2)
Spike−in proteins UPS1 Data Stable_LogLike (2, 2, 2, 2, 2 _ 2, 10, 25, 25, 25)
Spike−in proteins UPS1 Data Stable_LogLike (4, 4, 4, 4, 4 _ 2, 10, 25, 25, 25)
Spike−in proteins UPS1 Data Stable_LogLike (10, 10, 10, 10, 10 _ 2, 10, 25, 25, 25)
Spike−in proteins UPS1 Data Stable_LogLike (50, 50, 50, 50, 50 _ 2, 10, 25, 25, 25)
Normalized Intensity
Timepoint
Condition2
Condition1

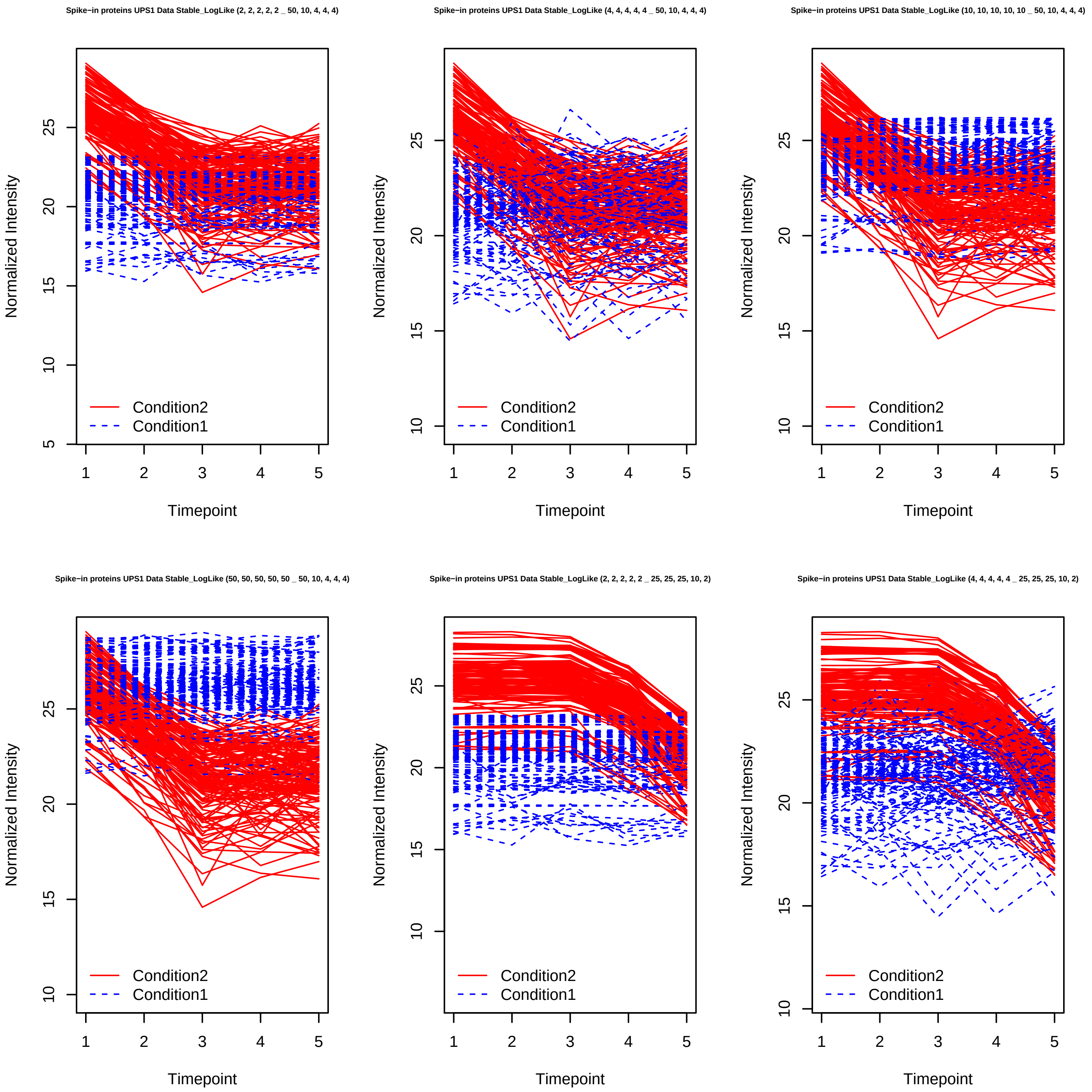
Spike−in proteins UPS1 Data Stable_LogLike (2, 2, 2, 2, 2 _ 50, 10, 4, 4, 4)
Spike−in proteins UPS1 Data Stable_LogLike (4, 4, 4, 4, 4 _ 50, 10, 4, 4, 4)
Spike−in proteins UPS1 Data Stable_LogLike (10, 10, 10, 10, 10 _ 50, 10, 4, 4, 4)
Spike−in proteins UPS1 Data Stable_LogLike (50, 50, 50, 50, 50 _ 50, 10, 4, 4, 4)
Spike−in proteins UPS1 Data Stable_LogLike (2, 2, 2, 2, 2 _ 25, 25, 25, 10, 2)
Spike−in proteins UPS1 Data Stable_LogLike (4, 4, 4, 4, 4 _ 25, 25, 25, 10, 2)
Normalized Intensity
Timepoint
Condition2
Condition1

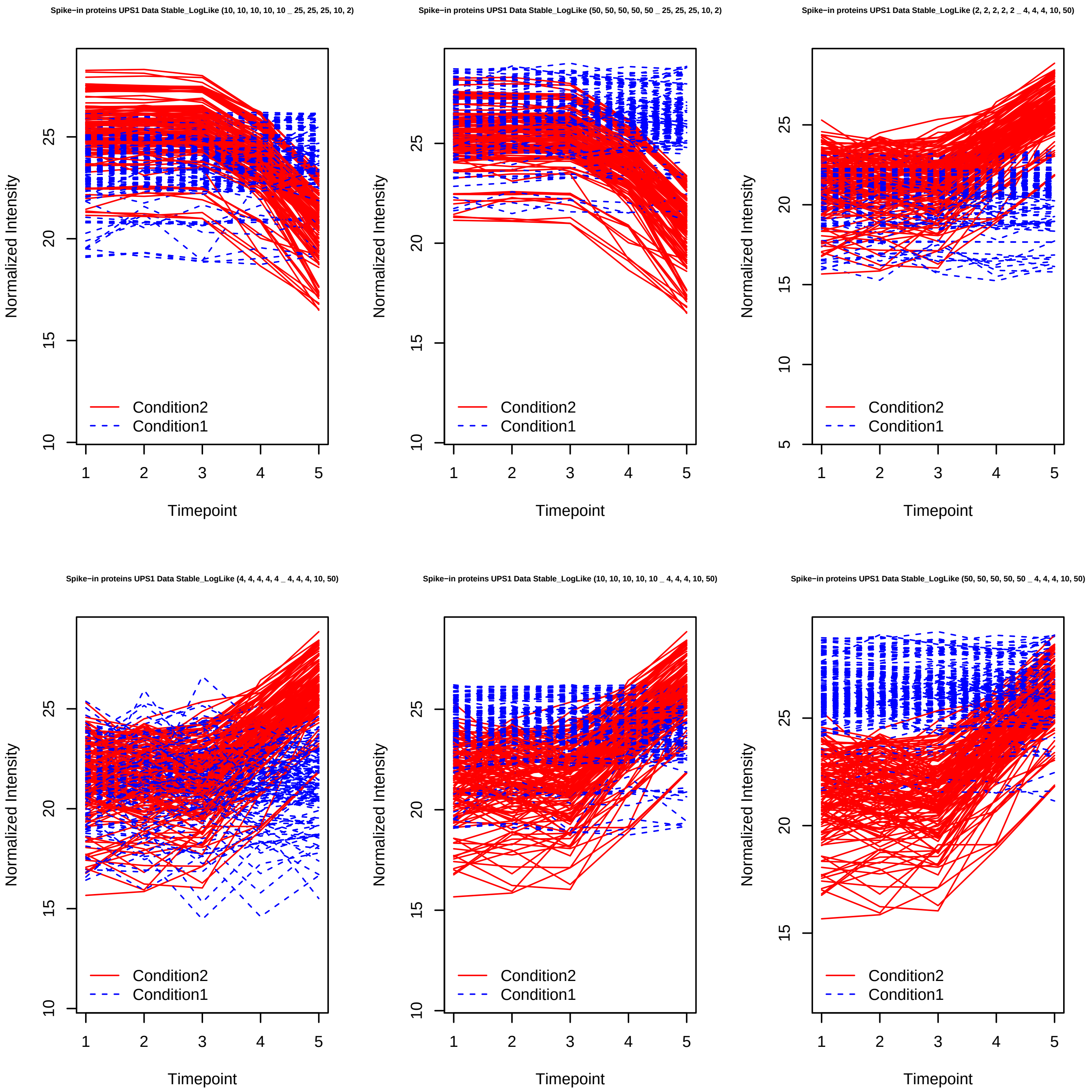
Spike−in proteins UPS1 Data Stable_LogLike (10, 10, 10, 10, 10 _ 25, 25, 25, 10, 2)
Spike−in proteins UPS1 Data Stable_LogLike (50, 50, 50, 50, 50 _ 25, 25, 25, 10, 2)
Spike−in proteins UPS1 Data Stable_LogLike (2, 2, 2, 2, 2 _ 4, 4, 4, 10, 50)
Spike−in proteins UPS1 Data Stable_LogLike (4, 4, 4, 4, 4 _ 4, 4, 4, 10, 50)
Spike−in proteins UPS1 Data Stable_LogLike (10, 10, 10, 10, 10 _ 4, 4, 4, 10, 50)
Spike−in proteins UPS1 Data Stable_LogLike (50, 50, 50, 50, 50 _ 4, 4, 4, 10, 50)
Normalized Intensity
Timepoint
Condition2
Condition1

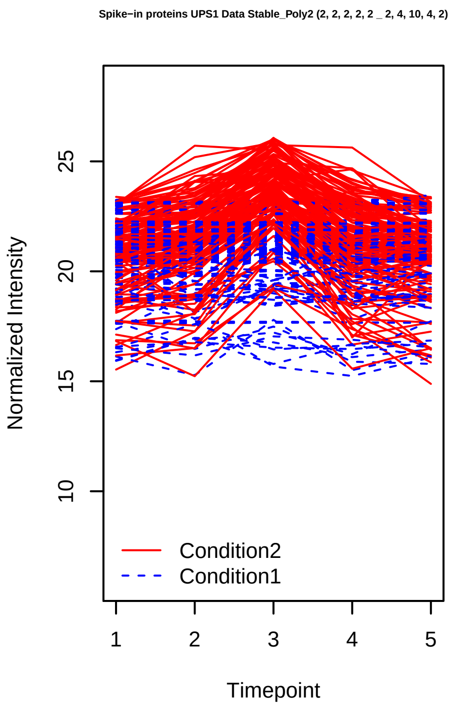
Spike−in proteins UPS1 Data Stable_Poly2 (2, 2, 2, 2, 2 _ 2, 4, 10, 4, 2)
Normalized Intensity
Timepoint
Condition2
Condition1

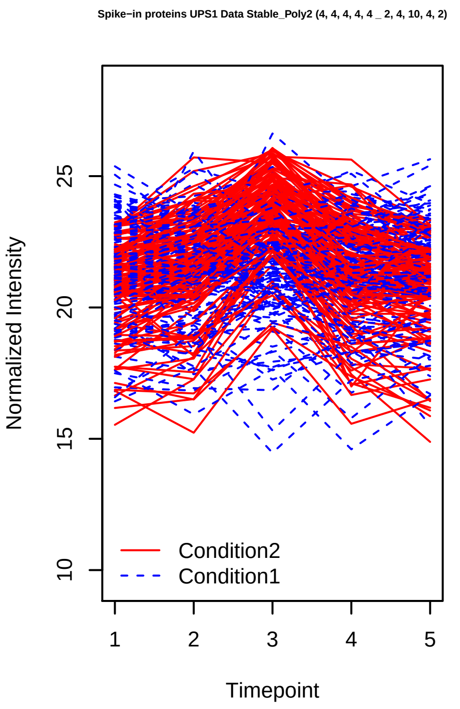
Spike−in proteins UPS1 Data Stable_Poly2 (4, 4, 4, 4, 4 _ 2, 4, 10, 4, 2)
Normalized Intensity
Timepoint
Condition2
Condition1

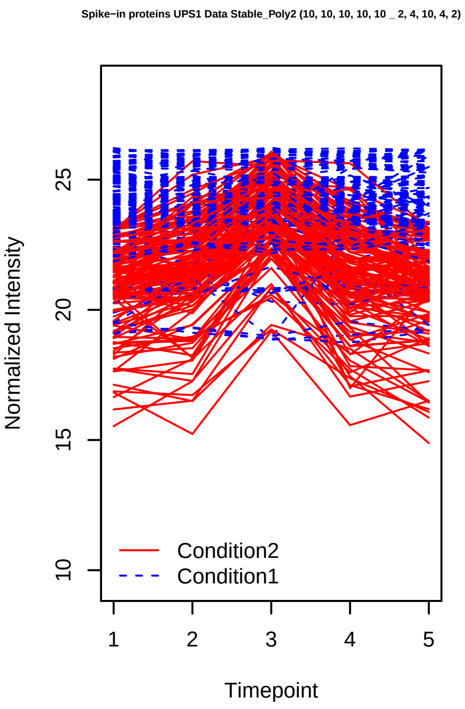
Spike−in proteins UPS1 Data Stable_Poly2 (10, 10, 10, 10, 10 _ 2, 4, 10, 4, 2)
Normalized Intensity
Timepoint
Condition2
Condition1

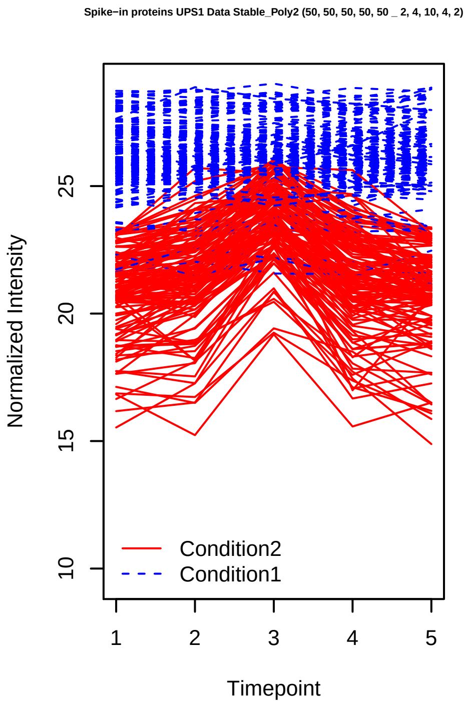
Spike−in proteins UPS1 Data Stable_Poly2 (50, 50, 50, 50, 50 _ 2, 4, 10, 4, 2)
Normalized Intensity
Timepoint
Condition2
Condition1

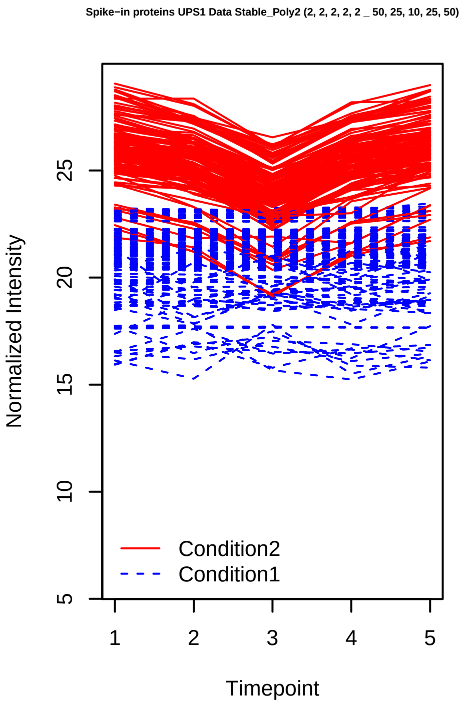
Spike−in proteins UPS1 Data Stable_Poly2 (2, 2, 2, 2, 2 _ 50, 25, 10, 25, 50)
Normalized Intensity
Timepoint
Condition2
Condition1

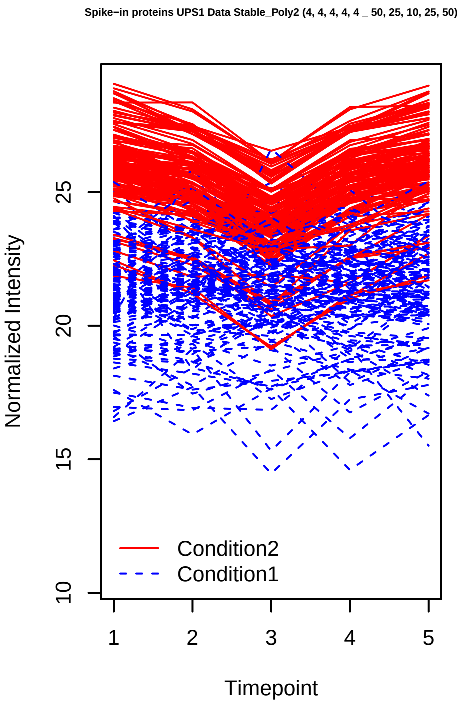
Spike−in proteins UPS1 Data Stable_Poly2 (4, 4, 4, 4, 4 _ 50, 25, 10, 25, 50)
Normalized Intensity
Timepoint
Condition2
Condition1

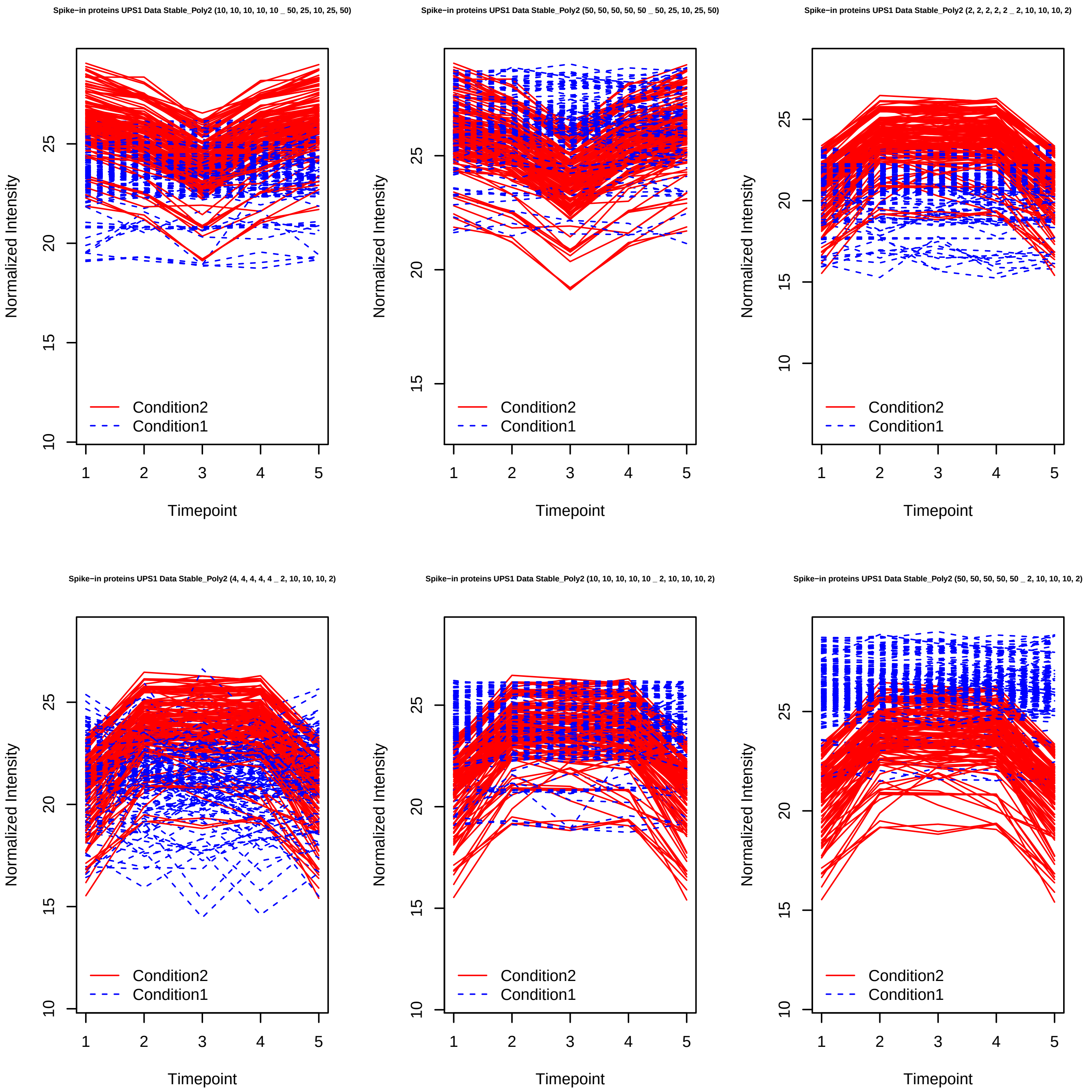

Spike−in proteins UPS1 Data Stable_Poly2 (10, 10, 10, 10, 10 _ 50, 25, 10, 25, 50)
Spike−in proteins UPS1 Data Stable_Poly2 (50, 50, 50, 50, 50 _ 50, 25, 10, 25, 50)
Spike−in proteins UPS1 Data Stable_Poly2 (2, 2, 2, 2, 2 _ 2, 10, 10, 10, 2)
Spike−in proteins UPS1 Data Stable_Poly2 (4, 4, 4, 4, 4 _ 2, 10, 10, 10, 2)
Spike−in proteins UPS1 Data Stable_Poly2 (10, 10, 10, 10, 10 _ 2, 10, 10, 10, 2)
Spike−in proteins UPS1 Data Stable_Poly2 (50, 50, 50, 50, 50 _ 2, 10, 10, 10, 2)
Normalized Intensity
Timepoint
Condition2
Condition1

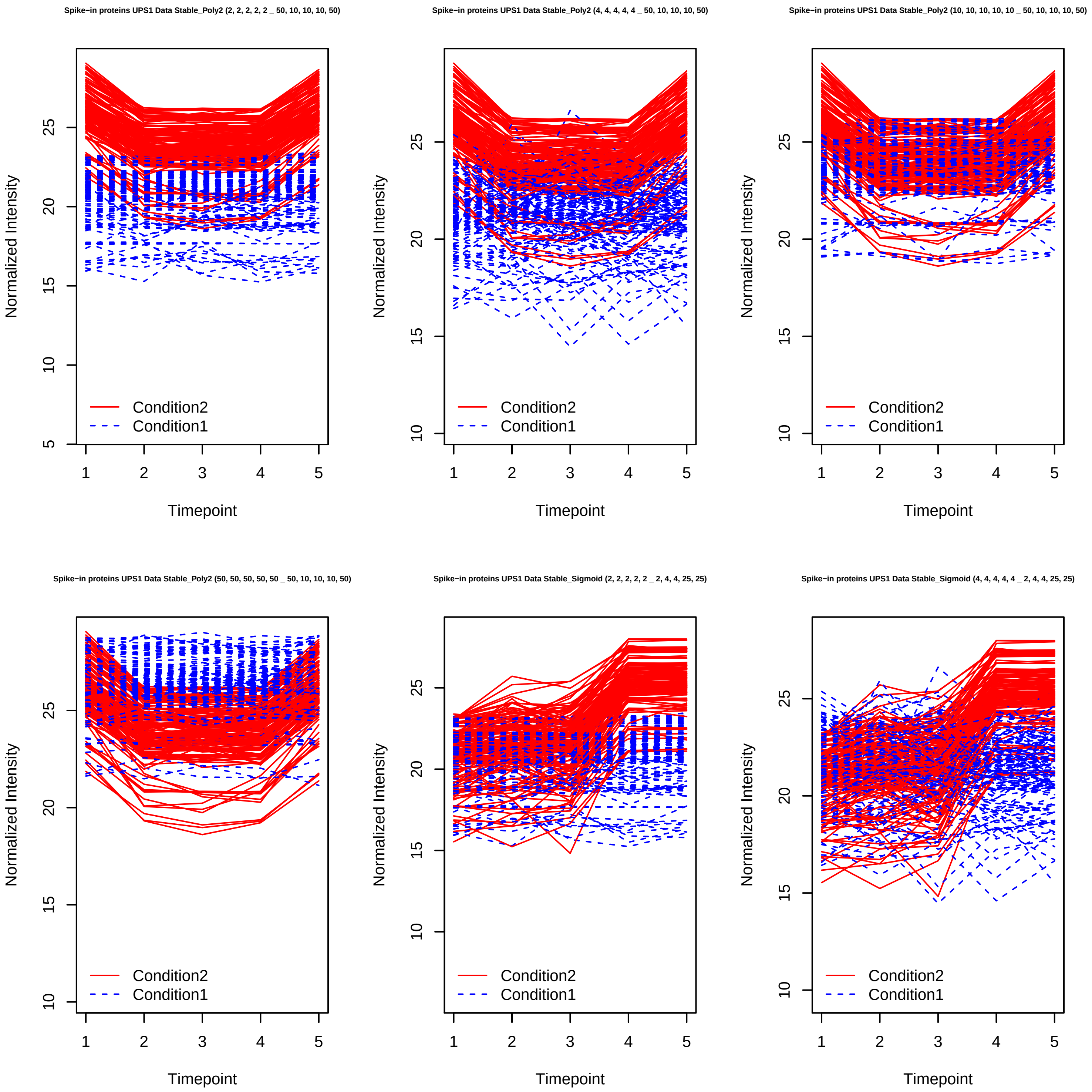
Spike−in proteins UPS1 Data Stable_Poly2 (2, 2, 2, 2, 2 _ 50, 10, 10, 10, 50)
Normalized Intensity
5
10
15
20
25
Condition2
Condition1
1
2
3
4
5
Timepoint
Spike−in proteins UPS1 Data Stable_Poly2 (4, 4, 4, 4, 4 _ 50, 10, 10, 10, 50)
Normalized Intensity
10
15
20
25
Condition2
Condition1
1
2
3
4
5
Timepoint
Spike−in proteins UPS1 Data Stable_Poly2 (10, 10, 10, 10, 10 _ 50, 10, 10, 10, 50)
Normalized Intensity
10
15
20
25
Condition2
Condition1
1
2
3
4
5
Timepoint
Spike−in proteins UPS1 Data Stable_Poly2 (50, 50, 50, 50, 50 _ 50, 10, 10, 10, 50)
Normalized Intensity
10
15
20
25
Condition2
Condition1
1
2
3
4
5
Timepoint
Spike−in proteins UPS1 Data Stable_Sigmoid (2, 2, 2, 2, 2 _ 2, 4, 4, 25, 25)
Normalized Intensity
10
15
20
25
Condition2
Condition1
1
2
3
4
5
Timepoint
Spike−in proteins UPS1 Data Stable_Sigmoid (4, 4, 4, 4, 4 _ 2, 4, 4, 25, 25)
Normalized Intensity
10
15
20
25
Condition2
Condition1
1
2
3
4
5
Timepoint

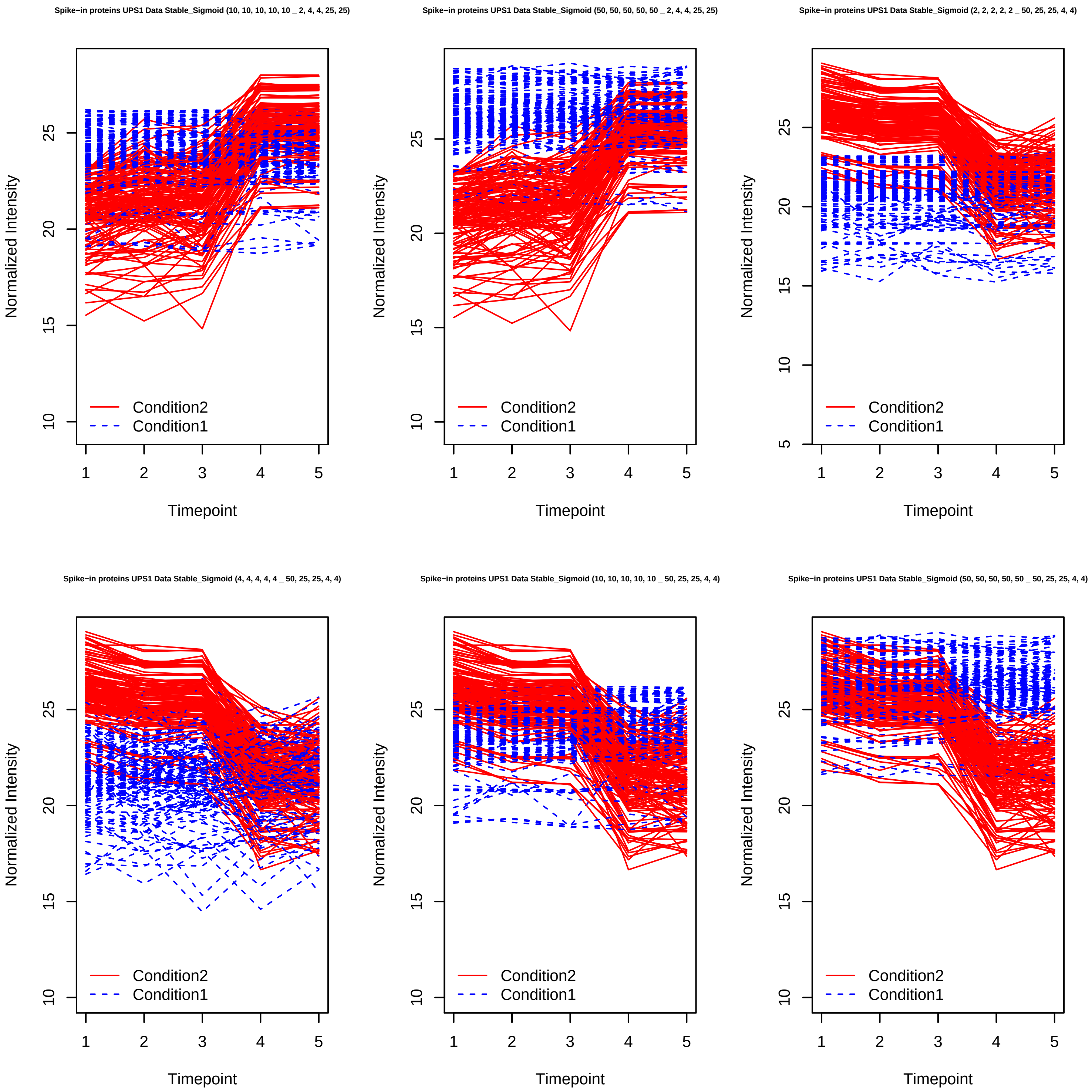

Spike−in proteins UPS1 Data Stable_Sigmoid (10, 10, 10, 10, 10 _ 2, 4, 4, 25, 25)
Spike−in proteins UPS1 Data Stable_Sigmoid (50, 50, 50, 50, 50 _ 2, 4, 4, 25, 25)
Spike−in proteins UPS1 Data Stable_Sigmoid (2, 2, 2, 2, 2 _ 50, 25, 25, 4, 4)
Spike−in proteins UPS1 Data Stable_Sigmoid (4, 4, 4, 4, 4 _ 50, 25, 25, 4, 4)
Spike−in proteins UPS1 Data Stable_Sigmoid (10, 10, 10, 10, 10 _ 50, 25, 25, 4, 4)
Spike−in proteins UPS1 Data Stable_Sigmoid (50, 50, 50, 50, 50 _ 50, 25, 25, 4, 4)
Normalized Intensity
Timepoint
Condition2
Condition1

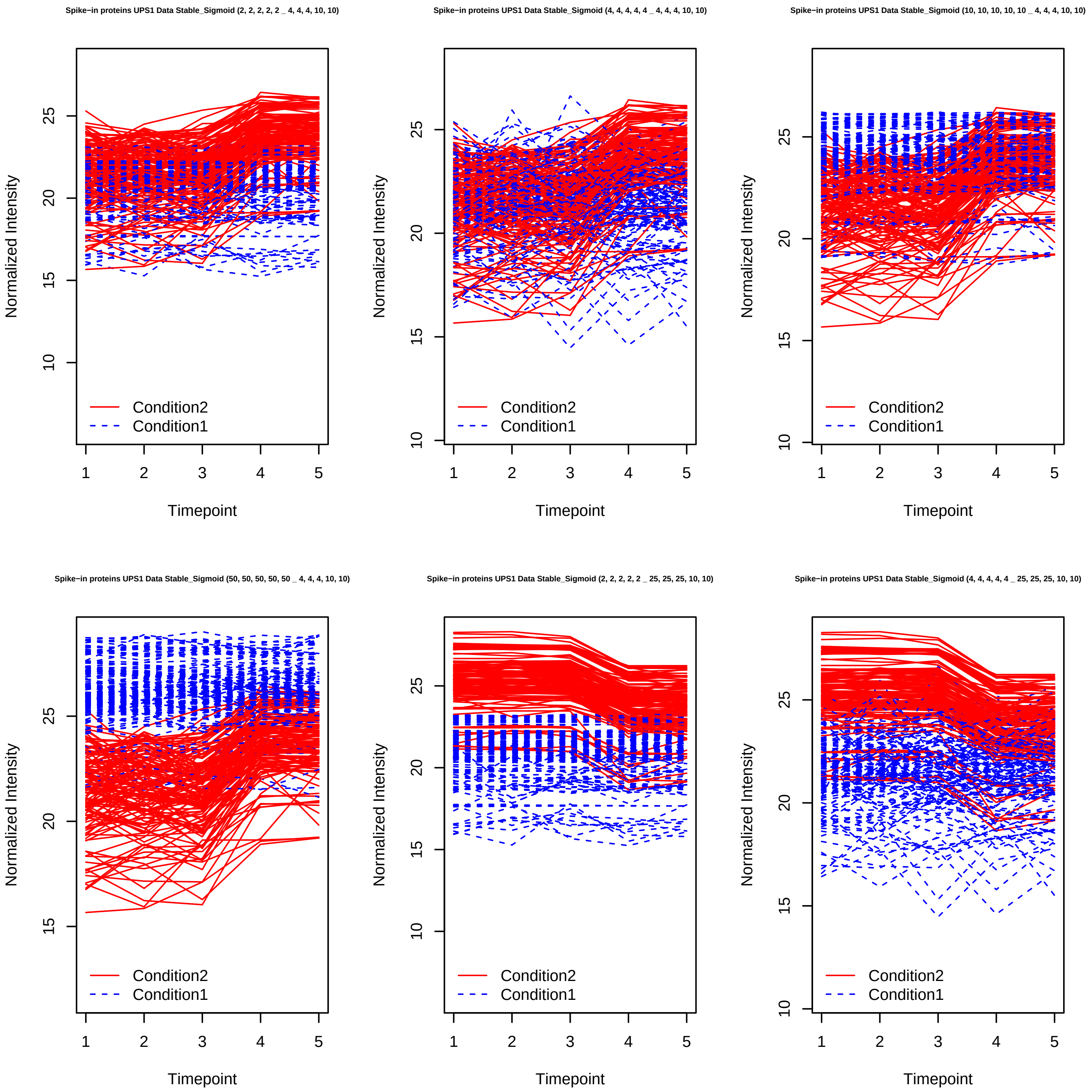
Spike−in proteins UPS1 Data Stable_Sigmoid (2, 2, 2, 2, 2 _ 4, 4, 4, 10, 10)
Spike−in proteins UPS1 Data Stable_Sigmoid (4, 4, 4, 4, 4 _ 4, 4, 4, 10, 10)
Spike−in proteins UPS1 Data Stable_Sigmoid (10, 10, 10, 10, 10 _ 4, 4, 4, 10, 10)
Spike−in proteins UPS1 Data Stable_Sigmoid (50, 50, 50, 50, 50 _ 4, 4, 4, 10, 10)
Spike−in proteins UPS1 Data Stable_Sigmoid (2, 2, 2, 2, 2 _ 25, 25, 25, 10, 10)
Spike−in proteins UPS1 Data Stable_Sigmoid (4, 4, 4, 4, 4 _ 25, 25, 25, 10, 10)
Normalized Intensity
Timepoint
Condition2
Condition1

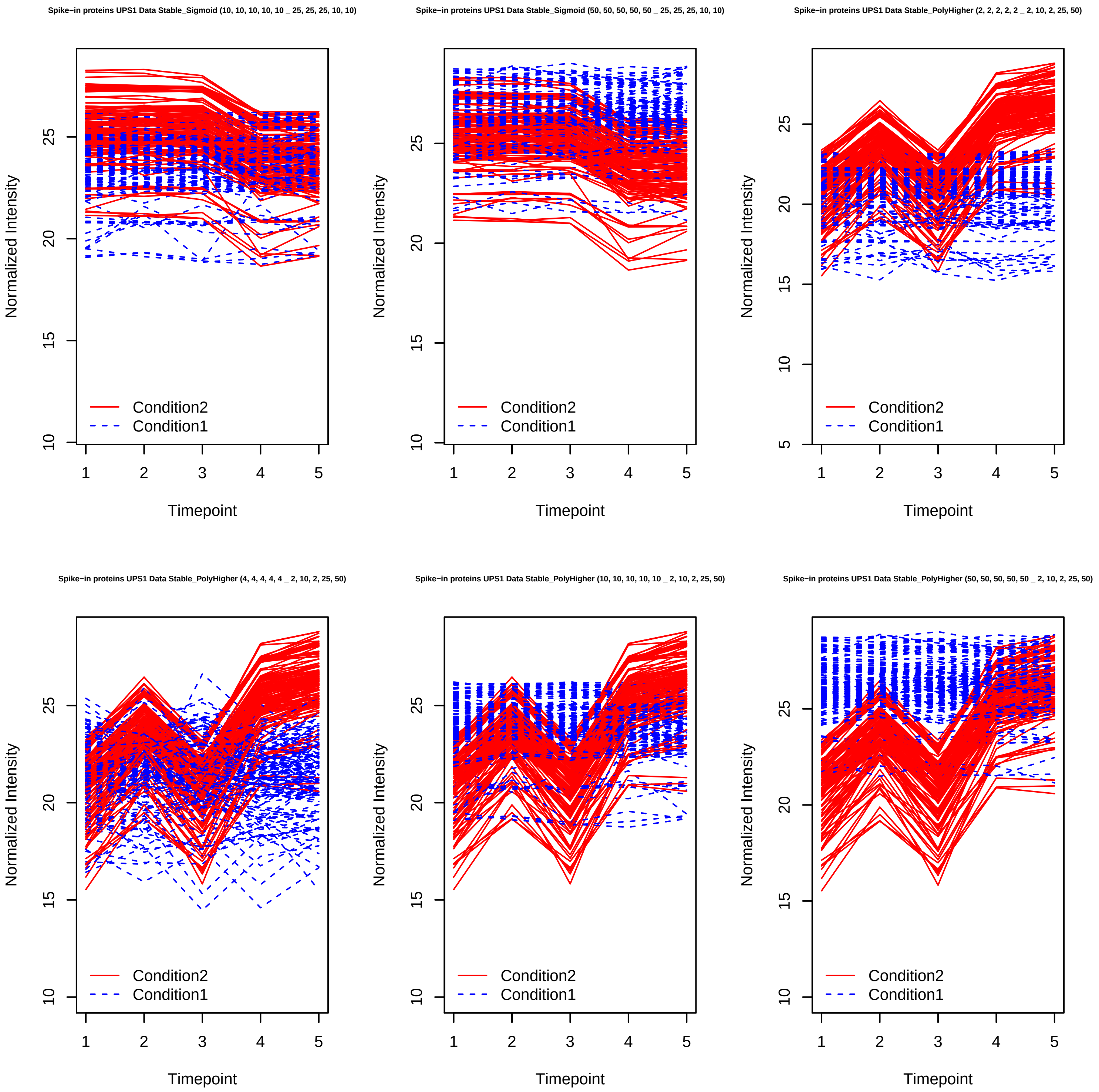
Spike−in proteins UPS1 Data Stable_Sigmoid (10, 10, 10, 10, 10 _ 25, 25, 25, 10, 10)
Spike−in proteins UPS1 Data Stable_Sigmoid (50, 50, 50, 50, 50 _ 25, 25, 25, 10, 10)
Spike−in proteins UPS1 Data Stable_PolyHigher (2, 2, 2, 2, 2 _ 2, 10, 2, 25, 50)
Spike−in proteins UPS1 Data Stable_PolyHigher (4, 4, 4, 4, 4 _ 2, 10, 2, 25, 50)
Spike−in proteins UPS1 Data Stable_PolyHigher (10, 10, 10, 10, 10 _ 2, 10, 2, 25, 50)
Spike−in proteins UPS1 Data Stable_PolyHigher (50, 50, 50, 50, 50 _ 2, 10, 2, 25, 50)
Normalized Intensity
Timepoint
Condition2
Condition1

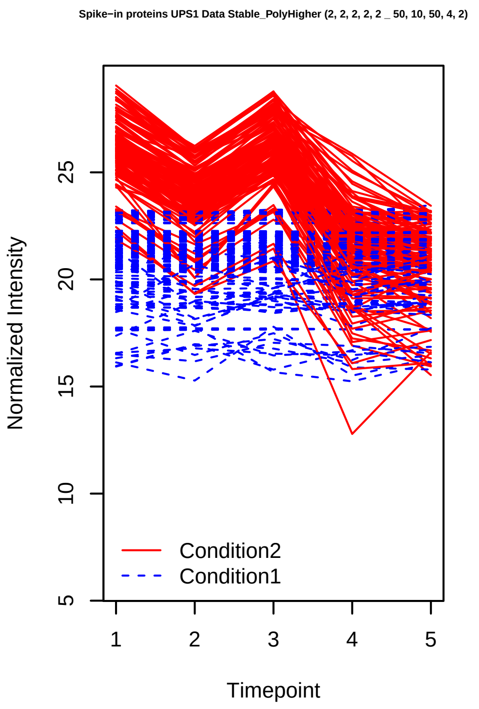
Spike-in proteins UPS1 Data Stable_PolyHigher (2, 2, 2, 2, 2 _ 50, 10, 50, 4, 2)
Normalized Intensity
Timepoint
Condition2
Condition1

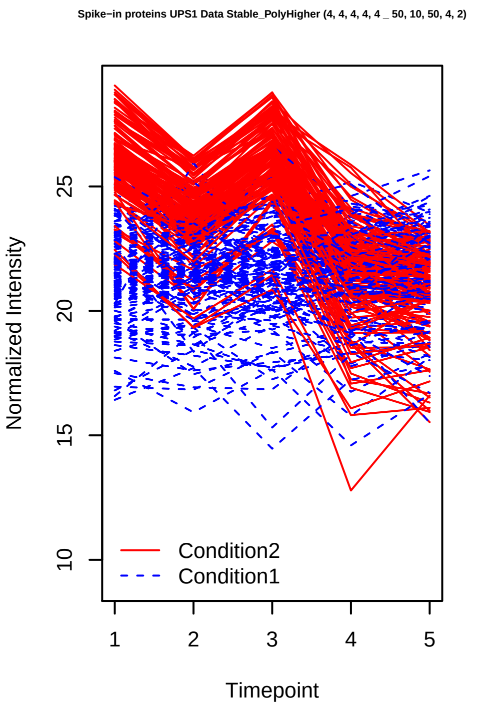
Spike-in proteins UPS1 Data Stable_PolyHigher (4, 4, 4, 4, 4 _ 50, 10, 50, 4, 2)
Normalized Intensity
Timepoint
Condition2
Condition1

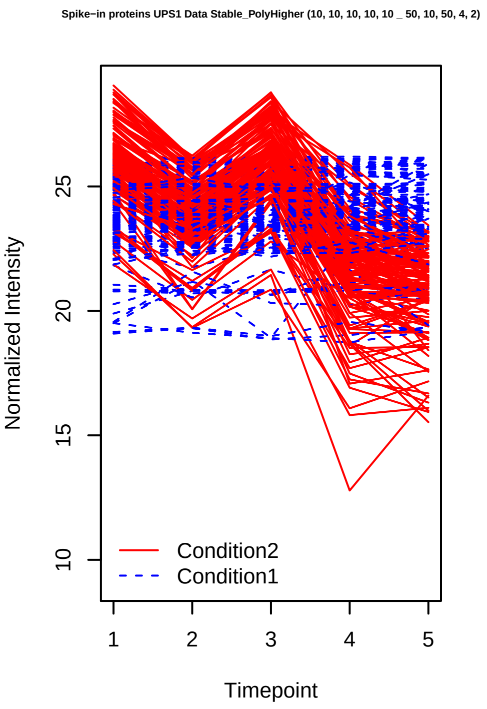
Spike-in proteins UPS1 Data Stable_PolyHigher (10, 10, 10, 10, 10 _ 50, 10, 50, 4, 2)
Normalized Intensity
Timepoint
Condition2
Condition1

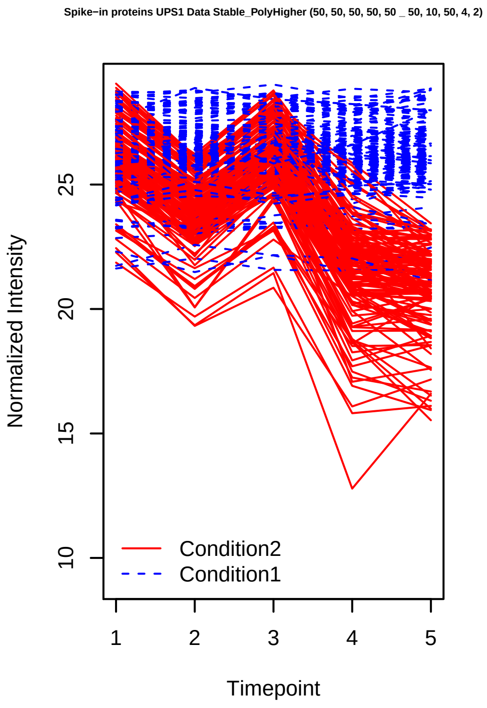
Spike-in proteins UPS1 Data Stable_PolyHigher (50, 50, 50, 50, 50 _ 50, 10, 50, 4, 2)
Normalized Intensity
Timepoint
Condition2
Condition1

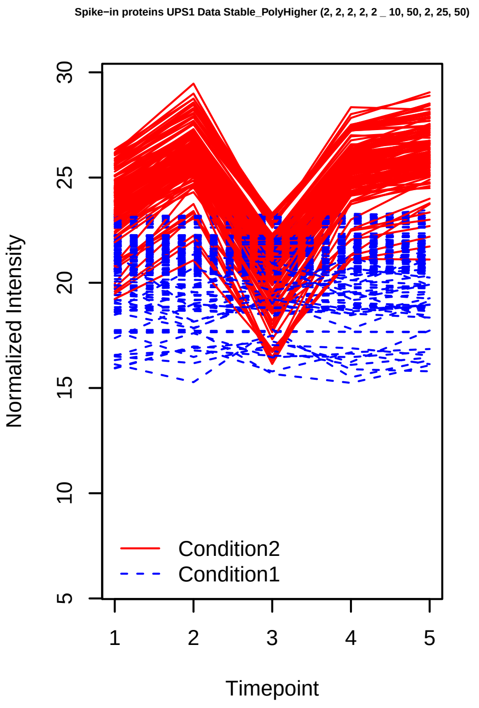
Spike-in proteins UPS1 Data Stable_PolyHigher (2, 2, 2, 2, 2 _ 10, 50, 2, 25, 50)
Normalized Intensity
Timepoint
Condition2
Condition1

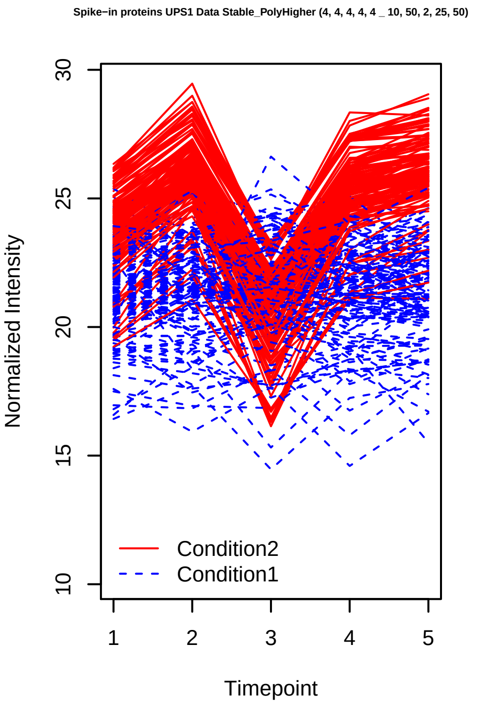
Spike-in proteins UPS1 Data Stable_PolyHigher (4, 4, 4, 4, 4 _ 10, 50, 2, 25, 50)
Normalized Intensity
Timepoint
Condition2
Condition1

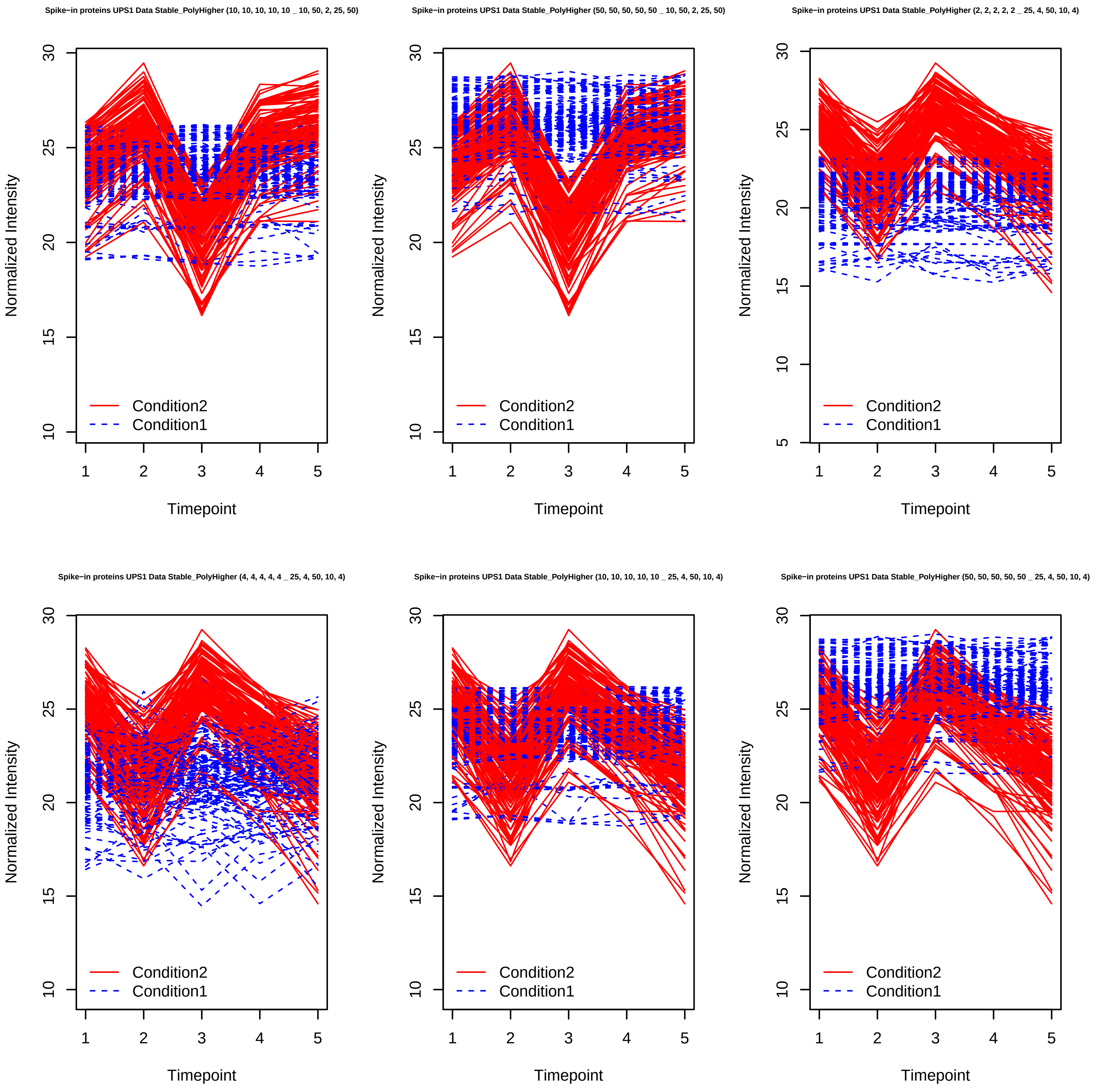
Spike−in proteins UPS1 Data Stable_PolyHigher (10, 10, 10, 10, 10 _ 10, 50, 2, 25, 50)
Normalized Intensity
Timepoint
Condition2
Condition1
Spike−in proteins UPS1 Data Stable_PolyHigher (50, 50, 50, 50, 50 _ 10, 50, 2, 25, 50)
Normalized Intensity
Timepoint
Condition2
Condition1
Spike−in proteins UPS1 Data Stable_PolyHigher (2, 2, 2, 2, 2 _ 25, 4, 50, 10, 4)
Normalized Intensity
Timepoint
Condition2
Condition1
Spike−in proteins UPS1 Data Stable_PolyHigher (4, 4, 4, 4, 4 _ 25, 4, 50, 10, 4)
Normalized Intensity
Timepoint
Condition2
Condition1
Spike−in proteins UPS1 Data Stable_PolyHigher (10, 10, 10, 10, 10 _ 25, 4, 50, 10, 4)
Normalized Intensity
Timepoint
Condition2
Condition1
Spike−in proteins UPS1 Data Stable_PolyHigher (50, 50, 50, 50, 50 _ 25, 4, 50, 10, 4)
Normalized Intensity
Timepoint
Condition2
Condition1

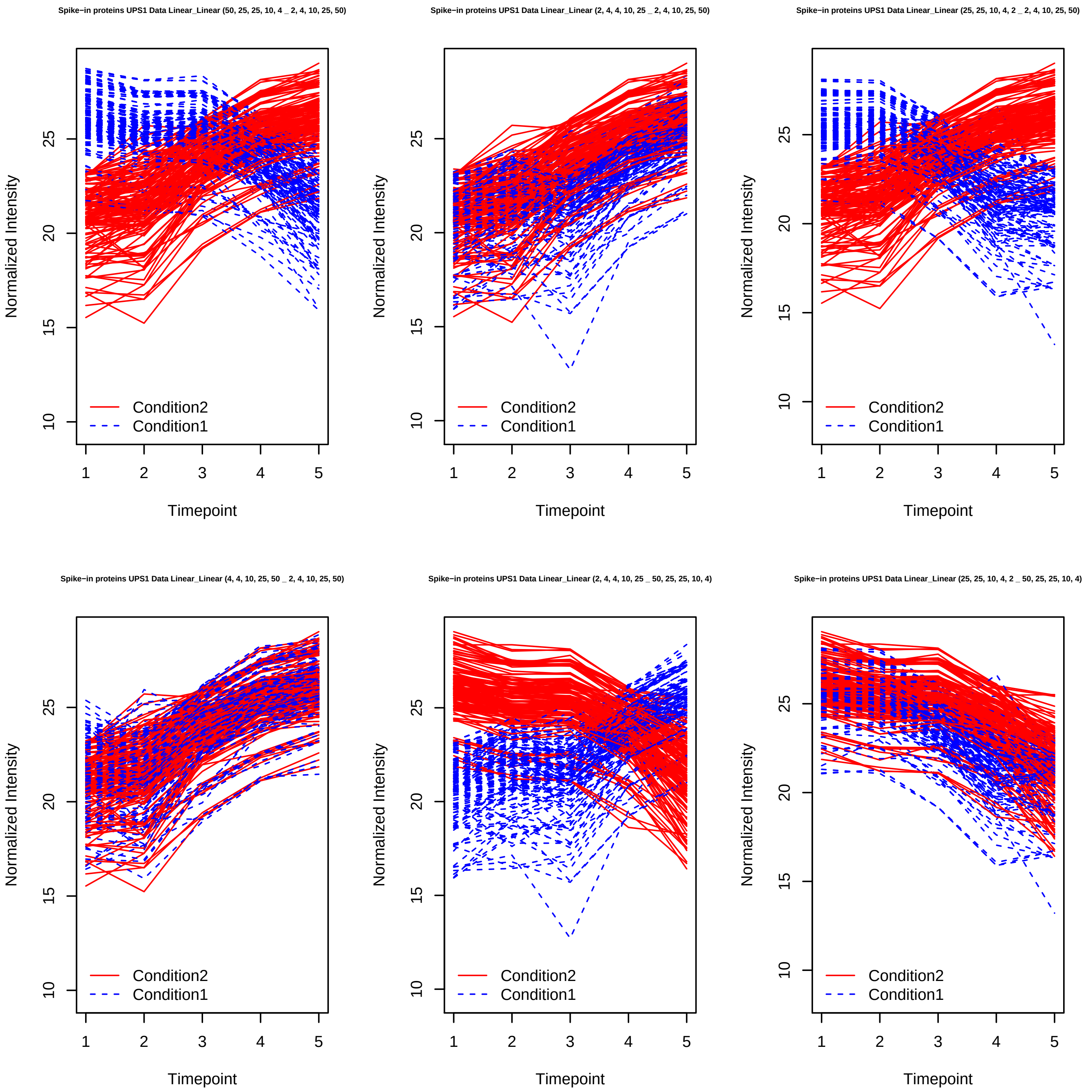
Spike−in proteins UPS1 Data Linear_Linear (50, 25, 25, 10, 4 _ 2, 4, 10, 25, 50)
Spike−in proteins UPS1 Data Linear_Linear (2, 4, 4, 10, 25 _ 2, 4, 10, 25, 50)
Spike−in proteins UPS1 Data Linear_Linear (25, 25, 10, 4, 2 _ 2, 4, 10, 25, 50)
Spike−in proteins UPS1 Data Linear_Linear (4, 4, 10, 25, 50 _ 2, 4, 10, 25, 50)
Spike−in proteins UPS1 Data Linear_Linear (2, 4, 4, 10, 25 _ 50, 25, 25, 10, 4)
Spike−in proteins UPS1 Data Linear_Linear (25, 25, 10, 4, 2 _ 50, 25, 25, 10, 4)
Normalized Intensity
Timepoint
Condition2
Condition1

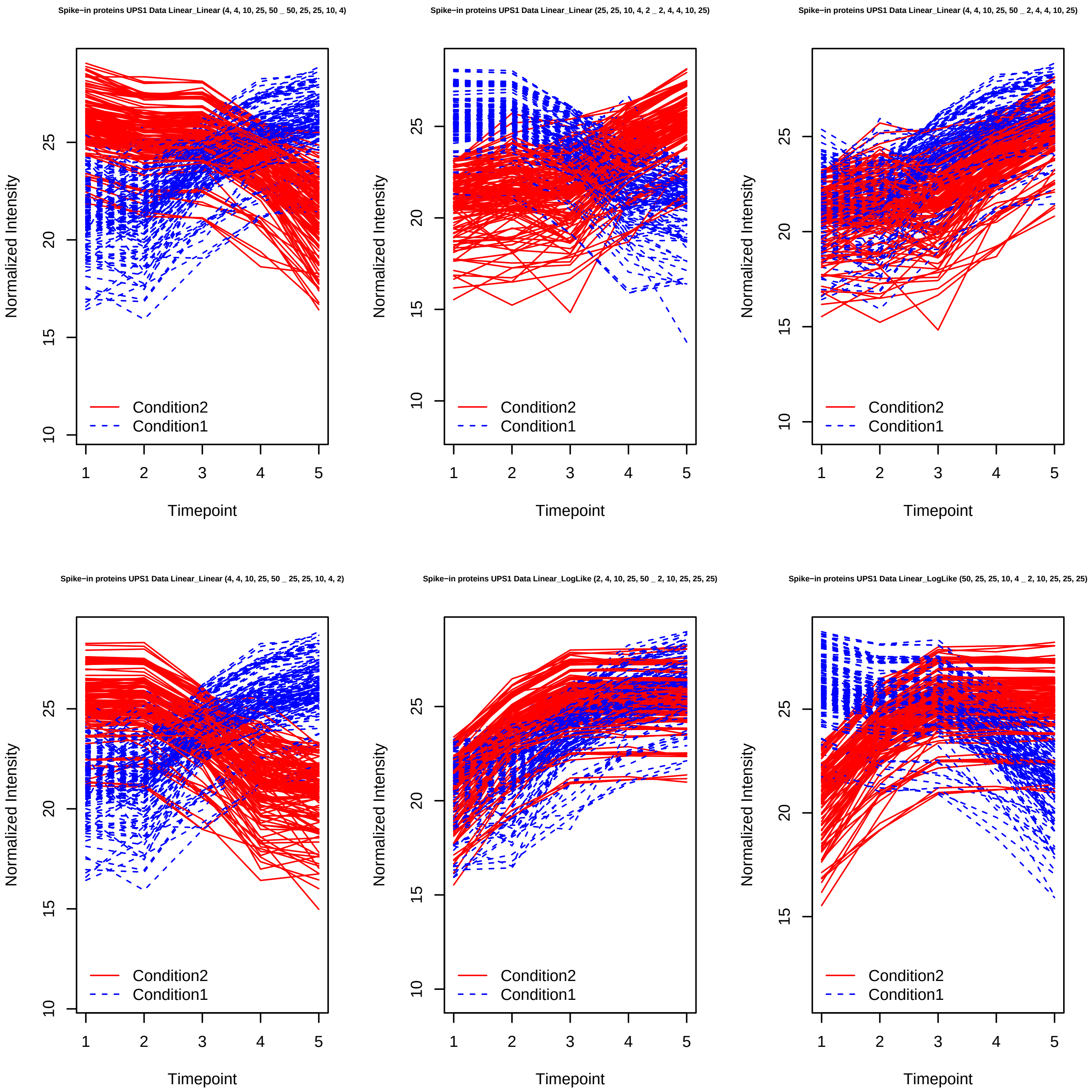

Spike−in proteins UPS1 Data Linear_Linear (4, 4, 10, 25, 50 _ 50, 25, 25, 10, 4)
Spike−in proteins UPS1 Data Linear_Linear (25, 25, 10, 4, 2 _ 2, 4, 4, 10, 25)
Spike−in proteins UPS1 Data Linear_Linear (4, 4, 10, 25, 50 _ 2, 4, 4, 10, 25)
Spike−in proteins UPS1 Data Linear_Linear (4, 4, 10, 25, 50 _ 25, 25, 10, 4, 2)
Spike−in proteins UPS1 Data Linear_LogLike (2, 4, 10, 25, 50 _ 2, 10, 25, 25, 25)
Spike−in proteins UPS1 Data Linear_LogLike (50, 25, 25, 10, 4 _ 2, 10, 25, 25, 25)
Normalized Intensity
Timepoint
Condition2
Condition1

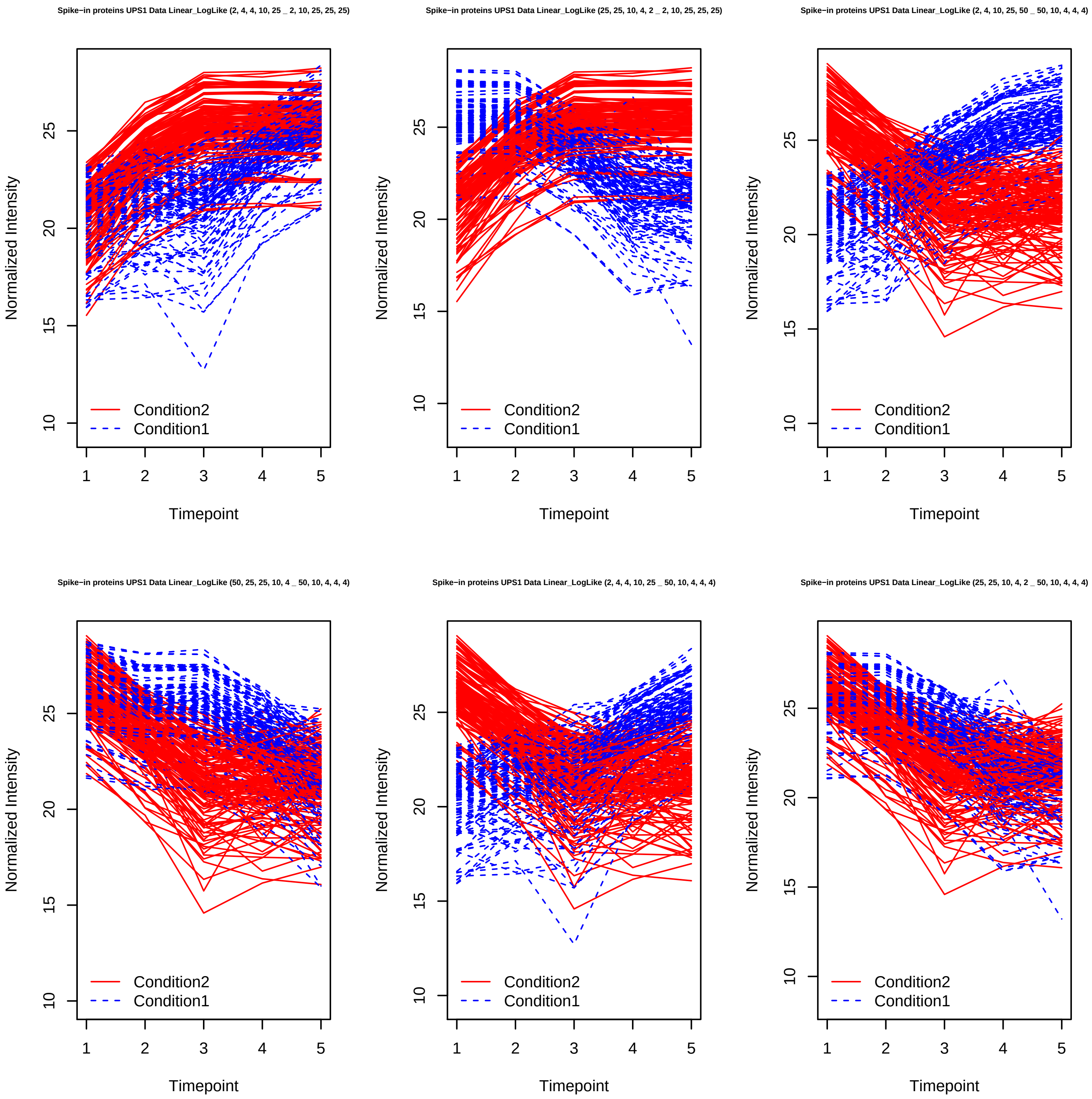
Spike−in proteins UPS1 Data Linear_LogLike (2, 4, 4, 10, 25 _ 2, 10, 25, 25, 25)
Spike−in proteins UPS1 Data Linear_LogLike (25, 25, 10, 4, 2 _ 2, 10, 25, 25, 25)
Spike−in proteins UPS1 Data Linear_LogLike (2, 4, 10, 25, 50 _ 50, 10, 4, 4, 4)
Spike−in proteins UPS1 Data Linear_LogLike (50, 25, 25, 10, 4 _ 50, 10, 4, 4, 4)
Spike−in proteins UPS1 Data Linear_LogLike (2, 4, 4, 10, 25 _ 50, 10, 4, 4, 4)
Spike−in proteins UPS1 Data Linear_LogLike (25, 25, 10, 4, 2 _ 50, 10, 4, 4, 4)
Normalized Intensity
Timepoint
Condition2
Condition1

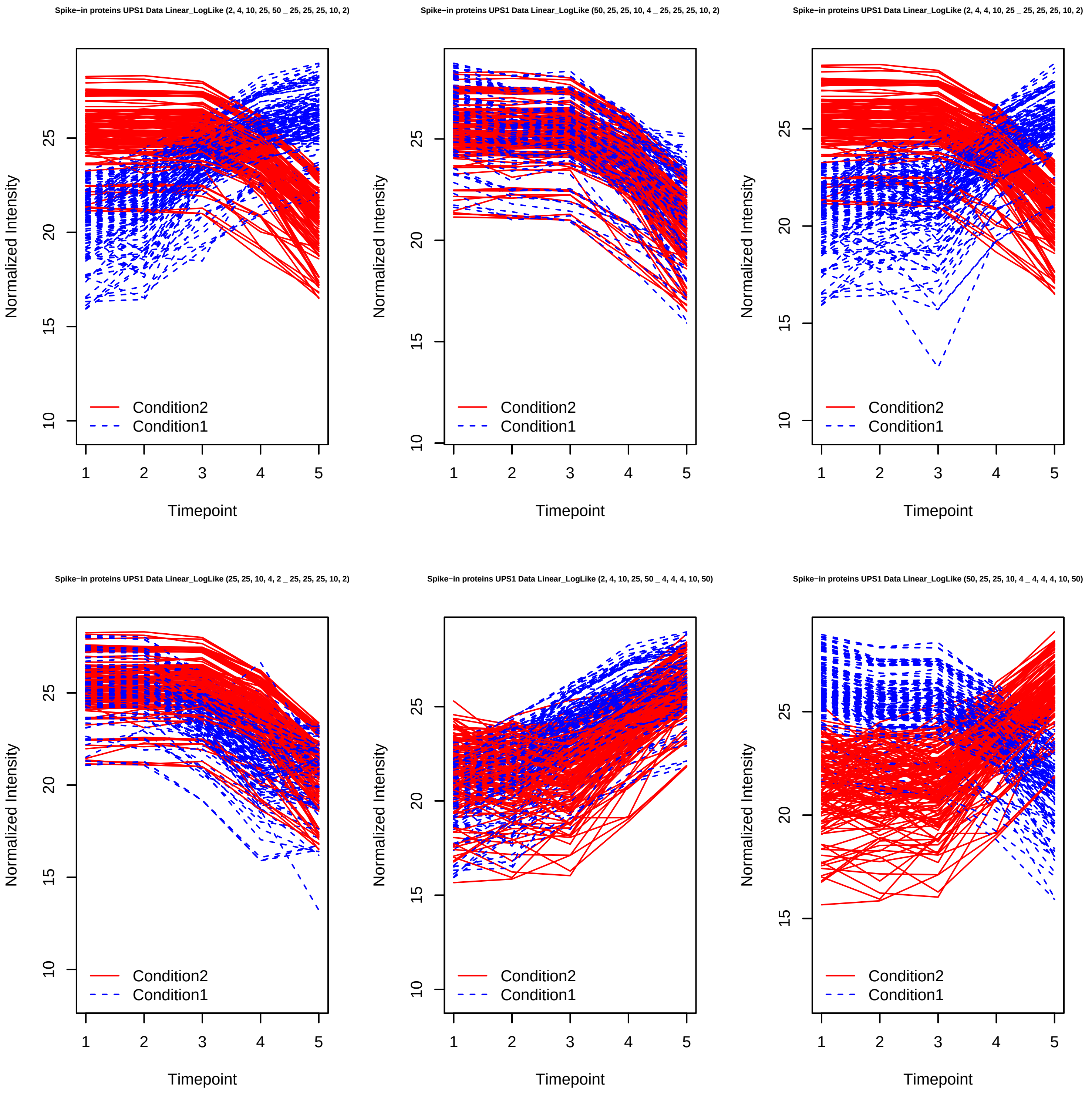
Spike−in proteins UPS1 Data Linear_LogLike (2, 4, 10, 25, 50 _ 25, 25, 25, 10, 2)
Spike−in proteins UPS1 Data Linear_LogLike (50, 25, 25, 10, 4 _ 25, 25, 25, 10, 2)
Spike−in proteins UPS1 Data Linear_LogLike (2, 4, 4, 10, 25 _ 25, 25, 25, 10, 2)
Spike−in proteins UPS1 Data Linear_LogLike (25, 25, 10, 4, 2 _ 25, 25, 25, 10, 2)
Spike−in proteins UPS1 Data Linear_LogLike (2, 4, 10, 25, 50 _ 4, 4, 4, 10, 50)
Spike−in proteins UPS1 Data Linear_LogLike (50, 25, 25, 10, 4 _ 4, 4, 4, 10, 50)
Normalized Intensity
Timepoint
Condition2
Condition1

Spike−in proteins UPS1 Data Linear_LogLike (2, 4, 4, 10, 25 _ 4, 4, 4, 10, 50)

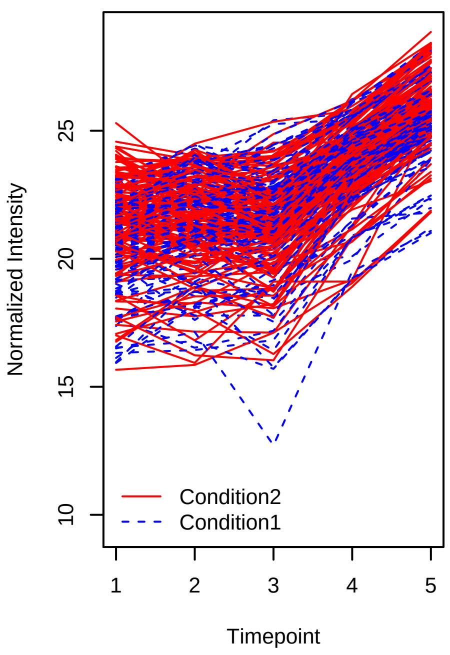


Spike−in proteins UPS1 Data Linear_LogLike (25, 25, 10, 4, 2 _ 4, 4, 4, 10, 50)

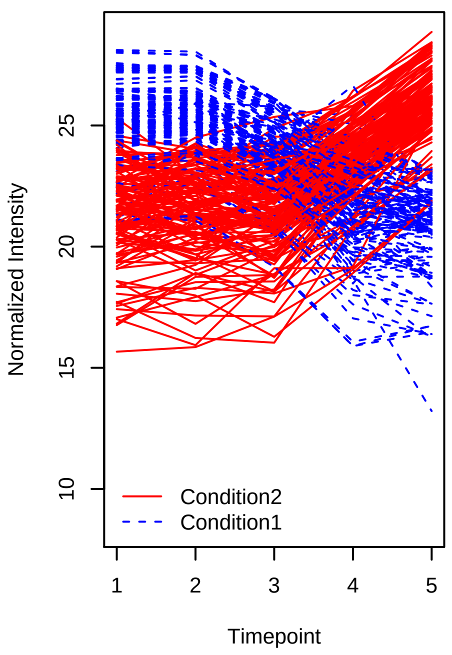


Spike−in proteins UPS1 Data Linear_Poly2 (2, 4, 10, 25, 50 _ 2, 4, 10, 4, 2)

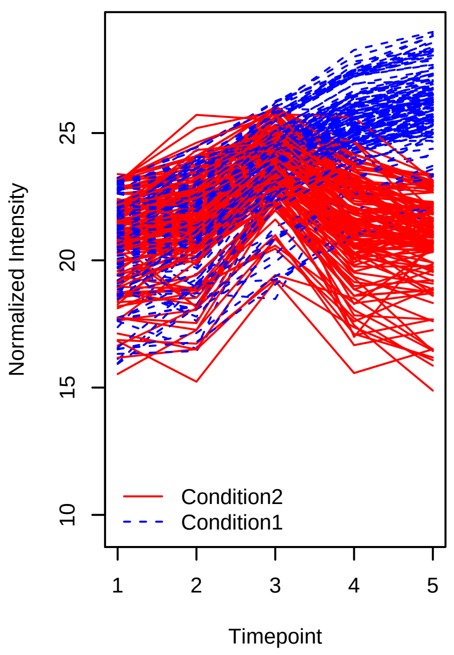


Spike−in proteins UPS1 Data Linear_Poly2 (50, 25, 25, 10, 4 _ 2, 4, 10, 4, 2)

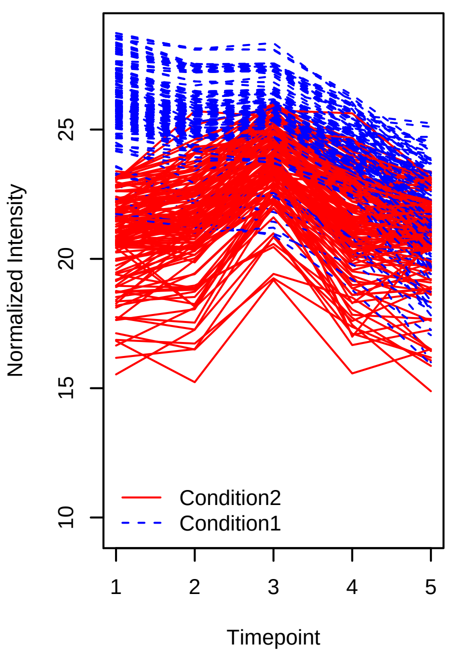


Spike−in proteins UPS1 Data Linear_Poly2 (2, 4, 4, 10, 25 _ 2, 4, 10, 4, 2)

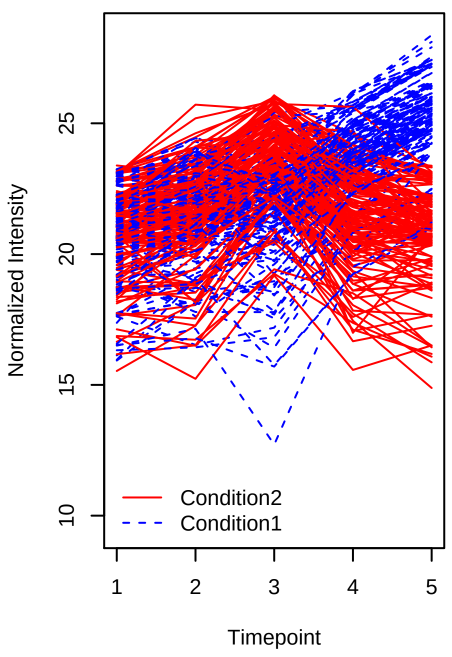


Spike−in proteins UPS1 Data Linear_Poly2 (25, 25, 10, 4, 2 _ 2, 4, 10, 4, 2)

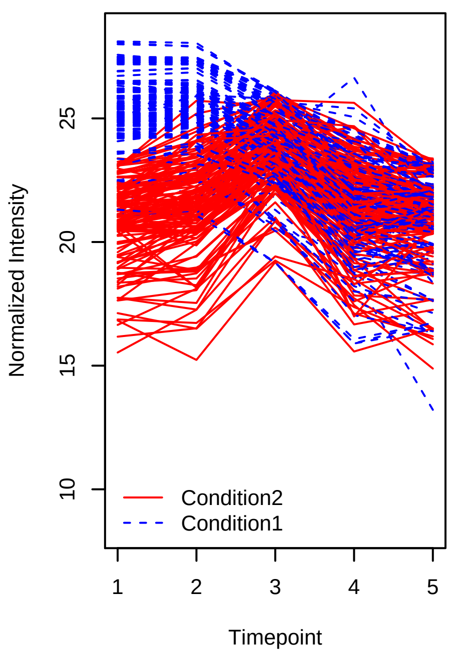

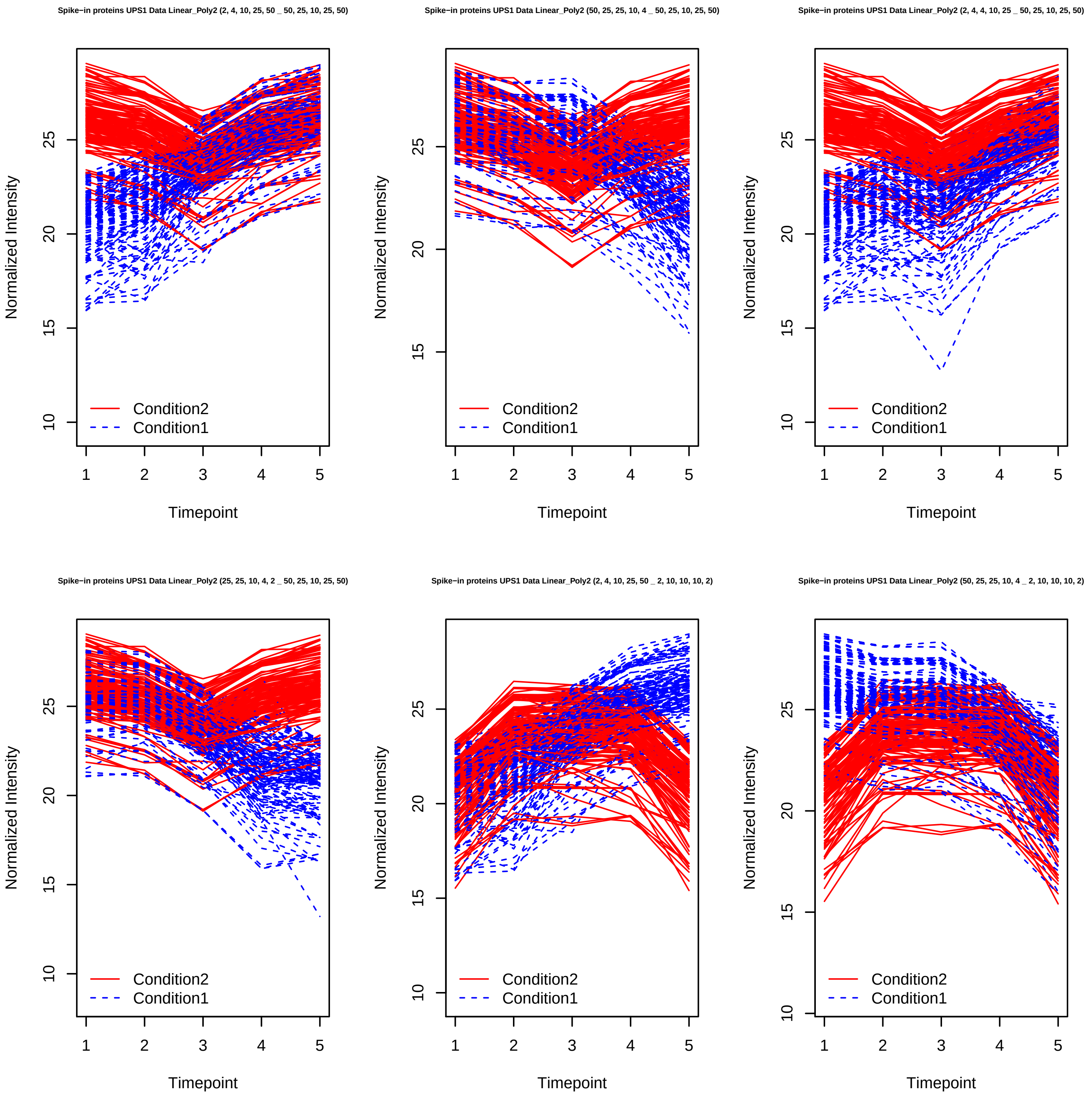
Spike−in proteins UPS1 Data Linear_Poly2 (2, 4, 10, 25, 50 _ 50, 25, 10, 25, 50)
Spike−in proteins UPS1 Data Linear_Poly2 (50, 25, 25, 10, 4 _ 50, 25, 10, 25, 50)
Spike−in proteins UPS1 Data Linear_Poly2 (2, 4, 4, 10, 25 _ 50, 25, 10, 25, 50)
Spike−in proteins UPS1 Data Linear_Poly2 (25, 25, 10, 4, 2 _ 50, 25, 10, 25, 50)
Spike−in proteins UPS1 Data Linear_Poly2 (2, 4, 10, 25, 50 _ 2, 10, 10, 10, 2)
Spike−in proteins UPS1 Data Linear_Poly2 (50, 25, 25, 10, 4 _ 2, 10, 10, 10, 2)
Normalized Intensity
Timepoint
Condition2
Condition1

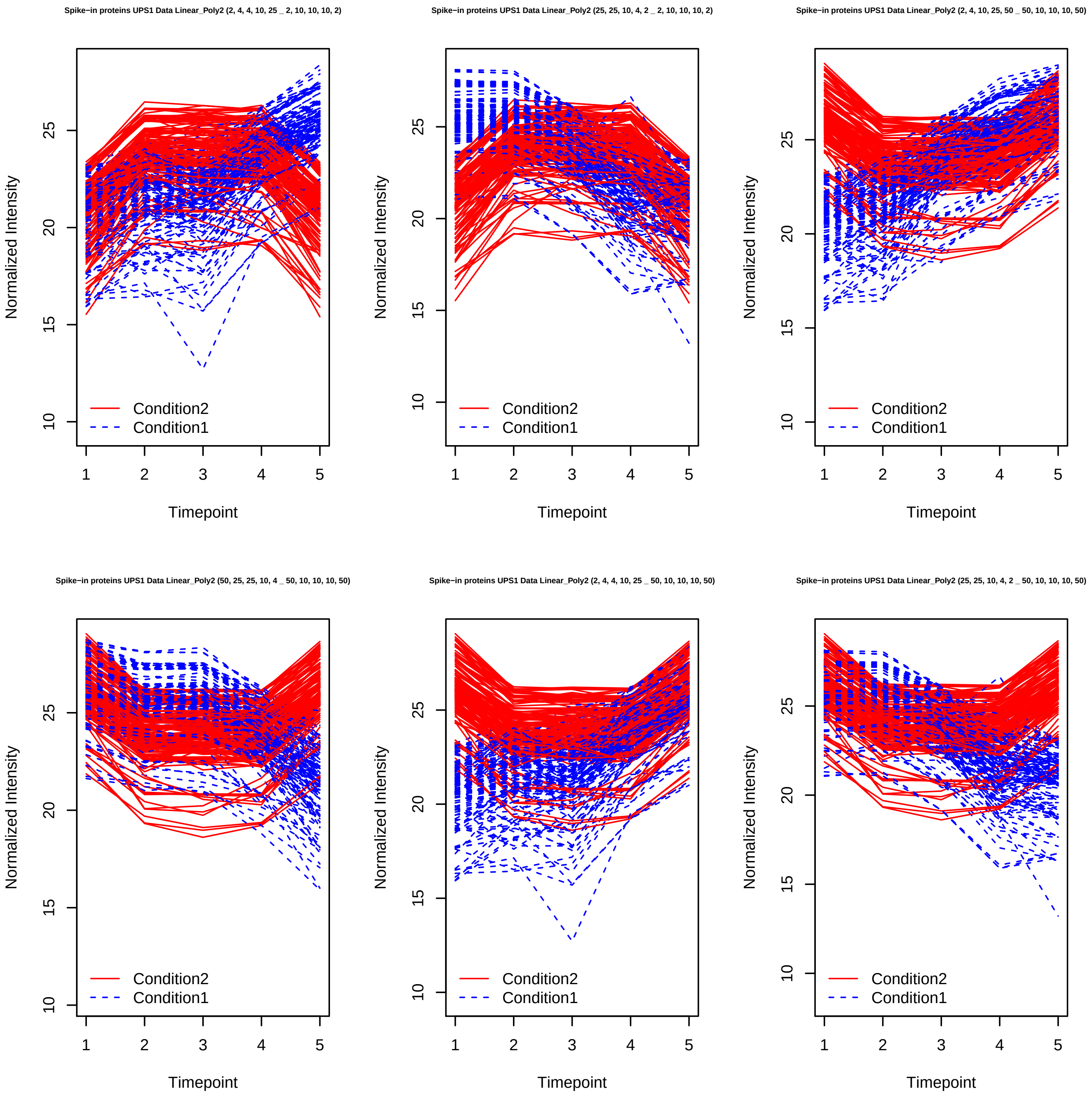
Spike−in proteins UPS1 Data Linear_Poly2 (2, 4, 4, 10, 25 _ 2, 10, 10, 10, 2)
Spike−in proteins UPS1 Data Linear_Poly2 (25, 25, 10, 4, 2 _ 2, 10, 10, 10, 2)
Spike−in proteins UPS1 Data Linear_Poly2 (2, 4, 10, 25, 50 _ 50, 10, 10, 10, 50)
Spike−in proteins UPS1 Data Linear_Poly2 (50, 25, 25, 10, 4 _ 50, 10, 10, 10, 50)
Spike−in proteins UPS1 Data Linear_Poly2 (2, 4, 4, 10, 25 _ 50, 10, 10, 10, 50)
Spike−in proteins UPS1 Data Linear_Poly2 (25, 25, 10, 4, 2 _ 50, 10, 10, 10, 50)
Normalized Intensity
Timepoint
Condition2
Condition1

Spike−in proteins UPS1 Data Linear_Sigmoid (2, 4, 10, 25, 50 _ 2, 4, 4, 25, 25)

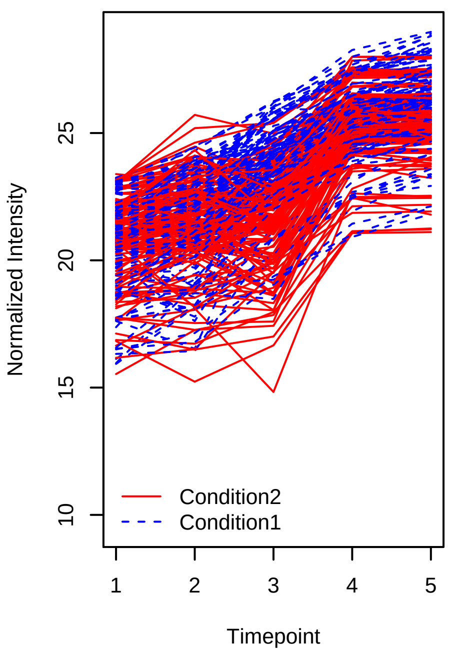


Spike−in proteins UPS1 Data Linear_Sigmoid (50, 25, 25, 10, 4 _ 2, 4, 4, 25, 25)

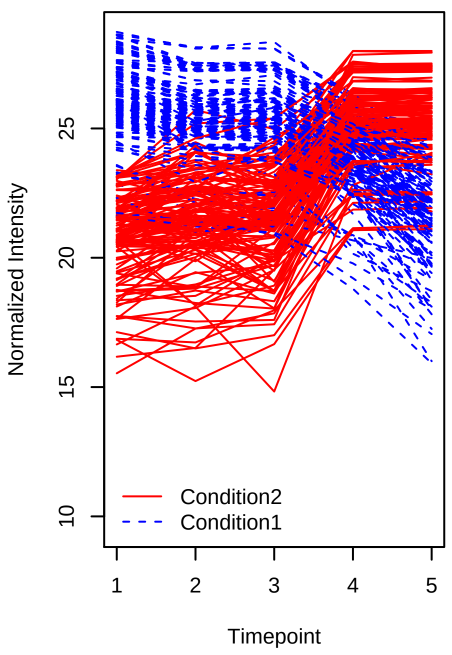


Spike−in proteins UPS1 Data Linear_Sigmoid (2, 4, 4, 10, 25 _ 2, 4, 4, 25, 25)

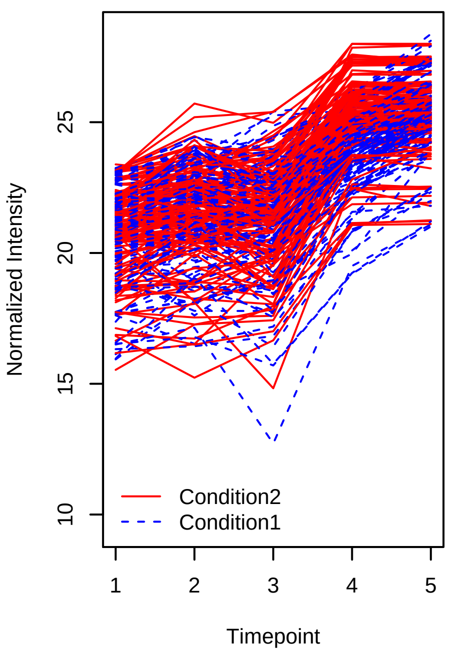


Spike−in proteins UPS1 Data Linear_Sigmoid (25, 25, 10, 4, 2 _ 2, 4, 4, 25, 25)

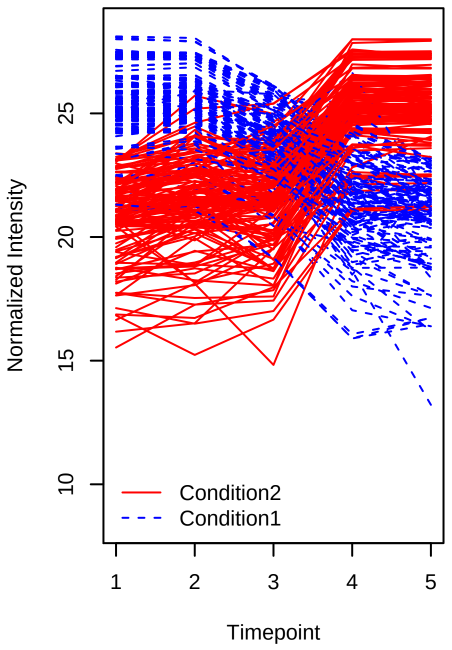


Spike−in proteins UPS1 Data Linear_Sigmoid (2, 4, 10, 25, 50 _ 50, 25, 25, 4, 4)

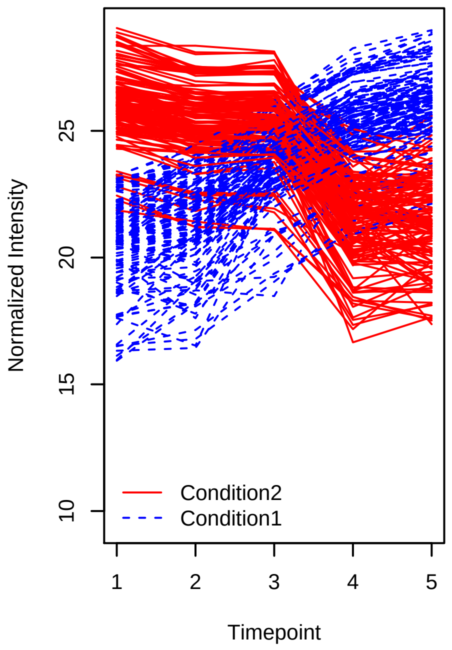


Spike−in proteins UPS1 Data Linear_Sigmoid (50, 25, 25, 10, 4 _ 50, 25, 25, 4, 4)

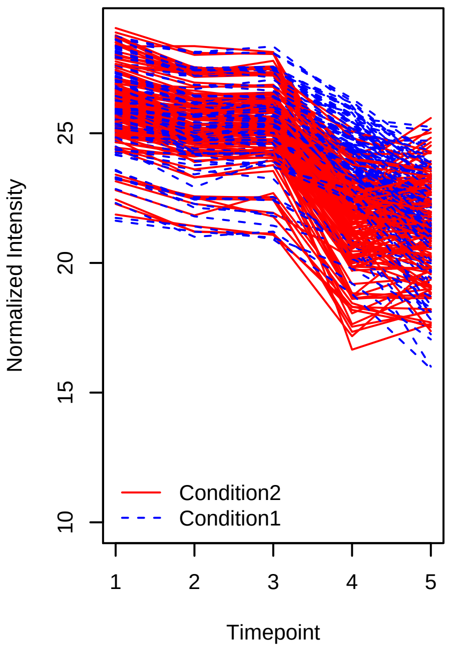

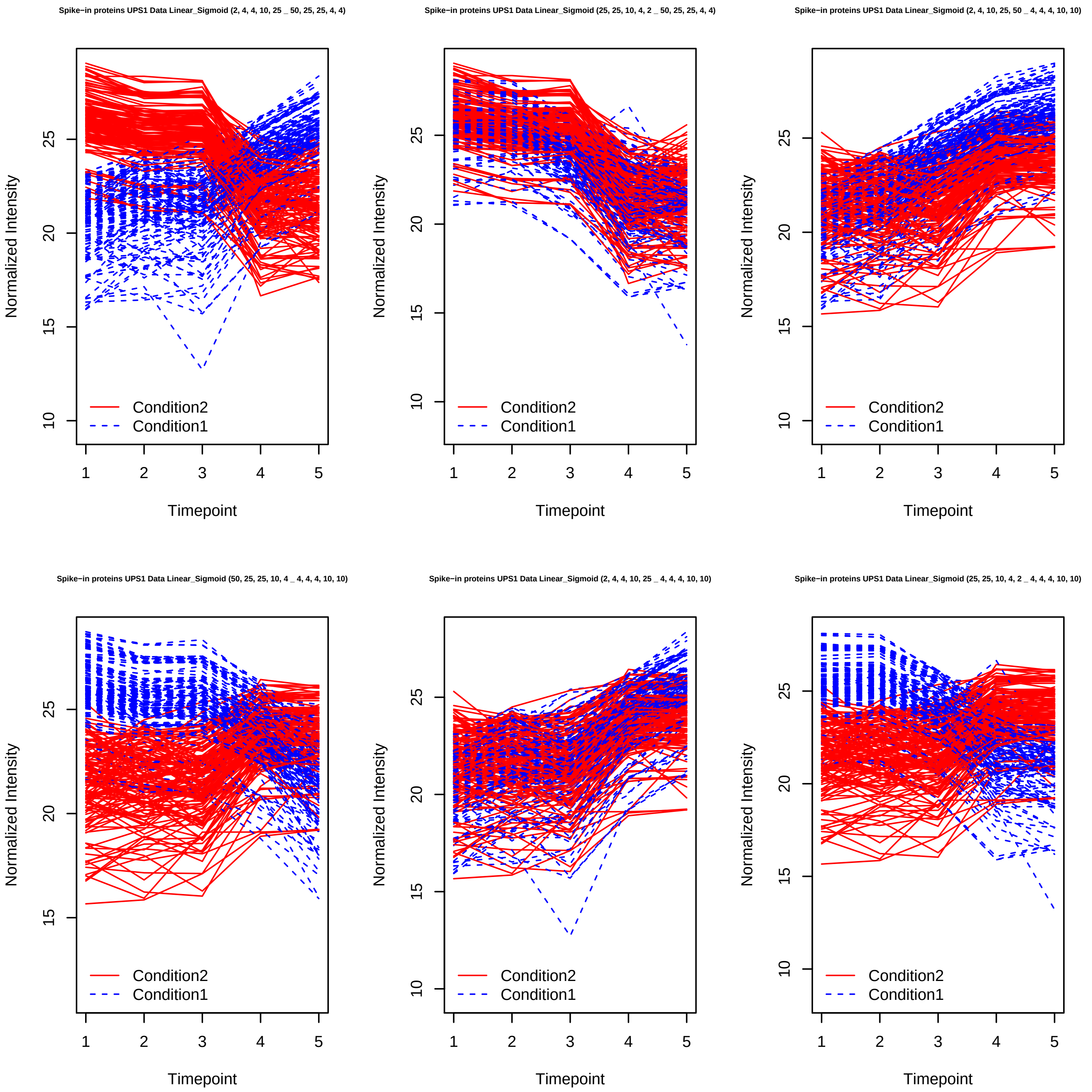
Spike−in proteins UPS1 Data Linear_Sigmoid (2, 4, 4, 10, 25 _ 50, 25, 25, 4, 4)
Spike−in proteins UPS1 Data Linear_Sigmoid (25, 25, 10, 4, 2 _ 50, 25, 25, 4, 4)
Spike−in proteins UPS1 Data Linear_Sigmoid (2, 4, 10, 25, 50 _ 4, 4, 4, 10, 10)
Spike−in proteins UPS1 Data Linear_Sigmoid (50, 25, 25, 10, 4 _ 4, 4, 4, 10, 10)
Spike−in proteins UPS1 Data Linear_Sigmoid (2, 4, 4, 10, 25 _ 4, 4, 4, 10, 10)
Spike−in proteins UPS1 Data Linear_Sigmoid (25, 25, 10, 4, 2 _ 4, 4, 4, 10, 10)
Normalized Intensity
Timepoint
Condition2
Condition1

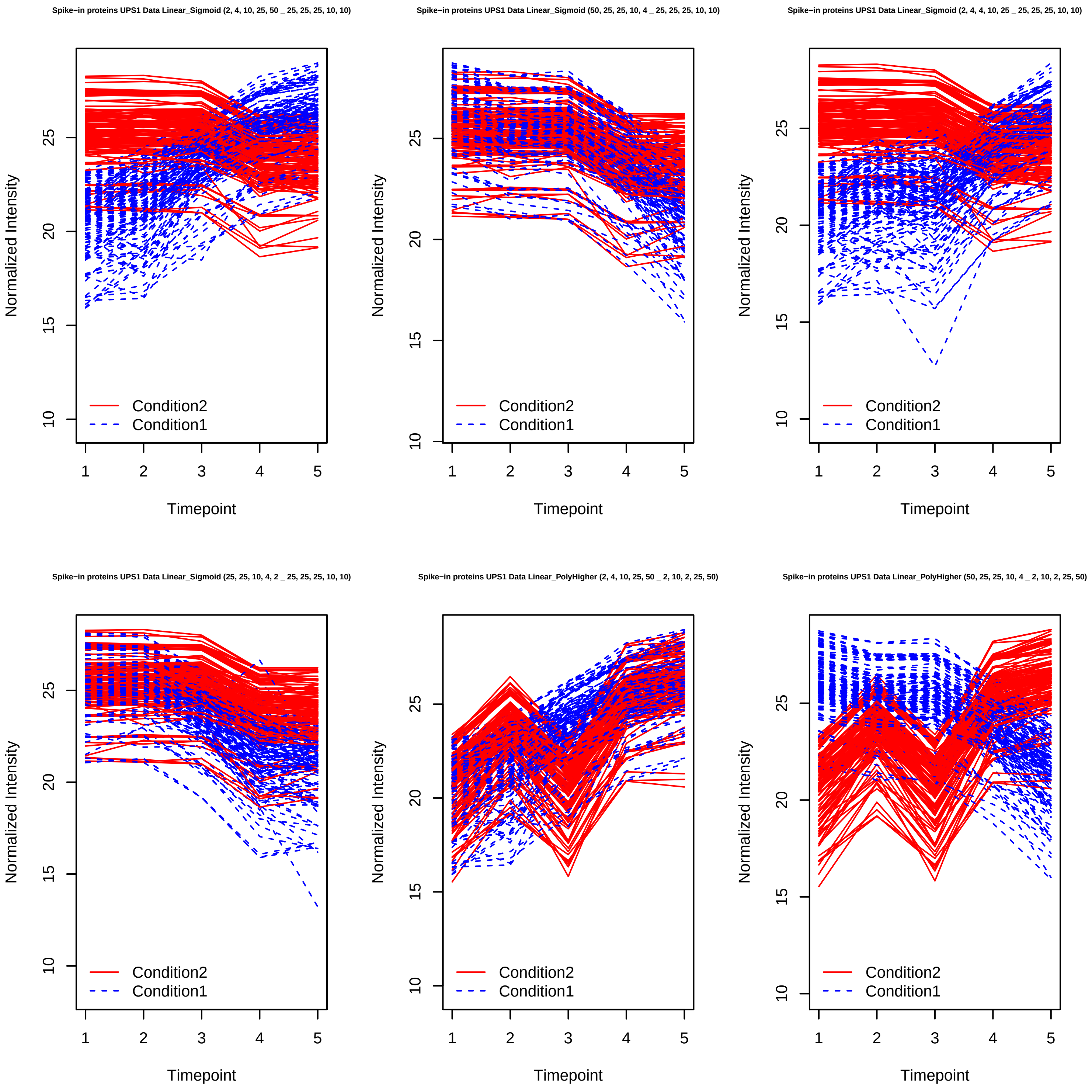
Spike−in proteins UPS1 Data Linear_Sigmoid (2, 4, 10, 25, 50 _ 25, 25, 25, 10, 10)
Spike−in proteins UPS1 Data Linear_Sigmoid (50, 25, 25, 10, 4 _ 25, 25, 25, 10, 10)
Spike−in proteins UPS1 Data Linear_Sigmoid (2, 4, 4, 10, 25 _ 25, 25, 25, 10, 10)
Spike−in proteins UPS1 Data Linear_Sigmoid (25, 25, 10, 4, 2 _ 25, 25, 25, 10, 10)
Spike−in proteins UPS1 Data Linear_PolyHigher (2, 4, 10, 25, 50 _ 2, 10, 2, 25, 50)
Spike−in proteins UPS1 Data Linear_PolyHigher (50, 25, 25, 10, 4 _ 2, 10, 2, 25, 50)
Normalized Intensity
Timepoint
Condition2
Condition1

Spike−in proteins UPS1 Data Linear_PolyHigher (2, 4, 4, 10, 25 _ 2, 10, 2, 25, 50)

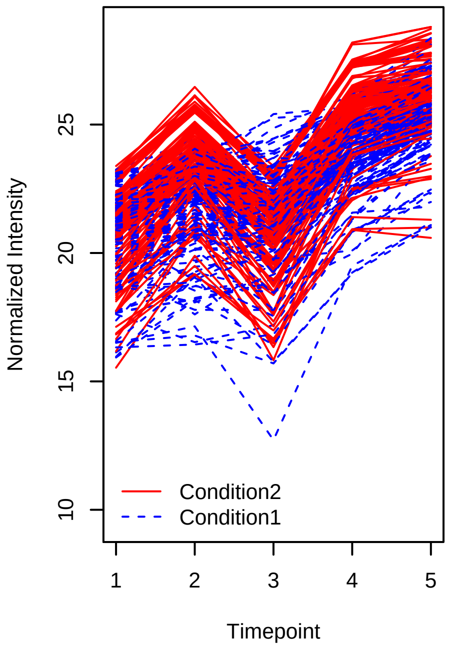


Spike−in proteins UPS1 Data Linear_PolyHigher (25, 25, 10, 4, 2 _ 2, 10, 2, 25, 50)

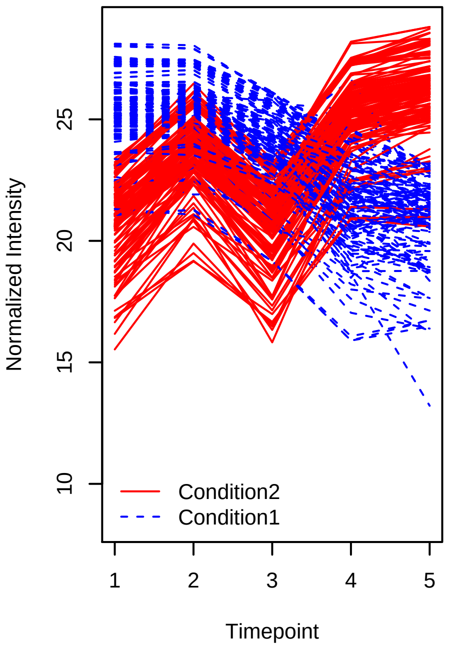


Spike−in proteins UPS1 Data Linear_PolyHigher (2, 4, 10, 25, 50 _ 50, 10, 50, 4, 2)

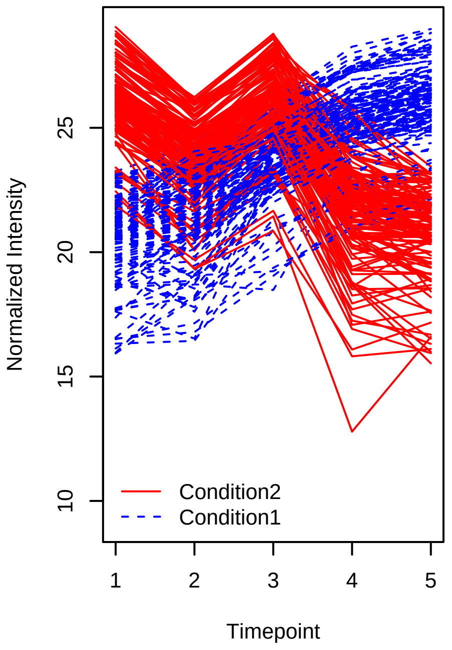


Spike−in proteins UPS1 Data Linear_PolyHigher (50, 25, 25, 10, 4 _ 50, 10, 50, 4, 2)

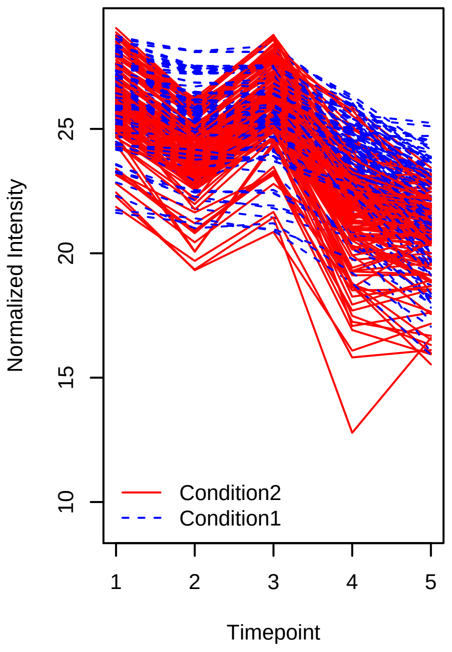


Spike−in proteins UPS1 Data Linear_PolyHigher (2, 4, 4, 10, 25 _ 50, 10, 50, 4, 2)

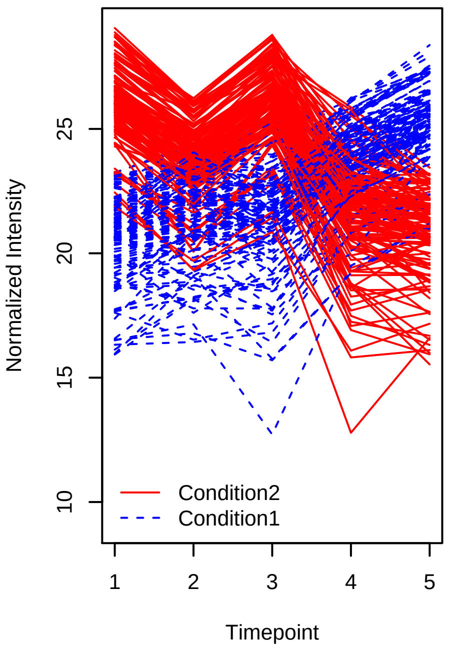


Spike−in proteins UPS1 Data Linear_PolyHigher (25, 25, 10, 4, 2 _ 50, 10, 50, 4, 2)

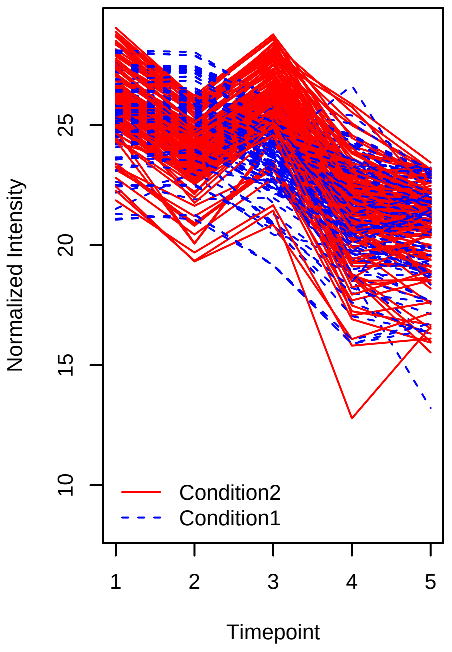

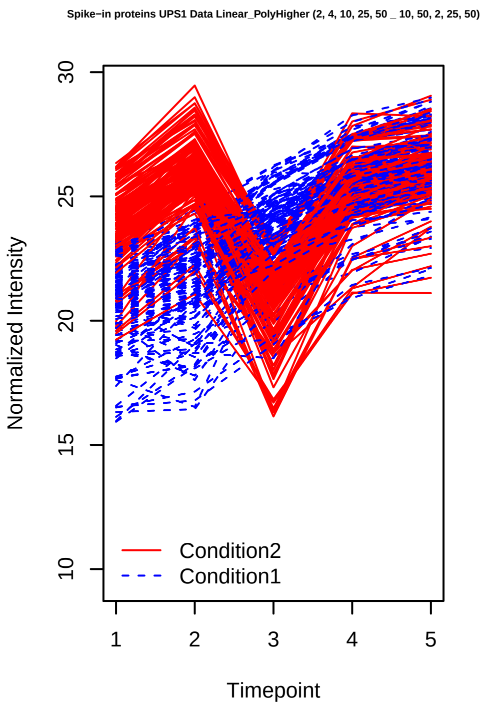
Spike−in proteins UPS1 Data Linear_PolyHigher (2, 4, 10, 25, 50 _ 10, 50, 2, 25, 50)
Normalized Intensity
Timepoint
Condition2
Condition1

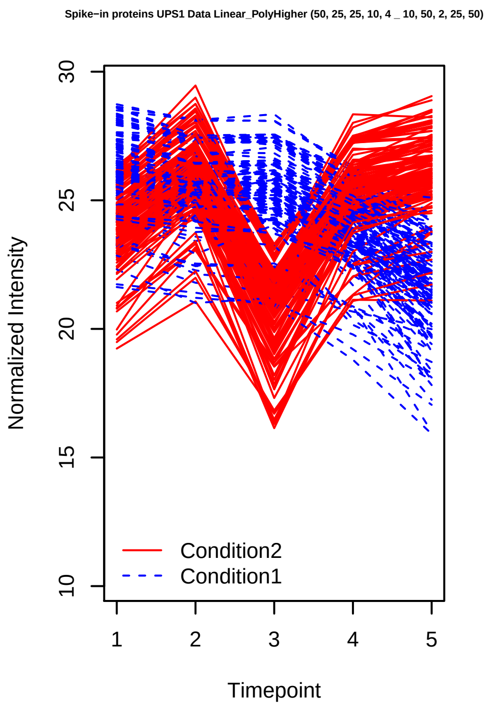
Spike−in proteins UPS1 Data Linear_PolyHigher (50, 25, 25, 10, 4 _ 10, 50, 2, 25, 50)
Normalized Intensity
Timepoint
Condition2
Condition1

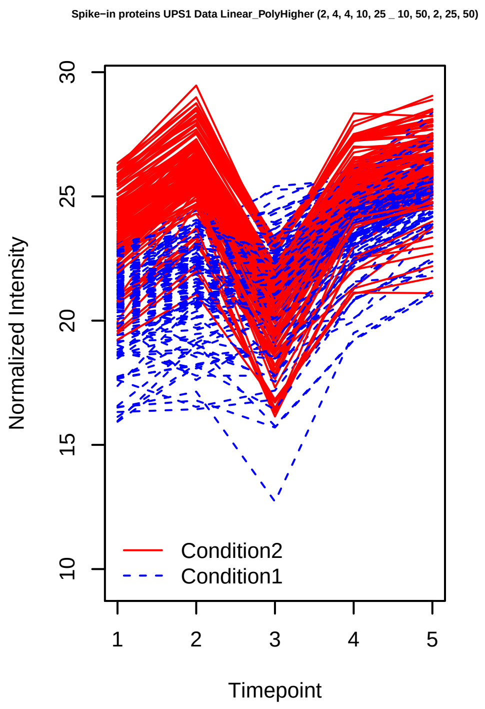
Spike−in proteins UPS1 Data Linear_PolyHigher (2, 4, 4, 10, 25 _ 10, 50, 2, 25, 50)
Normalized Intensity
Timepoint
Condition2
Condition1

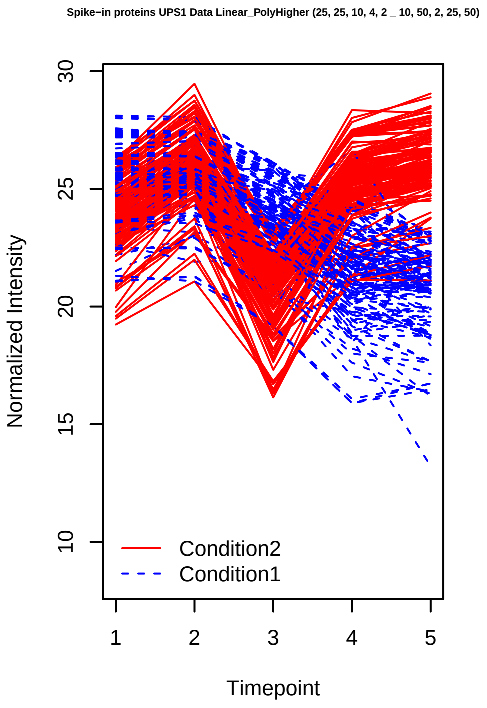
Spike−in proteins UPS1 Data Linear_PolyHigher (25, 25, 10, 4, 2 _ 10, 50, 2, 25, 50)
Normalized Intensity
Timepoint
Condition2
Condition1

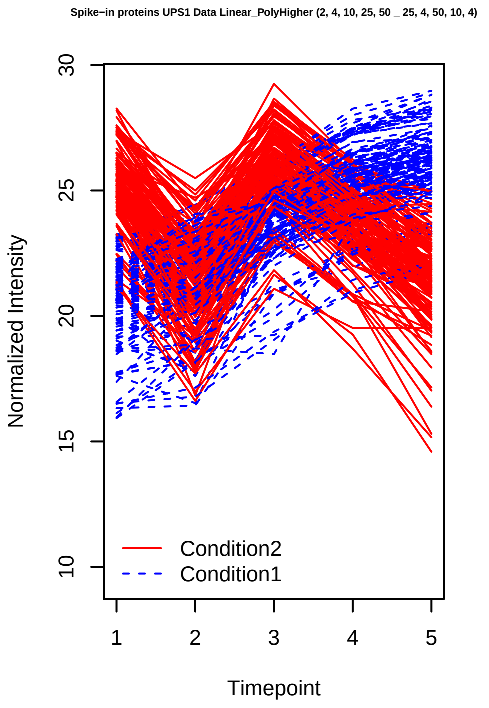
Spike−in proteins UPS1 Data Linear_PolyHigher (2, 4, 10, 25, 50 _ 25, 4, 50, 10, 4)
Normalized Intensity
Timepoint
Condition2
Condition1

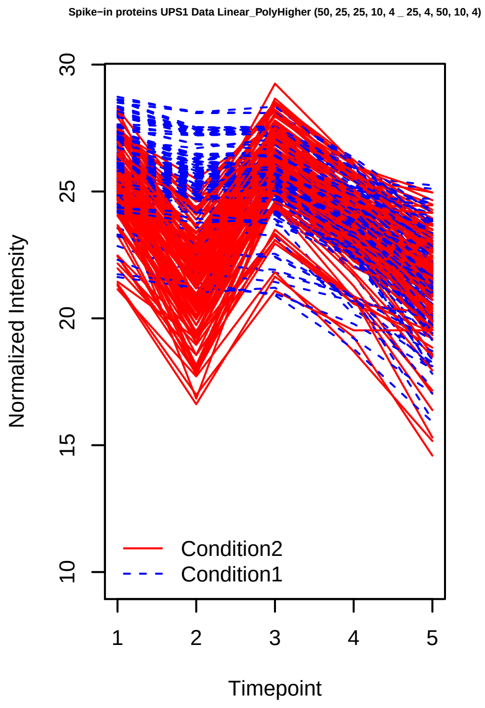
Spike−in proteins UPS1 Data Linear_PolyHigher (50, 25, 25, 10, 4 _ 25, 4, 50, 10, 4)
Normalized Intensity
Timepoint
Condition2
Condition1

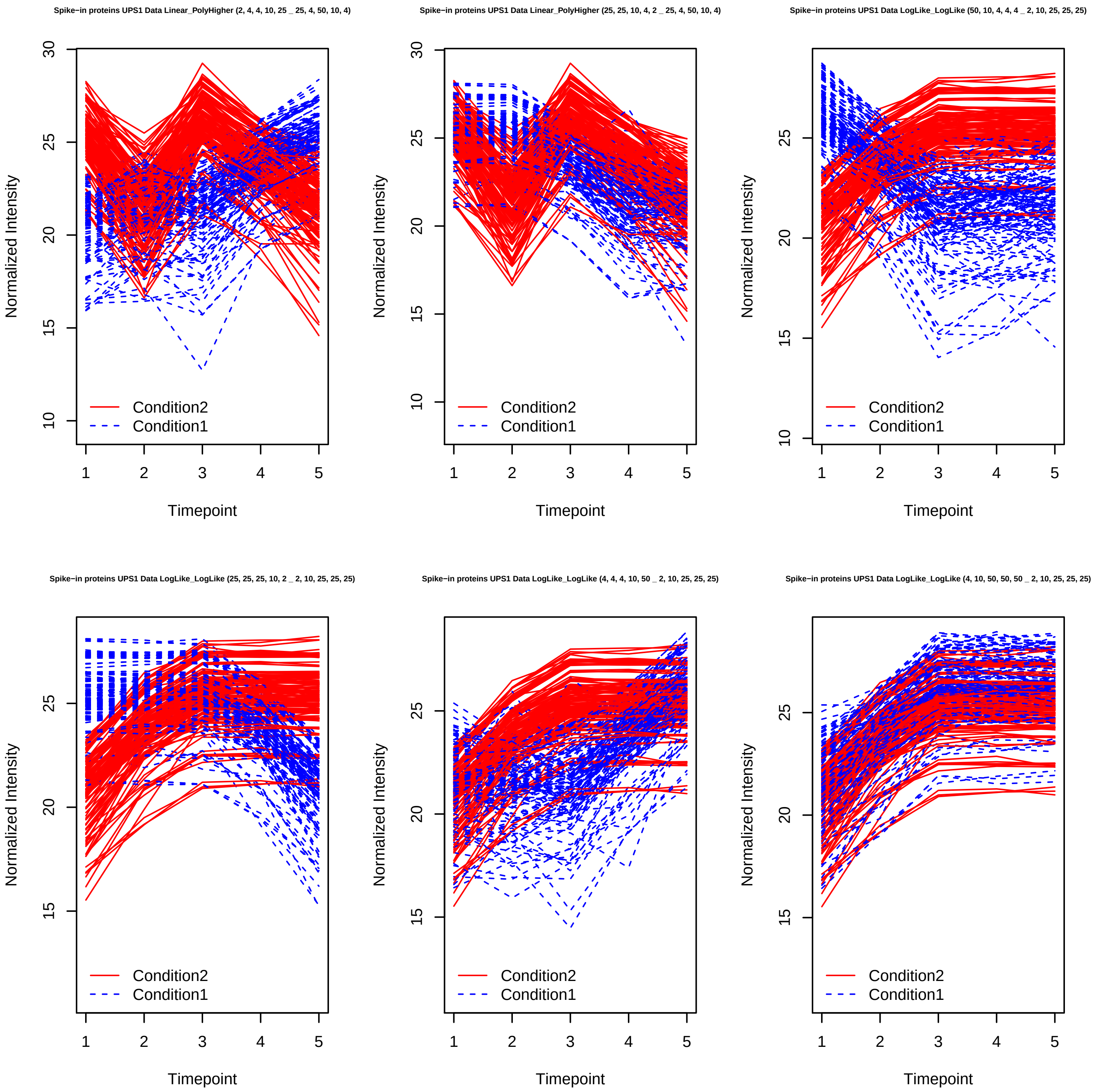

Spike−in proteins UPS1 Data Linear_PolyHigher (2, 4, 4, 10, 25 _ 25, 4, 50, 10, 4)
Spike−in proteins UPS1 Data Linear_PolyHigher (25, 25, 10, 4, 2 _ 25, 4, 50, 10, 4)
Spike−in proteins UPS1 Data LogLike_LogLike (50, 10, 4, 4, 4 _ 2, 10, 25, 25, 25)
Spike−in proteins UPS1 Data LogLike_LogLike (25, 25, 25, 10, 2 _ 2, 10, 25, 25, 25)
Spike−in proteins UPS1 Data LogLike_LogLike (4, 4, 4, 10, 50 _ 2, 10, 25, 25, 25)
Spike−in proteins UPS1 Data LogLike_LogLike (4, 10, 50, 50, 50 _ 2, 10, 25, 25, 25)
Normalized Intensity
Timepoint
Condition2
Condition1

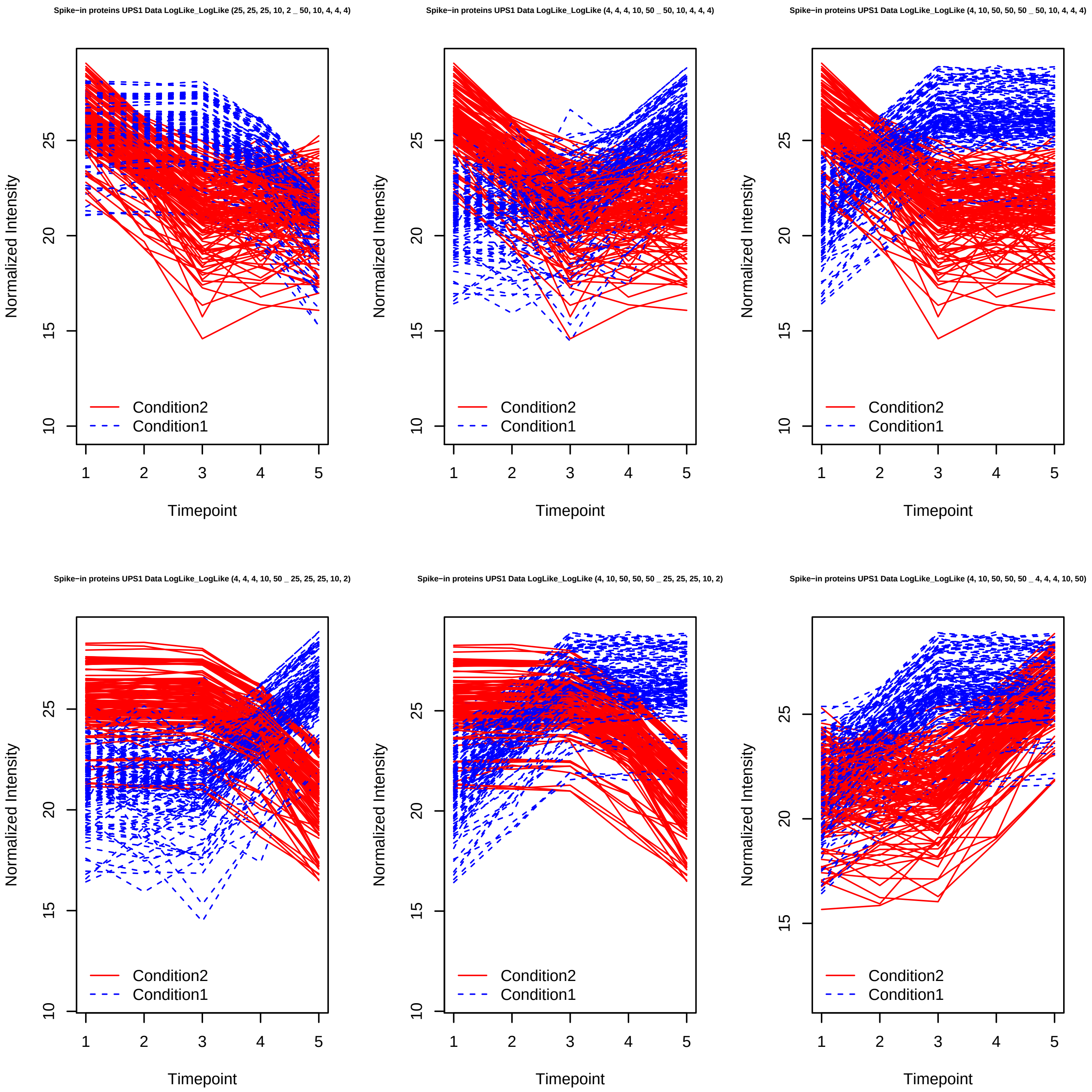

Spike−in proteins UPS1 Data LogLike_LogLike (25, 25, 25, 10, 2 _ 50, 10, 4, 4, 4)
Spike−in proteins UPS1 Data LogLike_LogLike (4, 4, 4, 10, 50 _ 50, 10, 4, 4, 4)
Spike−in proteins UPS1 Data LogLike_LogLike (4, 10, 50, 50, 50 _ 50, 10, 4, 4, 4)
Spike−in proteins UPS1 Data LogLike_LogLike (4, 4, 4, 10, 50 _ 25, 25, 25, 10, 2)
Spike−in proteins UPS1 Data LogLike_LogLike (4, 10, 50, 50, 50 _ 25, 25, 25, 10, 2)
Spike−in proteins UPS1 Data LogLike_LogLike (4, 10, 50, 50, 50 _ 4, 4, 4, 10, 50)
Normalized Intensity
Timepoint
Condition2
Condition1

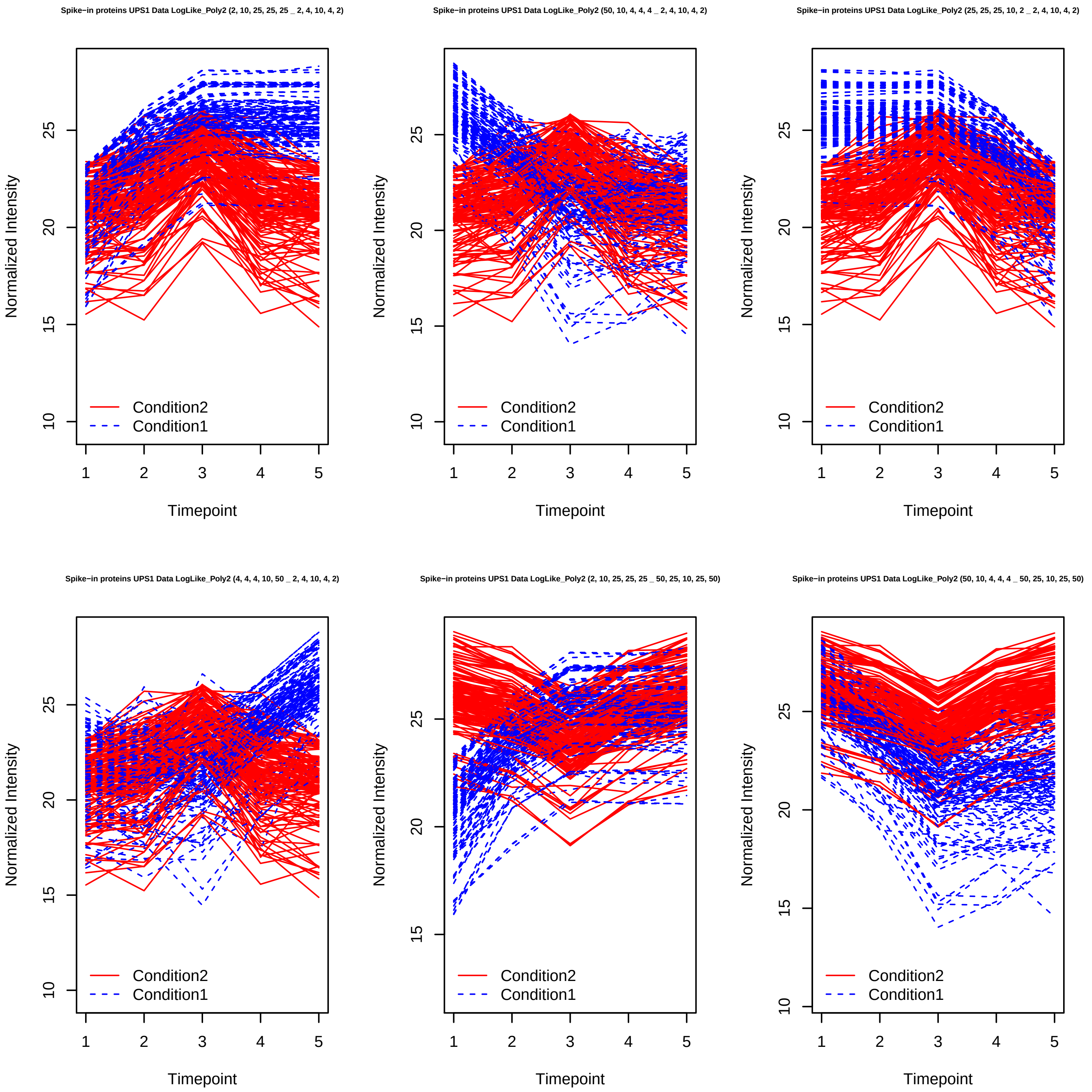
Spike−in proteins UPS1 Data LogLike_Poly2 (2, 10, 25, 25, 25 _ 2, 4, 10, 4, 2)
Spike−in proteins UPS1 Data LogLike_Poly2 (50, 10, 4, 4, 4 _ 2, 4, 10, 4, 2)
Spike−in proteins UPS1 Data LogLike_Poly2 (25, 25, 25, 10, 2 _ 2, 4, 10, 4, 2)
Spike−in proteins UPS1 Data LogLike_Poly2 (4, 4, 4, 10, 50 _ 2, 4, 10, 4, 2)
Spike−in proteins UPS1 Data LogLike_Poly2 (2, 10, 25, 25, 25 _ 50, 25, 10, 25, 50)
Spike−in proteins UPS1 Data LogLike_Poly2 (50, 10, 4, 4, 4 _ 50, 25, 10, 25, 50)
Normalized Intensity
Timepoint
Condition2
Condition1
10
15
20
25
1
2
3
4
5

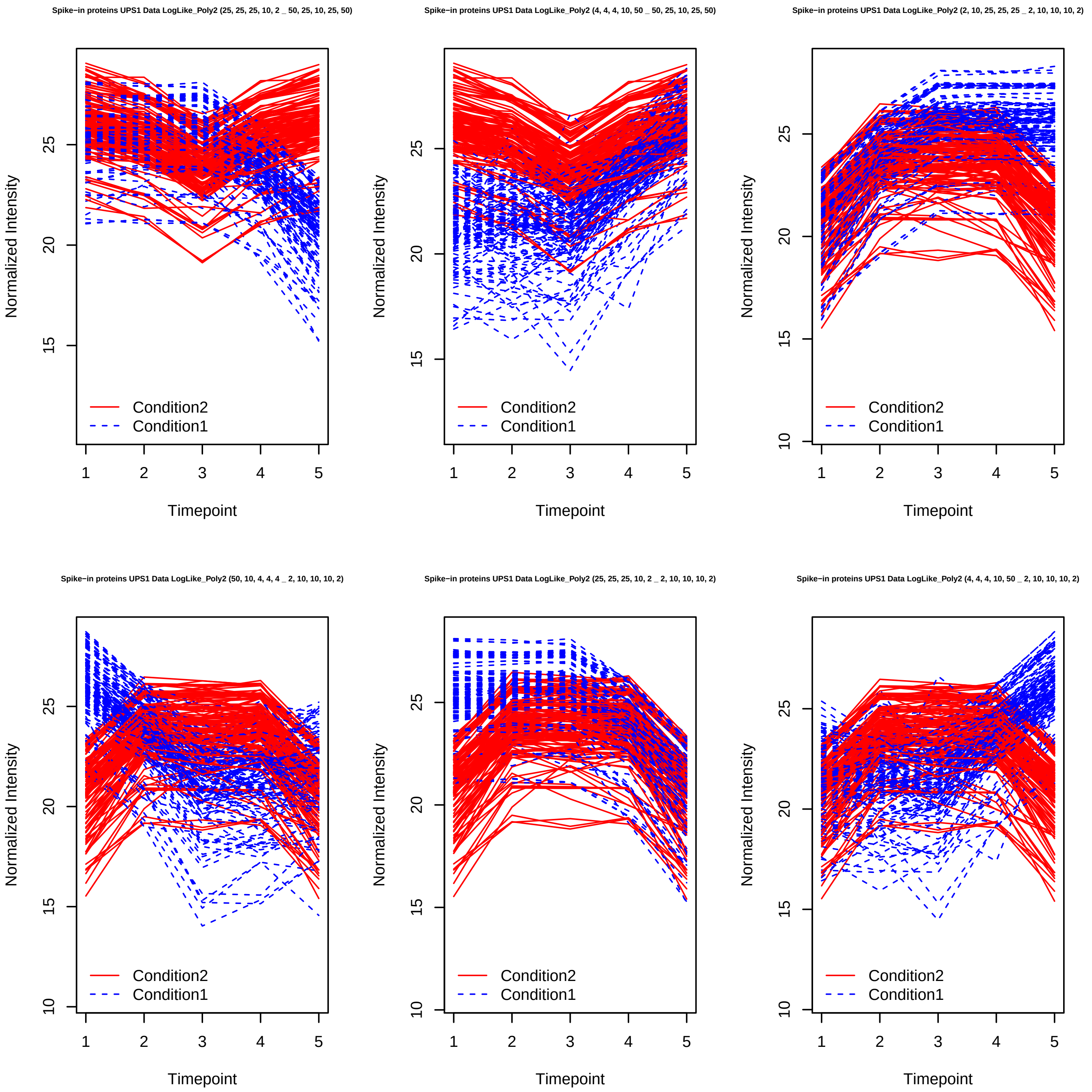
Spike−in proteins UPS1 Data LogLike_Poly2 (25, 25, 25, 10, 2 _ 50, 25, 10, 25, 50)
Spike−in proteins UPS1 Data LogLike_Poly2 (4, 4, 4, 10, 50 _ 50, 25, 10, 25, 50)
Spike−in proteins UPS1 Data LogLike_Poly2 (2, 10, 25, 25, 25 _ 2, 10, 10, 10, 2)
Spike−in proteins UPS1 Data LogLike_Poly2 (50, 10, 4, 4, 4 _ 2, 10, 10, 10, 2)
Spike−in proteins UPS1 Data LogLike_Poly2 (25, 25, 25, 10, 2 _ 2, 10, 10, 10, 2)
Spike−in proteins UPS1 Data LogLike_Poly2 (4, 4, 4, 10, 50 _ 2, 10, 10, 10, 2)
Normalized Intensity
Timepoint
Condition2
Condition1

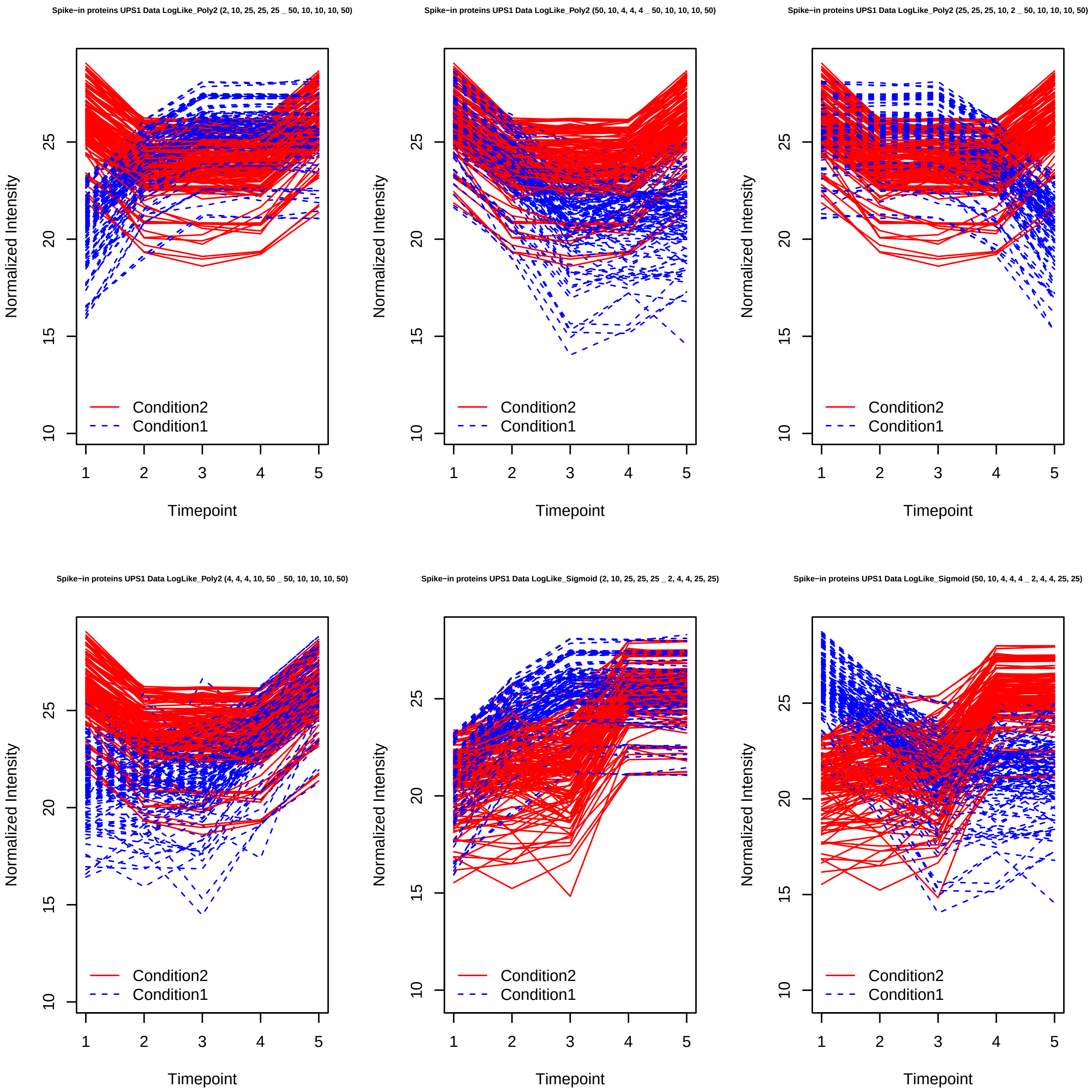
Spike−in proteins UPS1 Data LogLike_Poly2 (2, 10, 25, 25, 25 _ 50, 10, 10, 10, 50)
Spike−in proteins UPS1 Data LogLike_Poly2 (50, 10, 4, 4, 4 _ 50, 10, 10, 10, 50)
Spike−in proteins UPS1 Data LogLike_Poly2 (25, 25, 25, 10, 2 _ 50, 10, 10, 10, 50)
Spike−in proteins UPS1 Data LogLike_Poly2 (4, 4, 4, 10, 50 _ 50, 10, 10, 10, 50)
Spike−in proteins UPS1 Data LogLike_Sigmoid (2, 10, 25, 25, 25 _ 2, 4, 4, 25, 25)
Spike−in proteins UPS1 Data LogLike_Sigmoid (50, 10, 4, 4, 4 _ 2, 4, 4, 25, 25)
Normalized Intensity
Timepoint
Condition2
Condition1

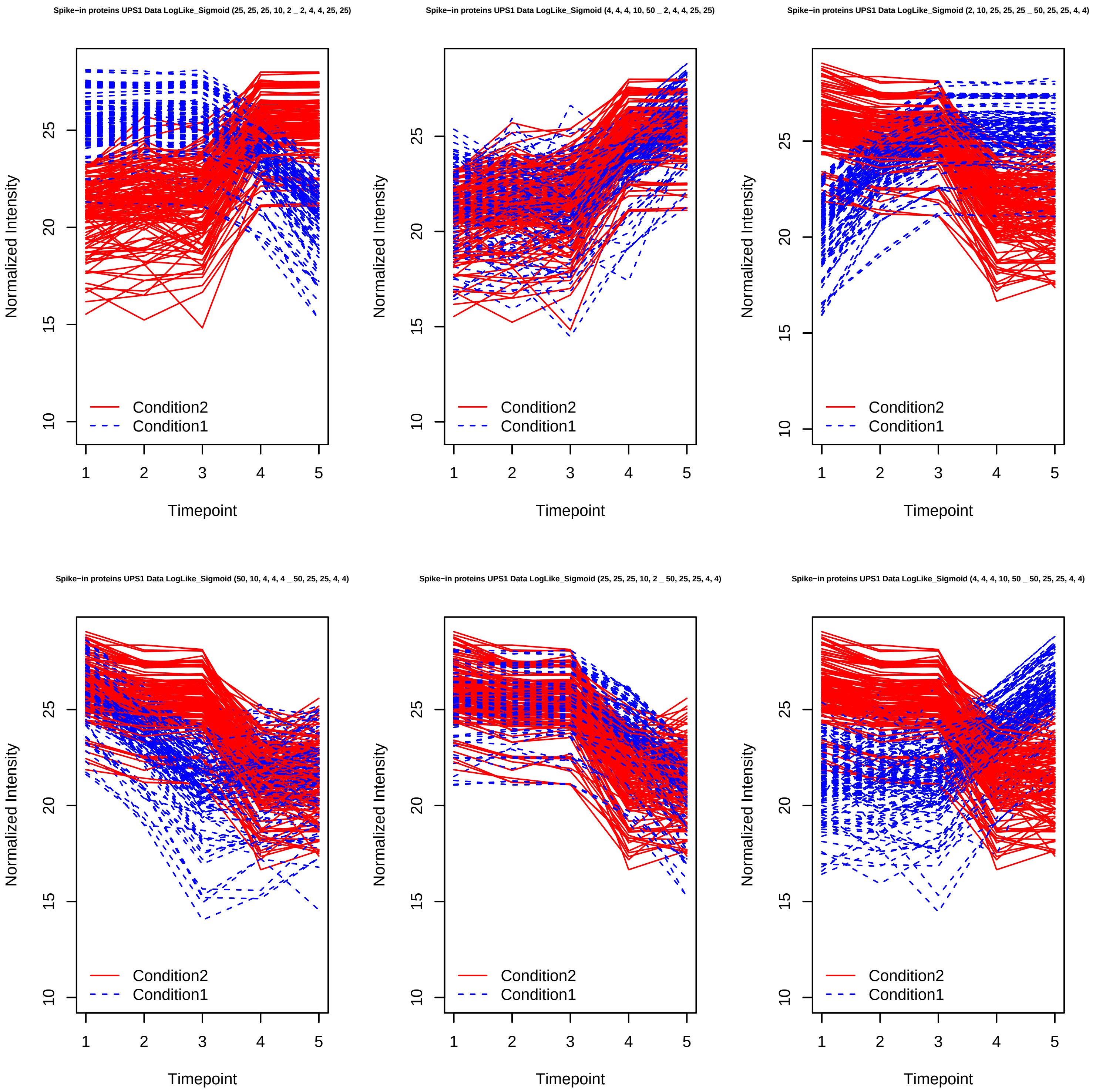

Spike−in proteins UPS1 Data LogLike_Sigmoid (25, 25, 25, 10, 2 _ 2, 4, 4, 25, 25)
Spike−in proteins UPS1 Data LogLike_Sigmoid (4, 4, 4, 10, 50 _ 2, 4, 4, 25, 25)
Spike−in proteins UPS1 Data LogLike_Sigmoid (2, 10, 25, 25, 25 _ 50, 25, 25, 4, 4)
Spike−in proteins UPS1 Data LogLike_Sigmoid (50, 10, 4, 4, 4 _ 50, 25, 25, 4, 4)
Spike−in proteins UPS1 Data LogLike_Sigmoid (25, 25, 25, 10, 2 _ 50, 25, 25, 4, 4)
Spike−in proteins UPS1 Data LogLike_Sigmoid (4, 4, 4, 10, 50 _ 50, 25, 25, 4, 4)
Normalized Intensity
Timepoint
10
15
20
25
1
2
3
4
5
Condition2
Condition1

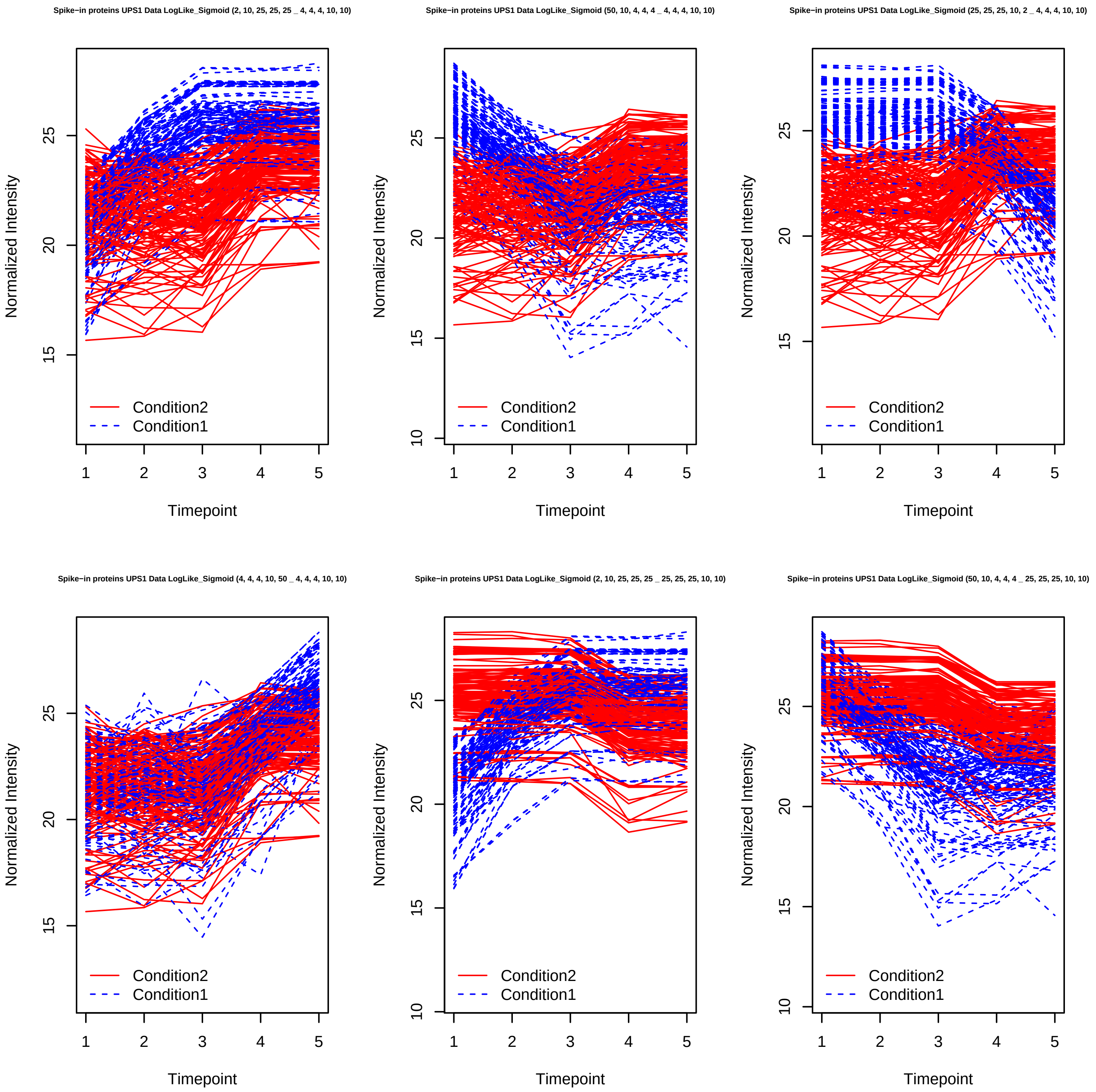
Spike−in proteins UPS1 Data LogLike_Sigmoid (2, 10, 25, 25, 25 _ 4, 4, 4, 10, 10)
Spike−in proteins UPS1 Data LogLike_Sigmoid (50, 10, 4, 4, 4 _ 4, 4, 4, 10, 10)
Spike−in proteins UPS1 Data LogLike_Sigmoid (25, 25, 25, 10, 2 _ 4, 4, 4, 10, 10)
Spike−in proteins UPS1 Data LogLike_Sigmoid (4, 4, 4, 10, 50 _ 4, 4, 4, 10, 10)
Spike−in proteins UPS1 Data LogLike_Sigmoid (2, 10, 25, 25, 25 _ 25, 25, 25, 10, 10)
Spike−in proteins UPS1 Data LogLike_Sigmoid (50, 10, 4, 4, 4 _ 25, 25, 25, 10, 10)
Normalized Intensity
Timepoint
Condition2
Condition1

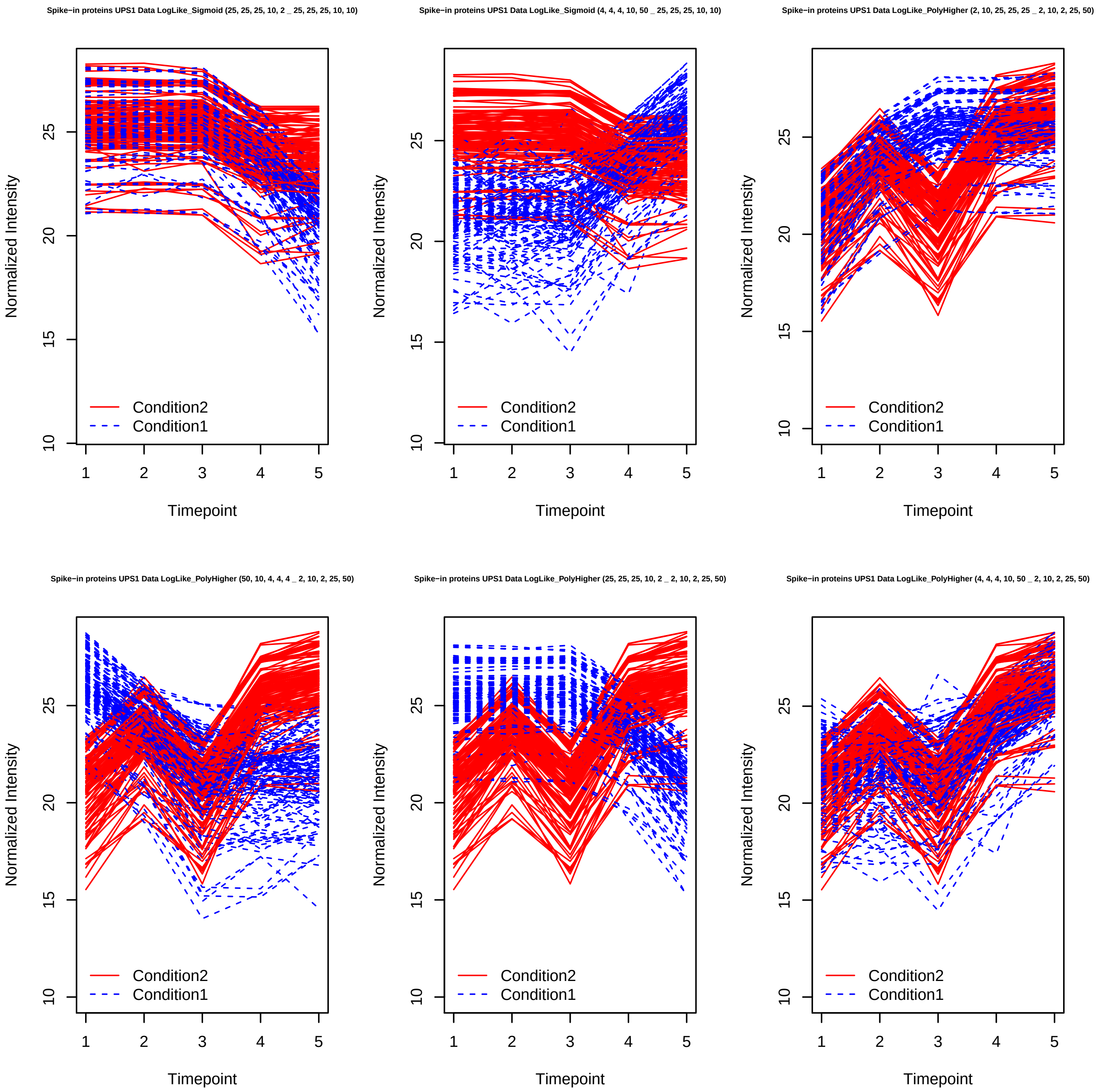
Spike−in proteins UPS1 Data LogLike_Sigmoid (25, 25, 25, 10, 2 _ 25, 25, 25, 10, 10)
Spike−in proteins UPS1 Data LogLike_Sigmoid (4, 4, 4, 10, 50 _ 25, 25, 25, 10, 10)
Spike−in proteins UPS1 Data LogLike_PolyHigher (2, 10, 25, 25, 25 _ 2, 10, 2, 25, 50)
Spike−in proteins UPS1 Data LogLike_PolyHigher (50, 10, 4, 4, 4 _ 2, 10, 2, 25, 50)
Spike−in proteins UPS1 Data LogLike_PolyHigher (25, 25, 25, 10, 2 _ 2, 10, 2, 25, 50)
Spike−in proteins UPS1 Data LogLike_PolyHigher (4, 4, 4, 10, 50 _ 2, 10, 2, 25, 50)
Normalized Intensity
Timepoint
Condition2
Condition1

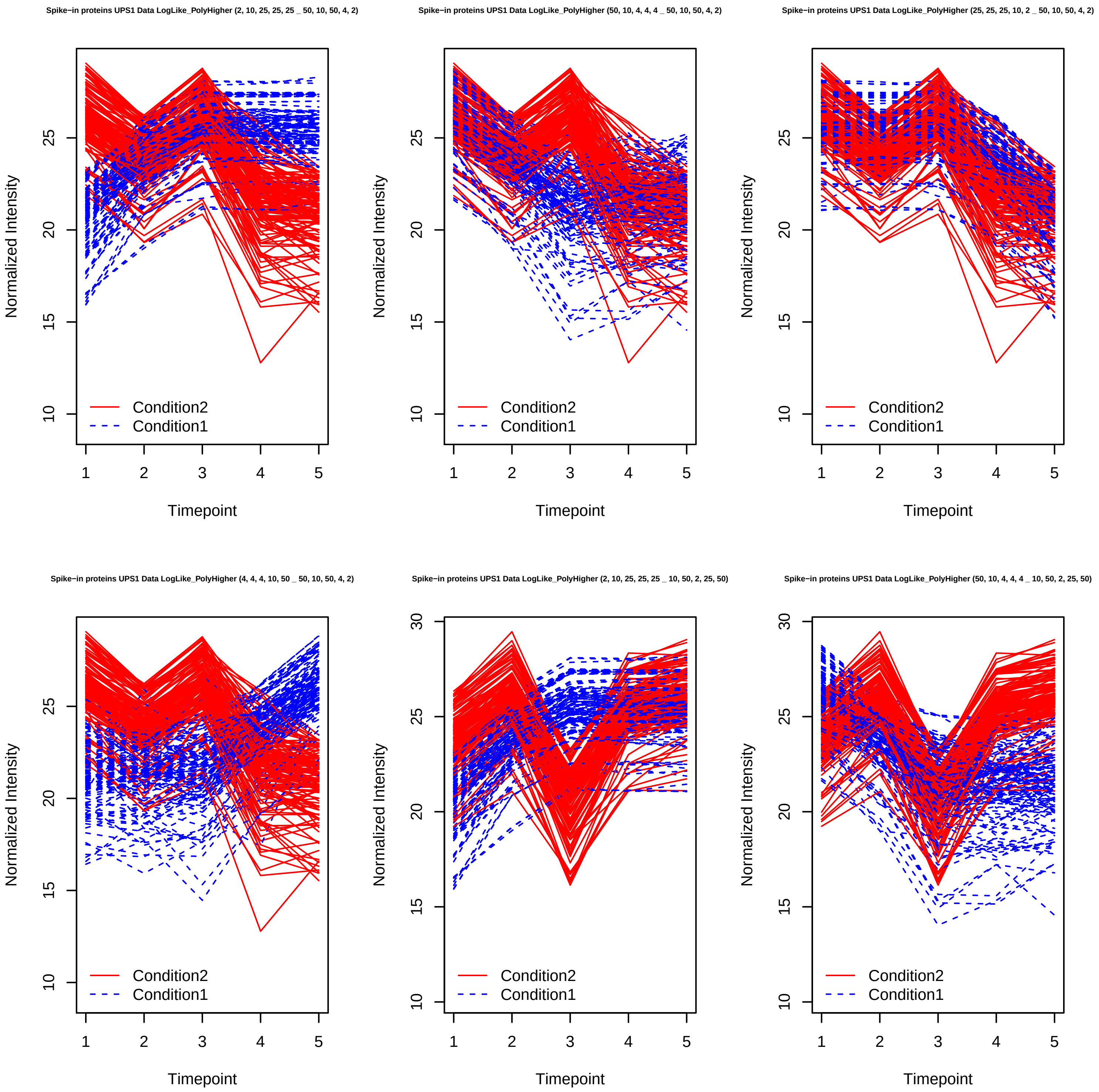
Spike−in proteins UPS1 Data LogLike_PolyHigher (2, 10, 25, 25, 25 _ 50, 10, 50, 4, 2)
Spike−in proteins UPS1 Data LogLike_PolyHigher (50, 10, 4, 4, 4 _ 50, 10, 50, 4, 2)
Spike−in proteins UPS1 Data LogLike_PolyHigher (25, 25, 25, 10, 2 _ 50, 10, 50, 4, 2)
Spike−in proteins UPS1 Data LogLike_PolyHigher (4, 4, 4, 10, 50 _ 50, 10, 50, 4, 2)
Spike−in proteins UPS1 Data LogLike_PolyHigher (2, 10, 25, 25, 25 _ 10, 50, 2, 25, 50)
Spike−in proteins UPS1 Data LogLike_PolyHigher (50, 10, 4, 4, 4 _ 10, 50, 2, 25, 50)
Normalized Intensity
Timepoint
Condition2
Condition1

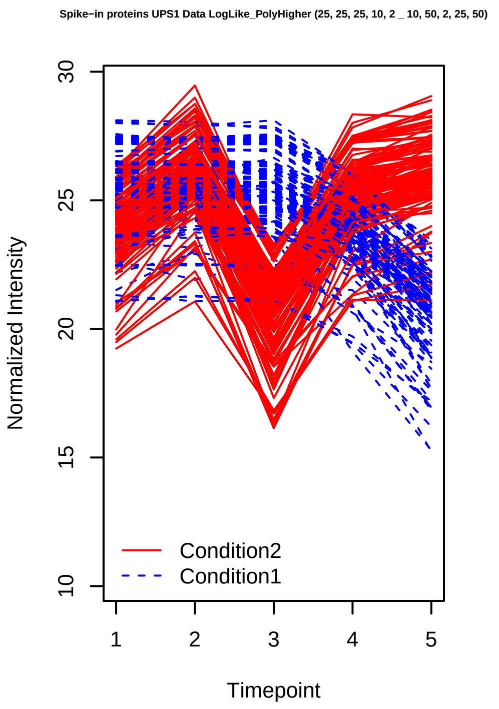
Spike−in proteins UPS1 Data LogLike_PolyHigher (25, 25, 25, 10, 2 _ 10, 50, 2, 25, 50)
Normalized Intensity
Timepoint
Condition2
Condition1

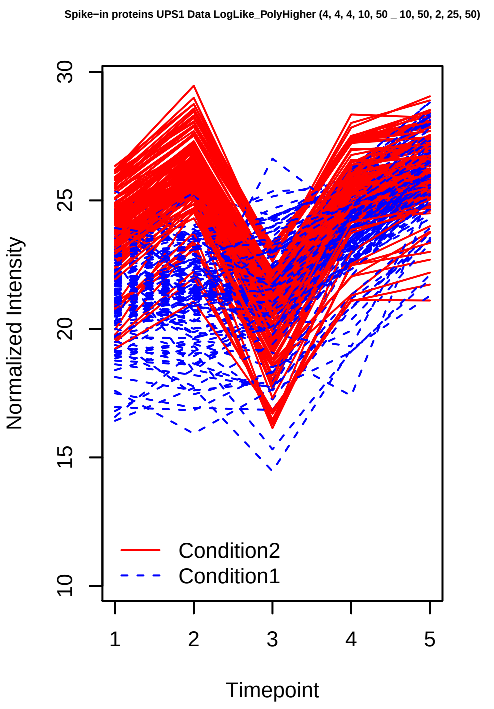
Spike−in proteins UPS1 Data LogLike_PolyHigher (4, 4, 4, 10, 50 _ 10, 50, 2, 25, 50)
Normalized Intensity
Timepoint
Condition2
Condition1

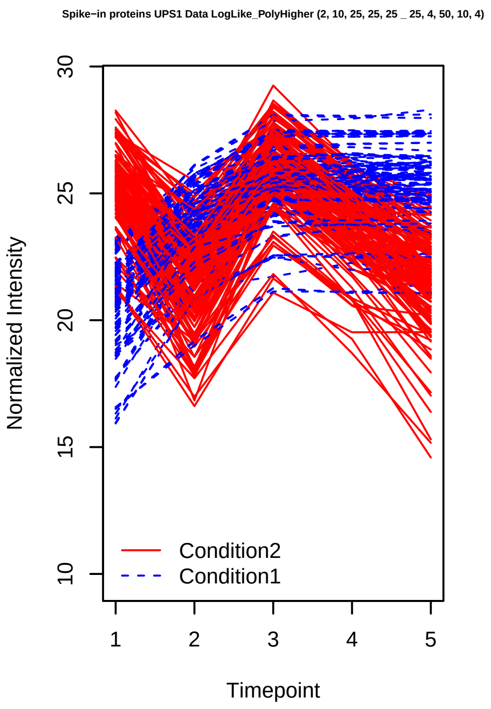
Spike−in proteins UPS1 Data LogLike_PolyHigher (2, 10, 25, 25, 25 _ 25, 4, 50, 10, 4)
Normalized Intensity
Timepoint
Condition2
Condition1

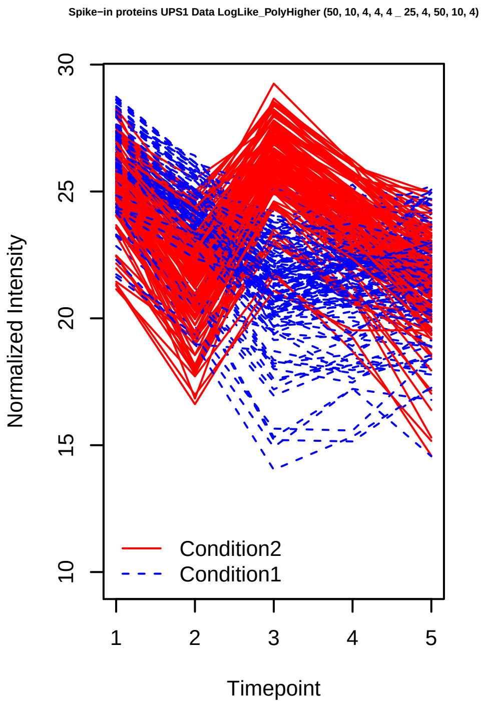
Spike−in proteins UPS1 Data LogLike_PolyHigher (50, 10, 4, 4, 4 _ 25, 4, 50, 10, 4)
Normalized Intensity
Timepoint
Condition2
Condition1

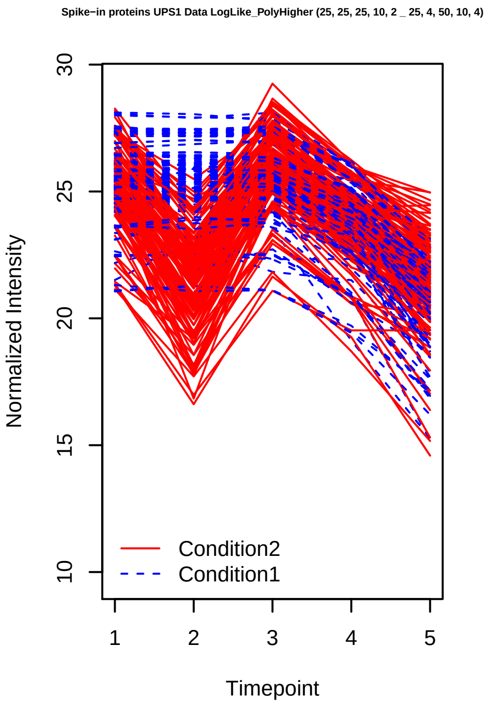
Spike−in proteins UPS1 Data LogLike_PolyHigher (25, 25, 25, 10, 2 _ 25, 4, 50, 10, 4)
Normalized Intensity
Timepoint
Condition2
Condition1

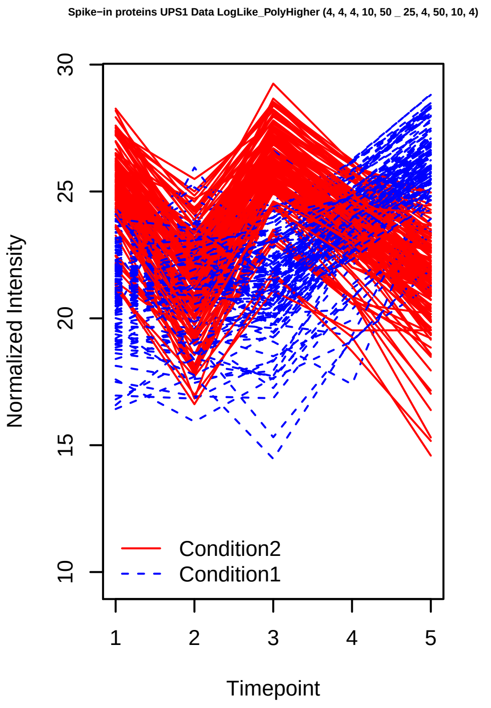
Spike−in proteins UPS1 Data LogLike_PolyHigher (4, 4, 4, 10, 50 _ 25, 4, 50, 10, 4)
Normalized Intensity
Timepoint
Condition2
Condition1

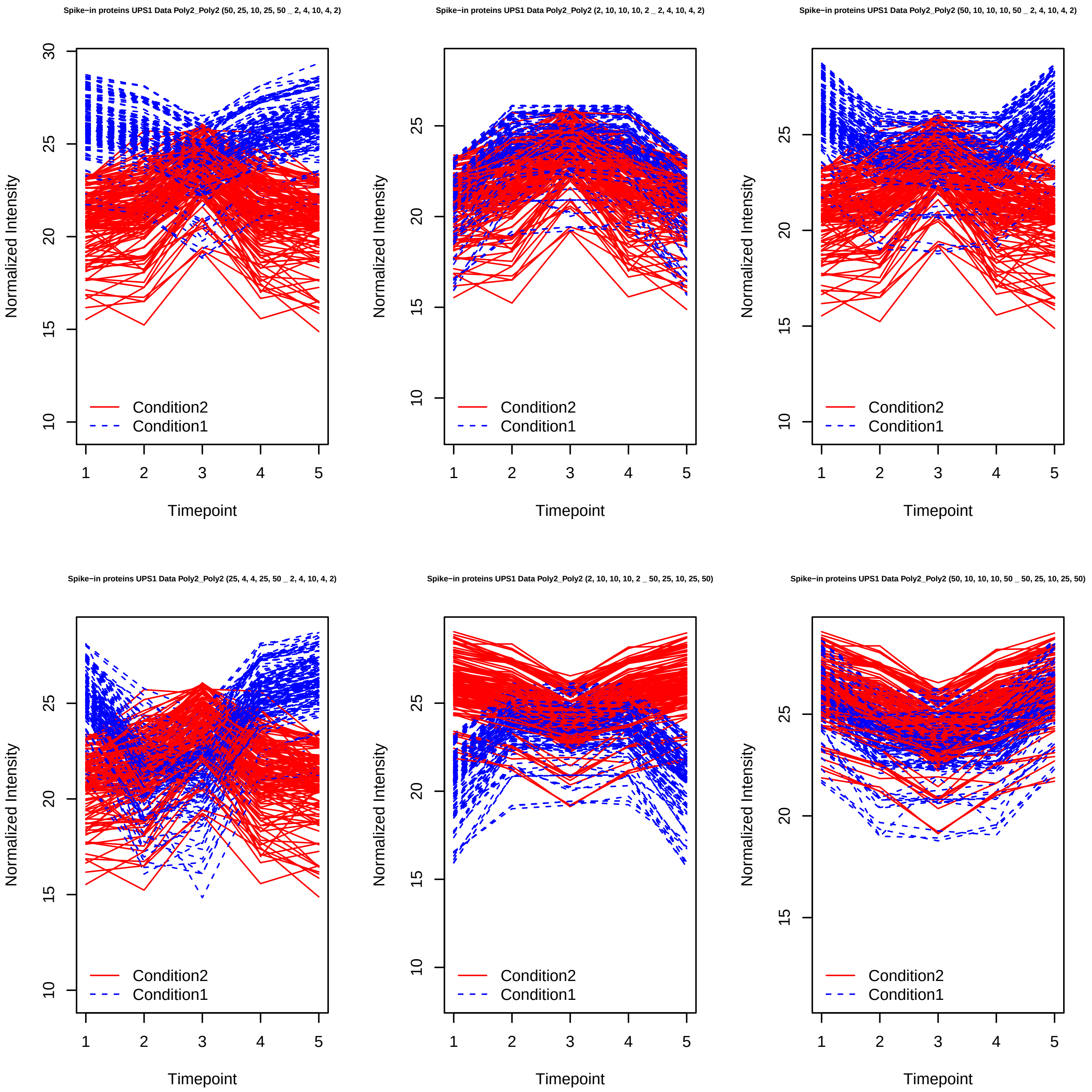
Spike−in proteins UPS1 Data Poly2_Poly2 (50, 25, 10, 25, 50 _ 2, 4, 10, 4, 2)
Spike−in proteins UPS1 Data Poly2_Poly2 (2, 10, 10, 10, 2 _ 2, 4, 10, 4, 2)
Spike−in proteins UPS1 Data Poly2_Poly2 (50, 10, 10, 10, 50 _ 2, 4, 10, 4, 2)
Spike−in proteins UPS1 Data Poly2_Poly2 (25, 4, 4, 25, 50 _ 2, 4, 10, 4, 2)
Spike−in proteins UPS1 Data Poly2_Poly2 (2, 10, 10, 10, 2 _ 50, 25, 10, 25, 50)
Spike−in proteins UPS1 Data Poly2_Poly2 (50, 10, 10, 10, 50 _ 50, 25, 10, 25, 50)
Normalized Intensity
Timepoint
Condition2
Condition1

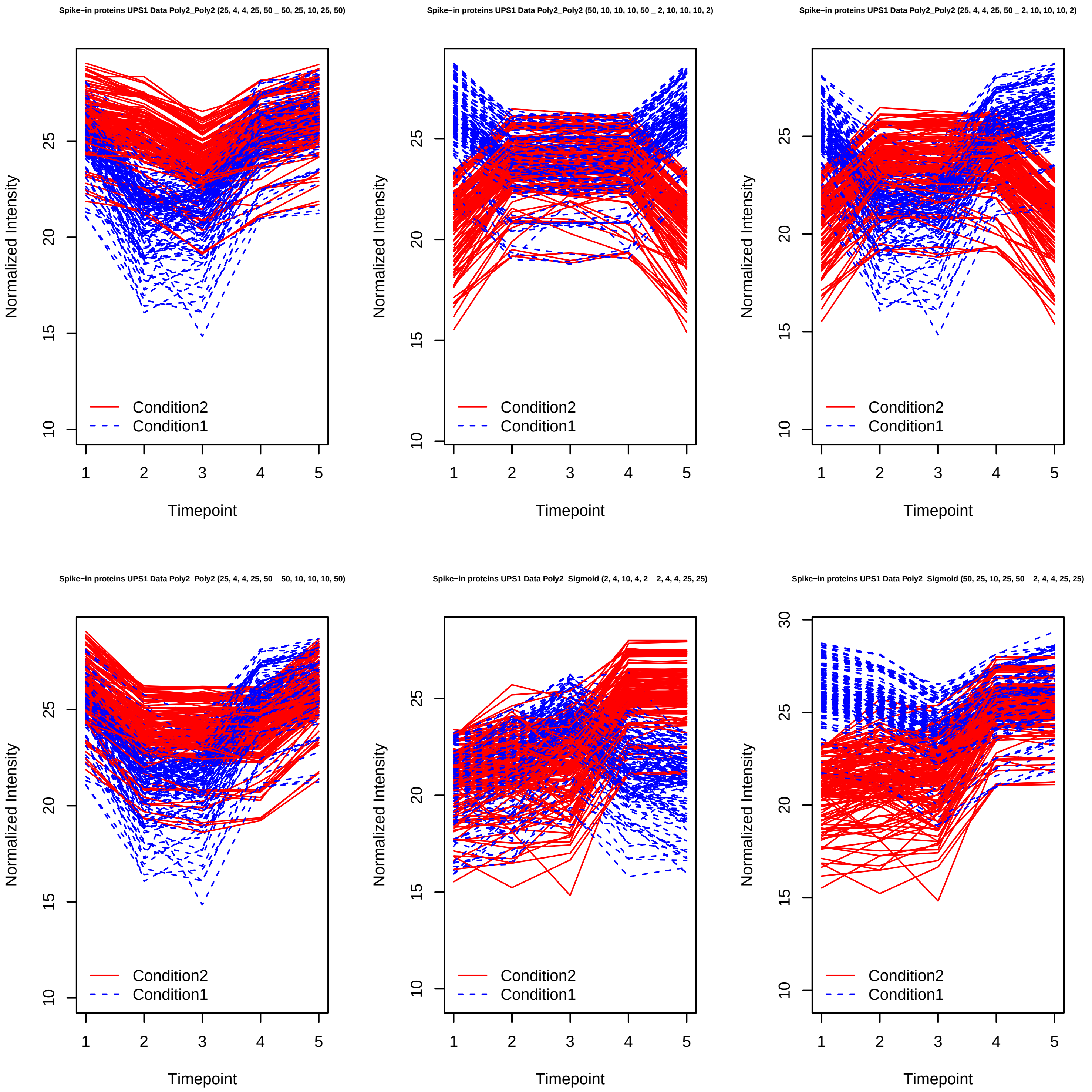

Spike−in proteins UPS1 Data Poly2_Poly2 (25, 4, 4, 25, 50 _ 50, 25, 10, 25, 50)
Spike−in proteins UPS1 Data Poly2_Poly2 (50, 10, 10, 10, 50 _ 2, 10, 10, 10, 2)
Spike−in proteins UPS1 Data Poly2_Poly2 (25, 4, 4, 25, 50 _ 2, 10, 10, 10, 2)
Spike−in proteins UPS1 Data Poly2_Poly2 (25, 4, 4, 25, 50 _ 50, 10, 10, 10, 50)
Spike−in proteins UPS1 Data Poly2_Sigmoid (2, 4, 10, 4, 2 _ 2, 4, 4, 25, 25)
Spike−in proteins UPS1 Data Poly2_Sigmoid (50, 25, 10, 25, 50 _ 2, 4, 4, 25, 25)
Normalized Intensity
Timepoint
Condition2
Condition1

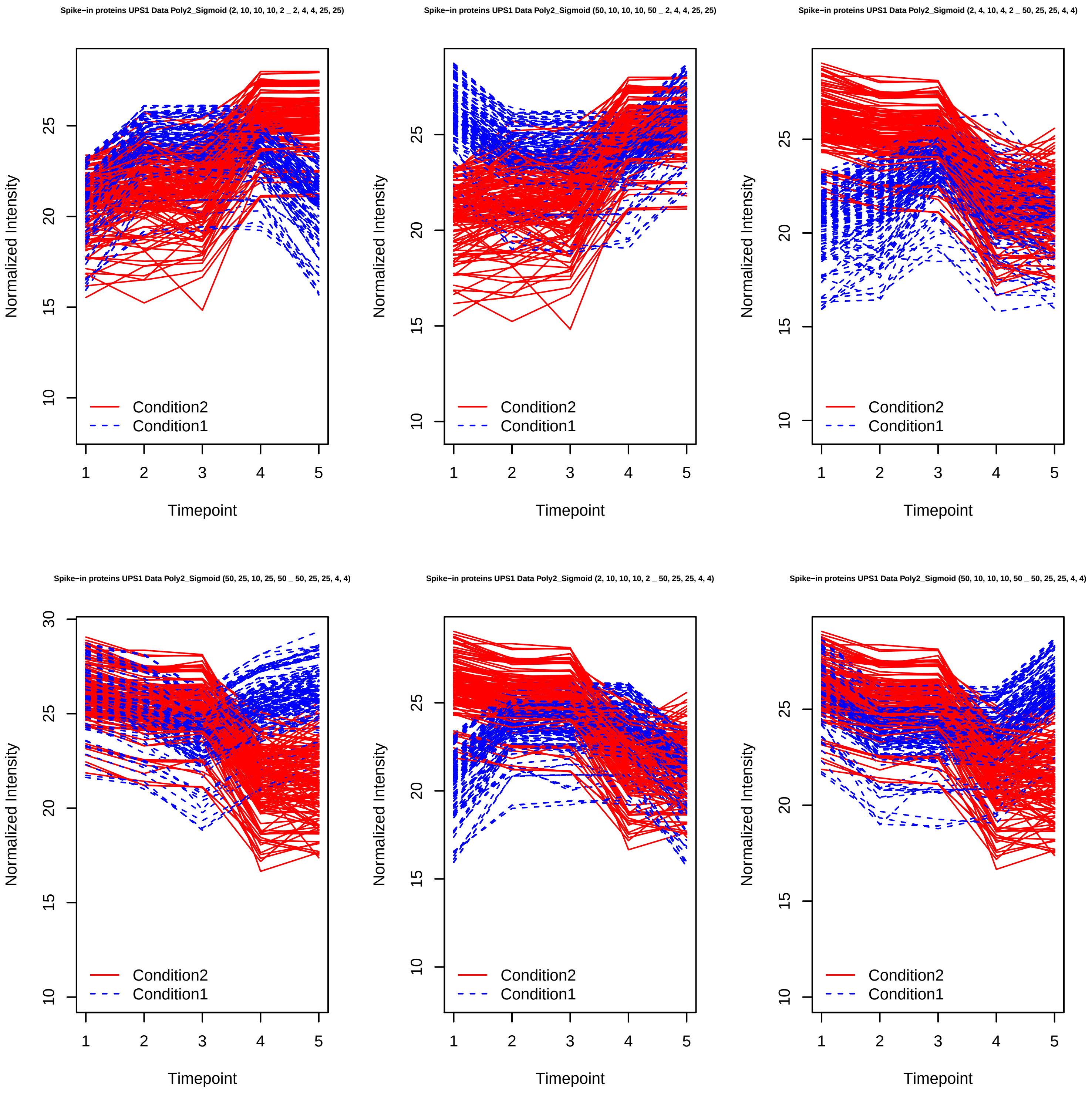
Spike−in proteins UPS1 Data Poly2_Sigmoid (2, 10, 10, 10, 2 _ 2, 4, 4, 25, 25)
Normalized Intensity
10
15
20
25
1
2
3
4
5
Timepoint
Condition2
Condition1
Spike−in proteins UPS1 Data Poly2_Sigmoid (50, 10, 10, 10, 50 _ 2, 4, 4, 25, 25)
Normalized Intensity
10
15
20
25
1
2
3
4
5
Timepoint
Condition2
Condition1
Spike−in proteins UPS1 Data Poly2_Sigmoid (2, 4, 10, 4, 2 _ 50, 25, 25, 4, 4)
Normalized Intensity
10
15
20
25
1
2
3
4
5
Timepoint
Condition2
Condition1
Spike−in proteins UPS1 Data Poly2_Sigmoid (50, 25, 10, 25, 50 _ 50, 25, 25, 4, 4)
Normalized Intensity
10
15
20
25
30
1
2
3
4
5
Timepoint
Condition2
Condition1
Spike−in proteins UPS1 Data Poly2_Sigmoid (2, 10, 10, 10, 2 _ 50, 25, 25, 4, 4)
Normalized Intensity
10
15
20
25
1
2
3
4
5
Timepoint
Condition2
Condition1
Spike−in proteins UPS1 Data Poly2_Sigmoid (50, 10, 10, 10, 50 _ 50, 25, 25, 4, 4)
Normalized Intensity
10
15
20
25
1
2
3
4
5
Timepoint
Condition2
Condition1

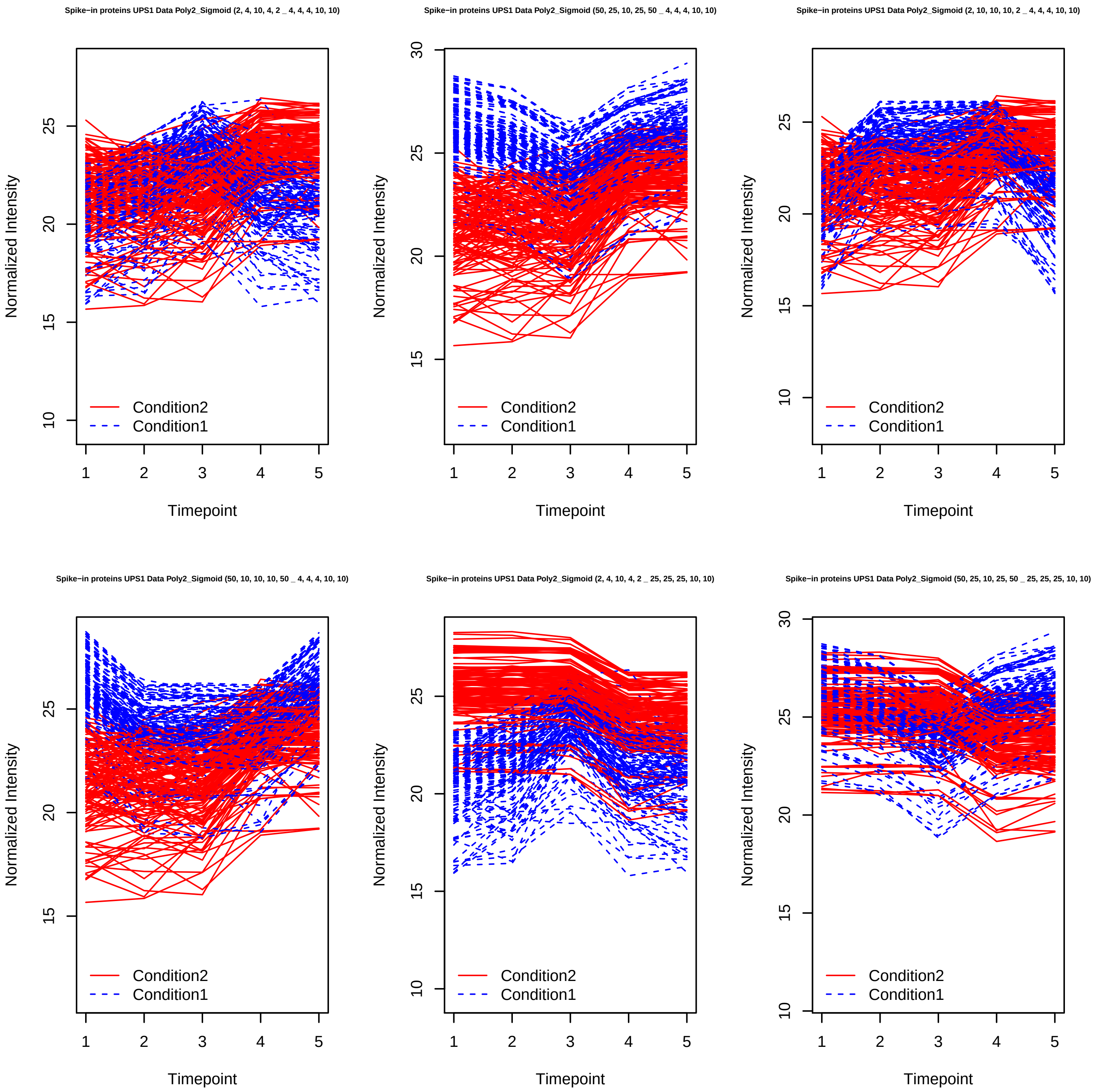

Spike−in proteins UPS1 Data Poly2_Sigmoid (2, 4, 10, 4, 2 _ 4, 4, 4, 10, 10)
Normalized Intensity
Timepoint
Condition2
Condition1
Spike−in proteins UPS1 Data Poly2_Sigmoid (50, 25, 10, 25, 50 _ 4, 4, 4, 10, 10)
Normalized Intensity
Timepoint
Condition2
Condition1
Spike−in proteins UPS1 Data Poly2_Sigmoid (2, 10, 10, 10, 2 _ 4, 4, 4, 10, 10)
Normalized Intensity
Timepoint
Condition2
Condition1
Spike−in proteins UPS1 Data Poly2_Sigmoid (50, 10, 10, 10, 50 _ 4, 4, 4, 10, 10)
Normalized Intensity
Timepoint
Condition2
Condition1
Spike−in proteins UPS1 Data Poly2_Sigmoid (2, 4, 10, 4, 2 _ 25, 25, 25, 10, 10)
Normalized Intensity
Timepoint
Condition2
Condition1
Spike−in proteins UPS1 Data Poly2_Sigmoid (50, 25, 10, 25, 50 _ 25, 25, 25, 10, 10)
Normalized Intensity
Timepoint
Condition2
Condition1

Spike−in proteins UPS1 Data Poly2_Sigmoid (2, 10, 10, 10, 2 _ 25, 25, 25, 10, 10)

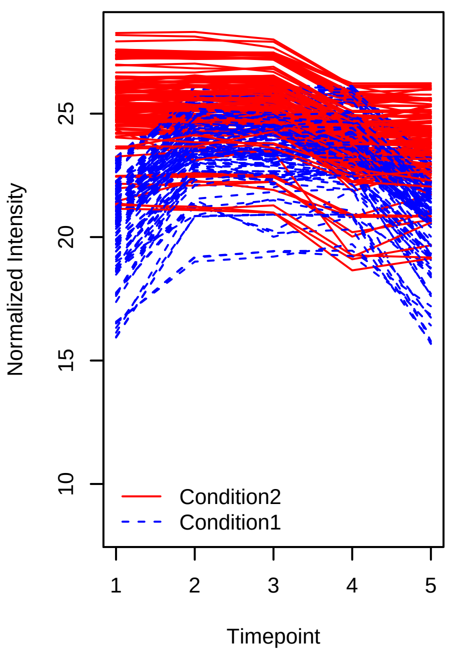


Spike−in proteins UPS1 Data Poly2_Sigmoid (50, 10, 10, 10, 50 _ 25, 25, 25, 10, 10)

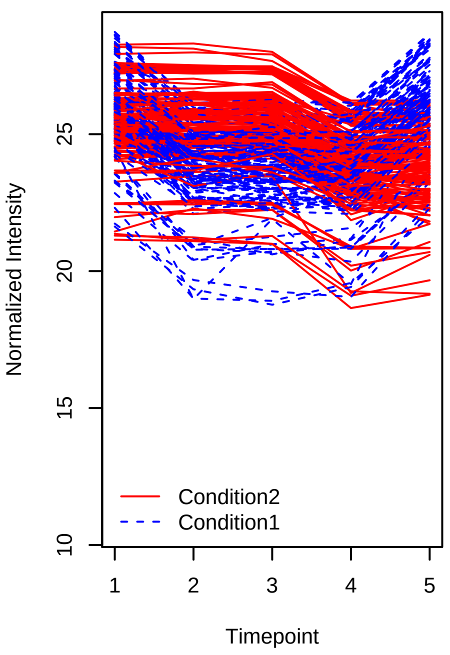


Spike−in proteins UPS1 Data Poly2_PolyHigher (2, 4, 10, 4, 2 _ 2, 10, 2, 25, 50)

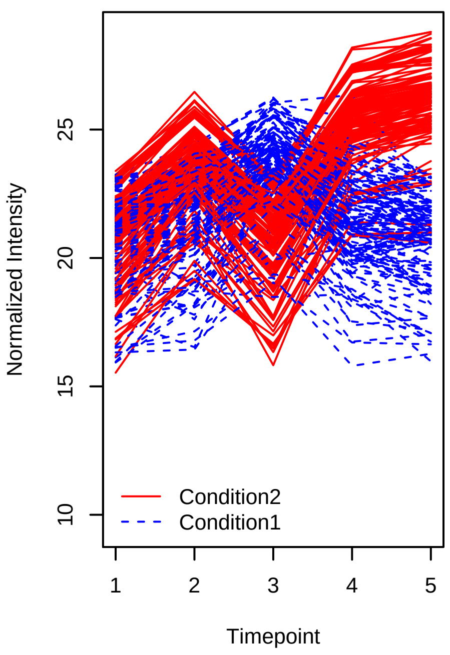


Spike−in proteins UPS1 Data Poly2_PolyHigher (50, 25, 10, 25, 50 _ 2, 10, 2, 25, 50)

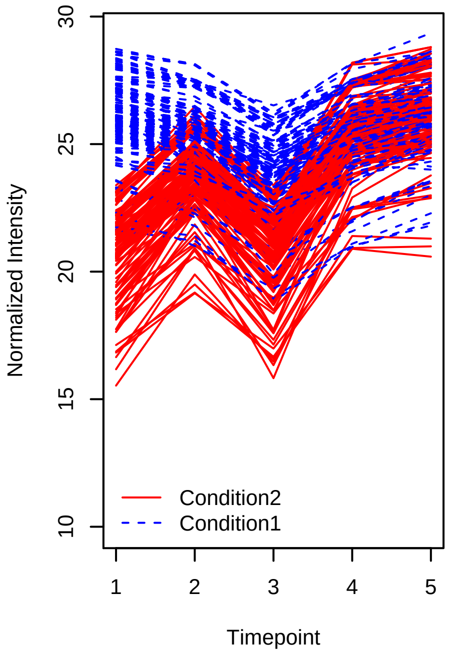


Spike−in proteins UPS1 Data Poly2_PolyHigher (2, 10, 10, 10, 2 _ 2, 10, 2, 25, 50)

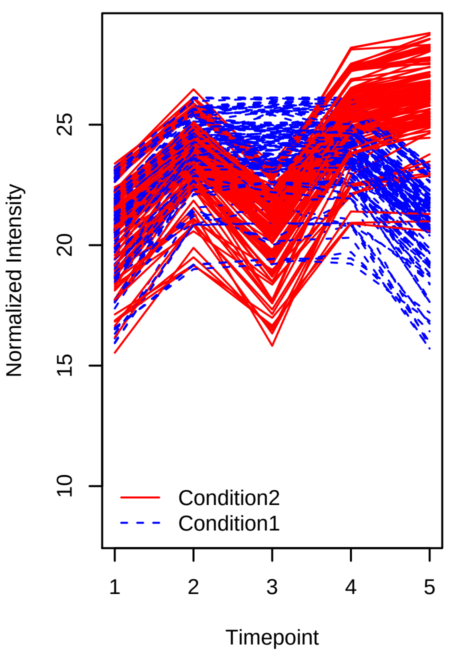


Spike−in proteins UPS1 Data Poly2_PolyHigher (50, 10, 10, 10, 50 _ 2, 10, 2, 25, 50)

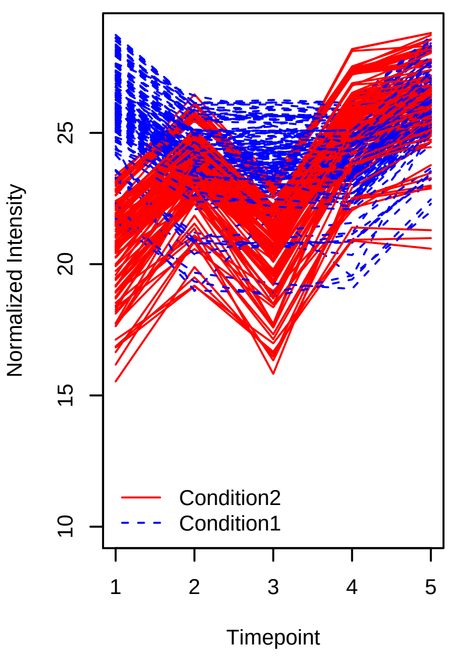

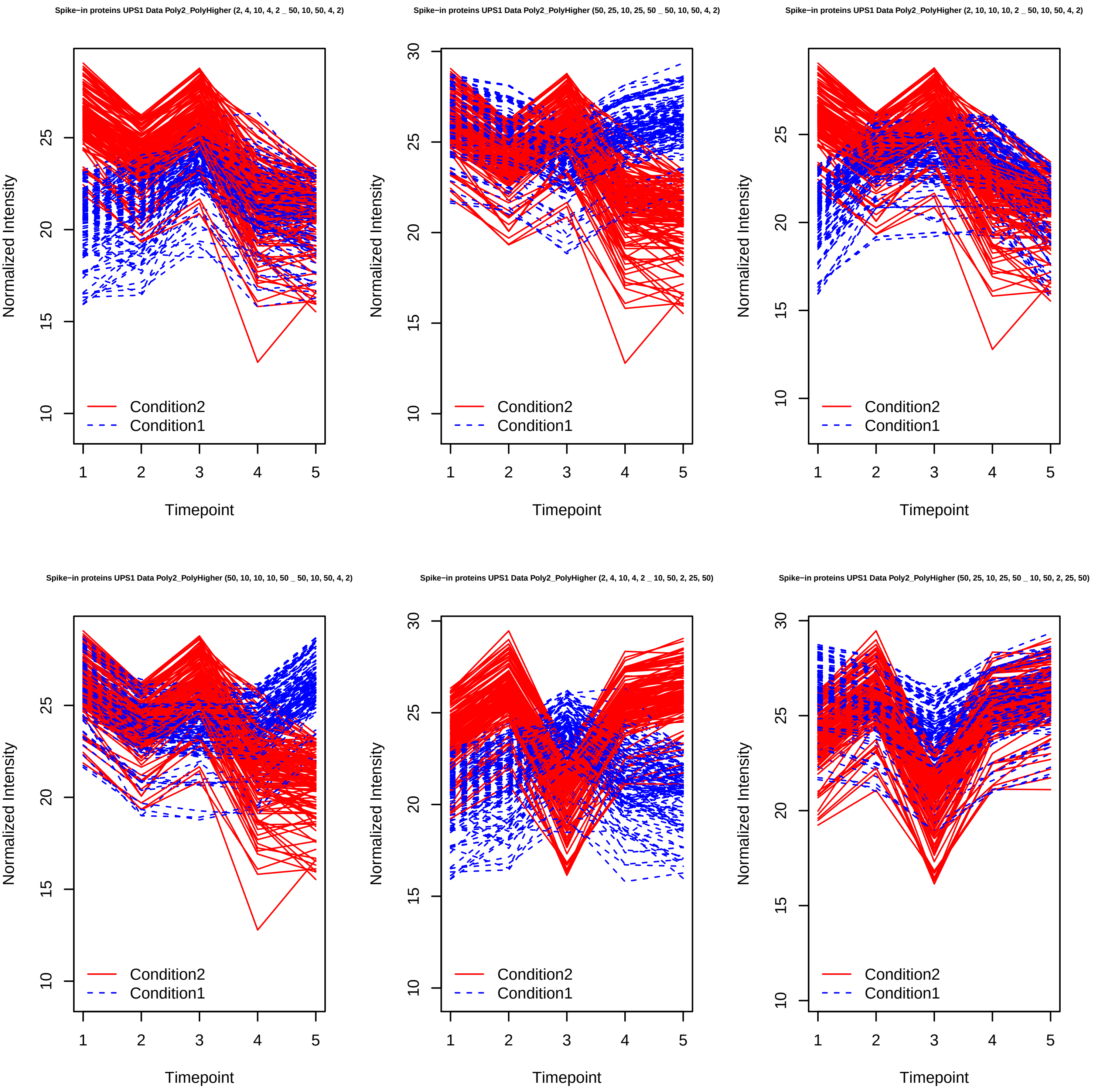
Spike−in proteins UPS1 Data Poly2_PolyHigher (2, 4, 10, 4, 2 _ 50, 10, 50, 4, 2)
Spike−in proteins UPS1 Data Poly2_PolyHigher (50, 25, 10, 25, 50 _ 50, 10, 50, 4, 2)
Spike−in proteins UPS1 Data Poly2_PolyHigher (2, 10, 10, 10, 2 _ 50, 10, 50, 4, 2)
Spike−in proteins UPS1 Data Poly2_PolyHigher (50, 10, 10, 10, 50 _ 50, 10, 50, 4, 2)
Spike−in proteins UPS1 Data Poly2_PolyHigher (2, 4, 10, 4, 2 _ 10, 50, 2, 25, 50)
Spike−in proteins UPS1 Data Poly2_PolyHigher (50, 25, 10, 25, 50 _ 10, 50, 2, 25, 50)
Normalized Intensity
Timepoint
Condition2
Condition1

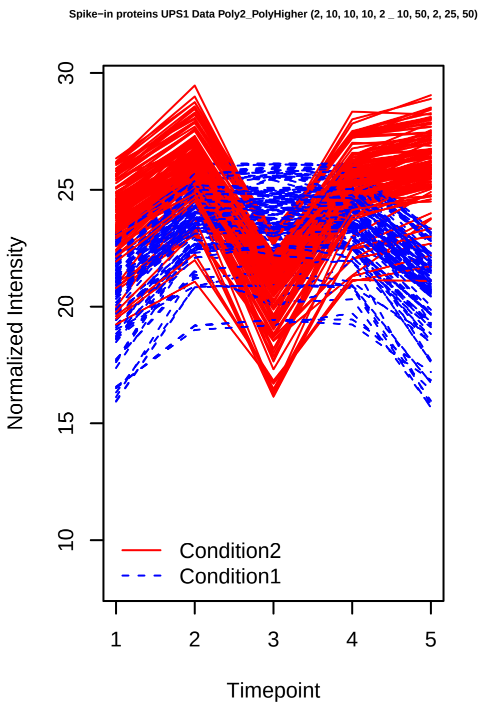
Spike−in proteins UPS1 Data Poly2_PolyHigher (2, 10, 10, 10, 2 _ 10, 50, 2, 25, 50)
Normalized Intensity
Timepoint
Condition2
Condition1

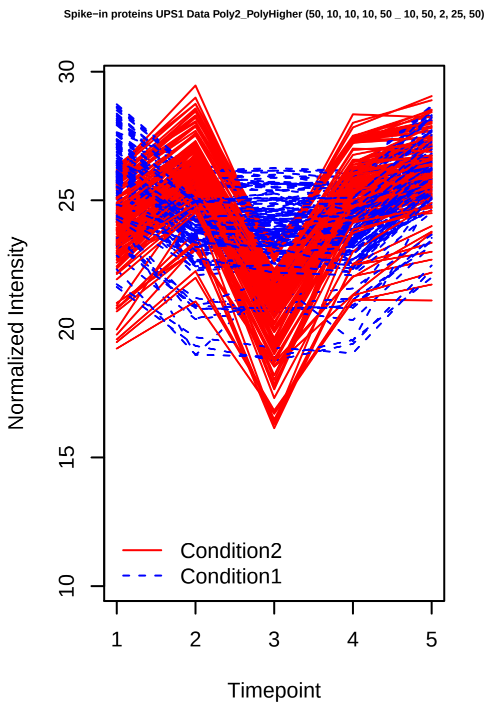
Spike−in proteins UPS1 Data Poly2_PolyHigher (50, 10, 10, 10, 50 _ 10, 50, 2, 25, 50)
Normalized Intensity
Timepoint
Condition2
Condition1

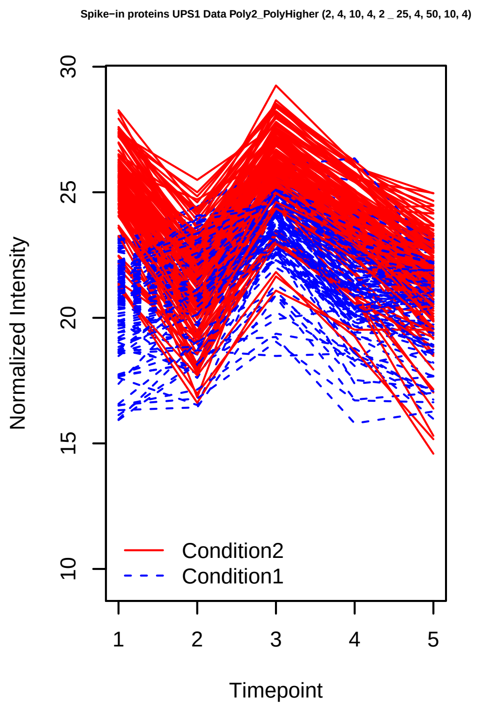
Spike−in proteins UPS1 Data Poly2_PolyHigher (2, 4, 10, 4, 2 _ 25, 4, 50, 10, 4)
Normalized Intensity
Timepoint
Condition2
Condition1

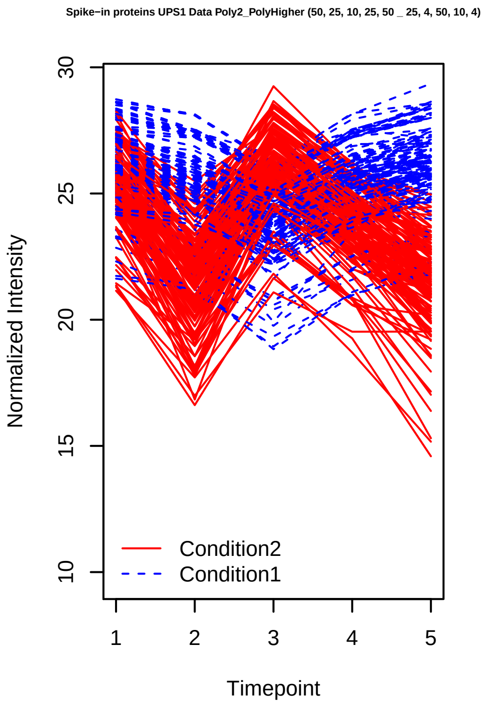
Spike−in proteins UPS1 Data Poly2_PolyHigher (50, 25, 10, 25, 50 _ 25, 4, 50, 10, 4)
Normalized Intensity
Timepoint
Condition2
Condition1

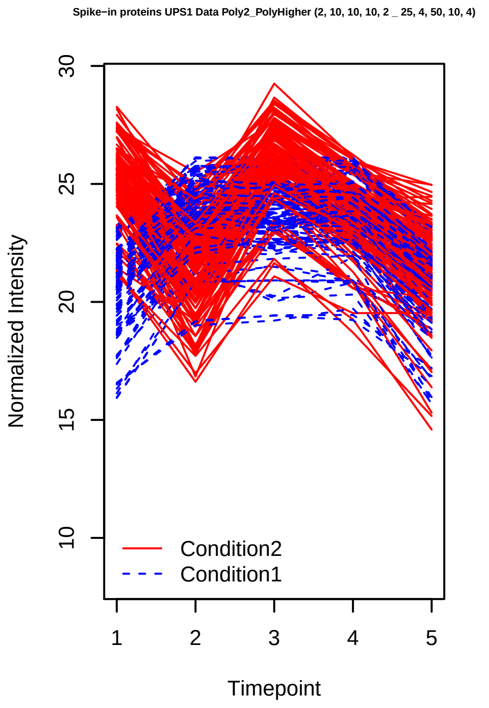
Spike−in proteins UPS1 Data Poly2_PolyHigher (2, 10, 10, 10, 2 _ 25, 4, 50, 10, 4)
Normalized Intensity
Timepoint
Condition2
Condition1

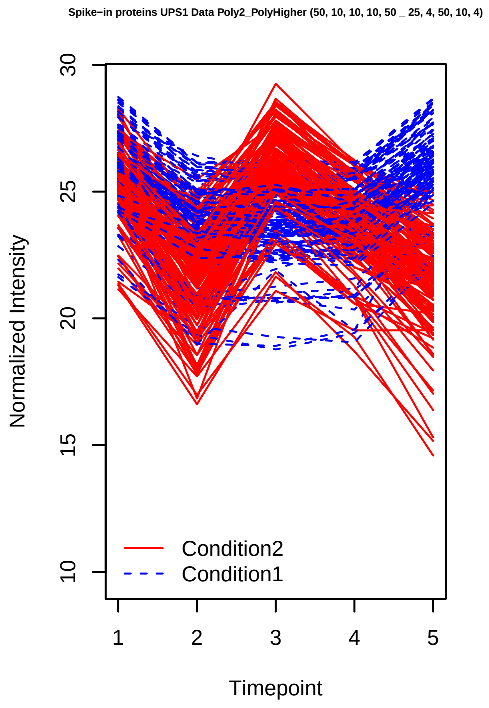
Spike−in proteins UPS1 Data Poly2_PolyHigher (50, 10, 10, 10, 50 _ 25, 4, 50, 10, 4)
Normalized Intensity
Timepoint
Condition2
Condition1

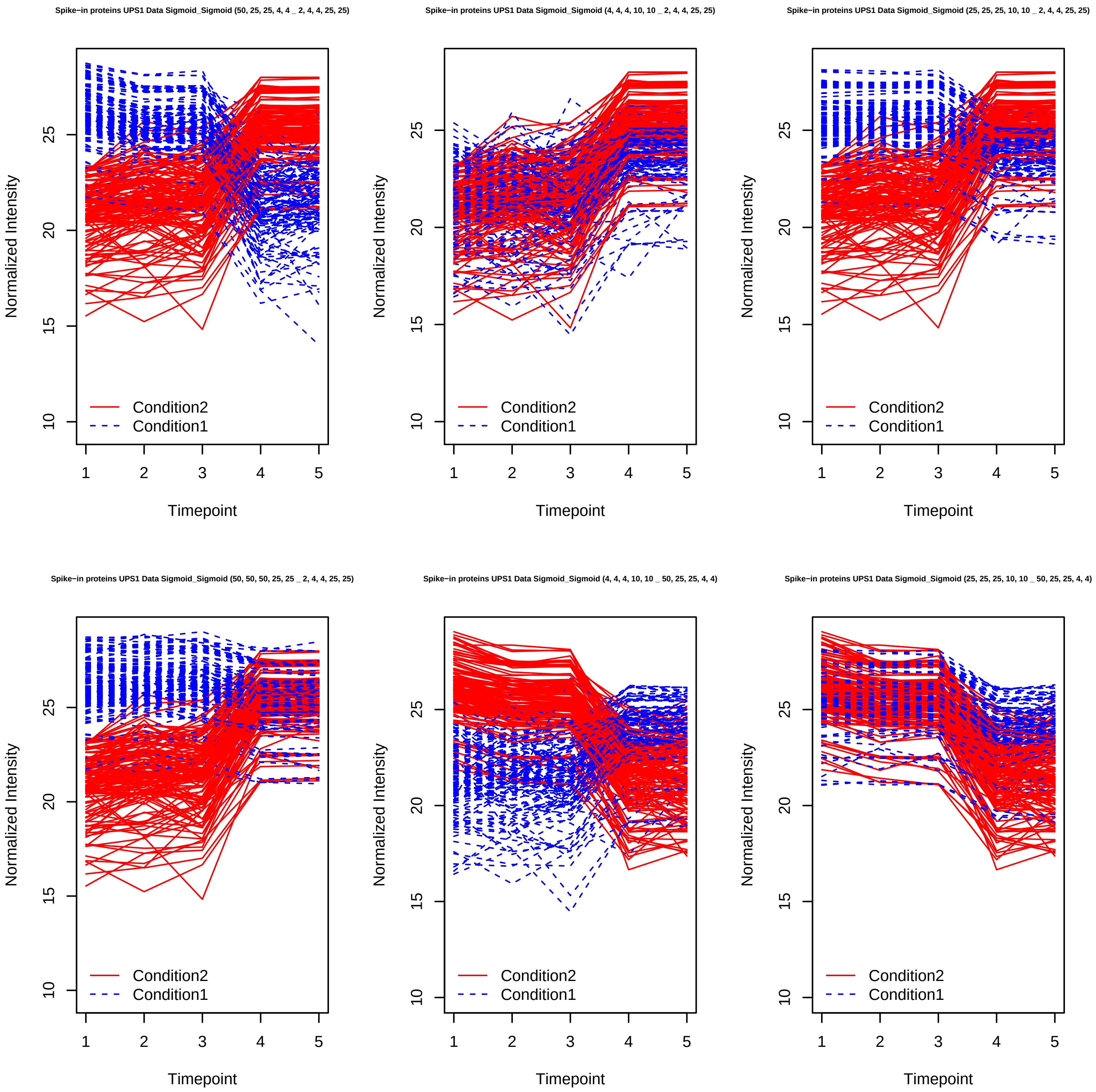
Spike−in proteins UPS1 Data Sigmoid_Sigmoid (50, 25, 25, 4, 4 _ 2, 4, 4, 25, 25)
Spike−in proteins UPS1 Data Sigmoid_Sigmoid (4, 4, 4, 10, 10 _ 2, 4, 4, 25, 25)
Spike−in proteins UPS1 Data Sigmoid_Sigmoid (25, 25, 25, 10, 10 _ 2, 4, 4, 25, 25)
Spike−in proteins UPS1 Data Sigmoid_Sigmoid (50, 50, 50, 25, 25 _ 2, 4, 4, 25, 25)
Spike−in proteins UPS1 Data Sigmoid_Sigmoid (4, 4, 4, 10, 10 _ 50, 25, 25, 4, 4)
Spike−in proteins UPS1 Data Sigmoid_Sigmoid (25, 25, 25, 10, 10 _ 50, 25, 25, 4, 4)
Normalized Intensity
Timepoint
Condition2
Condition1
10
15
20
25
1
2
3
4
5

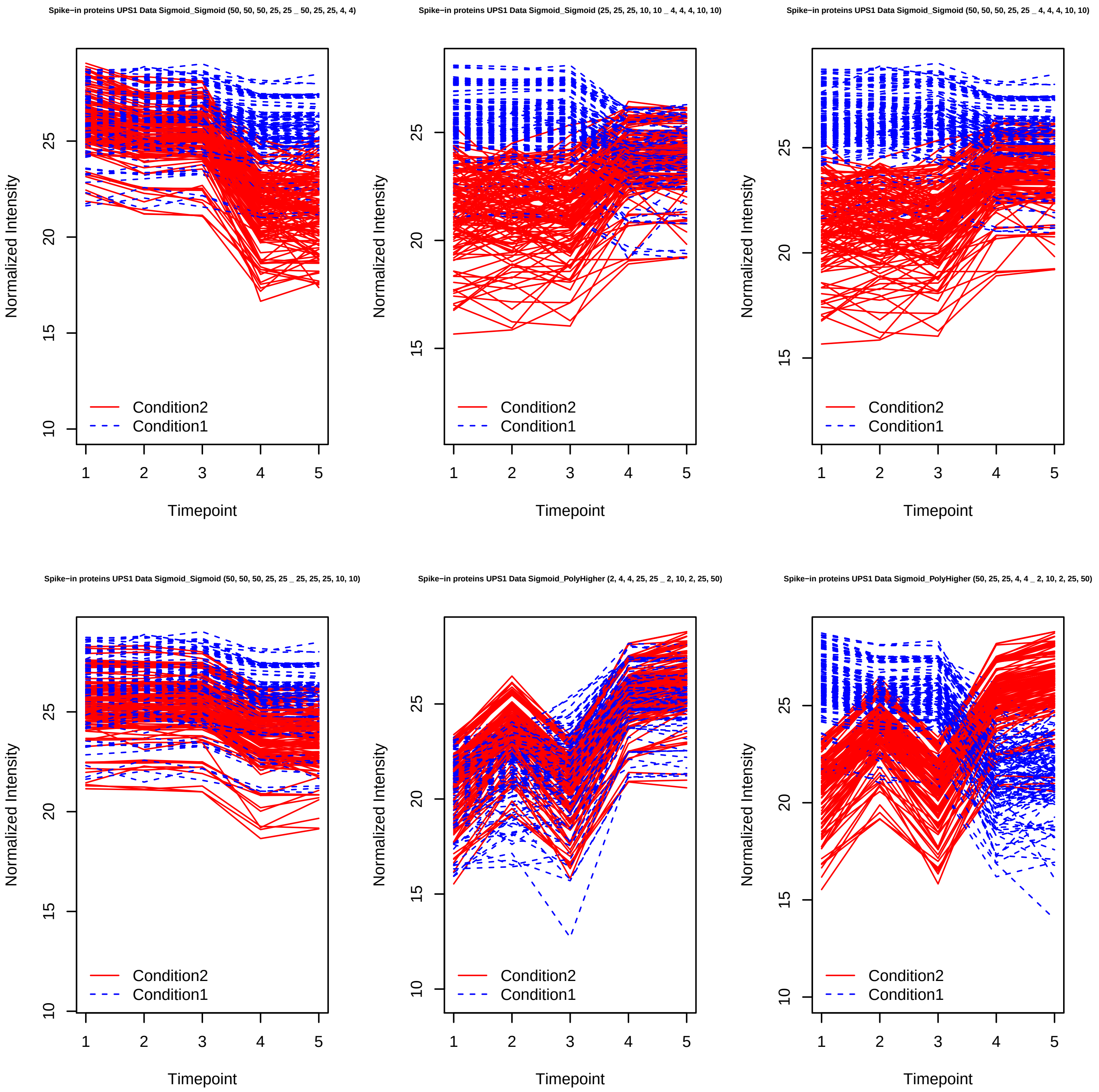
Spike−in proteins UPS1 Data Sigmoid_Sigmoid (50, 50, 50, 25, 25 _ 50, 25, 25, 4, 4)
Spike−in proteins UPS1 Data Sigmoid_Sigmoid (25, 25, 25, 10, 10 _ 4, 4, 4, 10, 10)
Spike−in proteins UPS1 Data Sigmoid_Sigmoid (50, 50, 50, 25, 25 _ 4, 4, 4, 10, 10)
Spike−in proteins UPS1 Data Sigmoid_Sigmoid (50, 50, 50, 25, 25 _ 25, 25, 25, 10, 10)
Spike−in proteins UPS1 Data Sigmoid_PolyHigher (2, 4, 4, 25, 25 _ 2, 10, 2, 25, 50)
Spike−in proteins UPS1 Data Sigmoid_PolyHigher (50, 25, 25, 4, 4 _ 2, 10, 2, 25, 50)
Normalized Intensity
Timepoint
Condition2
Condition1
10
15
20
25
1
2
3
4
5

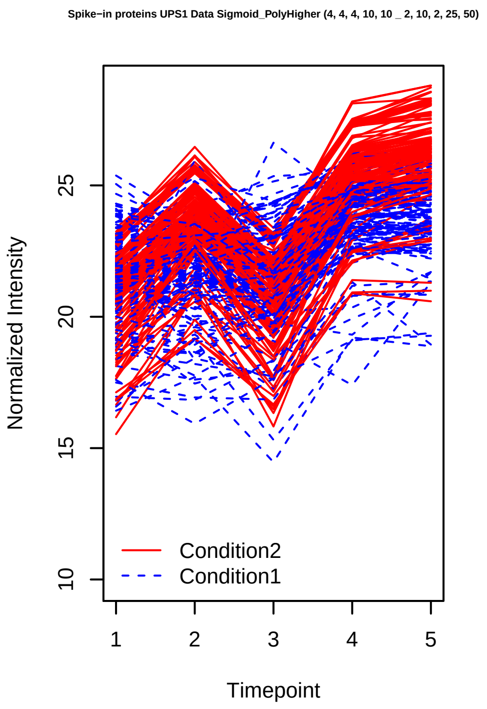
Spike−in proteins UPS1 Data Sigmoid_PolyHigher (4, 4, 4, 10, 10 _ 2, 10, 2, 25, 50)
Normalized Intensity
10
15
20
25
1
2
3
4
5
Timepoint
Condition2
Condition1

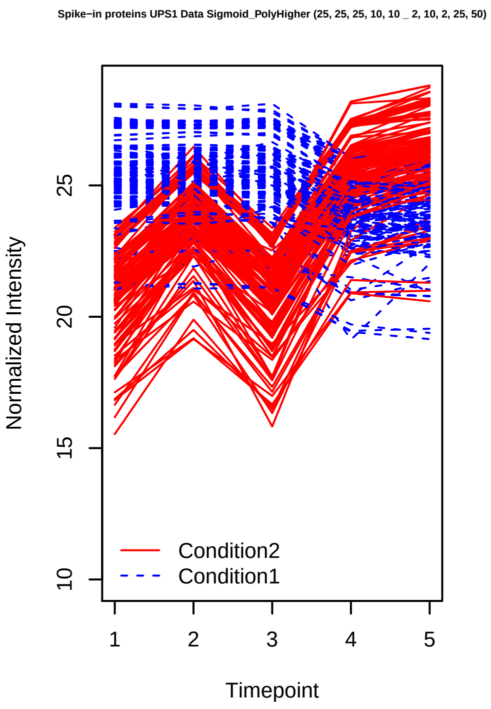
Spike−in proteins UPS1 Data Sigmoid_PolyHigher (25, 25, 25, 10, 10 _ 2, 10, 2, 25, 50)
Normalized Intensity
10
15
20
25
1
2
3
4
5
Timepoint
Condition2
Condition1

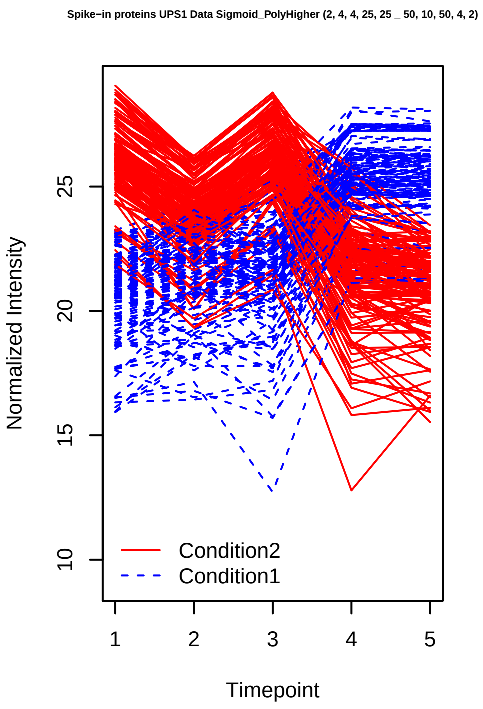
Spike−in proteins UPS1 Data Sigmoid_PolyHigher (2, 4, 4, 25, 25 _ 50, 10, 50, 4, 2)
Normalized Intensity
10
15
20
25
1
2
3
4
5
Timepoint
Condition2
Condition1

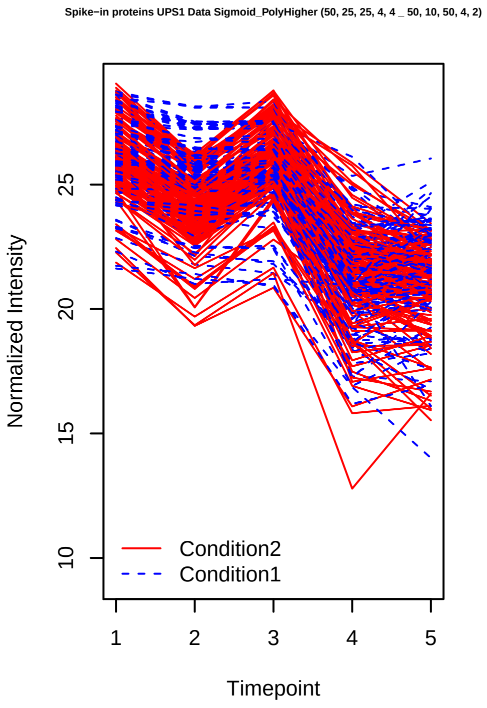
Spike−in proteins UPS1 Data Sigmoid_PolyHigher (50, 25, 25, 4, 4 _ 50, 10, 50, 4, 2)
Normalized Intensity
10
15
20
25
1
2
3
4
5
Timepoint
Condition2
Condition1

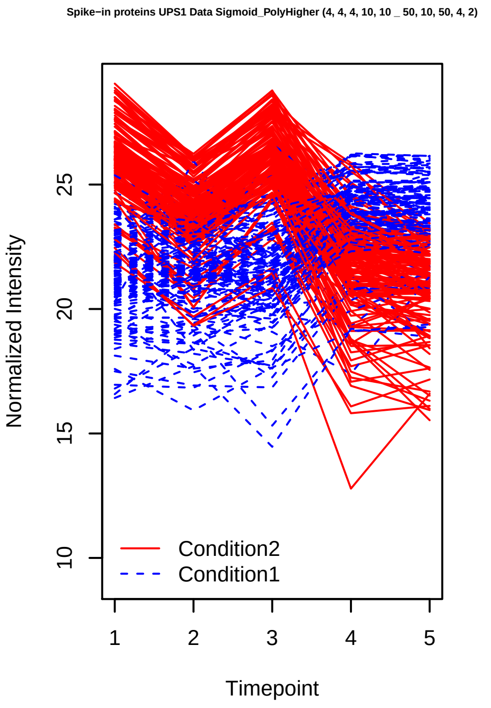
Spike−in proteins UPS1 Data Sigmoid_PolyHigher (4, 4, 4, 10, 10 _ 50, 10, 50, 4, 2)
Normalized Intensity
10
15
20
25
1
2
3
4
5
Timepoint
Condition2
Condition1

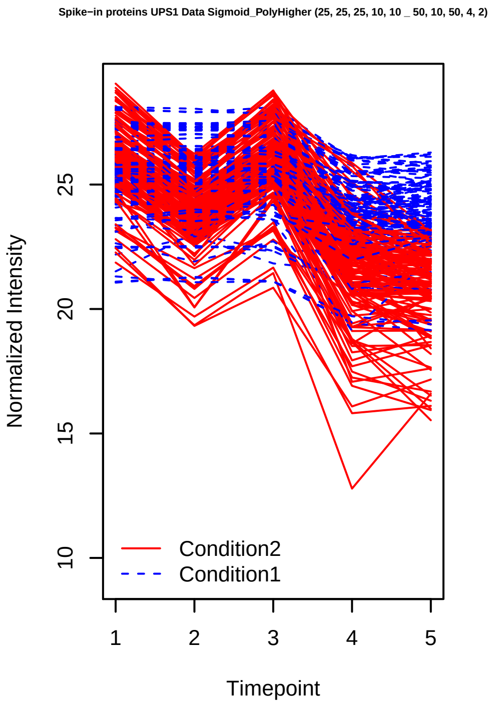
Spike−in proteins UPS1 Data Sigmoid_PolyHigher (25, 25, 25, 10, 10 _ 50, 10, 50, 4, 2)
Normalized Intensity
10
15
20
25
1
2
3
4
5
Timepoint
Condition2
Condition1

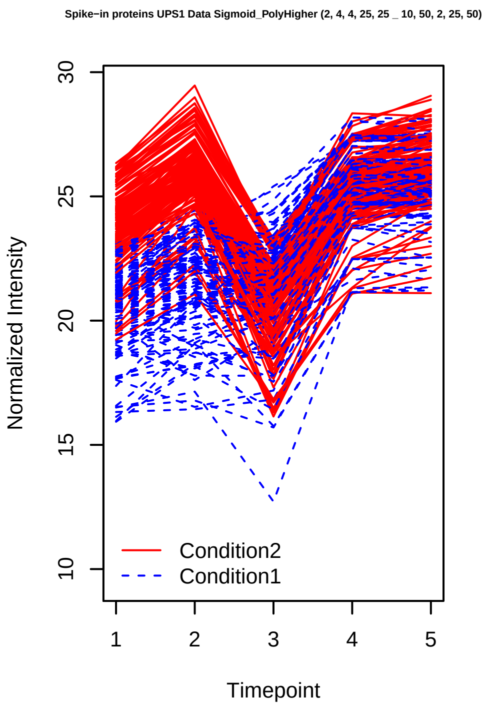
Spike−in proteins UPS1 Data Sigmoid_PolyHigher (2, 4, 4, 25, 25 _ 10, 50, 2, 25, 50)
Normalized Intensity
Timepoint
Condition2
Condition1

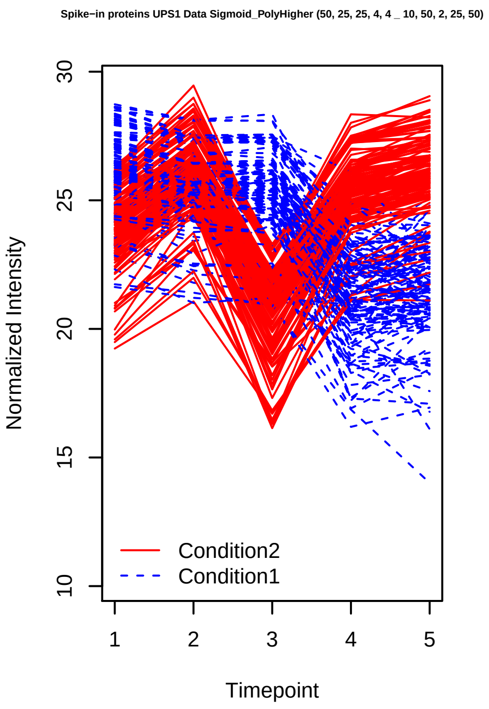
Spike−in proteins UPS1 Data Sigmoid_PolyHigher (50, 25, 25, 4, 4 _ 10, 50, 2, 25, 50)
Normalized Intensity
Timepoint
Condition2
Condition1

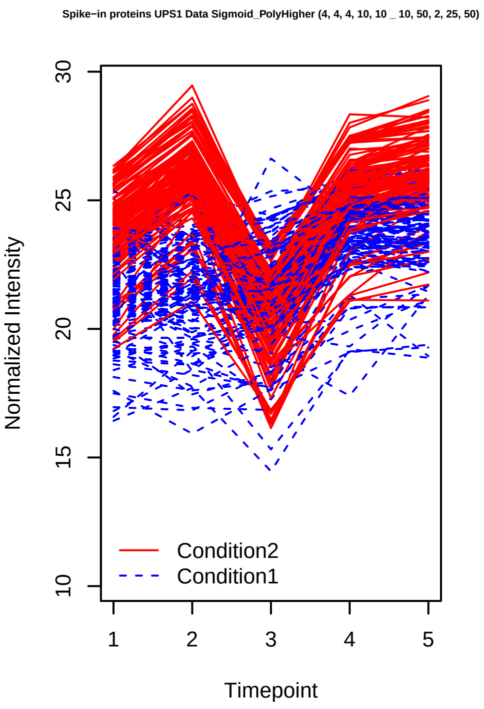
Spike−in proteins UPS1 Data Sigmoid_PolyHigher (4, 4, 4, 10, 10 _ 10, 50, 2, 25, 50)
Normalized Intensity
Timepoint
Condition2
Condition1

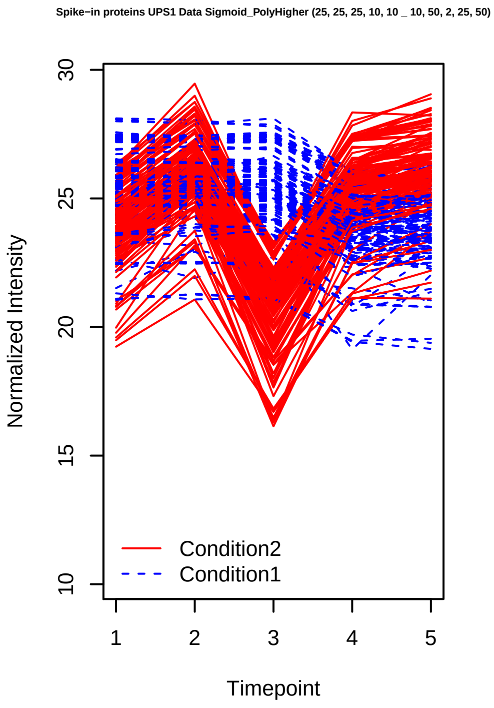
Spike−in proteins UPS1 Data Sigmoid_PolyHigher (25, 25, 25, 10, 10 _ 10, 50, 2, 25, 50)
Normalized Intensity
Timepoint
Condition2
Condition1

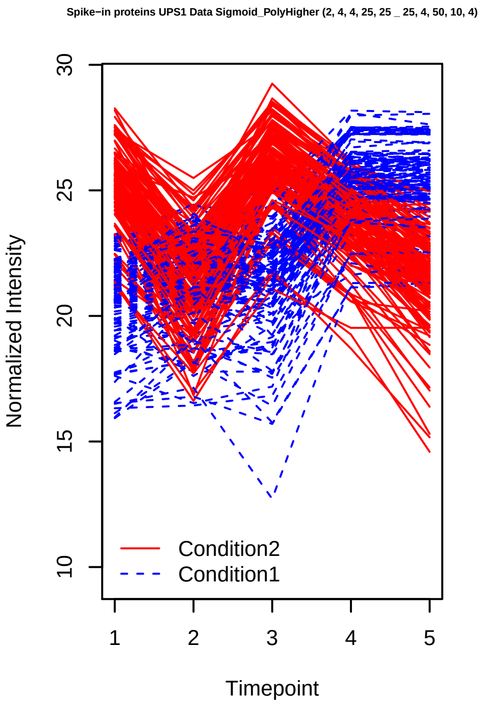
Spike−in proteins UPS1 Data Sigmoid_PolyHigher (2, 4, 4, 25, 25 _ 25, 4, 50, 10, 4)
Normalized Intensity
Timepoint
Condition2
Condition1

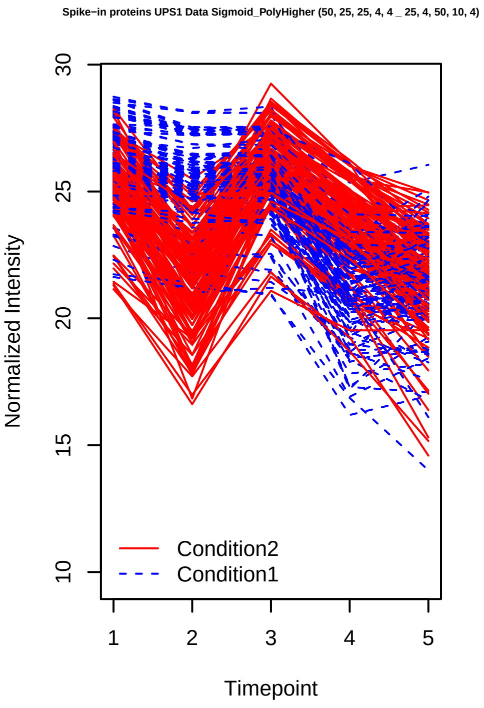
Spike−in proteins UPS1 Data Sigmoid_PolyHigher (50, 25, 25, 4, 4 _ 25, 4, 50, 10, 4)
Normalized Intensity
Timepoint
Condition2
Condition1

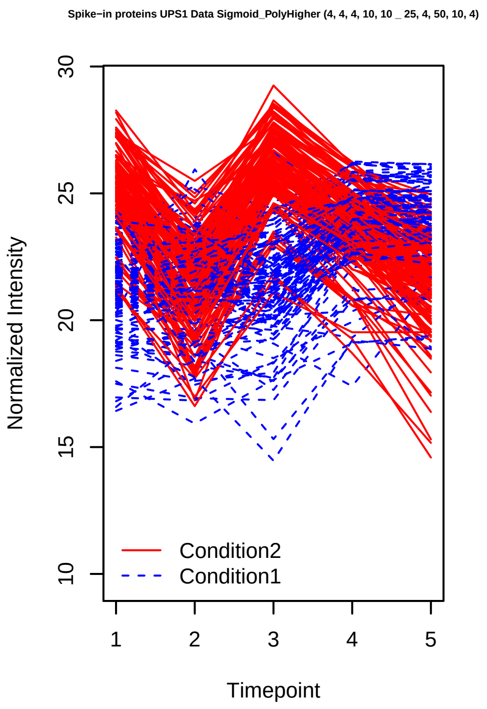
Spike−in proteins UPS1 Data Sigmoid_PolyHigher (4, 4, 4, 10, 10 _ 25, 4, 50, 10, 4)
Normalized Intensity
Timepoint
Condition2
Condition1

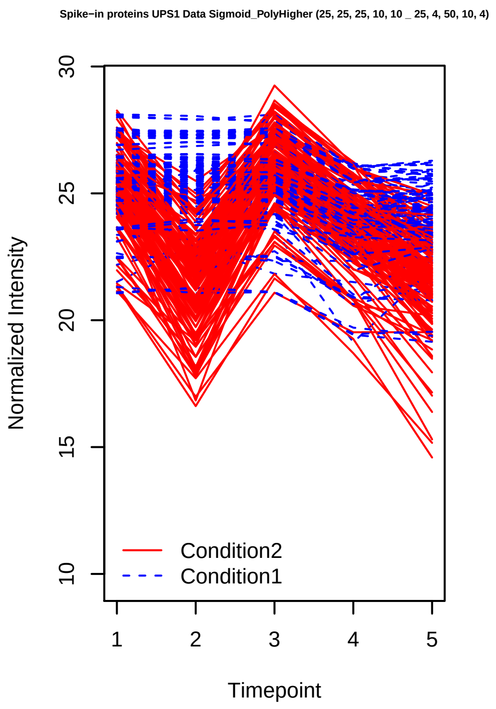
Spike−in proteins UPS1 Data Sigmoid_PolyHigher (25, 25, 25, 10, 10 _ 25, 4, 50, 10, 4)
Normalized Intensity
Timepoint
Condition2
Condition1

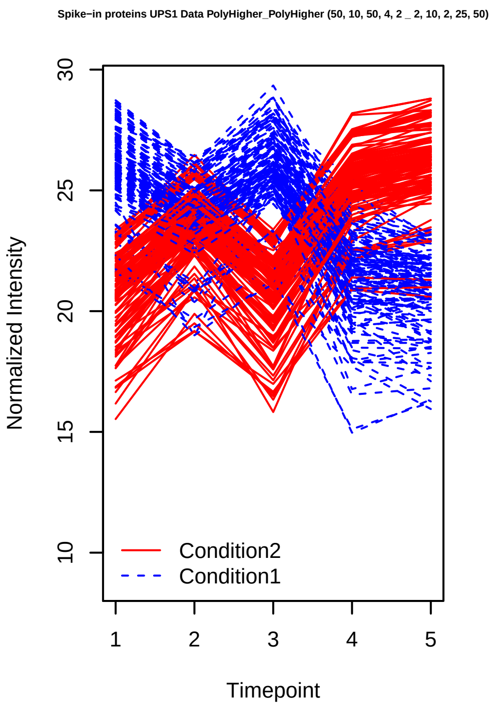
Spike−in proteins UPS1 Data PolyHigher_PolyHigher (50, 10, 50, 4, 2 _ 2, 10, 2, 25, 50)
Normalized Intensity
Timepoint
Condition2
Condition1

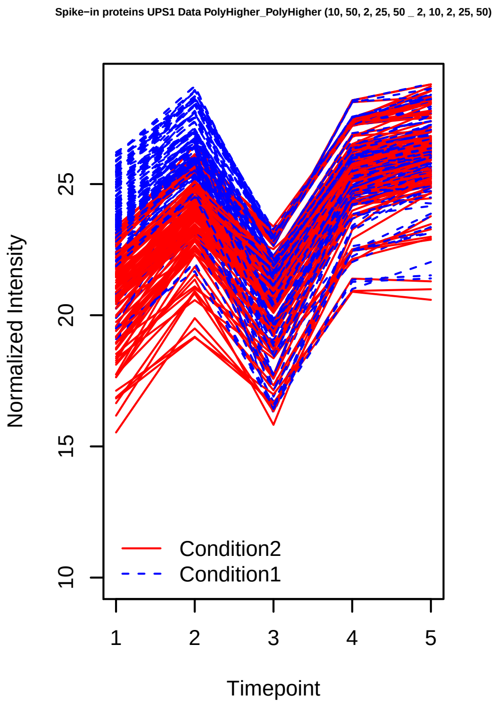
Spike−in proteins UPS1 Data PolyHigher_PolyHigher (10, 50, 2, 25, 50 _ 2, 10, 2, 25, 50)
Normalized Intensity
Timepoint
Condition2
Condition1

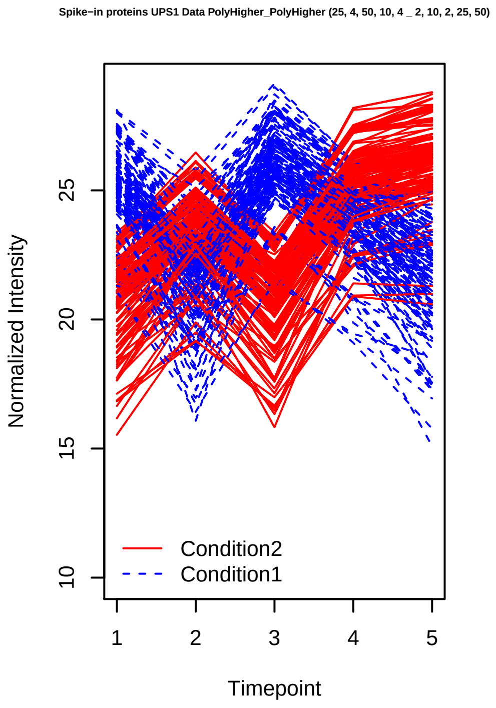
Spike−in proteins UPS1 Data PolyHigher_PolyHigher (25, 4, 50, 10, 4 _ 2, 10, 2, 25, 50)
Normalized Intensity
Timepoint
Condition2
Condition1

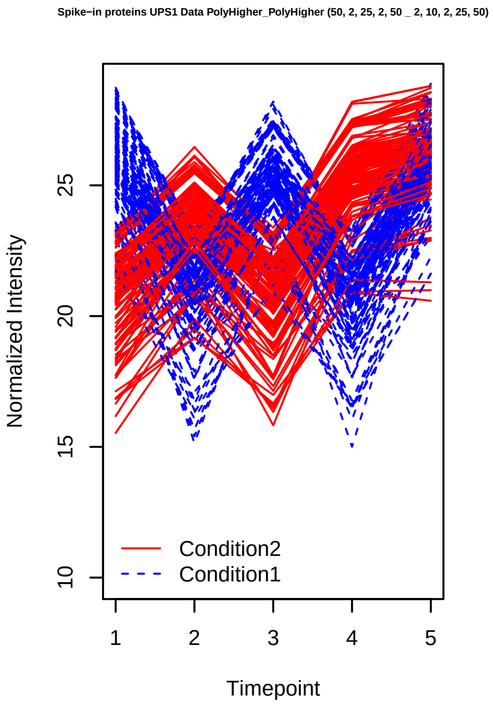
Spike−in proteins UPS1 Data PolyHigher_PolyHigher (50, 2, 25, 2, 50 _ 2, 10, 2, 25, 50)
Normalized Intensity
Timepoint
Condition2
Condition1

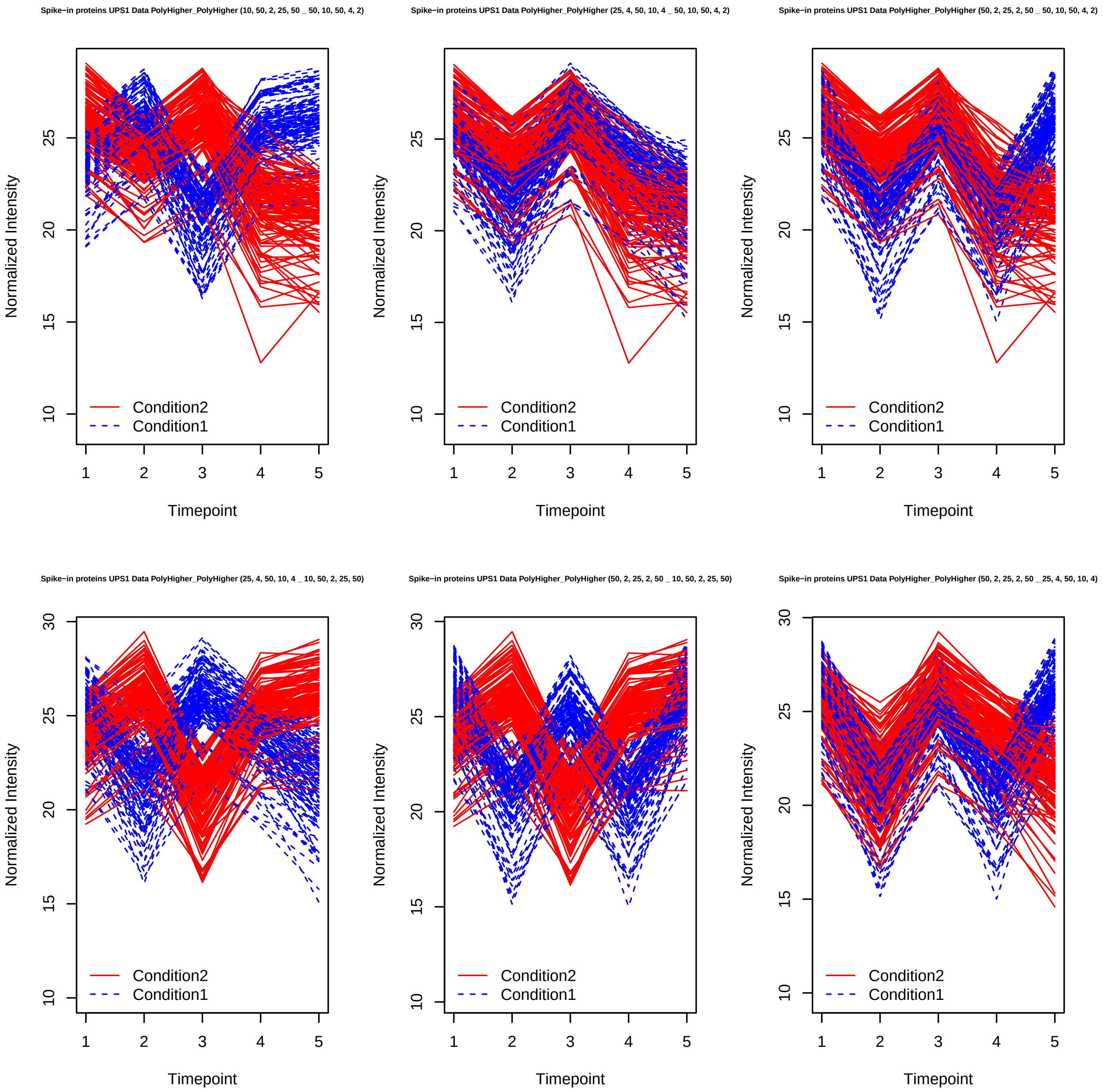
Spike−in proteins UPS1 Data PolyHigher_PolyHigher (10, 50, 2, 25, 50 _ 50, 10, 50, 4, 2)
Spike−in proteins UPS1 Data PolyHigher_PolyHigher (25, 4, 50, 10, 4 _ 50, 10, 50, 4, 2)
Spike−in proteins UPS1 Data PolyHigher_PolyHigher (50, 2, 25, 2, 50 _ 50, 10, 50, 4, 2)
Spike−in proteins UPS1 Data PolyHigher_PolyHigher (25, 4, 50, 10, 4 _ 10, 50, 2, 25, 50)
Spike−in proteins UPS1 Data PolyHigher_PolyHigher (50, 2, 25, 2, 50 _ 10, 50, 2, 25, 50)
Spike−in proteins UPS1 Data PolyHigher_PolyHigher (50, 2, 25, 2, 50 _ 25, 4, 50, 10, 4)
Normalized Intensity
Timepoint
Condition2
Condition1

Spike−in proteins SGSDS Data Stable_Stable (3, 3, 3, 3, 3, 3, 3, 3 _ 1, 1, 1, 1, 1, 1, 1, 1)

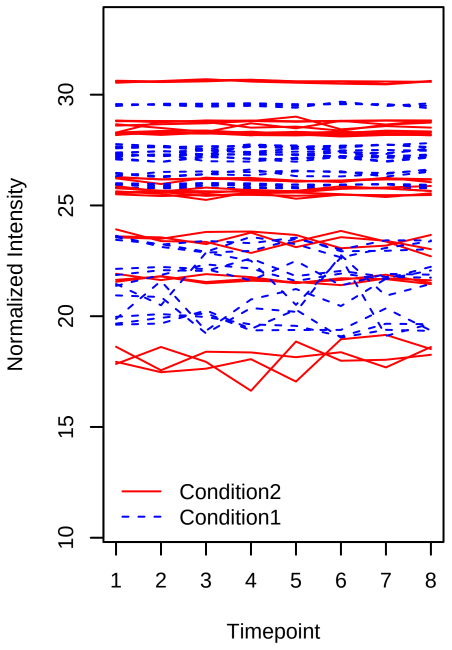


Spike−in proteins SGSDS Data Stable_Stable (5, 5, 5, 5, 5, 5, 5, 5 _ 1, 1, 1, 1, 1, 1, 1, 1)

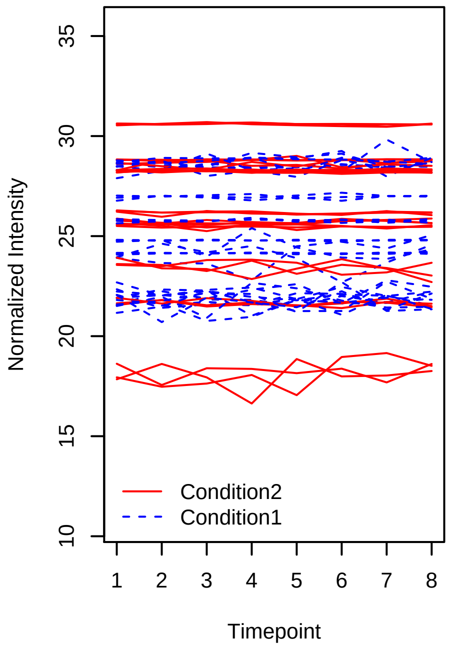


Spike−in proteins SGSDS Data Stable_Stable (7, 7, 7, 7, 7, 7, 7, 7 _ 1, 1, 1, 1, 1, 1, 1, 1)

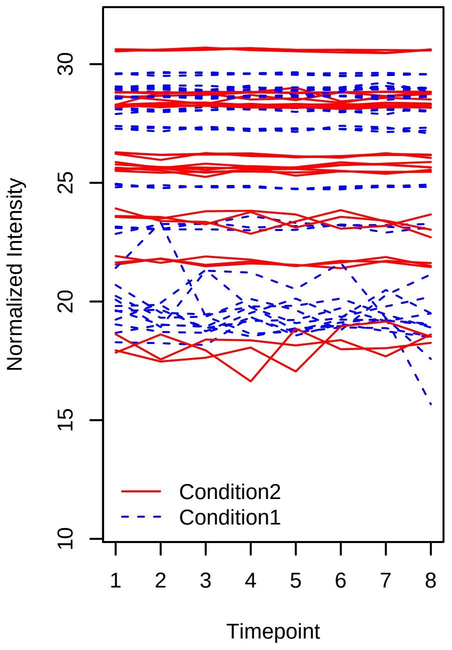


Spike−in proteins SGSDS Data Stable_Stable (8, 8, 8, 8, 8, 8, 8, 8 _ 1, 1, 1, 1, 1, 1, 1, 1)

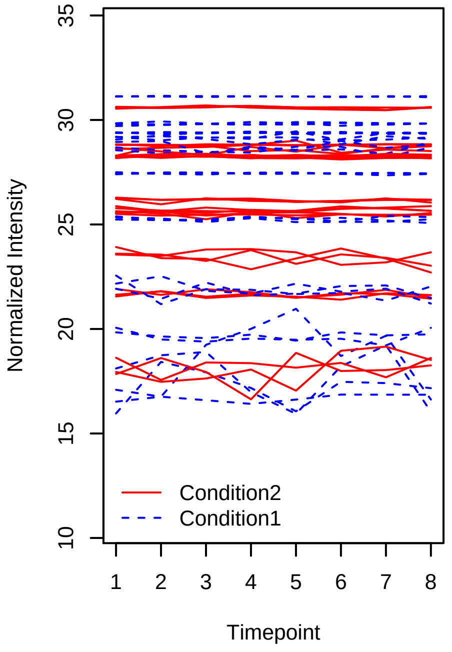


Spike−in proteins SGSDS Data Stable_Stable (5, 5, 5, 5, 5, 5, 5, 5 _ 3, 3, 3, 3, 3, 3, 3, 3)

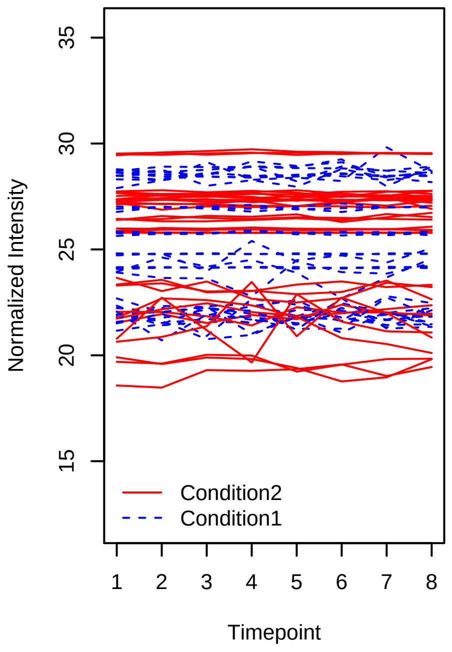


Spike−in proteins SGSDS Data Stable_Stable (7, 7, 7, 7, 7, 7, 7, 7 _ 3, 3, 3, 3, 3, 3, 3, 3)

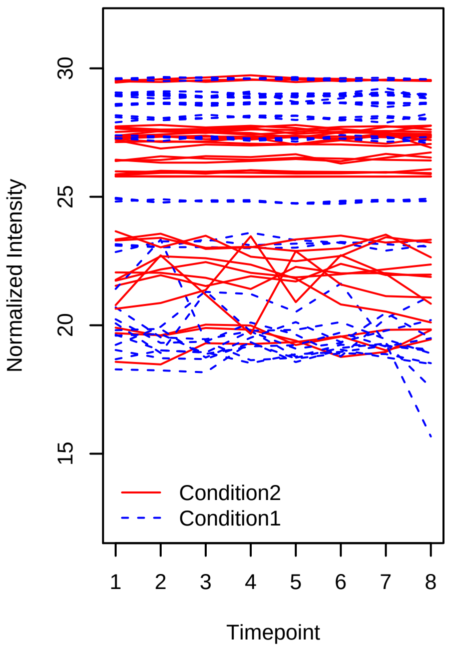

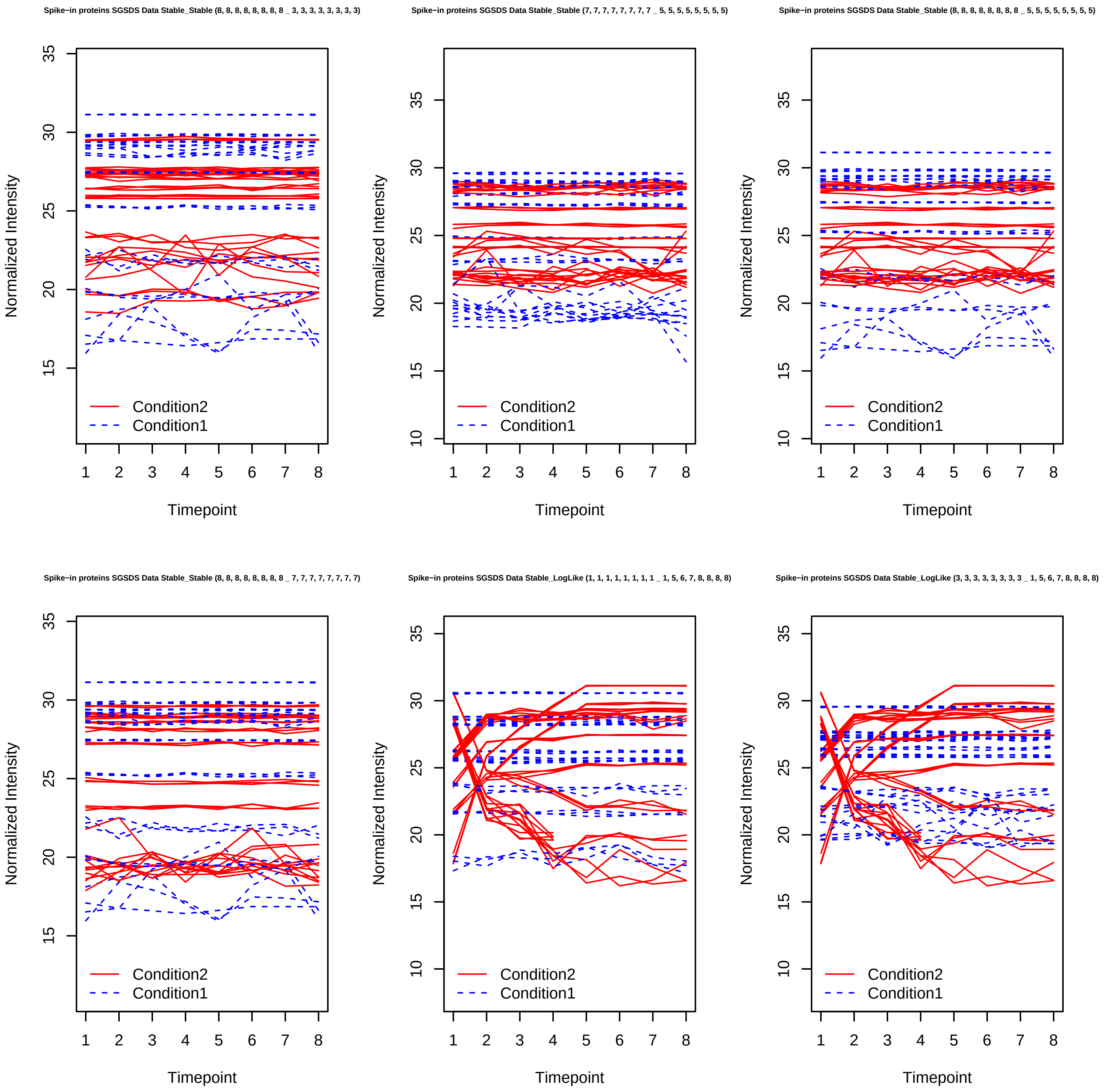

Spike−in proteins SGSDS Data Stable_Stable (8, 8, 8, 8, 8, 8, 8, 8 _ 3, 3, 3, 3, 3, 3, 3, 3)
Spike−in proteins SGSDS Data Stable_Stable (7, 7, 7, 7, 7, 7, 7, 7 _ 5, 5, 5, 5, 5, 5, 5, 5)
Spike−in proteins SGSDS Data Stable_Stable (8, 8, 8, 8, 8, 8, 8, 8 _ 5, 5, 5, 5, 5, 5, 5, 5)
Spike−in proteins SGSDS Data Stable_Stable (8, 8, 8, 8, 8, 8, 8, 8 _ 7, 7, 7, 7, 7, 7, 7, 7)
Spike−in proteins SGSDS Data Stable_LogLike (1, 1, 1, 1, 1, 1, 1, 1 _ 1, 5, 6, 7, 8, 8, 8, 8)
Spike−in proteins SGSDS Data Stable_LogLike (3, 3, 3, 3, 3, 3, 3, 3 _ 1, 5, 6, 7, 8, 8, 8, 8)
Normalized Intensity
Timepoint
Condition2
Condition1

Spike−in proteins SGSDS Data Stable_LogLike (5, 5, 5, 5, 5, 5, 5, 5 _ 1, 5, 6, 7, 8, 8, 8, 8)

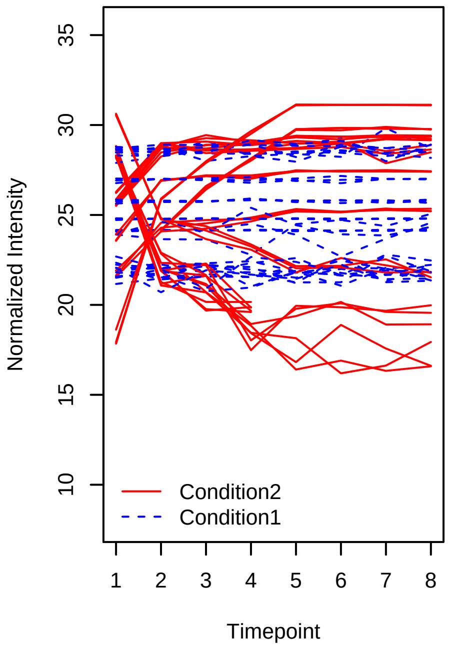


Spike−in proteins SGSDS Data Stable_LogLike (7, 7, 7, 7, 7, 7, 7, 7 _ 1, 5, 6, 7, 8, 8, 8, 8)

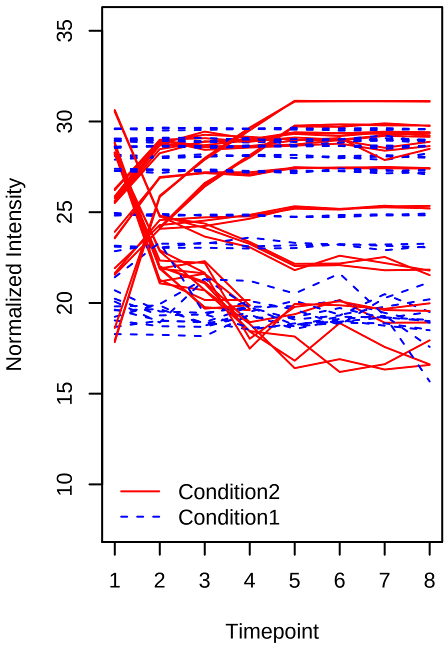


Spike−in proteins SGSDS Data Stable_LogLike (1, 1, 1, 1, 1, 1, 1, 1 _ 8, 4, 3, 2, 1, 1, 1, 1)

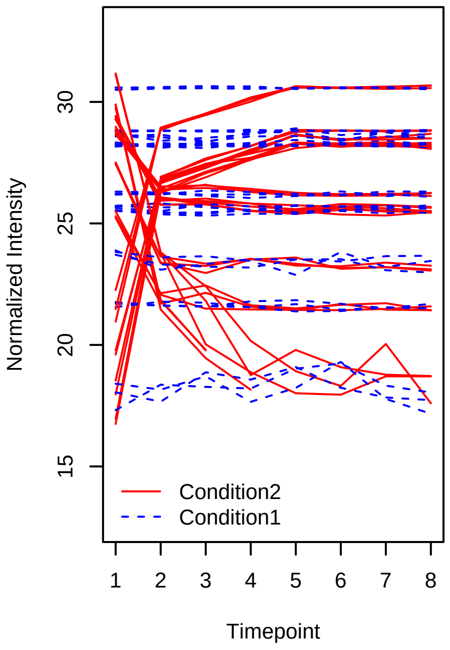


Spike−in proteins SGSDS Data Stable_LogLike (3, 3, 3, 3, 3, 3, 3, 3 _ 8, 4, 3, 2, 1, 1, 1, 1)

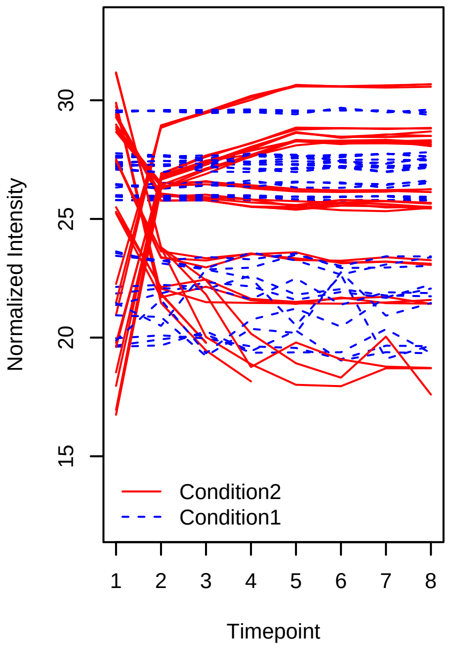


Spike−in proteins SGSDS Data Stable_LogLike (5, 5, 5, 5, 5, 5, 5, 5 _ 8, 4, 3, 2, 1, 1, 1, 1)

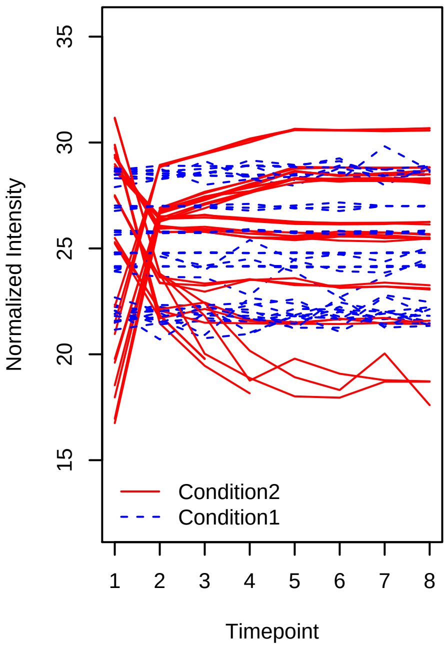


Spike−in proteins SGSDS Data Stable_LogLike (7, 7, 7, 7, 7, 7, 7, 7 _ 8, 4, 3, 2, 1, 1, 1, 1)

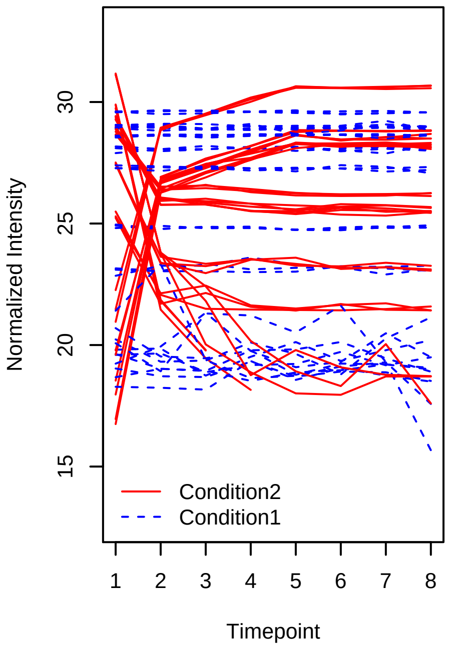

Spike−in proteins SGSDS Data Stable_LogLike (1, 1, 1, 1, 1, 1, 1, 1 _ 8, 7, 7, 6, 5, 5, 4, 1)

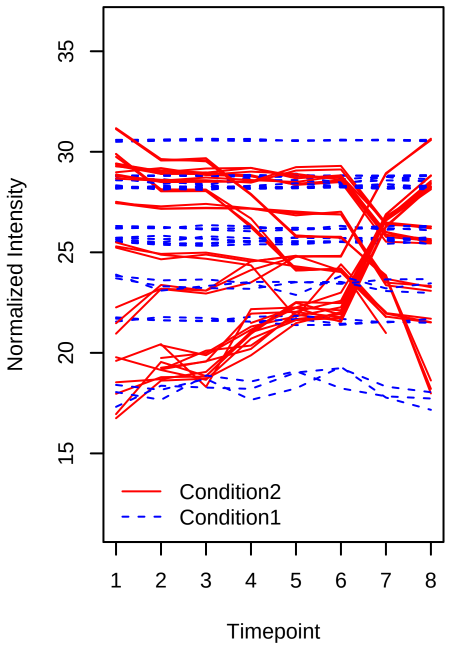


Spike−in proteins SGSDS Data Stable_LogLike (3, 3, 3, 3, 3, 3, 3, 3 _ 8, 7, 7, 6, 5, 5, 4, 1)

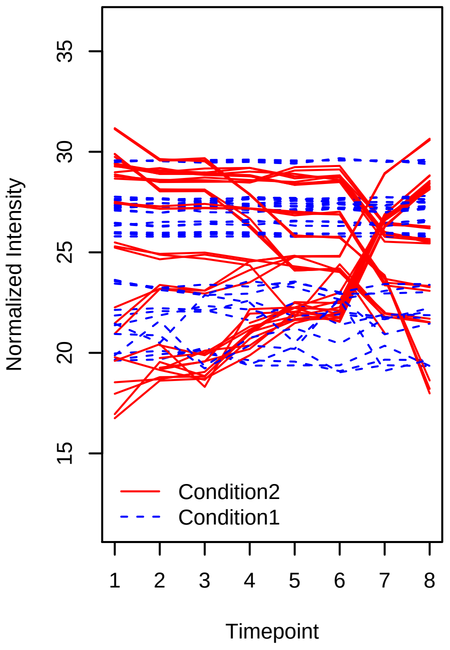


Spike−in proteins SGSDS Data Stable_LogLike (5, 5, 5, 5, 5, 5, 5, 5 _ 8, 7, 7, 6, 5, 5, 4, 1)

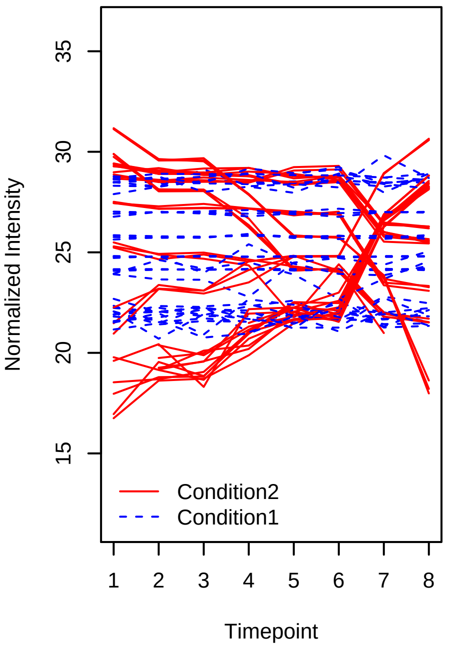


Spike−in proteins SGSDS Data Stable_LogLike (7, 7, 7, 7, 7, 7, 7, 7 _ 8, 7, 7, 6, 5, 5, 4, 1)

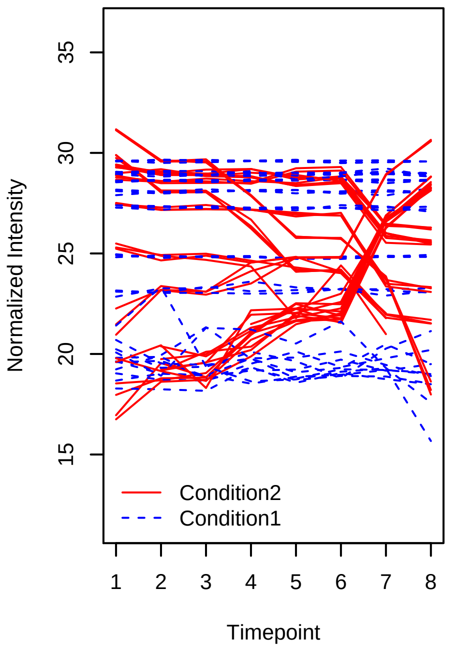


Spike−in proteins SGSDS Data Stable_LogLike (1, 1, 1, 1, 1, 1, 1, 1 _ 1, 1, 2, 2, 3, 6, 7, 8)

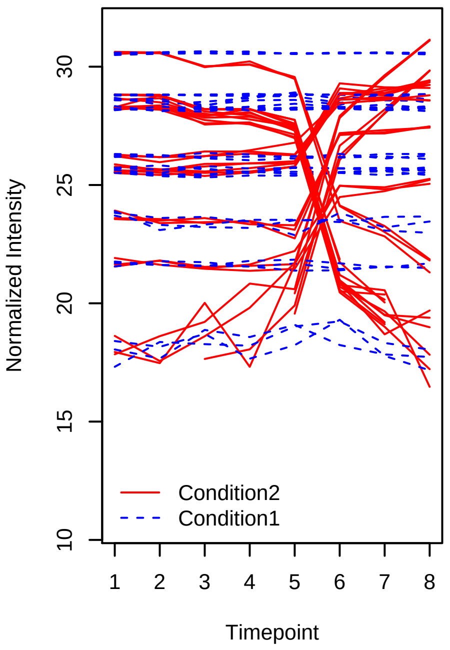


Spike−in proteins SGSDS Data Stable_LogLike (3, 3, 3, 3, 3, 3, 3, 3 _ 1, 1, 2, 2, 3, 6, 7, 8)

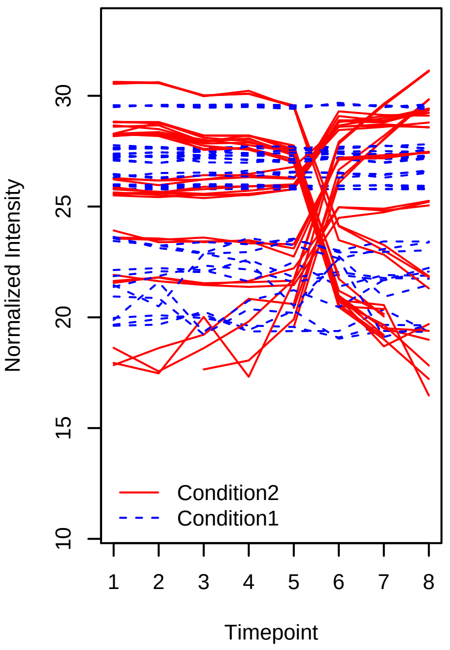

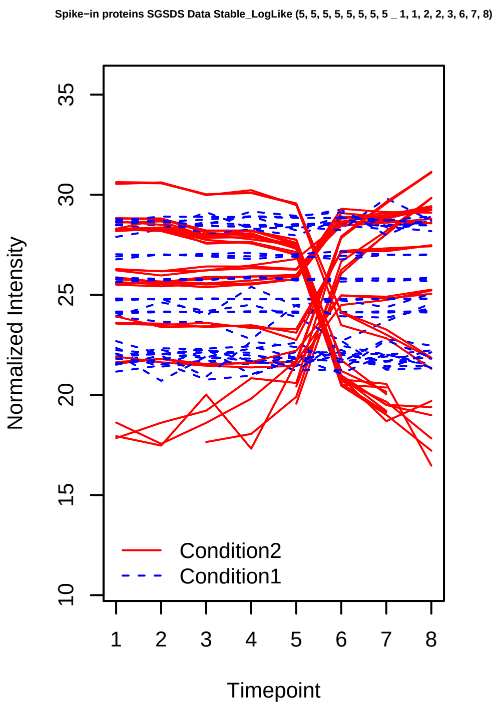
Spike−in proteins SGSDS Data Stable_LogLike (5, 5, 5, 5, 5, 5, 5, 5 _ 1, 1, 2, 2, 3, 6, 7, 8)
Normalized Intensity
10
15
20
25
30
35
Condition2
Condition1
1
2
3
4
5
6
7
8
Timepoint

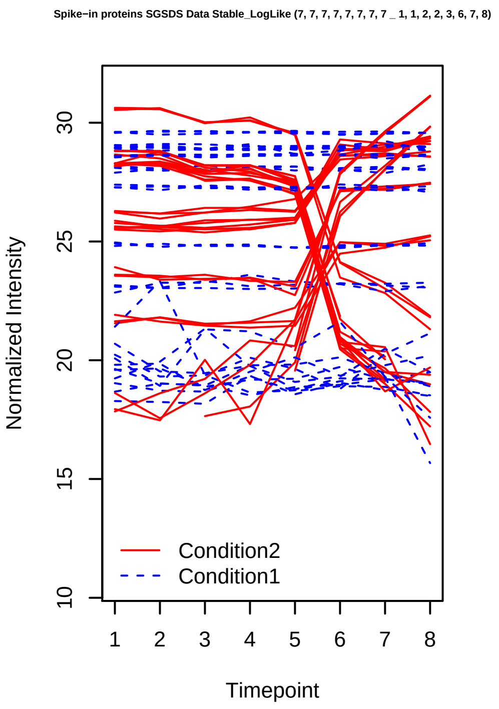
Spike−in proteins SGSDS Data Stable_LogLike (7, 7, 7, 7, 7, 7, 7, 7 _ 1, 1, 2, 2, 3, 6, 7, 8)
Normalized Intensity
10
15
20
25
30
Condition2
Condition1
1
2
3
4
5
6
7
8
Timepoint

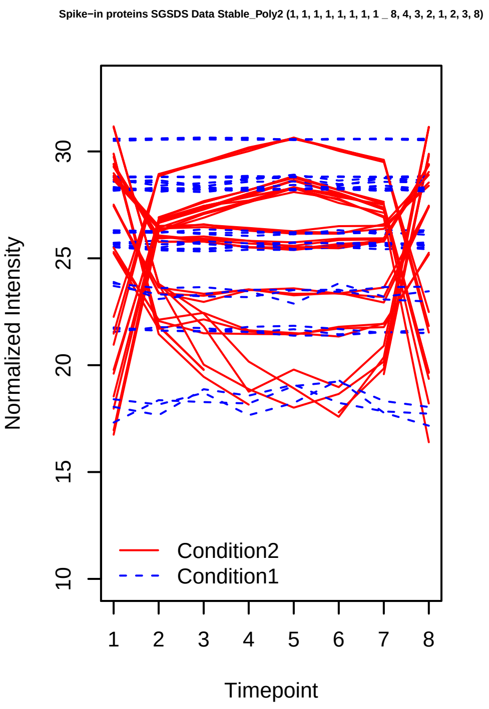
Spike−in proteins SGSDS Data Stable_Poly2 (1, 1, 1, 1, 1, 1, 1, 1 _ 8, 4, 3, 2, 1, 2, 3, 8)
Normalized Intensity
10
15
20
25
30
Condition2
Condition1
1
2
3
4
5
6
7
8
Timepoint

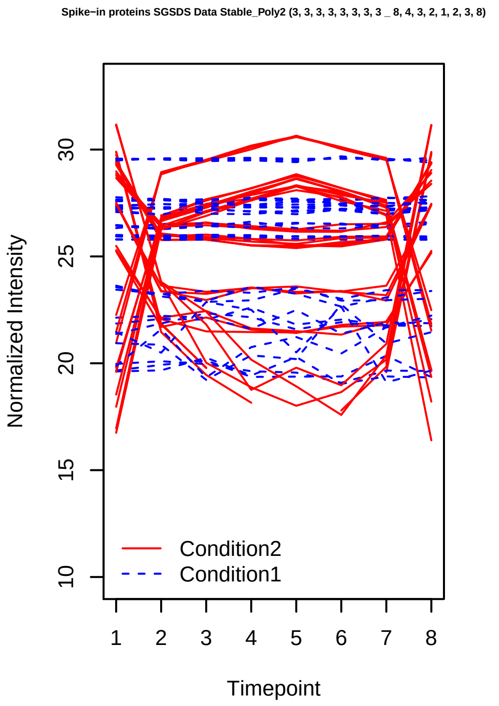
Spike−in proteins SGSDS Data Stable_Poly2 (3, 3, 3, 3, 3, 3, 3, 3 _ 8, 4, 3, 2, 1, 2, 3, 8)
Normalized Intensity
10
15
20
25
30
Condition2
Condition1
1
2
3
4
5
6
7
8
Timepoint

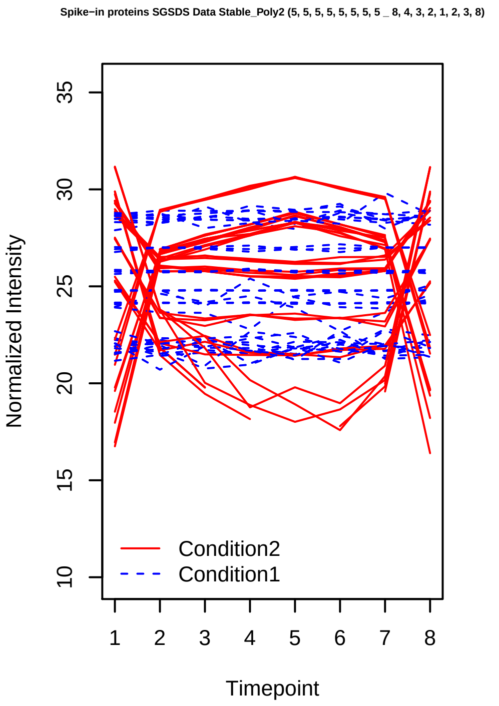
Spike−in proteins SGSDS Data Stable_Poly2 (5, 5, 5, 5, 5, 5, 5, 5 _ 8, 4, 3, 2, 1, 2, 3, 8)
Normalized Intensity
10
15
20
25
30
35
Condition2
Condition1
1
2
3
4
5
6
7
8
Timepoint

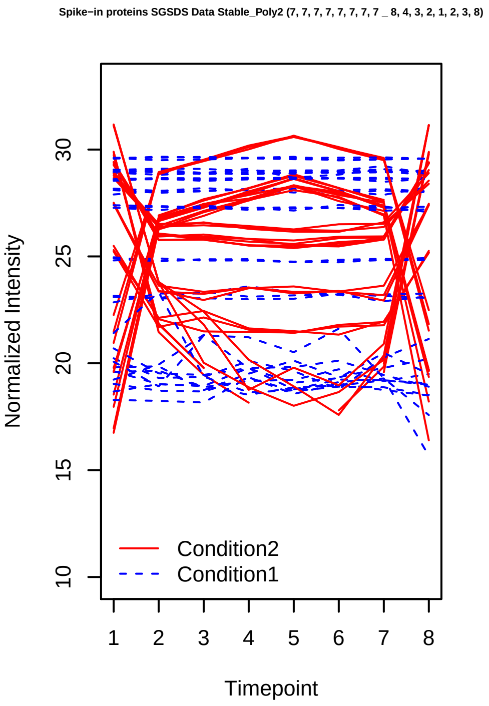
Spike−in proteins SGSDS Data Stable_Poly2 (7, 7, 7, 7, 7, 7, 7, 7 _ 8, 4, 3, 2, 1, 2, 3, 8)
Normalized Intensity
10
15
20
25
30
Condition2
Condition1
1
2
3
4
5
6
7
8
Timepoint

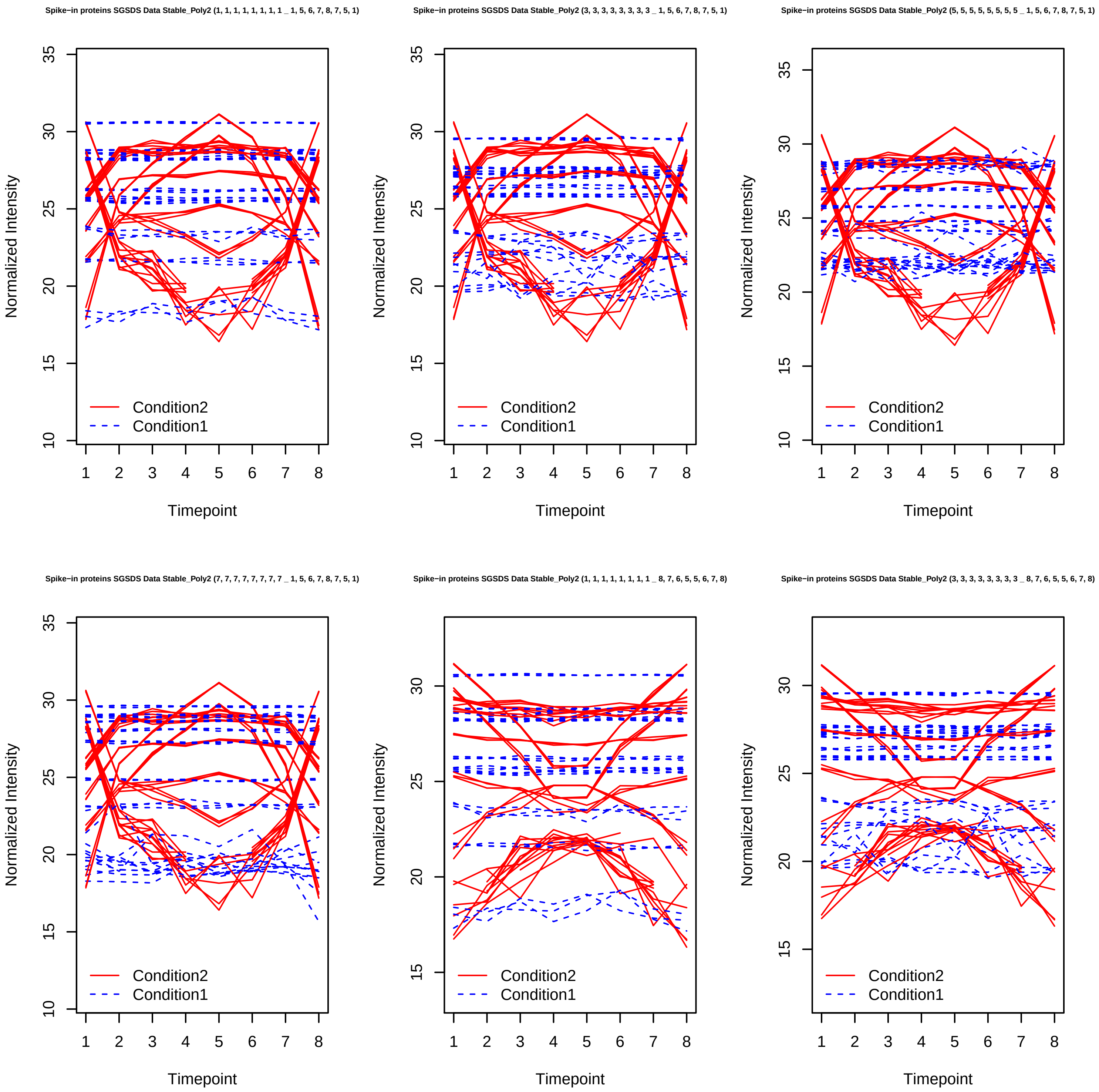
Spike−in proteins SGSDS Data Stable_Poly2 (1, 1, 1, 1, 1, 1, 1, 1 _ 1, 5, 6, 7, 8, 7, 5, 1)
Spike−in proteins SGSDS Data Stable_Poly2 (3, 3, 3, 3, 3, 3, 3, 3 _ 1, 5, 6, 7, 8, 7, 5, 1)
Spike−in proteins SGSDS Data Stable_Poly2 (5, 5, 5, 5, 5, 5, 5, 5 _ 1, 5, 6, 7, 8, 7, 5, 1)
Spike−in proteins SGSDS Data Stable_Poly2 (7, 7, 7, 7, 7, 7, 7, 7 _ 1, 5, 6, 7, 8, 7, 5, 1)
Spike−in proteins SGSDS Data Stable_Poly2 (1, 1, 1, 1, 1, 1, 1, 1 _ 8, 7, 6, 5, 5, 6, 7, 8)
Spike−in proteins SGSDS Data Stable_Poly2 (3, 3, 3, 3, 3, 3, 3, 3 _ 8, 7, 6, 5, 5, 6, 7, 8)
Normalized Intensity
Timepoint
Condition2
Condition1

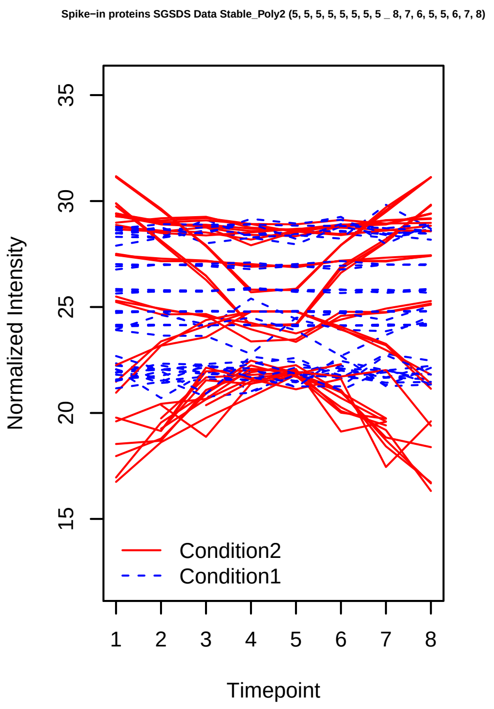
Spike−in proteins SGSDS Data Stable_Poly2 (5, 5, 5, 5, 5, 5, 5, 5 _ 8, 7, 6, 5, 5, 6, 7, 8)
Normalized Intensity
15
20
25
30
35
Condition2
Condition1
1 2 3 4 5 6 7 8
Timepoint

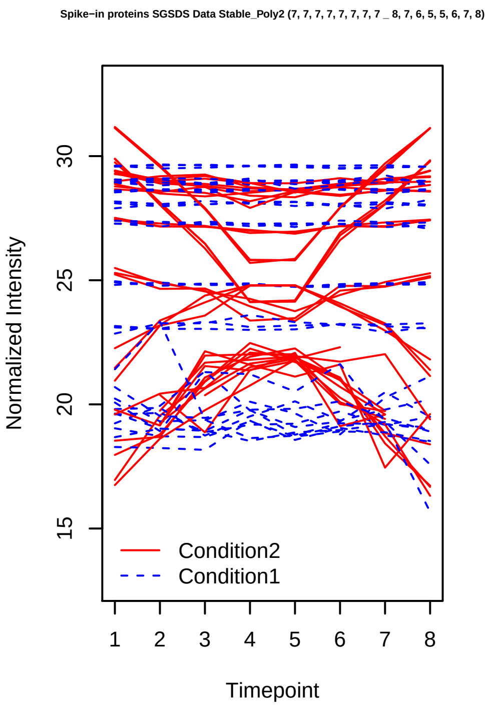
Spike−in proteins SGSDS Data Stable_Poly2 (7, 7, 7, 7, 7, 7, 7, 7 _ 8, 7, 6, 5, 5, 6, 7, 8)
Normalized Intensity
15
20
25
30
Condition2
Condition1
1 2 3 4 5 6 7 8
Timepoint

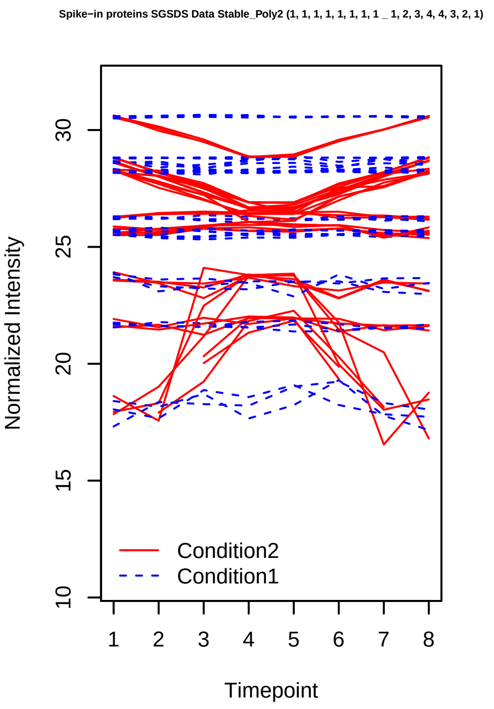
Spike−in proteins SGSDS Data Stable_Poly2 (1, 1, 1, 1, 1, 1, 1, 1 _ 1, 2, 3, 4, 4, 3, 2, 1)
Normalized Intensity
10
15
20
25
30
Condition2
Condition1
1 2 3 4 5 6 7 8
Timepoint

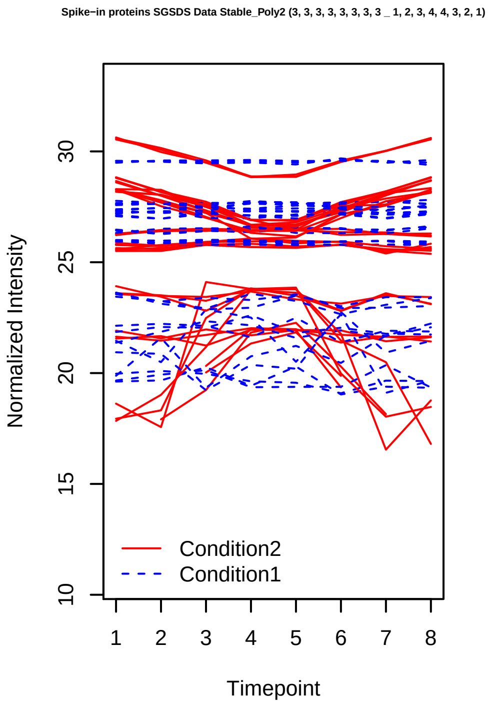
Spike−in proteins SGSDS Data Stable_Poly2 (3, 3, 3, 3, 3, 3, 3, 3 _ 1, 2, 3, 4, 4, 3, 2, 1)
Normalized Intensity
10
15
20
25
30
Condition2
Condition1
1 2 3 4 5 6 7 8
Timepoint

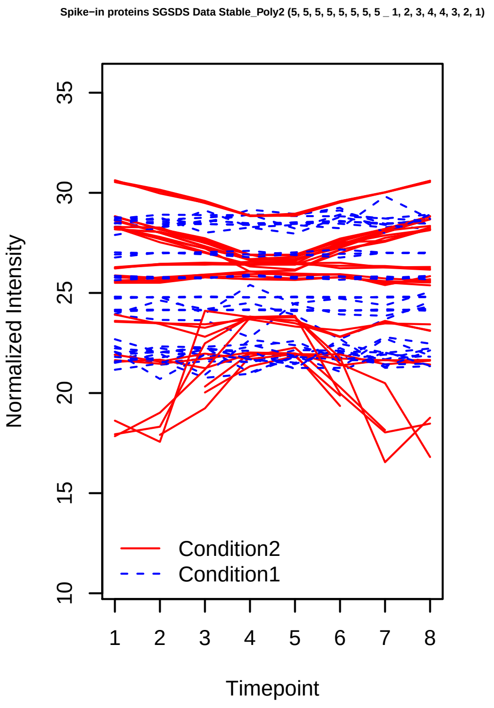
Spike−in proteins SGSDS Data Stable_Poly2 (5, 5, 5, 5, 5, 5, 5, 5 _ 1, 2, 3, 4, 4, 3, 2, 1)
Normalized Intensity
10
15
20
25
30
35
Condition2
Condition1
1 2 3 4 5 6 7 8
Timepoint

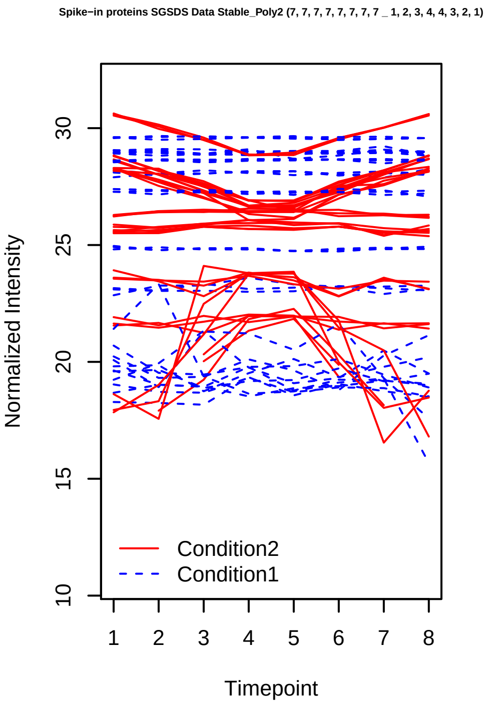
Spike−in proteins SGSDS Data Stable_Poly2 (7, 7, 7, 7, 7, 7, 7, 7 _ 1, 2, 3, 4, 4, 3, 2, 1)
Normalized Intensity
10
15
20
25
30
Condition2
Condition1
1 2 3 4 5 6 7 8
Timepoint

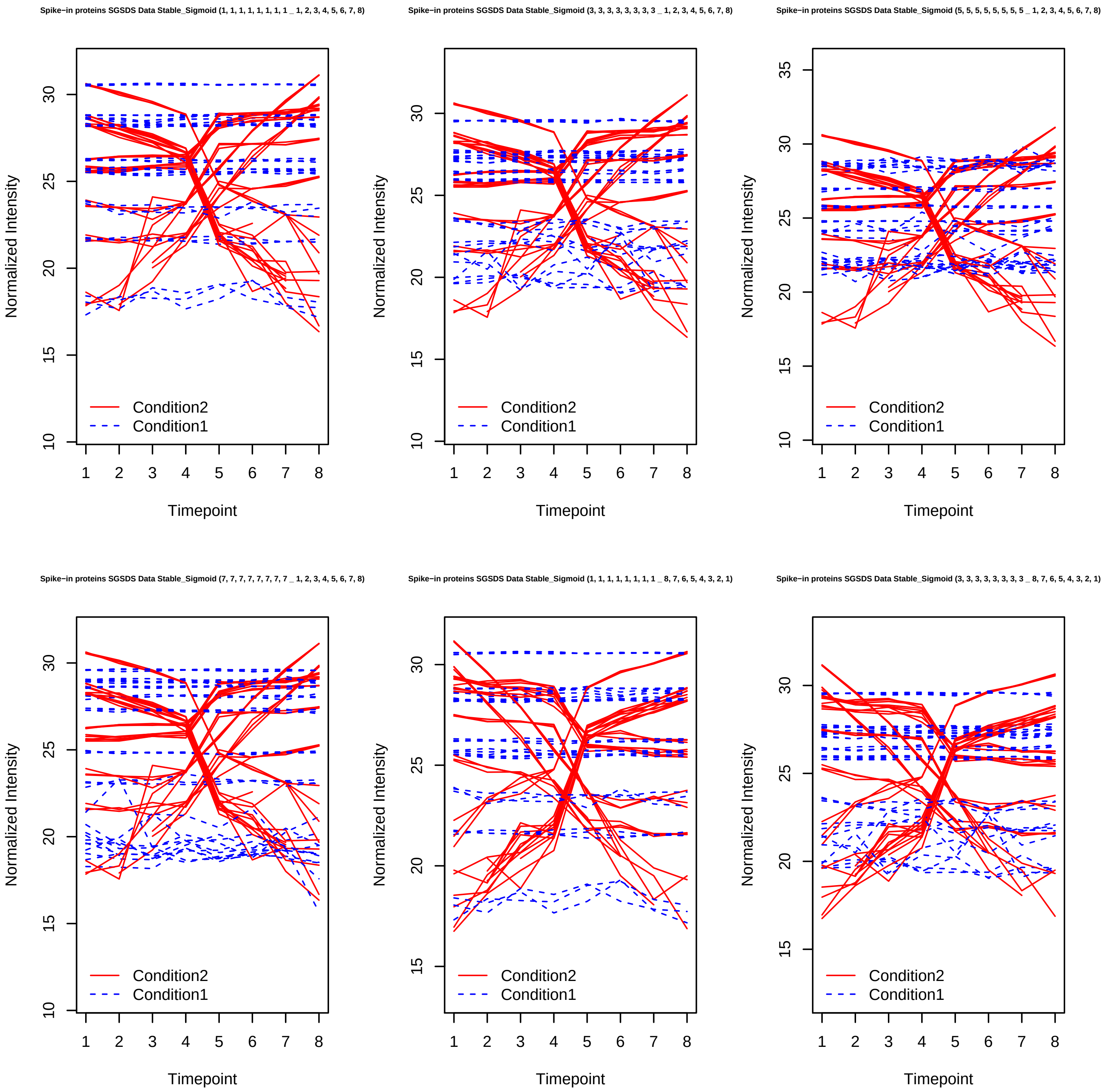
Spike−in proteins SGSDS Data Stable_Sigmoid (1, 1, 1, 1, 1, 1, 1, 1 _ 1, 2, 3, 4, 5, 6, 7, 8)
Spike−in proteins SGSDS Data Stable_Sigmoid (3, 3, 3, 3, 3, 3, 3, 3 _ 1, 2, 3, 4, 5, 6, 7, 8)
Spike−in proteins SGSDS Data Stable_Sigmoid (5, 5, 5, 5, 5, 5, 5, 5 _ 1, 2, 3, 4, 5, 6, 7, 8)
Spike−in proteins SGSDS Data Stable_Sigmoid (7, 7, 7, 7, 7, 7, 7, 7 _ 1, 2, 3, 4, 5, 6, 7, 8)
Spike−in proteins SGSDS Data Stable_Sigmoid (1, 1, 1, 1, 1, 1, 1, 1 _ 8, 7, 6, 5, 4, 3, 2, 1)
Spike−in proteins SGSDS Data Stable_Sigmoid (3, 3, 3, 3, 3, 3, 3, 3 _ 8, 7, 6, 5, 4, 3, 2, 1)
Normalized Intensity
Timepoint
Condition2
Condition1

Spike−in proteins SGSDS Data Stable_Sigmoid (5, 5, 5, 5, 5, 5, 5, 5 _ 8, 7, 6, 5, 4, 3, 2, 1)

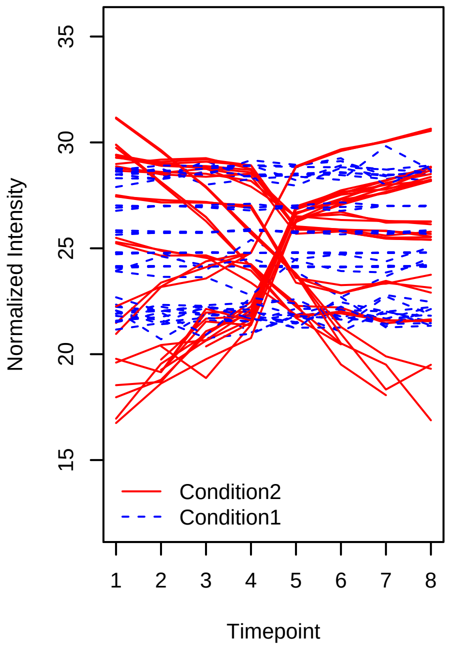


Spike−in proteins SGSDS Data Stable_Sigmoid (7, 7, 7, 7, 7, 7, 7, 7 _ 8, 7, 6, 5, 4, 3, 2, 1)

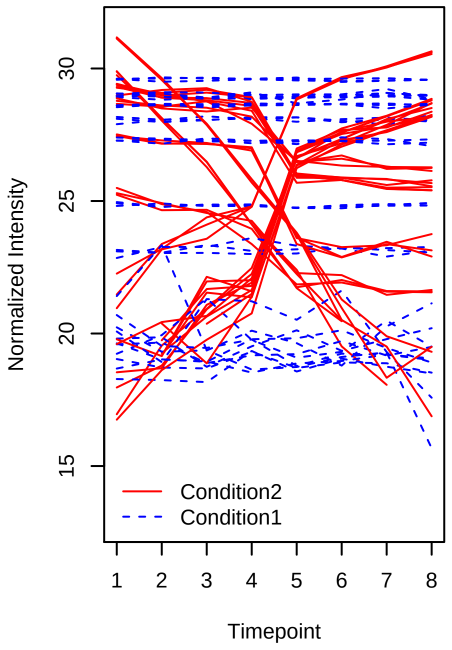


Spike−in proteins SGSDS Data Stable_Sigmoid (1, 1, 1, 1, 1, 1, 1, 1 _ 1, 1, 2, 5, 6, 7, 7, 8)

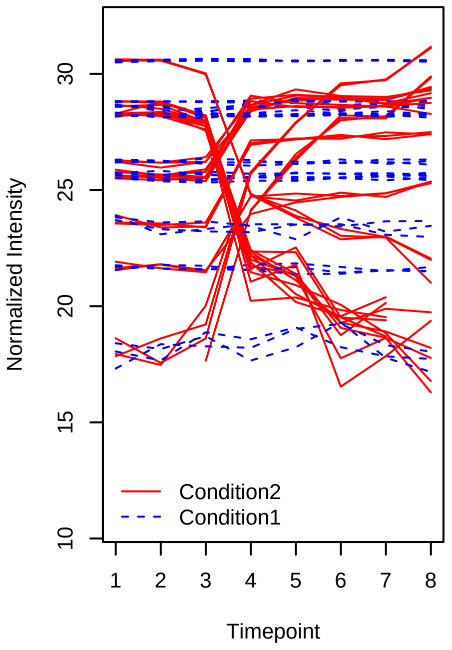


Spike−in proteins SGSDS Data Stable_Sigmoid (3, 3, 3, 3, 3, 3, 3, 3 _ 1, 1, 2, 5, 6, 7, 7, 8)

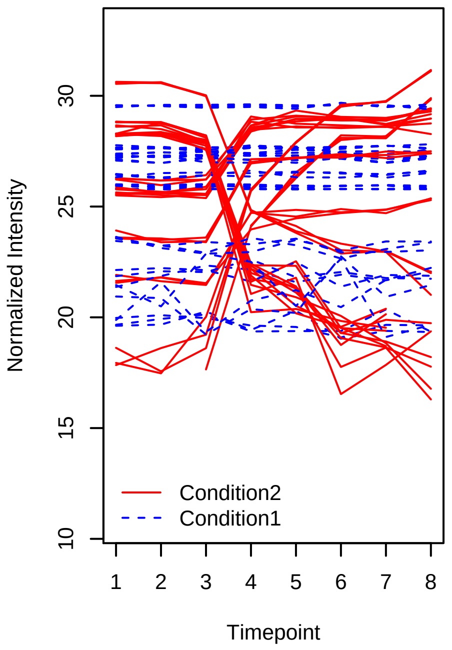


Spike−in proteins SGSDS Data Stable_Sigmoid (5, 5, 5, 5, 5, 5, 5, 5 _ 1, 1, 2, 5, 6, 7, 7, 8)

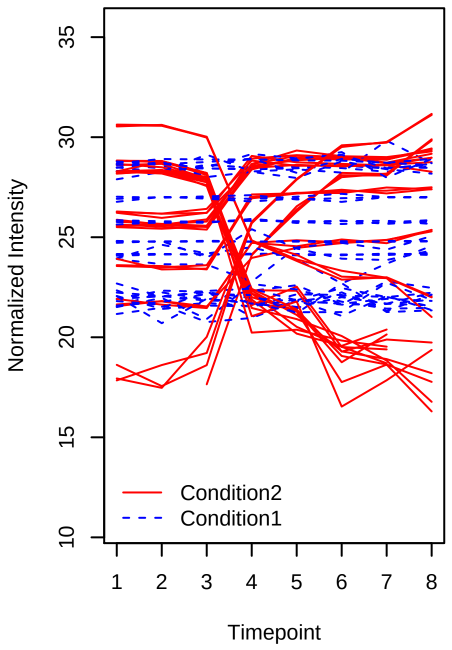


Spike−in proteins SGSDS Data Stable_Sigmoid (7, 7, 7, 7, 7, 7, 7, 7 _ 1, 1, 2, 5, 6, 7, 7, 8)

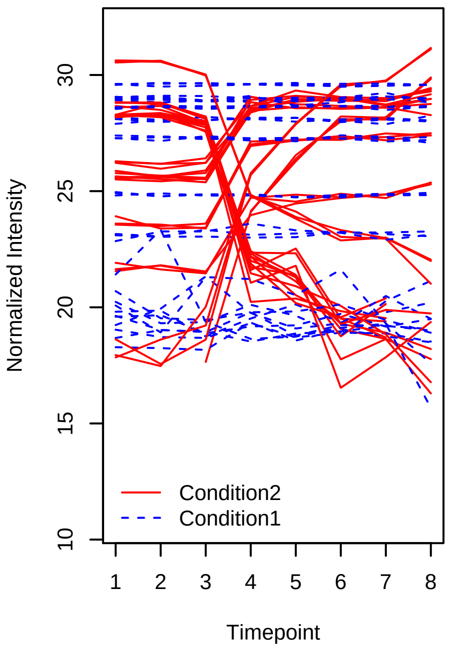

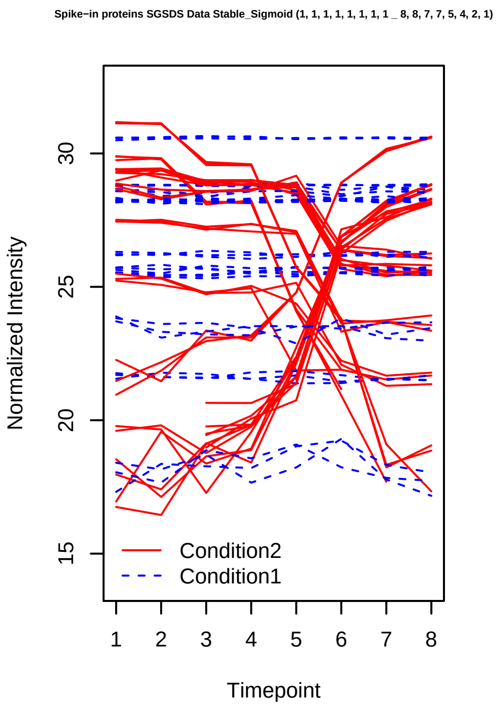
Spike−in proteins SGSDS Data Stable_Sigmoid (1, 1, 1, 1, 1, 1, 1, 1 _ 8, 8, 7, 7, 5, 4, 2, 1)
Normalized Intensity
15
20
25
30
Timepoint
1 2 3 4 5 6 7 8
Condition2
Condition1

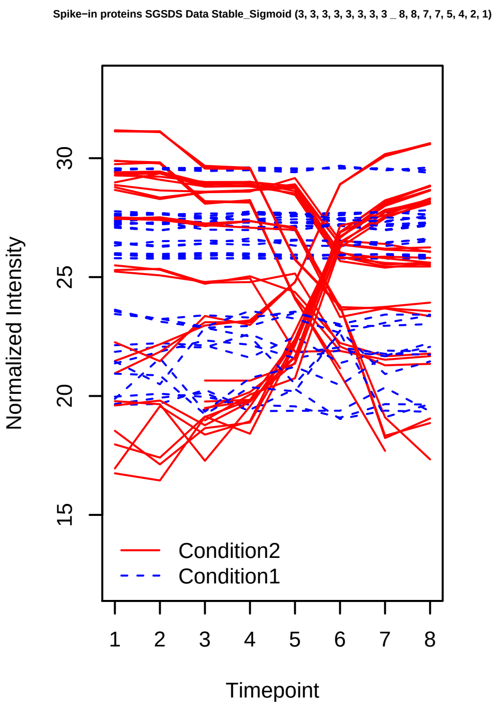
Spike−in proteins SGSDS Data Stable_Sigmoid (3, 3, 3, 3, 3, 3, 3, 3 _ 8, 8, 7, 7, 5, 4, 2, 1)
Normalized Intensity
15
20
25
30
Timepoint
1 2 3 4 5 6 7 8
Condition2
Condition1

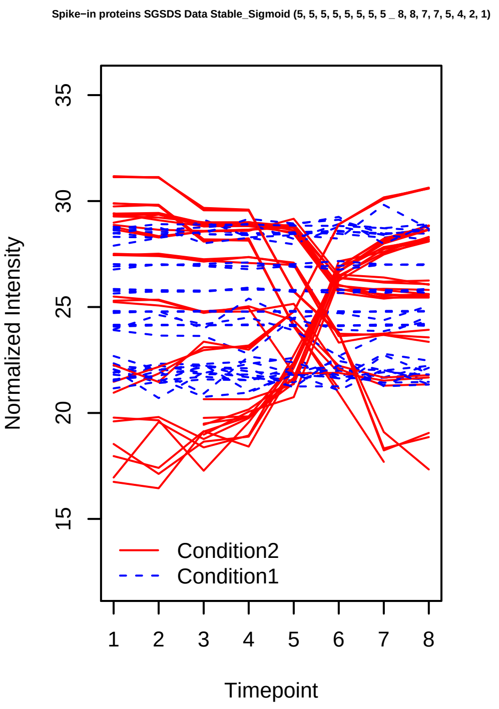
Spike−in proteins SGSDS Data Stable_Sigmoid (5, 5, 5, 5, 5, 5, 5, 5 _ 8, 8, 7, 7, 5, 4, 2, 1)
Normalized Intensity
15
20
25
30
35
Timepoint
1 2 3 4 5 6 7 8
Condition2
Condition1

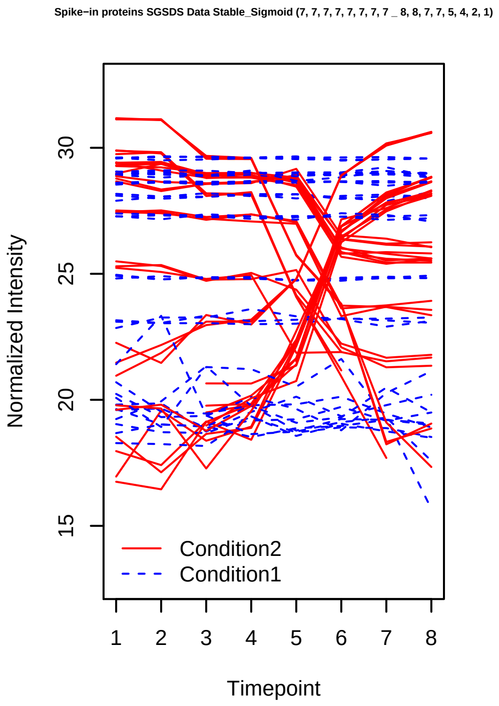
Spike−in proteins SGSDS Data Stable_Sigmoid (7, 7, 7, 7, 7, 7, 7, 7 _ 8, 8, 7, 7, 5, 4, 2, 1)
Normalized Intensity
15
20
25
30
Timepoint
1 2 3 4 5 6 7 8
Condition2
Condition1

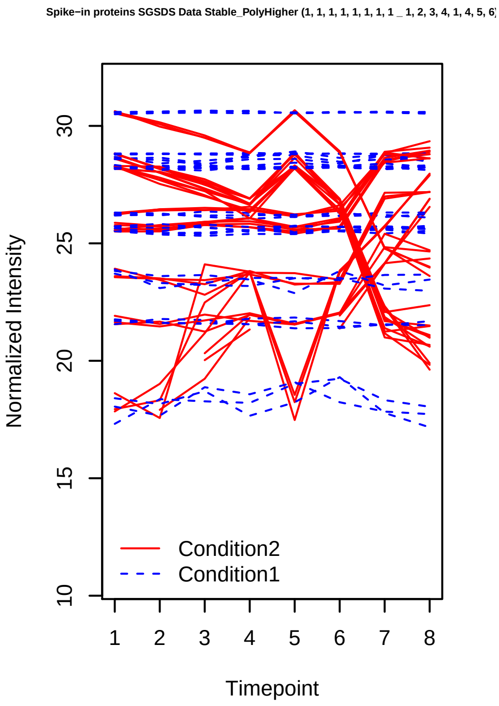
Spike−in proteins SGSDS Data Stable_PolyHigher (1, 1, 1, 1, 1, 1, 1, 1 _ 1, 2, 3, 4, 1, 4, 5, 6)
Normalized Intensity
10
15
20
25
30
Timepoint
1 2 3 4 5 6 7 8
Condition2
Condition1

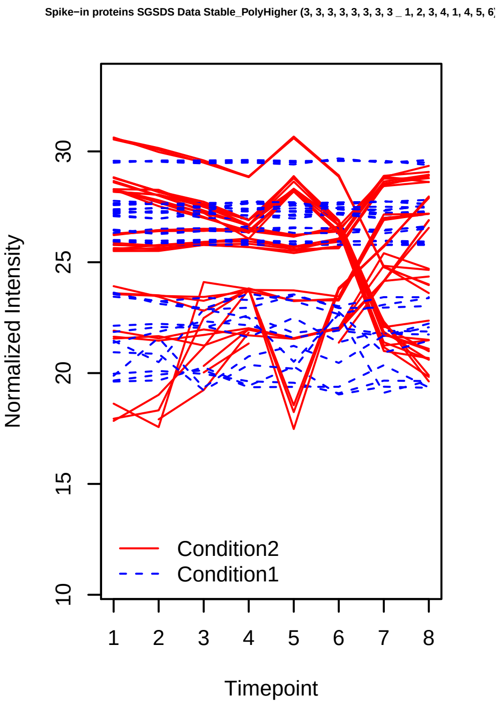
Spike−in proteins SGSDS Data Stable_PolyHigher (3, 3, 3, 3, 3, 3, 3, 3 _ 1, 2, 3, 4, 1, 4, 5, 6)
Normalized Intensity
10
15
20
25
30
Timepoint
1 2 3 4 5 6 7 8
Condition2
Condition1

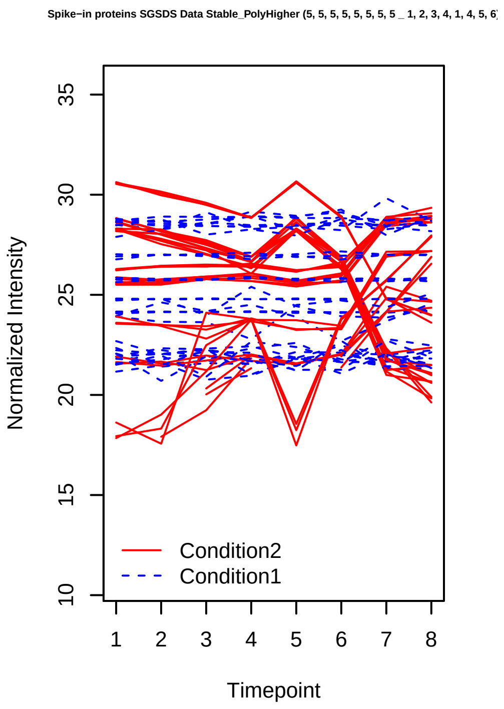
Spike−in proteins SGSDS Data Stable_PolyHigher (5, 5, 5, 5, 5, 5, 5, 5 _ 1, 2, 3, 4, 1, 4, 5, 6)
Normalized Intensity
Timepoint
Condition2
Condition1

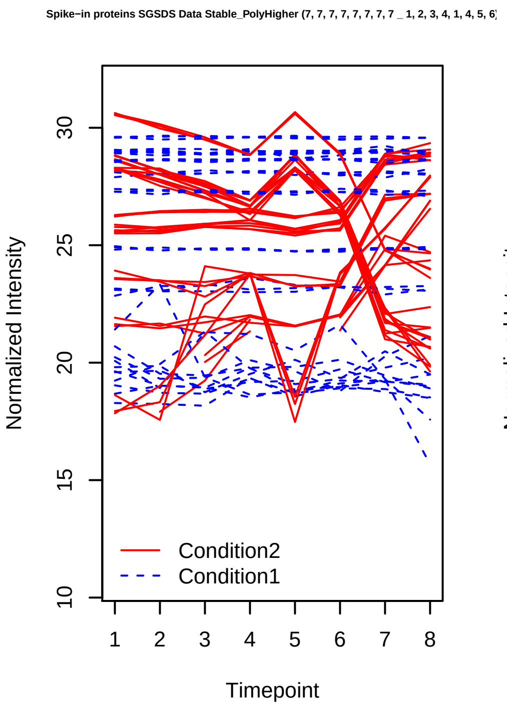
Spike−in proteins SGSDS Data Stable_PolyHigher (7, 7, 7, 7, 7, 7, 7, 7 _ 1, 2, 3, 4, 1, 4, 5, 6)
Normalized Intensity
Timepoint
Condition2
Condition1

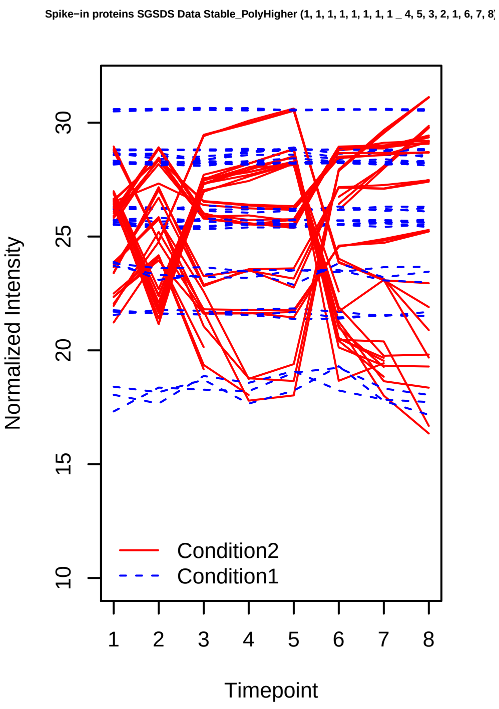
Spike−in proteins SGSDS Data Stable_PolyHigher (1, 1, 1, 1, 1, 1, 1, 1 _ 4, 5, 3, 2, 1, 6, 7, 8)
Normalized Intensity
Timepoint
Condition2
Condition1

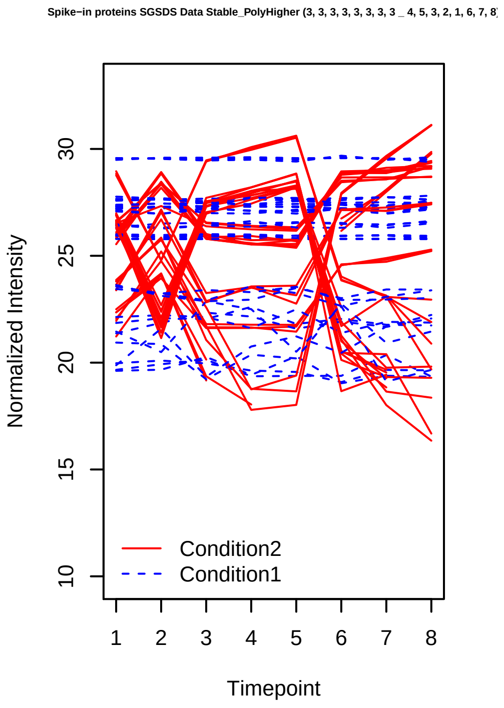
Spike−in proteins SGSDS Data Stable_PolyHigher (3, 3, 3, 3, 3, 3, 3, 3 _ 4, 5, 3, 2, 1, 6, 7, 8)
Normalized Intensity
Timepoint
Condition2
Condition1

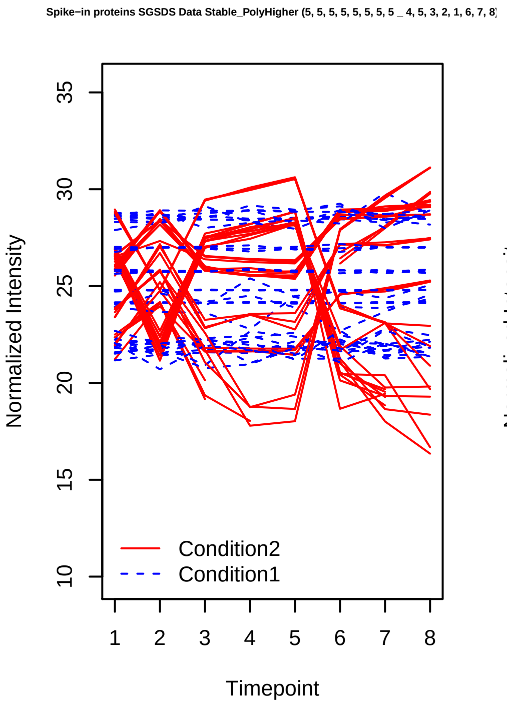
Spike−in proteins SGSDS Data Stable_PolyHigher (5, 5, 5, 5, 5, 5, 5, 5 _ 4, 5, 3, 2, 1, 6, 7, 8)
Normalized Intensity
Timepoint
Condition2
Condition1

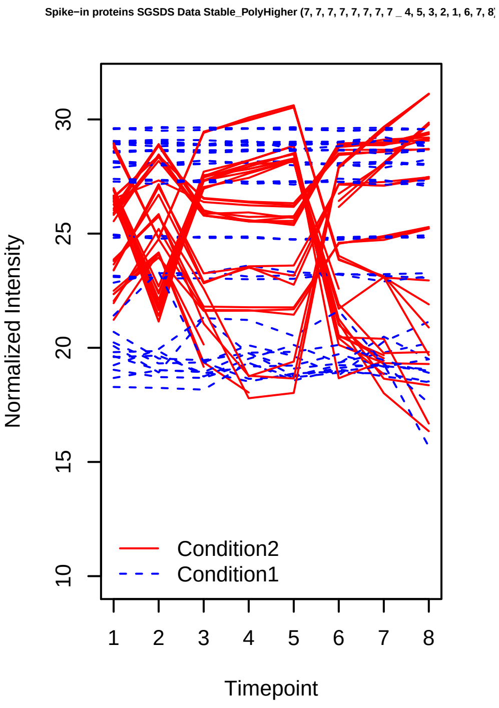
Spike−in proteins SGSDS Data Stable_PolyHigher (7, 7, 7, 7, 7, 7, 7, 7 _ 4, 5, 3, 2, 1, 6, 7, 8)
Normalized Intensity
Timepoint
Condition2
Condition1

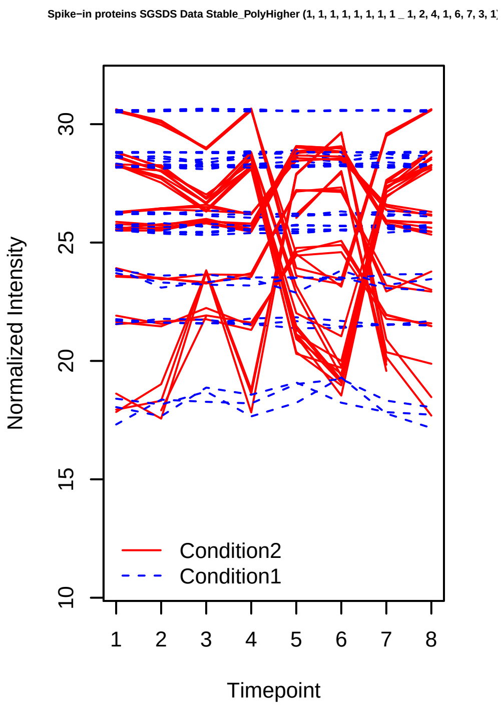
Spike−in proteins SGSDS Data Stable_PolyHigher (1, 1, 1, 1, 1, 1, 1, 1 _ 1, 2, 4, 1, 6, 7, 3, 1)
Normalized Intensity
10
15
20
25
30
1 2 3 4 5 6 7 8
Timepoint
Condition2
Condition1

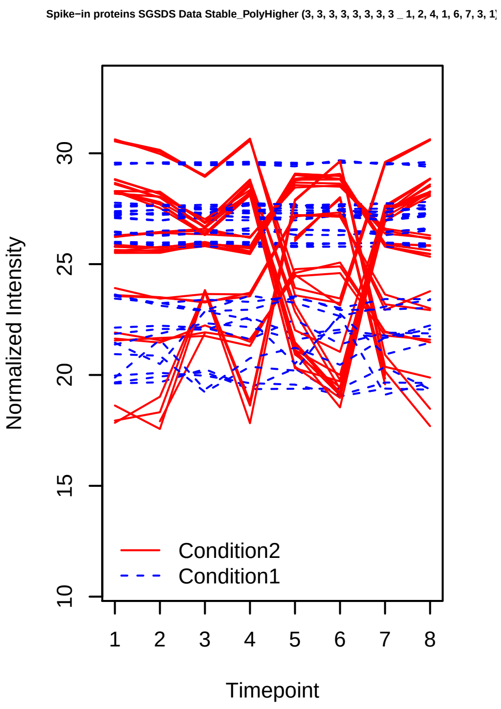
Spike−in proteins SGSDS Data Stable_PolyHigher (3, 3, 3, 3, 3, 3, 3, 3 _ 1, 2, 4, 1, 6, 7, 3, 1)
Normalized Intensity
10
15
20
25
30
1 2 3 4 5 6 7 8
Timepoint
Condition2
Condition1

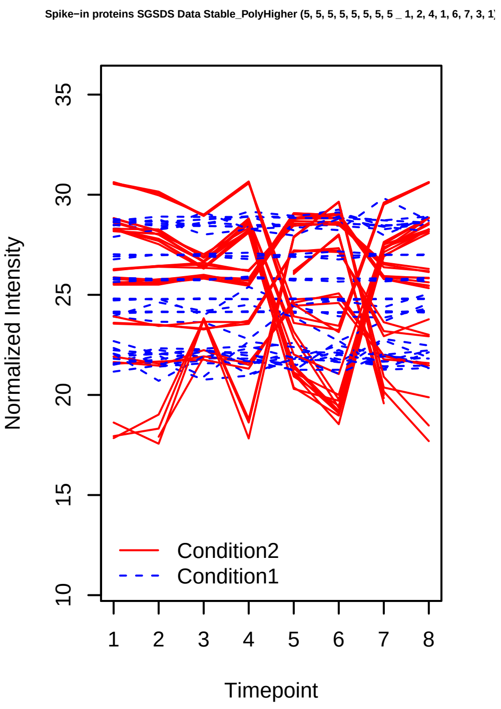
Spike−in proteins SGSDS Data Stable_PolyHigher (5, 5, 5, 5, 5, 5, 5, 5 _ 1, 2, 4, 1, 6, 7, 3, 1)
Normalized Intensity
10
15
20
25
30
35
1 2 3 4 5 6 7 8
Timepoint
Condition2
Condition1

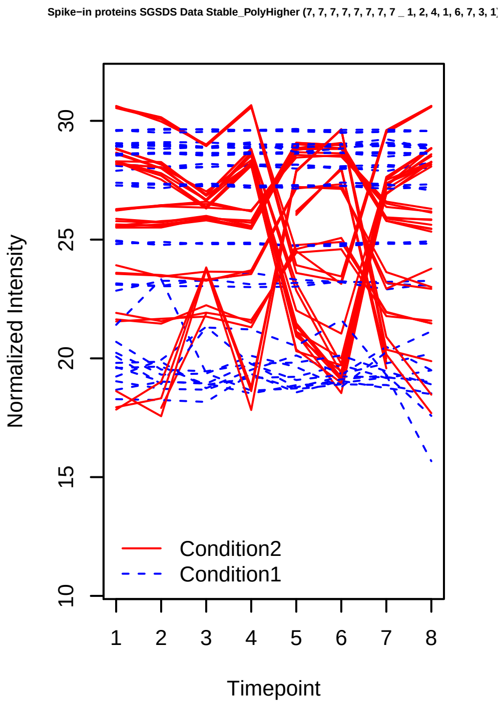
Spike−in proteins SGSDS Data Stable_PolyHigher (7, 7, 7, 7, 7, 7, 7, 7 _ 1, 2, 4, 1, 6, 7, 3, 1)
Normalized Intensity
10
15
20
25
30
1 2 3 4 5 6 7 8
Timepoint
Condition2
Condition1

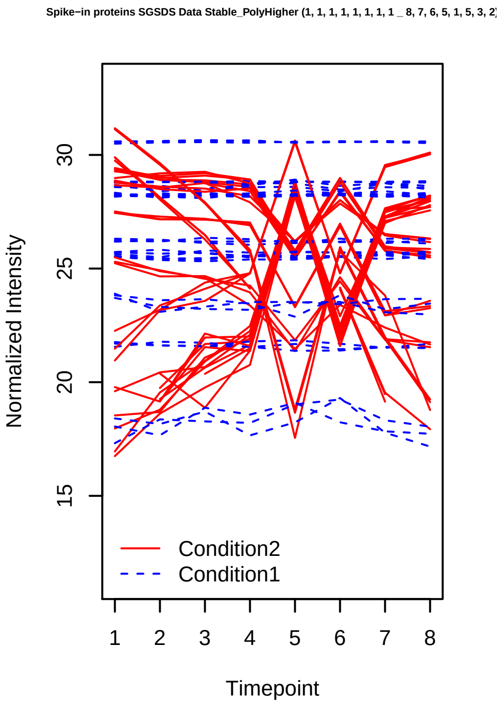
Spike−in proteins SGSDS Data Stable_PolyHigher (1, 1, 1, 1, 1, 1, 1, 1 _ 8, 7, 6, 5, 1, 5, 3, 2)
Normalized Intensity
15
20
25
30
1 2 3 4 5 6 7 8
Timepoint
Condition2
Condition1

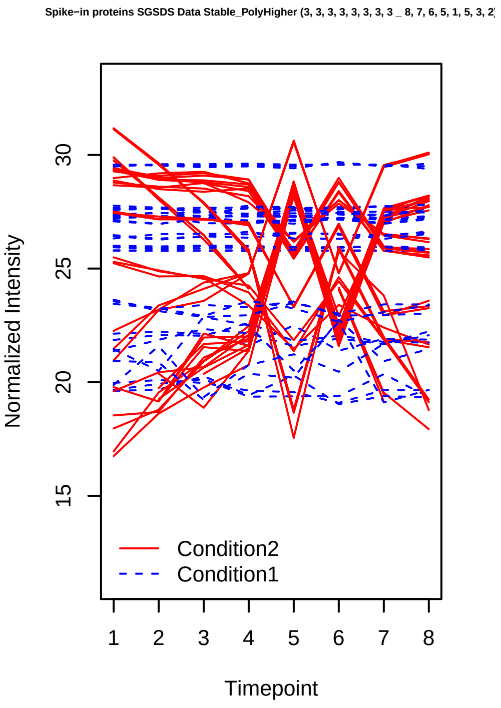
Spike−in proteins SGSDS Data Stable_PolyHigher (3, 3, 3, 3, 3, 3, 3, 3 _ 8, 7, 6, 5, 1, 5, 3, 2)
Normalized Intensity
15
20
25
30
1 2 3 4 5 6 7 8
Timepoint
Condition2
Condition1

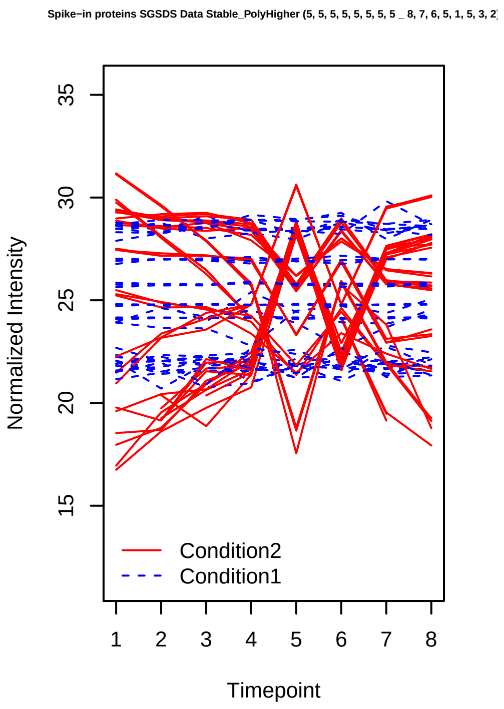
Spike−in proteins SGSDS Data Stable_PolyHigher (5, 5, 5, 5, 5, 5, 5, 5 _ 8, 7, 6, 5, 1, 5, 3, 2)
Normalized Intensity
Timepoint
Condition2
Condition1

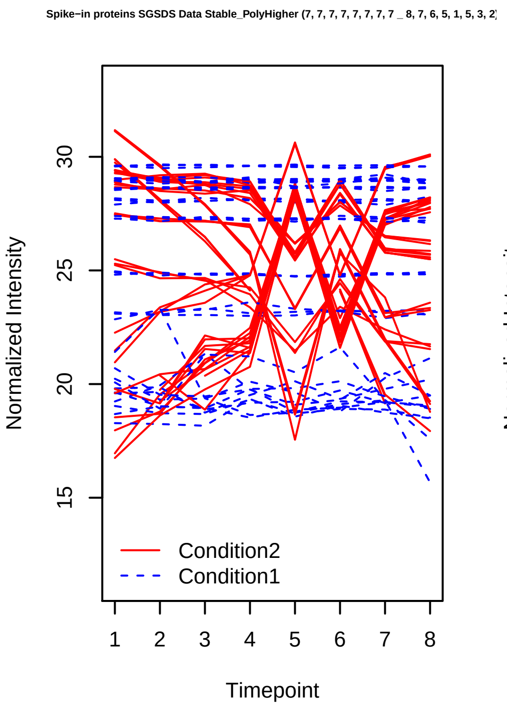
Spike−in proteins SGSDS Data Stable_PolyHigher (7, 7, 7, 7, 7, 7, 7, 7 _ 8, 7, 6, 5, 1, 5, 3, 2)
Normalized Intensity
Timepoint
Condition2
Condition1

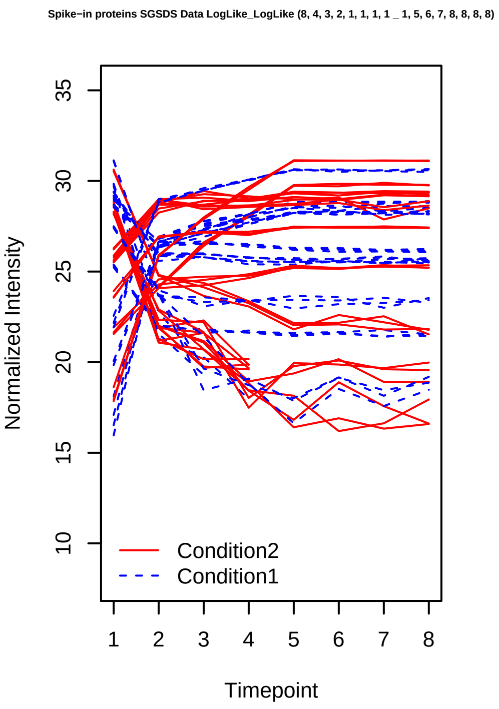
Spike−in proteins SGSDS Data LogLike_LogLike (8, 4, 3, 2, 1, 1, 1, 1 _ 1, 5, 6, 7, 8, 8, 8, 8)
Normalized Intensity
Timepoint
Condition2
Condition1

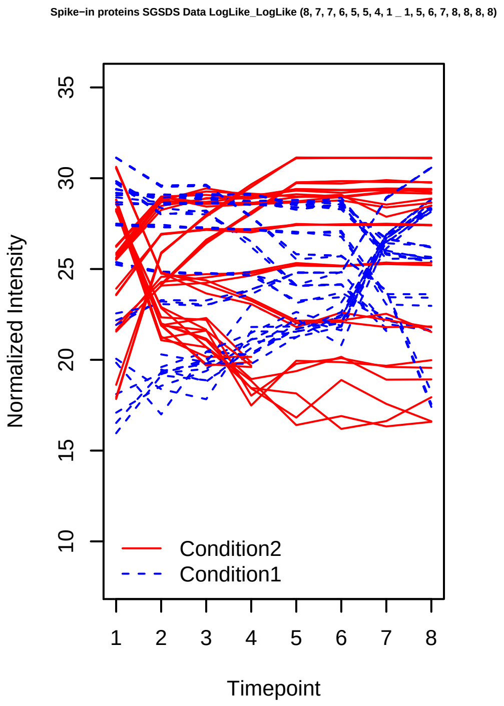
Spike−in proteins SGSDS Data LogLike_LogLike (8, 7, 7, 6, 5, 5, 4, 1 _ 1, 5, 6, 7, 8, 8, 8, 8)
Normalized Intensity
Timepoint
Condition2
Condition1

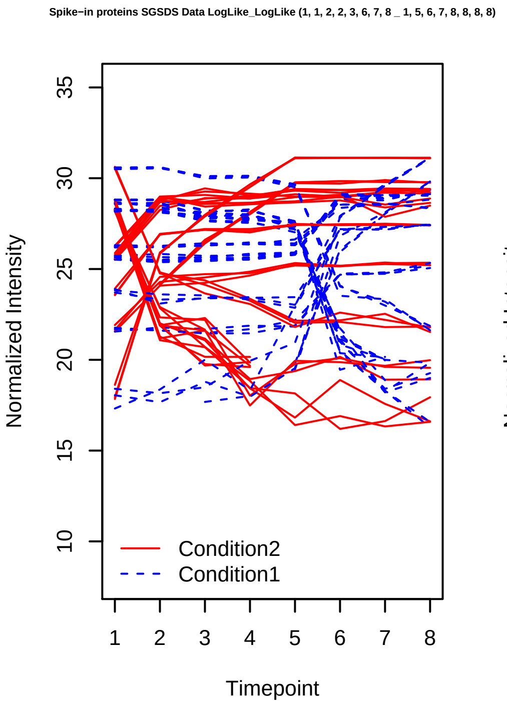
Spike−in proteins SGSDS Data LogLike_LogLike (1, 1, 2, 2, 3, 6, 7, 8 _ 1, 5, 6, 7, 8, 8, 8, 8)
Normalized Intensity
Timepoint
Condition2
Condition1

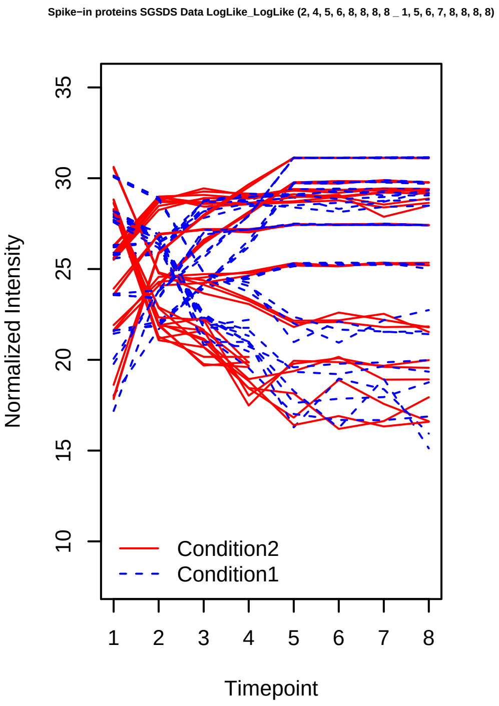
Spike−in proteins SGSDS Data LogLike_LogLike (2, 4, 5, 6, 8, 8, 8, 8 _ 1, 5, 6, 7, 8, 8, 8, 8)
Normalized Intensity
Timepoint
Condition2
Condition1

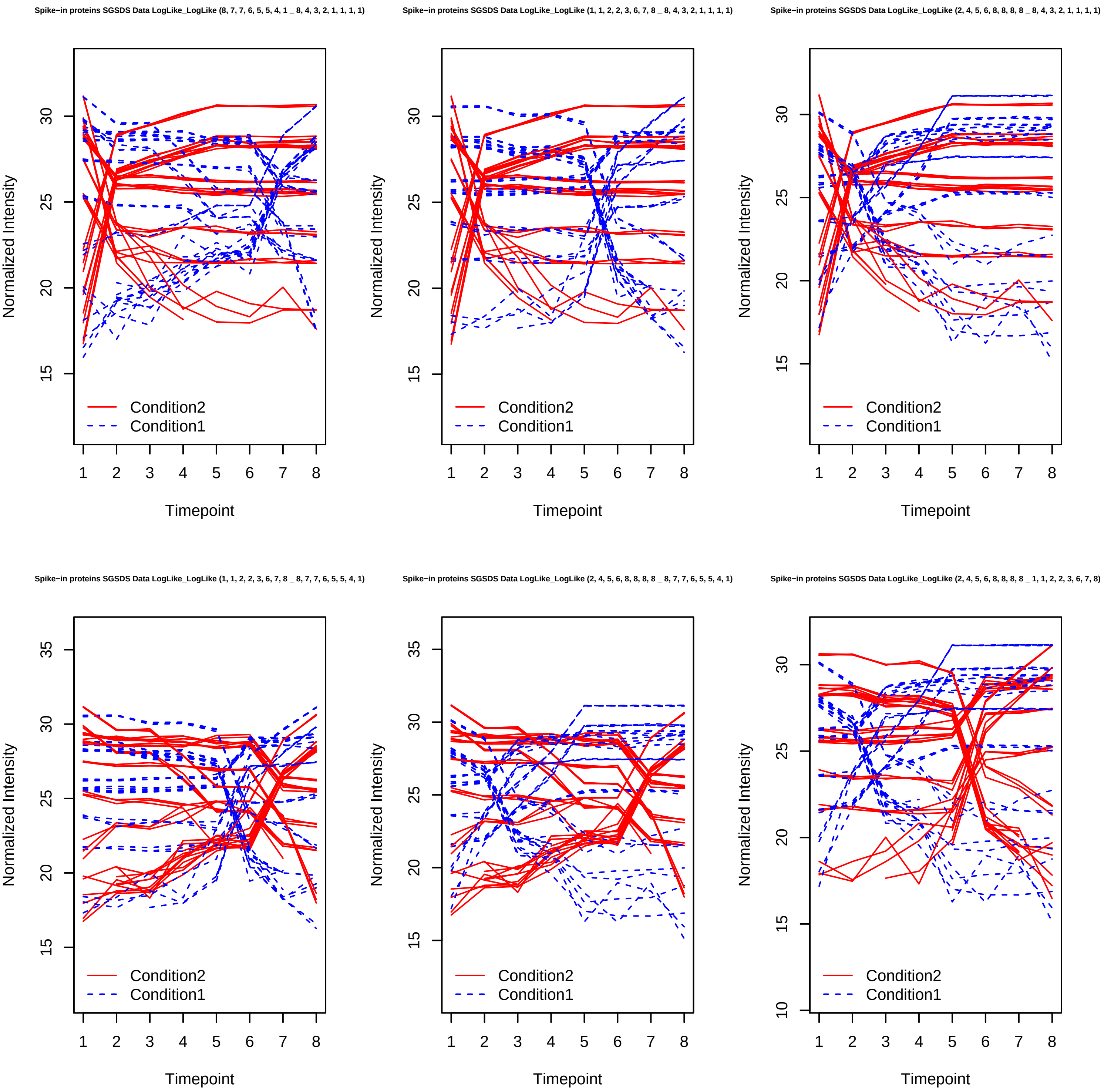
Spike−in proteins SGSDS Data LogLike_LogLike (8, 7, 7, 6, 5, 5, 4, 1 _ 8, 4, 3, 2, 1, 1, 1, 1)
Spike−in proteins SGSDS Data LogLike_LogLike (1, 1, 2, 2, 3, 6, 7, 8 _ 8, 4, 3, 2, 1, 1, 1, 1)
Spike−in proteins SGSDS Data LogLike_LogLike (2, 4, 5, 6, 8, 8, 8, 8 _ 8, 4, 3, 2, 1, 1, 1, 1)
Spike−in proteins SGSDS Data LogLike_LogLike (1, 1, 2, 2, 3, 6, 7, 8 _ 8, 7, 7, 6, 5, 5, 4, 1)
Spike−in proteins SGSDS Data LogLike_LogLike (2, 4, 5, 6, 8, 8, 8, 8 _ 8, 7, 7, 6, 5, 5, 4, 1)
Spike−in proteins SGSDS Data LogLike_LogLike (2, 4, 5, 6, 8, 8, 8, 8 _ 1, 1, 2, 2, 3, 6, 7, 8)
Normalized Intensity
Timepoint
Condition2
Condition1

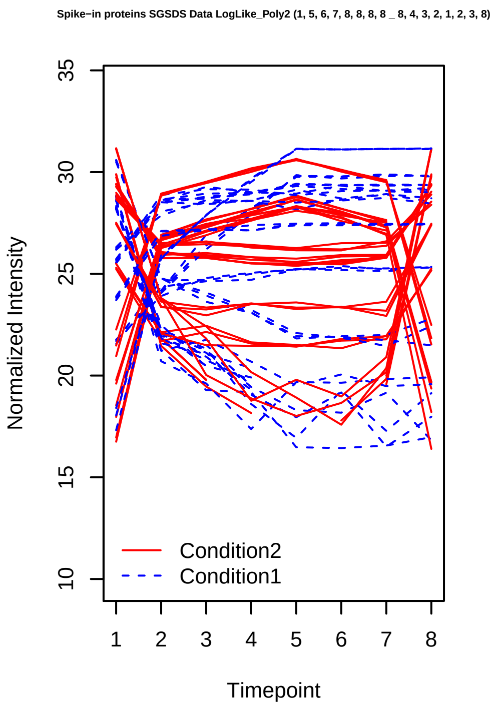
Spike−in proteins SGSDS Data LogLike_Poly2 (1, 5, 6, 7, 8, 8, 8, 8 _ 8, 4, 3, 2, 1, 2, 3, 8)
Normalized Intensity
Timepoint
Condition2
Condition1

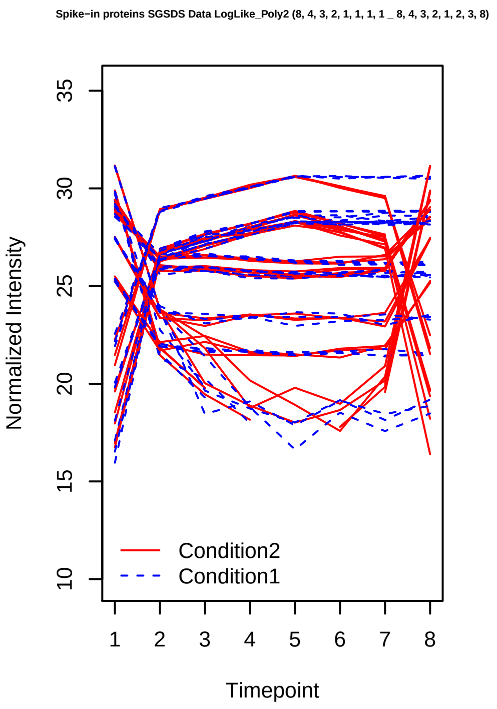
Spike−in proteins SGSDS Data LogLike_Poly2 (8, 4, 3, 2, 1, 1, 1, 1 _ 8, 4, 3, 2, 1, 2, 3, 8)
Normalized Intensity
Timepoint
Condition2
Condition1

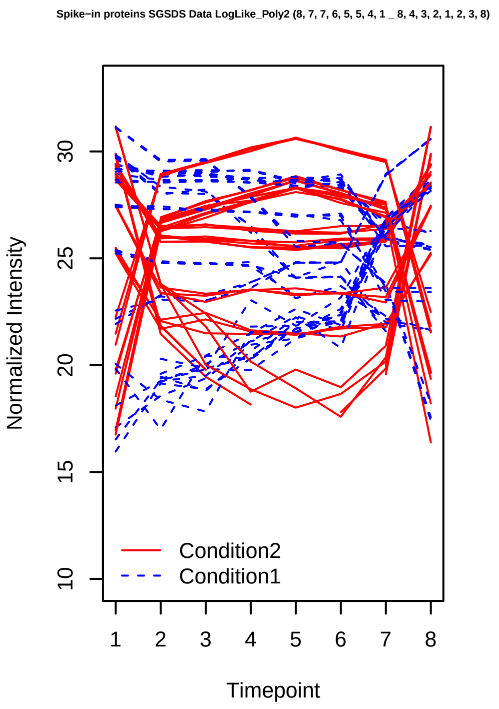
Spike−in proteins SGSDS Data LogLike_Poly2 (8, 7, 7, 6, 5, 5, 4, 1 _ 8, 4, 3, 2, 1, 2, 3, 8)
Normalized Intensity
Timepoint
Condition2
Condition1

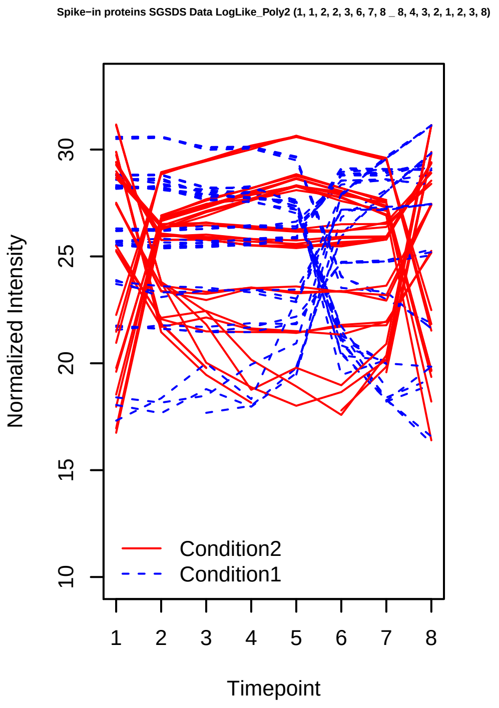
Spike−in proteins SGSDS Data LogLike_Poly2 (1, 1, 2, 2, 3, 6, 7, 8 _ 8, 4, 3, 2, 1, 2, 3, 8)
Normalized Intensity
Timepoint
Condition2
Condition1

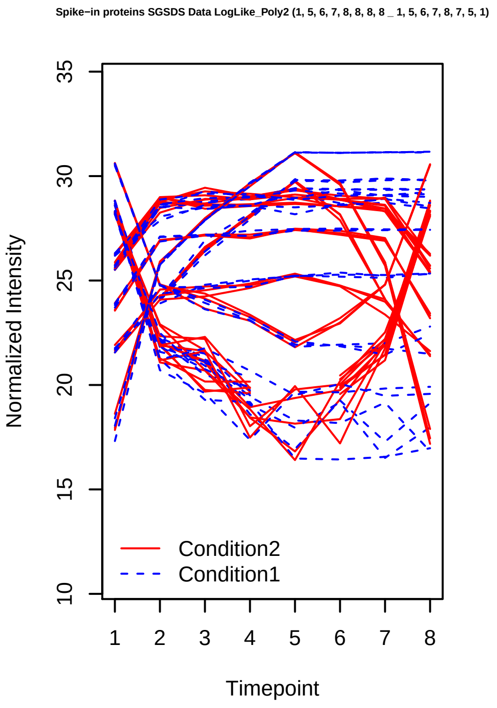
Spike−in proteins SGSDS Data LogLike_Poly2 (1, 5, 6, 7, 8, 8, 8, 8 _ 1, 5, 6, 7, 8, 7, 5, 1)
Normalized Intensity
Timepoint
Condition2
Condition1

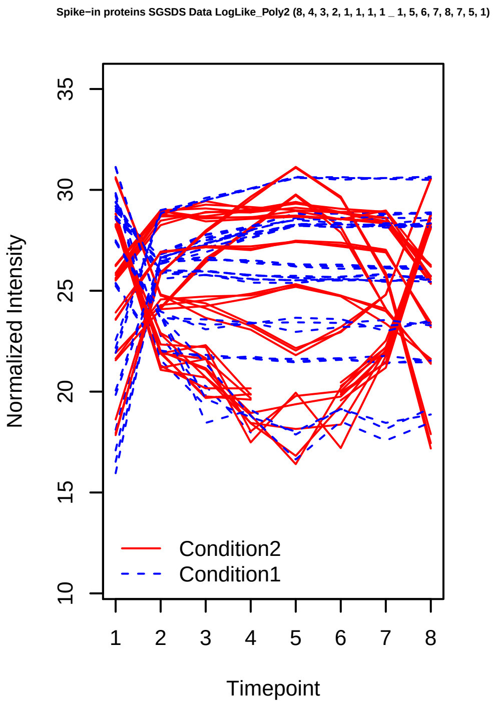
Spike−in proteins SGSDS Data LogLike_Poly2 (8, 4, 3, 2, 1, 1, 1, 1 _ 1, 5, 6, 7, 8, 7, 5, 1)
Normalized Intensity
Timepoint
Condition2
Condition1

Spike−in proteins SGSDS Data LogLike_Poly2 (8, 7, 7, 6, 5, 5, 4, 1 _ 1, 5, 6, 7, 8, 7, 5, 1)

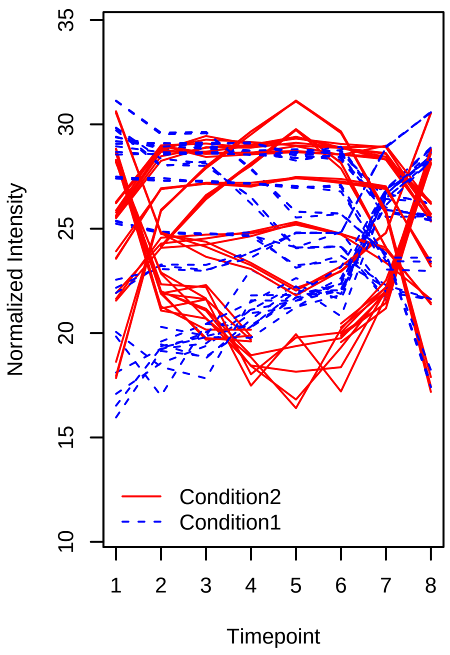


Spike−in proteins SGSDS Data LogLike_Poly2 (1, 1, 2, 2, 3, 6, 7, 8 _ 1, 5, 6, 7, 8, 7, 5, 1)

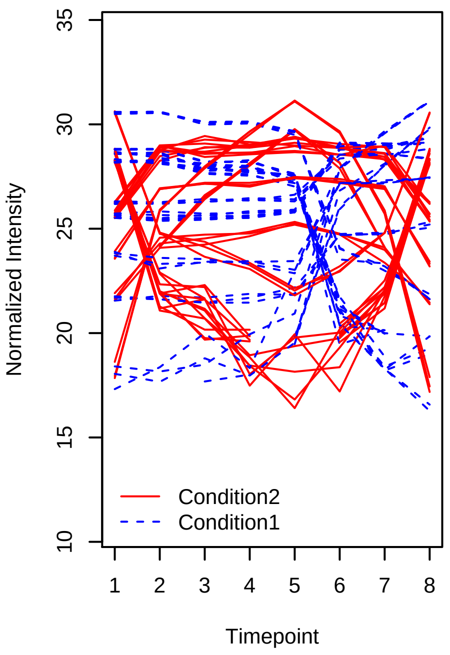


Spike−in proteins SGSDS Data LogLike_Poly2 (1, 5, 6, 7, 8, 8, 8, 8 _ 8, 7, 6, 5, 5, 6, 7, 8)

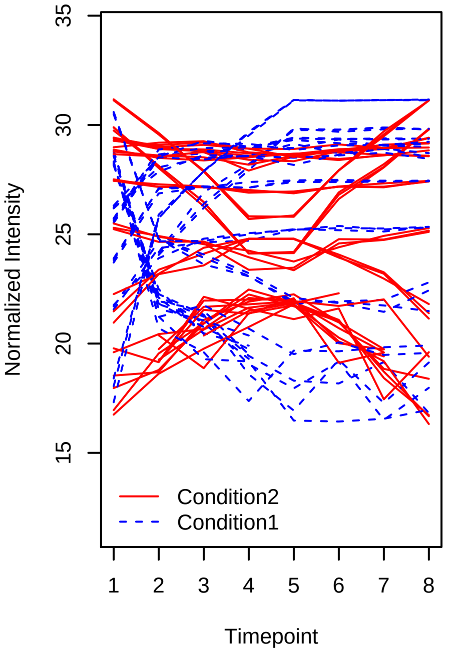


Spike−in proteins SGSDS Data LogLike_Poly2 (8, 4, 3, 2, 1, 1, 1, 1 _ 8, 7, 6, 5, 5, 6, 7, 8)

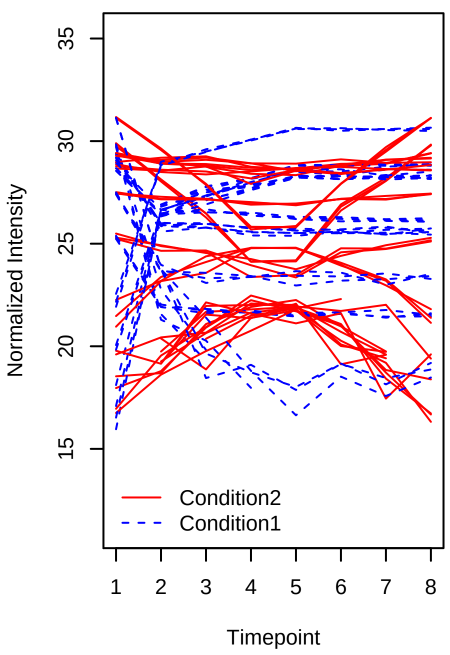


Spike−in proteins SGSDS Data LogLike_Poly2 (8, 7, 7, 6, 5, 5, 4, 1 _ 8, 7, 6, 5, 5, 6, 7, 8)

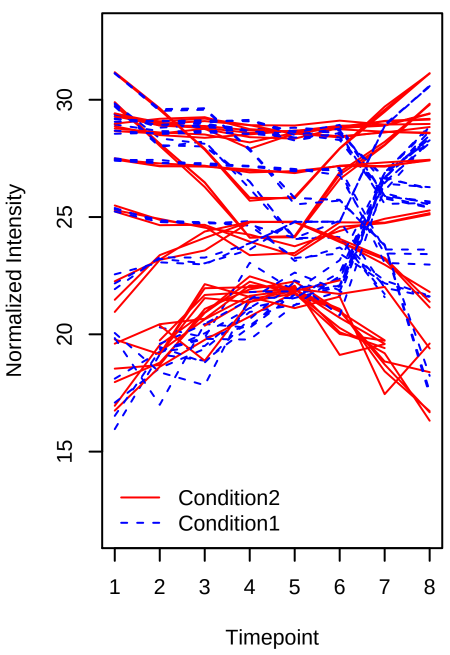


Spike−in proteins SGSDS Data LogLike_Poly2 (1, 1, 2, 2, 3, 6, 7, 8 _ 8, 7, 6, 5, 5, 6, 7, 8)

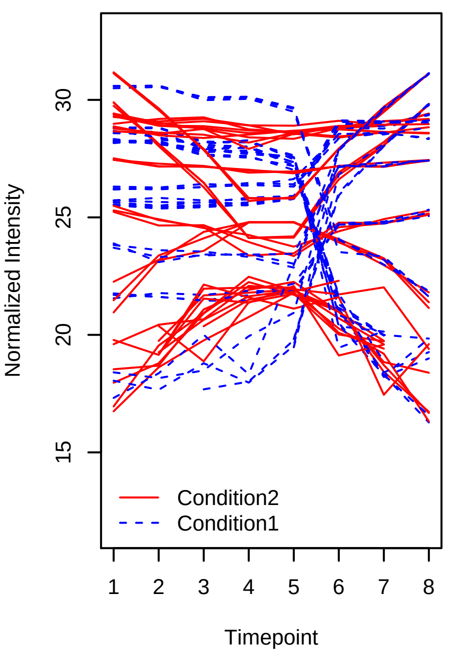

Spike−in proteins SGSDS Data LogLike_Poly2 (1, 5, 6, 7, 8, 8, 8, 8 _ 1, 2, 3, 4, 4, 3, 2, 1)

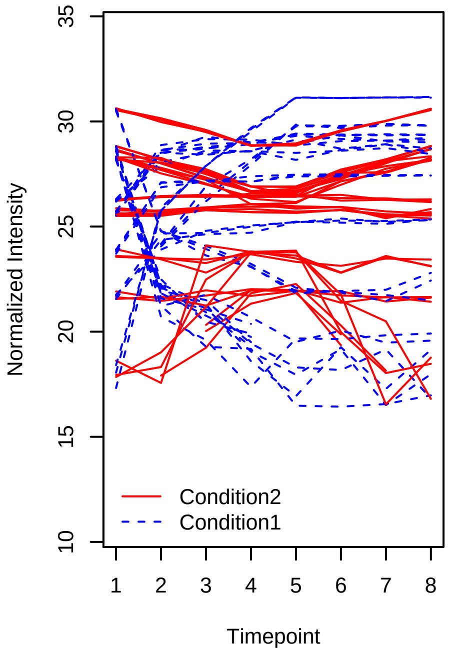


Spike−in proteins SGSDS Data LogLike_Poly2 (8, 4, 3, 2, 1, 1, 1, 1 _ 1, 2, 3, 4, 4, 3, 2, 1)

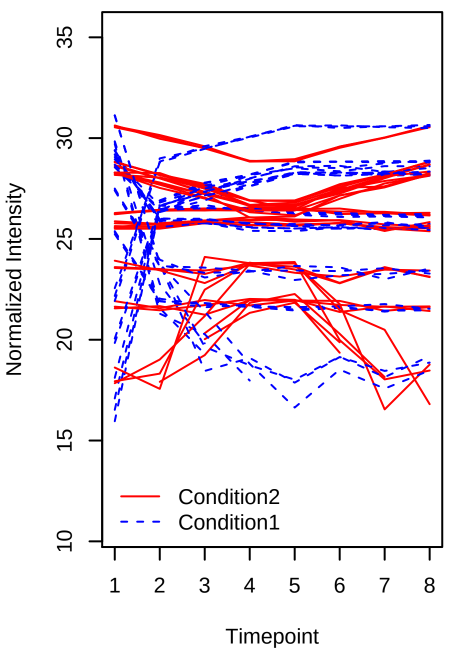


Spike−in proteins SGSDS Data LogLike_Poly2 (8, 7, 7, 6, 5, 5, 4, 1 _ 1, 2, 3, 4, 4, 3, 2, 1)

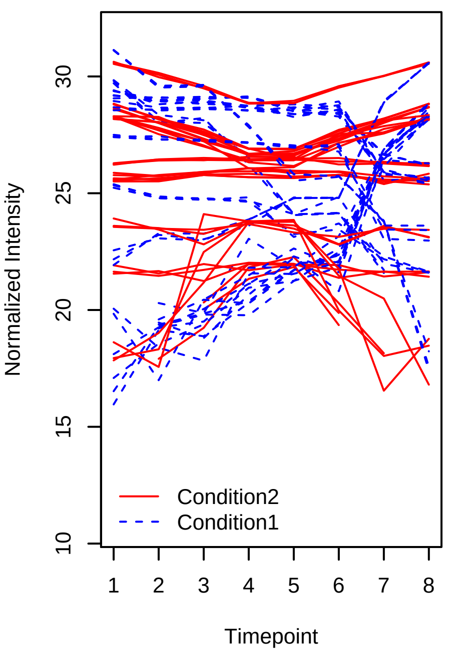


Spike−in proteins SGSDS Data LogLike_Poly2 (1, 1, 2, 2, 3, 6, 7, 8 _ 1, 2, 3, 4, 4, 3, 2, 1)

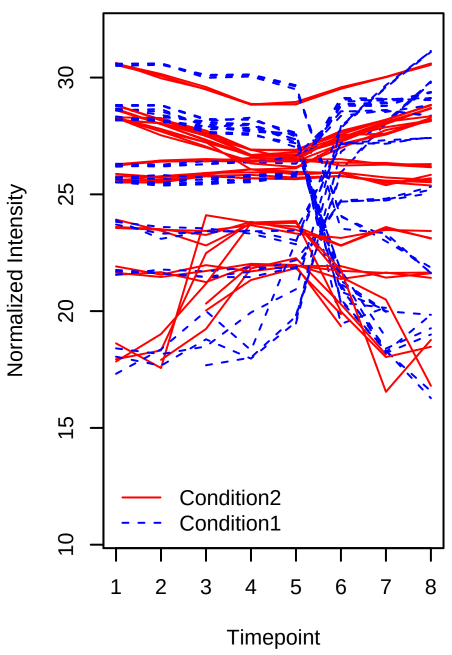


Spike−in proteins SGSDS Data LogLike_Sigmoid (1, 5, 6, 7, 8, 8, 8, 8 _ 1, 2, 3, 4, 5, 6, 7, 8)

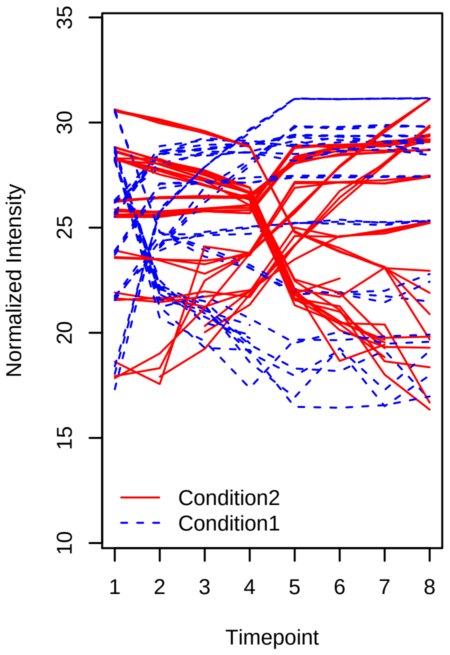


Spike−in proteins SGSDS Data LogLike_Sigmoid (8, 4, 3, 2, 1, 1, 1, 1 _ 1, 2, 3, 4, 5, 6, 7, 8)

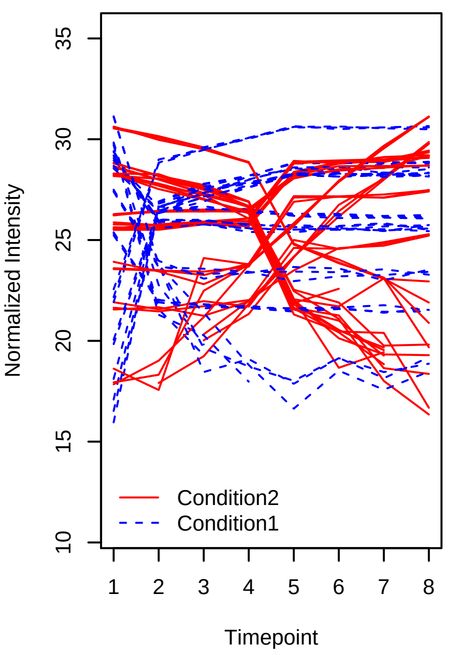

Spike−in proteins SGSDS Data LogLike_Sigmoid (8, 7, 7, 6, 5, 5, 4, 1 _ 1, 2, 3, 4, 5, 6, 7, 8)

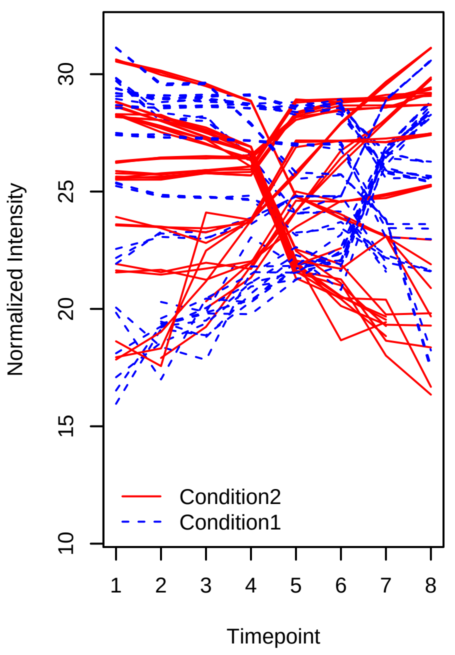


Spike−in proteins SGSDS Data LogLike_Sigmoid (1, 1, 2, 2, 3, 6, 7, 8 _ 1, 2, 3, 4, 5, 6, 7, 8)

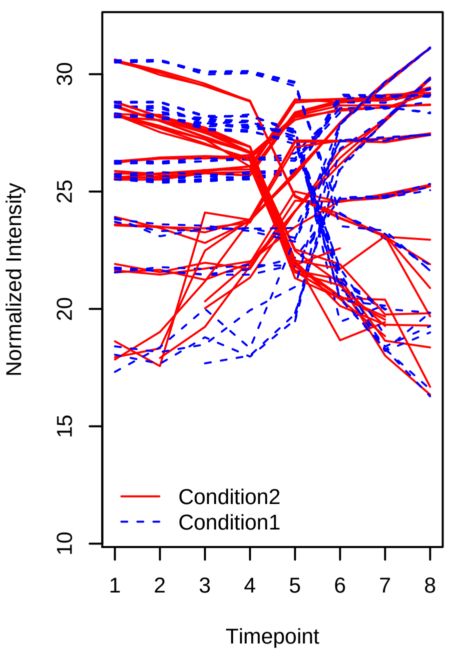


Spike−in proteins SGSDS Data LogLike_Sigmoid (1, 5, 6, 7, 8, 8, 8, 8 _ 8, 7, 6, 5, 4, 3, 2, 1)

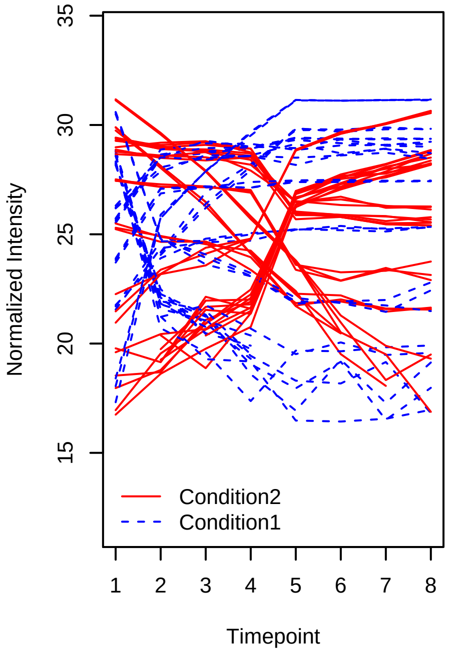


Spike−in proteins SGSDS Data LogLike_Sigmoid (8, 4, 3, 2, 1, 1, 1, 1 _ 8, 7, 6, 5, 4, 3, 2, 1)

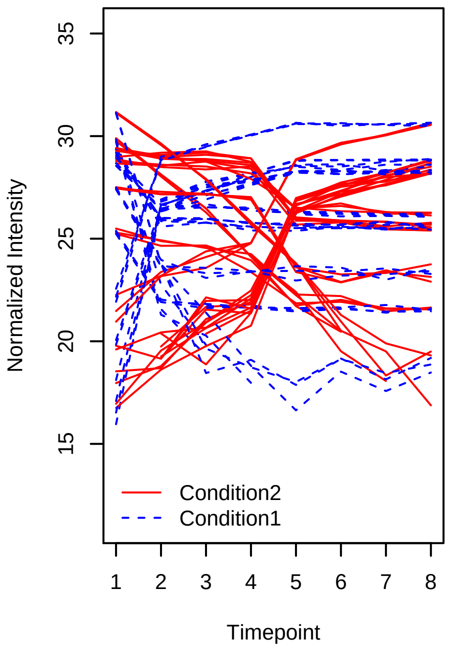


Spike−in proteins SGSDS Data LogLike_Sigmoid (8, 7, 7, 6, 5, 5, 4, 1 _ 8, 7, 6, 5, 4, 3, 2, 1)

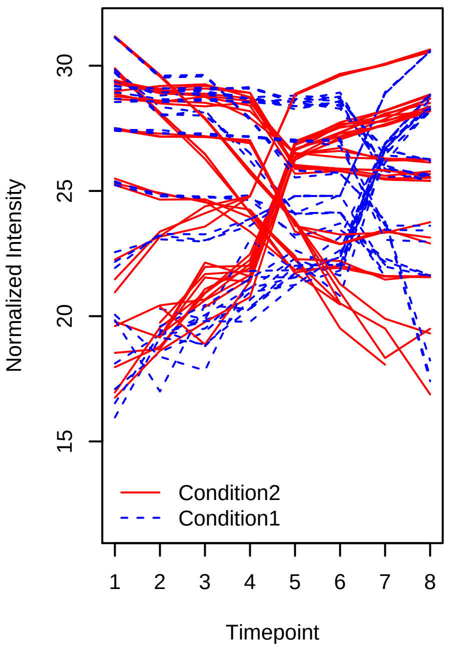


Spike−in proteins SGSDS Data LogLike_Sigmoid (1, 1, 2, 2, 3, 6, 7, 8 _ 8, 7, 6, 5, 4, 3, 2, 1)

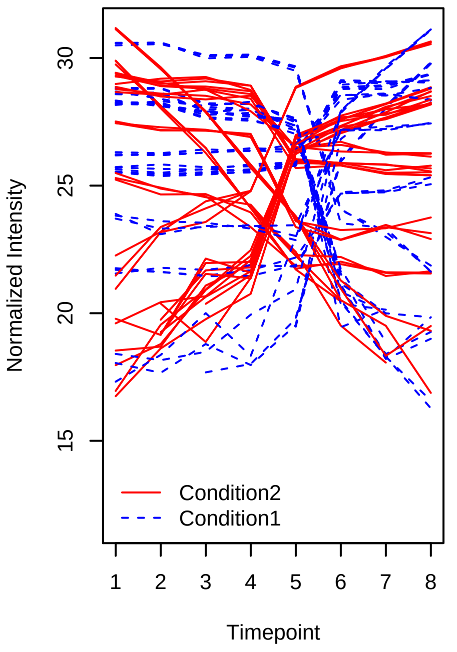

Spike−in proteins SGSDS Data LogLike_Sigmoid (1, 5, 6, 7, 8, 8, 8, 8 _ 1, 1, 2, 5, 6, 7, 7, 8)

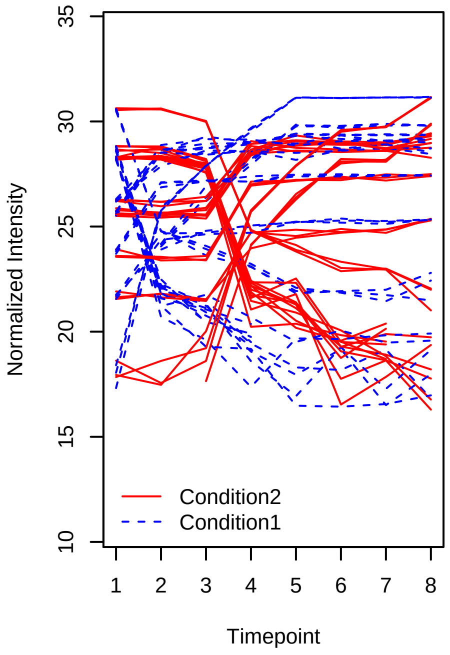


Spike−in proteins SGSDS Data LogLike_Sigmoid (8, 4, 3, 2, 1, 1, 1, 1 _ 1, 1, 2, 5, 6, 7, 7, 8)

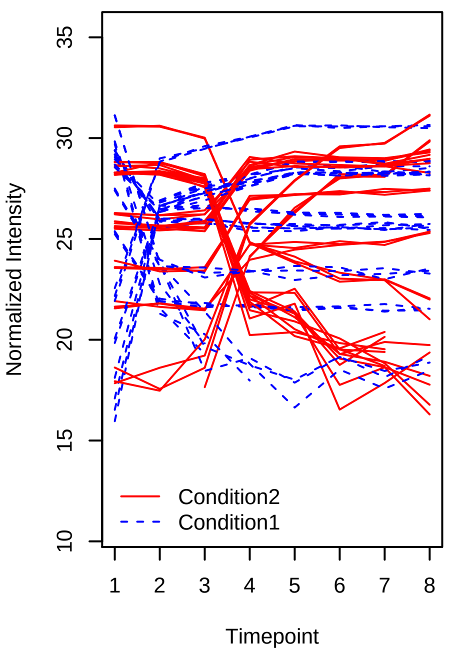


Spike−in proteins SGSDS Data LogLike_Sigmoid (8, 7, 7, 6, 5, 5, 4, 1 _ 1, 1, 2, 5, 6, 7, 7, 8)

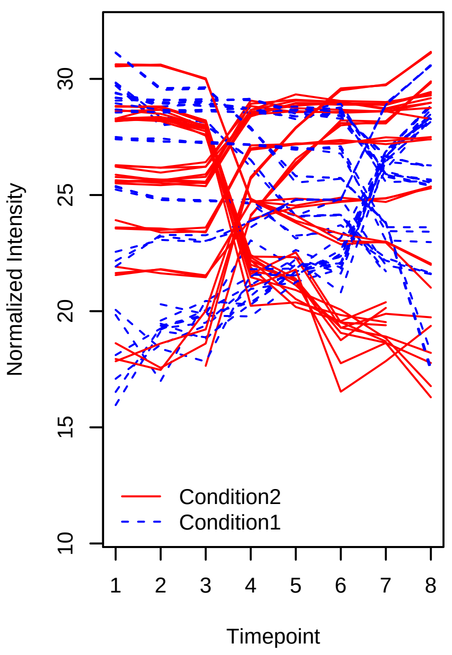


Spike−in proteins SGSDS Data LogLike_Sigmoid (1, 1, 2, 2, 3, 6, 7, 8 _ 1, 1, 2, 5, 6, 7, 7, 8)

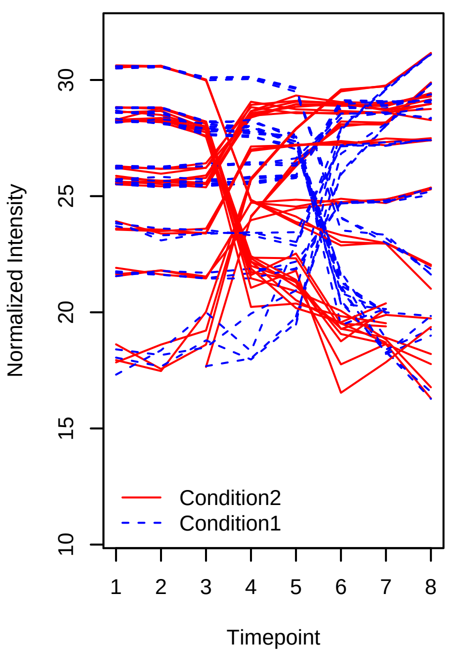


Spike−in proteins SGSDS Data LogLike_Sigmoid (1, 5, 6, 7, 8, 8, 8, 8 _ 8, 8, 7, 7, 5, 4, 2, 1)

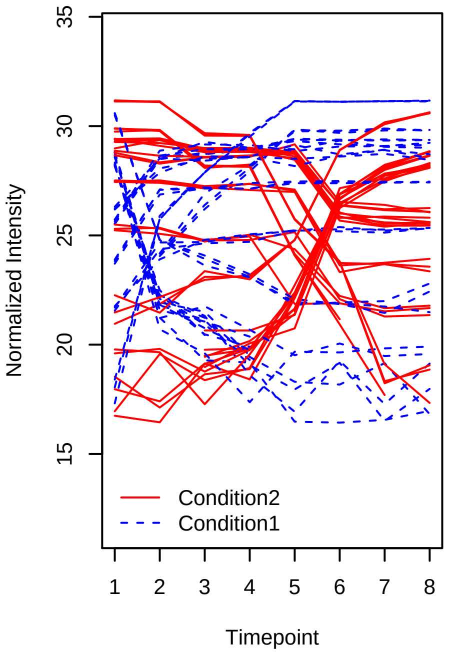


Spike−in proteins SGSDS Data LogLike_Sigmoid (8, 4, 3, 2, 1, 1, 1, 1 _ 8, 8, 7, 7, 5, 4, 2, 1)

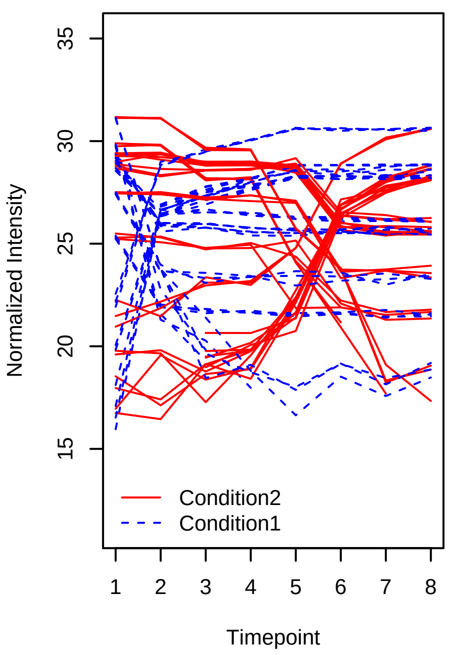

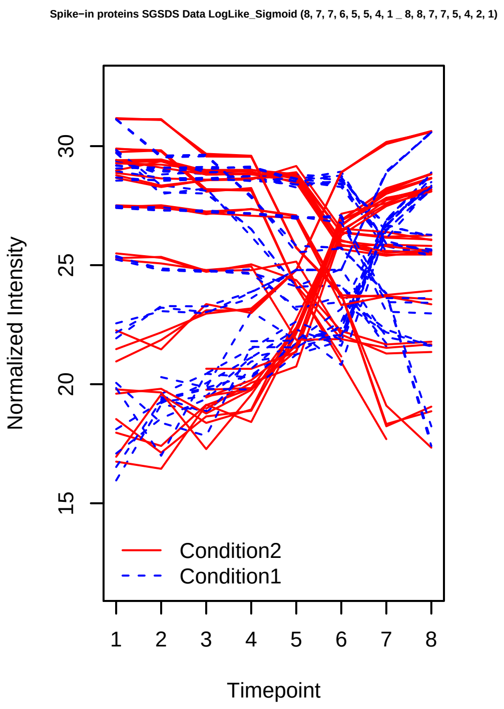
Spike−in proteins SGSDS Data LogLike_Sigmoid (8, 7, 7, 6, 5, 5, 4, 1 _ 8, 8, 7, 7, 5, 4, 2, 1)
Normalized Intensity
Timepoint
Condition2
Condition1
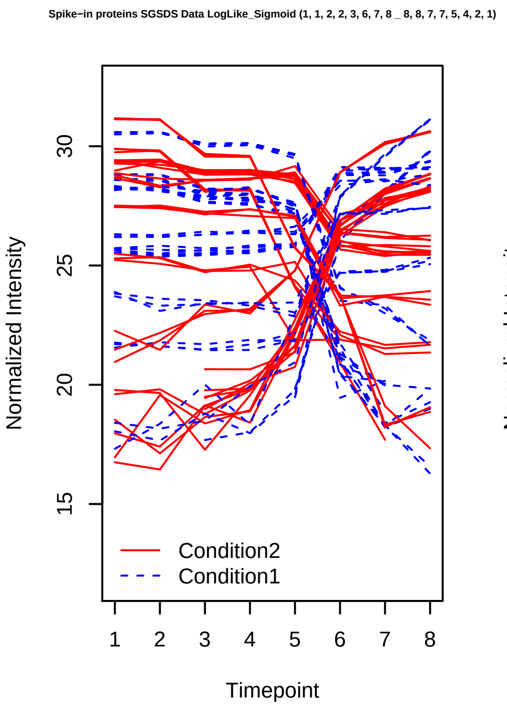
Spike−in proteins SGSDS Data LogLike_Sigmoid (1, 1, 2, 2, 3, 6, 7, 8 _ 8, 8, 7, 7, 5, 4, 2, 1)
Normalized Intensity
Timepoint
Condition2
Condition1
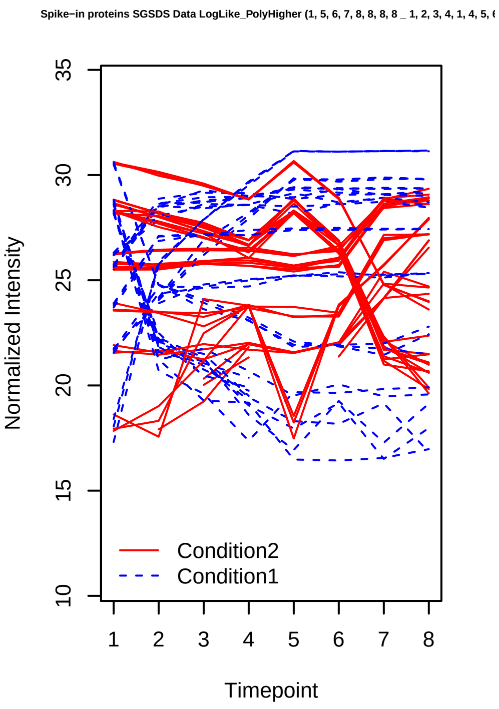
Spike−in proteins SGSDS Data LogLike_PolyHigher (1, 5, 6, 7, 8, 8, 8, 8 _ 1, 2, 3, 4, 1, 4, 5,
Normalized Intensity
Timepoint
Condition2
Condition1
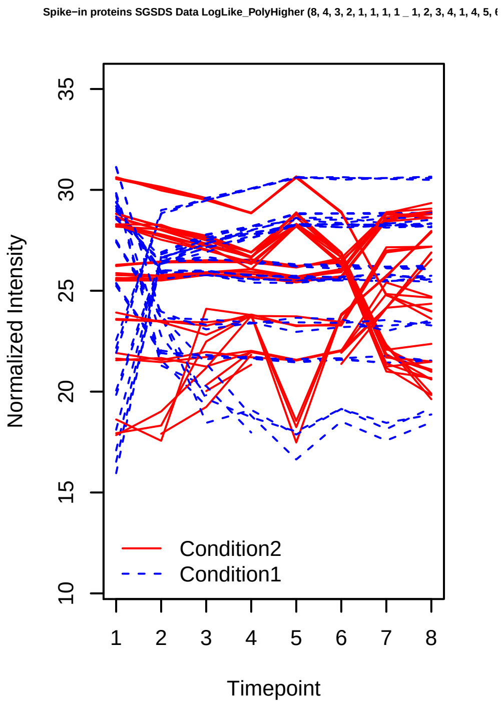
Spike−in proteins SGSDS Data LogLike_PolyHigher (8, 4, 3, 2, 1, 1, 1, 1 _ 1, 2, 3, 4, 1, 4, 5,
Normalized Intensity
Timepoint
Condition2
Condition1
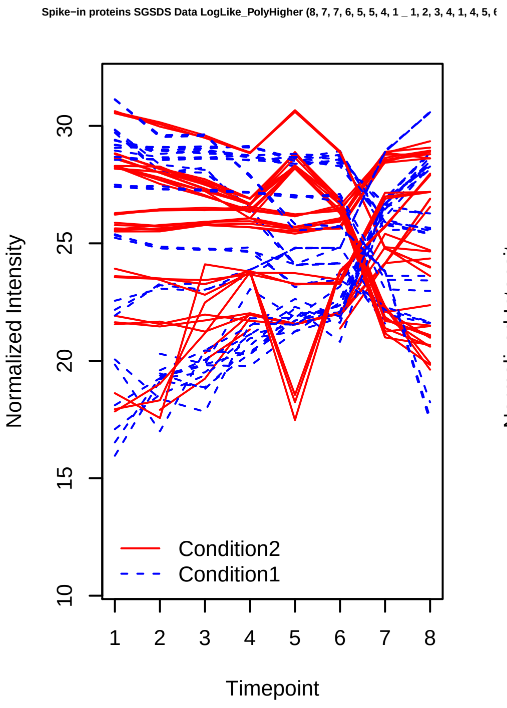
Spike−in proteins SGSDS Data LogLike_PolyHigher (8, 7, 7, 6, 5, 5, 4, 1 _ 1, 2, 3, 4, 1, 4, 5,
Normalized Intensity
Timepoint
Condition2
Condition1
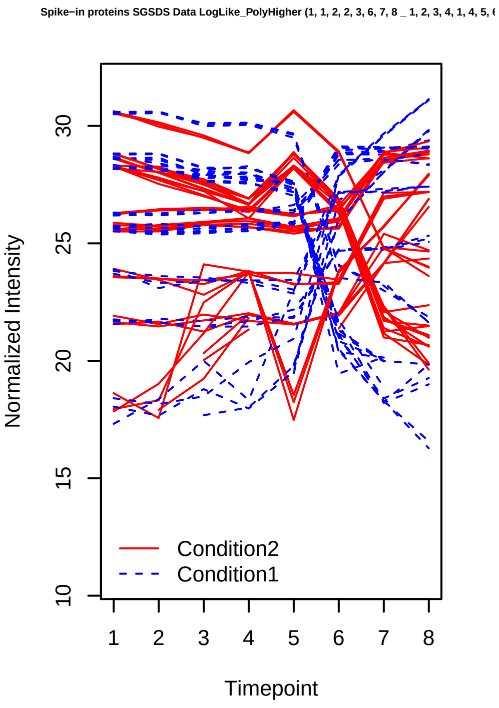
Spike−in proteins SGSDS Data LogLike_PolyHigher (1, 1, 2, 2, 3, 6, 7, 8 _ 1, 2, 3, 4, 1, 4, 5,
Normalized Intensity
Timepoint
Condition2
Condition1

Spike−in proteins SGSDS Data LogLike_PolyHigher (1, 5, 6, 7, 8, 8, 8, 8 _ 4, 5, 3, 2, 1, 6, 7,

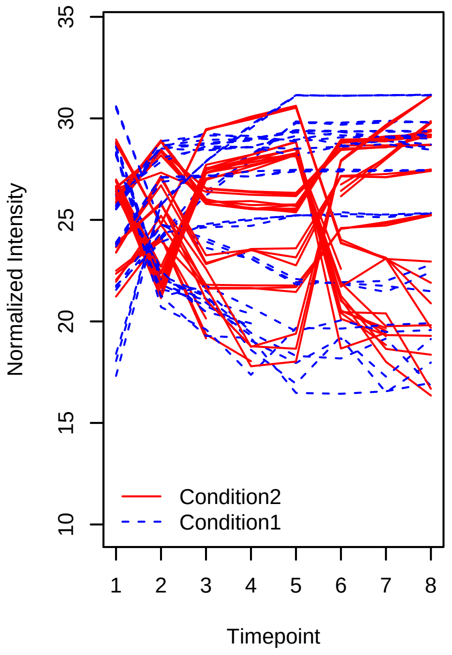


Spike−in proteins SGSDS Data LogLike_PolyHigher (8, 4, 3, 2, 1, 1, 1, 1 _ 4, 5, 3, 2, 1, 6, 7,

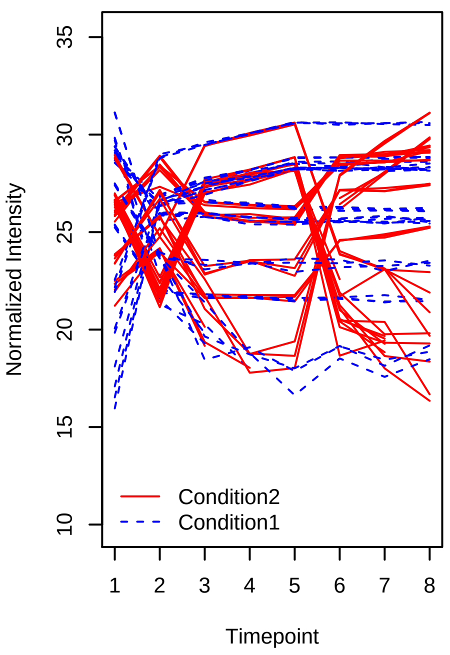


Spike−in proteins SGSDS Data LogLike_PolyHigher (8, 7, 7, 6, 5, 5, 4, 1 _ 4, 5, 3, 2, 1, 6, 7,

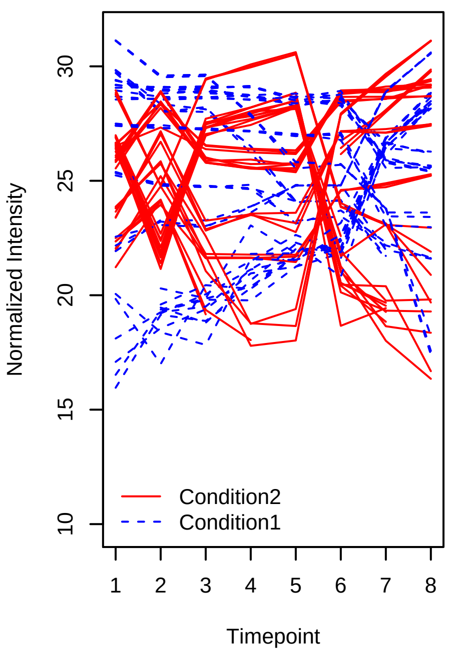


Spike−in proteins SGSDS Data LogLike_PolyHigher (1, 1, 2, 2, 3, 6, 7, 8 _ 4, 5, 3, 2, 1, 6, 7,

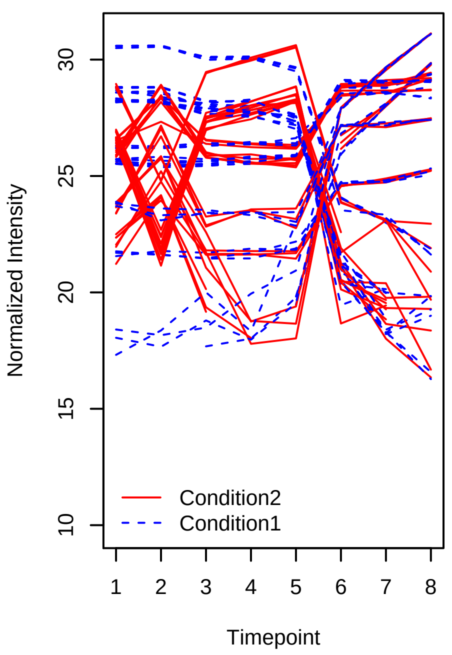


Spike−in proteins SGSDS Data LogLike_PolyHigher (1, 5, 6, 7, 8, 8, 8, 8 _ 1, 2, 4, 1, 6, 7, 3,

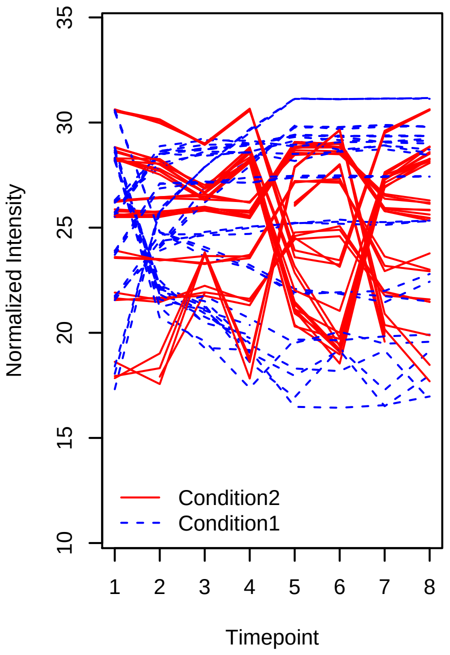


Spike−in proteins SGSDS Data LogLike_PolyHigher (8, 4, 3, 2, 1, 1, 1, 1 _ 1, 2, 4, 1, 6, 7, 3,

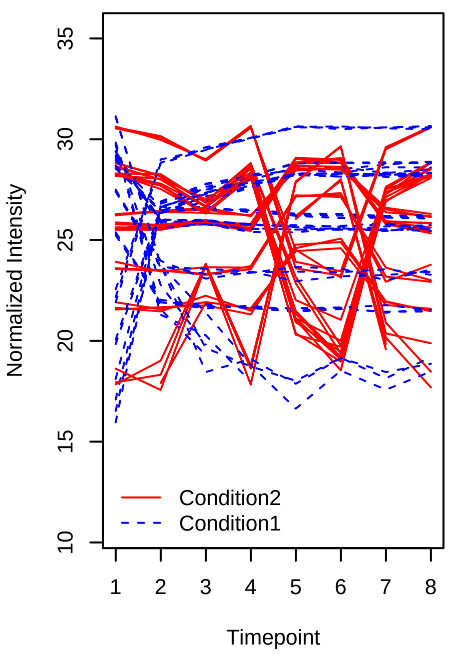

Spike−in proteins SGSDS Data LogLike_PolyHigher (8, 7, 7, 6, 5, 5, 4, 1 _ 1, 2, 4, 1, 6, 7, 3, 1

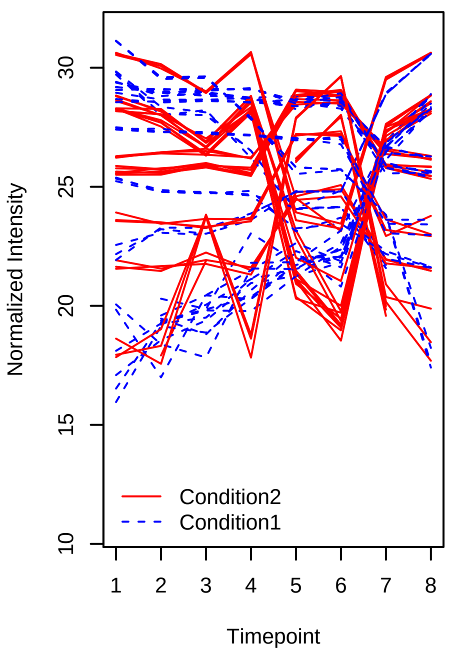


Spike−in proteins SGSDS Data LogLike_PolyHigher (1, 1, 2, 2, 3, 6, 7, 8 _ 1, 2, 4, 1, 6, 7, 3, 1

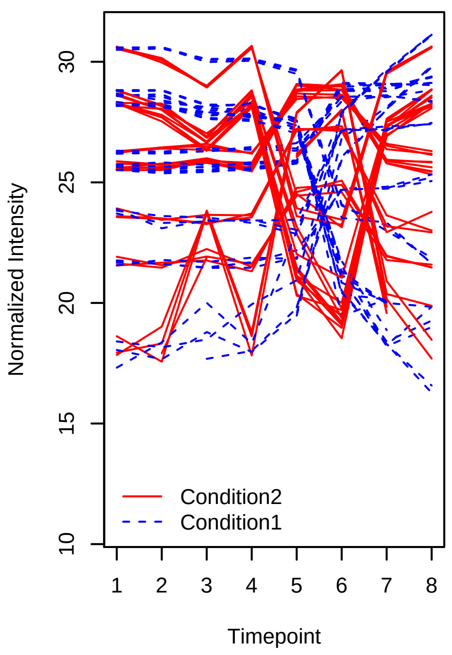


Spike−in proteins SGSDS Data LogLike_PolyHigher (1, 5, 6, 7, 8, 8, 8, 8 _ 8, 7, 6, 5, 1, 5, 3, 2

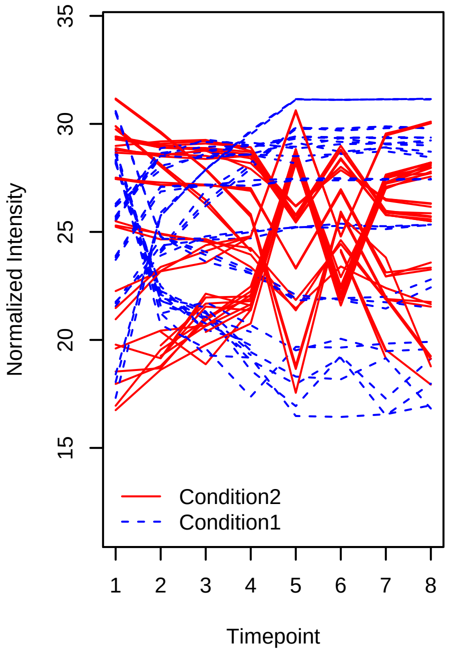


Spike−in proteins SGSDS Data LogLike_PolyHigher (8, 4, 3, 2, 1, 1, 1, 1 _ 8, 7, 6, 5, 1, 5, 3, 2

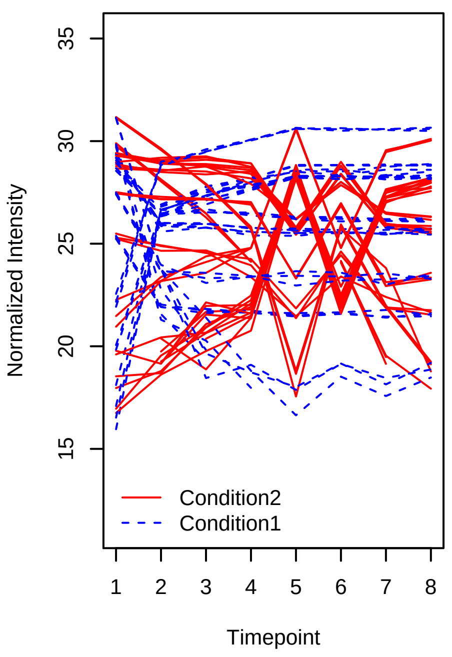


Spike−in proteins SGSDS Data LogLike_PolyHigher (8, 7, 7, 6, 5, 5, 4, 1 _ 8, 7, 6, 5, 1, 5, 3, 2

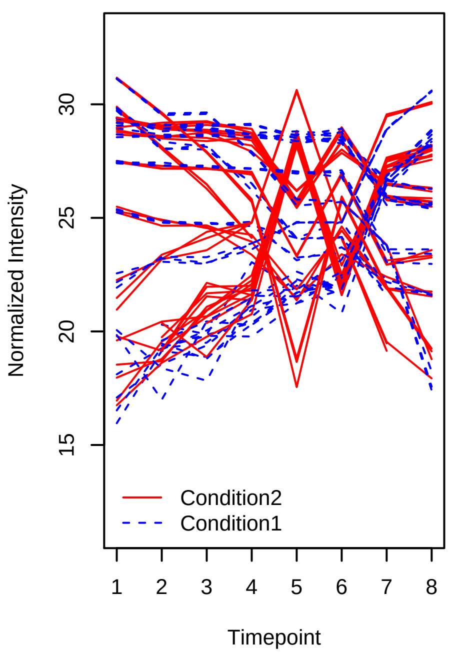


Spike−in proteins SGSDS Data LogLike_PolyHigher (1, 1, 2, 2, 3, 6, 7, 8 _ 8, 7, 6, 5, 1, 5, 3, 2

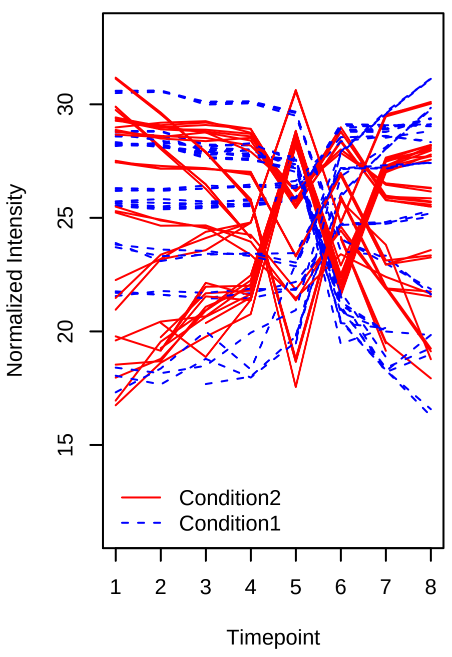

Spike−in proteins SGSDS Data Poly2_Poly2 (1, 5, 6, 7, 8, 7, 5, 1 _ 8, 4, 3, 2, 1, 2, 3, 8)

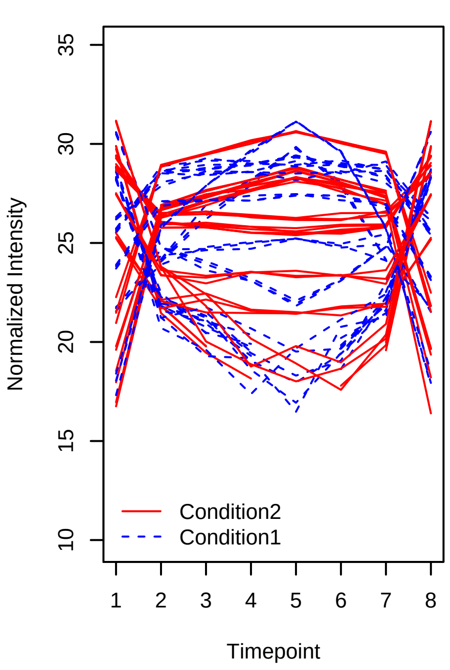


Spike−in proteins SGSDS Data Poly2_Poly2 (8, 7, 6, 5, 5, 6, 7, 8 _ 8, 4, 3, 2, 1, 2, 3, 8)

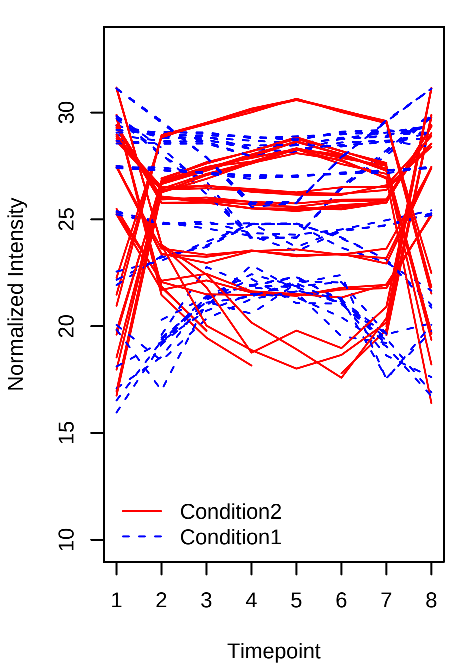


Spike−in proteins SGSDS Data Poly2_Poly2 (1, 2, 3, 4, 4, 3, 2, 1 _ 8, 4, 3, 2, 1, 2, 3, 8)

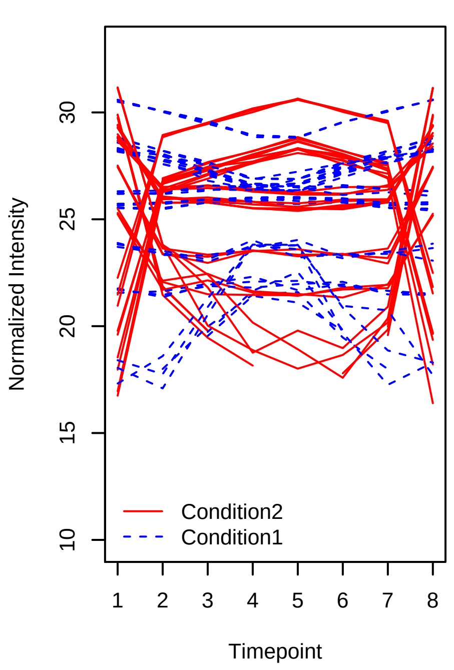


Spike−in proteins SGSDS Data Poly2_Poly2 (8, 5, 3, 3, 3, 3, 6, 8 _ 8, 4, 3, 2, 1, 2, 3, 8)

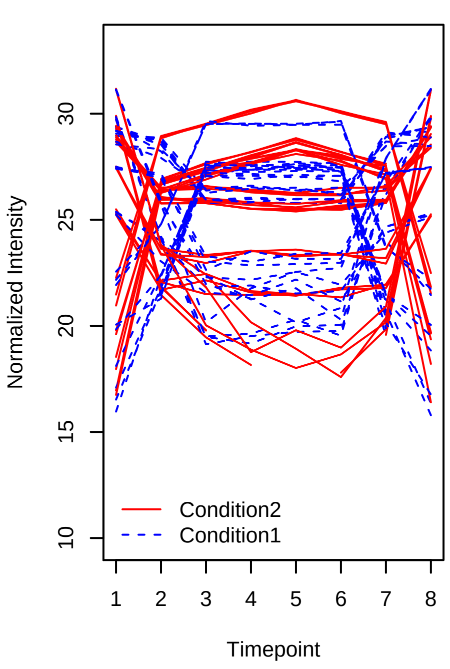


Spike−in proteins SGSDS Data Poly2_Poly2 (8, 7, 6, 5, 5, 6, 7, 8 _ 1, 5, 6, 7, 8, 7, 5, 1)

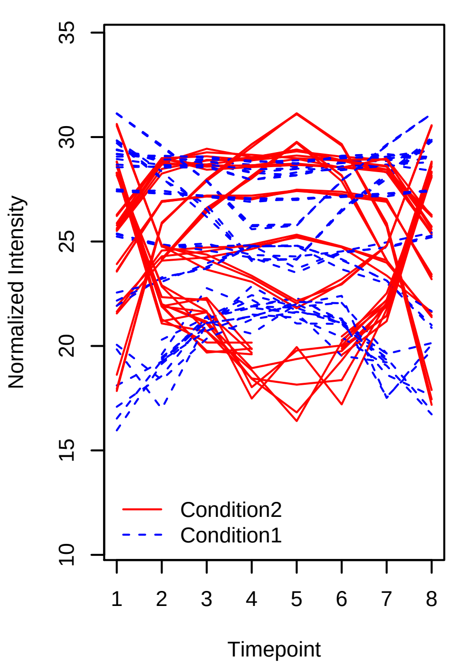


Spike−in proteins SGSDS Data Poly2_Poly2 (1, 2, 3, 4, 4, 3, 2, 1 _ 1, 5, 6, 7, 8, 7, 5, 1)

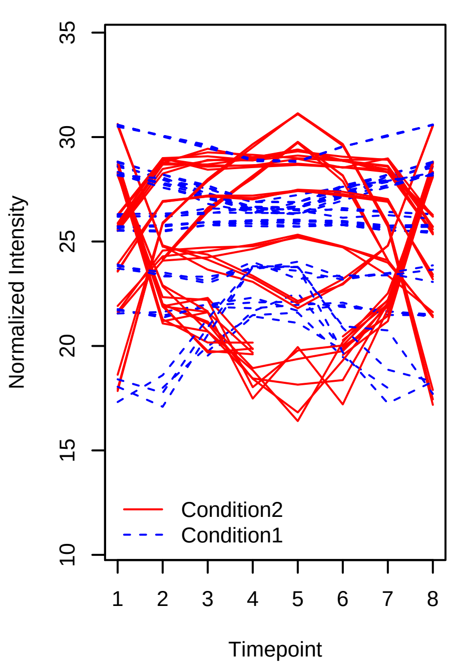

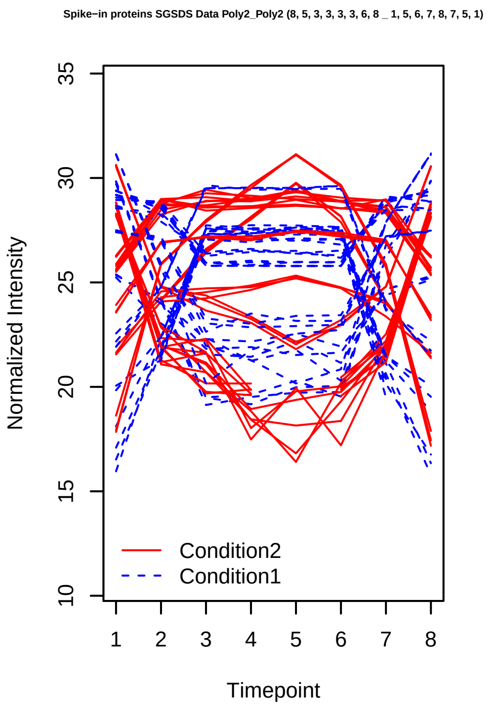
Spike−in proteins SGSDS Data Poly2_Poly2 (8, 5, 3, 3, 3, 3, 6, 8 _ 1, 5, 6, 7, 8, 7, 5, 1)
Normalized Intensity
10 15 20 25 30 35
1 2 3 4 5 6 7 8
Timepoint
Condition2
Condition1

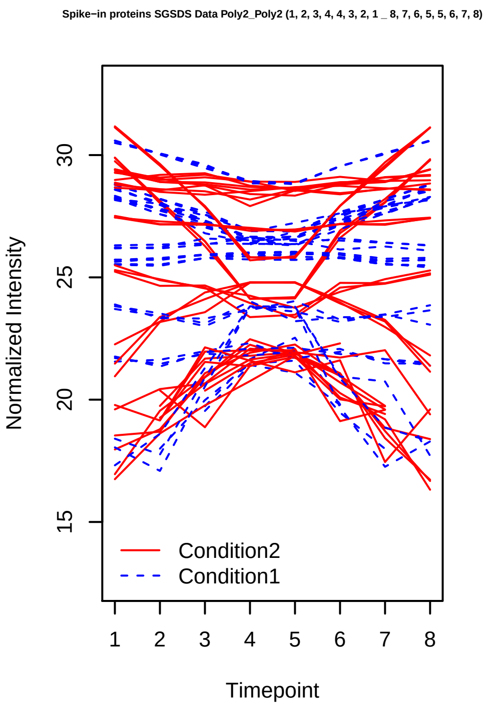
Spike−in proteins SGSDS Data Poly2_Poly2 (1, 2, 3, 4, 4, 3, 2, 1 _ 8, 7, 6, 5, 5, 6, 7, 8)
Normalized Intensity
15 20 25 30
1 2 3 4 5 6 7 8
Timepoint
Condition2
Condition1

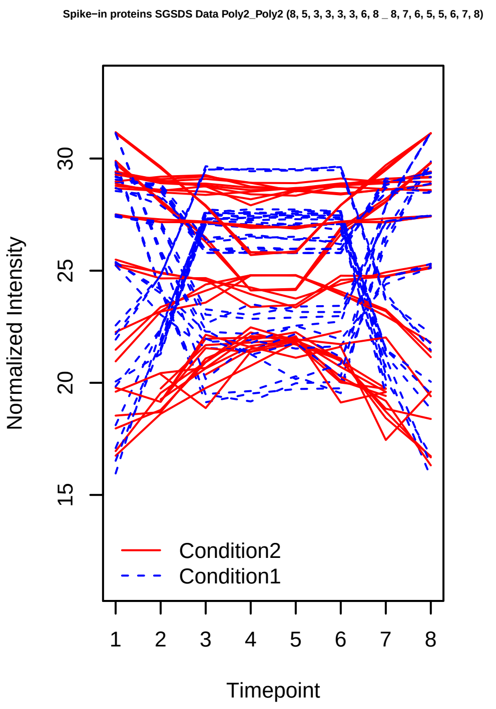
Spike−in proteins SGSDS Data Poly2_Poly2 (8, 5, 3, 3, 3, 3, 6, 8 _ 8, 7, 6, 5, 5, 6, 7, 8)
Normalized Intensity
15 20 25 30
1 2 3 4 5 6 7 8
Timepoint
Condition2
Condition1

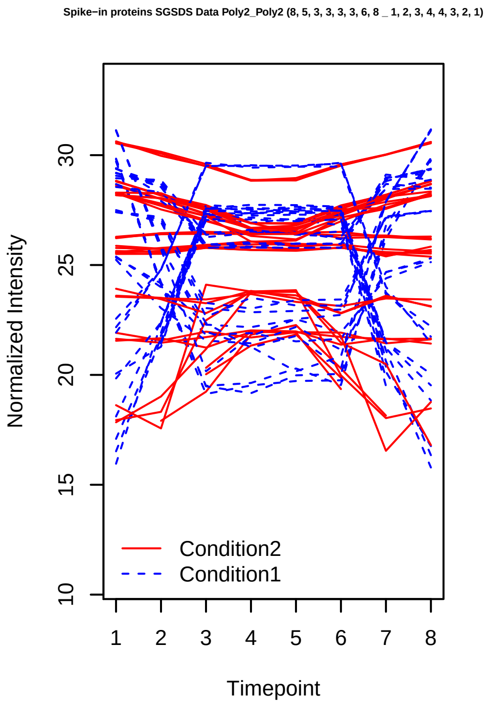
Spike−in proteins SGSDS Data Poly2_Poly2 (8, 5, 3, 3, 3, 3, 6, 8 _ 1, 2, 3, 4, 4, 3, 2, 1)
Normalized Intensity
10 15 20 25 30
1 2 3 4 5 6 7 8
Timepoint
Condition2
Condition1

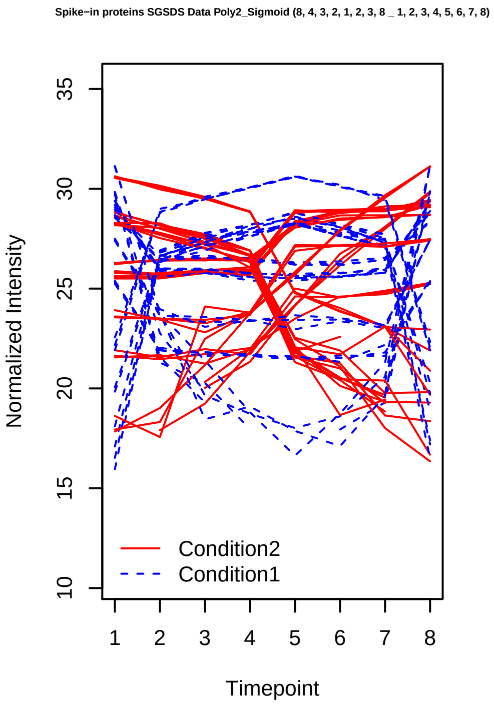
Spike−in proteins SGSDS Data Poly2_Sigmoid (8, 4, 3, 2, 1, 2, 3, 8 _ 1, 2, 3, 4, 5, 6, 7, 8)
Normalized Intensity
10 15 20 25 30 35
1 2 3 4 5 6 7 8
Timepoint
Condition2
Condition1

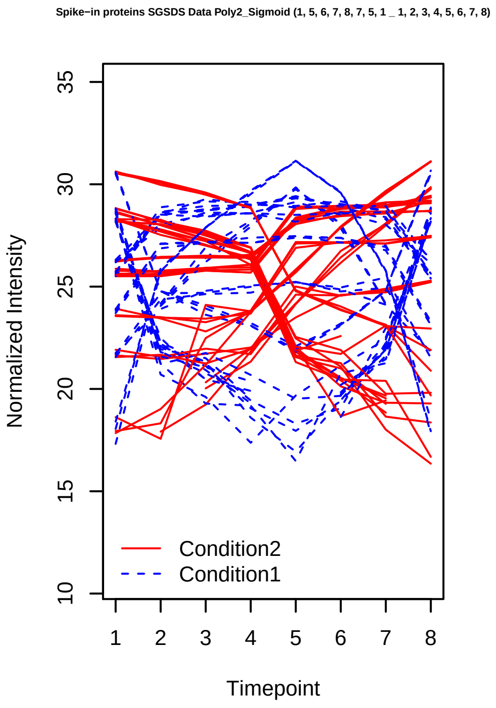
Spike−in proteins SGSDS Data Poly2_Sigmoid (1, 5, 6, 7, 8, 7, 5, 1 _ 1, 2, 3, 4, 5, 6, 7, 8)
Normalized Intensity
10 15 20 25 30 35
1 2 3 4 5 6 7 8
Timepoint
Condition2
Condition1

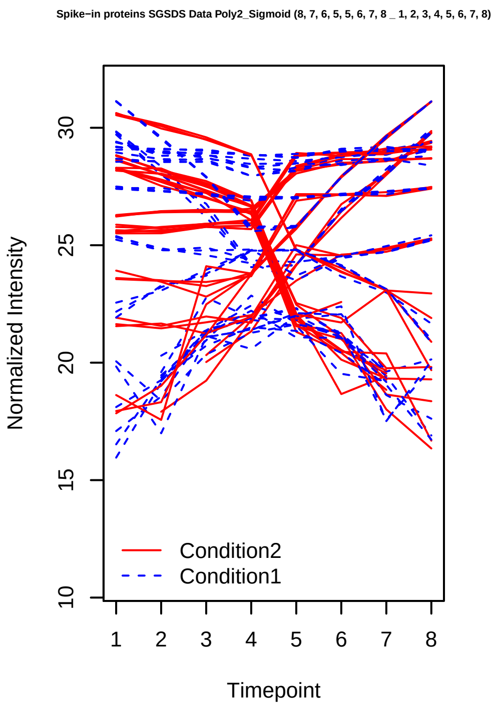
Spike−in proteins SGSDS Data Poly2_Sigmoid (8, 7, 6, 5, 5, 6, 7, 8 _ 1, 2, 3, 4, 5, 6, 7, 8)
Normalized Intensity
10
15
20
25
30
1 2 3 4 5 6 7 8
Timepoint
Condition2
Condition1

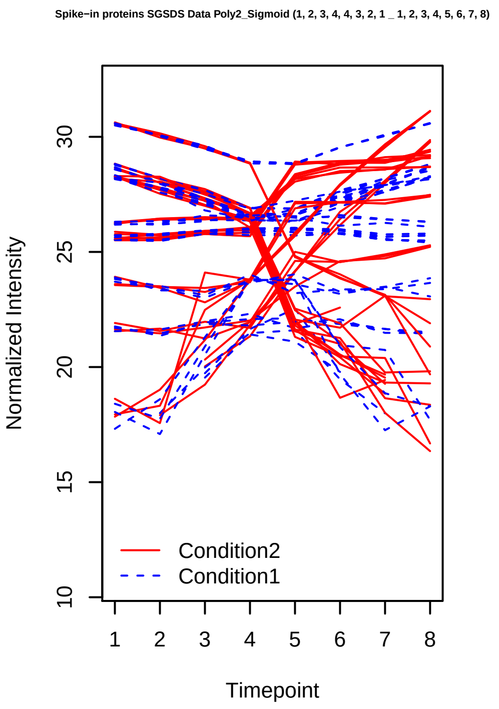
Spike−in proteins SGSDS Data Poly2_Sigmoid (1, 2, 3, 4, 4, 3, 2, 1 _ 1, 2, 3, 4, 5, 6, 7, 8)
Normalized Intensity
10
15
20
25
30
1 2 3 4 5 6 7 8
Timepoint
Condition2
Condition1

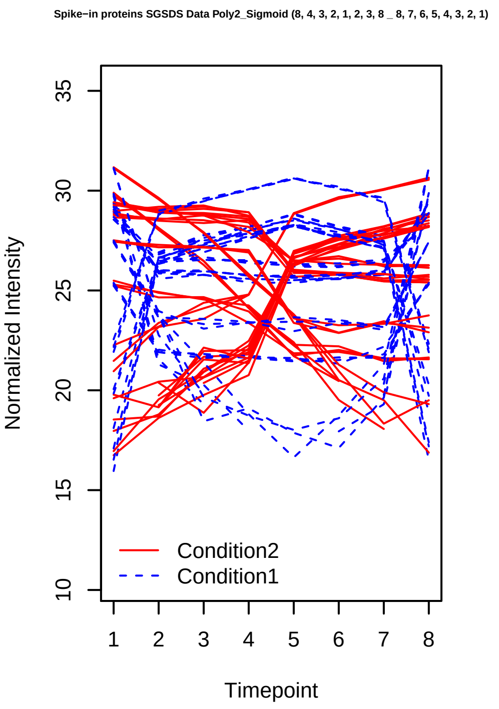
Spike−in proteins SGSDS Data Poly2_Sigmoid (8, 4, 3, 2, 1, 2, 3, 8 _ 8, 7, 6, 5, 4, 3, 2, 1)
Normalized Intensity
10
15
20
25
30
35
1 2 3 4 5 6 7 8
Timepoint
Condition2
Condition1

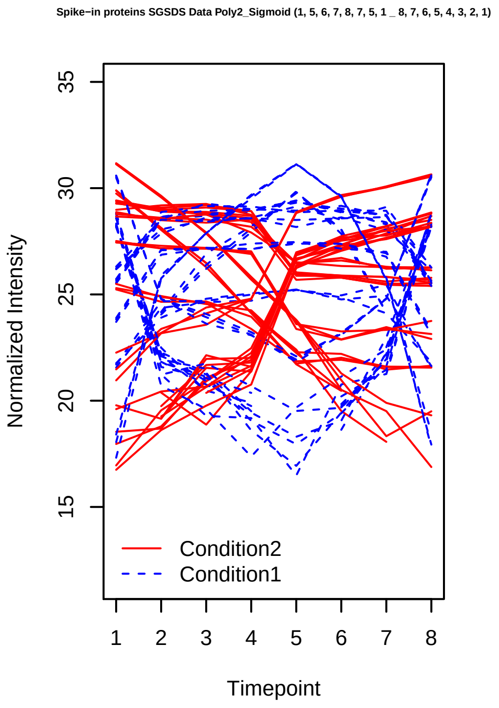
Spike−in proteins SGSDS Data Poly2_Sigmoid (1, 5, 6, 7, 8, 7, 5, 1 _ 8, 7, 6, 5, 4, 3, 2, 1)
Normalized Intensity
15
20
25
30
35
1 2 3 4 5 6 7 8
Timepoint
Condition2
Condition1

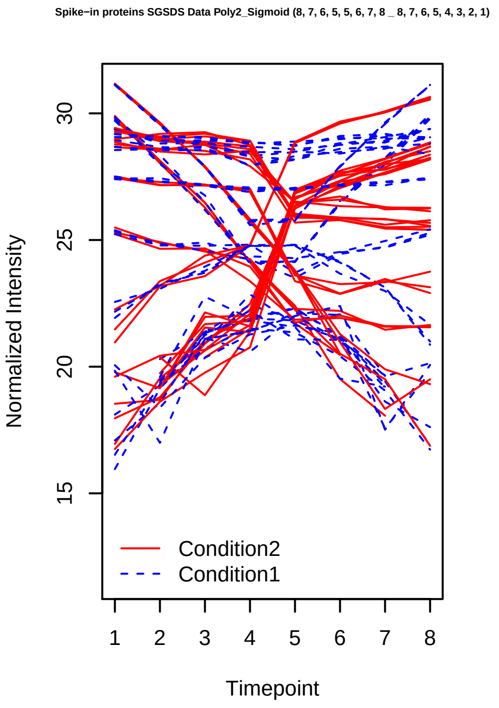
Spike−in proteins SGSDS Data Poly2_Sigmoid (8, 7, 6, 5, 5, 6, 7, 8 _ 8, 7, 6, 5, 4, 3, 2, 1)
Normalized Intensity
15
20
25
30
1 2 3 4 5 6 7 8
Timepoint
Condition2
Condition1

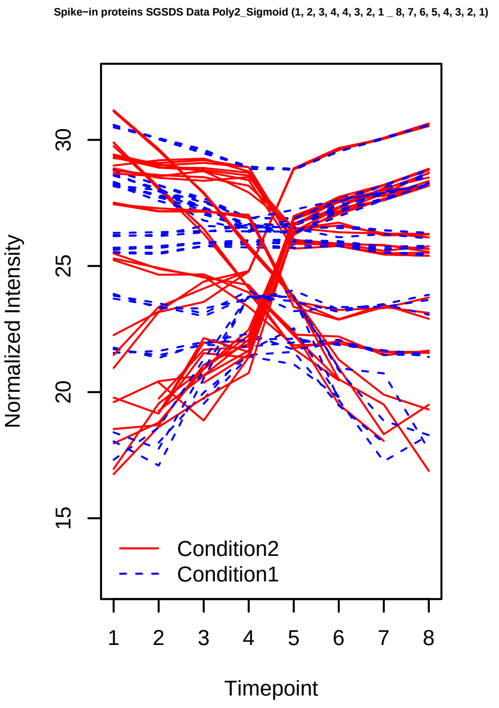
Spike−in proteins SGSDS Data Poly2_Sigmoid (1, 2, 3, 4, 4, 3, 2, 1 _ 8, 7, 6, 5, 4, 3, 2, 1)
Normalized Intensity
15
20
25
30
1 2 3 4 5 6 7 8
Timepoint
Condition2
Condition1

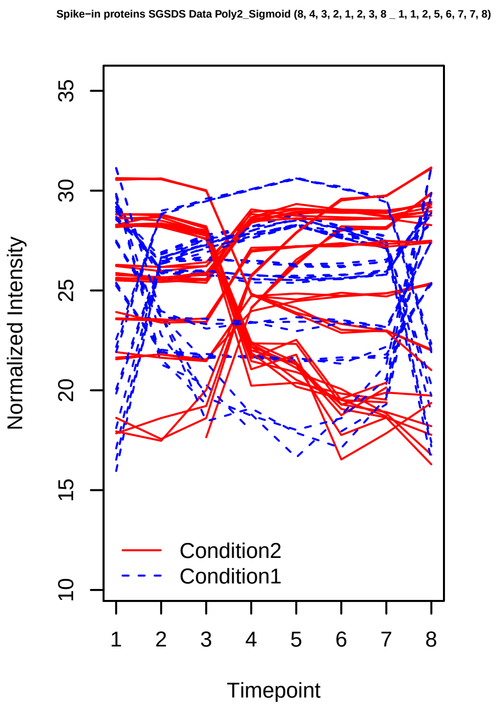
Spike−in proteins SGSDS Data Poly2_Sigmoid (8, 4, 3, 2, 1, 2, 3, 8 _ 1, 1, 2, 5, 6, 7, 7, 8)
Normalized Intensity
Timepoint
Condition2
Condition1

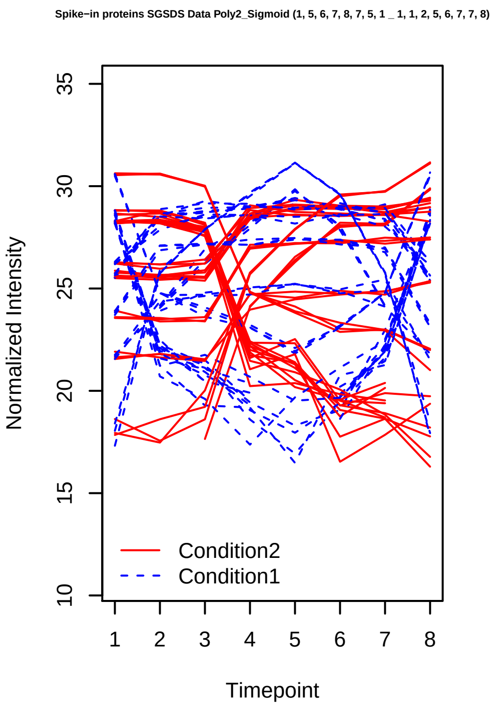
Spike−in proteins SGSDS Data Poly2_Sigmoid (1, 5, 6, 7, 8, 7, 5, 1 _ 1, 1, 2, 5, 6, 7, 7, 8)
Normalized Intensity
Timepoint
Condition2
Condition1

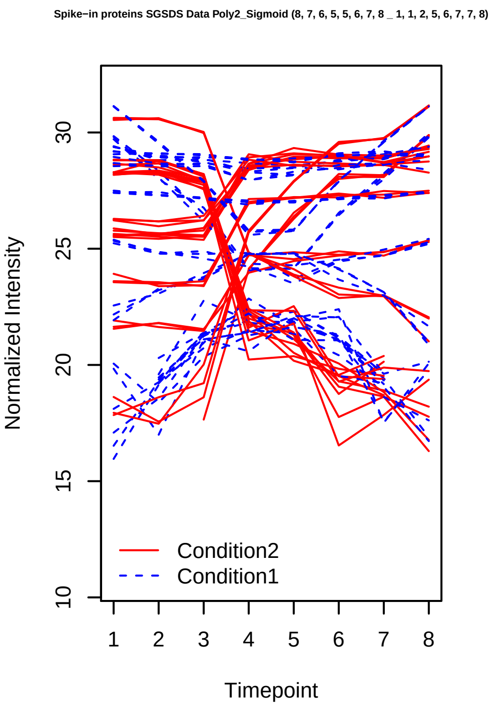
Spike−in proteins SGSDS Data Poly2_Sigmoid (8, 7, 6, 5, 5, 6, 7, 8 _ 1, 1, 2, 5, 6, 7, 7, 8)
Normalized Intensity
Timepoint
Condition2
Condition1

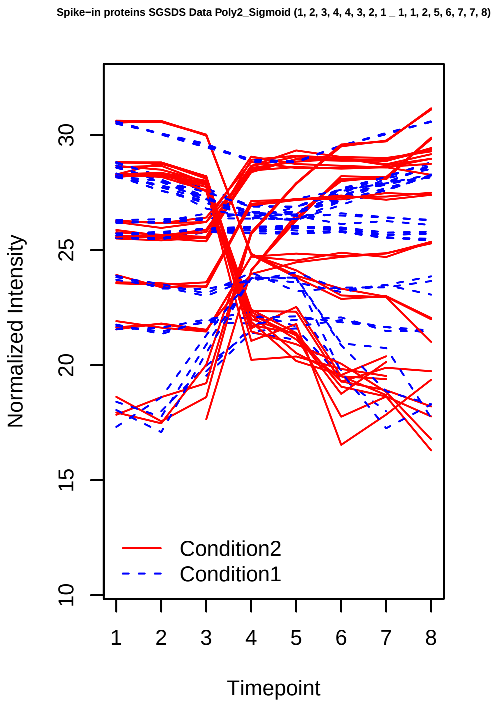
Spike−in proteins SGSDS Data Poly2_Sigmoid (1, 2, 3, 4, 4, 3, 2, 1 _ 1, 1, 2, 5, 6, 7, 7, 8)
Normalized Intensity
Timepoint
Condition2
Condition1

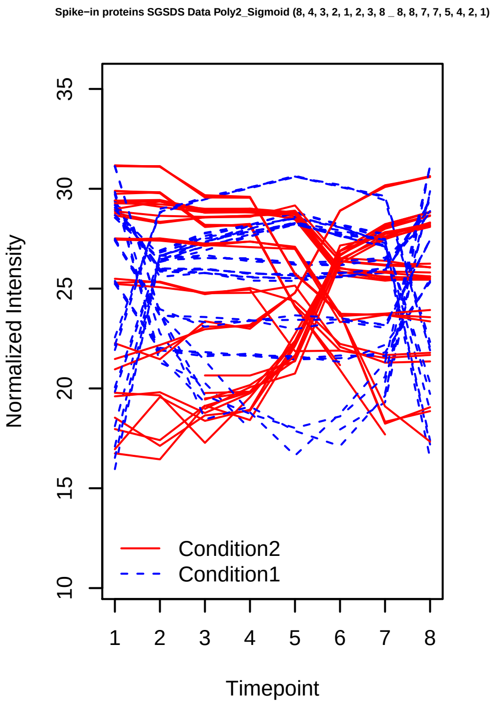
Spike−in proteins SGSDS Data Poly2_Sigmoid (8, 4, 3, 2, 1, 2, 3, 8 _ 8, 8, 7, 7, 5, 4, 2, 1)
Normalized Intensity
Timepoint
Condition2
Condition1

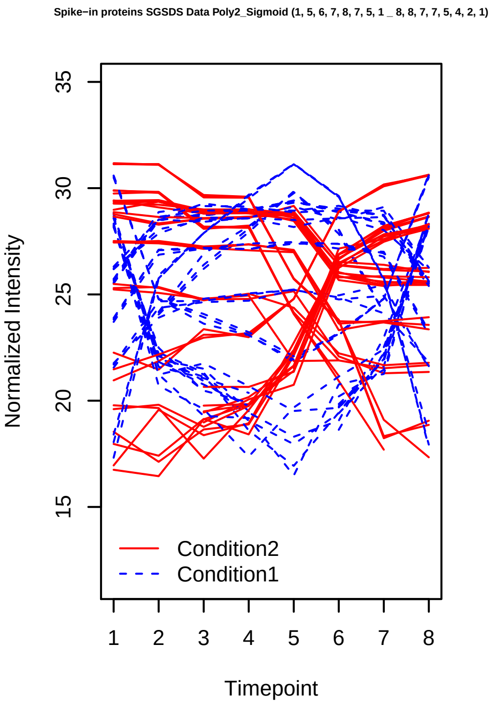
Spike−in proteins SGSDS Data Poly2_Sigmoid (1, 5, 6, 7, 8, 7, 5, 1 _ 8, 8, 7, 7, 5, 4, 2, 1)
Normalized Intensity
Timepoint
Condition2
Condition1

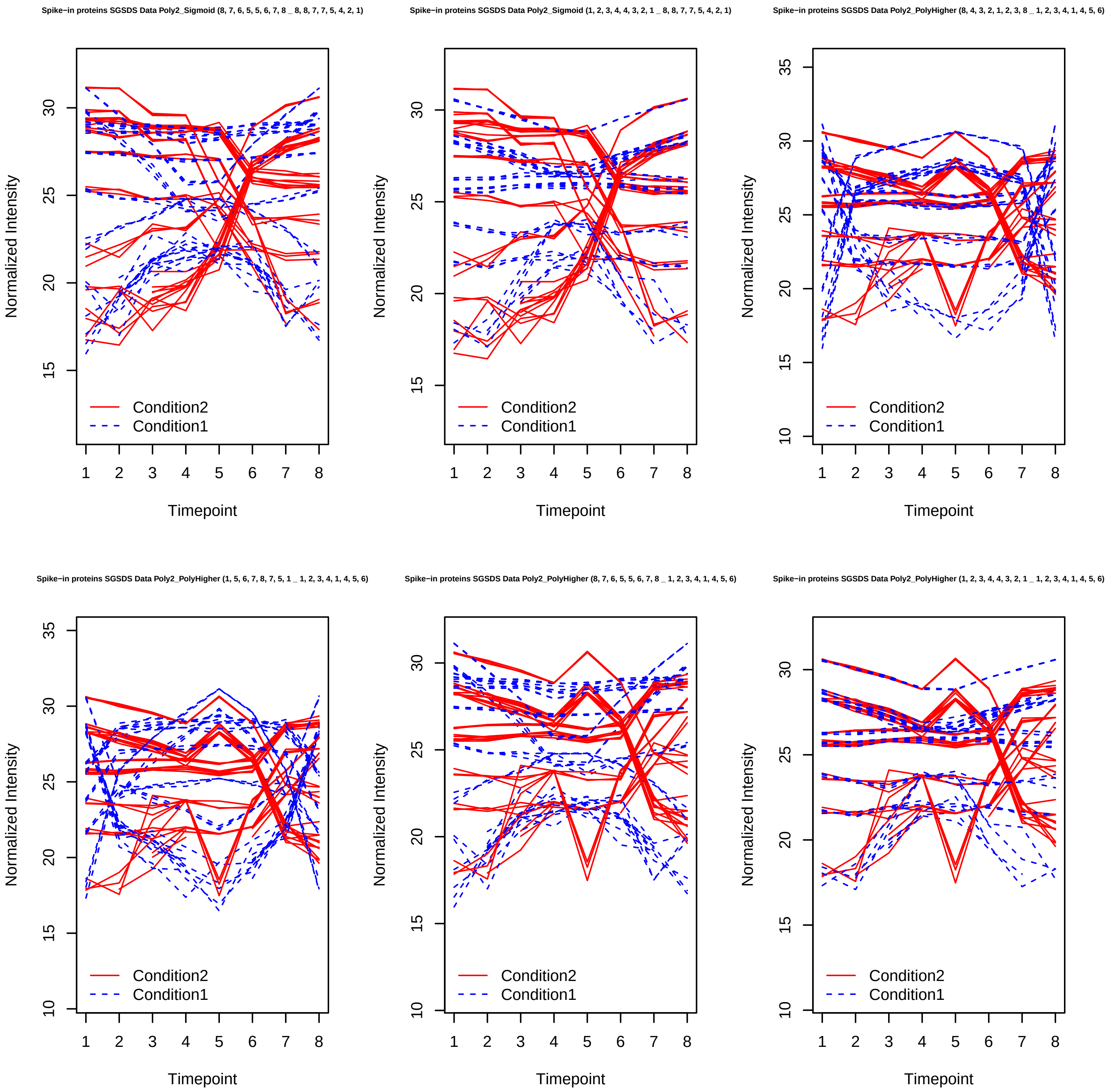

Spike−in proteins SGSDS Data Poly2_Sigmoid (8, 7, 6, 5, 5, 6, 7, 8 _ 8, 8, 7, 7, 5, 4, 2, 1)
Spike−in proteins SGSDS Data Poly2_Sigmoid (1, 2, 3, 4, 4, 3, 2, 1 _ 8, 8, 7, 7, 5, 4, 2, 1)
Spike−in proteins SGSDS Data Poly2_PolyHigher (8, 4, 3, 2, 1, 2, 3, 8 _ 1, 2, 3, 4, 1, 4, 5, 6)
Spike−in proteins SGSDS Data Poly2_PolyHigher (1, 5, 6, 7, 8, 7, 5, 1 _ 1, 2, 3, 4, 1, 4, 5, 6)
Spike−in proteins SGSDS Data Poly2_PolyHigher (8, 7, 6, 5, 5, 6, 7, 8 _ 1, 2, 3, 4, 1, 4, 5, 6)
Spike−in proteins SGSDS Data Poly2_PolyHigher (1, 2, 3, 4, 4, 3, 2, 1 _ 1, 2, 3, 4, 1, 4, 5, 6)
Normalized Intensity
Timepoint
Condition2
Condition1

Spike−in proteins SGSDS Data Poly2_PolyHigher (8, 4, 3, 2, 1, 2, 3, 8 _ 4, 5, 3, 2, 1, 6, 7, 8)

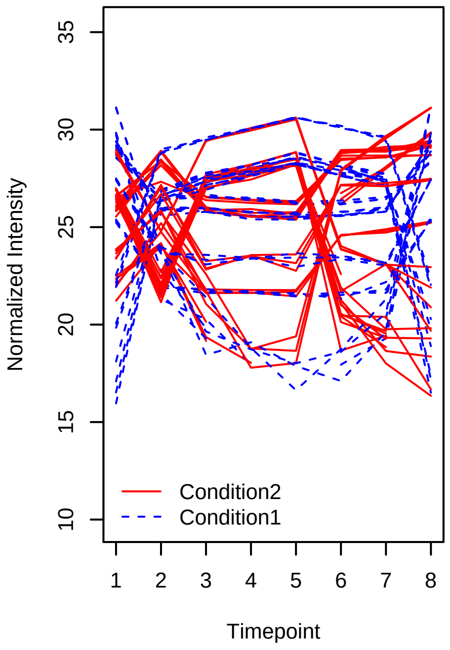


Spike−in proteins SGSDS Data Poly2_PolyHigher (1, 5, 6, 7, 8, 7, 5, 1 _ 4, 5, 3, 2, 1, 6, 7, 8)

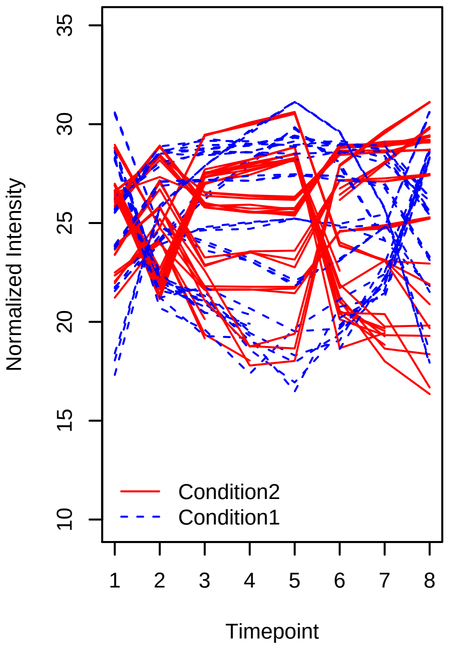


Spike−in proteins SGSDS Data Poly2_PolyHigher (8, 7, 6, 5, 5, 6, 7, 8 _ 4, 5, 3, 2, 1, 6, 7, 8)

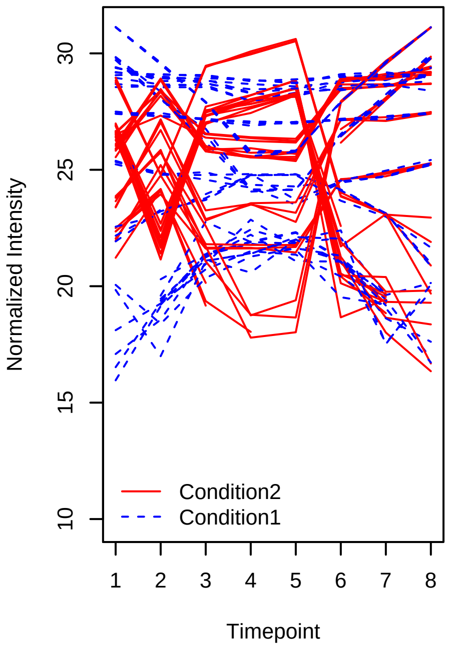


Spike−in proteins SGSDS Data Poly2_PolyHigher (1, 2, 3, 4, 4, 3, 2, 1 _ 4, 5, 3, 2, 1, 6, 7, 8)

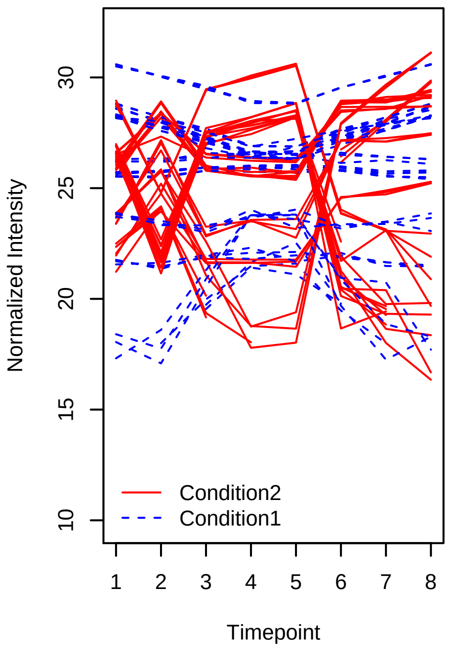


Spike−in proteins SGSDS Data Poly2_PolyHigher (8, 4, 3, 2, 1, 2, 3, 8 _ 1, 2, 4, 1, 6, 7, 3, 1)

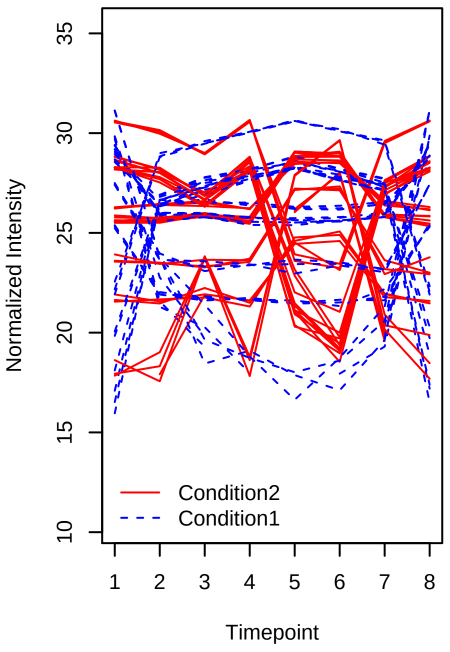


Spike−in proteins SGSDS Data Poly2_PolyHigher (1, 5, 6, 7, 8, 7, 5, 1 _ 1, 2, 4, 1, 6, 7, 3, 1)

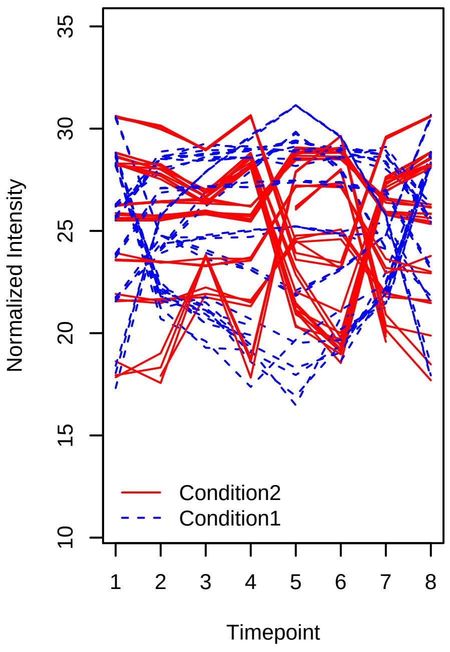

Spike−in proteins SGSDS Data Poly2_PolyHigher (8, 7, 6, 5, 5, 6, 7, 8 _ 1, 2, 4, 1, 6, 7, 3, 1)

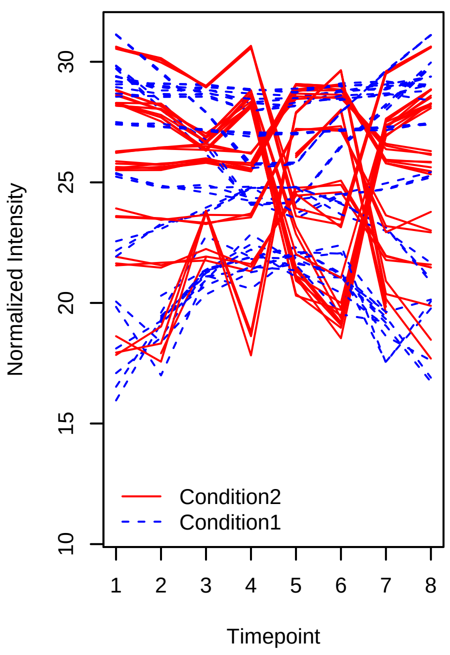


Spike−in proteins SGSDS Data Poly2_PolyHigher (1, 2, 3, 4, 4, 3, 2, 1 _ 1, 2, 4, 1, 6, 7, 3, 1)

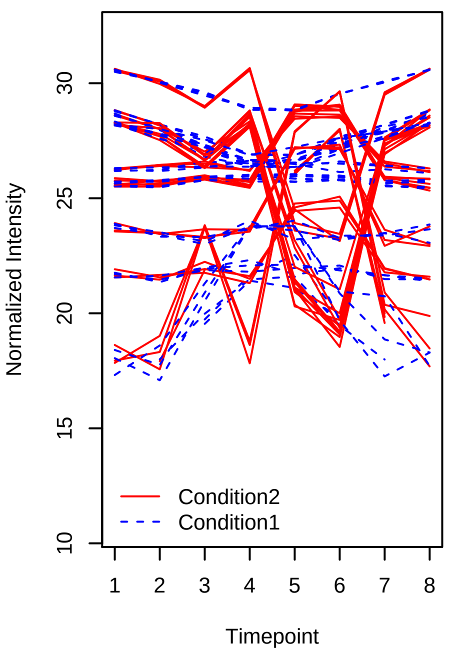


Spike−in proteins SGSDS Data Poly2_PolyHigher (8, 4, 3, 2, 1, 2, 3, 8 _ 8, 7, 6, 5, 1, 5, 3, 2)

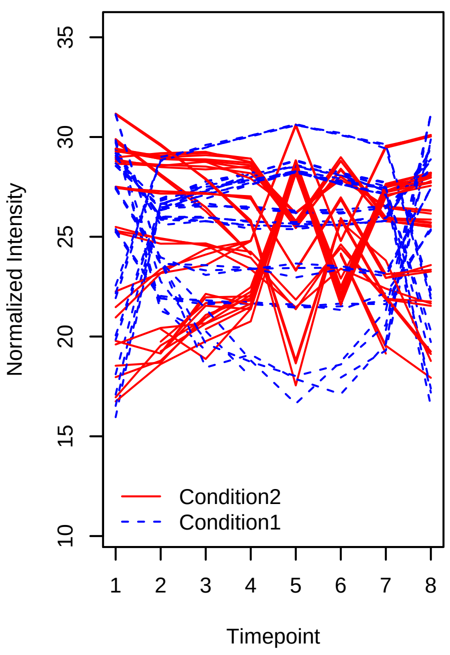


Spike−in proteins SGSDS Data Poly2_PolyHigher (1, 5, 6, 7, 8, 7, 5, 1 _ 8, 7, 6, 5, 1, 5, 3, 2)

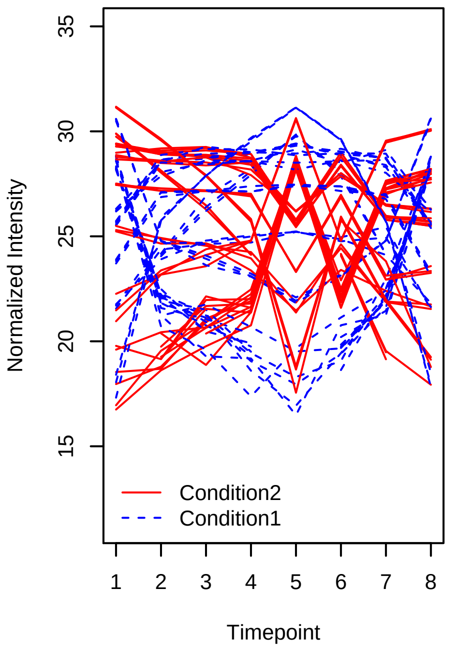


Spike−in proteins SGSDS Data Poly2_PolyHigher (8, 7, 6, 5, 5, 6, 7, 8 _ 8, 7, 6, 5, 1, 5, 3, 2)

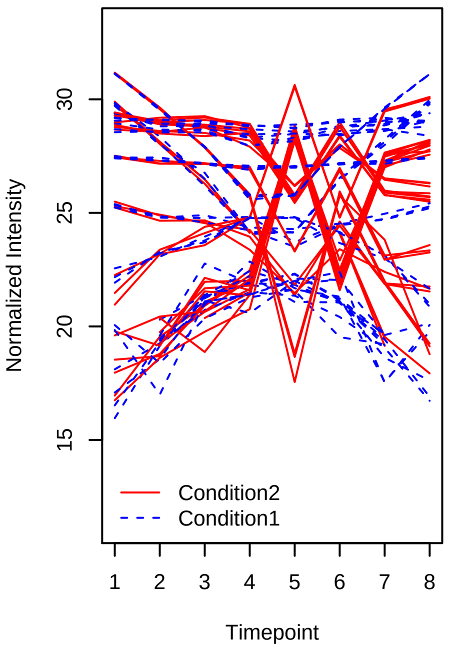


Spike−in proteins SGSDS Data Poly2_PolyHigher (1, 2, 3, 4, 4, 3, 2, 1 _ 8, 7, 6, 5, 1, 5, 3, 2)

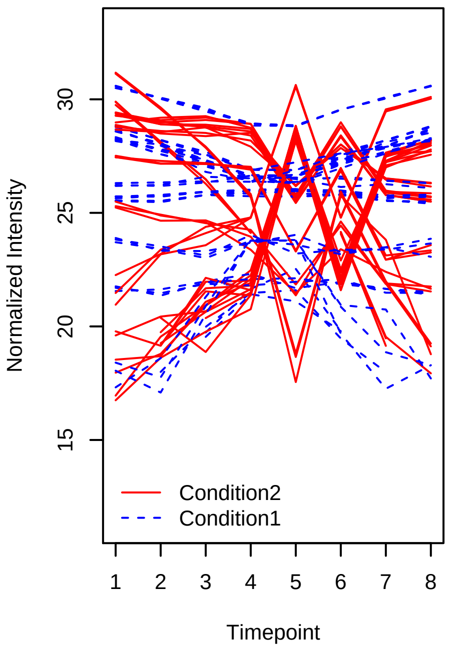

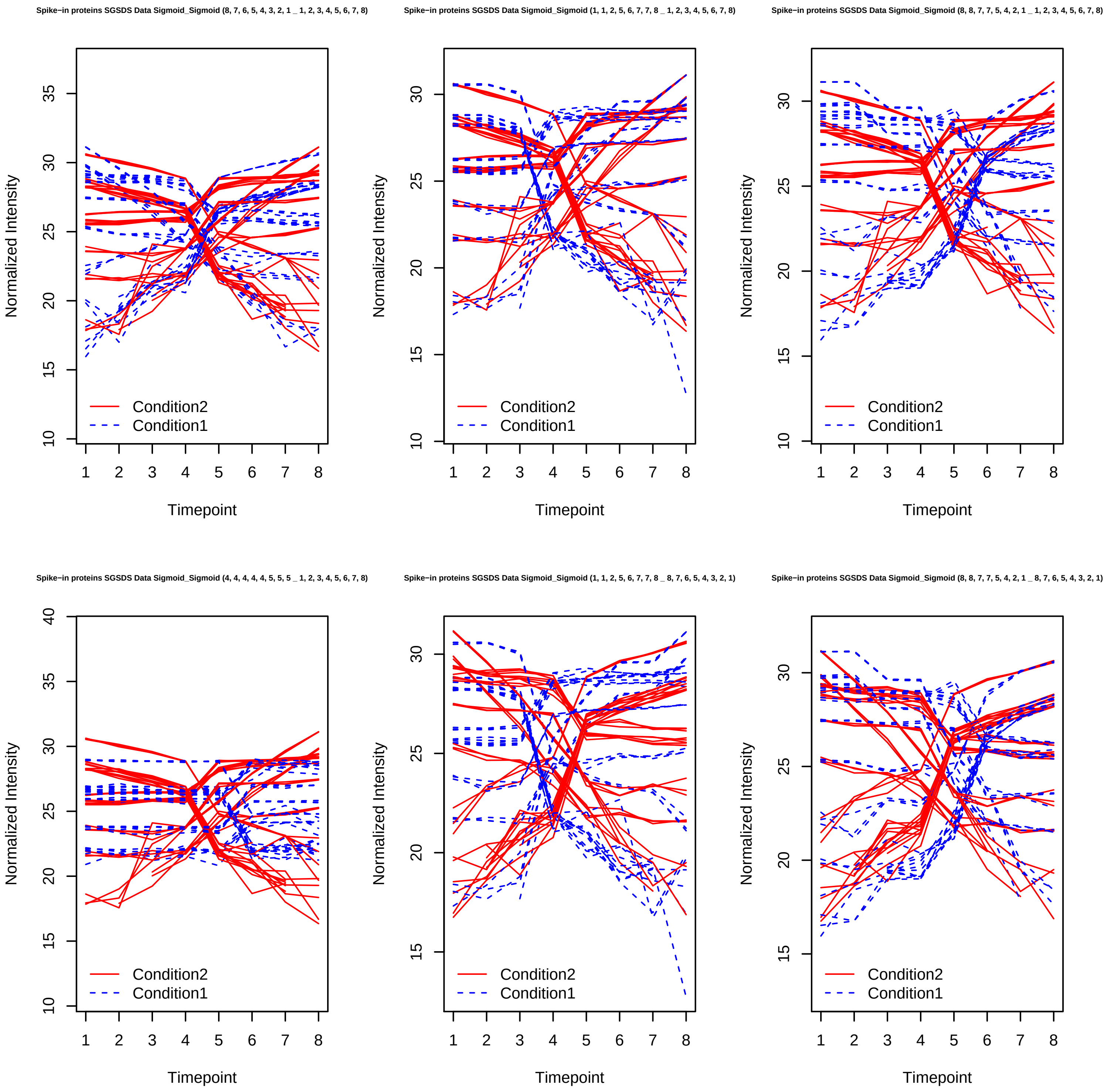
Spike−in proteins SGSDS Data Sigmoid_Sigmoid (8, 7, 6, 5, 4, 3, 2, 1 _ 1, 2, 3, 4, 5, 6, 7, 8)
Spike−in proteins SGSDS Data Sigmoid_Sigmoid (1, 1, 2, 5, 6, 7, 7, 8 _ 1, 2, 3, 4, 5, 6, 7, 8)
Spike−in proteins SGSDS Data Sigmoid_Sigmoid (8, 8, 7, 7, 5, 4, 2, 1 _ 1, 2, 3, 4, 5, 6, 7, 8)
Spike−in proteins SGSDS Data Sigmoid_Sigmoid (4, 4, 4, 4, 4, 5, 5, 5 _ 1, 2, 3, 4, 5, 6, 7, 8)
Spike−in proteins SGSDS Data Sigmoid_Sigmoid (1, 1, 2, 5, 6, 7, 7, 8 _ 8, 7, 6, 5, 4, 3, 2, 1)
Spike−in proteins SGSDS Data Sigmoid_Sigmoid (8, 8, 7, 7, 5, 4, 2, 1 _ 8, 7, 6, 5, 4, 3, 2, 1)
Normalized Intensity
Timepoint
Condition2
Condition1

Spike−in proteins SGSDS Data Sigmoid_Sigmoid (4, 4, 4, 4, 4, 5, 5, 5 _ 8, 7, 6, 5, 4, 3, 2, 1)

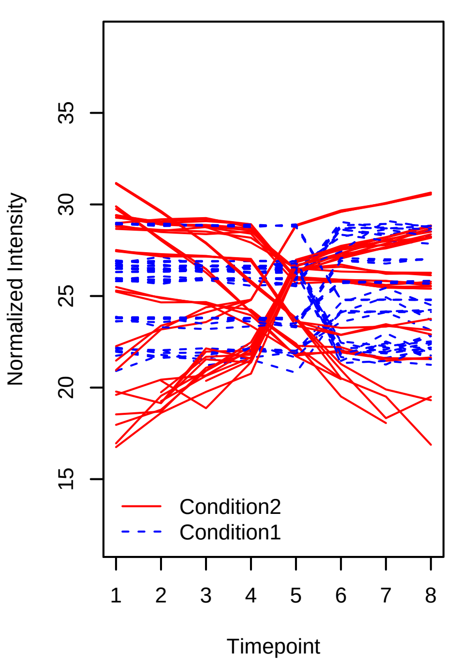


Spike−in proteins SGSDS Data Sigmoid_Sigmoid (8, 8, 7, 7, 5, 4, 2, 1 _ 1, 1, 2, 5, 6, 7, 7, 8)

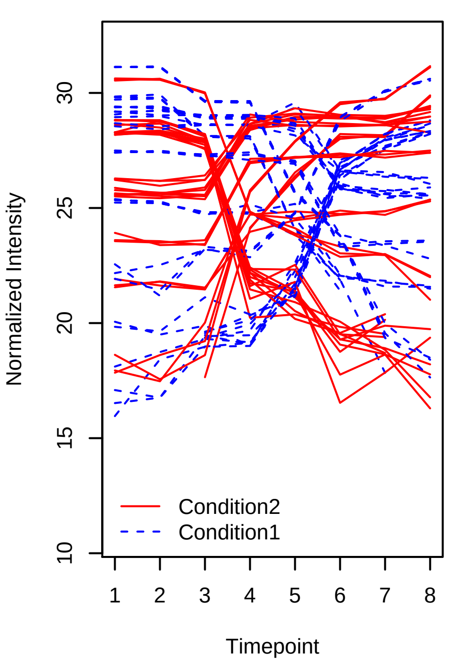


Spike−in proteins SGSDS Data Sigmoid_Sigmoid (4, 4, 4, 4, 4, 5, 5, 5 _ 1, 1, 2, 5, 6, 7, 7, 8)

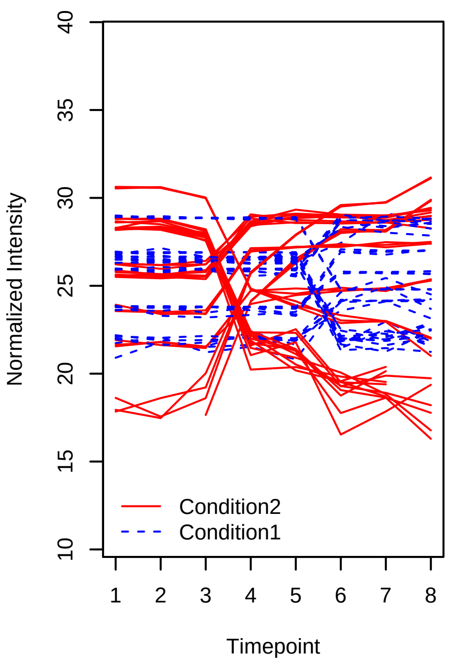


Spike−in proteins SGSDS Data Sigmoid_Sigmoid (4, 4, 4, 4, 4, 5, 5, 5 _ 8, 8, 7, 7, 5, 4, 2, 1)

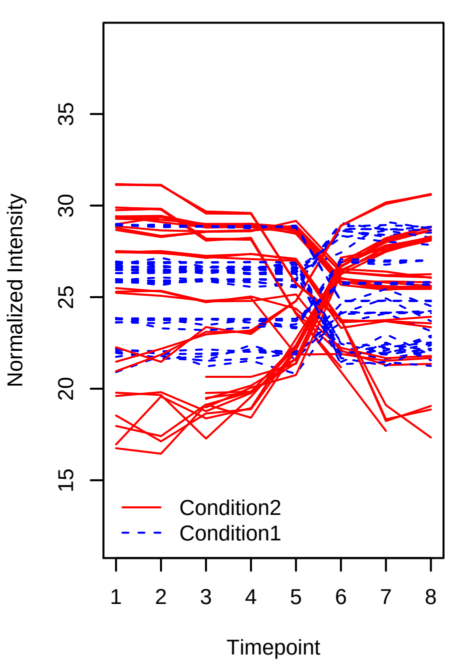


Spike−in proteins SGSDS Data Sigmoid_PolyHigher (1, 2, 3, 4, 5, 6, 7, 8 _ 1, 2, 3, 4, 1, 4, 5, (

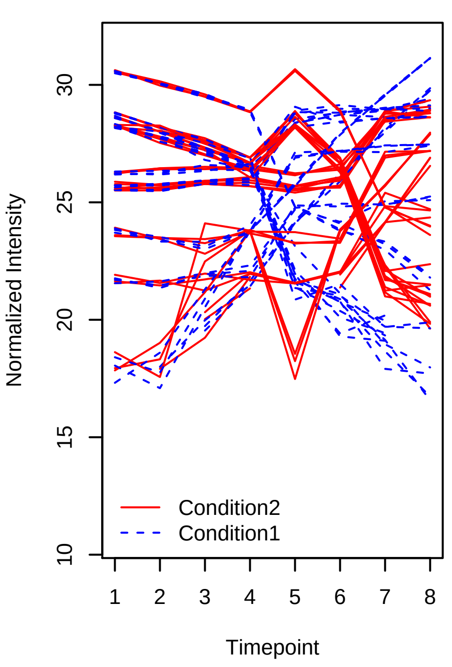


Spike−in proteins SGSDS Data Sigmoid_PolyHigher (8, 7, 6, 5, 4, 3, 2, 1 _ 1, 2, 3, 4, 1, 4, 5, (

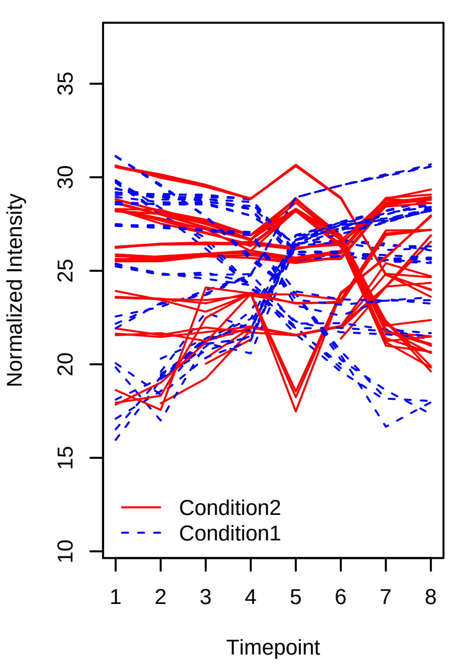

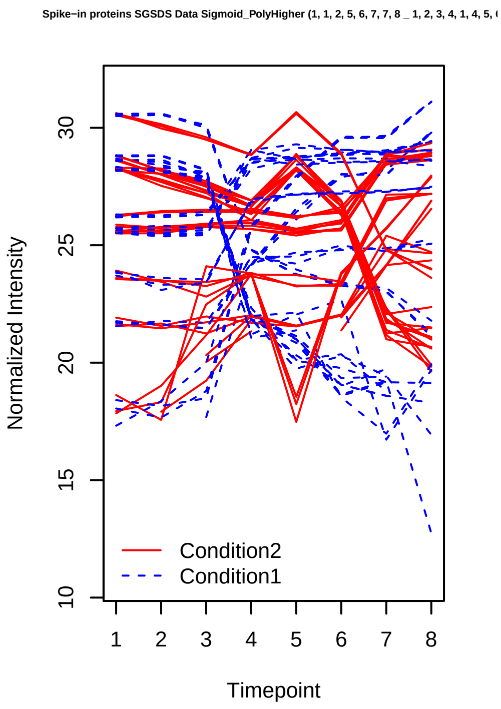
Spike−in proteins SGSDS Data Sigmoid_PolyHigher (1, 1, 2, 5, 6, 7, 7, 8 _ 1, 2, 3, 4, 1, 4, 5,
Normalized Intensity
10
15
20
25
30
1 2 3 4 5 6 7 8
Timepoint
Condition2
Condition1

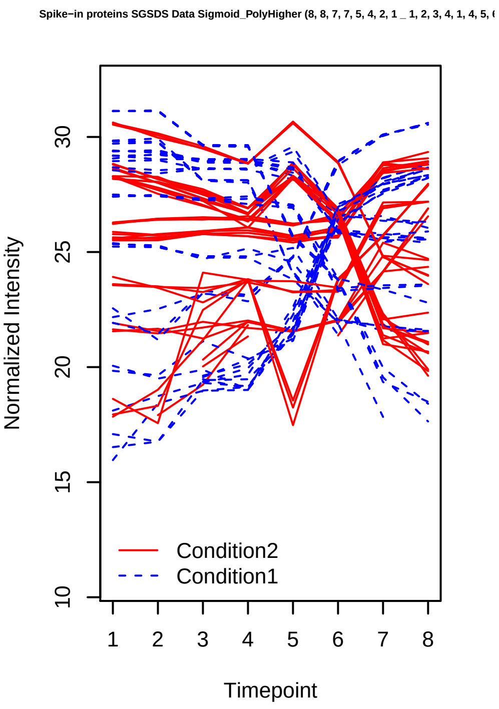
Spike−in proteins SGSDS Data Sigmoid_PolyHigher (8, 8, 7, 7, 5, 4, 2, 1 _ 1, 2, 3, 4, 1, 4, 5,
Normalized Intensity
10
15
20
25
30
1 2 3 4 5 6 7 8
Timepoint
Condition2
Condition1

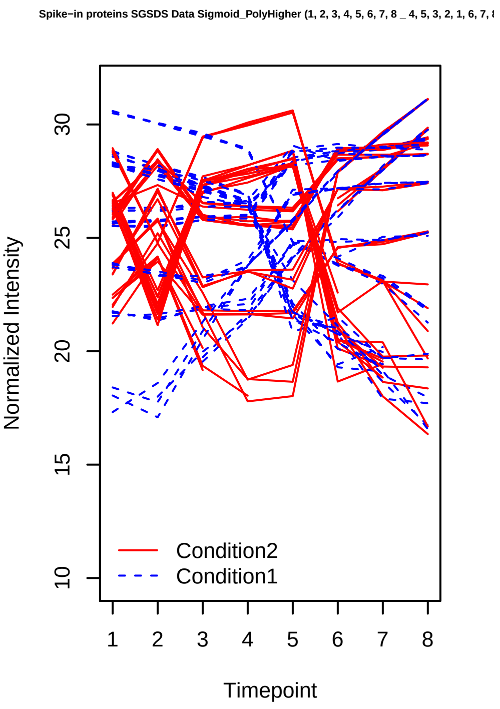
Spike−in proteins SGSDS Data Sigmoid_PolyHigher (1, 2, 3, 4, 5, 6, 7, 8 _ 4, 5, 3, 2, 1, 6, 7,
Normalized Intensity
10
15
20
25
30
1 2 3 4 5 6 7 8
Timepoint
Condition2
Condition1

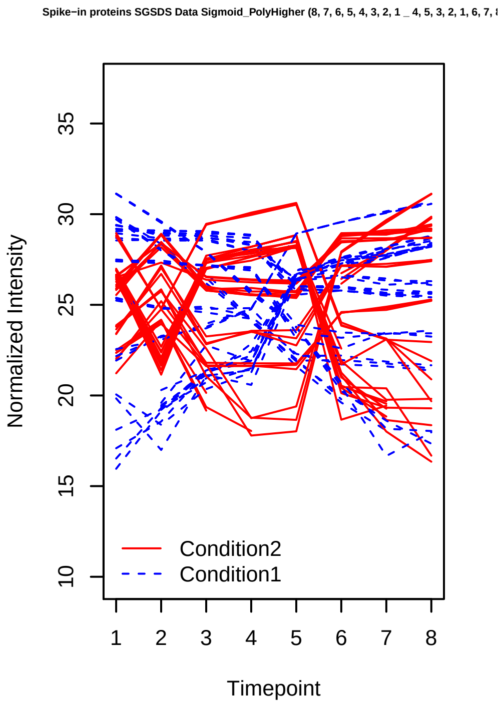
Spike−in proteins SGSDS Data Sigmoid_PolyHigher (8, 7, 6, 5, 4, 3, 2, 1 _ 4, 5, 3, 2, 1, 6, 7,
Normalized Intensity
10
15
20
25
30
35
1 2 3 4 5 6 7 8
Timepoint
Condition2
Condition1

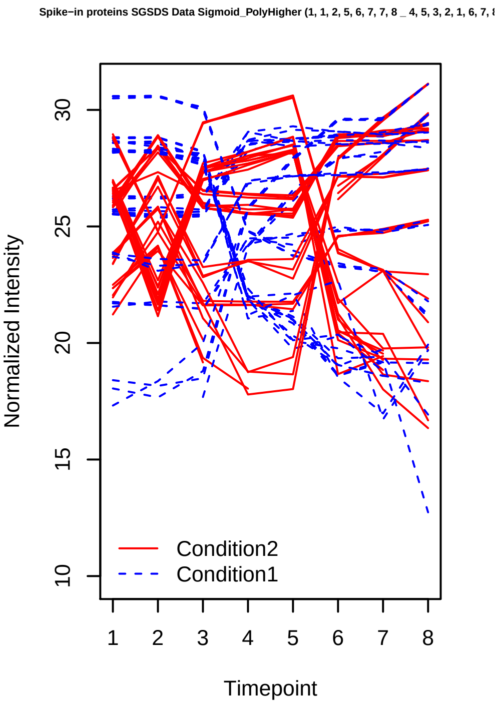
Spike−in proteins SGSDS Data Sigmoid_PolyHigher (1, 1, 2, 5, 6, 7, 7, 8 _ 4, 5, 3, 2, 1, 6, 7,
Normalized Intensity
10
15
20
25
30
1 2 3 4 5 6 7 8
Timepoint
Condition2
Condition1

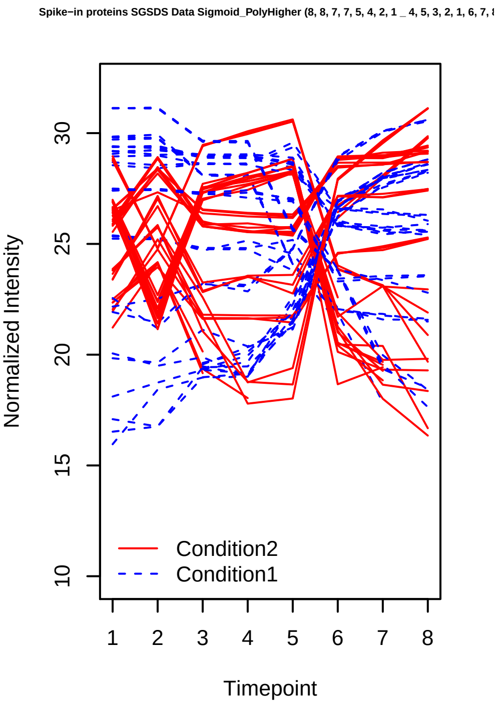
Spike−in proteins SGSDS Data Sigmoid_PolyHigher (8, 8, 7, 7, 5, 4, 2, 1 _ 4, 5, 3, 2, 1, 6, 7,
Normalized Intensity
10
15
20
25
30
1 2 3 4 5 6 7 8
Timepoint
Condition2
Condition1

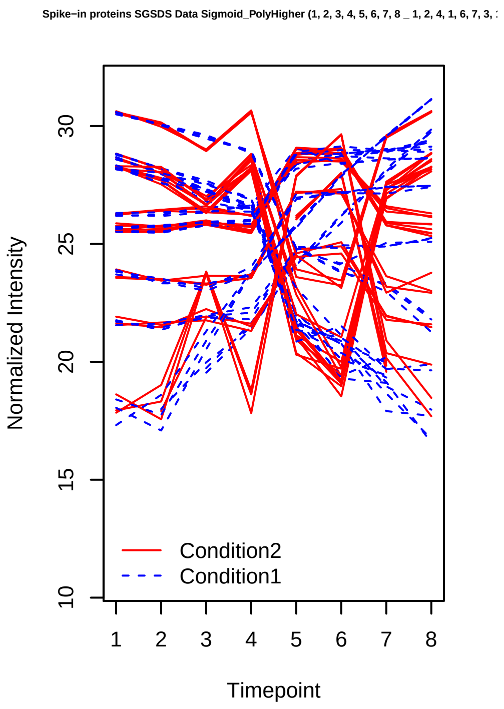
Spike−in proteins SGSDS Data Sigmoid_PolyHigher (1, 2, 3, 4, 5, 6, 7, 8 _ 1, 2, 4, 1, 6, 7, 3,
Normalized Intensity
Timepoint
Condition2
Condition1

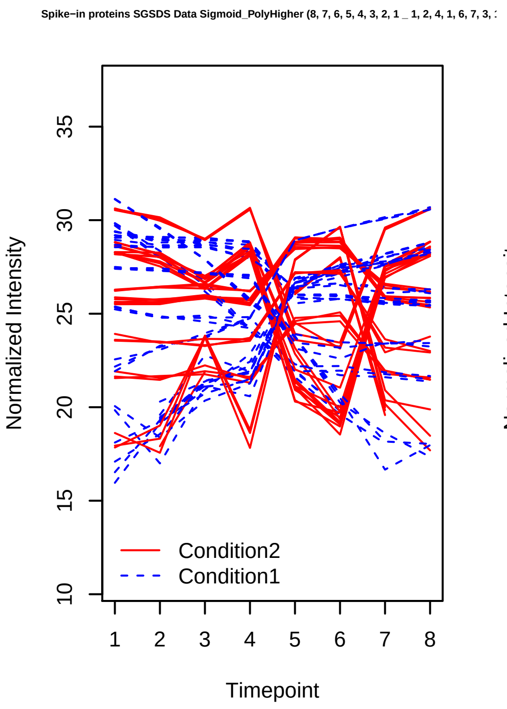
Spike−in proteins SGSDS Data Sigmoid_PolyHigher (8, 7, 6, 5, 4, 3, 2, 1 _ 1, 2, 4, 1, 6, 7, 3,
Normalized Intensity
Timepoint
Condition2
Condition1

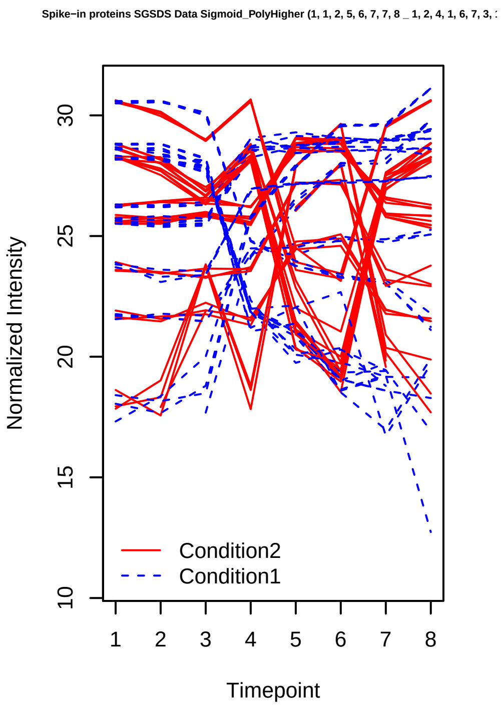
Spike−in proteins SGSDS Data Sigmoid_PolyHigher (1, 1, 2, 5, 6, 7, 7, 8 _ 1, 2, 4, 1, 6, 7, 3,
Normalized Intensity
Timepoint
Condition2
Condition1

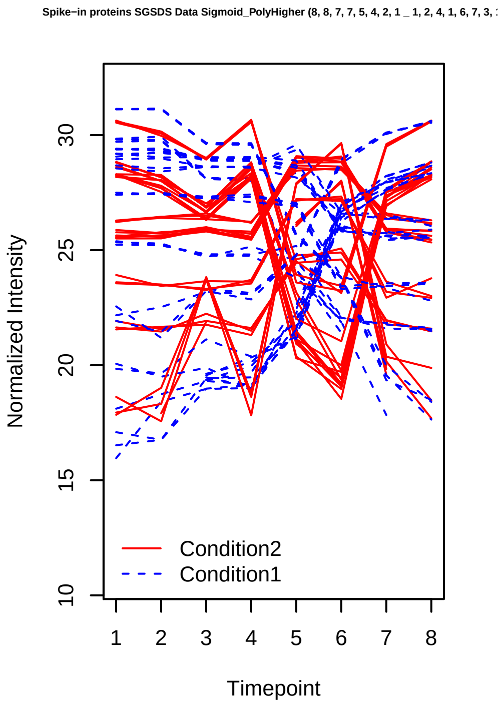
Spike−in proteins SGSDS Data Sigmoid_PolyHigher (8, 8, 7, 7, 5, 4, 2, 1 _ 1, 2, 4, 1, 6, 7, 3,
Normalized Intensity
Timepoint
Condition2
Condition1

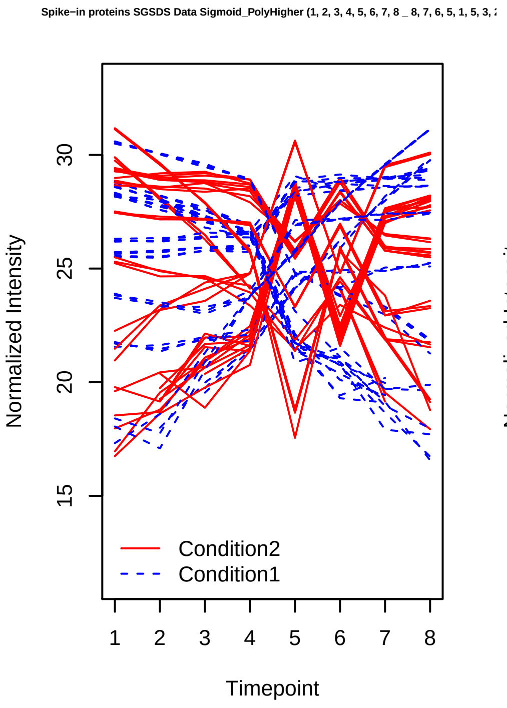
Spike−in proteins SGSDS Data Sigmoid_PolyHigher (1, 2, 3, 4, 5, 6, 7, 8 _ 8, 7, 6, 5, 1, 5, 3,
Normalized Intensity
Timepoint
Condition2
Condition1

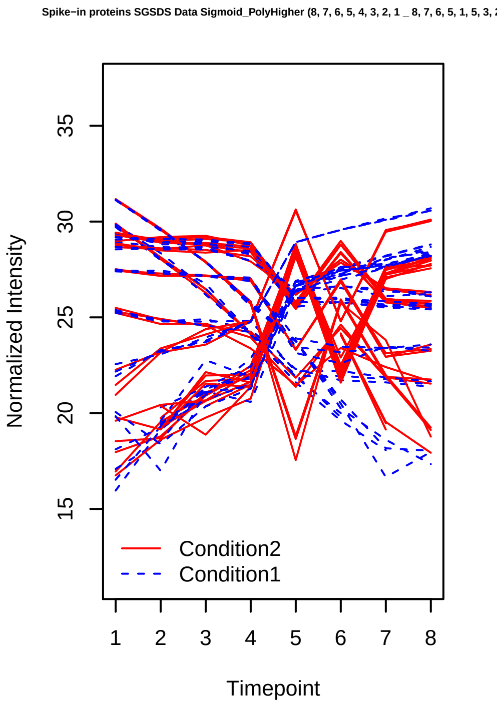
Spike−in proteins SGSDS Data Sigmoid_PolyHigher (8, 7, 6, 5, 4, 3, 2, 1 _ 8, 7, 6, 5, 1, 5, 3,
Normalized Intensity
Timepoint
Condition2
Condition1

Spike−in proteins SGSDS Data Sigmoid_PolyHigher (1, 1, 2, 5, 6, 7, 7, 8 _ 8, 7, 6, 5, 1, 5, 3,

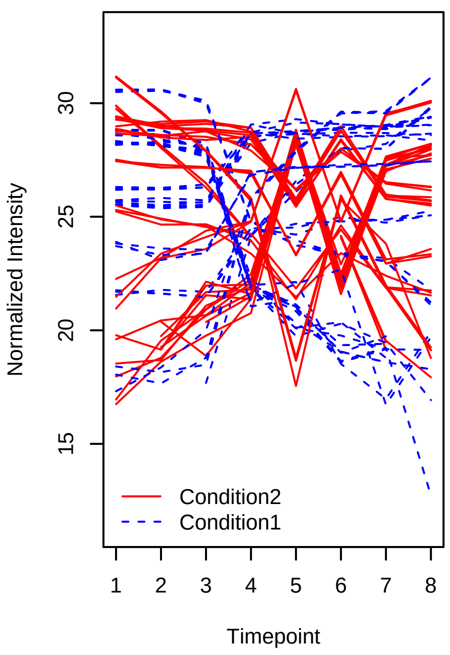


Spike−in proteins SGSDS Data Sigmoid_PolyHigher (8, 8, 7, 7, 5, 4, 2, 1 _ 8, 7, 6, 5, 1, 5, 3,

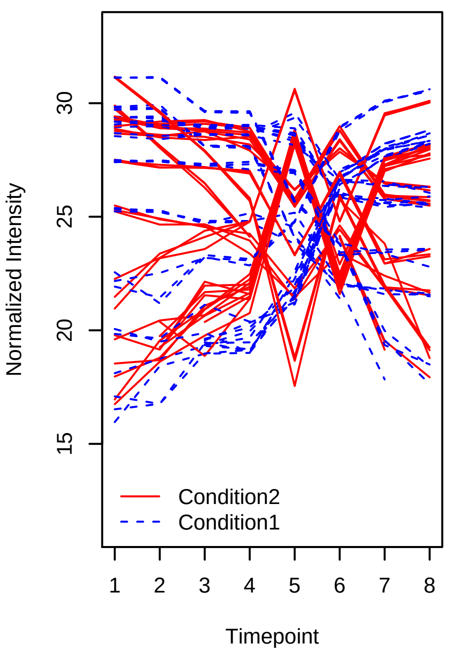


Spike−in proteins SGSDS Data PolyHigher_PolyHigher (4, 5, 3, 2, 1, 6, 7, 8 _ 1, 2, 3, 4, 1, 4, 5,

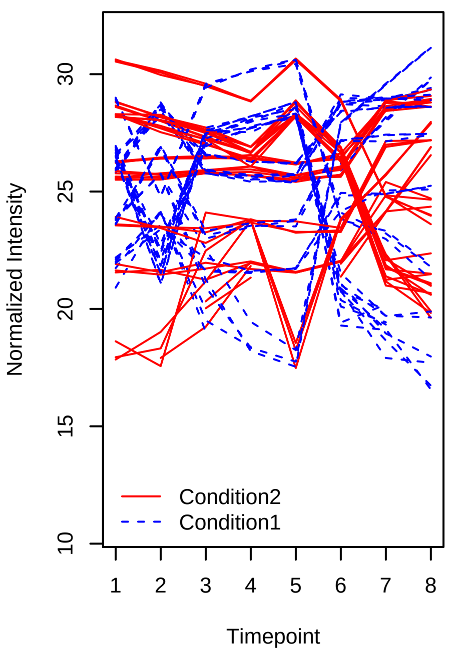


Spike−in proteins SGSDS Data PolyHigher_PolyHigher (1, 2, 4, 1, 6, 7, 3, 1 _ 1, 2, 3, 4, 1, 4, 5,

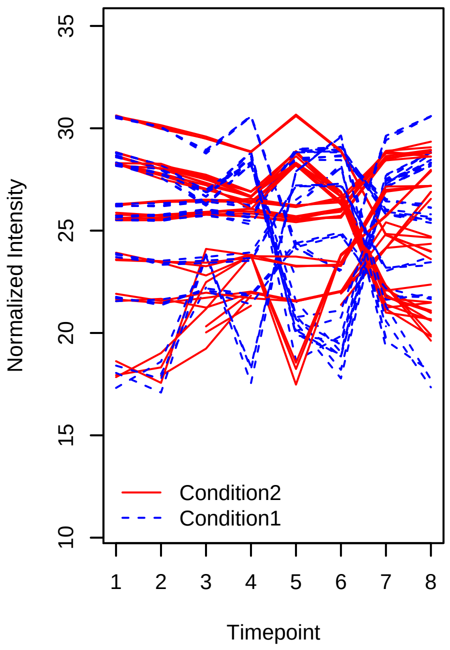


Spike−in proteins SGSDS Data PolyHigher_PolyHigher (8, 7, 6, 5, 1, 5, 3, 2 _ 1, 2, 3, 4, 1, 4, 5,

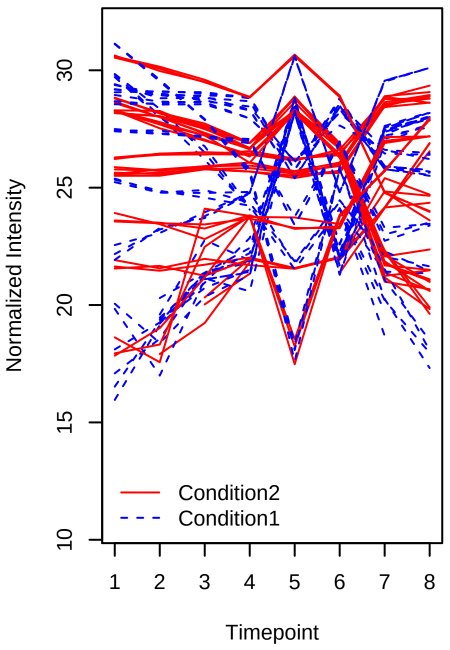


Spike−in proteins SGSDS Data PolyHigher_PolyHigher (5, 4, 6, 7, 8, 3, 2, 1 _ 1, 2, 3, 4, 1, 4, 5,

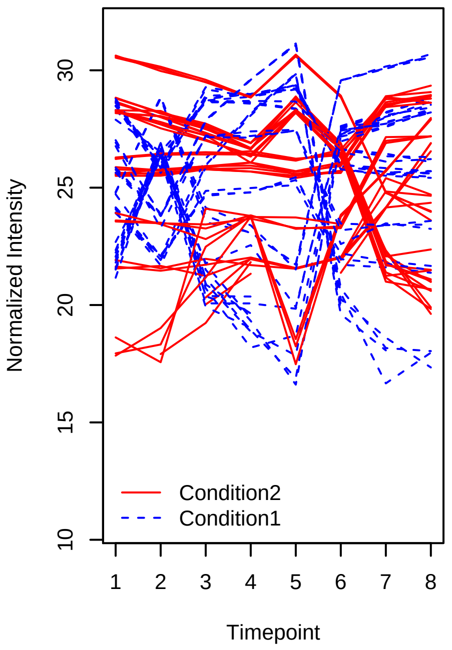

Spike−in proteins SGSDS Data PolyHigher_PolyHigher (1, 2, 4, 1, 6, 7, 3, 1 _ 4, 5, 3, 2, 1, 6, 7,

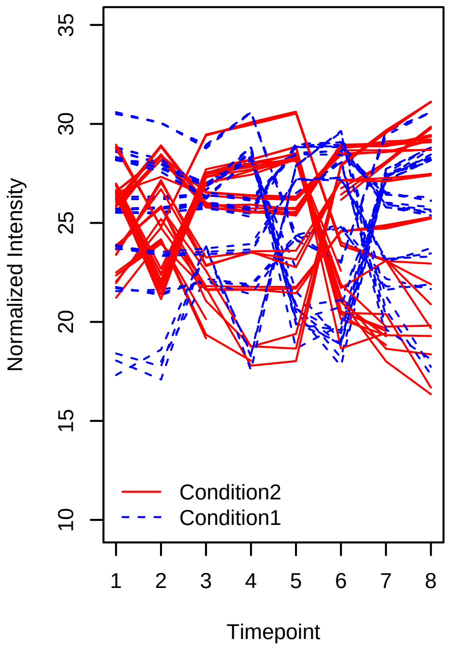


Spike−in proteins SGSDS Data PolyHigher_PolyHigher (8, 7, 6, 5, 1, 5, 3, 2 _ 4, 5, 3, 2, 1, 6, 7,

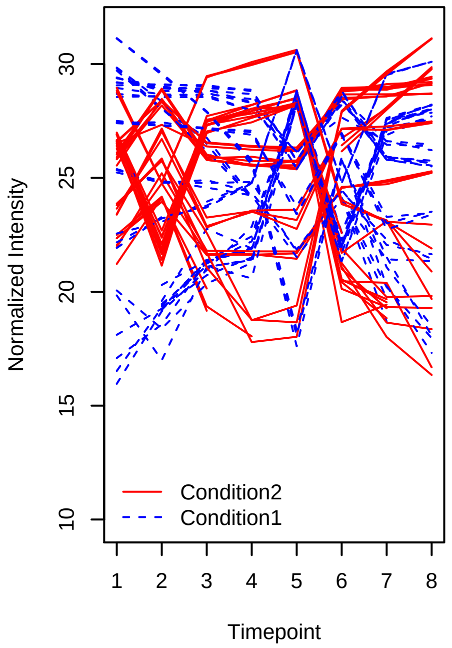


Spike−in proteins SGSDS Data PolyHigher_PolyHigher (5, 4, 6, 7, 8, 3, 2, 1 _ 4, 5, 3, 2, 1, 6, 7,

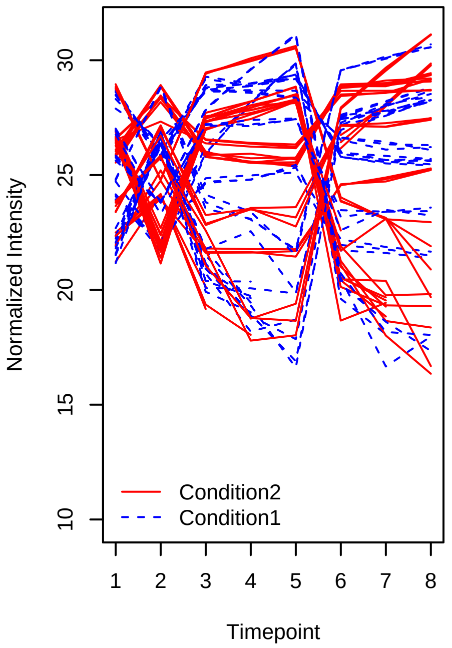


Spike−in proteins SGSDS Data PolyHigher_PolyHigher (8, 7, 6, 5, 1, 5, 3, 2 _ 1, 2, 4, 1, 6, 7, 3,

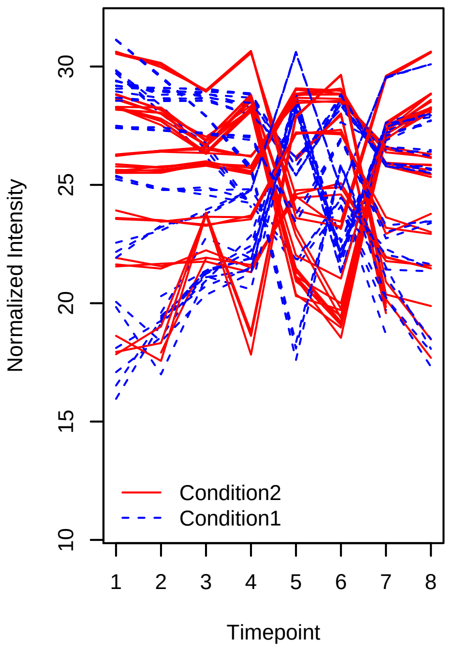


Spike−in proteins SGSDS Data PolyHigher_PolyHigher (5, 4, 6, 7, 8, 3, 2, 1 _ 1, 2, 4, 1, 6, 7, 3,

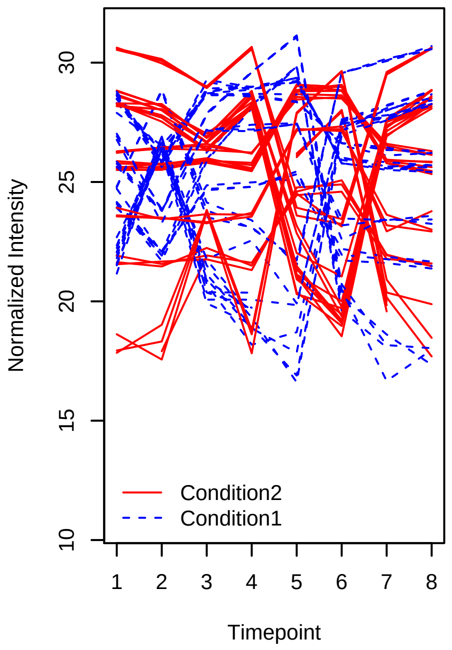


Spike−in proteins SGSDS Data PolyHigher_PolyHigher (5, 4, 6, 7, 8, 3, 2, 1 _ 8, 7, 6, 5, 1, 5, 3,

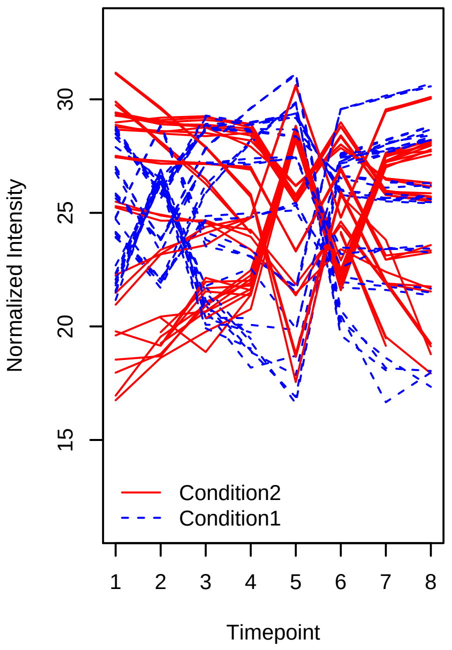

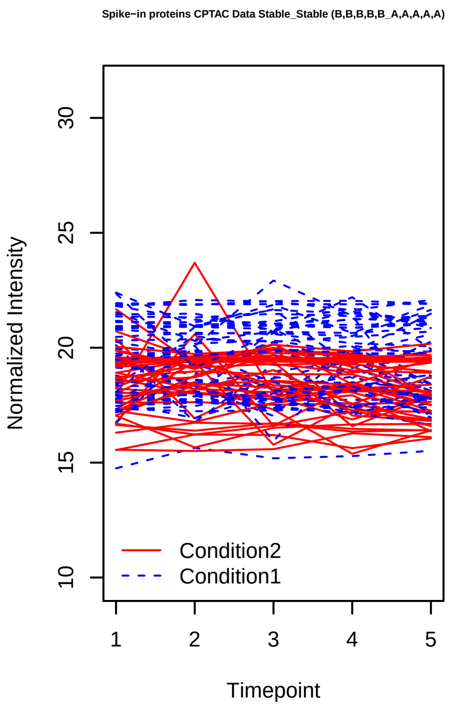
Spike−in proteins CPTAC Data Stable_Stable (B,B,B,B,B_A,A,A,A,A)
Normalized Intensity
10
15
20
25
30
Condition2
Condition1
1
2
3
4
5
Timepoint

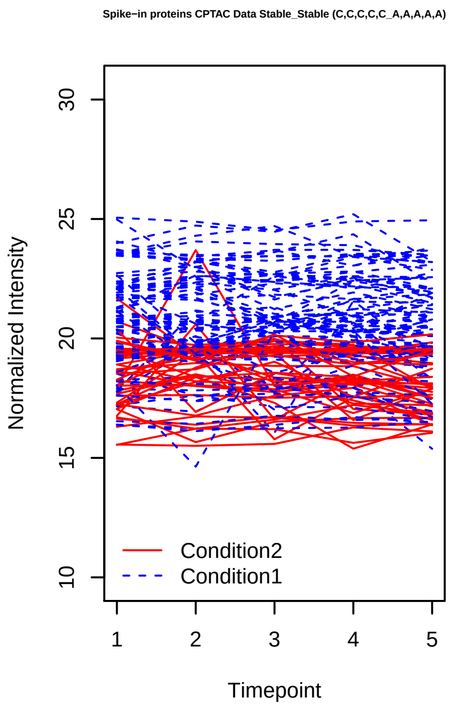
Spike−in proteins CPTAC Data Stable_Stable (C,C,C,C,C_A,A,A,A,A)
Normalized Intensity
10
15
20
25
30
Condition2
Condition1
1
2
3
4
5
Timepoint

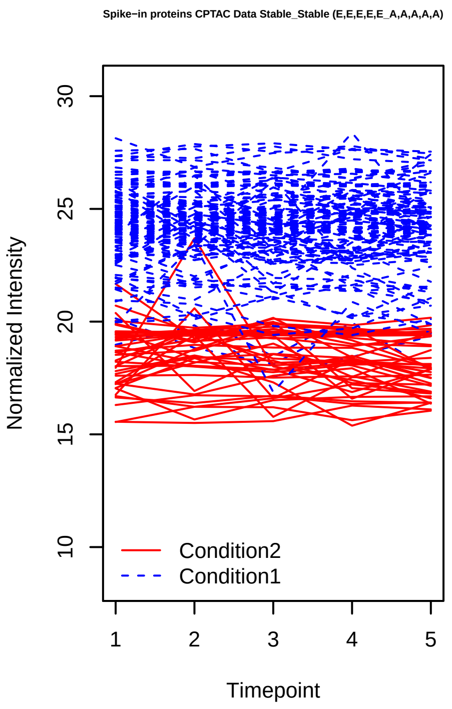
Spike−in proteins CPTAC Data Stable_Stable (E,E,E,E,E_A,A,A,A,A)
Normalized Intensity
10
15
20
25
30
Condition2
Condition1
1
2
3
4
5
Timepoint

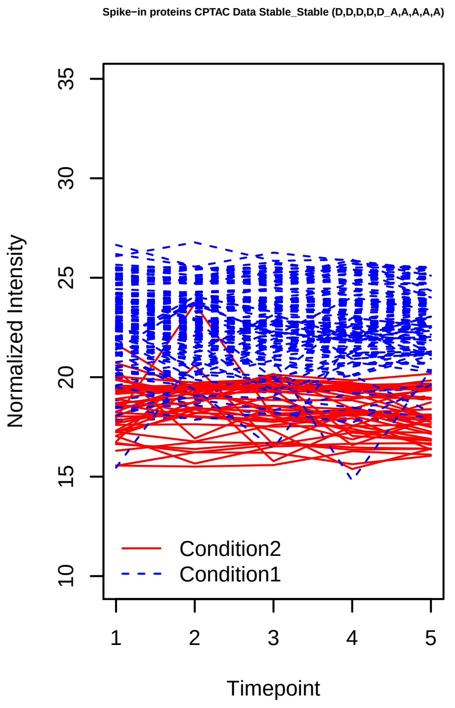
Spike−in proteins CPTAC Data Stable_Stable (D,D,D,D,D_A,A,A,A,A)
Normalized Intensity
10
15
20
25
30
35
Condition2
Condition1
1
2
3
4
5
Timepoint

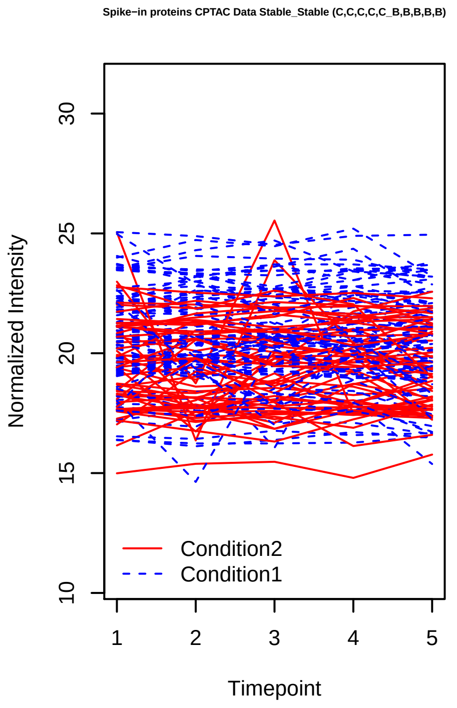
Spike−in proteins CPTAC Data Stable_Stable (C,C,C,C,C_B,B,B,B,B)
Normalized Intensity
10
15
20
25
30
Condition2
Condition1
1
2
3
4
5
Timepoint

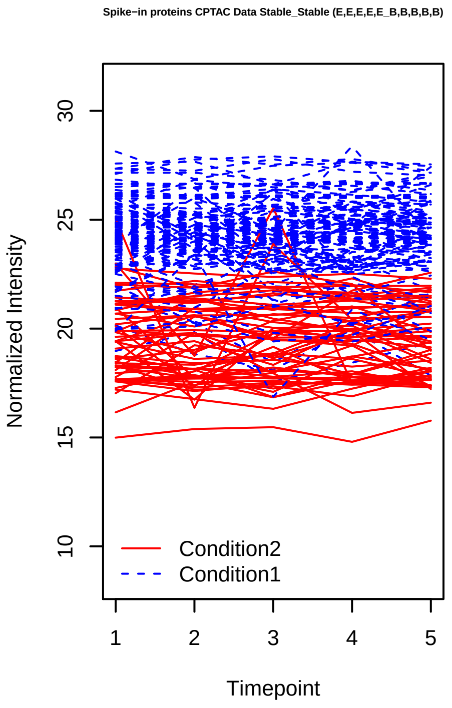
Spike−in proteins CPTAC Data Stable_Stable (E,E,E,E,E_B,B,B,B,B)
Normalized Intensity
10
15
20
25
30
Condition2
Condition1
1
2
3
4
5
Timepoint

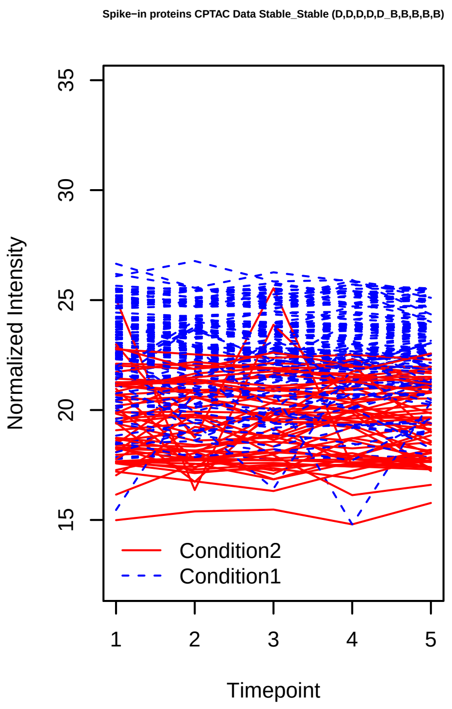
Spike−in proteins CPTAC Data Stable_Stable (D,D,D,D,D_B,B,B,B,B)
Normalized Intensity
15
20
25
30
35
Timepoint
1
2
3
4
5
Condition2
Condition1

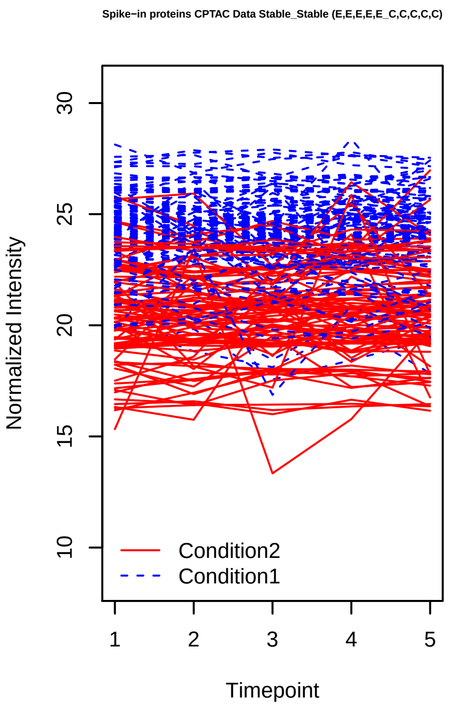
Spike−in proteins CPTAC Data Stable_Stable (E,E,E,E,E_C,C,C,C,C)
Normalized Intensity
10
15
20
25
30
Timepoint
1
2
3
4
5
Condition2
Condition1

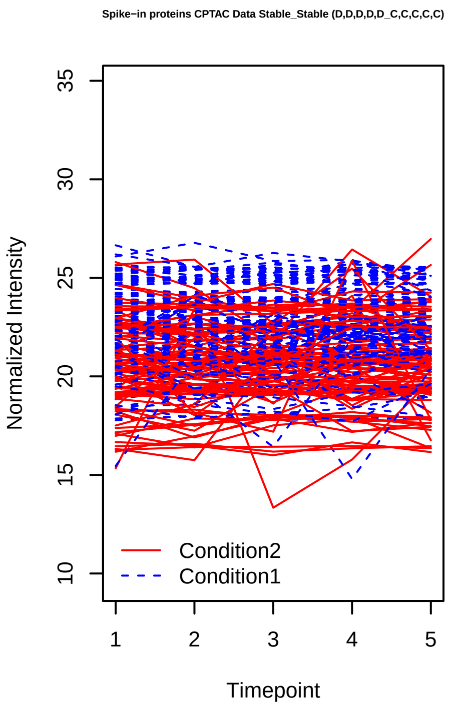
Spike−in proteins CPTAC Data Stable_Stable (D,D,D,D,D_C,C,C,C,C)
Normalized Intensity
10
15
20
25
30
35
Timepoint
1
2
3
4
5
Condition2
Condition1

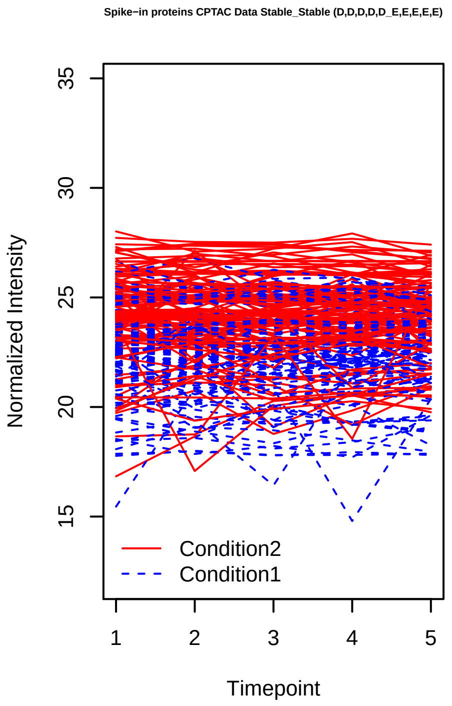
Spike−in proteins CPTAC Data Stable_Stable (D,D,D,D,D_E,E,E,E,E)
Normalized Intensity
15
20
25
30
35
Timepoint
1
2
3
4
5
Condition2
Condition1

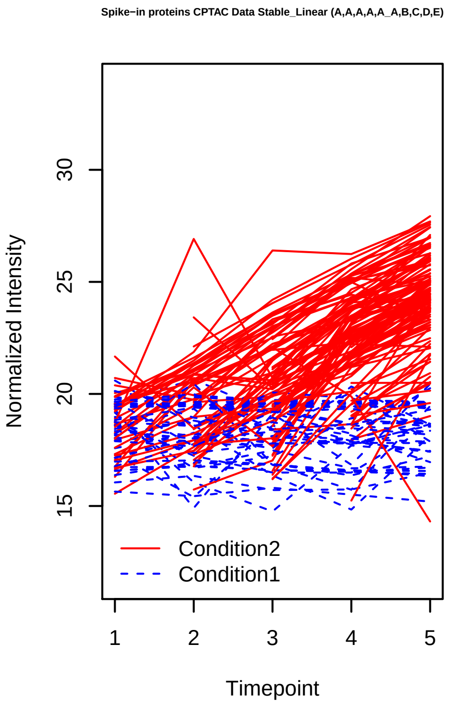
Spike−in proteins CPTAC Data Stable_Linear (A,A,A,A,A_A,B,C,D,E)
Normalized Intensity
15
20
25
30
Timepoint
1
2
3
4
5
Condition2
Condition1

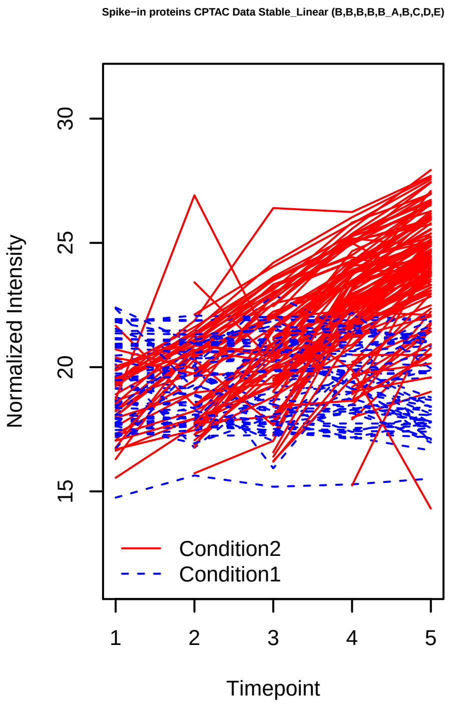
Spike−in proteins CPTAC Data Stable_Linear (B,B,B,B,B_A,B,C,D,E)
Normalized Intensity
15
20
25
30
Timepoint
1
2
3
4
5
Condition2
Condition1

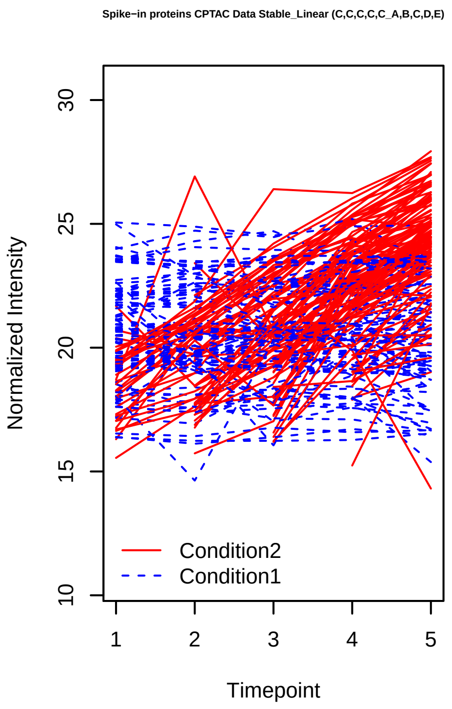
Spike−in proteins CPTAC Data Stable_Linear (C,C,C,C,C_A,B,C,D,E)
Normalized Intensity
10
15
20
25
30
1
2
3
4
5
Timepoint
Condition2
Condition1

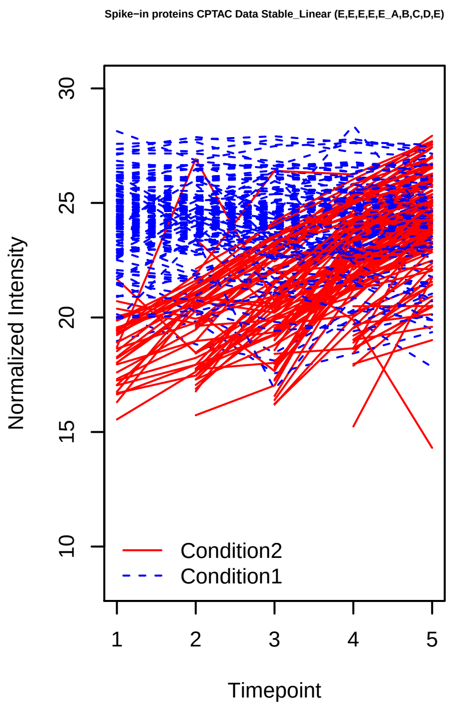
Spike−in proteins CPTAC Data Stable_Linear (E,E,E,E,E_A,B,C,D,E)
Normalized Intensity
10
15
20
25
30
1
2
3
4
5
Timepoint
Condition2
Condition1

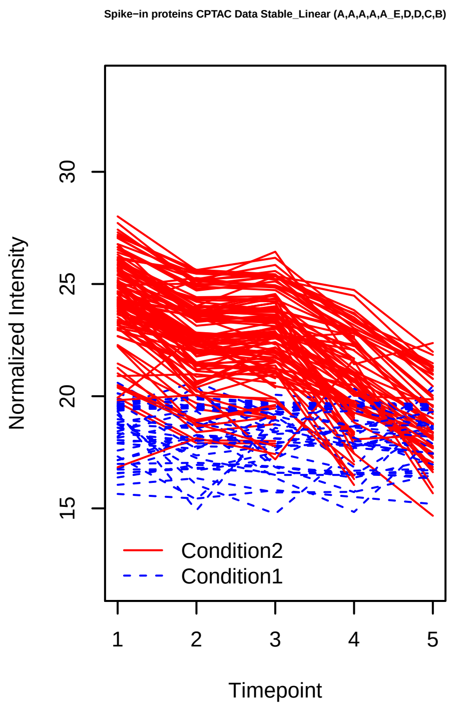
Spike−in proteins CPTAC Data Stable_Linear (A,A,A,A,A_E,D,D,C,B)
Normalized Intensity
15
20
25
30
1
2
3
4
5
Timepoint
Condition2
Condition1

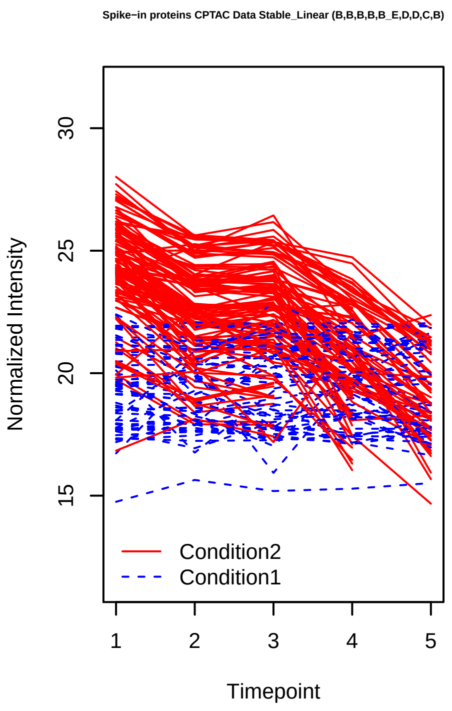
Spike−in proteins CPTAC Data Stable_Linear (B,B,B,B,B_E,D,D,C,B)
Normalized Intensity
15
20
25
30
1
2
3
4
5
Timepoint
Condition2
Condition1

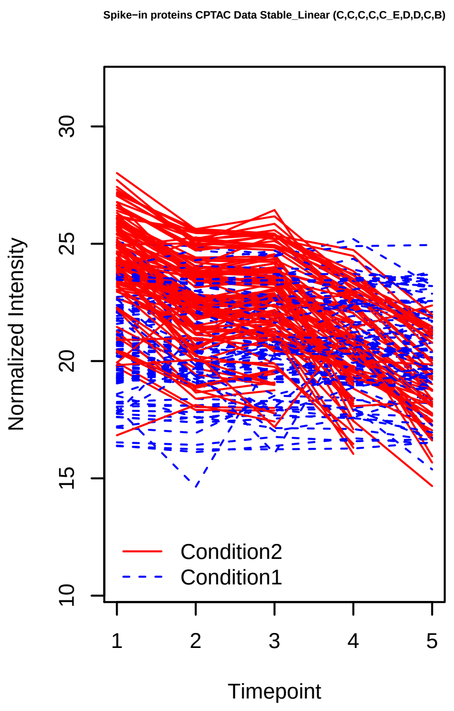
Spike−in proteins CPTAC Data Stable_Linear (C,C,C,C,C_E,D,D,C,B)
Normalized Intensity
10
15
20
25
30
1
2
3
4
5
Timepoint
Condition2
Condition1

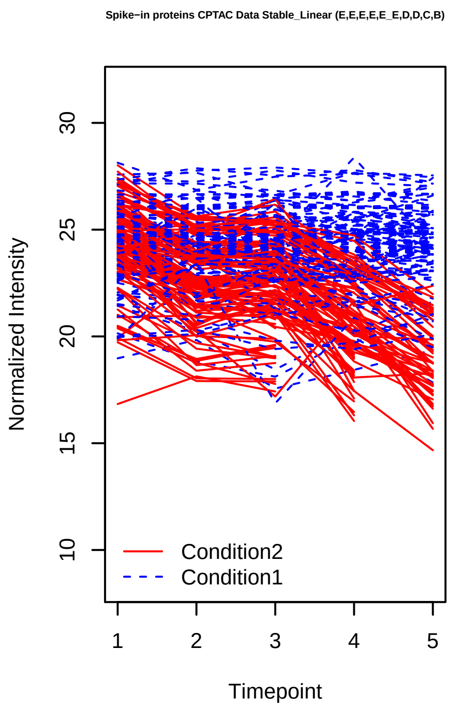
Spike−in proteins CPTAC Data Stable_Linear (E,E,E,E,E_E,D,D,C,B)
Normalized Intensity
10
15
20
25
30
1
2
3
4
5
Timepoint
Condition2
Condition1

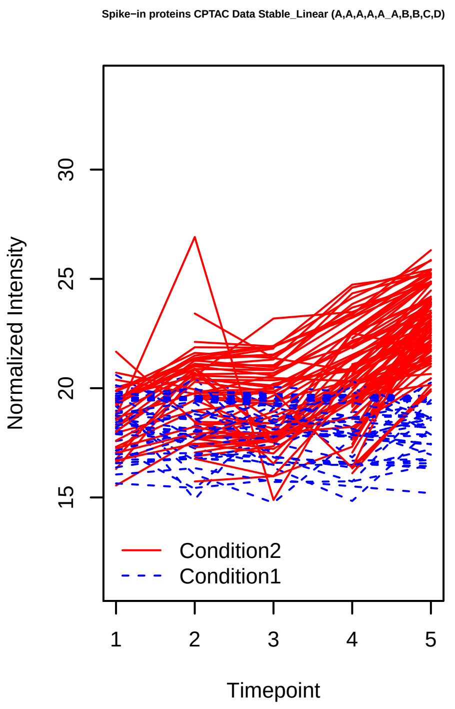
Spike−in proteins CPTAC Data Stable_Linear (A,A,A,A,A_A,B,B,C,D)
Normalized Intensity
Timepoint
Condition2
Condition1

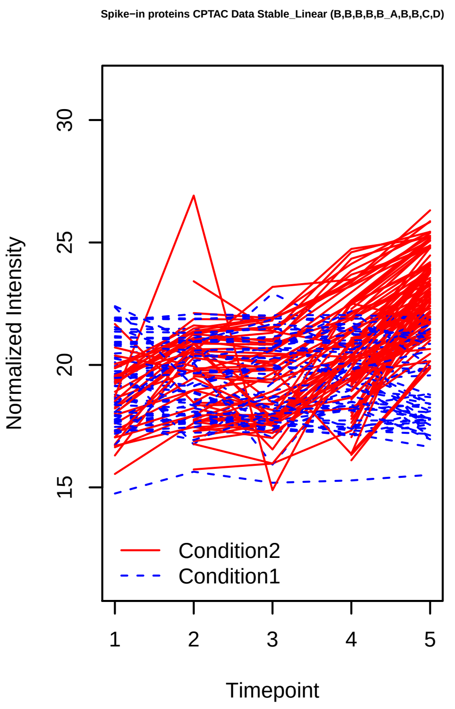
Spike−in proteins CPTAC Data Stable_Linear (B,B,B,B,B_A,B,B,C,D)
Normalized Intensity
Timepoint
Condition2
Condition1

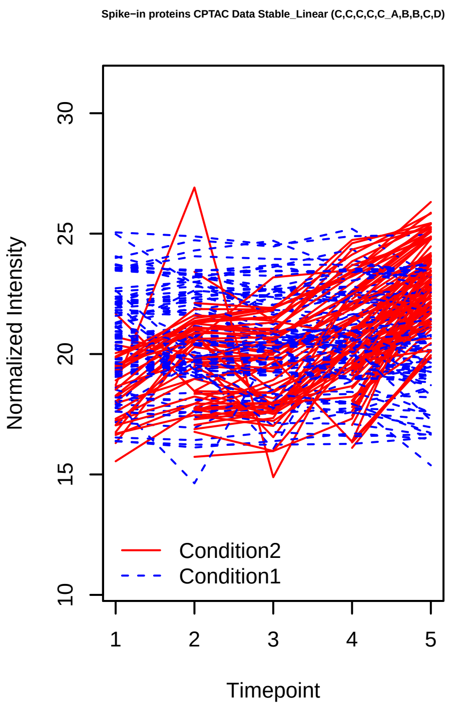
Spike−in proteins CPTAC Data Stable_Linear (C,C,C,C,C_A,B,B,C,D)
Normalized Intensity
Timepoint
Condition2
Condition1

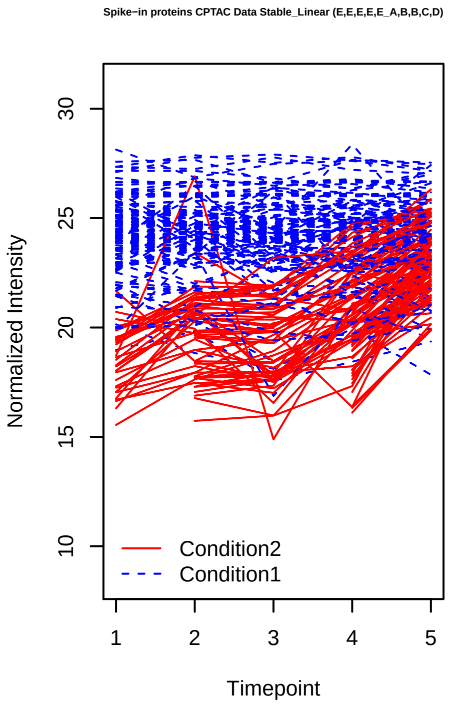
Spike−in proteins CPTAC Data Stable_Linear (E,E,E,E,E_A,B,B,C,D)
Normalized Intensity
Timepoint
Condition2
Condition1

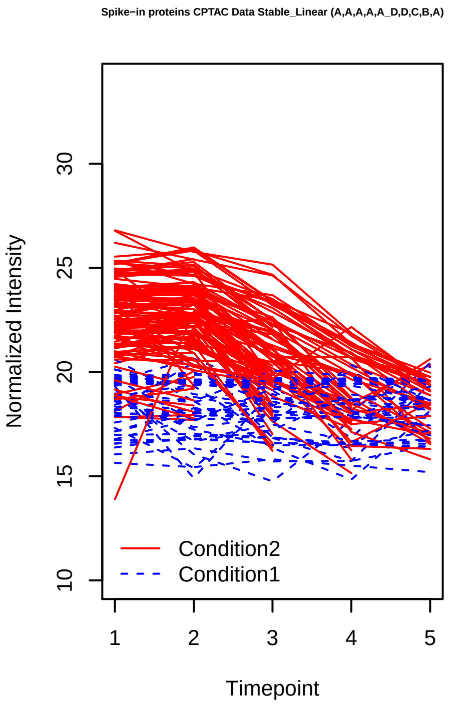
Spike−in proteins CPTAC Data Stable_Linear (A,A,A,A,A_D,D,C,B,A)
Normalized Intensity
Timepoint
Condition2
Condition1

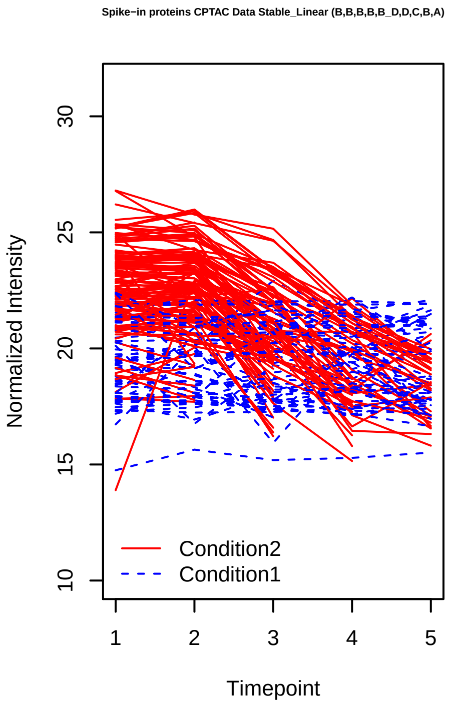
Spike−in proteins CPTAC Data Stable_Linear (B,B,B,B,B_D,D,C,B,A)
Normalized Intensity
Timepoint
Condition2
Condition1

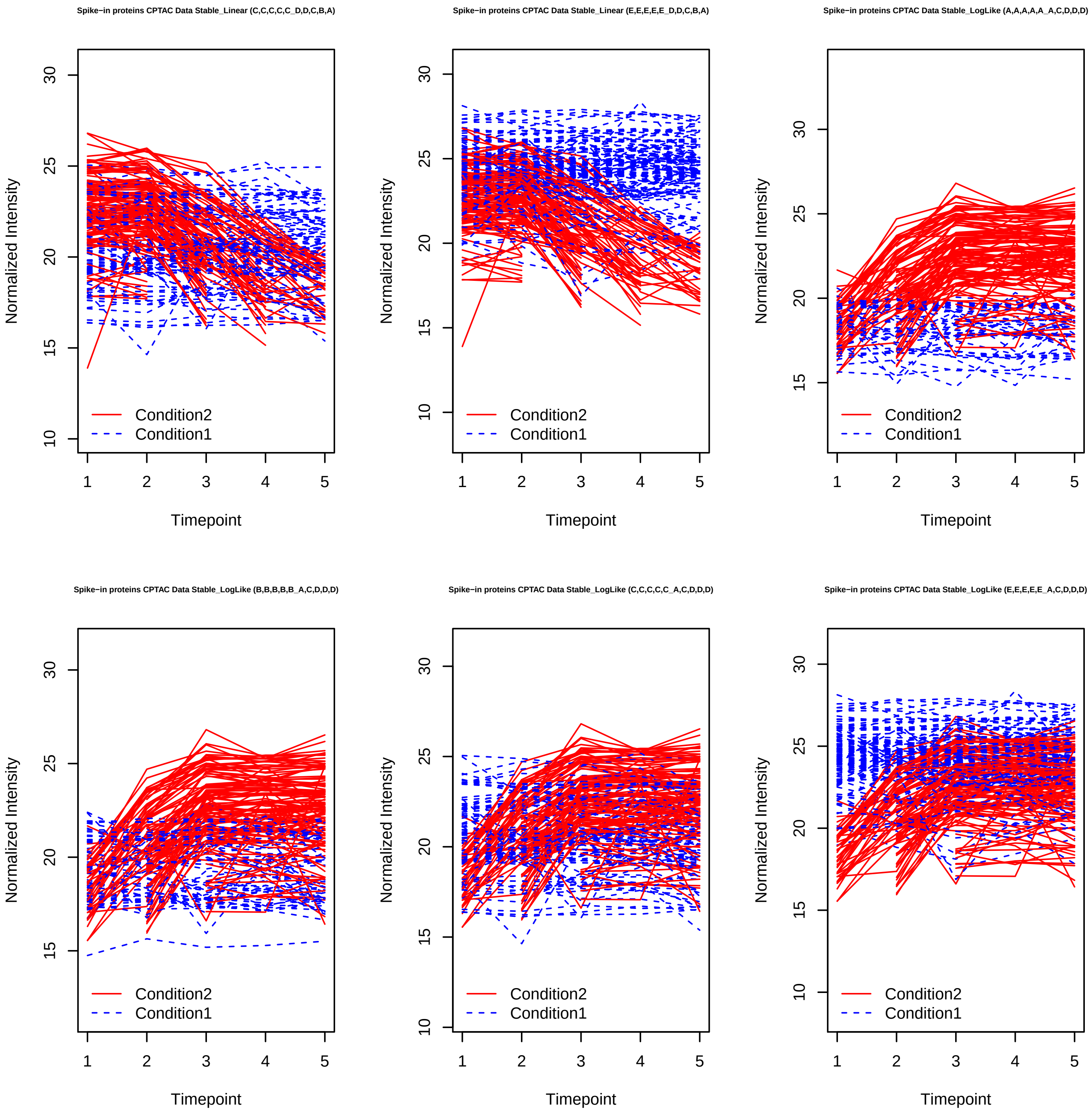
Spike−in proteins CPTAC Data Stable_Linear (C,C,C,C,C_D,D,C,B,A)
Spike−in proteins CPTAC Data Stable_Linear (E,E,E,E,E_D,D,C,B,A)
Spike−in proteins CPTAC Data Stable_LogLike (A,A,A,A,A_A,C,D,D,D)
Spike−in proteins CPTAC Data Stable_LogLike (B,B,B,B,B_A,C,D,D,D)
Spike−in proteins CPTAC Data Stable_LogLike (C,C,C,C,C_A,C,D,D,D)
Spike−in proteins CPTAC Data Stable_LogLike (E,E,E,E,E_A,C,D,D,D)
Normalized Intensity
Timepoint
Condition2
Condition1

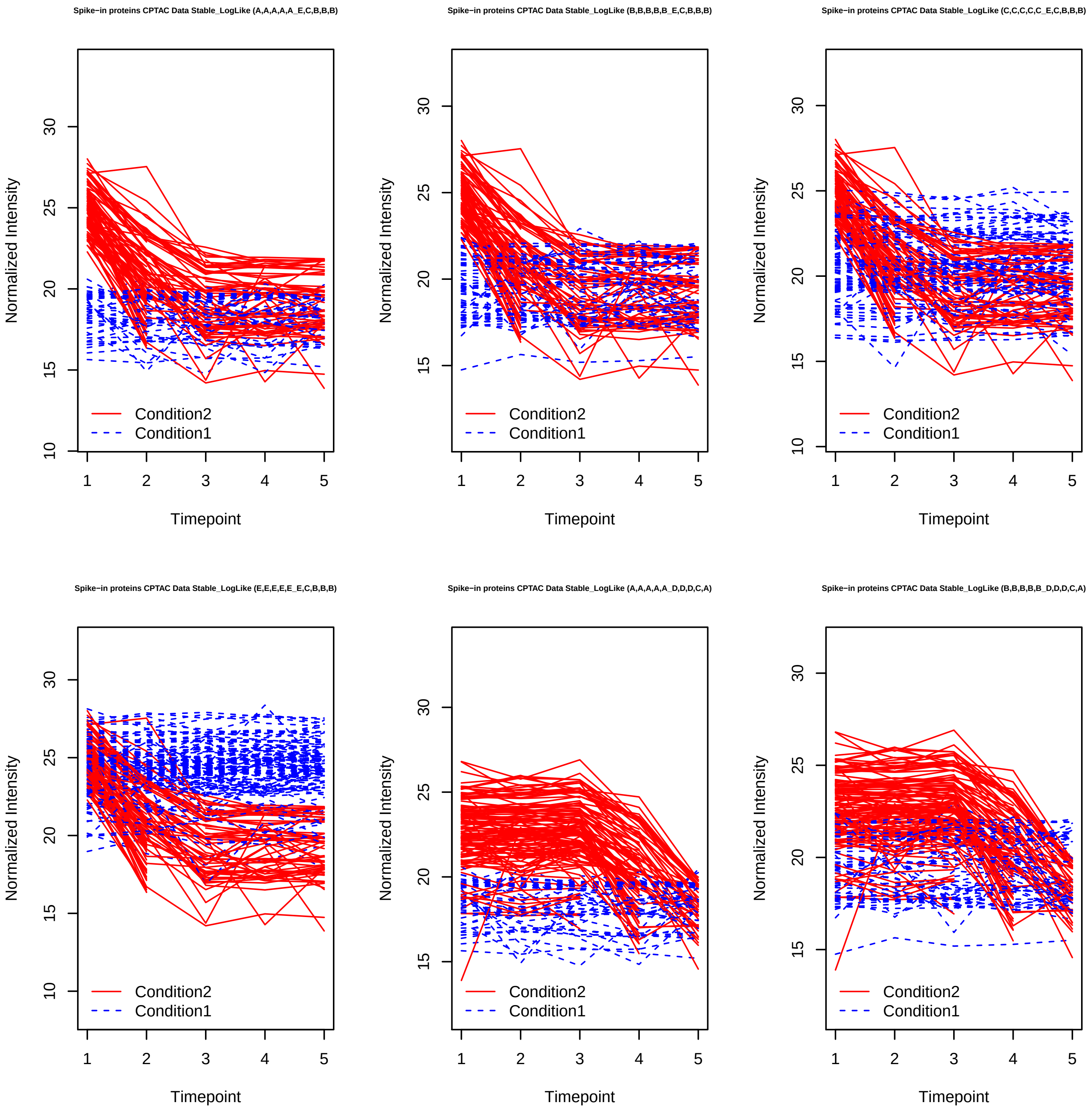
Spike−in proteins CPTAC Data Stable_LogLike (A,A,A,A,A_E,C,B,B,B)
Spike−in proteins CPTAC Data Stable_LogLike (B,B,B,B,B_E,C,B,B,B)
Spike−in proteins CPTAC Data Stable_LogLike (C,C,C,C,C_E,C,B,B,B)
Spike−in proteins CPTAC Data Stable_LogLike (E,E,E,E,E_E,C,B,B,B)
Spike−in proteins CPTAC Data Stable_LogLike (A,A,A,A,A_D,D,D,C,A)
Spike−in proteins CPTAC Data Stable_LogLike (B,B,B,B,B_D,D,D,C,A)
Normalized Intensity
Timepoint
Condition2
Condition1

Spike−in proteins CPTAC Data Stable_LogLike (C,C,C,C,C_D,D,D,C,A)

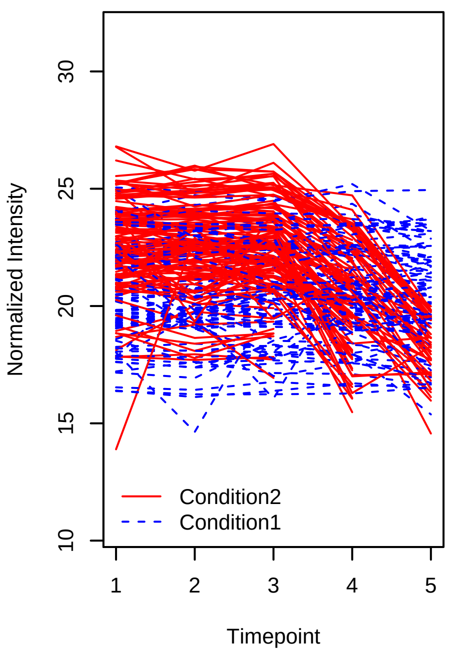


Spike−in proteins CPTAC Data Stable_LogLike (E,E,E,E,E_D,D,D,C,A)

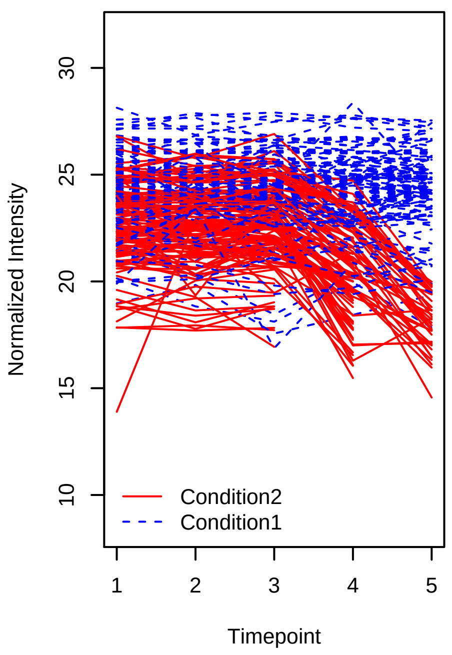


Spike−in proteins CPTAC Data Stable_LogLike (A,A,A,A,A_B,B,B,C,E)

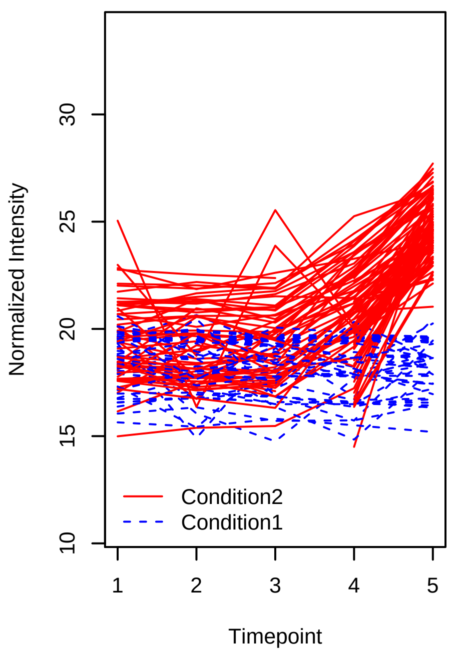


Spike−in proteins CPTAC Data Stable_LogLike (B,B,B,B,B_B,B,B,C,E)

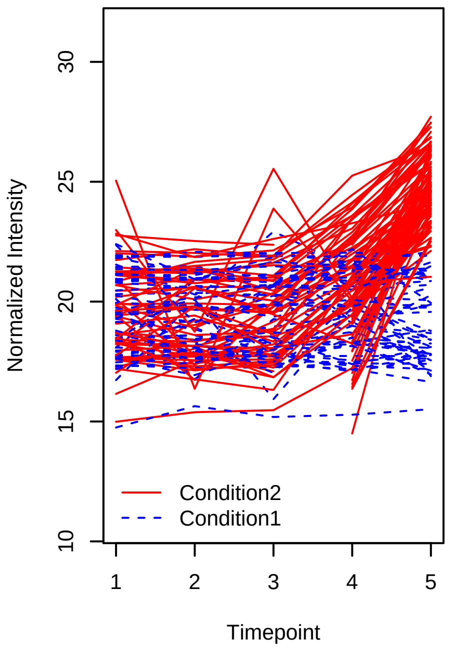


Spike−in proteins CPTAC Data Stable_LogLike (C,C,C,C,C_B,B,B,C,E)

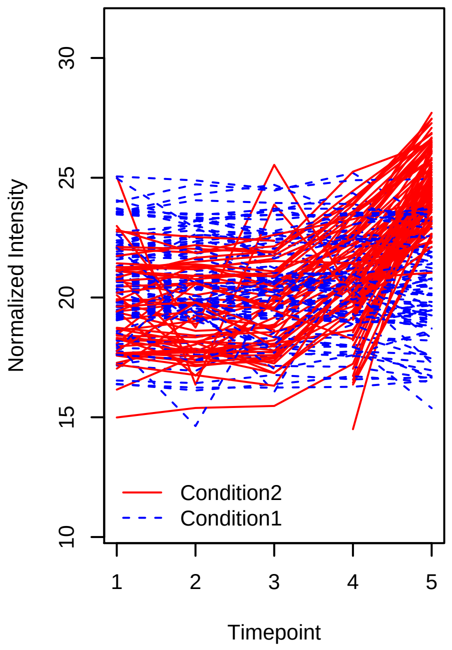


Spike−in proteins CPTAC Data Stable_LogLike (E,E,E,E,E_B,B,B,C,E)

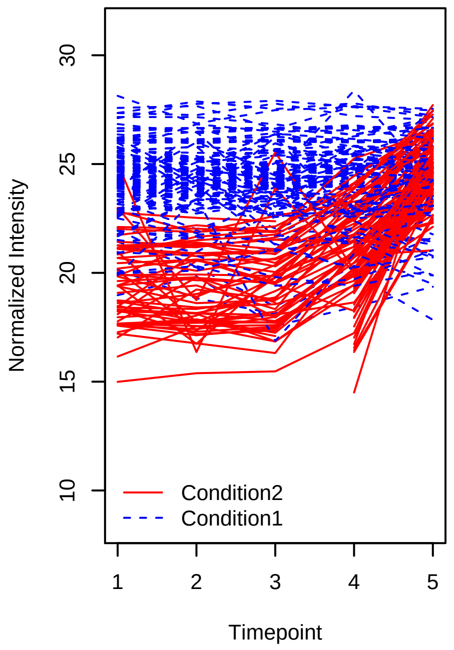

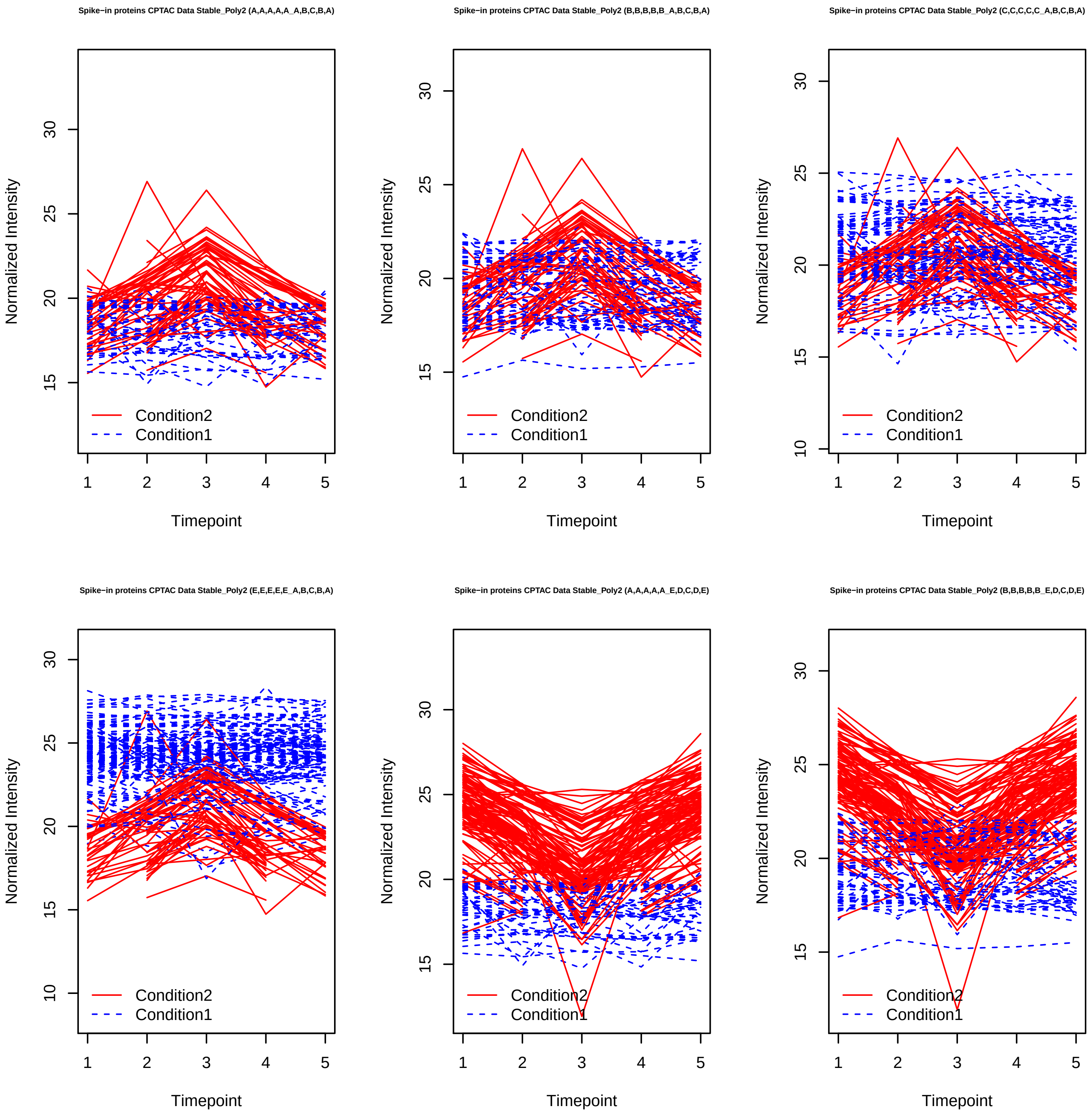
Spike−in proteins CPTAC Data Stable_Poly2 (A,A,A,A,A_A,B,C,B,A)
Normalized Intensity
Condition2
Condition1
Timepoint
Spike−in proteins CPTAC Data Stable_Poly2 (B,B,B,B,B_A,B,C,B,A)
Normalized Intensity
Condition2
Condition1
Timepoint
Spike−in proteins CPTAC Data Stable_Poly2 (C,C,C,C,C_A,B,C,B,A)
Normalized Intensity
Condition2
Condition1
Timepoint
Spike−in proteins CPTAC Data Stable_Poly2 (E,E,E,E,E_A,B,C,B,A)
Normalized Intensity
Condition2
Condition1
Timepoint
Spike−in proteins CPTAC Data Stable_Poly2 (A,A,A,A,A_E,D,C,D,E)
Normalized Intensity
Condition2
Condition1
Timepoint
Spike−in proteins CPTAC Data Stable_Poly2 (B,B,B,B,B_E,D,C,D,E)
Normalized Intensity
Condition2
Condition1
Timepoint

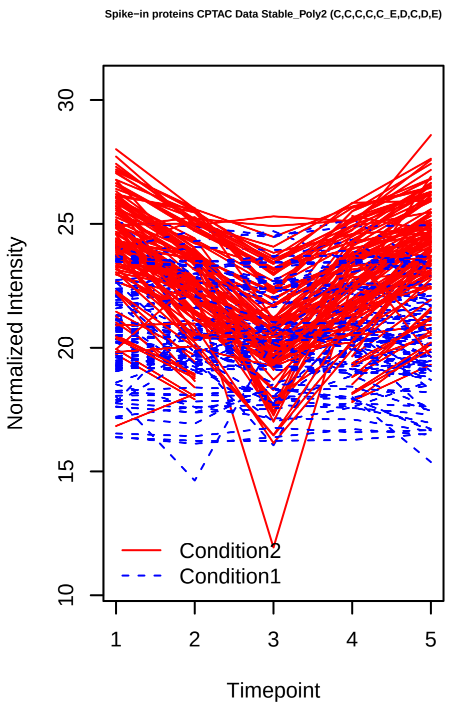

Spike−in proteins CPTAC Data Stable_Poly2 (C,C,C,C,C_E,D,C,D,E)
Normalized Intensity
10
15
20
25
30
1
2
3
4
5
Timepoint
Condition2
Condition1


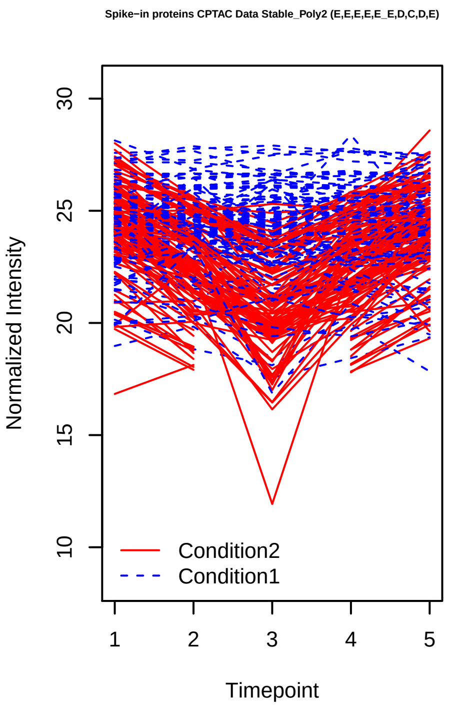

Spike−in proteins CPTAC Data Stable_Poly2 (E,E,E,E,E_E,D,C,D,E)
Normalized Intensity
10
15
20
25
30
1
2
3
4
5
Timepoint
Condition2
Condition1


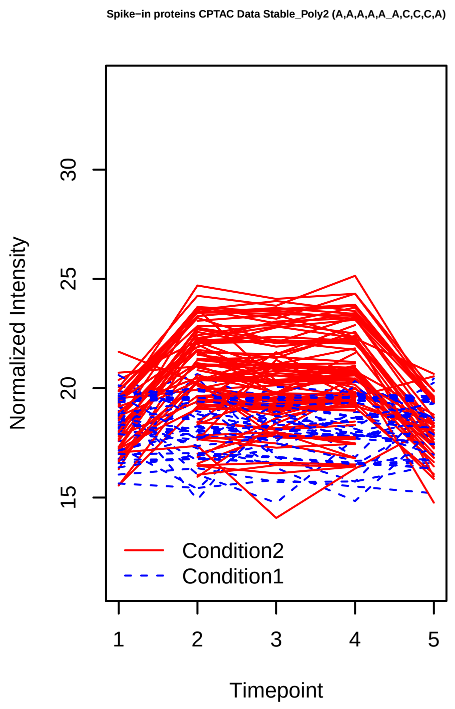

Spike−in proteins CPTAC Data Stable_Poly2 (A,A,A,A,A_A,C,C,C,A)
Normalized Intensity
15
20
25
30
1
2
3
4
5
Timepoint
Condition2
Condition1


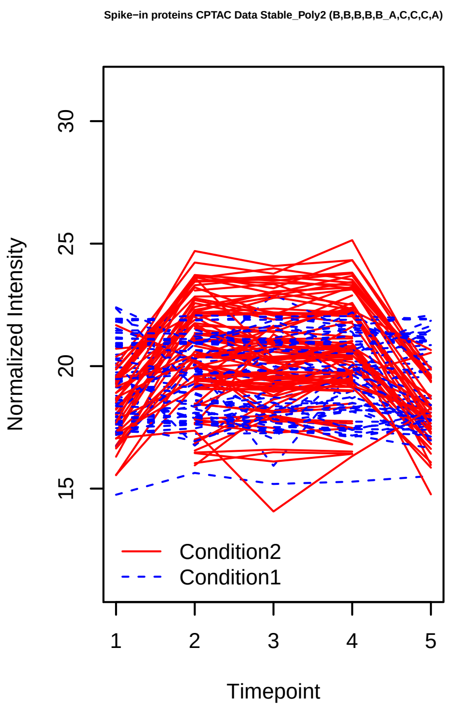

Spike−in proteins CPTAC Data Stable_Poly2 (B,B,B,B,B_A,C,C,C,A)
Normalized Intensity
15
20
25
30
1
2
3
4
5
Timepoint
Condition2
Condition1


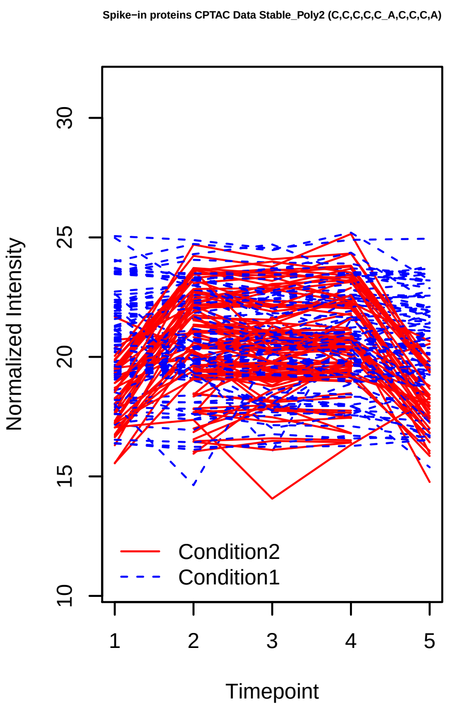

Spike−in proteins CPTAC Data Stable_Poly2 (C,C,C,C,C_A,C,C,C,A)
Normalized Intensity
10
15
20
25
30
1
2
3
4
5
Timepoint
Condition2
Condition1


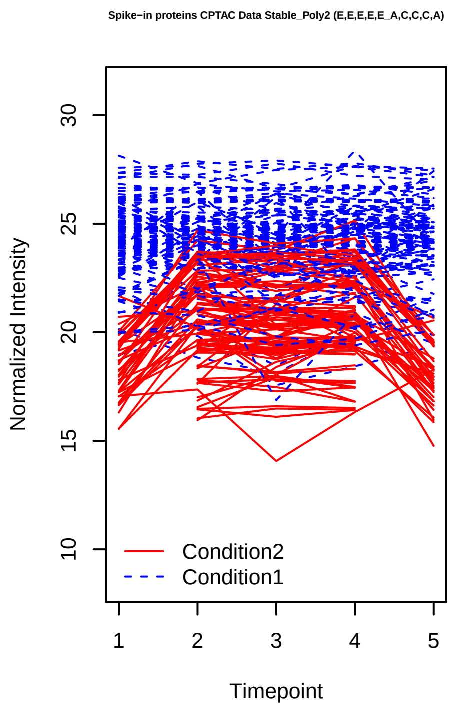

Spike−in proteins CPTAC Data Stable_Poly2 (E,E,E,E,E_A,C,C,C,A)
Normalized Intensity
10
15
20
25
30
1
2
3
4
5
Timepoint
Condition2
Condition1

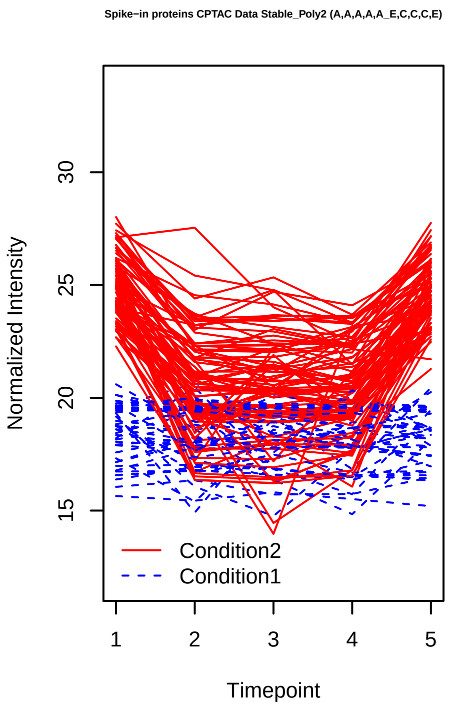
Spike−in proteins CPTAC Data Stable_Poly2 (A,A,A,A,A_E,C,C,C,E)
Normalized Intensity
Timepoint
Condition2
Condition1

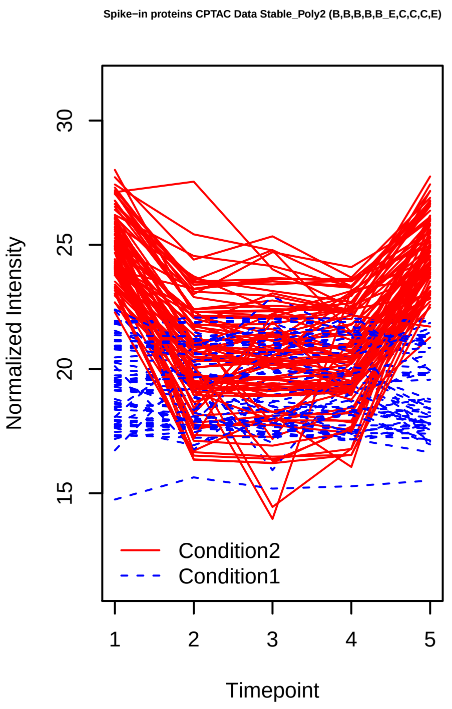
Spike−in proteins CPTAC Data Stable_Poly2 (B,B,B,B,B_E,C,C,C,E)
Normalized Intensity
Timepoint
Condition2
Condition1

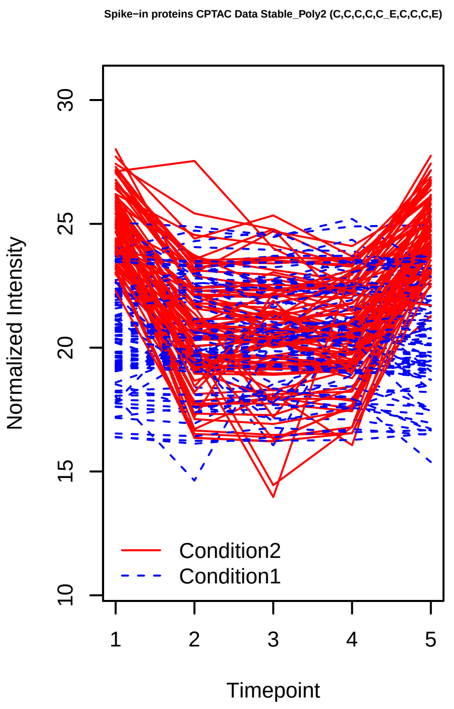
Spike−in proteins CPTAC Data Stable_Poly2 (C,C,C,C,C_E,C,C,C,E)
Normalized Intensity
Timepoint
Condition2
Condition1

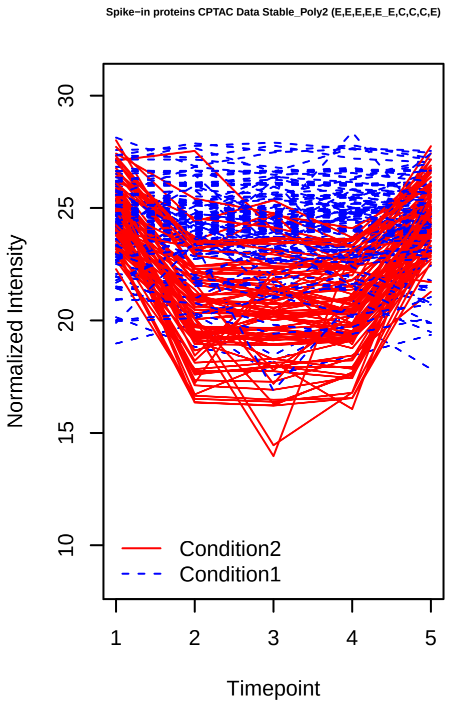
Spike−in proteins CPTAC Data Stable_Poly2 (E,E,E,E,E_E,C,C,C,E)
Normalized Intensity
Timepoint
Condition2
Condition1

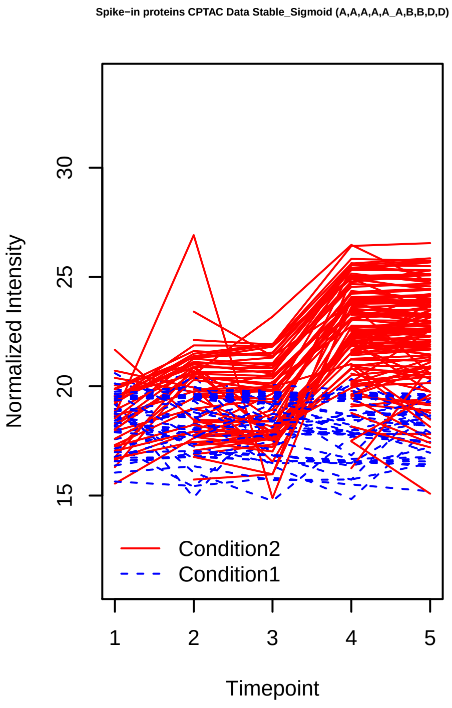
Spike−in proteins CPTAC Data Stable_Sigmoid (A,A,A,A,A_A,B,B,D,D)
Normalized Intensity
Timepoint
Condition2
Condition1

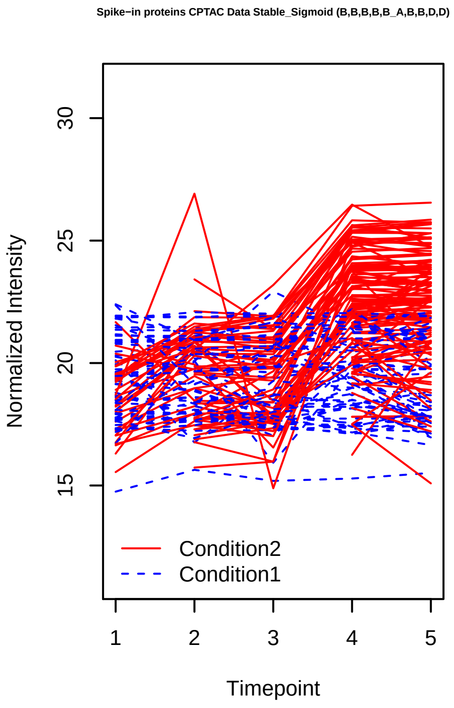
Spike−in proteins CPTAC Data Stable_Sigmoid (B,B,B,B,B_A,B,B,D,D)
Normalized Intensity
Timepoint
Condition2
Condition1

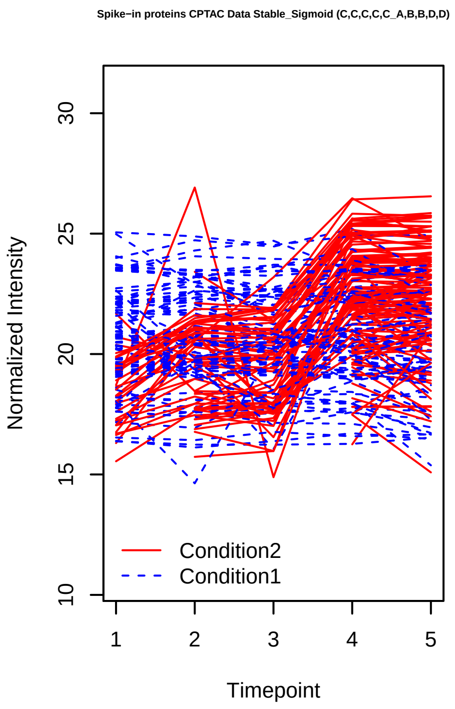
Spike−in proteins CPTAC Data Stable_Sigmoid (C,C,C,C,C_A,B,B,D,D)
Normalized Intensity
10
15
20
25
30
1
2
3
4
5
Timepoint
Condition2
Condition1

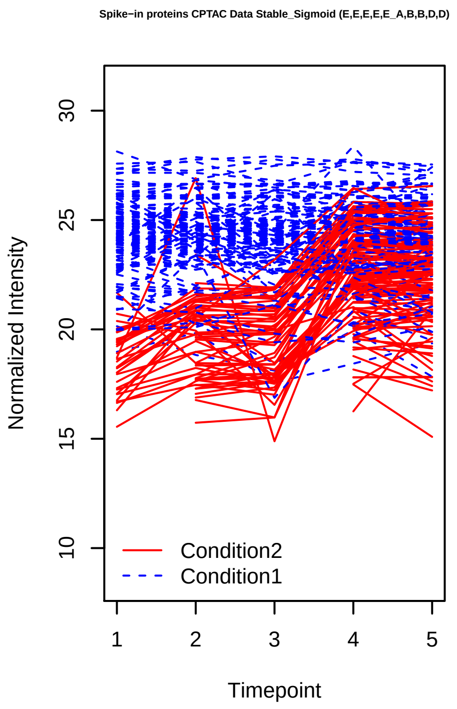
Spike−in proteins CPTAC Data Stable_Sigmoid (E,E,E,E,E_A,B,B,D,D)
Normalized Intensity
10
15
20
25
30
1
2
3
4
5
Timepoint
Condition2
Condition1

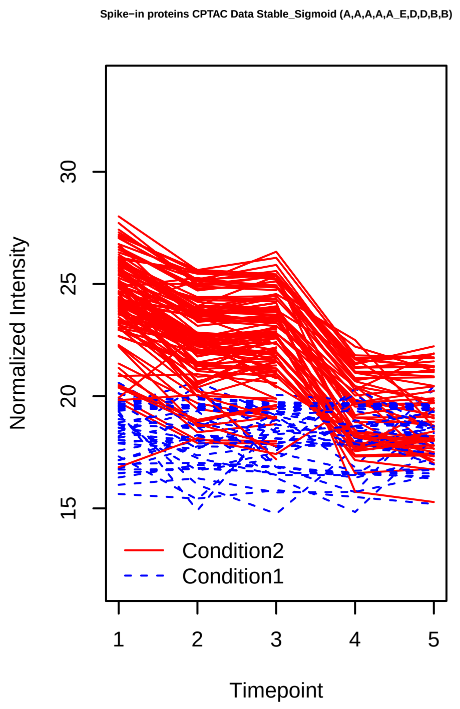
Spike−in proteins CPTAC Data Stable_Sigmoid (A,A,A,A,A_E,D,D,B,B)
Normalized Intensity
15
20
25
30
1
2
3
4
5
Timepoint
Condition2
Condition1

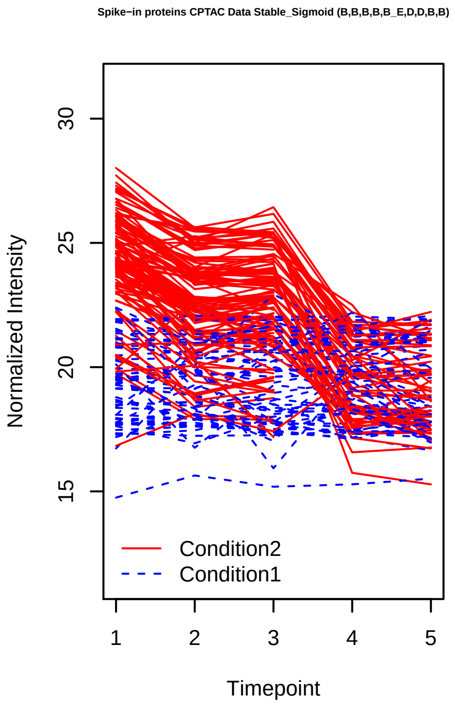
Spike−in proteins CPTAC Data Stable_Sigmoid (B,B,B,B,B_E,D,D,B,B)
Normalized Intensity
15
20
25
30
1
2
3
4
5
Timepoint
Condition2
Condition1

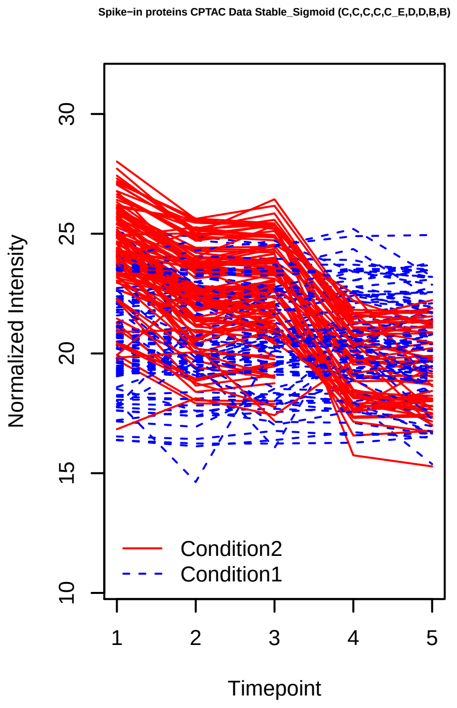
Spike−in proteins CPTAC Data Stable_Sigmoid (C,C,C,C,C_E,D,D,B,B)
Normalized Intensity
10
15
20
25
30
1
2
3
4
5
Timepoint
Condition2
Condition1

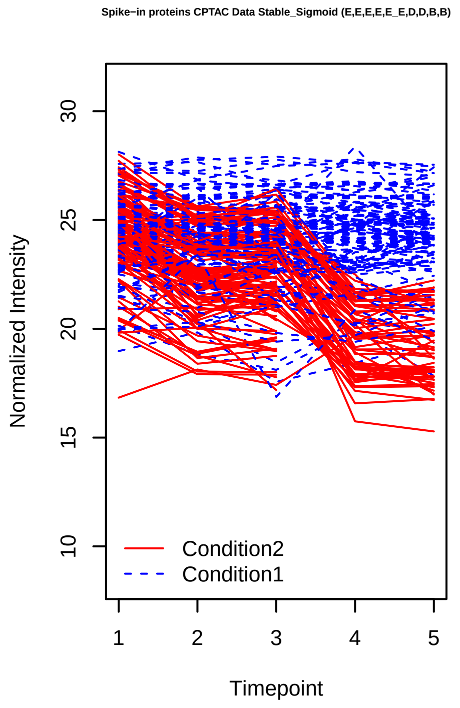
Spike−in proteins CPTAC Data Stable_Sigmoid (E,E,E,E,E_E,D,D,B,B)
Normalized Intensity
10
15
20
25
30
1
2
3
4
5
Timepoint
Condition2
Condition1

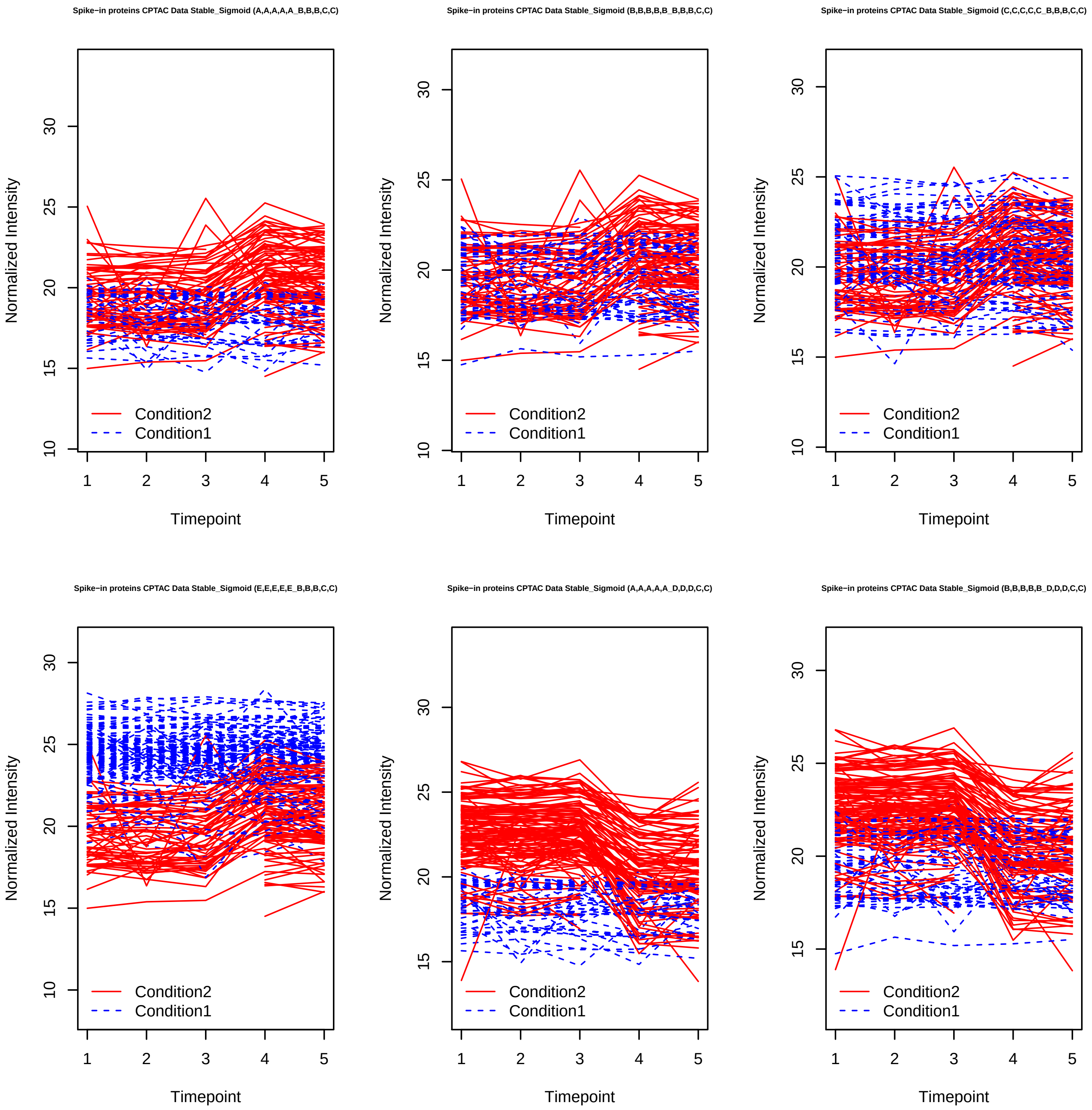

Spike−in proteins CPTAC Data Stable_Sigmoid (A,A,A,A,A_B,B,B,C,C)
Spike−in proteins CPTAC Data Stable_Sigmoid (B,B,B,B,B_B,B,B,C,C)
Spike−in proteins CPTAC Data Stable_Sigmoid (C,C,C,C,C_B,B,B,C,C)
Spike−in proteins CPTAC Data Stable_Sigmoid (E,E,E,E,E_B,B,B,C,C)
Spike−in proteins CPTAC Data Stable_Sigmoid (A,A,A,A,A_D,D,D,C,C)
Spike−in proteins CPTAC Data Stable_Sigmoid (B,B,B,B,B_D,D,D,C,C)
Normalized Intensity
Timepoint
Condition2
Condition1

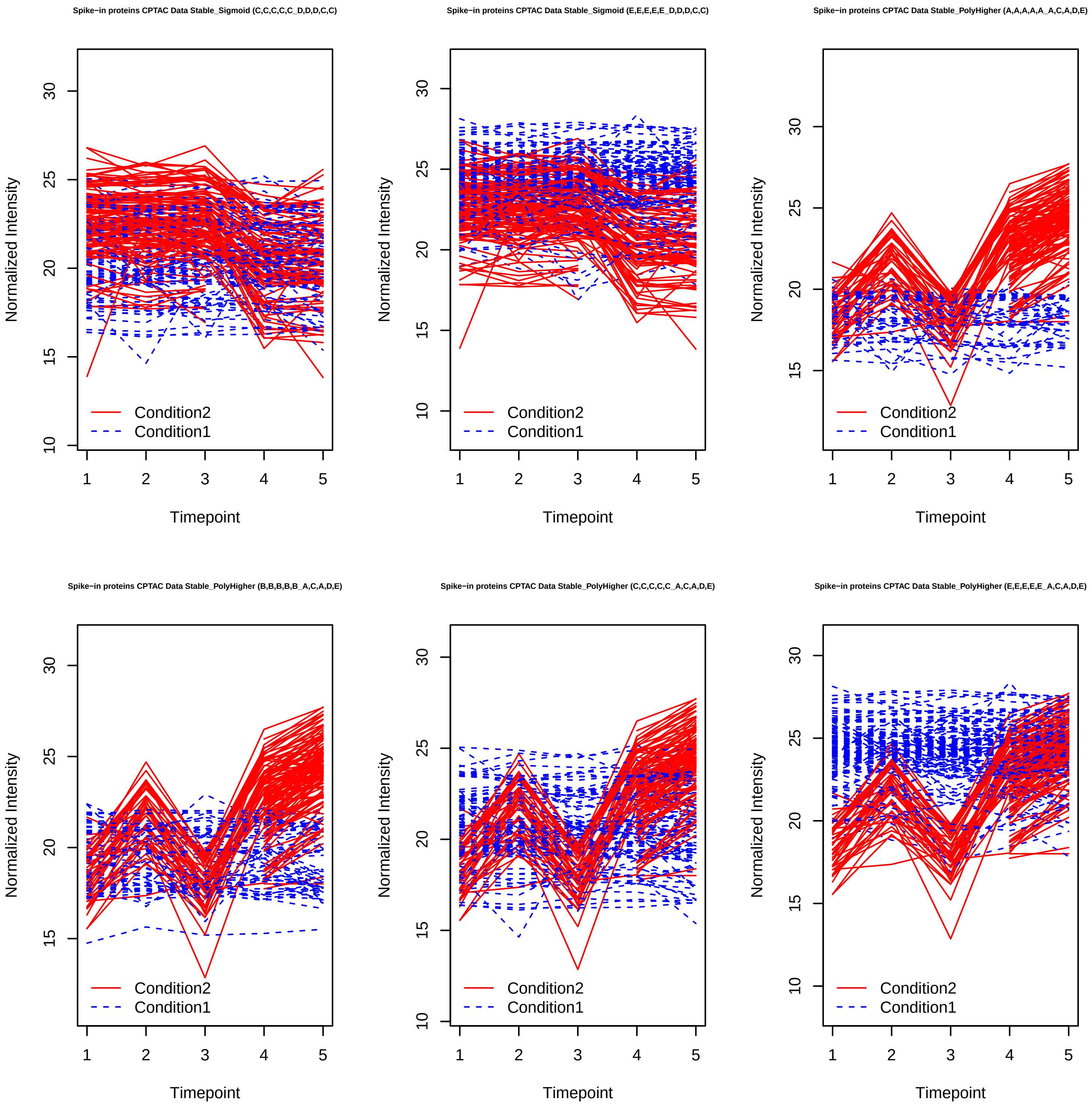
Spike−in proteins CPTAC Data Stable_Sigmoid (C,C,C,C,C_D,D,D,C,C)
Spike−in proteins CPTAC Data Stable_Sigmoid (E,E,E,E,E_D,D,D,C,C)
Spike−in proteins CPTAC Data Stable_PolyHigher (A,A,A,A,A_A,C,A,D,E)
Spike−in proteins CPTAC Data Stable_PolyHigher (B,B,B,B,B_A,C,A,D,E)
Spike−in proteins CPTAC Data Stable_PolyHigher (C,C,C,C,C_A,C,A,D,E)
Spike−in proteins CPTAC Data Stable_PolyHigher (E,E,E,E,E_A,C,A,D,E)
Normalized Intensity
Timepoint
Condition2
Condition1

Spike−in proteins CPTAC Data Stable_PolyHigher (A,A,A,A,A_E,C,E,B,A)

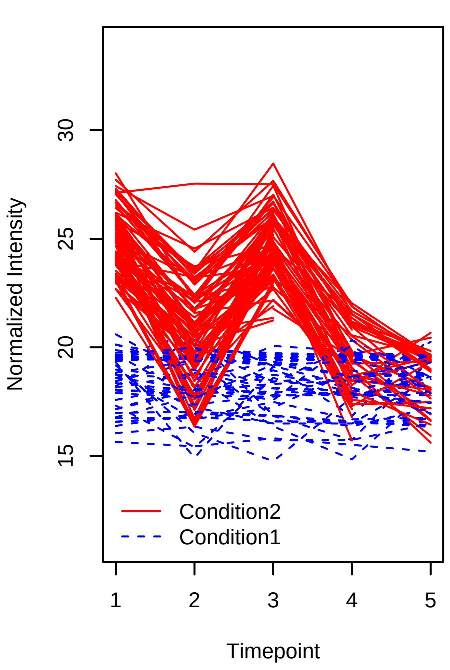


Spike−in proteins CPTAC Data Stable_PolyHigher (B,B,B,B,B_E,C,E,B,A)

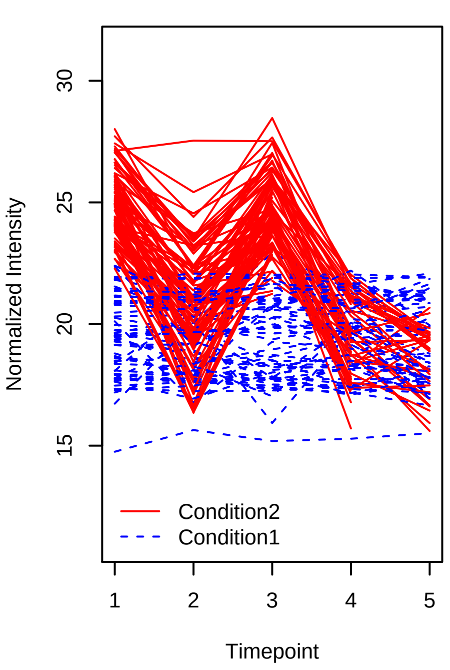


Spike−in proteins CPTAC Data Stable_PolyHigher (C,C,C,C,C_E,C,E,B,A)

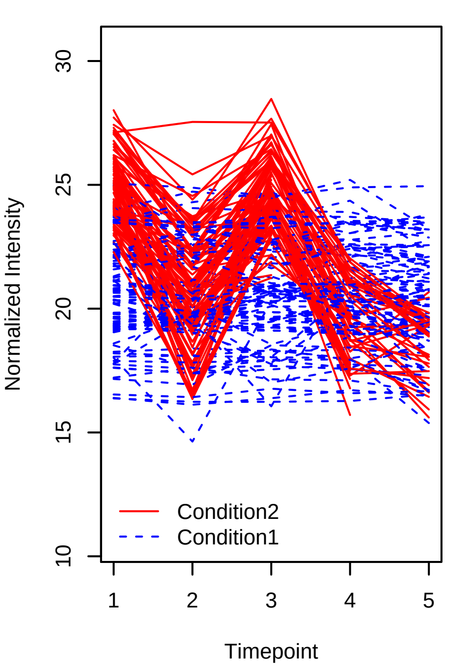


Spike−in proteins CPTAC Data Stable_PolyHigher (E,E,E,E,E_E,C,E,B,A)

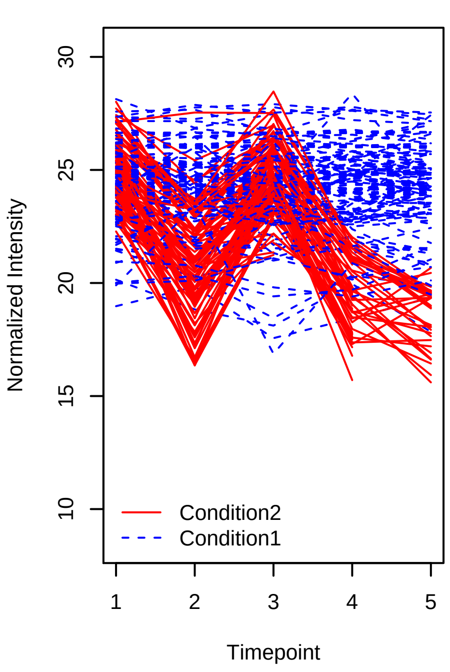


Spike−in proteins CPTAC Data Stable_PolyHigher (A,A,A,A,A_C,E,A,D,E)

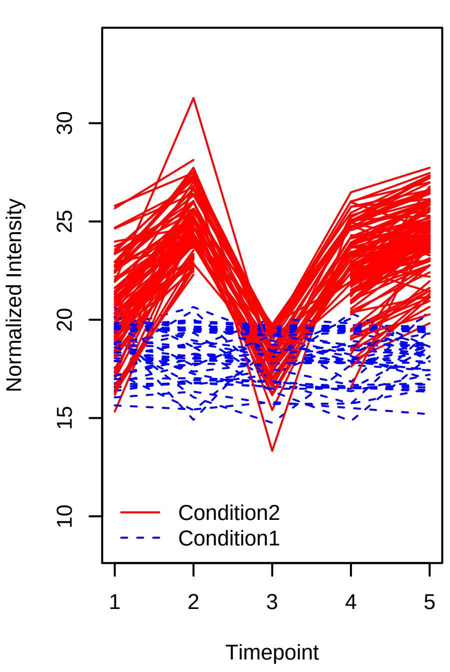


Spike−in proteins CPTAC Data Stable_PolyHigher (B,B,B,B,B_C,E,A,D,E)

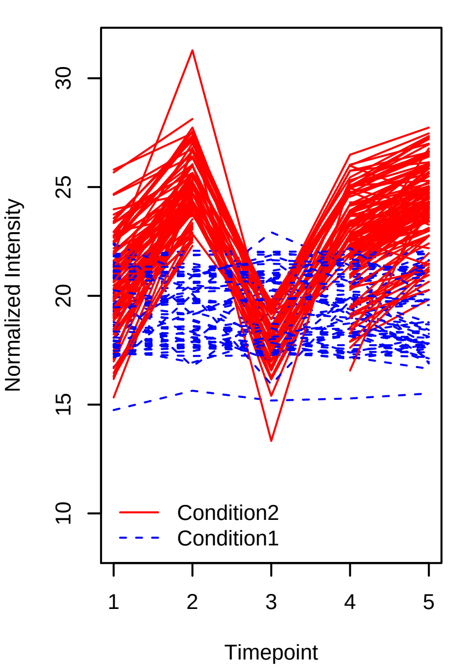

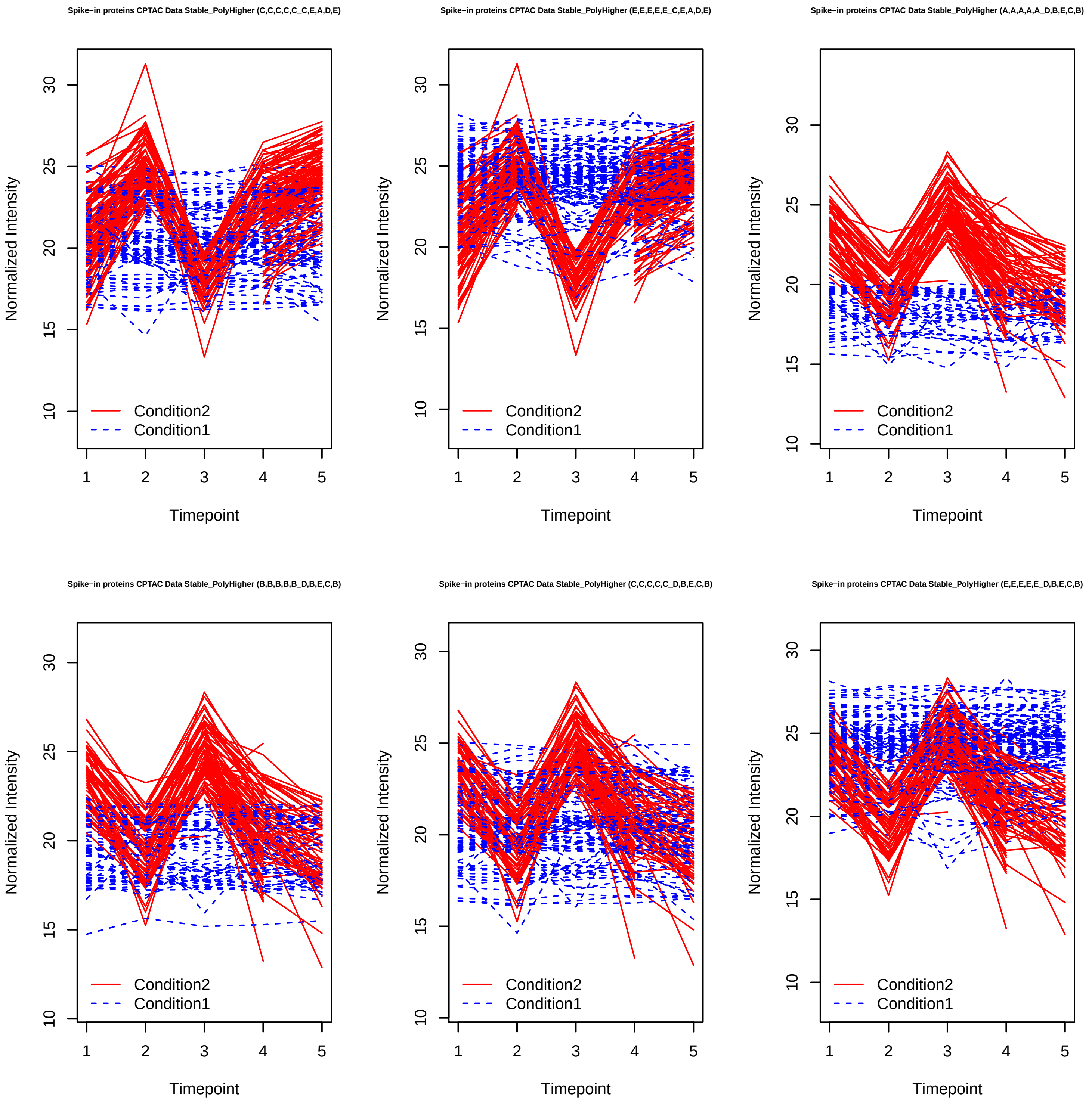

Spike−in proteins CPTAC Data Stable_PolyHigher (C,C,C,C,C_C,E,A,D,E)
Spike−in proteins CPTAC Data Stable_PolyHigher (E,E,E,E,E_C,E,A,D,E)
Spike−in proteins CPTAC Data Stable_PolyHigher (A,A,A,A,A_D,B,E,C,B)
Spike−in proteins CPTAC Data Stable_PolyHigher (B,B,B,B,B_D,B,E,C,B)
Spike−in proteins CPTAC Data Stable_PolyHigher (C,C,C,C,C_D,B,E,C,B)
Spike−in proteins CPTAC Data Stable_PolyHigher (E,E,E,E,E_D,B,E,C,B)
Normalized Intensity
Timepoint
Condition2
Condition1

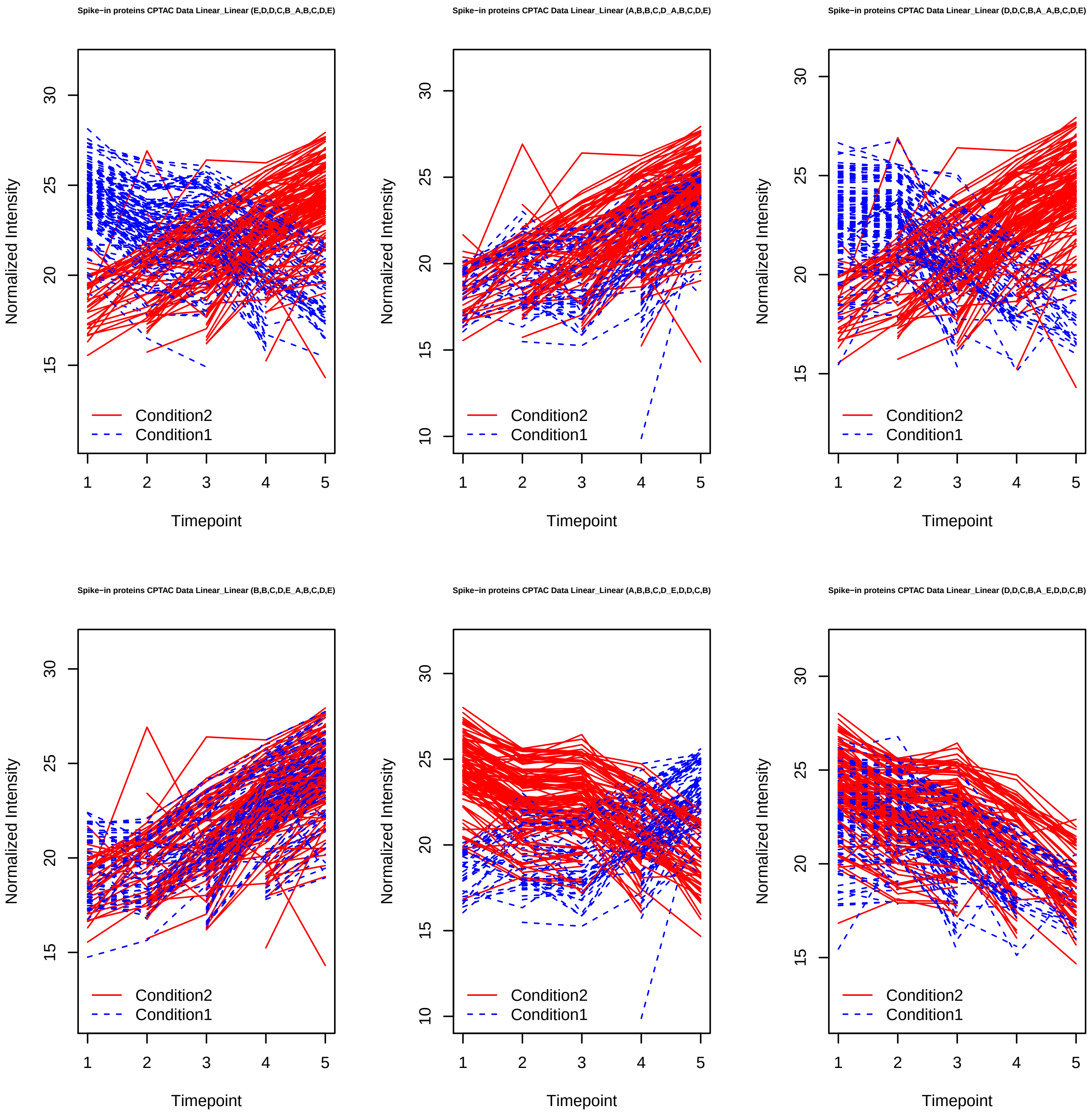
Spike−in proteins CPTAC Data Linear_Linear (E,D,D,C,B_A,B,C,D,E)
Spike−in proteins CPTAC Data Linear_Linear (A,B,B,C,D_A,B,C,D,E)
Spike−in proteins CPTAC Data Linear_Linear (D,D,C,B,A_A,B,C,D,E)
Spike−in proteins CPTAC Data Linear_Linear (B,B,C,D,E_A,B,C,D,E)
Spike−in proteins CPTAC Data Linear_Linear (A,B,B,C,D_E,D,D,C,B)
Spike−in proteins CPTAC Data Linear_Linear (D,D,C,B,A_E,D,D,C,B)
Normalized Intensity
Timepoint
Condition2
Condition1

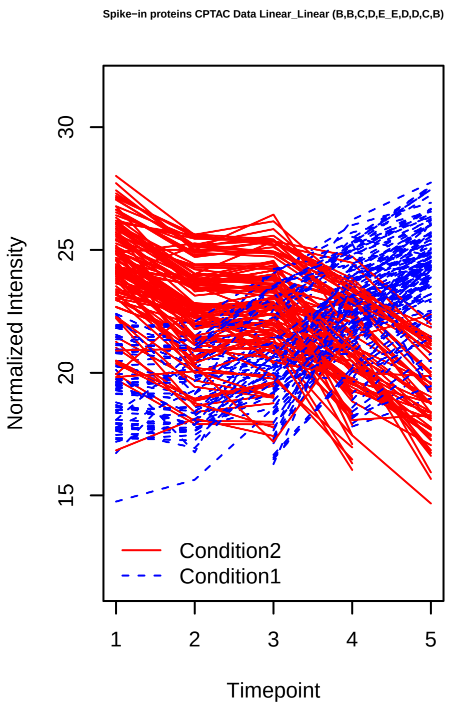
Spike−in proteins CPTAC Data Linear_Linear (B,B,C,D,E_E,D,D,C,B)
Normalized Intensity
Timepoint
Condition2
Condition1
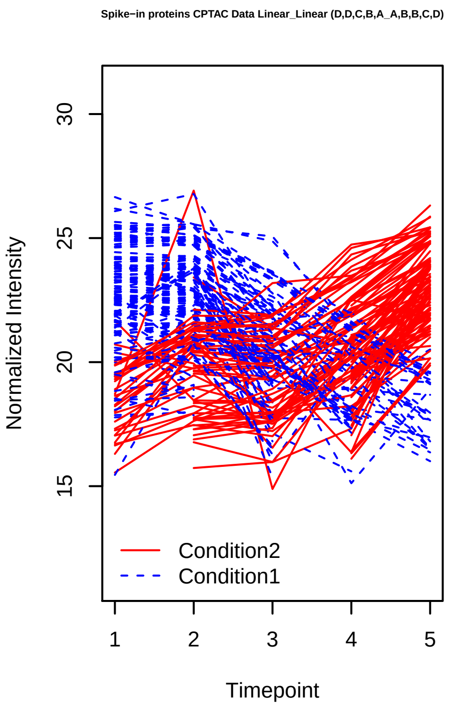
Spike−in proteins CPTAC Data Linear_Linear (D,D,C,B,A_A,B,B,C,D)
Normalized Intensity
Timepoint
Condition2
Condition1
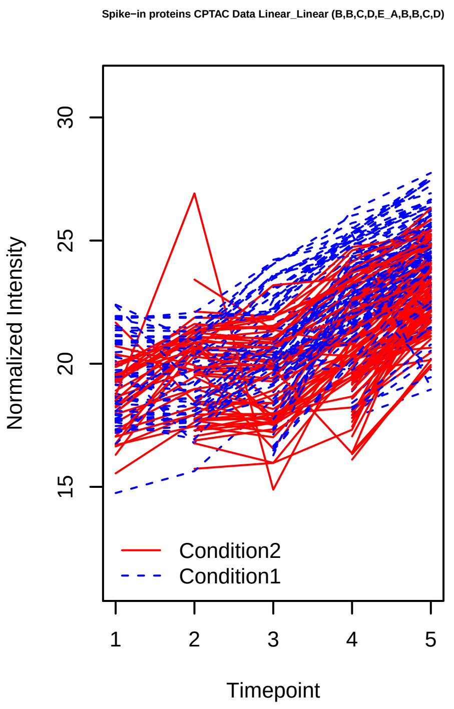
Spike−in proteins CPTAC Data Linear_Linear (B,B,C,D,E_A,B,B,C,D)
Normalized Intensity
Timepoint
Condition2
Condition1
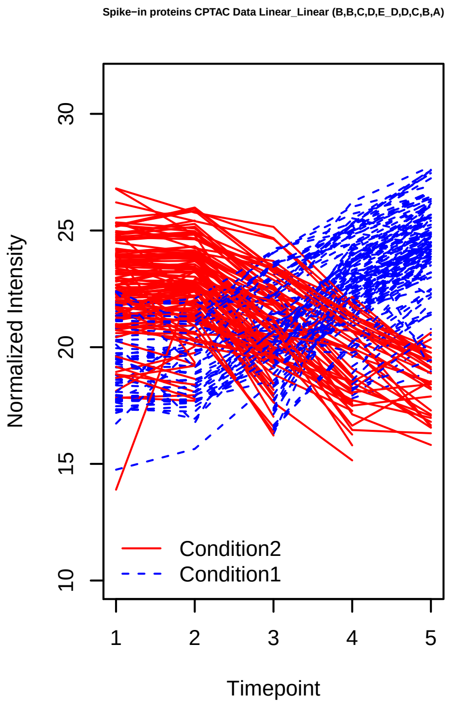
Spike−in proteins CPTAC Data Linear_Linear (B,B,C,D,E_D,D,C,B,A)
Normalized Intensity
Timepoint
Condition2
Condition1
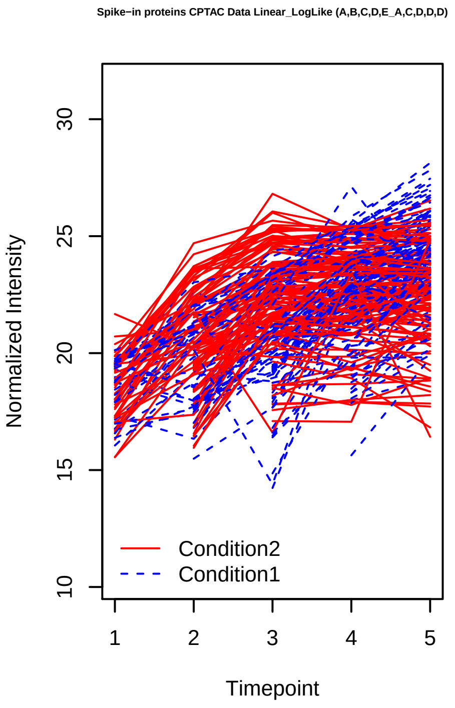
Spike−in proteins CPTAC Data Linear_LogLike (A,B,C,D,E_A,C,D,D,D)
Normalized Intensity
Timepoint
Condition2
Condition1
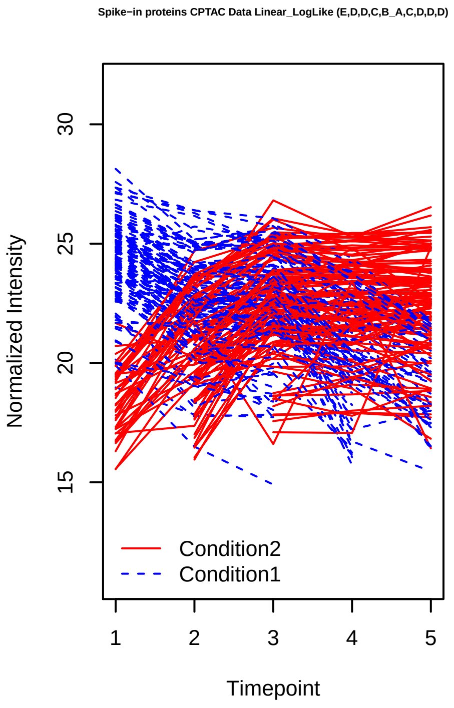
Spike−in proteins CPTAC Data Linear_LogLike (E,D,D,C,B_A,C,D,D,D)
Normalized Intensity
Timepoint
Condition2
Condition1

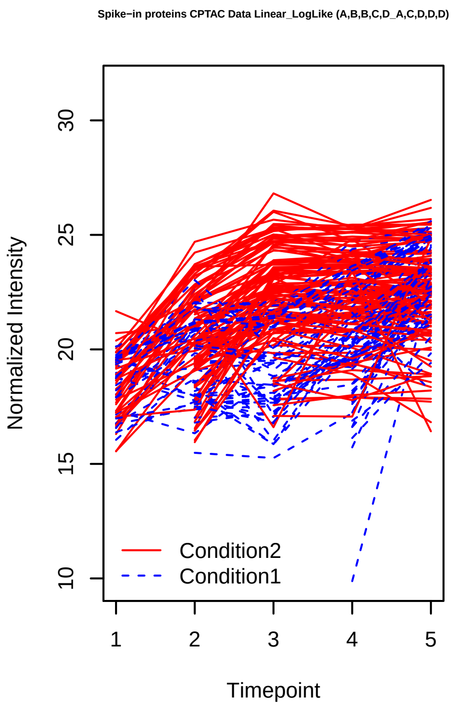
Spike−in proteins CPTAC Data Linear_LogLike (A,B,B,C,D_A,C,D,D,D)
Normalized Intensity
10
15
20
25
30
1
2
3
4
5
Timepoint
Condition2
Condition1

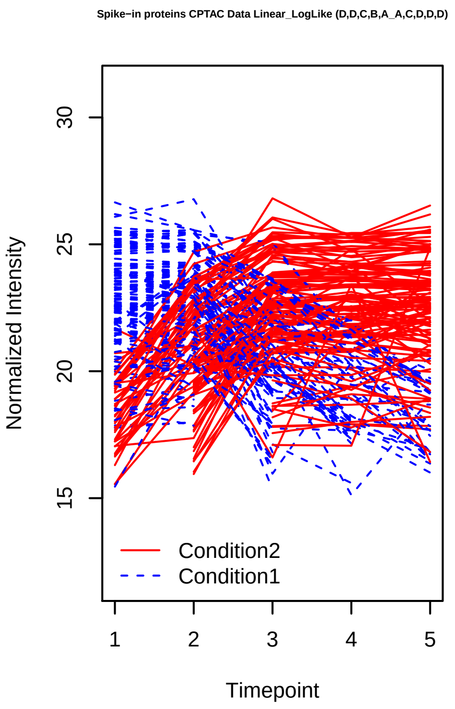
Spike−in proteins CPTAC Data Linear_LogLike (D,D,C,B,A_A,C,D,D,D)
Normalized Intensity
15
20
25
30
1
2
3
4
5
Timepoint
Condition2
Condition1

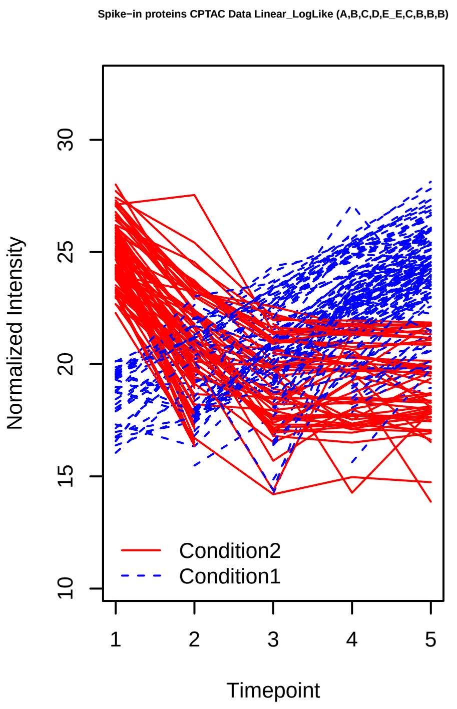
Spike−in proteins CPTAC Data Linear_LogLike (A,B,C,D,E_E,C,B,B,B)
Normalized Intensity
10
15
20
25
30
1
2
3
4
5
Timepoint
Condition2
Condition1

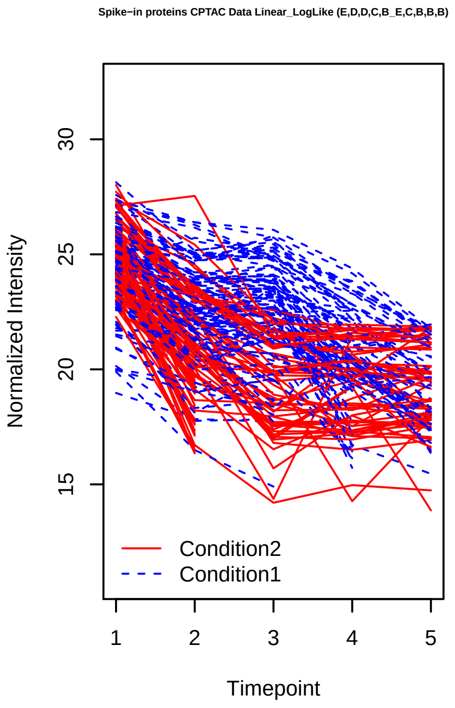
Spike−in proteins CPTAC Data Linear_LogLike (E,D,D,C,B_E,C,B,B,B)
Normalized Intensity
15
20
25
30
1
2
3
4
5
Timepoint
Condition2
Condition1

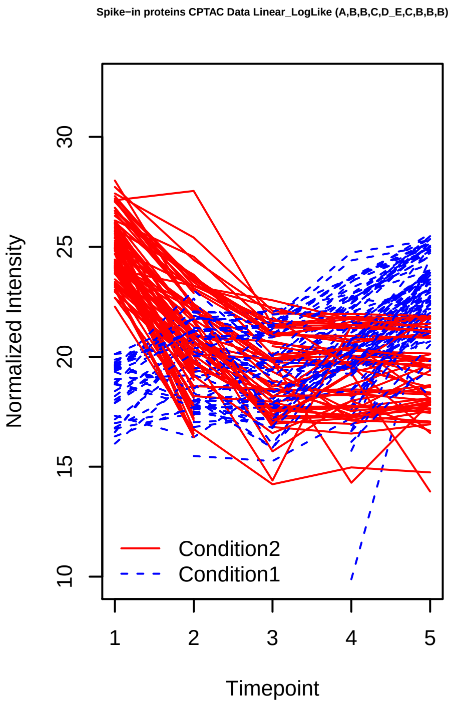
Spike−in proteins CPTAC Data Linear_LogLike (A,B,B,C,D_E,C,B,B,B)
Normalized Intensity
10
15
20
25
30
1
2
3
4
5
Timepoint
Condition2
Condition1

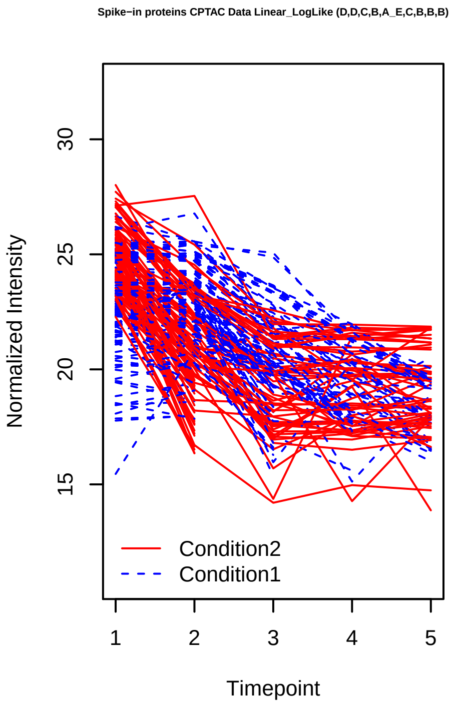
Spike−in proteins CPTAC Data Linear_LogLike (D,D,C,B,A_E,C,B,B,B)
Normalized Intensity
15
20
25
30
1
2
3
4
5
Timepoint
Condition2
Condition1

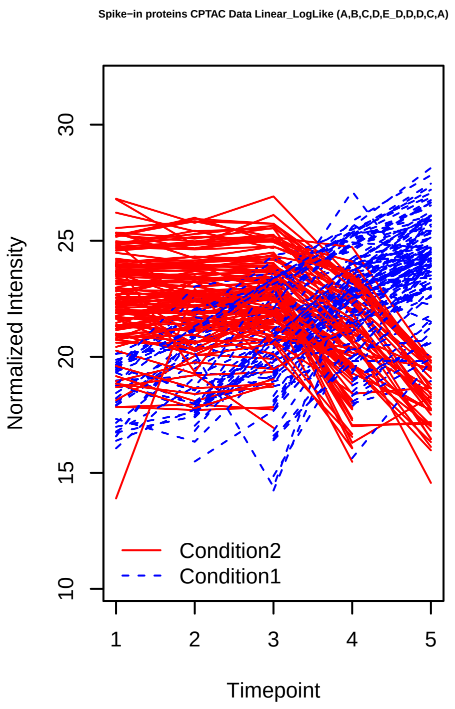
Spike−in proteins CPTAC Data Linear_LogLike (A,B,C,D,E_D,D,D,C,A)
Normalized Intensity
10
15
20
25
30
1
2
3
4
5
Timepoint
Condition2
Condition1

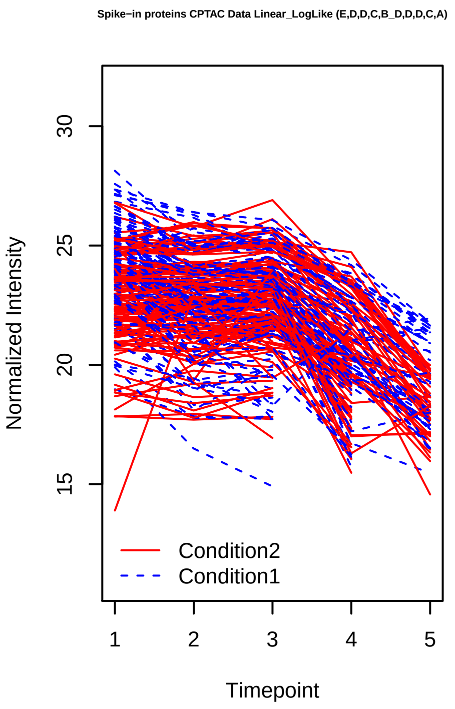
Spike−in proteins CPTAC Data Linear_LogLike (E,D,D,C,B_D,D,D,C,A)
Normalized Intensity
15
20
25
30
1
2
3
4
5
Timepoint
Condition2
Condition1

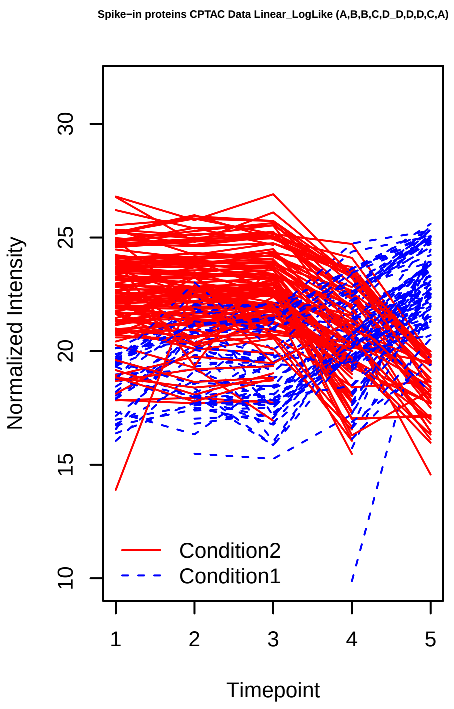
Spike−in proteins CPTAC Data Linear_LogLike (A,B,B,C,D_D,D,D,C,A)
Normalized Intensity
10
15
20
25
30
1
2
3
4
5
Timepoint
Condition2
Condition1

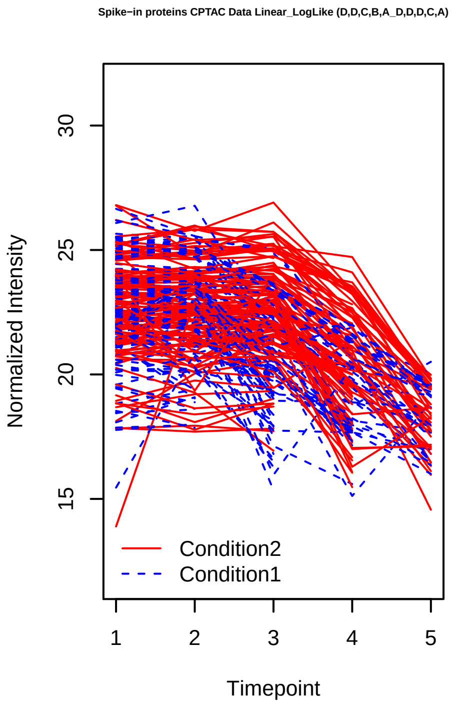
Spike−in proteins CPTAC Data Linear_LogLike (D,D,C,B,A_D,D,D,C,A)
Normalized Intensity
15
20
25
30
1
2
3
4
5
Timepoint
Condition2
Condition1

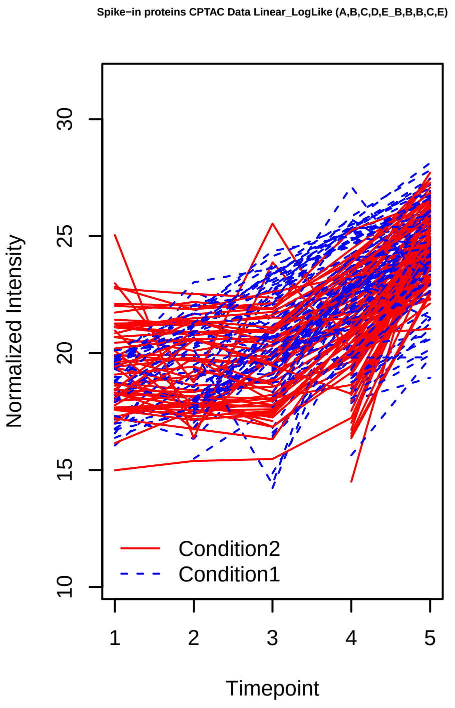
Spike−in proteins CPTAC Data Linear_LogLike (A,B,C,D,E_B,B,B,C,E)
Normalized Intensity
10
15
20
25
30
1
2
3
4
5
Timepoint
Condition2
Condition1

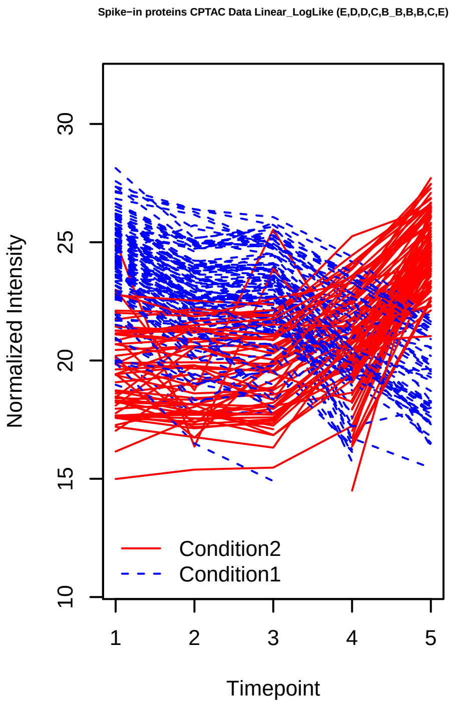
Spike−in proteins CPTAC Data Linear_LogLike (E,D,D,C,B_B,B,B,C,E)
Normalized Intensity
10
15
20
25
30
1
2
3
4
5
Timepoint
Condition2
Condition1

Spike−in proteins CPTAC Data Linear_LogLike (A,B,B,C,D_B,B,B,C,E)

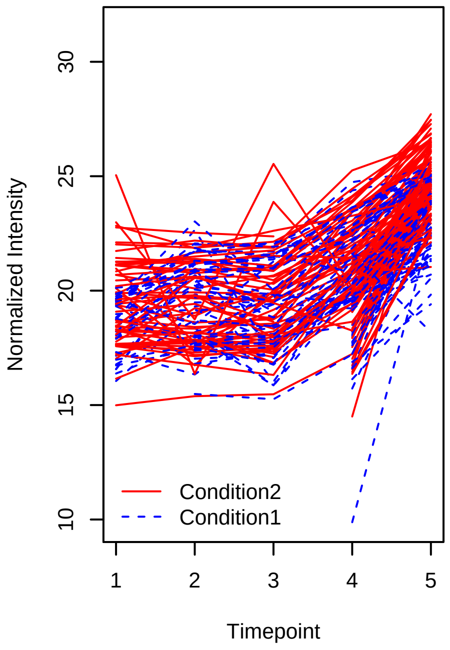


Spike−in proteins CPTAC Data Linear_LogLike (D,D,C,B,A_B,B,B,C,E)

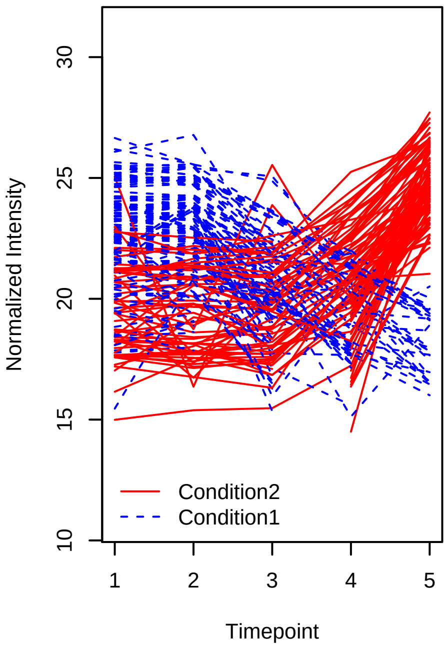


Spike−in proteins CPTAC Data Linear_Poly2 (A,B,C,D,E_A,B,C,B,A)

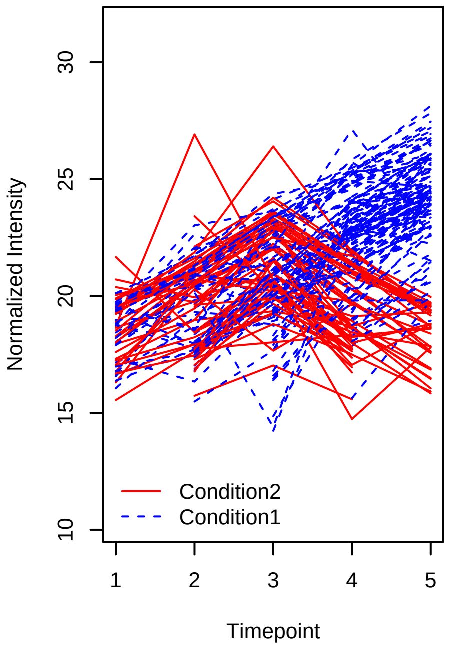


Spike−in proteins CPTAC Data Linear_Poly2 (E,D,D,C,B_A,B,C,B,A)

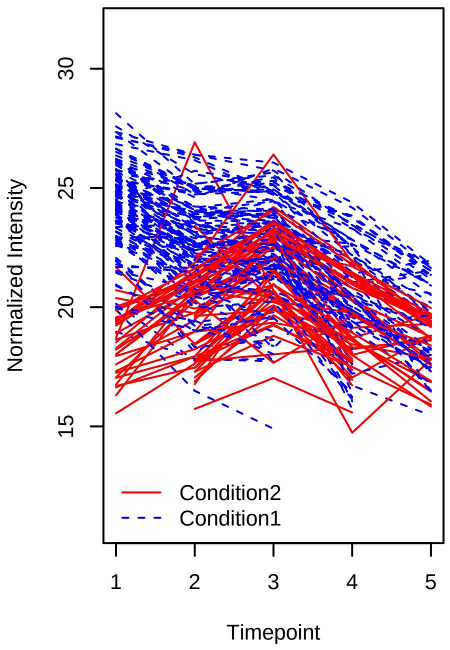


Spike−in proteins CPTAC Data Linear_Poly2 (A,B,B,C,D_A,B,C,B,A)

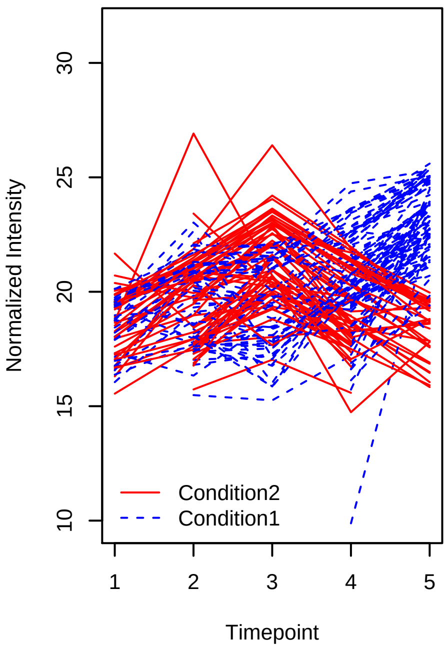


Spike−in proteins CPTAC Data Linear_Poly2 (D,D,C,B,A_A,B,C,B,A)

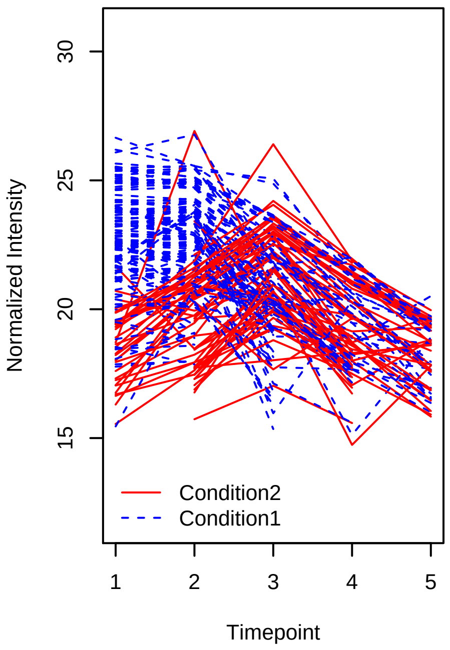

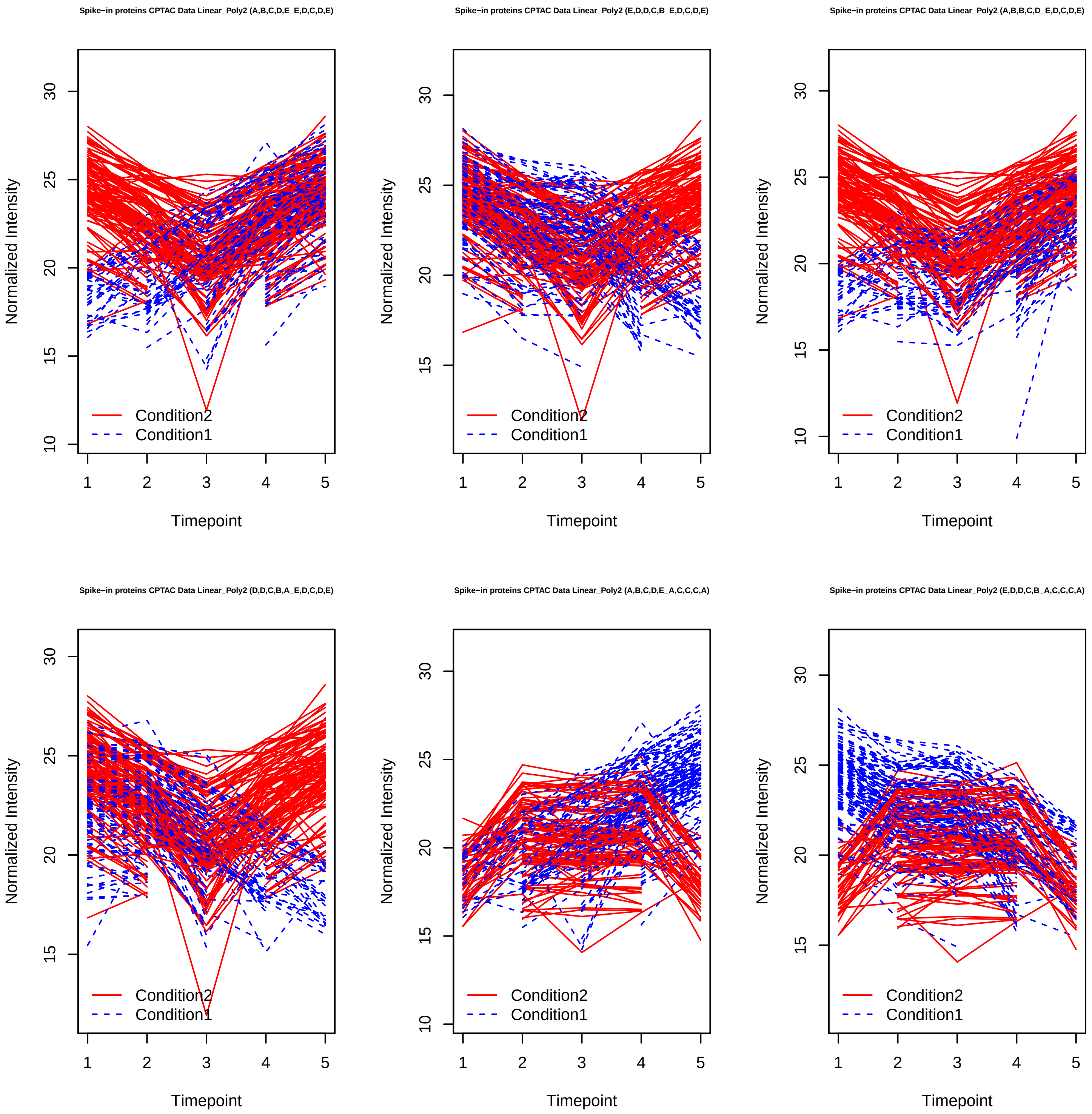
Spike−in proteins CPTAC Data Linear_Poly2 (A,B,C,D,E_E,D,C,D,E)
Spike−in proteins CPTAC Data Linear_Poly2 (E,D,D,C,B_E,D,C,D,E)
Spike−in proteins CPTAC Data Linear_Poly2 (A,B,B,C,D_E,D,C,D,E)
Spike−in proteins CPTAC Data Linear_Poly2 (D,D,C,B,A_E,D,C,D,E)
Spike−in proteins CPTAC Data Linear_Poly2 (A,B,C,D,E_A,C,C,C,A)
Spike−in proteins CPTAC Data Linear_Poly2 (E,D,D,C,B_A,C,C,C,A)
Normalized Intensity
Timepoint
Condition2
Condition1

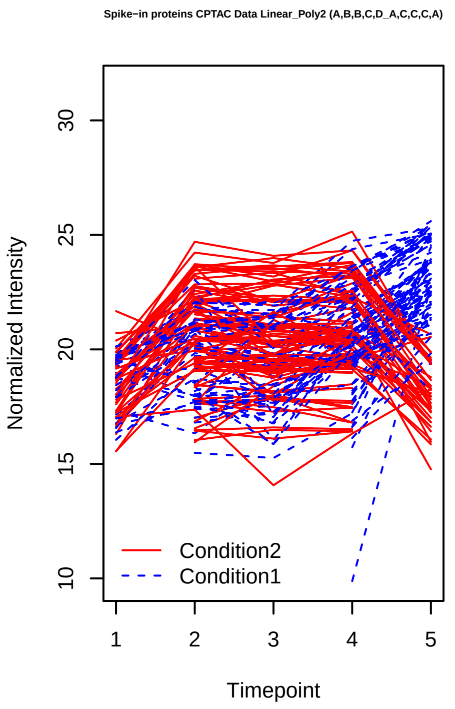
Spike−in proteins CPTAC Data Linear_Poly2 (A,B,B,C,D_A,C,C,C,A)
Normalized Intensity
30
25
20
15
10
Condition2
Condition1
1
2
3
4
5
Timepoint

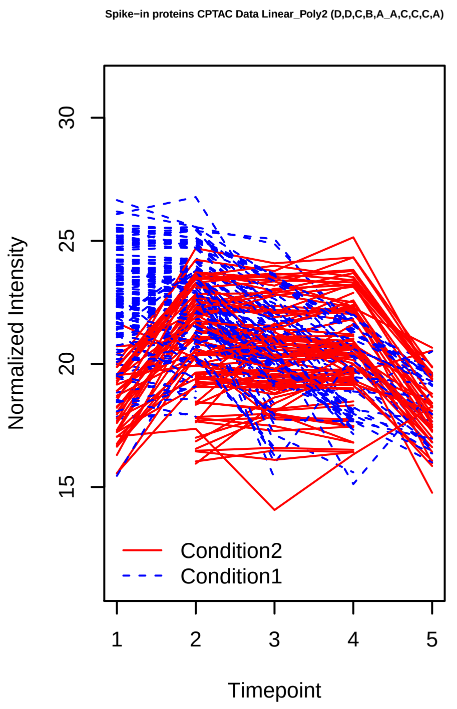
Spike−in proteins CPTAC Data Linear_Poly2 (D,D,C,B,A_A,C,C,C,A)
Normalized Intensity
30
25
20
15
Condition2
Condition1
1
2
3
4
5
Timepoint

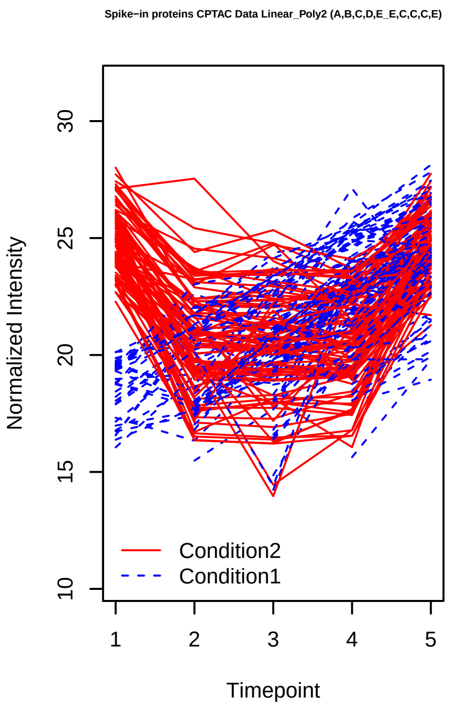
Spike−in proteins CPTAC Data Linear_Poly2 (A,B,C,D,E_E,C,C,C,E)
Normalized Intensity
30
25
20
15
10
Condition2
Condition1
1
2
3
4
5
Timepoint

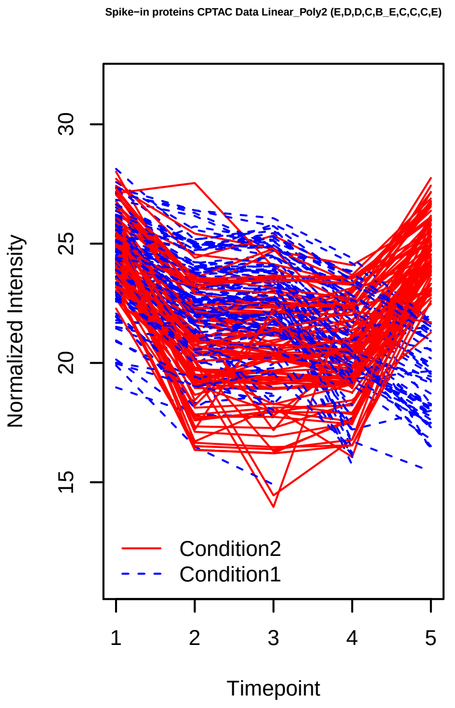
Spike−in proteins CPTAC Data Linear_Poly2 (E,D,D,C,B_E,C,C,C,E)
Normalized Intensity
30
25
20
15
Condition2
Condition1
1
2
3
4
5
Timepoint

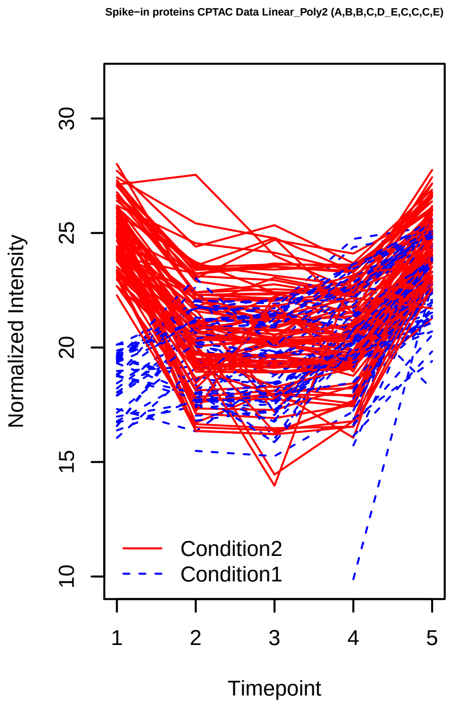
Spike−in proteins CPTAC Data Linear_Poly2 (A,B,B,C,D_E,C,C,C,E)
Normalized Intensity
30
25
20
15
10
Condition2
Condition1
1
2
3
4
5
Timepoint

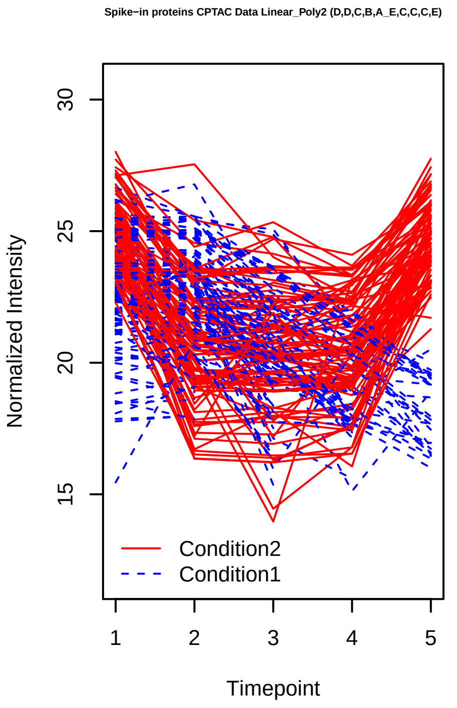
Spike−in proteins CPTAC Data Linear_Poly2 (D,D,C,B,A_E,C,C,C,E)
Normalized Intensity
30
25
20
15
Condition2
Condition1
1
2
3
4
5
Timepoint

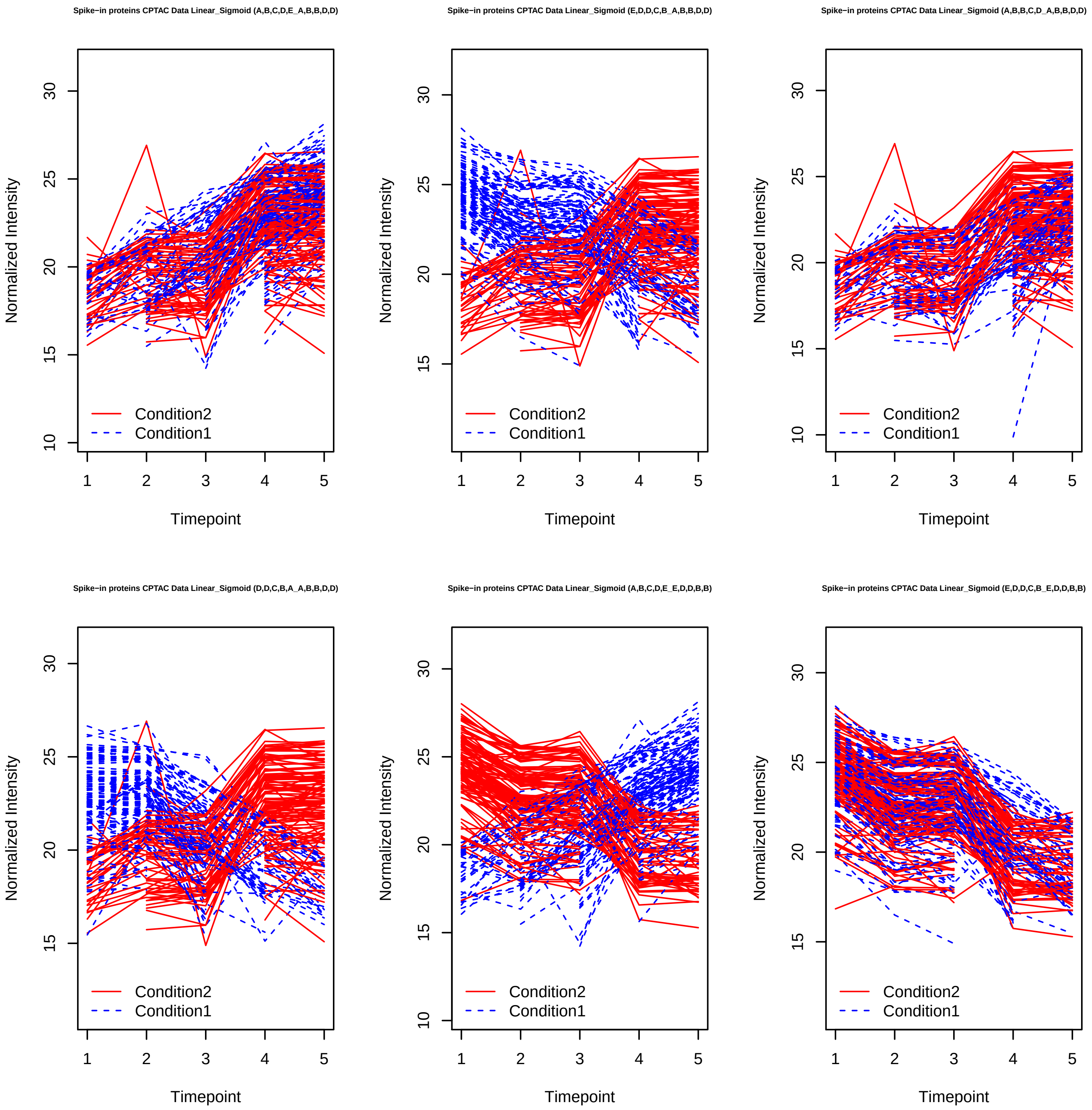

Spike−in proteins CPTAC Data Linear_Sigmoid (A,B,C,D,E_A,B,B,D,D)
Spike−in proteins CPTAC Data Linear_Sigmoid (E,D,D,C,B_A,B,B,D,D)
Spike−in proteins CPTAC Data Linear_Sigmoid (A,B,B,C,D_A,B,B,D,D)
Spike−in proteins CPTAC Data Linear_Sigmoid (D,D,C,B,A_A,B,B,D,D)
Spike−in proteins CPTAC Data Linear_Sigmoid (A,B,C,D,E_E,D,D,B,B)
Spike−in proteins CPTAC Data Linear_Sigmoid (E,D,D,C,B_E,D,D,B,B)
Normalized Intensity
Timepoint
Condition2
Condition1

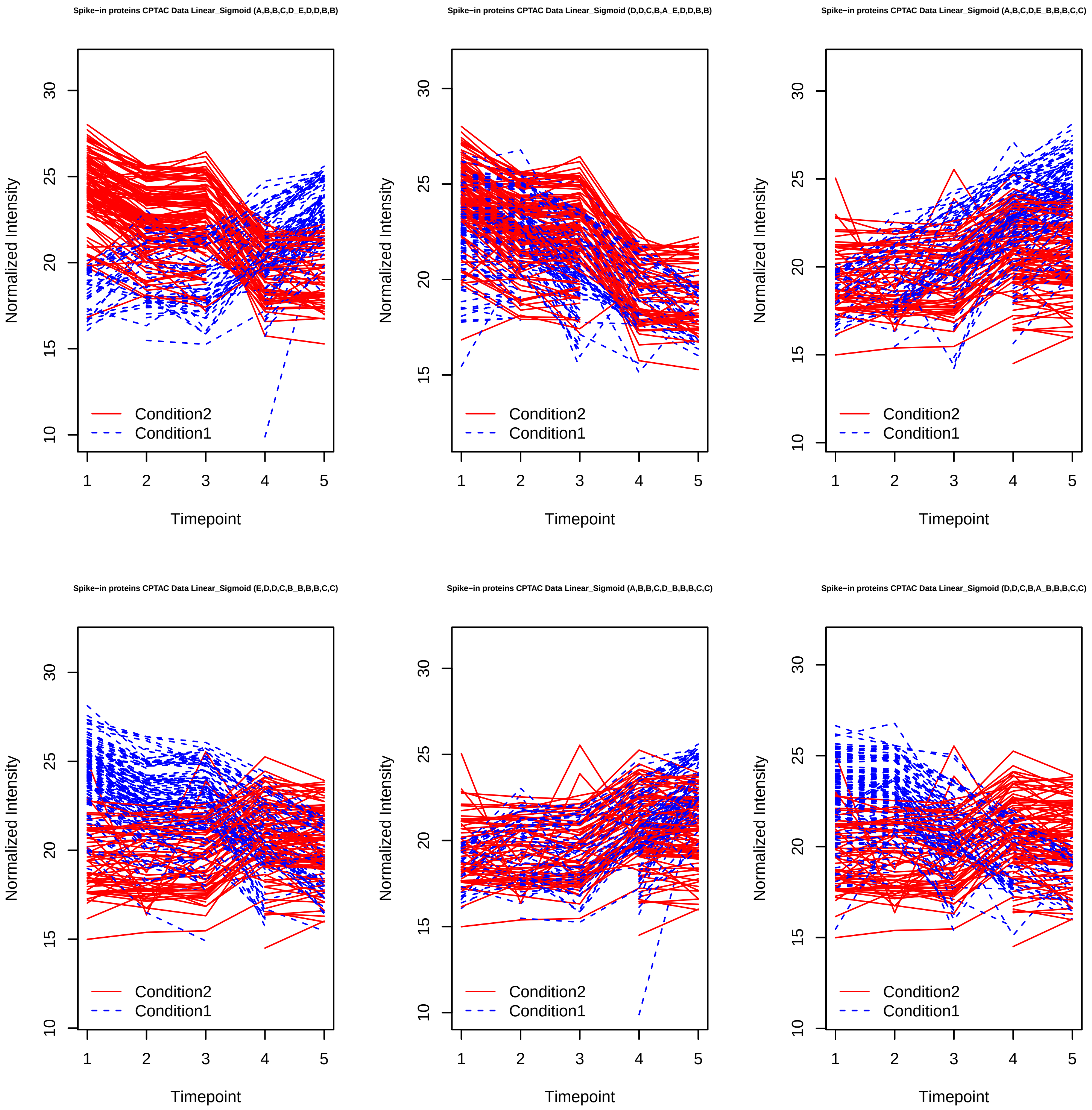
Spike−in proteins CPTAC Data Linear_Sigmoid (A,B,B,C,D_E,D,D,B,B)
Spike−in proteins CPTAC Data Linear_Sigmoid (D,D,C,B,A_E,D,D,B,B)
Spike−in proteins CPTAC Data Linear_Sigmoid (A,B,C,D,E_B,B,B,C,C)
Spike−in proteins CPTAC Data Linear_Sigmoid (E,D,D,C,B_B,B,B,C,C)
Spike−in proteins CPTAC Data Linear_Sigmoid (A,B,B,C,D_B,B,B,C,C)
Spike−in proteins CPTAC Data Linear_Sigmoid (D,D,C,B,A_B,B,B,C,C)
Normalized Intensity
Timepoint
Condition2
Condition1

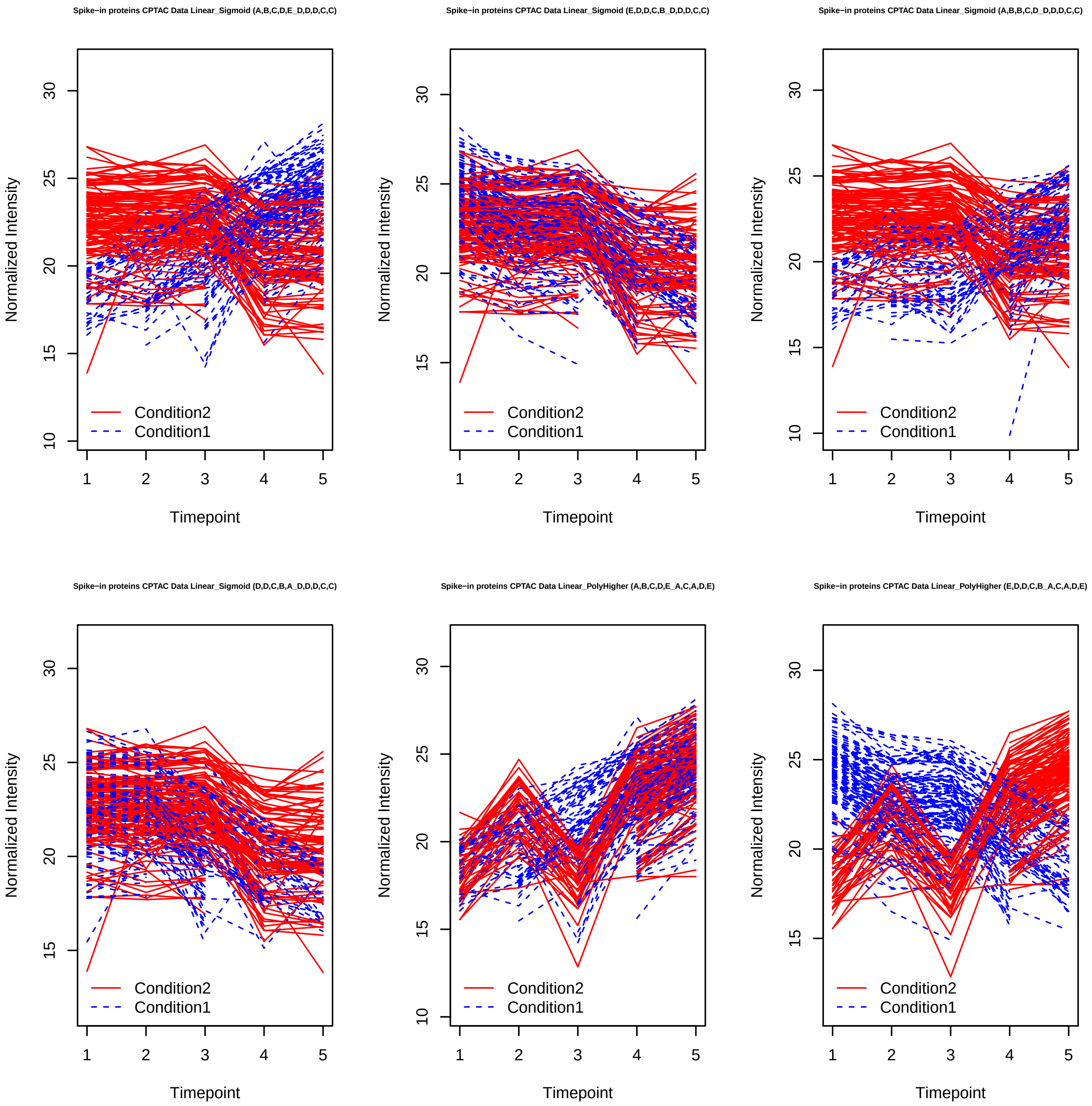
Spike−in proteins CPTAC Data Linear_Sigmoid (A,B,C,D,E_D,D,D,C,C)
Spike−in proteins CPTAC Data Linear_Sigmoid (E,D,D,C,B_D,D,D,C,C)
Spike−in proteins CPTAC Data Linear_Sigmoid (A,B,B,C,D_D,D,D,C,C)
Spike−in proteins CPTAC Data Linear_Sigmoid (D,D,C,B,A_D,D,D,C,C)
Spike−in proteins CPTAC Data Linear_PolyHigher (A,B,C,D,E_A,C,A,D,E)
Spike−in proteins CPTAC Data Linear_PolyHigher (E,D,D,C,B_A,C,A,D,E)
Normalized Intensity
Timepoint
Condition2
Condition1

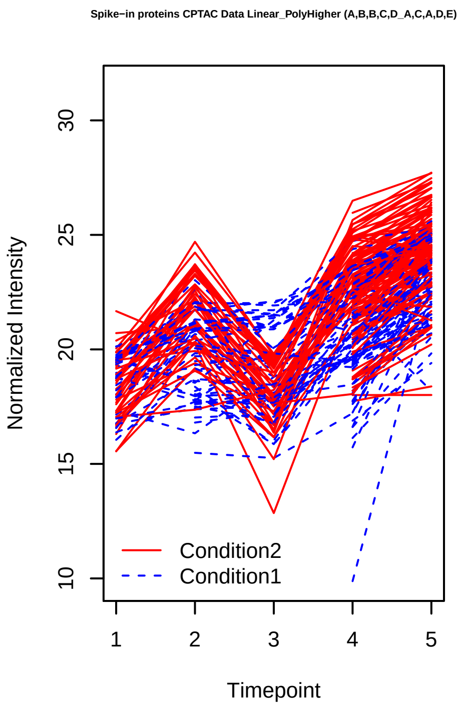
Spike−in proteins CPTAC Data Linear_PolyHigher (A,B,B,C,D_A,C,A,D,E)
Normalized Intensity
Timepoint
Condition2
Condition1

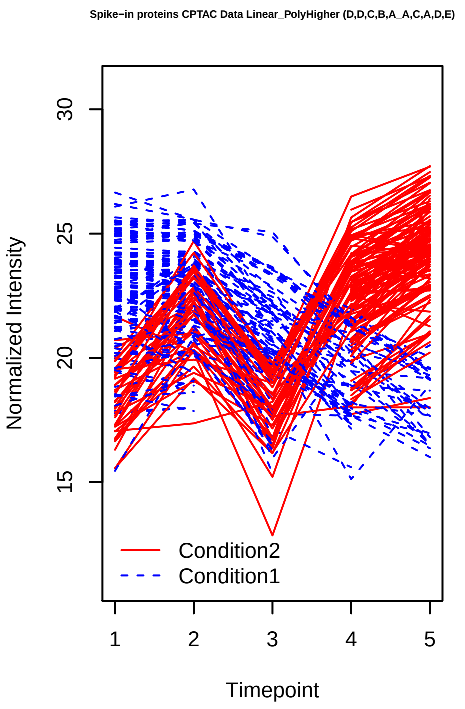
Spike−in proteins CPTAC Data Linear_PolyHigher (D,D,C,B,A_A,C,A,D,E)
Normalized Intensity
Timepoint
Condition2
Condition1

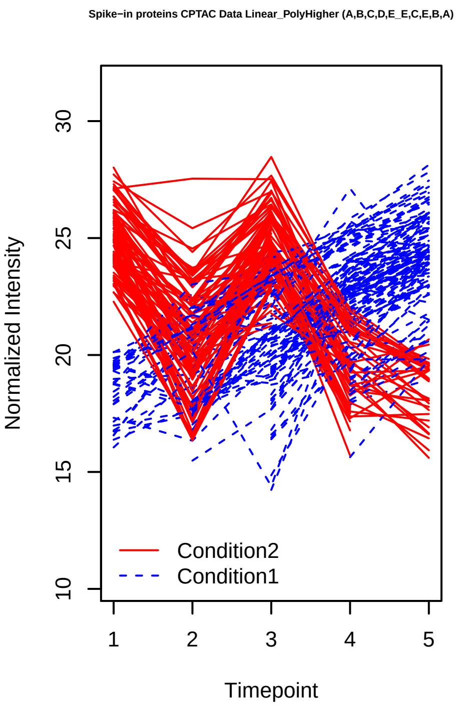
Spike−in proteins CPTAC Data Linear_PolyHigher (A,B,C,D,E_E,C,E,B,A)
Normalized Intensity
Timepoint
Condition2
Condition1

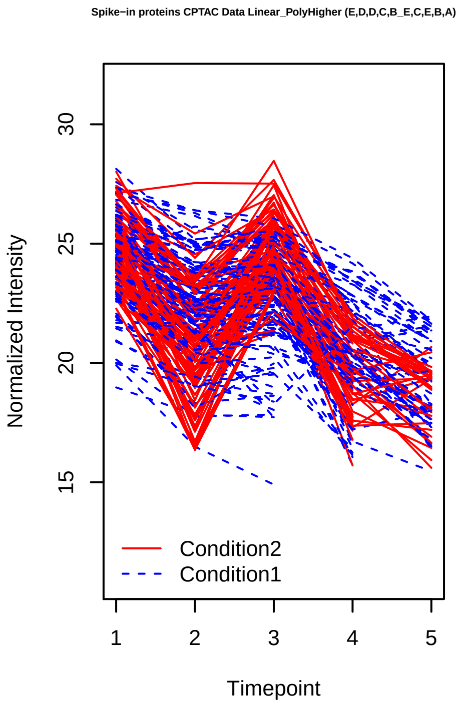
Spike−in proteins CPTAC Data Linear_PolyHigher (E,D,D,C,B_E,C,E,B,A)
Normalized Intensity
Timepoint
Condition2
Condition1

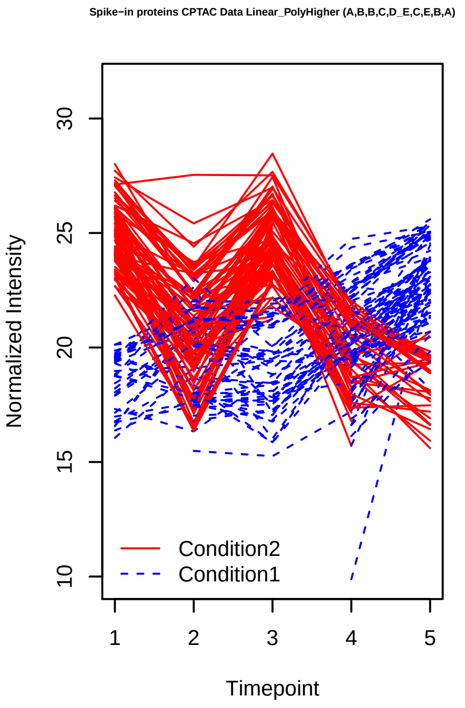
Spike−in proteins CPTAC Data Linear_PolyHigher (A,B,B,C,D_E,C,E,B,A)
Normalized Intensity
Timepoint
Condition2
Condition1

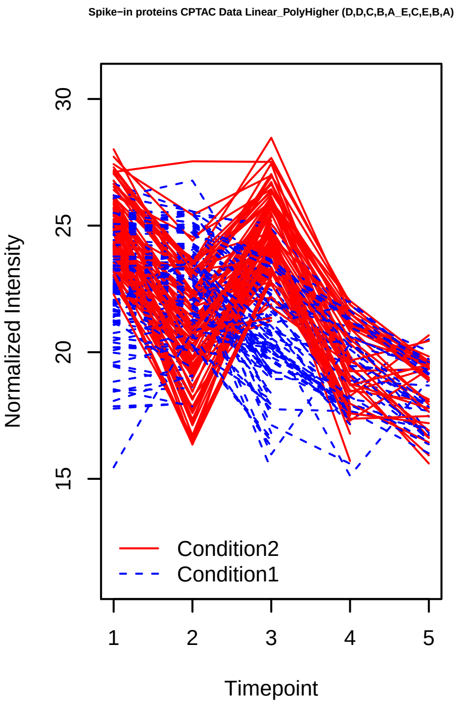
Spike−in proteins CPTAC Data Linear_PolyHigher (D,D,C,B,A_E,C,E,B,A)
Normalized Intensity
Timepoint
Condition2
Condition1

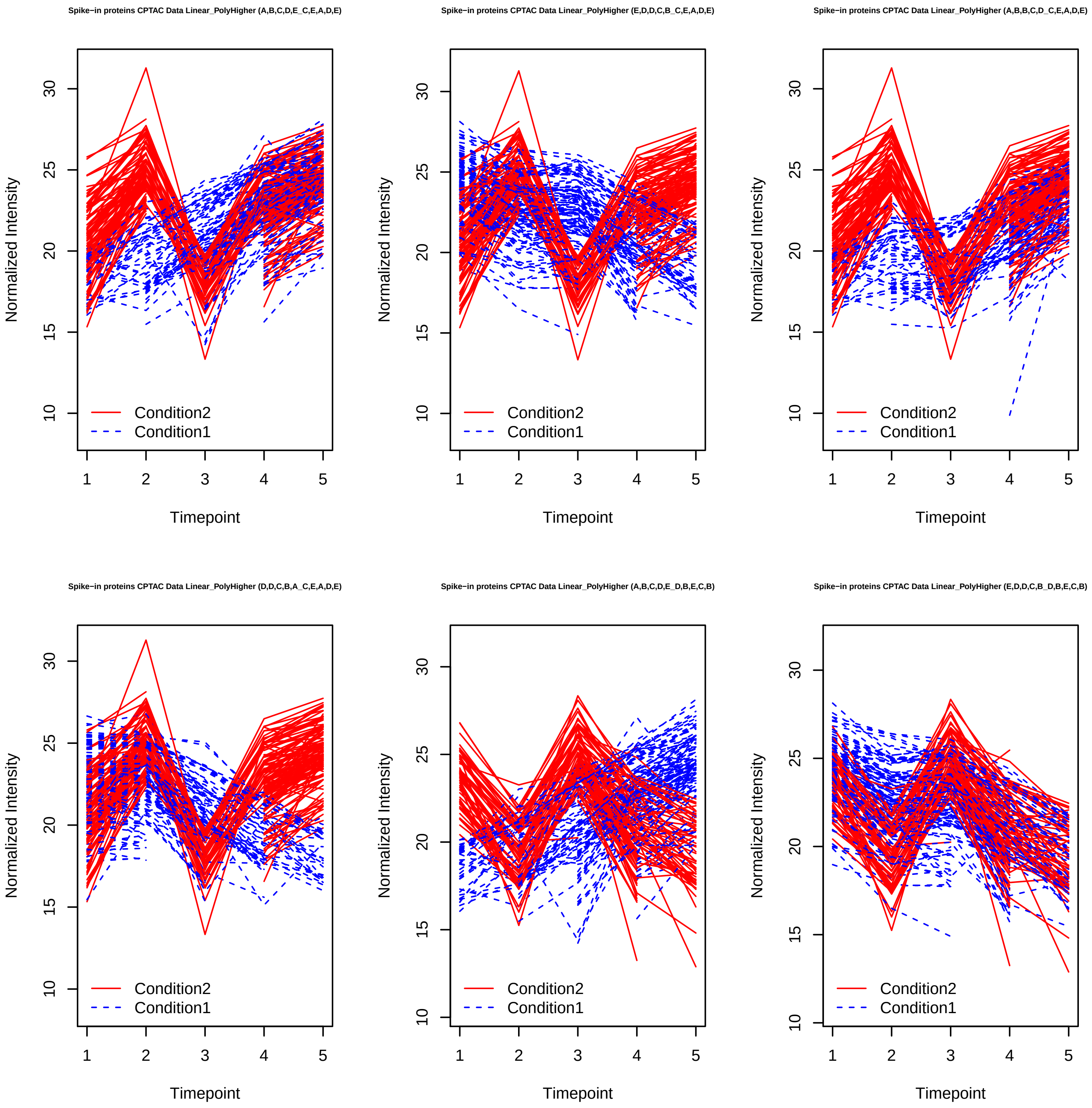
Spike−in proteins CPTAC Data Linear_PolyHigher (A,B,C,D,E_C,E,A,D,E)
Spike−in proteins CPTAC Data Linear_PolyHigher (E,D,D,C,B_C,E,A,D,E)
Spike−in proteins CPTAC Data Linear_PolyHigher (A,B,B,C,D_C,E,A,D,E)
Spike−in proteins CPTAC Data Linear_PolyHigher (D,D,C,B,A_C,E,A,D,E)
Spike−in proteins CPTAC Data Linear_PolyHigher (A,B,C,D,E_D,B,E,C,B)
Spike−in proteins CPTAC Data Linear_PolyHigher (E,D,D,C,B_D,B,E,C,B)
Normalized Intensity
Timepoint
Condition2
Condition1
10
15
20
25
30
1
2
3
4
5

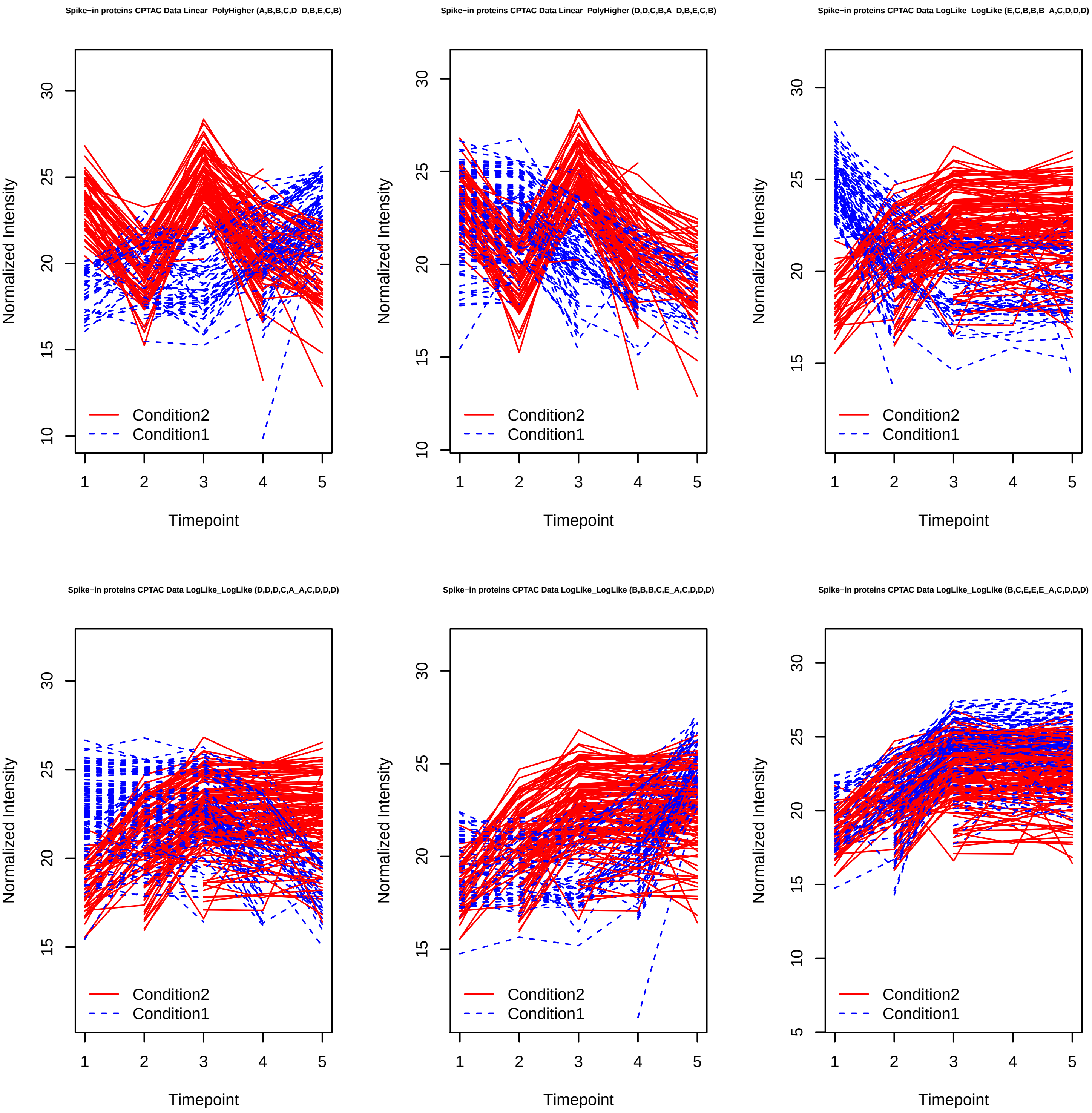
Spike−in proteins CPTAC Data Linear_PolyHigher (A,B,B,C,D_D,B,E,C,B)
Spike−in proteins CPTAC Data Linear_PolyHigher (D,D,C,B,A_D,B,E,C,B)
Spike−in proteins CPTAC Data LogLike_LogLike (E,C,B,B,B_A,C,D,D,D)
Spike−in proteins CPTAC Data LogLike_LogLike (D,D,D,C,A_A,C,D,D,D)
Spike−in proteins CPTAC Data LogLike_LogLike (B,B,B,C,E_A,C,D,D,D)
Spike−in proteins CPTAC Data LogLike_LogLike (B,C,E,E,E_A,C,D,D,D)
Normalized Intensity
Timepoint
Condition2
Condition1

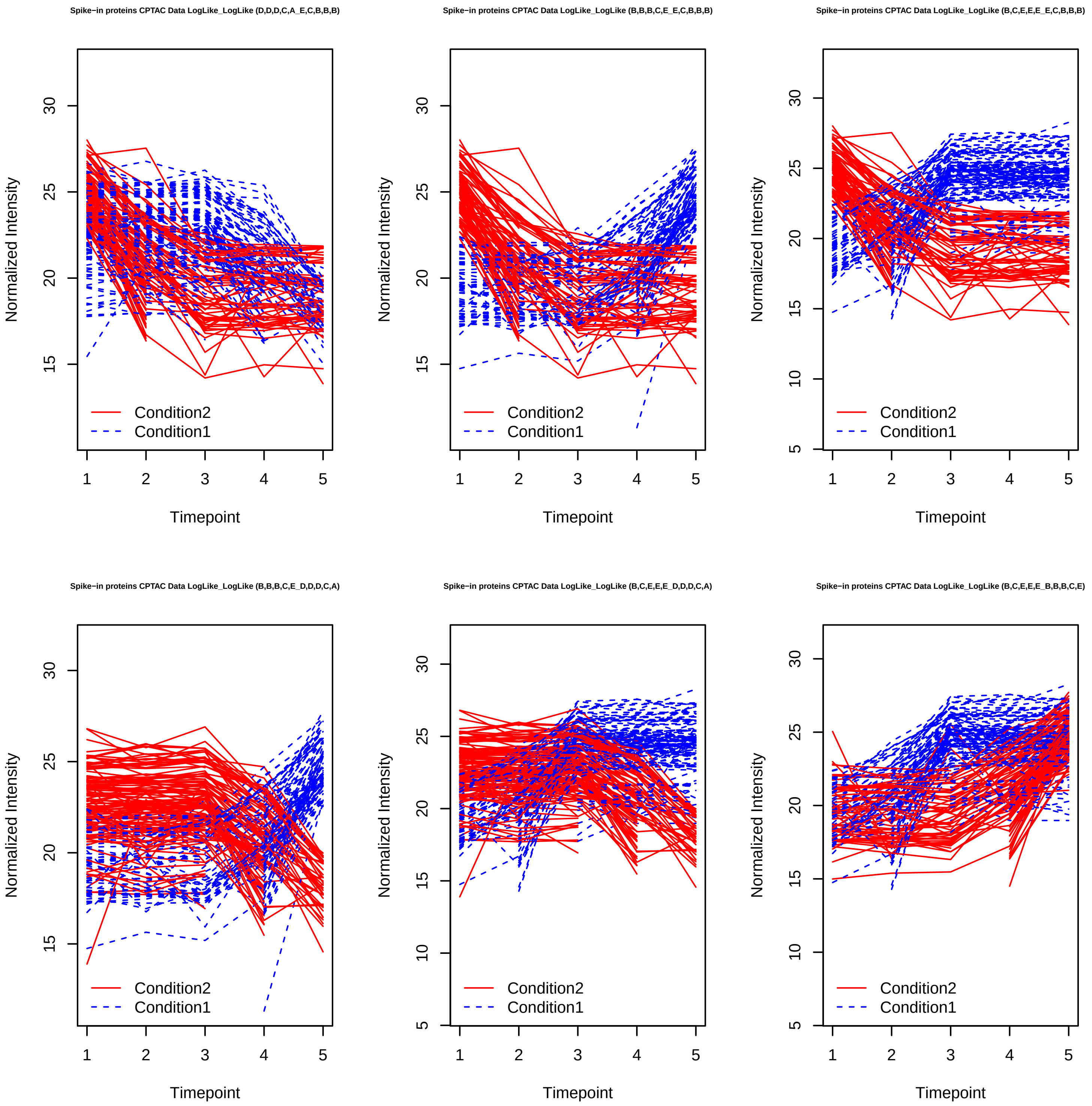
Spike−in proteins CPTAC Data LogLike_LogLike (D,D,D,C,A_E,C,B,B,B)
Spike−in proteins CPTAC Data LogLike_LogLike (B,B,B,C,E_E,C,B,B,B)
Spike−in proteins CPTAC Data LogLike_LogLike (B,C,E,E,E_E,C,B,B,B)
Spike−in proteins CPTAC Data LogLike_LogLike (B,B,B,C,E_D,D,D,C,A)
Spike−in proteins CPTAC Data LogLike_LogLike (B,C,E,E,E_D,D,D,C,A)
Spike−in proteins CPTAC Data LogLike_LogLike (B,C,E,E,E_B,B,B,C,E)
Normalized Intensity
Timepoint
Condition2
Condition1

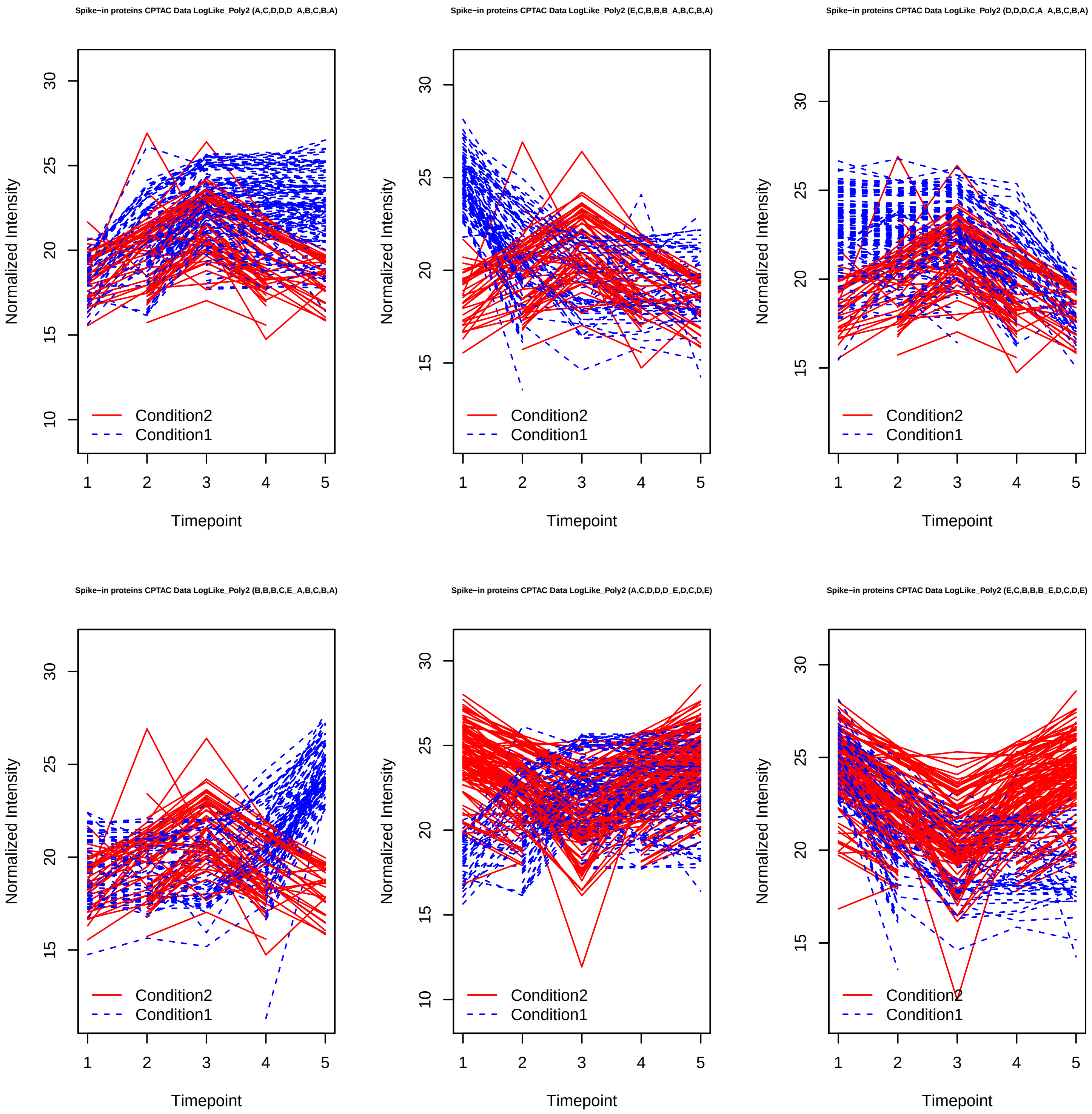
Spike−in proteins CPTAC Data LogLike_Poly2 (A,C,D,D,D_A,B,C,B,A)
Spike−in proteins CPTAC Data LogLike_Poly2 (E,C,B,B,B_A,B,C,B,A)
Spike−in proteins CPTAC Data LogLike_Poly2 (D,D,D,C,A_A,B,C,B,A)
Spike−in proteins CPTAC Data LogLike_Poly2 (B,B,B,C,E_A,B,C,B,A)
Spike−in proteins CPTAC Data LogLike_Poly2 (A,C,D,D,D_E,D,C,D,E)
Spike−in proteins CPTAC Data LogLike_Poly2 (E,C,B,B,B_E,D,C,D,E)
Normalized Intensity
Timepoint
Condition2
Condition1
10
15
20
25
30
1
2
3
4
5

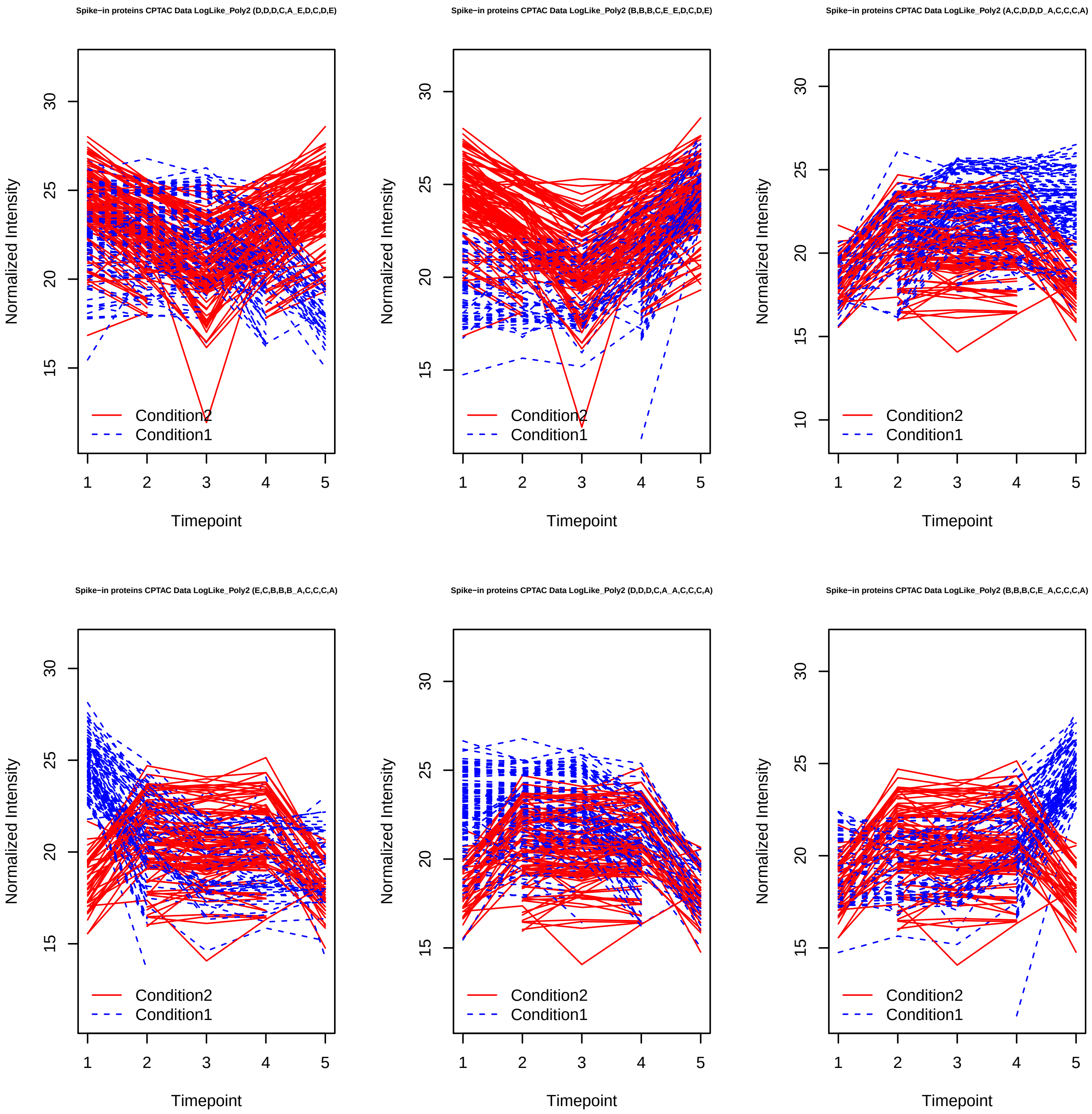
Spike−in proteins CPTAC Data LogLike_Poly2 (D,D,D,C,A_E,D,C,D,E)
Spike−in proteins CPTAC Data LogLike_Poly2 (B,B,B,C,E_E,D,C,D,E)
Spike−in proteins CPTAC Data LogLike_Poly2 (A,C,D,D,D_A,C,C,C,A)
Spike−in proteins CPTAC Data LogLike_Poly2 (E,C,B,B,B_A,C,C,C,A)
Spike−in proteins CPTAC Data LogLike_Poly2 (D,D,D,C,A_A,C,C,C,A)
Spike−in proteins CPTAC Data LogLike_Poly2 (B,B,B,C,E_A,C,C,C,A)
Normalized Intensity
Timepoint
Condition2
Condition1

Spike−in proteins CPTAC Data LogLike_Poly2 (A,C,D,D,D_E,C,C,C,E)

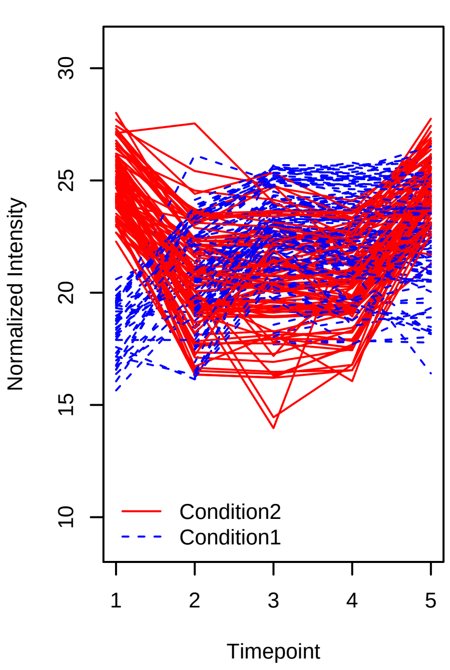


Spike−in proteins CPTAC Data LogLike_Poly2 (E,C,B,B,B_E,C,C,C,E)

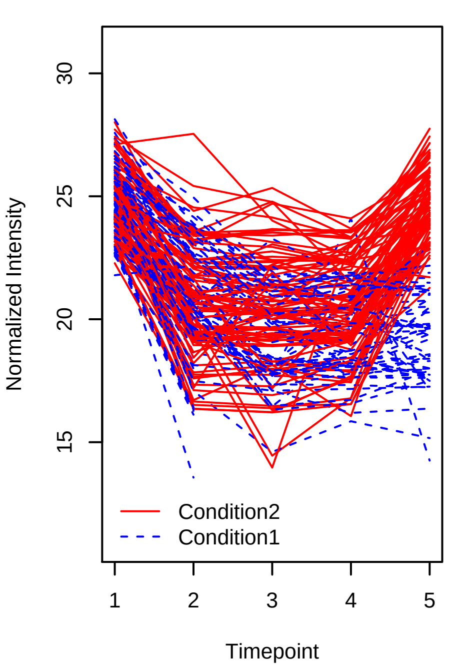


Spike−in proteins CPTAC Data LogLike_Poly2 (D,D,D,C,A_E,C,C,C,E)

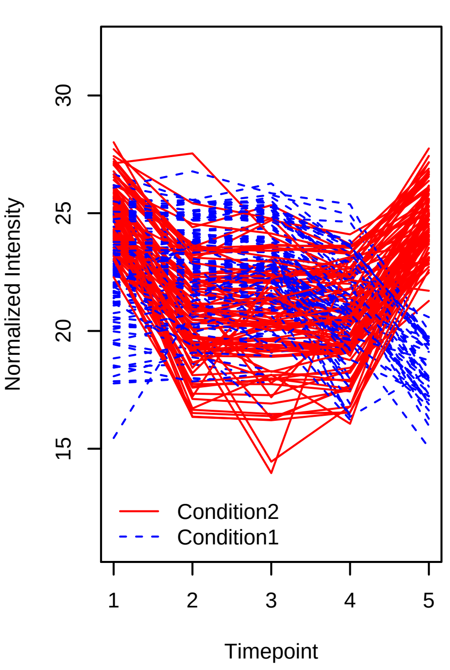


Spike−in proteins CPTAC Data LogLike_Poly2 (B,B,B,C,E_E,C,C,C,E)

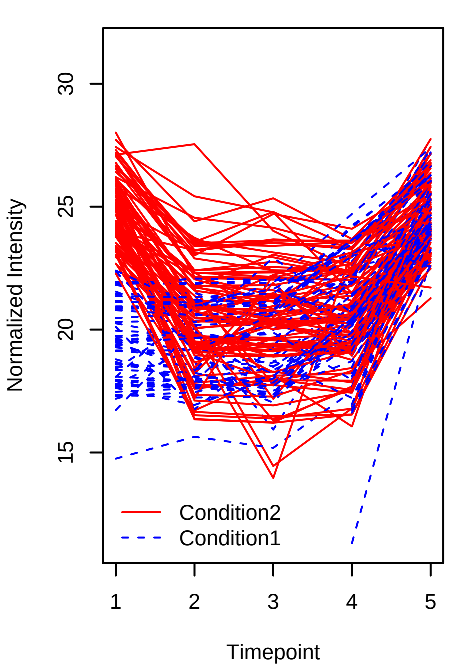


Spike−in proteins CPTAC Data LogLike_Sigmoid (A,C,D,D,D_A,B,B,D,D)

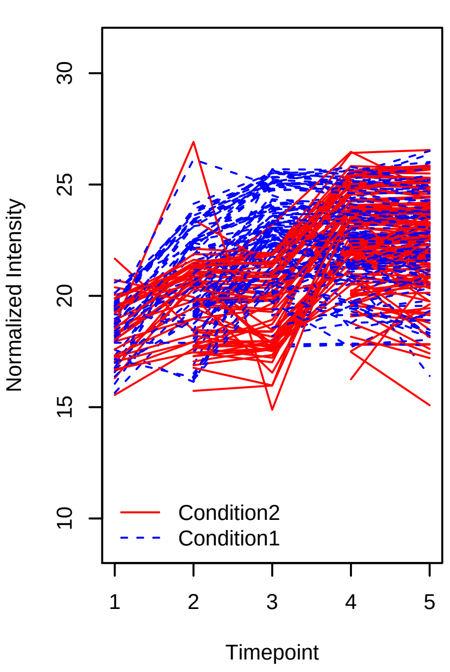


Spike−in proteins CPTAC Data LogLike_Sigmoid (E,C,B,B,B_A,B,B,D,D)

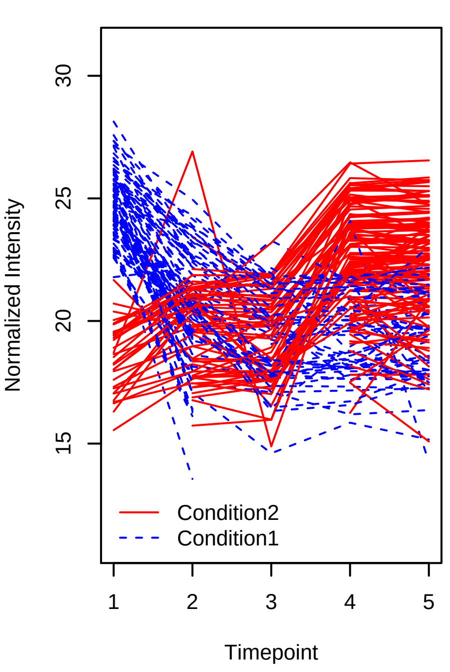

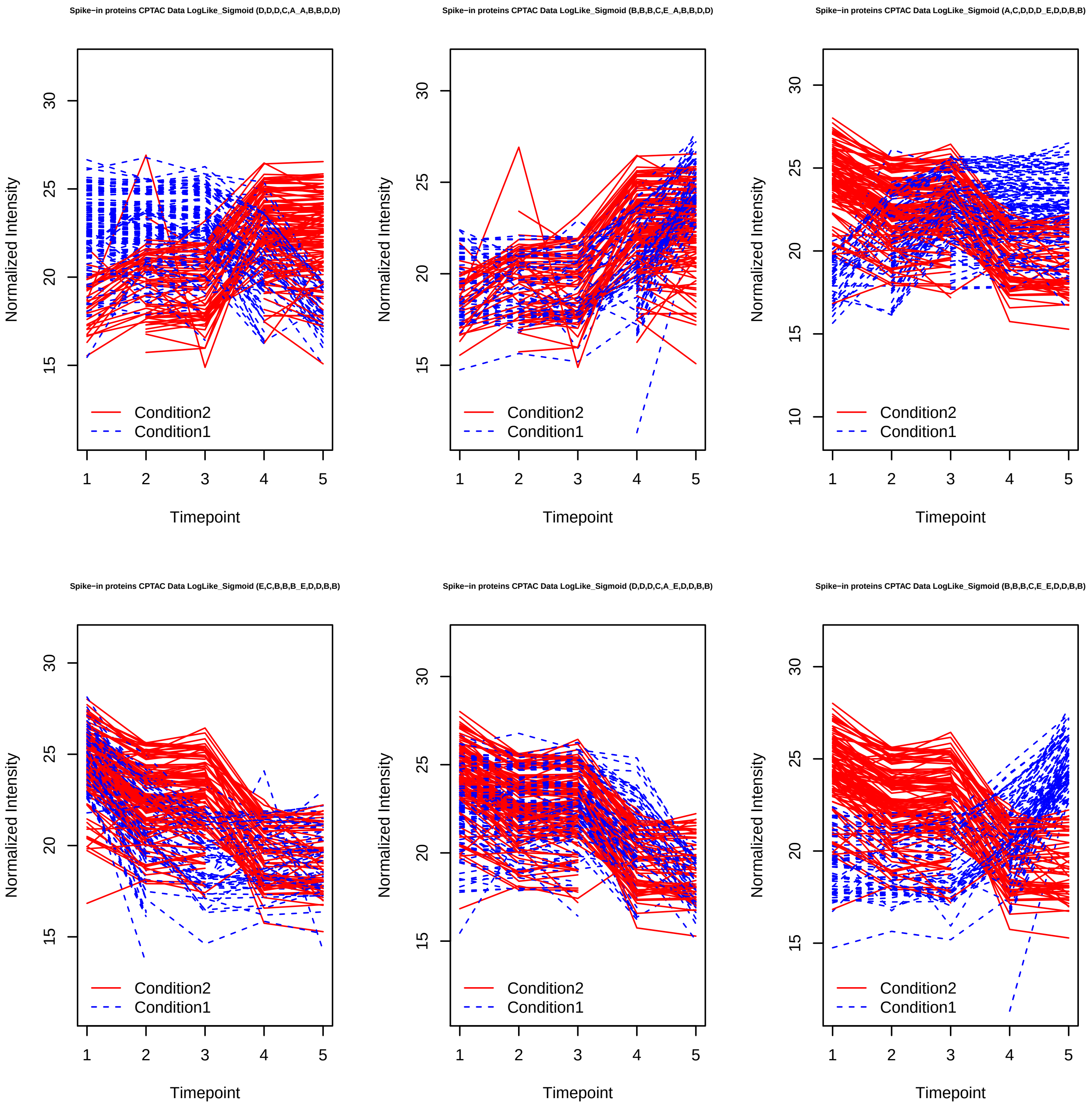
Spike−in proteins CPTAC Data LogLike_Sigmoid (D,D,D,C,A_A,B,B,D,D)
Spike−in proteins CPTAC Data LogLike_Sigmoid (B,B,B,C,E_A,B,B,D,D)
Spike−in proteins CPTAC Data LogLike_Sigmoid (A,C,D,D,D_E,D,D,B,B)
Spike−in proteins CPTAC Data LogLike_Sigmoid (E,C,B,B,B_E,D,D,B,B)
Spike−in proteins CPTAC Data LogLike_Sigmoid (D,D,D,C,A_E,D,D,B,B)
Spike−in proteins CPTAC Data LogLike_Sigmoid (B,B,B,C,E_E,D,D,B,B)
Normalized Intensity
Timepoint
Condition2
Condition1

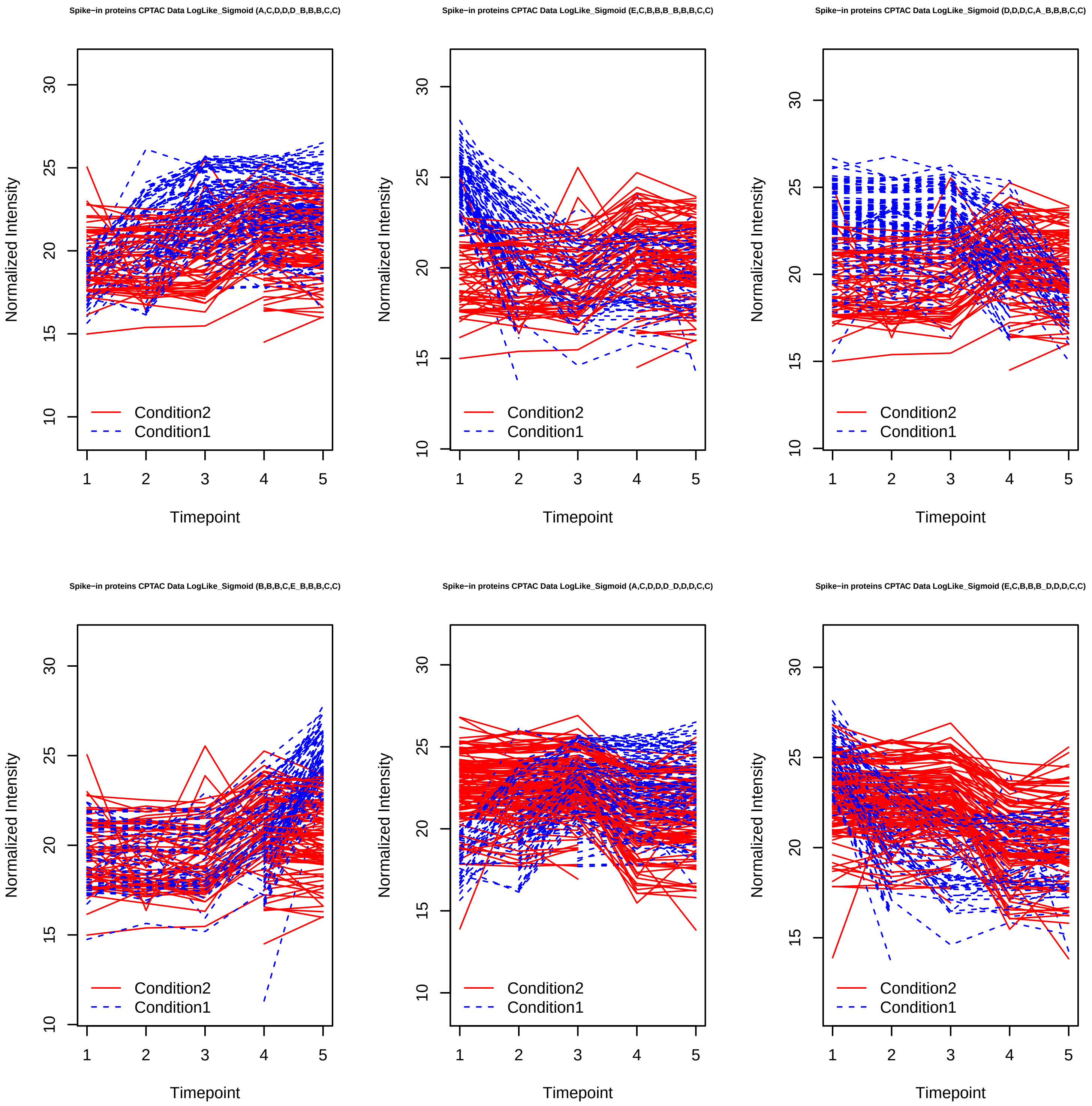
Spike−in proteins CPTAC Data LogLike_Sigmoid (A,C,D,D,D_B,B,B,C,C)
Spike−in proteins CPTAC Data LogLike_Sigmoid (E,C,B,B,B_B,B,B,C,C)
Spike−in proteins CPTAC Data LogLike_Sigmoid (D,D,D,C,A_B,B,B,C,C)
Spike−in proteins CPTAC Data LogLike_Sigmoid (B,B,B,C,E_B,B,B,C,C)
Spike−in proteins CPTAC Data LogLike_Sigmoid (A,C,D,D,D_D,D,D,C,C)
Spike−in proteins CPTAC Data LogLike_Sigmoid (E,C,B,B,B_D,D,D,C,C)
Normalized Intensity
Timepoint
Condition2
Condition1

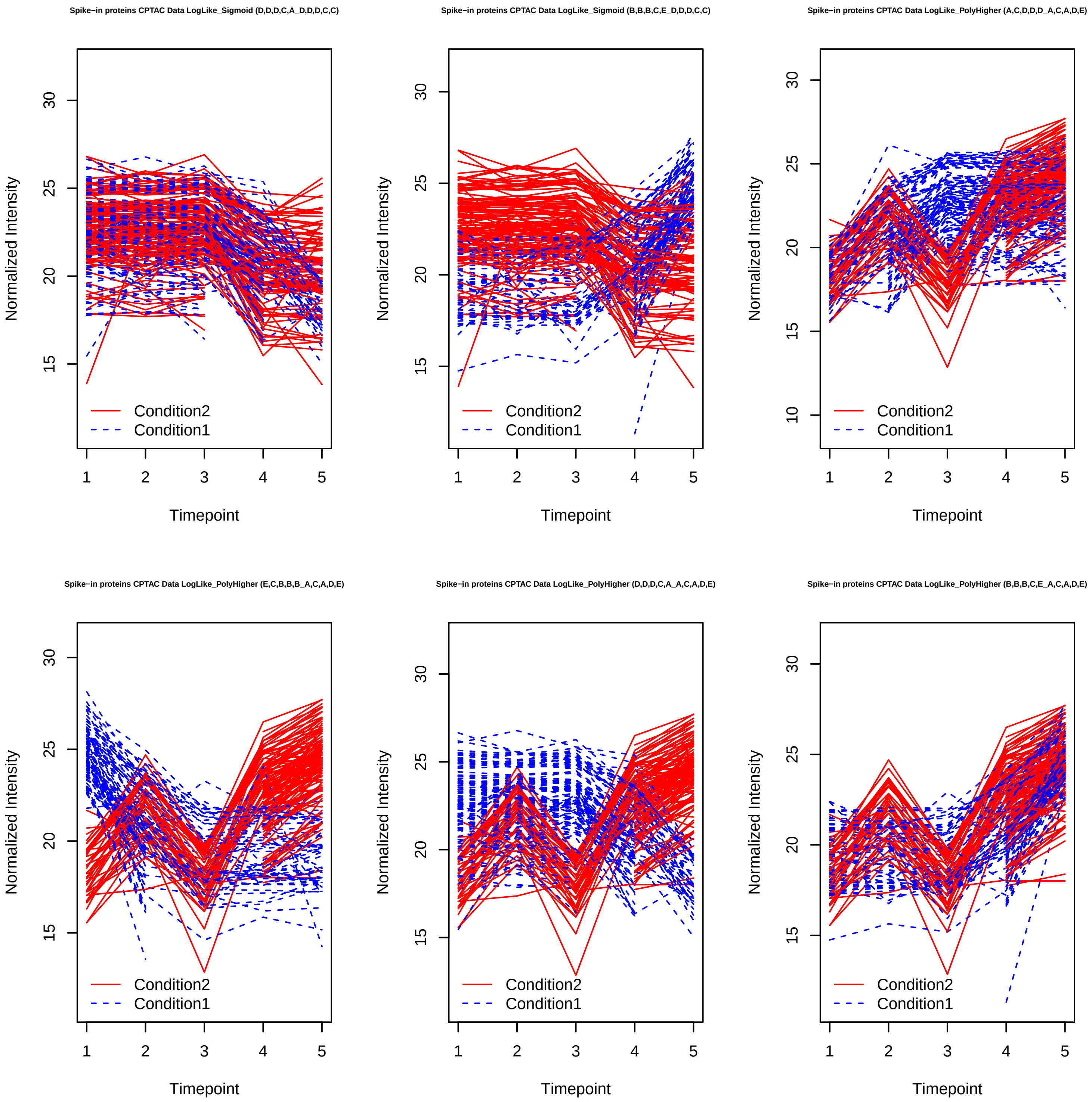
Spike−in proteins CPTAC Data LogLike_Sigmoid (D,D,D,C,A_D,D,D,C,C)
Spike−in proteins CPTAC Data LogLike_Sigmoid (B,B,B,C,E_D,D,D,C,C)
Spike−in proteins CPTAC Data LogLike_PolyHigher (A,C,D,D,D_A,C,A,D,E)
Spike−in proteins CPTAC Data LogLike_PolyHigher (E,C,B,B,B_A,C,A,D,E)
Spike−in proteins CPTAC Data LogLike_PolyHigher (D,D,D,C,A_A,C,A,D,E)
Spike−in proteins CPTAC Data LogLike_PolyHigher (B,B,B,C,E_A,C,A,D,E)
Normalized Intensity
Timepoint
Condition2
Condition1

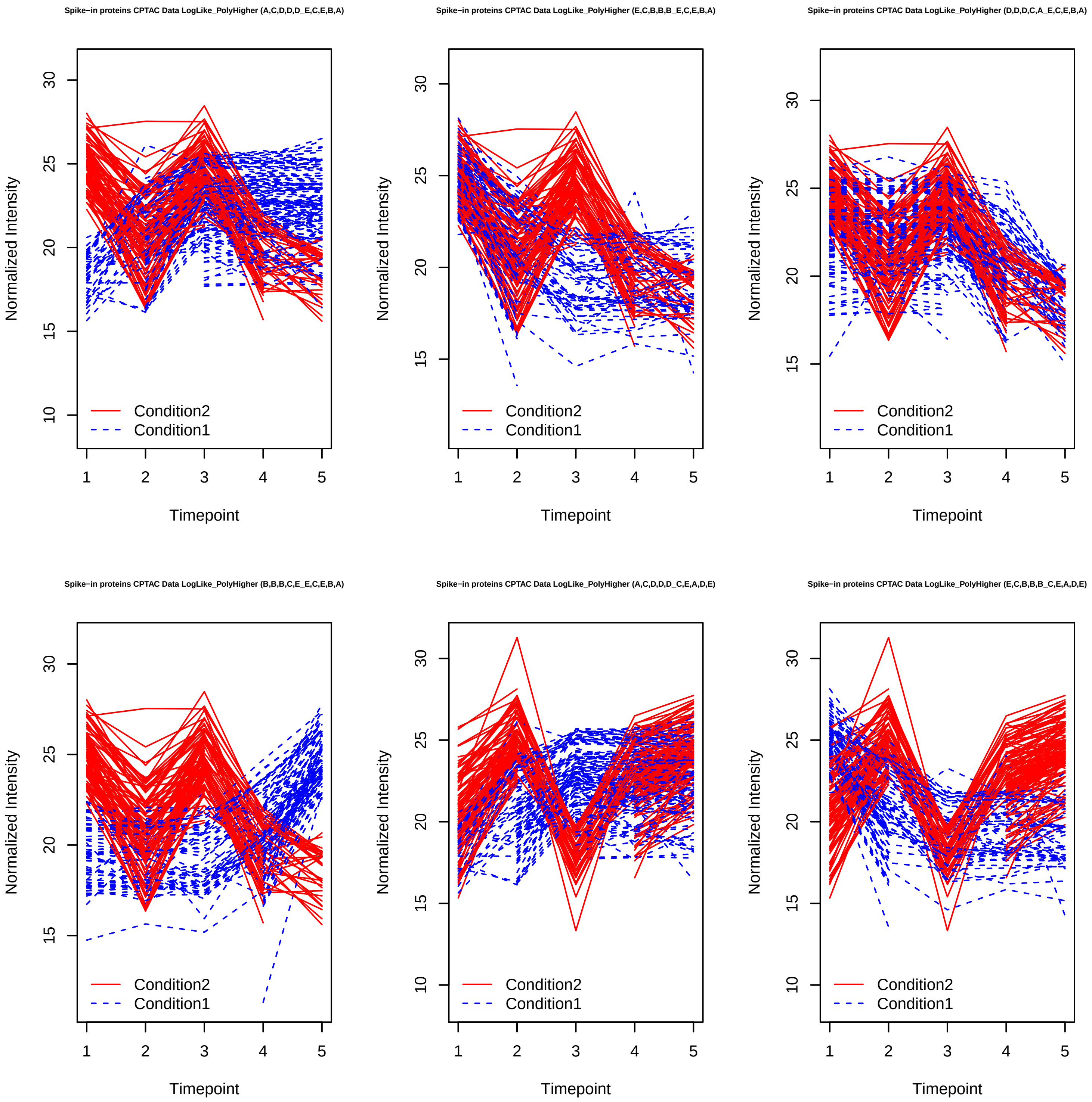
Spike−in proteins CPTAC Data LogLike_PolyHigher (A,C,D,D,D_E,C,E,B,A)
Spike−in proteins CPTAC Data LogLike_PolyHigher (E,C,B,B,B_E,C,E,B,A)
Spike−in proteins CPTAC Data LogLike_PolyHigher (D,D,D,C,A_E,C,E,B,A)
Spike−in proteins CPTAC Data LogLike_PolyHigher (B,B,B,C,E_E,C,E,B,A)
Spike−in proteins CPTAC Data LogLike_PolyHigher (A,C,D,D,D_C,E,A,D,E)
Spike−in proteins CPTAC Data LogLike_PolyHigher (E,C,B,B,B_C,E,A,D,E)
Normalized Intensity
Timepoint
Condition2
Condition1

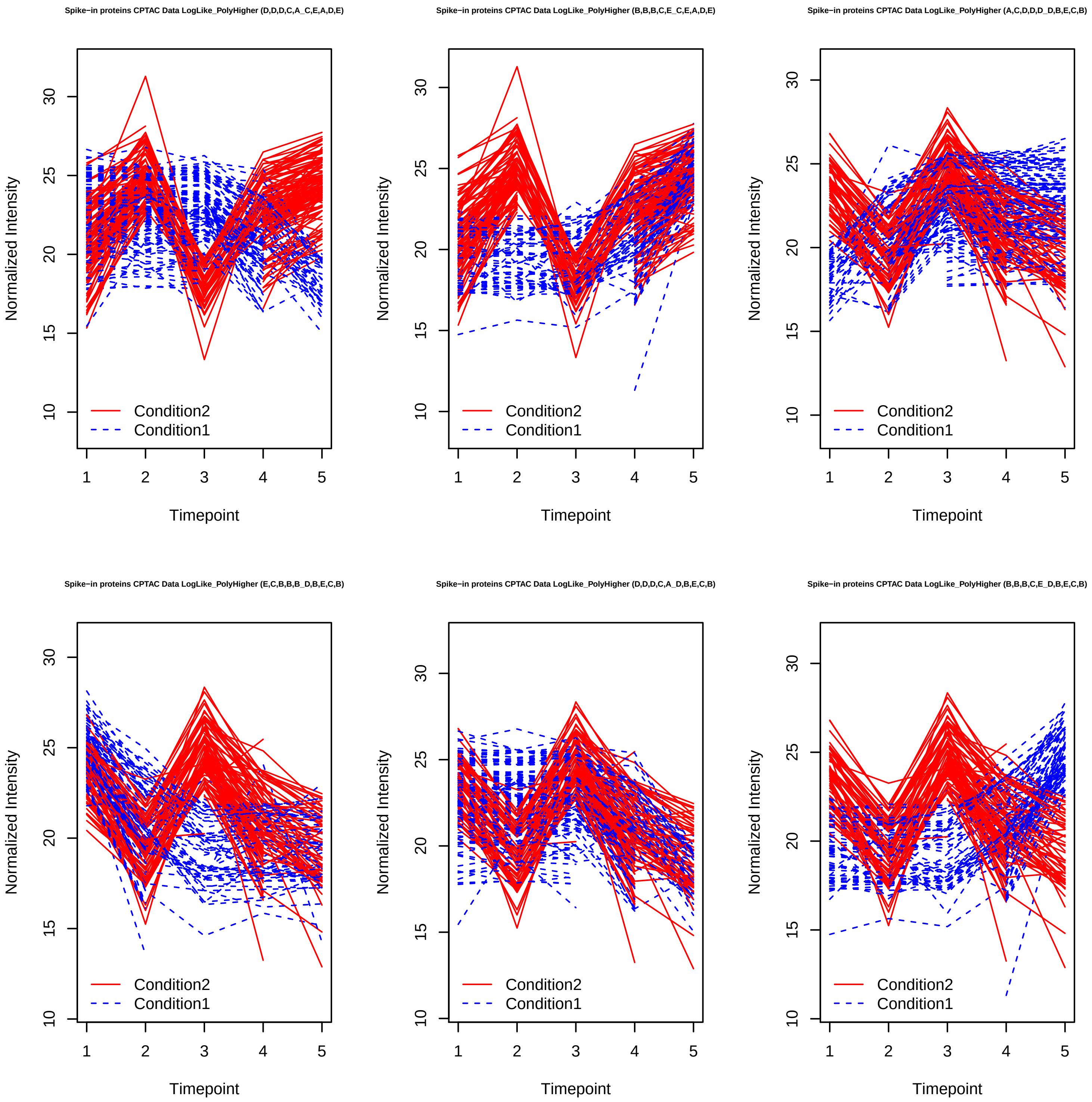
Spike−in proteins CPTAC Data LogLike_PolyHigher (D,D,D,C,A_C,E,A,D,E)
Spike−in proteins CPTAC Data LogLike_PolyHigher (B,B,B,C,E_C,E,A,D,E)
Spike−in proteins CPTAC Data LogLike_PolyHigher (A,C,D,D,D_D,B,E,C,B)
Spike−in proteins CPTAC Data LogLike_PolyHigher (E,C,B,B,B_D,B,E,C,B)
Spike−in proteins CPTAC Data LogLike_PolyHigher (D,D,D,C,A_D,B,E,C,B)
Spike−in proteins CPTAC Data LogLike_PolyHigher (B,B,B,C,E_D,B,E,C,B)
Normalized Intensity
Timepoint
Condition2
Condition1

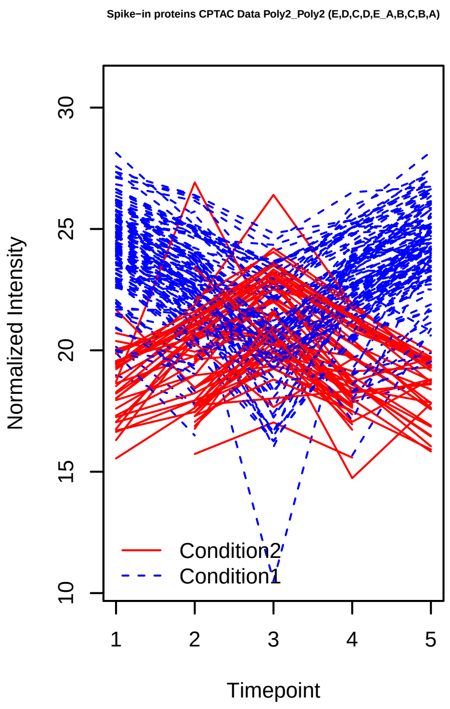
Spike−in proteins CPTAC Data Poly2_Poly2 (E,D,C,D,E_A,B,C,B,A)
Normalized Intensity
Timepoint
Condition2
Condition1

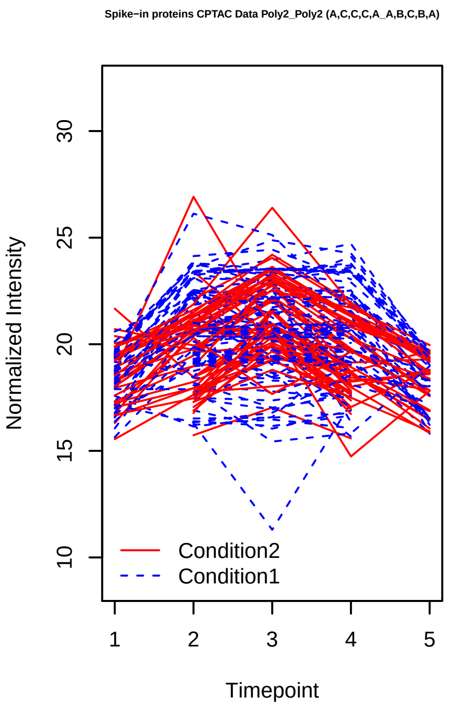
Spike−in proteins CPTAC Data Poly2_Poly2 (A,C,C,C,A_A,B,C,B,A)
Normalized Intensity
Timepoint
Condition2
Condition1

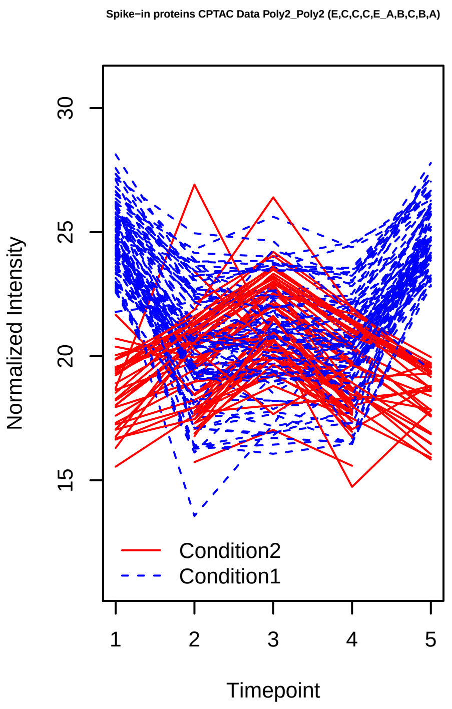
Spike−in proteins CPTAC Data Poly2_Poly2 (E,C,C,C,E_A,B,C,B,A)
Normalized Intensity
Timepoint
Condition2
Condition1

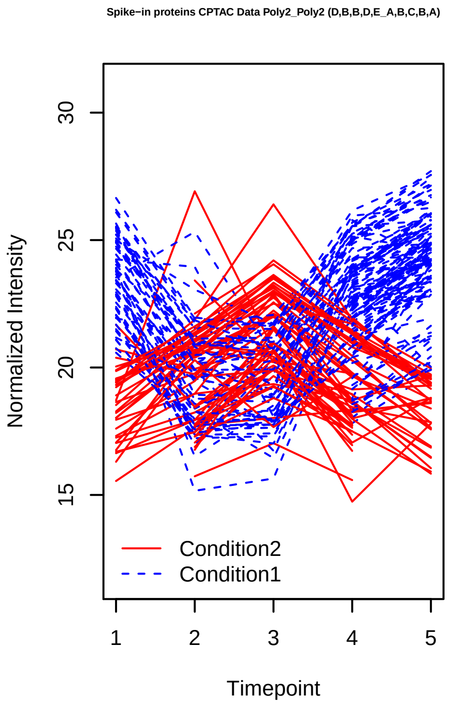
Spike−in proteins CPTAC Data Poly2_Poly2 (D,B,B,D,E_A,B,C,B,A)
Normalized Intensity
Timepoint
Condition2
Condition1

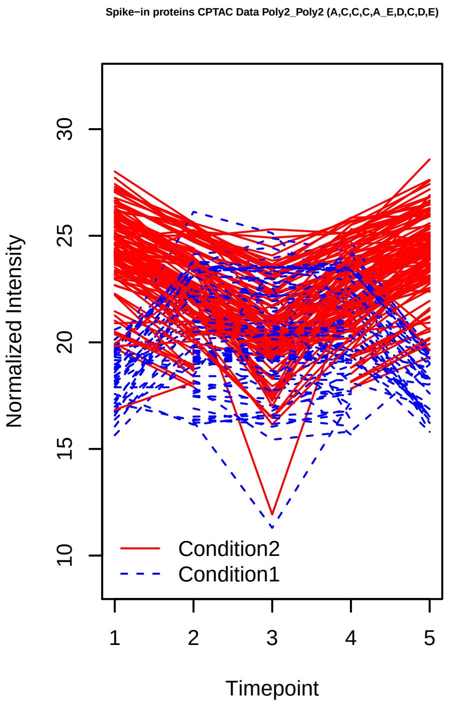
Spike−in proteins CPTAC Data Poly2_Poly2 (A,C,C,C,A_E,D,C,D,E)
Normalized Intensity
Timepoint
Condition2
Condition1

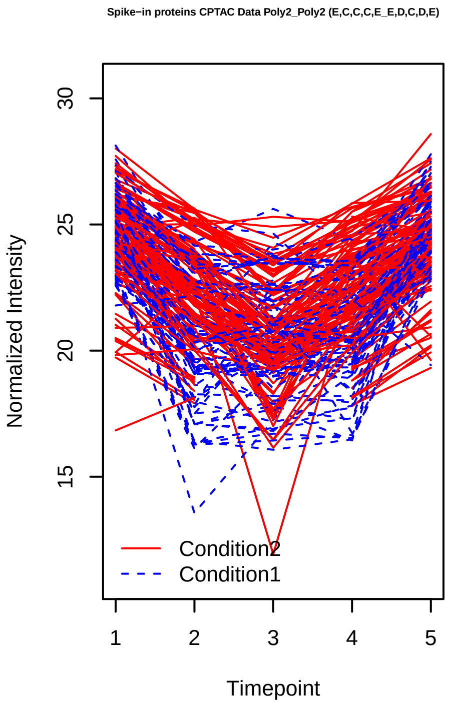
Spike−in proteins CPTAC Data Poly2_Poly2 (E,C,C,C,E_E,D,C,D,E)
Normalized Intensity
Timepoint
Condition2
Condition1

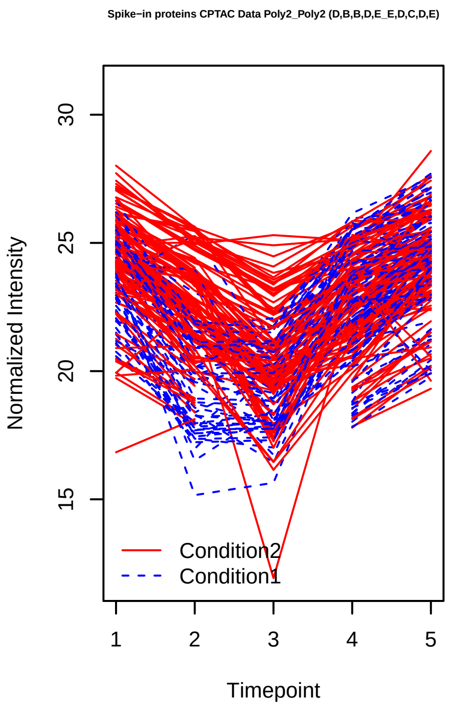
Spike−in proteins CPTAC Data Poly2_Poly2 (D,B,B,D,E_E,D,C,D,E)
Normalized Intensity
Timepoint
Condition2
Condition1

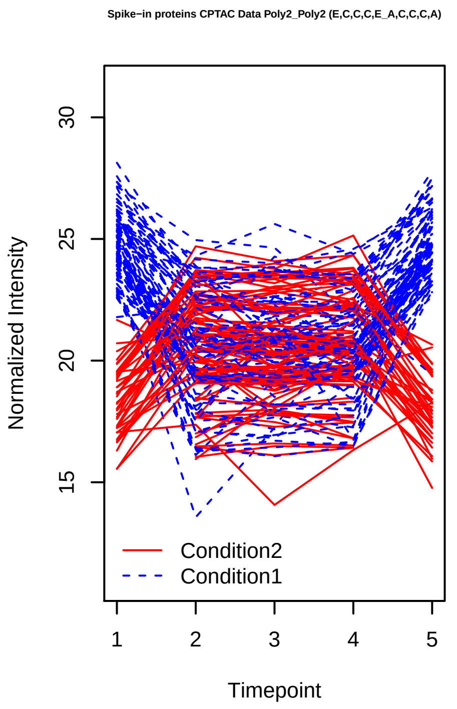
Spike−in proteins CPTAC Data Poly2_Poly2 (E,C,C,C,E_A,C,C,C,A)
Normalized Intensity
Timepoint
Condition2
Condition1

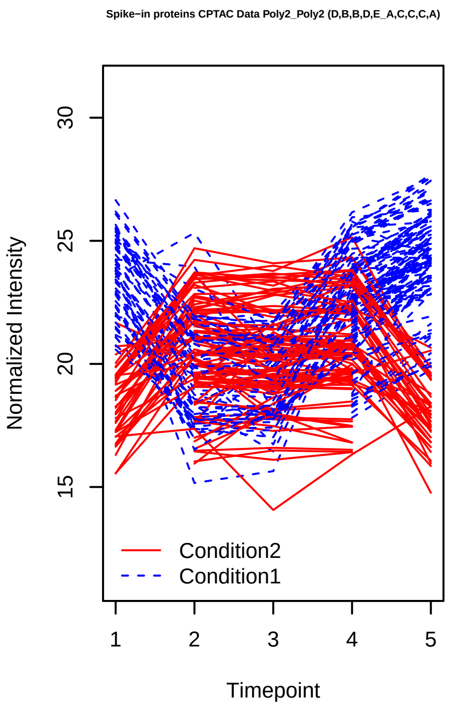
Spike−in proteins CPTAC Data Poly2_Poly2 (D,B,B,D,E_A,C,C,C,A)
Normalized Intensity
Timepoint
Condition2
Condition1

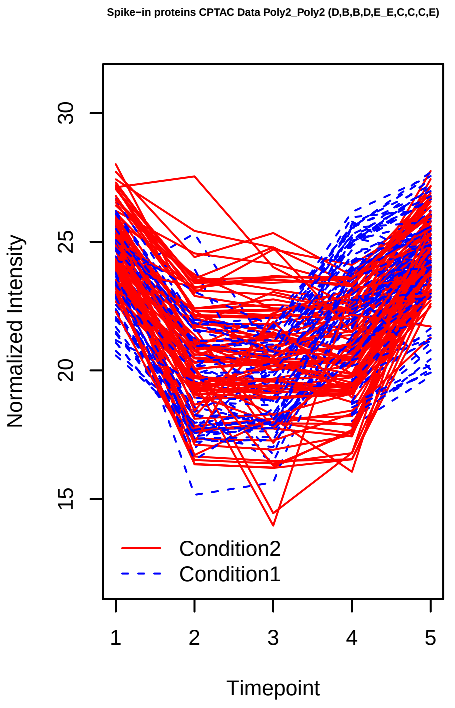
Spike−in proteins CPTAC Data Poly2_Poly2 (D,B,B,D,E_E,C,C,C,E)
Normalized Intensity
Timepoint
Condition2
Condition1

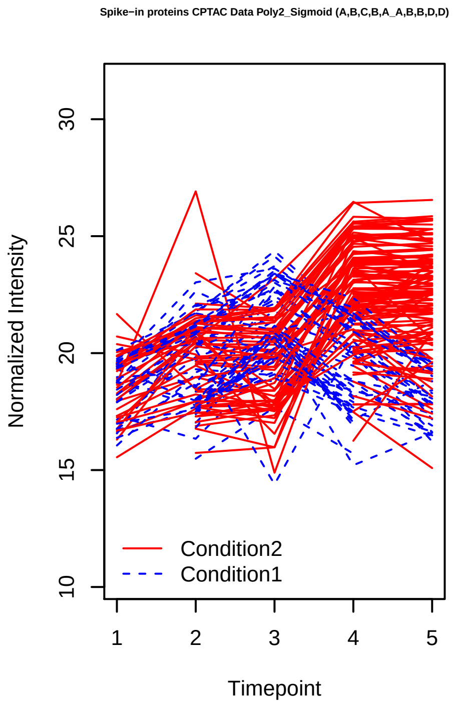
Spike−in proteins CPTAC Data Poly2_Sigmoid (A,B,C,B,A_A,B,B,D,D)
Normalized Intensity
Timepoint
Condition2
Condition1

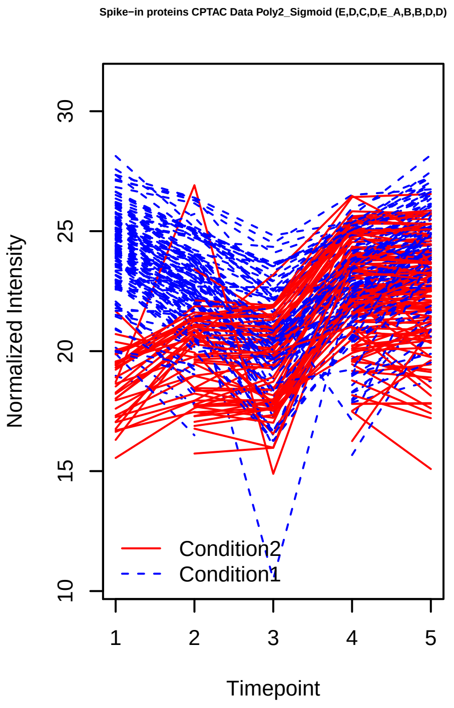
Spike−in proteins CPTAC Data Poly2_Sigmoid (E,D,C,D,E_A,B,B,D,D)
Normalized Intensity
Timepoint
Condition2
Condition1

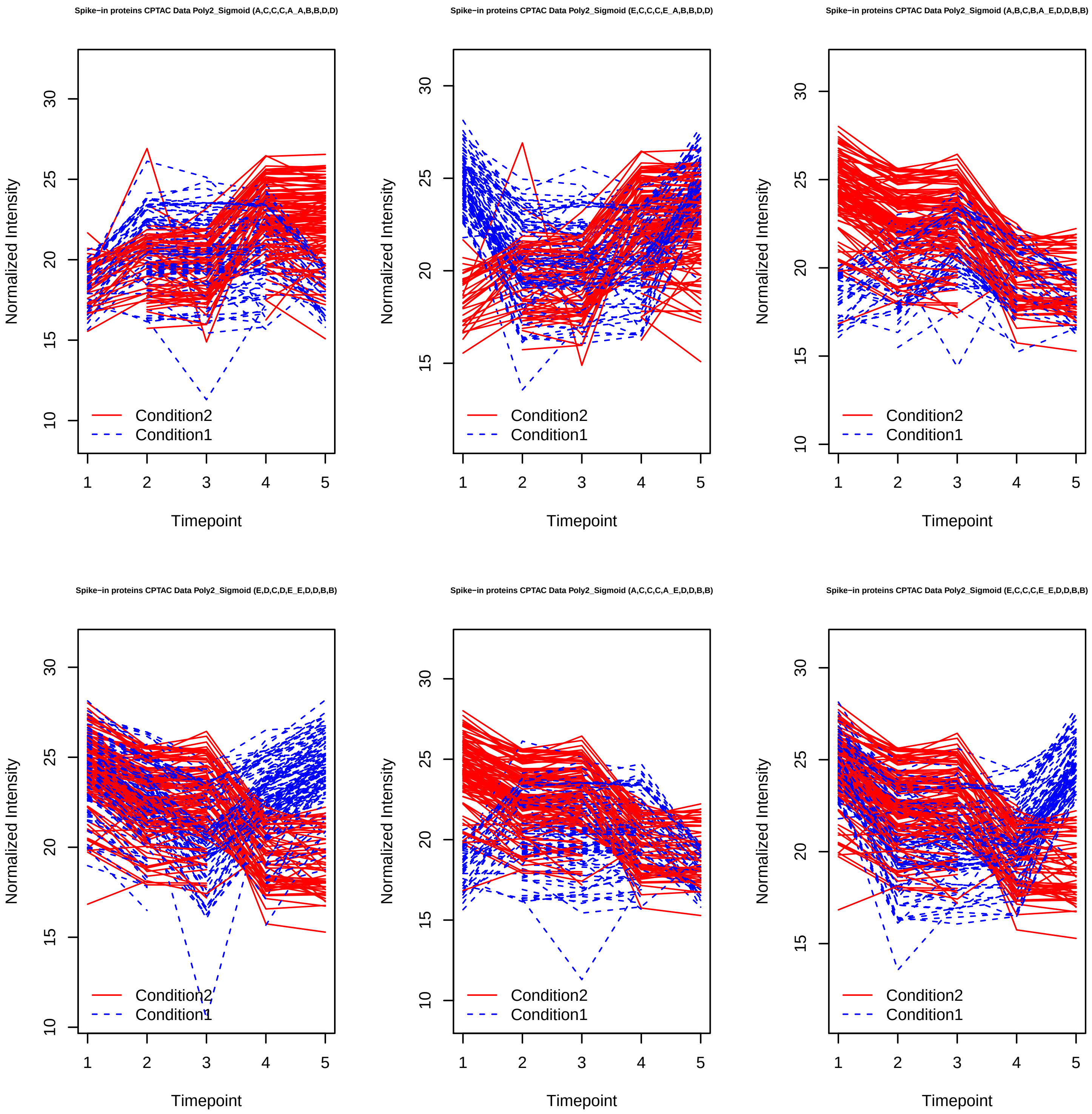

Spike−in proteins CPTAC Data Poly2_Sigmoid (A,C,C,C,A_A,B,B,D,D)
Spike−in proteins CPTAC Data Poly2_Sigmoid (E,C,C,C,E_A,B,B,D,D)
Spike−in proteins CPTAC Data Poly2_Sigmoid (A,B,C,B,A_E,D,D,B,B)
Spike−in proteins CPTAC Data Poly2_Sigmoid (E,D,C,D,E_E,D,D,B,B)
Spike−in proteins CPTAC Data Poly2_Sigmoid (A,C,C,C,A_E,D,D,B,B)
Spike−in proteins CPTAC Data Poly2_Sigmoid (E,C,C,C,E_E,D,D,B,B)
Normalized Intensity
Timepoint
Condition2
Condition1

Spike−in proteins CPTAC Data Poly2_Sigmoid (A,B,C,B,A_B,B,B,C,C)

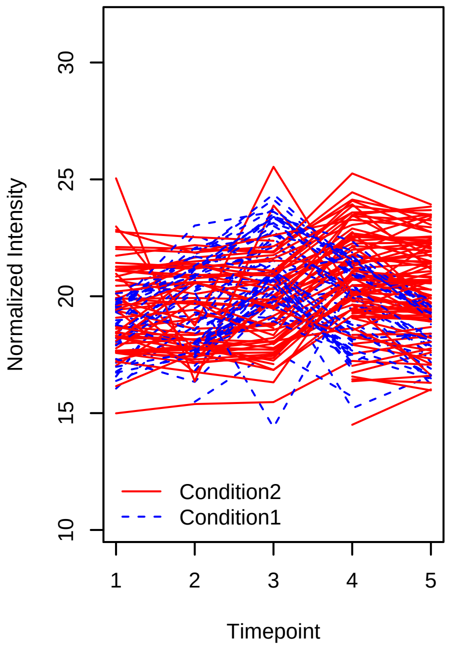


Spike−in proteins CPTAC Data Poly2_Sigmoid (E,D,C,D,E_B,B,B,C,C)

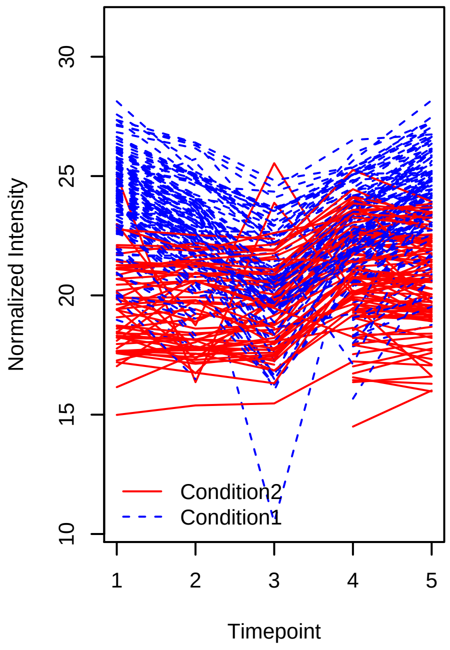


Spike−in proteins CPTAC Data Poly2_Sigmoid (A,C,C,C,A_B,B,B,C,C)

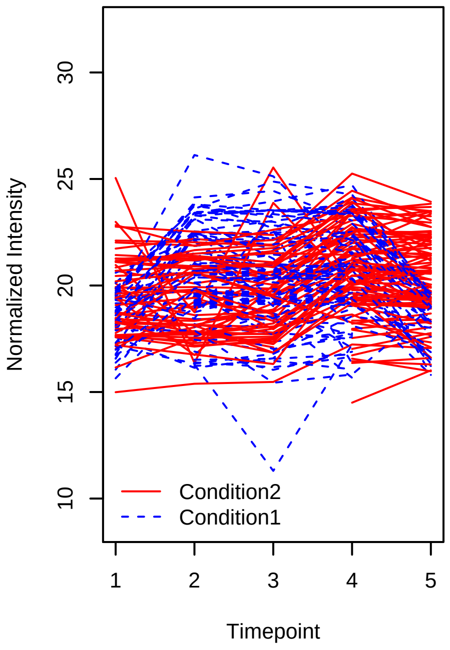


Spike−in proteins CPTAC Data Poly2_Sigmoid (E,C,C,C,E_B,B,B,C,C)

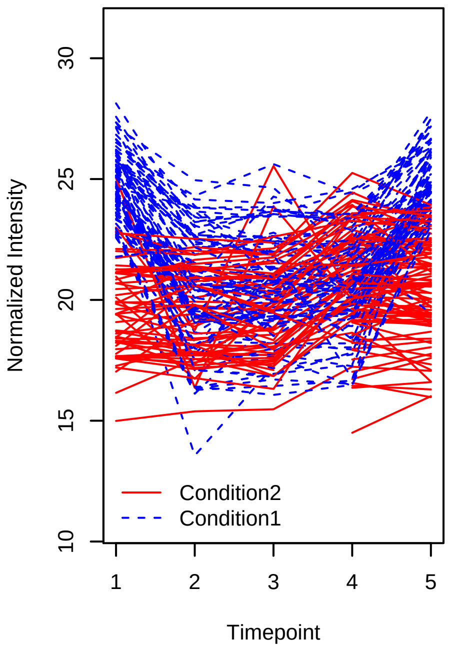


Spike−in proteins CPTAC Data Poly2_Sigmoid (A,B,C,B,A_D,D,D,C,C)

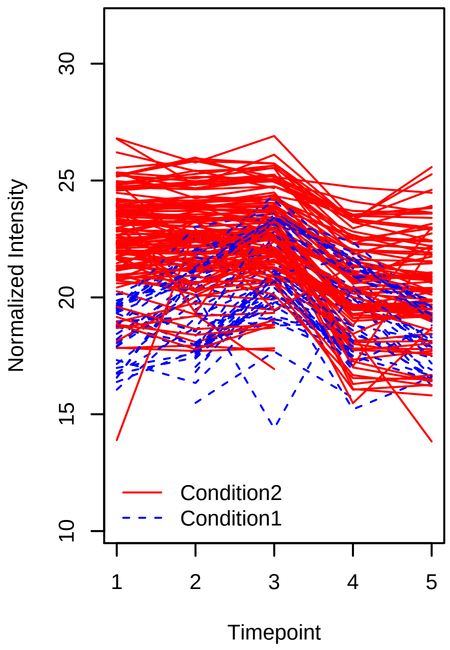


Spike−in proteins CPTAC Data Poly2_Sigmoid (E,D,C,D,E_D,D,D,C,C)

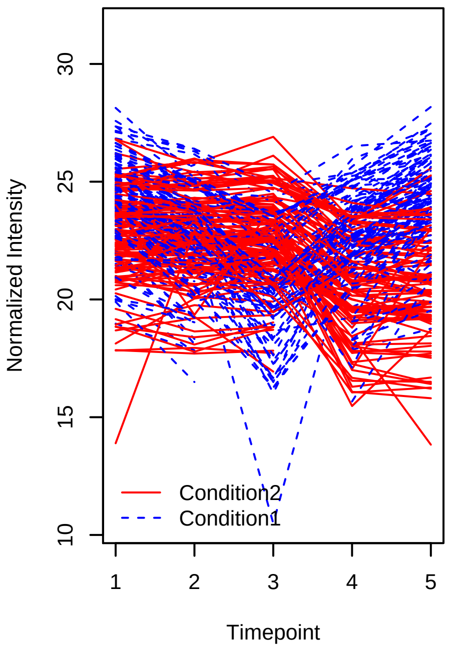

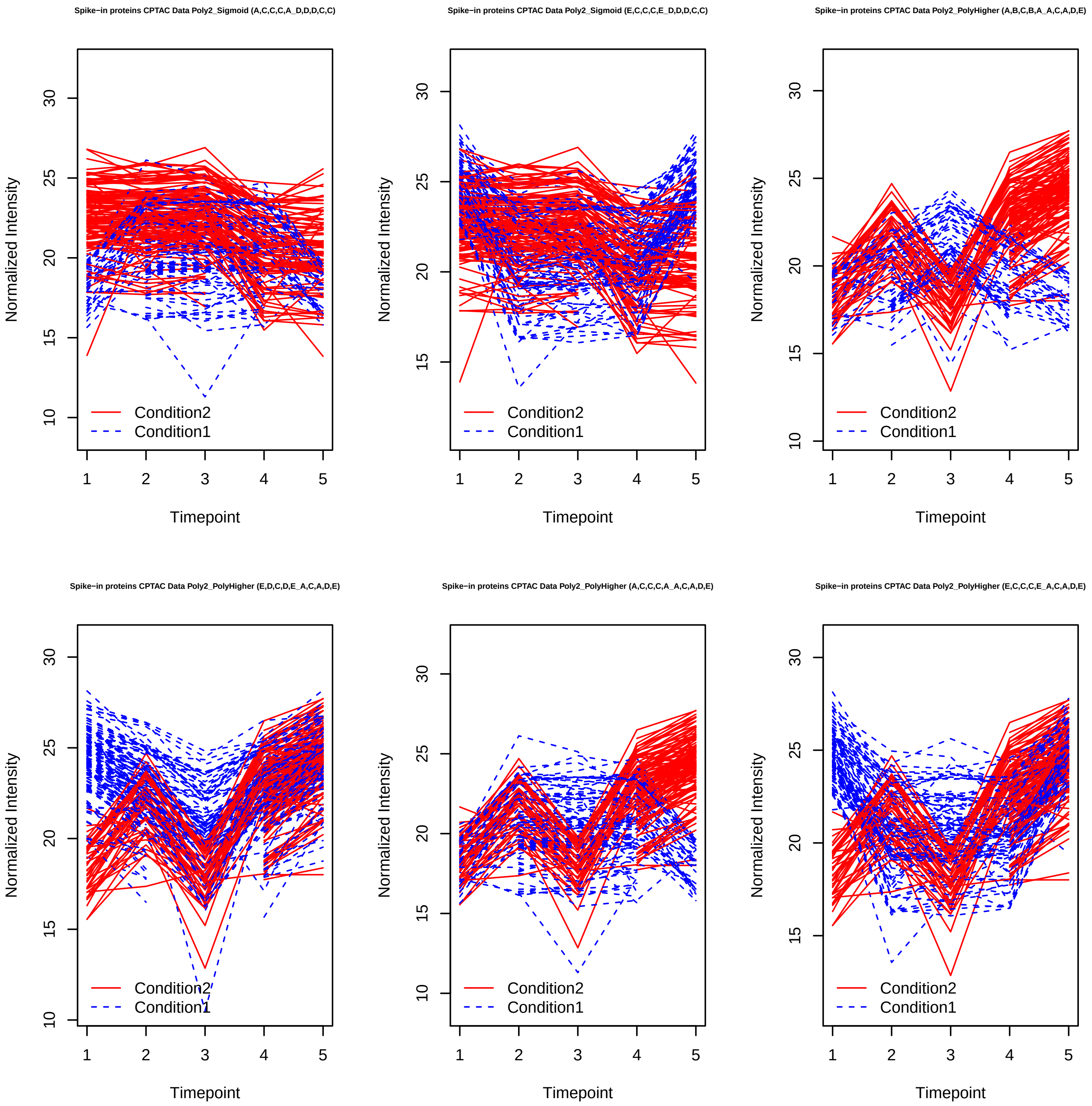
Spike−in proteins CPTAC Data Poly2_Sigmoid (A,C,C,C,A_D,D,D,C,C)
Spike−in proteins CPTAC Data Poly2_Sigmoid (E,C,C,C,E_D,D,D,C,C)
Spike−in proteins CPTAC Data Poly2_PolyHigher (A,B,C,B,A_A,C,A,D,E)
Spike−in proteins CPTAC Data Poly2_PolyHigher (E,D,C,D,E_A,C,A,D,E)
Spike−in proteins CPTAC Data Poly2_PolyHigher (A,C,C,C,A_A,C,A,D,E)
Spike−in proteins CPTAC Data Poly2_PolyHigher (E,C,C,C,E_A,C,A,D,E)
Normalized Intensity
Timepoint
Condition2
Condition1
10
15
20
25
30
1
2
3
4
5

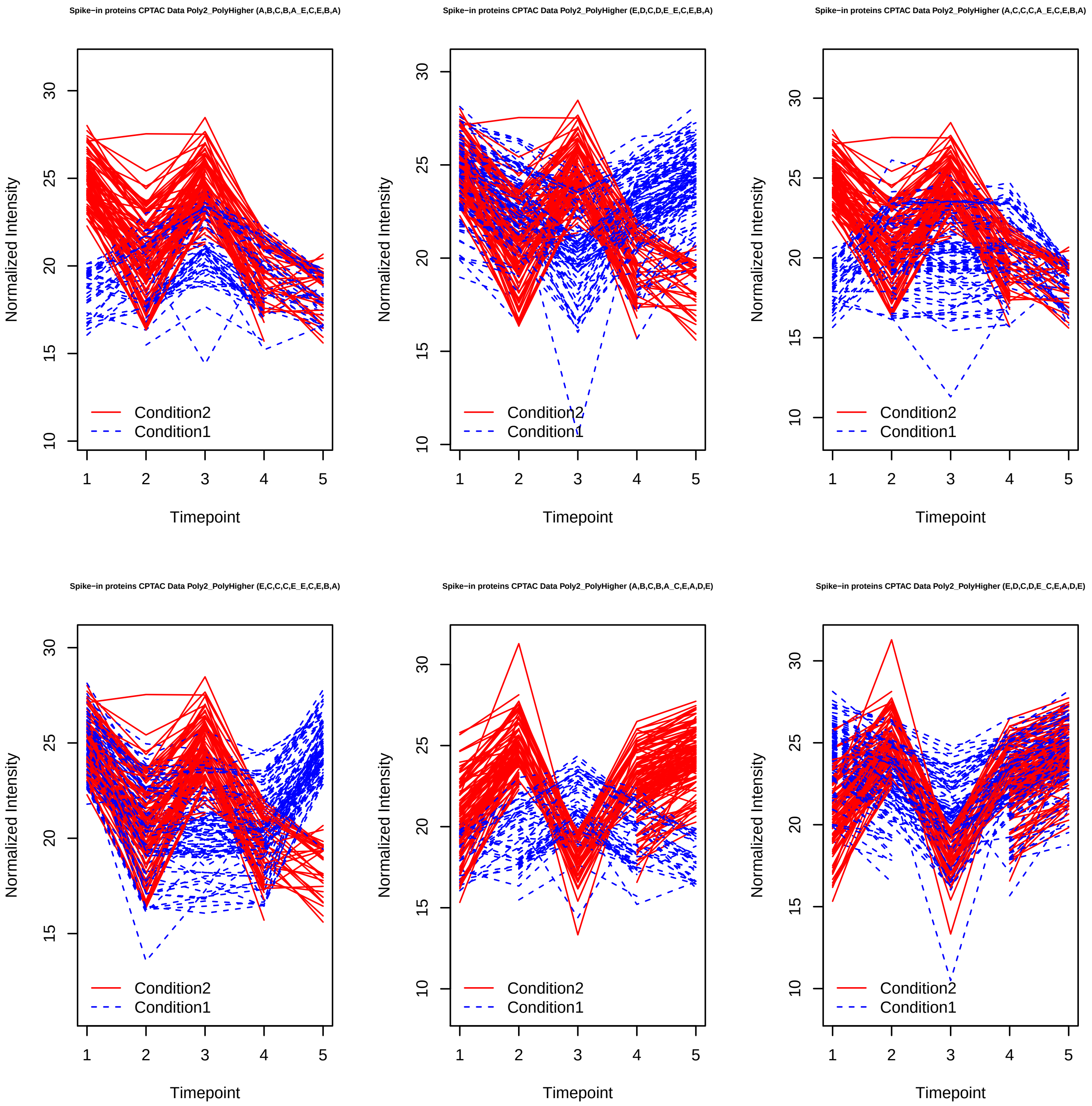
Spike−in proteins CPTAC Data Poly2_PolyHigher (A,B,C,B,A_E,C,E,B,A)
Spike−in proteins CPTAC Data Poly2_PolyHigher (E,D,C,D,E_E,C,E,B,A)
Spike−in proteins CPTAC Data Poly2_PolyHigher (A,C,C,C,A_E,C,E,B,A)
Spike−in proteins CPTAC Data Poly2_PolyHigher (E,C,C,C,E_E,C,E,B,A)
Spike−in proteins CPTAC Data Poly2_PolyHigher (A,B,C,B,A_C,E,A,D,E)
Spike−in proteins CPTAC Data Poly2_PolyHigher (E,D,C,D,E_C,E,A,D,E)
Normalized Intensity
Timepoint
Condition2
Condition1

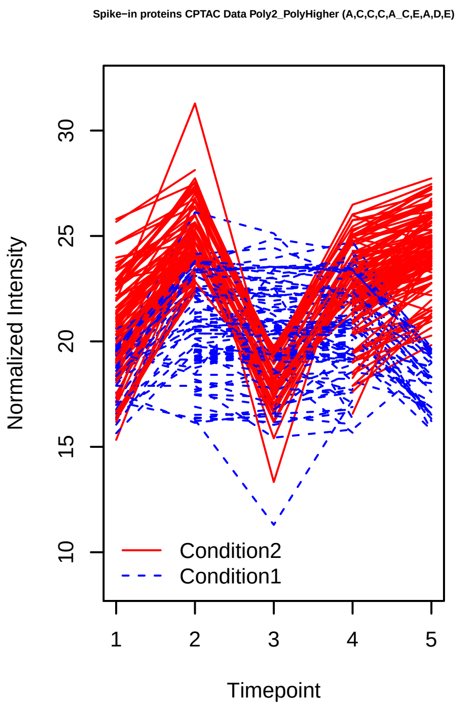
Spike−in proteins CPTAC Data Poly2_PolyHigher (A,C,C,C,A_C,E,A,D,E)
Normalized Intensity
Timepoint
Condition2
Condition1

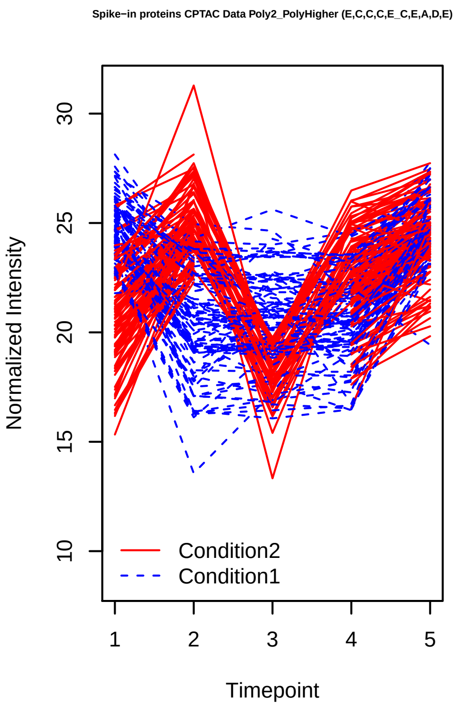
Spike−in proteins CPTAC Data Poly2_PolyHigher (E,C,C,C,E_C,E,A,D,E)
Normalized Intensity
Timepoint
Condition2
Condition1

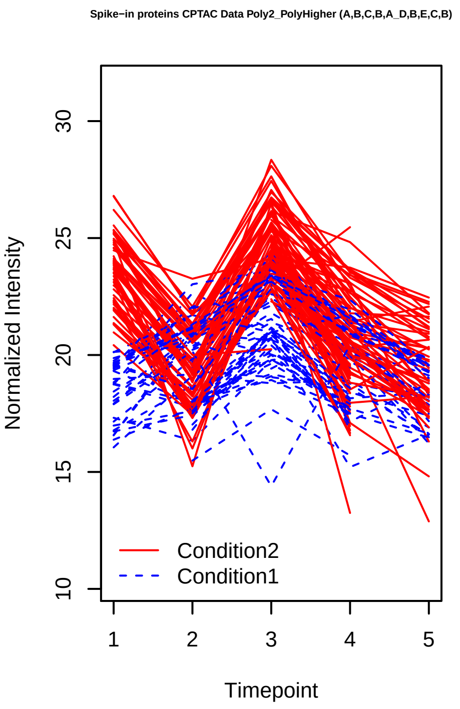
Spike−in proteins CPTAC Data Poly2_PolyHigher (A,B,C,B,A_D,B,E,C,B)
Normalized Intensity
Timepoint
Condition2
Condition1

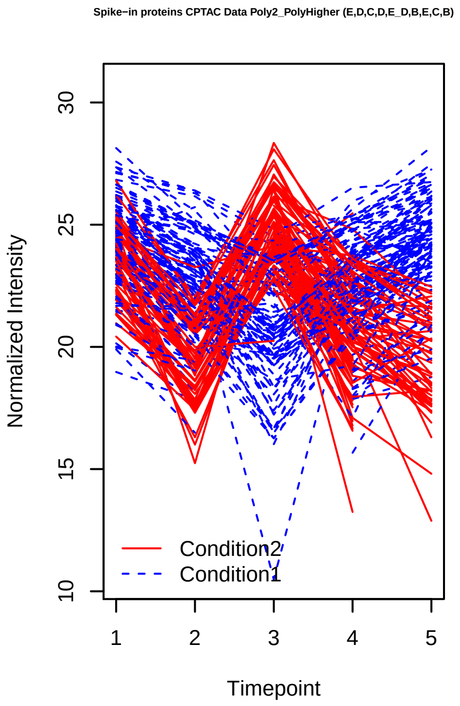
Spike−in proteins CPTAC Data Poly2_PolyHigher (E,D,C,D,E_D,B,E,C,B)
Normalized Intensity
Timepoint
Condition2
Condition1

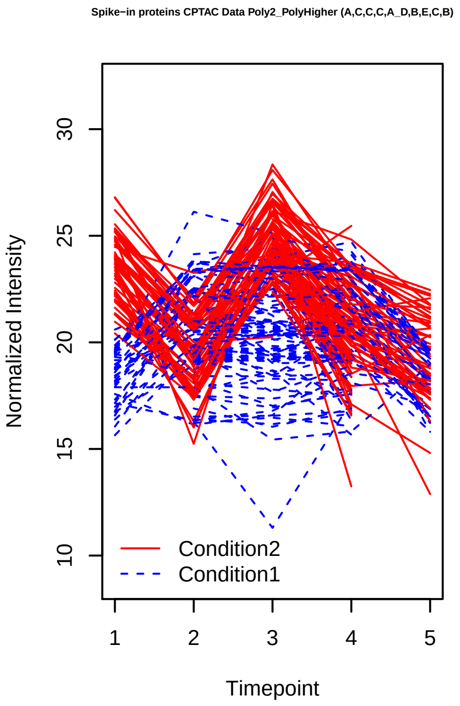
Spike−in proteins CPTAC Data Poly2_PolyHigher (A,C,C,C,A_D,B,E,C,B)
Normalized Intensity
Timepoint
Condition2
Condition1

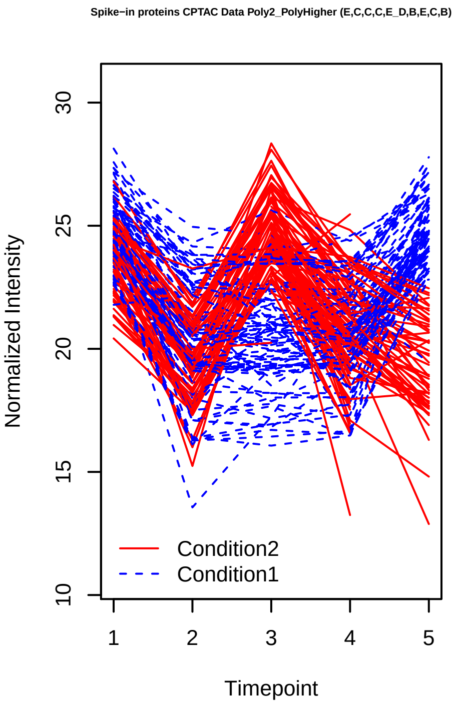
Spike−in proteins CPTAC Data Poly2_PolyHigher (E,C,C,C,E_D,B,E,C,B)
Normalized Intensity
Timepoint
Condition2
Condition1

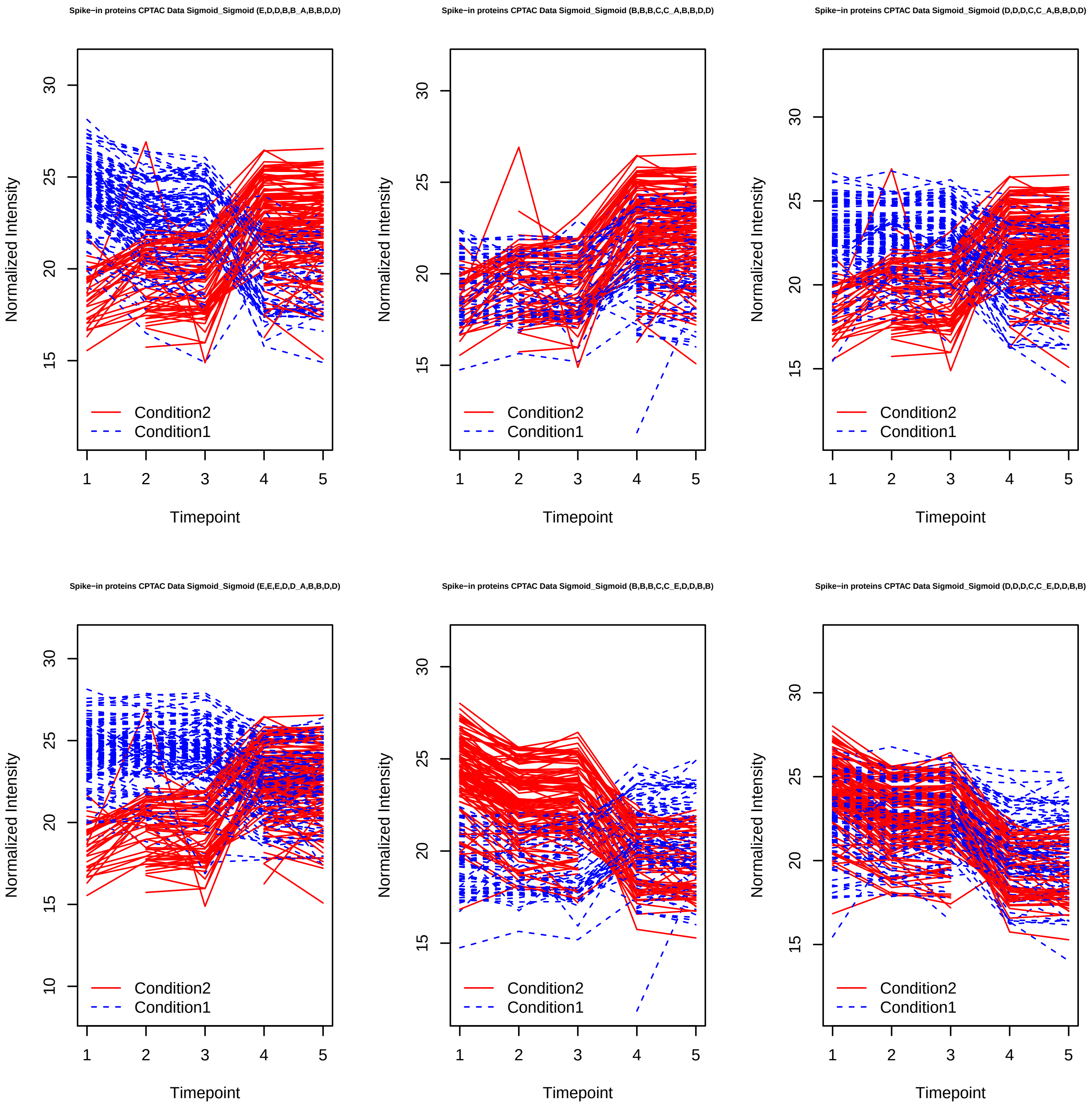
Spike−in proteins CPTAC Data Sigmoid_Sigmoid (E,D,D,B,B_A,B,B,D,D)
Normalized Intensity
Timepoint
Condition2
Condition1
Spike−in proteins CPTAC Data Sigmoid_Sigmoid (B,B,B,C,C_A,B,B,D,D)
Normalized Intensity
Timepoint
Condition2
Condition1
Spike−in proteins CPTAC Data Sigmoid_Sigmoid (D,D,D,C,C_A,B,B,D,D)
Normalized Intensity
Timepoint
Condition2
Condition1
Spike−in proteins CPTAC Data Sigmoid_Sigmoid (E,E,E,D,D_A,B,B,D,D)
Normalized Intensity
Timepoint
Condition2
Condition1
Spike−in proteins CPTAC Data Sigmoid_Sigmoid (B,B,B,C,C_E,D,D,B,B)
Normalized Intensity
Timepoint
Condition2
Condition1
Spike−in proteins CPTAC Data Sigmoid_Sigmoid (D,D,D,C,C_E,D,D,B,B)
Normalized Intensity
Timepoint
Condition2
Condition1

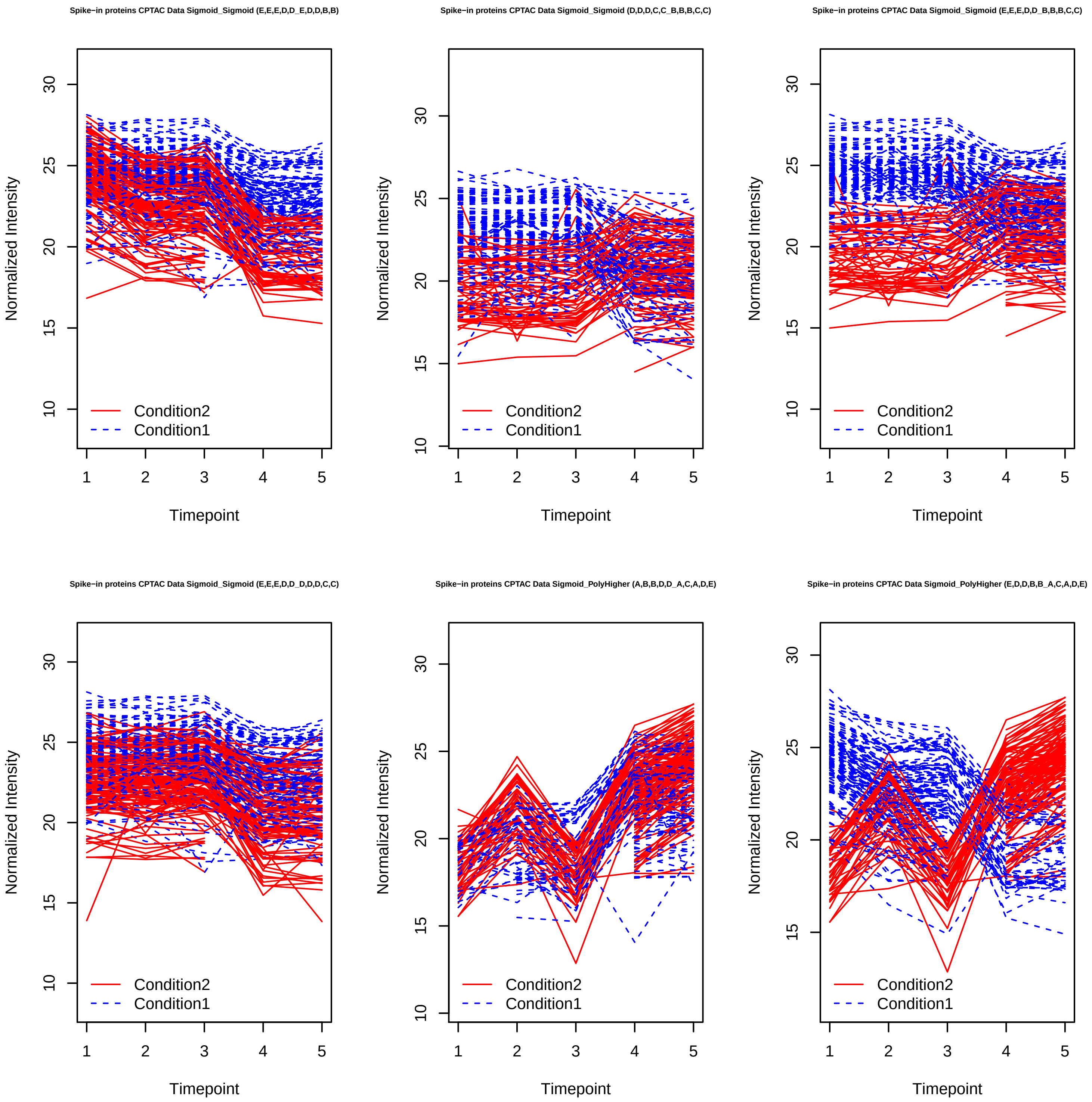
Spike−in proteins CPTAC Data Sigmoid_Sigmoid (E,E,E,D,D_E,D,D,B,B)
Spike−in proteins CPTAC Data Sigmoid_Sigmoid (D,D,D,C,C_B,B,B,C,C)
Spike−in proteins CPTAC Data Sigmoid_Sigmoid (E,E,E,D,D_B,B,B,C,C)
Spike−in proteins CPTAC Data Sigmoid_Sigmoid (E,E,E,D,D_D,D,D,C,C)
Spike−in proteins CPTAC Data Sigmoid_PolyHigher (A,B,B,D,D_A,C,A,D,E)
Spike−in proteins CPTAC Data Sigmoid_PolyHigher (E,D,D,B,B_A,C,A,D,E)
Normalized Intensity
Timepoint
Condition2
Condition1

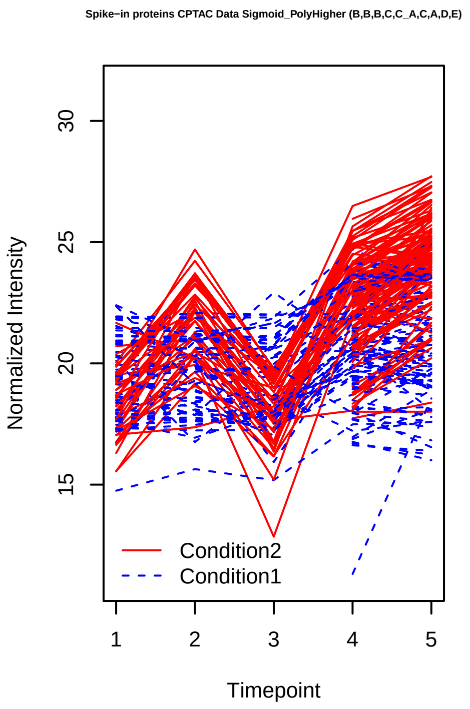
Spike−in proteins CPTAC Data Sigmoid_PolyHigher (B,B,B,C,C_A,C,A,D,E)
Normalized Intensity
15
20
25
30
Condition2
Condition1
1
2
3
4
5
Timepoint

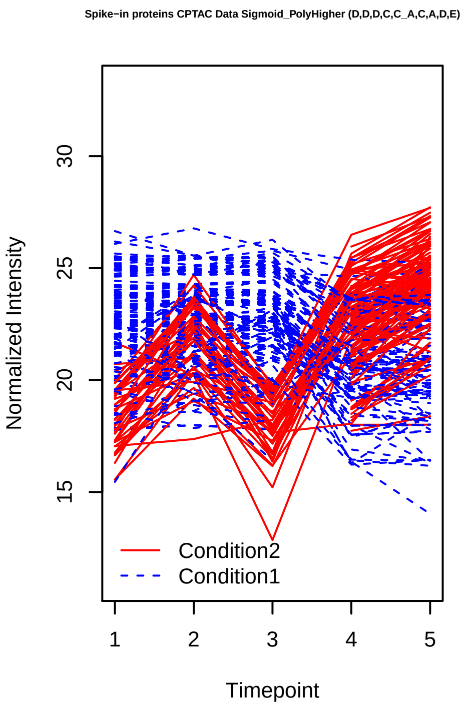
Spike−in proteins CPTAC Data Sigmoid_PolyHigher (D,D,D,C,C_A,C,A,D,E)
Normalized Intensity
15
20
25
30
Condition2
Condition1
1
2
3
4
5
Timepoint

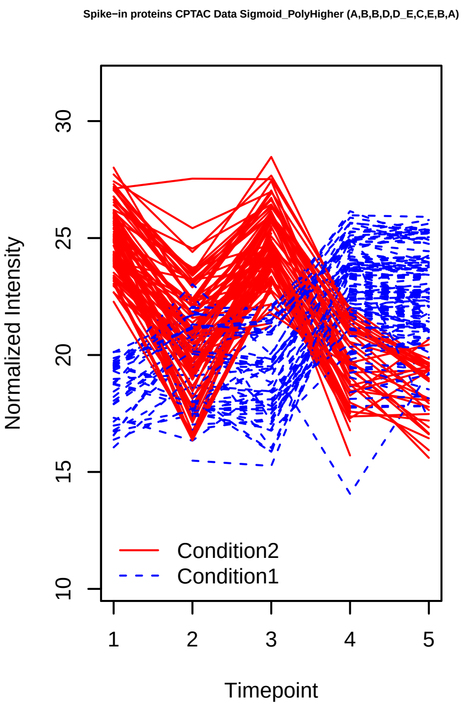
Spike−in proteins CPTAC Data Sigmoid_PolyHigher (A,B,B,D,D_E,C,E,B,A)
Normalized Intensity
10
15
20
25
30
Condition2
Condition1
1
2
3
4
5
Timepoint

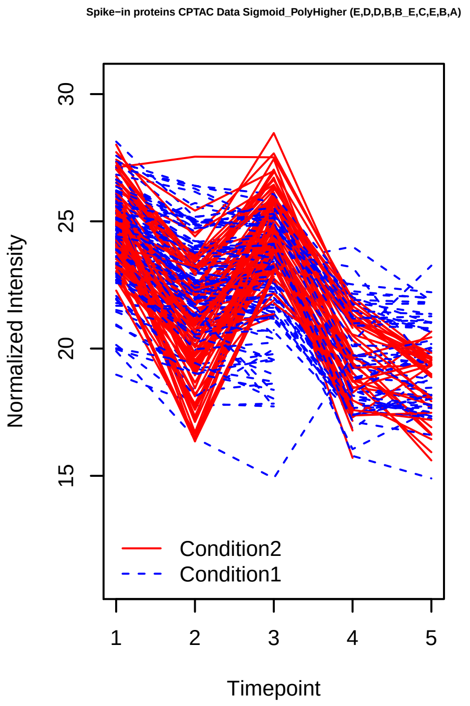
Spike−in proteins CPTAC Data Sigmoid_PolyHigher (E,D,D,B,B_E,C,E,B,A)
Normalized Intensity
15
20
25
30
Condition2
Condition1
1
2
3
4
5
Timepoint

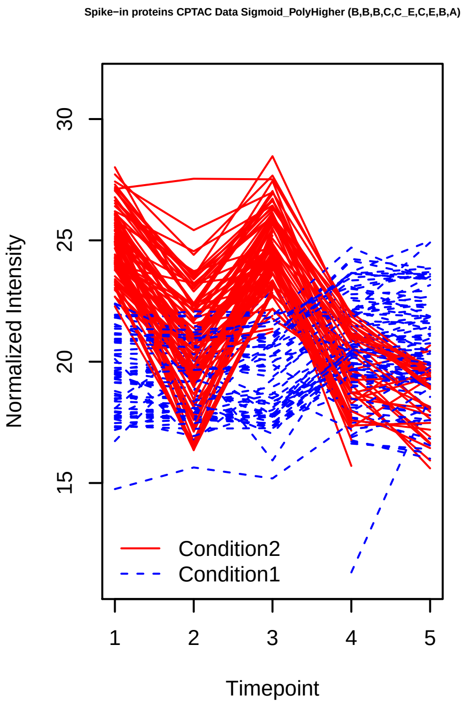
Spike−in proteins CPTAC Data Sigmoid_PolyHigher (B,B,B,C,C_E,C,E,B,A)
Normalized Intensity
15
20
25
30
Condition2
Condition1
1
2
3
4
5
Timepoint

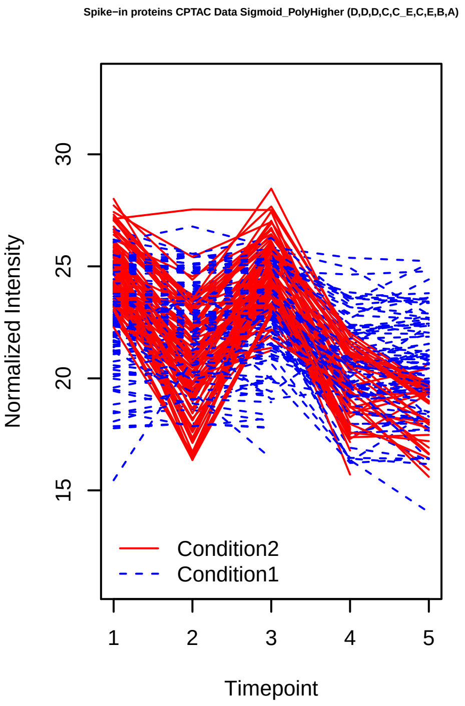
Spike−in proteins CPTAC Data Sigmoid_PolyHigher (D,D,D,C,C_E,C,E,B,A)
Normalized Intensity
15
20
25
30
Condition2
Condition1
1
2
3
4
5
Timepoint

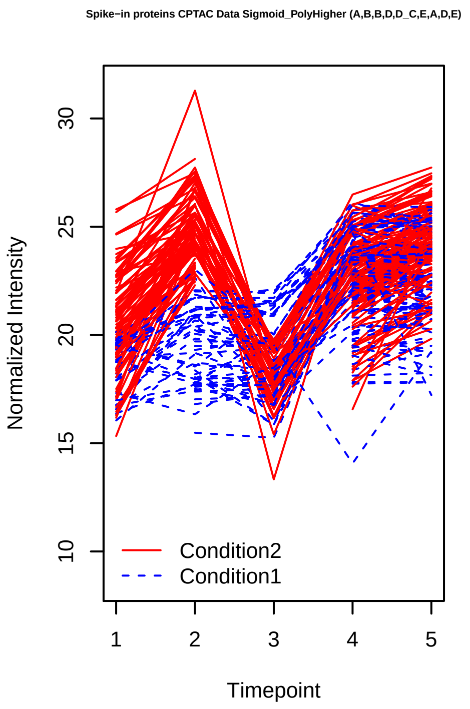
Spike−in proteins CPTAC Data Sigmoid_PolyHigher (A,B,B,D,D_C,E,A,D,E)
Normalized Intensity
10 15 20 25 30
1 2 3 4 5
Timepoint
Condition2
Condition1

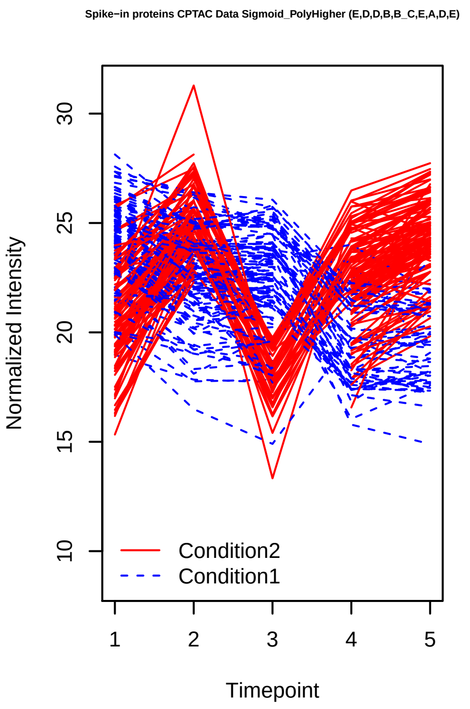
Spike−in proteins CPTAC Data Sigmoid_PolyHigher (E,D,D,B,B_C,E,A,D,E)
Normalized Intensity
10 15 20 25 30
1 2 3 4 5
Timepoint
Condition2
Condition1

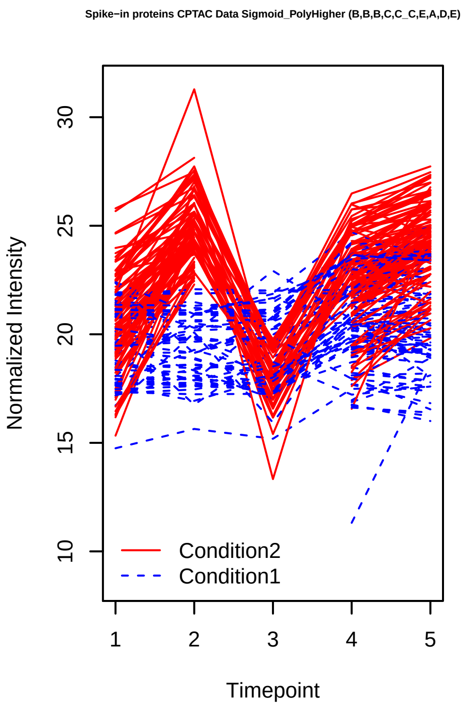
Spike−in proteins CPTAC Data Sigmoid_PolyHigher (B,B,B,C,C_C,E,A,D,E)
Normalized Intensity
10 15 20 25 30
1 2 3 4 5
Timepoint
Condition2
Condition1

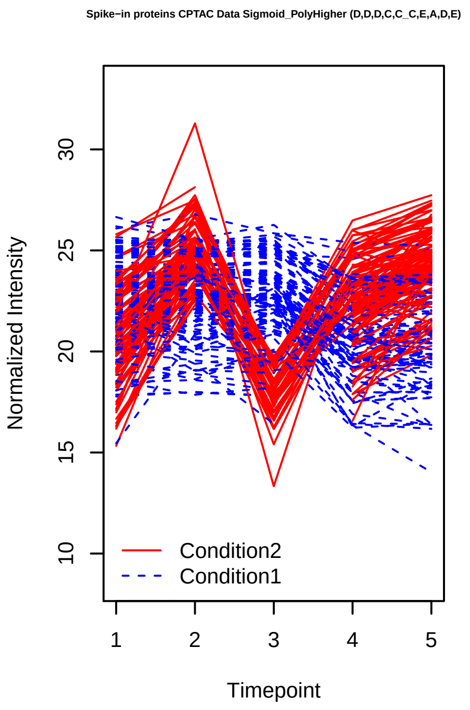
Spike−in proteins CPTAC Data Sigmoid_PolyHigher (D,D,D,C,C_C,E,A,D,E)
Normalized Intensity
10 15 20 25 30
1 2 3 4 5
Timepoint
Condition2
Condition1

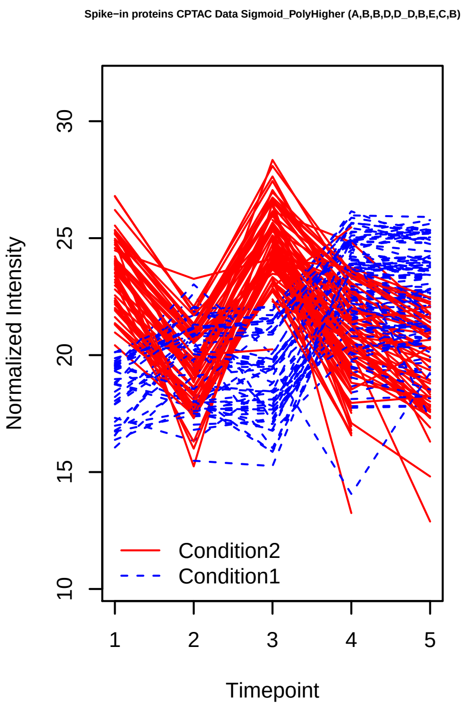
Spike−in proteins CPTAC Data Sigmoid_PolyHigher (A,B,B,D,D_D,B,E,C,B)
Normalized Intensity
10 15 20 25 30
1 2 3 4 5
Timepoint
Condition2
Condition1

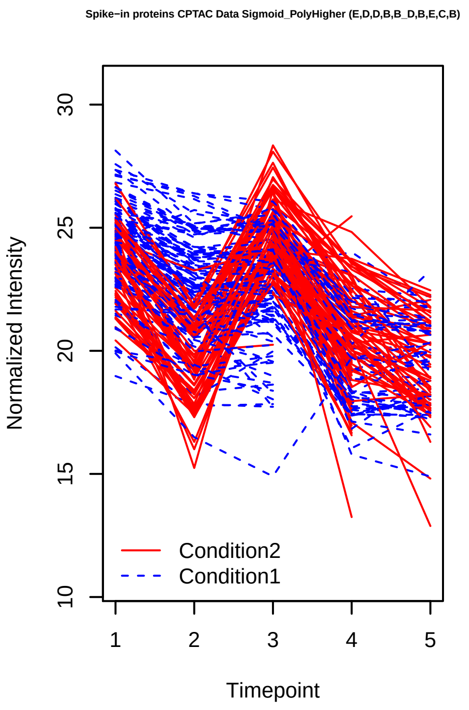
Spike−in proteins CPTAC Data Sigmoid_PolyHigher (E,D,D,B,B_D,B,E,C,B)
Normalized Intensity
10 15 20 25 30
1 2 3 4 5
Timepoint
Condition2
Condition1

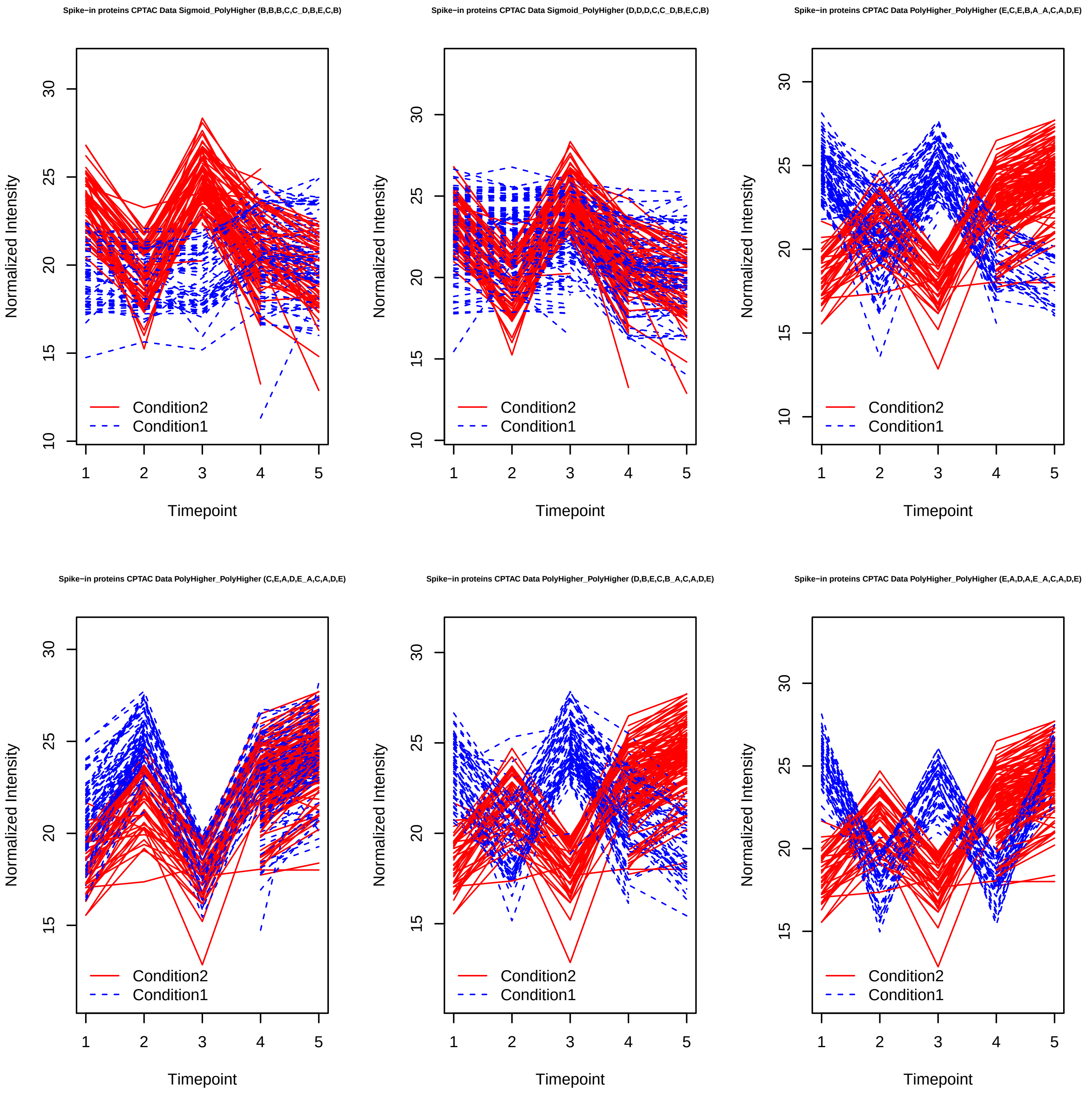
Spike−in proteins CPTAC Data Sigmoid_PolyHigher (B,B,B,C,C_D,B,E,C,B)
Spike−in proteins CPTAC Data Sigmoid_PolyHigher (D,D,D,C,C_D,B,E,C,B)
Spike−in proteins CPTAC Data PolyHigher_PolyHigher (E,C,E,B,A_A,C,A,D,E)
Spike−in proteins CPTAC Data PolyHigher_PolyHigher (C,E,A,D,E_A,C,A,D,E)
Spike−in proteins CPTAC Data PolyHigher_PolyHigher (D,B,E,C,B_A,C,A,D,E)
Spike−in proteins CPTAC Data PolyHigher_PolyHigher (E,A,D,A,E_A,C,A,D,E)
Normalized Intensity
Timepoint
Condition2
Condition1

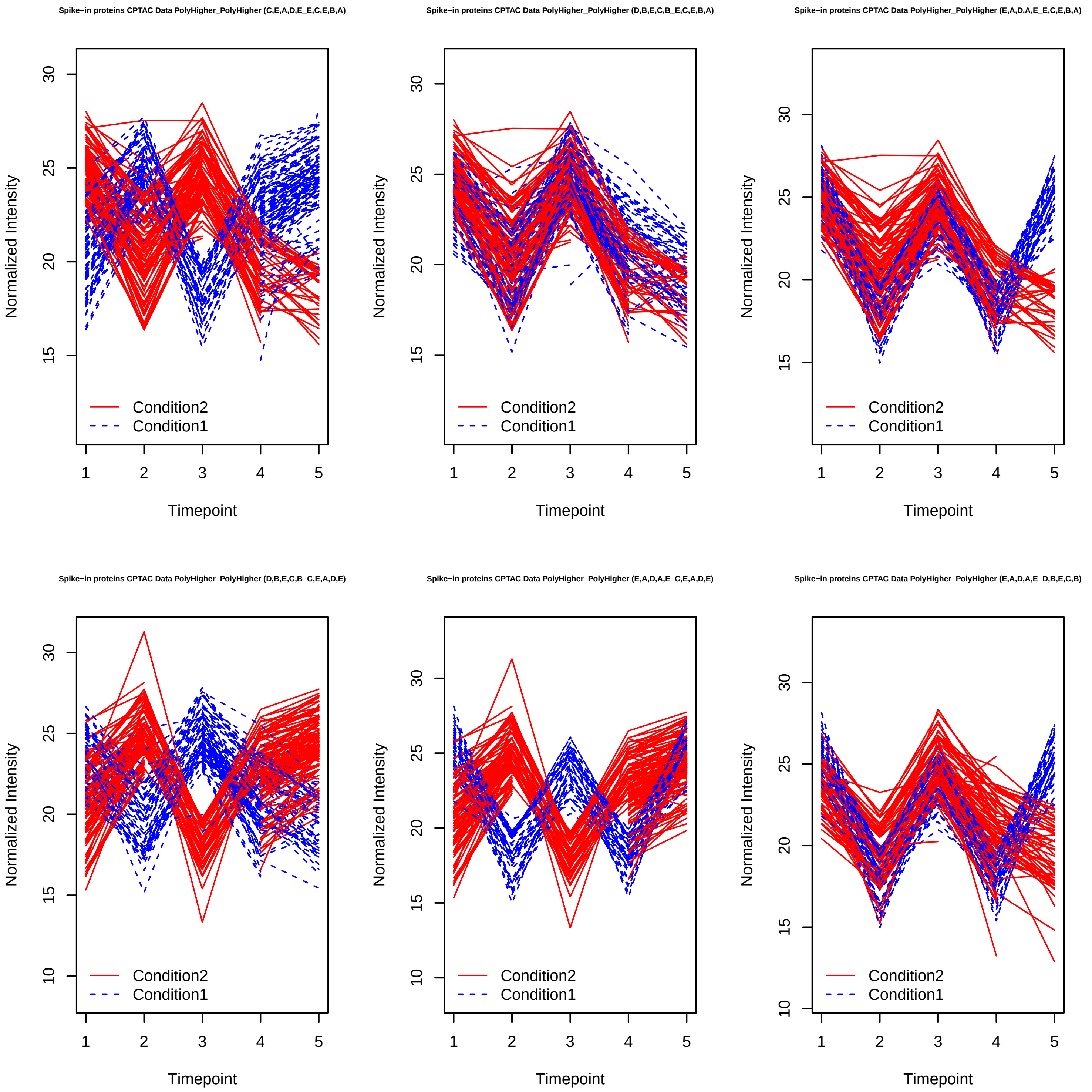
Spike−in proteins CPTAC Data PolyHigher_PolyHigher (C,E,A,D,E_E,C,E,B,A)
Spike−in proteins CPTAC Data PolyHigher_PolyHigher (D,B,E,C,B_E,C,E,B,A)
Spike−in proteins CPTAC Data PolyHigher_PolyHigher (E,A,D,A,E_E,C,E,B,A)
Spike−in proteins CPTAC Data PolyHigher_PolyHigher (D,B,E,C,B_C,E,A,D,E)
Spike−in proteins CPTAC Data PolyHigher_PolyHigher (E,A,D,A,E_C,E,A,D,E)
Spike−in proteins CPTAC Data PolyHigher_PolyHigher (E,A,D,A,E_D,B,E,C,B)
Normalized Intensity
Timepoint
Condition2
Condition1
